# Robust clustering and interpretation of scRNA-seq data using reference component analysis

**DOI:** 10.1101/2021.02.16.431527

**Authors:** Florian Schmidt, Bobby Ranjan, Quy Xiao Xuan Lin, Vaidehi Krishnan, Ignasius Joanito, Mohammad Amin Honardoost, Zahid Nawaz, Prasanna Nori Venkatesh, Joanna Tan, Nirmala Arul Rayan, S.Tiong Ong, Shyam Prabhakar

**Author notes:** contributed equally (Florian Schmidt, Bobby Ranjan).

## Abstract

**Motivation:** The transcriptomic diversity of the hundreds of cell types in the human body can be analysed in unprecedented detail using single cell (SC) technologies. Though clustering of cellular transcriptomes is the default technique for defining cell types and subtypes, single cell clustering can be strongly influenced by technical variation. In fact, the prevalent unsupervised clustering algorithms can cluster cells by technical, rather than biological, variation.

**Results:** Compared to *de novo* (unsupervised) clustering methods, we demonstrate using multiple benchmarks that supervised clustering, which uses reference transcriptomes as a guide, is robust to batch effects. To leverage the advantages of supervised clustering, we present RCA2, a new, scalable, and broadly applicable version of our RCA algorithm. RCA2 provides a user-friendly framework for supervised clustering and downstream analysis of large scRNA-seq data sets. RCA2 can be seamlessly incorporated into existing algorithmic pipelines. It incorporates various new reference panels for human and mouse, supports generation of custom panels and uses efficient graph-based clustering and sparse data structures to ensure scalability. We demonstrate the applicability of RCA2 on SC data from human bone marrow, healthy PBMCs and PBMCs from COVID-19 patients. Importantly, RCA2 facilitates cell-type-specific QC, which we show is essential for accurate clustering of SC data from heterogeneous tissues. In the era of cohort-scale SC analysis, supervised clustering methods such as RCA2 will facilitate unified analysis of diverse SC datasets.

**Availability:** RCA2 is implemented in R and is available at github.com/prabhakarlab/RCAv2

## Introduction

Since its first usage in 2009 (1), single cell (SC) RNA sequencing (scRNA-seq) has quickly become the method of choice for profiling gene expression in complex samples (2). Due to the unprecedented resolution of scRNA-seq data, cell-type-specific analysis of gene expression can now be performed easily and at low cost. SCTs are well-suited to characterize heterogeneous biological specimens, e.g. tumors (3). Clustering is an essential step in SC data analysis, since each cell cluster in transcriptome space represents a distinct cell type or state. There are two established paradigms to address the SC clustering problem: [i] unsupervised (*de novo*) clustering (4) and [ii] supervised approaches that use reference data sets to either cluster cells or to classify cells into cell types (5, 6). The graph-based clustering methods Seurat (7) and Scanpy (8) are among the most prevalent *de novo* clustering approaches.

However, SC clustering is a challenging algorithmic problem: 1) cells may cluster by technical variation and batch effects rather than biological properties (4), 2) scRNA-seq data tend to be noisy, primarily due to sampling noise and 3) the gene expression matrix can be very large, since modern datasets commonly include > 100, 000 cells. Consequently, different algorithms can return highly divergent clusterings, i.e. partitions of cells into clusters, of the same input dataset (9). Moreover, *de novo* clustering requires an error-prone, time-consuming manual step of assigning cell clusters to cell types (annotation) based on subjective evaluation of the expression of literature marker genes. Supervised clustering and supervised cell type annotation algorithms have been developed to address these limitations (6).

Previously, we proposed *Reference Component Analysis (RCA)* for supervised clustering of scRNA-seq data guided by a panel of reference transcriptomes (5). Unlike methods such as SingleR (10) or scMatch (11), the aim of RCA is not cell type annotation. Rather, RCA was designed to cluster cells in the space of reference transcriptome projections. As we show below, this reference-based clustering approach is applicable even in situations where the dataset contains cell states not present in the reference panel. To the best of our knowledge, RCA is the only supervised clustering algorithm for scRNA-seq data. However, the original version of RCA cannot scale to datasets larger than 20, 000*cells* on a high-end laptop, used only a single reference panel, did not implement differential expression and Gene Ontology (GO) enrichment analysis, was benchmarked on only a single Smart-seq dataset and could not be easily integrated into existing data analysis workflows.

To fully leverage the merits of supervised clustering, we present RCA2, a new implementation of reference-based clustering that addresses the above limitations. We show the unique advantages of RCA2 by analyzing diverse scRNA-seq data sets and show that supervised clustering can detect novel disease/condition-specific cell states. Importantly, we demonstrate that supervised clustering using RCA2 is exceptionally robust against batch effects on large scRNA-seq data sets, rendering it the method of choice for the joint analysis of heterogeneous cohort-scale datasets.

## MATERIALS AND METHODS

### Projection to a reference

Given a reference data set 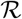 containing *n* cell types and 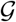 marker genes as well as a query data set 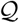 containing *i* scs and 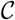 genes, we determine a marker gene set 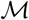 by intersecting 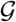 and 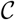:

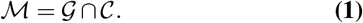

The reference matrix 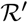 and set 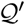 are generated by extracting the gene set 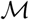 from 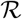 and 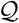, respectively. Next, RCA2 computes the correlation (default: Pearson(*r*)) between 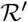 and 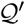 yielding a reference projection 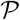 for *i* scs to *n* cell types:

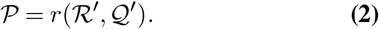

The projection matrix 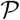 is modified according to

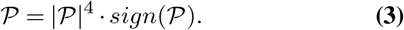

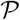 is scaled to zero mean and unit variance. All matrices are represented as *sparse matrix* R objects. The projection is computed using the *fastcor* package (12). 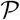 can be visualized as a 2D and 3D UMAP.

### Clustering and interpreting the projection

RCA2 offers three clustering algorithms: i) hierarchical clustering using the memory efficient *fastcluster* (13) package, ii) shared-nearest neighbour (SNN) clustering using *dbscan* (14) and iii) graph based clustering using the Louvain algorithm (7). The depth to cut the dendrogram in hierarchical clustering is a parameter (default 1). The SNN algorithm used in *db-scan* has three parameters: *k* (neighborhood size of the SNN graph), *eps* (two cells are only reachable from each other if they share at least *eps* nearest cells) and *min pts* (minimum number of nearest neighbours for a cell to be considered a core cell). To guide the users choice on parameters for graph based clustering, a 3D figure illustrating how the final number of clusters depends to the used parameters can be generated. The Louvain algorithm requires only the *resolution* parameter. A line-plot illustrating how the resolution influences the number of identified clusters can be generated. As input, all clustering methods use either a distance matrix 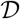 computed from 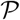 according to

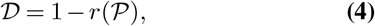

where 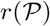 is the cell-to-cell similarity using correlation (Pearson(default in this manuscript), Spearman or Kendal) as a metric in the cell type space or an embedding of cells in PC space computed on the reference projection (not available with hierarchical clustering). The clustering result is visualized in a heatmap, including quality control (QC) metrics: number of detected genes (NODG), the percentage of mitochondrial genes (pMito) and the number of unique molecular identifiers (NUMI). Reference cell types with a low variance across all query cells are not shown.

### Reference panels

RCA2 includes ten human reference panels as well as two mouse reference panels (Sup. Section 1). Multiple panels can be used for reference projections simultaneously. Furthermore, RCA2 provides users with the option to generate their own reference panel: the *buildReferencePanel* function considers a bulk gene expression matrix (genes as rows and replicates as columns) of raw counts and returns a reference panel that can be used with RCA2. Details are provided in Sup. Section 2.1.

### Annotation of cell types

RCA2 implements cell type assignment at the SC level following a strategy inspired by SINGLER (Sup. Section 2.2). To annotate cell types on the cluster level, we consider the cell type composition for each cluster based on the SC cell type assignment described above. If the cell type distribution within a cluster is heterogeneous and the proportion of the major cell type is below a user defined threshold (default 50%), the cluster is labelled as *Unknown*.

### Cluster specific quality control

Quality of scRNA-seq data is usually assessed using NODG, nUMI and pMito metrics. RCA2 provides cluster-specific QC, allowing to impose upper/lower bounds on QC metrics for each cluster independently.

### Differentially expressed gene computation and enrichment analysis

Differentially expressed genes (DEGs) are calculated between clusters, either in a *1 vs. all* (default) or a *pairwise* fashion using a modified version of Seurat’s DEG calling function. We incorporated a mean expression threshold, which is either a user defined value or automatically determined as a trimmed mean excluding the top *n* (default: 5) genes with the highest expression. Gene’s with a cluster specific expression below the threshold are not considered for the DE test. DEGs computed in this way are used to perform enrichment analysis of GeneOntology (GO) terms or KEGG Pathways, for which RCA2 incorporates the ClusterPro-filer R-package (15). Details are provided in Sup. Section 2.3.

### Considered scRNA-seq data sets and data processing

#### 10X PBMC data sets

We downloaded scRNA-seq data of 5025 PBMCs generated using Chromium SC 3’ Reagent Kits v3 from 10X (single-cell-gene-expression/datasets/3.0.2/5k_pbmc_protein_v3). In total, 4249 cells passed QC (Sup. Table 1). The data set was projected against the *Novosthern* reference panel comprised of 15 hematopoietic cell types (16) (Sup. Sec. 1).The resolution parameter used for Louvain clustering was determined using a grid-search with a step-size of 0.05. DEGs between clusters are computed in a pairwise-manner using the parameters:*min.pct*= 0.5*, logfc.threshold*= 0.5 *and p_val_adj*≤0.05. GO terms were computed using Clus-terProfiler utilising the *org.Hs.eg.db* database, and a q-value threshold of 0.05.

#### CITE-seq PBMC data sets

A Drop-Seq data set with 29, 929 genes profiled in 7, 985 cells and 10 antibodies was obtained from (17). A 10x CITE-seq data set with 7, 865 cells profiled on 33, 538 genes and 17 antibodies was obtained from 10x (single-cell-gene-expression/datasets/3.0.0/pbmc_10k_protein_v3).

Both data sets were processed using Seurat. To obtain a ground truth, we clustered cells in antibody-derived tags (ADT) space. ADT data was normalized using the centered log ratio transformation (sati-jalab.org/seurat/v3.2/multimodal_vignette.html). All PCs were selected for clustering using Louvain clustering. Since the Drop-Seq data did not include a control for antibody detection, clusters exhibiting noisy antibody detection or those clusters not representing known immune cell type signature (16, 18) were removed. In the 10x data, clusters showing *IgG1*, *IgG2a* or *IgG2b* and clusters showing promiscuous antibody expression were discarded. After QC (Sup. Table 1), the Drop-Seq and 10x datasets contained 5, 925 and 6, 744 cells, respectively. The Drop-Seq and 10x data sets were next merged with respect to their common genes (13, 267). Using Seurat’s FindMarker function, we computed batch specific marker genes for each ADT cluster in the merged data set using the parameters: *min.pct*= 0.5*, logfc.threshold*= 1.5 *and p_val_adj*≤0.05. GO terms for batch specific clusters are computed using ClusterPro-filer utilising the *org.Hs.eg.db* database, and a q-value threshold of 0.05.

#### Rheumatoid arthritis scRNA-seq data set

The scRNA-seq data set of 10, 001 cells from Rheumatoid Arthritis (RA) samples, obtained from (19), was processed using Seurat. Cells were filtered based on the QC criteria provided in Sup. Table 1. This data was generated using CEL-Seq2 (20) after sorting for B cells (CD45+CD3-CD19+), T cells (CD45+CD3+), monocytes (CD45+CD14+), and stromal fibroblasts (CD45-CD31-PDPN+) from synovial tissues of ultrasound-guided biopsies or joint replacements of RA patients. Cell type annotation based on the authors cell sorting strategy is used as a ground truth.

#### Bone-marrow scRNA-seq data set

We obtained eight human bone marrow (BM) specimens from *STEMCELL technologies* and generated ten scRNA-seq data sets using the 10X 5’scRNA-seq protocol (see Sup. Section 2.4) by separating the cells into CD34+ and CD34-cell fractions followed by sequencing on a HiSeq4000. Preprocessing was done using the 10X CellRanger pipeline (3.0.1) using the *hg38* reference genome resulting in a data set comprised of 45, 363 cells, capturing 24, 206 genes. Considering only an initial requirement of at least 1, 000 nUMI, and a pMito rate between 2.5% and 10%, we projected the data against RCA’s global panel obtaining a classification into major groups (resolution 0.1) to perform cluster specific QC (Sup. Table 2). Final cell types were identified upon QC using a resolution of 0.5. Doublets were removed using DoubletFinder (21). DoubletFinder was run separately on the CD34+ and CD34-populations using 20 PCs and a pNN value of 0.25 as well as pk values of 0.005 and 0.01, respectively. The obtained *pANN* values were merged to rank cells based on their doublet neighbourhood. A *pANN* threshold was derived considering both the expected number of doublets (≈ 1, 560) and by examining the proportion of possible doublets in each cluster.

#### COVID-19 PBMCs

A Seurat object with scRNA-seq data studied by (22) was obtained from the COVID-19 Cell Atlas (covid19cellatlas.org). All 44, 721 cells were projected against the global, Monaco, Novershtern, and CITE-seq panel and clustered using the Louvain algorithm (resolution 1.3). Clusters were annotated according to reference projection profiles and marker gene expression. Marker genes for each subset of developing neutrophils are computed using the following DEG parameters: *min.pct*= 0.25*, logfc.threshold*= 1*, and p_val_adj*≤0.05. Sub-clustering and cell embedding of CD14 monocytes, intermediate monocytes, CD16 monocytes, myeloid dendritic cells (mDC), myelocytes, neutrophils and plasmablasts was conducted using Seurat. The union set of pair-wise DEGs between these cell types was used as feature genes for PCA. DEGs were determined using the parameters: *min.pct*= 0.25*, logfc.threshold*= 1.5*, and p_val_adj*≤0.05. For downstream clustering we used 18 PCs and annotated clusters using marker genes.

#### AML dataset

The AML data scRNA-seq data set *809653* was obtained from the zenodo archive of (23) at 10.5281/zen-odo.3345981. No additional QC was performed. The data was projected using RCA2s Multi Panel Projection function with default parameters. The data was clustered using hierarchical clustering at deep split 1.

### Methods used for batch effect benchmarking

To benchmark the batch effect robustness of RCA2, we considered Seurat, Scran (24), Scanpy, Seurat with CCA, MNNCorrect (25) and Scanorama (26). Seurat (with and without CCA) (3.2) was used as recommended in its documentation (satijalab.org/seurat/vignettes.html). For Scran (1.18.1), we followed the tutorial from bioconductor (bio-conductor.org/packages/ release/bioc/html/scran.html). Highly variable genes are used as features (FDR ≤ 0.05) and the walktrap community detection algorithm was used for clustering. We used MNNCorrect (batchelor R package (1.6)) according to biocon-ductor.org/packages/release/bioc/html/batchelor.html.

Furthermore, we used Scanpy (1.5.1) (scanpy.readthedocs.io/en/stable/) and Scanorama (1.6) (nbisweden.github.io/workshop-scRNAseq). We also exploit SCTransform (27) to normalize data before analyzing it with Seurat or Seurat with CCA.

### Silhouette Index for quantifying batch effect

The Silhouette Index (SI) of a cell measures how similar a cell is to other cells within its own cluster, relative to cells in other clusters (28). We compute SI *S*(*x*) for each cell *x* in the DE gene-space defined by CITE-Seq antibody tags:

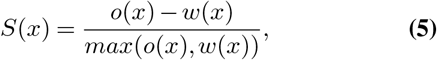

where *o*(*x*) is the smallest mean between-cluster distance and *w*(*x*) is the mean within-cluster distance for cell *x* defined as

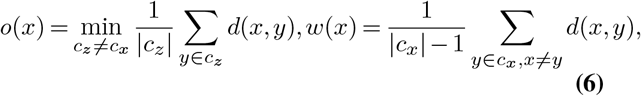

w here we use Euclidean distance to compute the distance *d*(*x, y*) between cell *x* and cell *y*,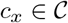 is the cluster assigned to cell *x* and |*c_x_*| is the size of that cluster. We obtain the average SI for each cluster by averaging the SI values over all cells in that cluster. Thereby, each cell type is given equal weight in the final SI score.

For Seurat, Seurat Integration (CCA), scran, MNNCorrect and Scanpy, cell-cell distances are calculated in principal component (PC) space considering the top 20 PCs. For Scanorama, the dimensionality was fixed to 100, as recommended by the authors. For RCA, cell-cell distances were calculated in the reference projection space.

### Implementation and Availability

RCA2 is freely available at www.github.com/prabhakarlab/RCAv2. It is extensively tested with 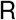 versions ≥ 3.6 on Windows, Linux and Mac devices. Scripts to create the main figures and RDS files with R objects for the batch effect benchmarking, the 10X PBMC data, the BM use case and the COVID-19 data are available at Zenodo (10.5281/zenodo.4021967). Fastq files for the BM data are available upon request.

## Results

### RCA2 is a comprehensive software solution for supervised clustering and analysis of scRNA-seq data

The RCA2 workflow is shown in Fig.1. As input, RCA2 takes either raw or pre-processed scRNA-seq data and facilitates QC either as a single operation on all cells or in a cluster-specific manner (Sup. Fig. 1, 2).

**Fig. 1.**
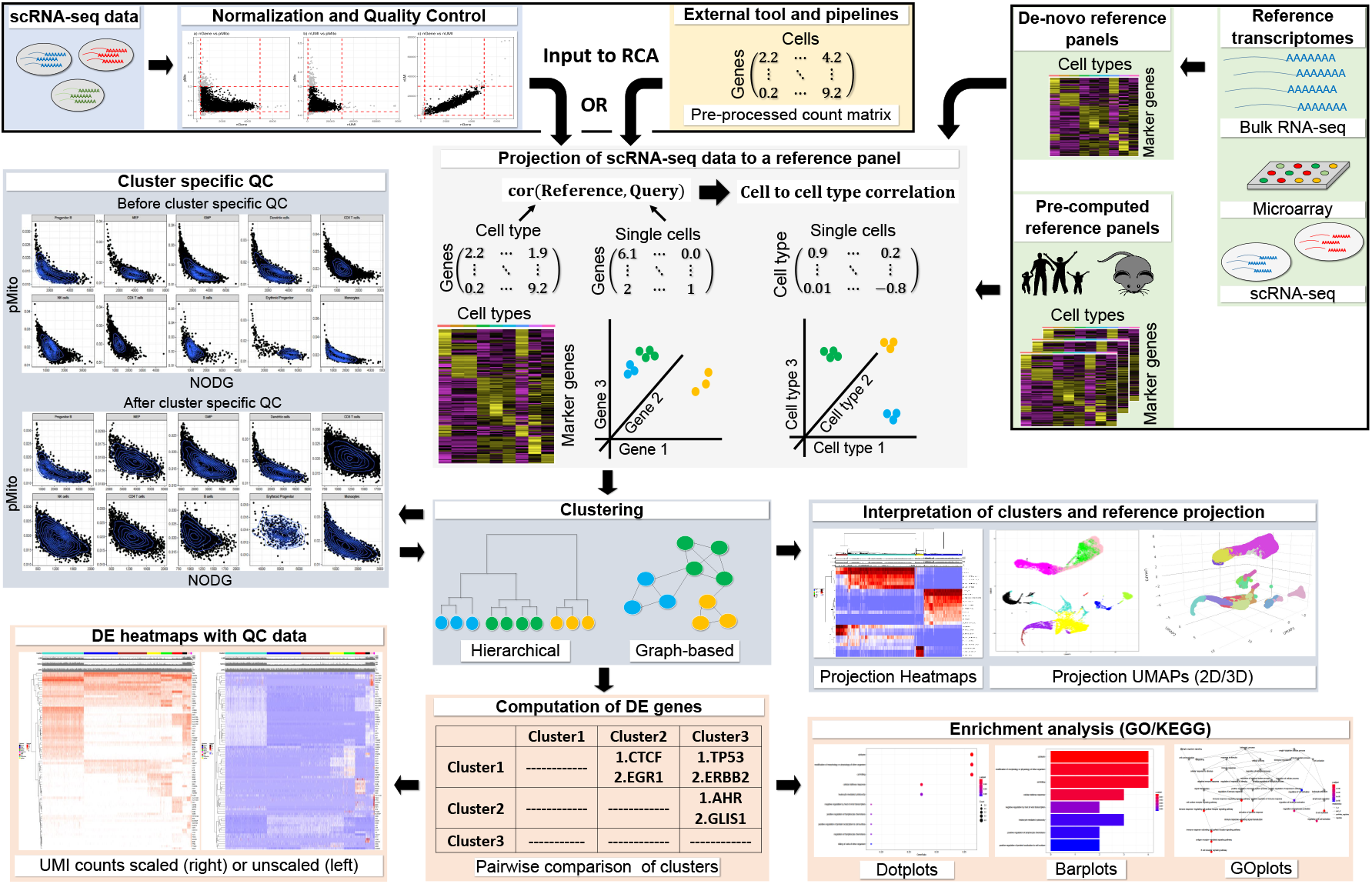
RCA2 takes two types of scRNA-seq data as input: 1) CellRanger output files and 2) data preprocessed elsewhere, which can be loaded as a gene x cell count matrix. Reference datasets in RCA2 for human and mouse are based on bulk RNA-seq, microarray and scRNA-seq assays. RCA2 can also generate custom reference panels from user-supplied raw count matrices. RCA2 computes a correlation matrix representing the similarity of each SC transcriptome to each reference transcriptome. Correlations are calculated using marker (DE) genes from the reference panel. Cells are clustered and visualized in the space of reference projections After DE gene analysis, enriched GO terms and KEGG pathways can be identified.

**Fig. 2.**
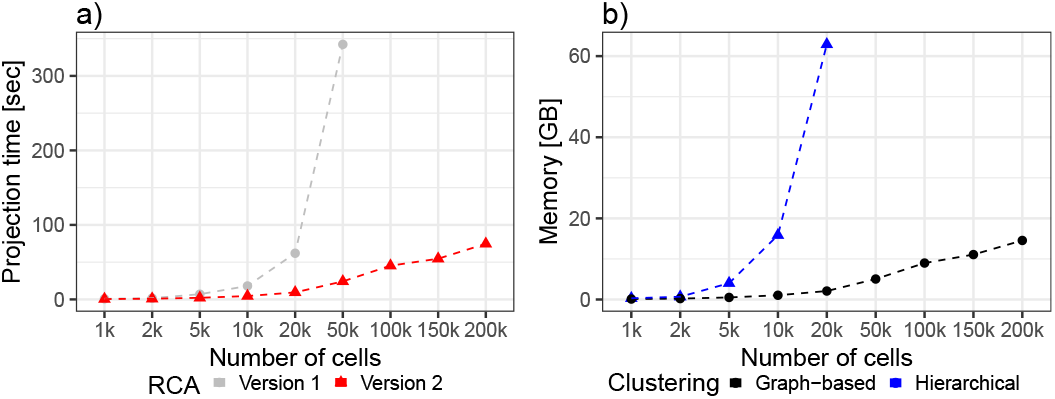
a) Speedup of the reference projection step. b) Memory consumption of graph based clustering compared to hierarchical clustering. Benchmarking was performed with a notebook using an Intel i9-9980 CPU(2.40GHz) and 64GB RAM. Projecting using RCAv1 and hierarchical clustering ran out of memory using 100*k* cells and 50*k* cell, respectively.

While the first release of RCA provided one reference panel only, RCA2 includes twelve panels, for instance a microarray-based human cord blood cell panel with 15 immune cell types (16), one related RNA-seq panel with 28 human immune cell types (18), a panel based on CITE-seq data containing 34 primary human cell types (29), one human primary cell type panel based on ENCODE (30) RNA-seq data containing 97 cell types and a mouse ENCODE panel with 15 cell types. A list of all panels is provided in Sup. Section 1. Also, RCA2 offers means for de-novo panel generation from user-provided transcriptomes (Sup. Sec. 2.4). Unlike the original RCA software, RCA2 allows SC data to be projected against several reference panels at the same time and offers a significant speed up of several folds in computing the projection (Fig. 2a, Sup. Sec. 2.5).

Rising cell numbers cause memory consumption to rise, hence RCA2 uses a memory-efficient implementation of hierarchical clustering (31) and provides two graph-based clustering methods as an alternative: SNN clustering (DBSCAN) and Louvain clustering as used in Seurat. The latter two are not only faster than hierarchical clustering, but also require less memory 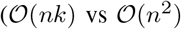, where *n* is the number of cells and *k* is the number of nearest neighbours; Fig. 2b (Sup. Sec. 2.5). To aid in parameter selection RCA2 provides visualizations on how parameter settings influence cluster numbers (Sup. Fig. 3, 4). These modifications render RCA2 to be applicable even to datasets comprising millions of cells. Note that unlike other SC clustering frameworks, RCA utilizes the reference projection to cluster cells in a cell type space instead of a high-dimensional gene-space. Clustering results can be visualised as a heatmap and as 2D as well as 3D UMAPs (Sup. Fig. 5, 6). Compared to the previous visualization, RCA2 shows additional QC information (NODG, nUMI, pMito) and removes cell types not showing significant variation in the correlation score to facilitate data interpretation and outlier detection.

**Fig. 3.**
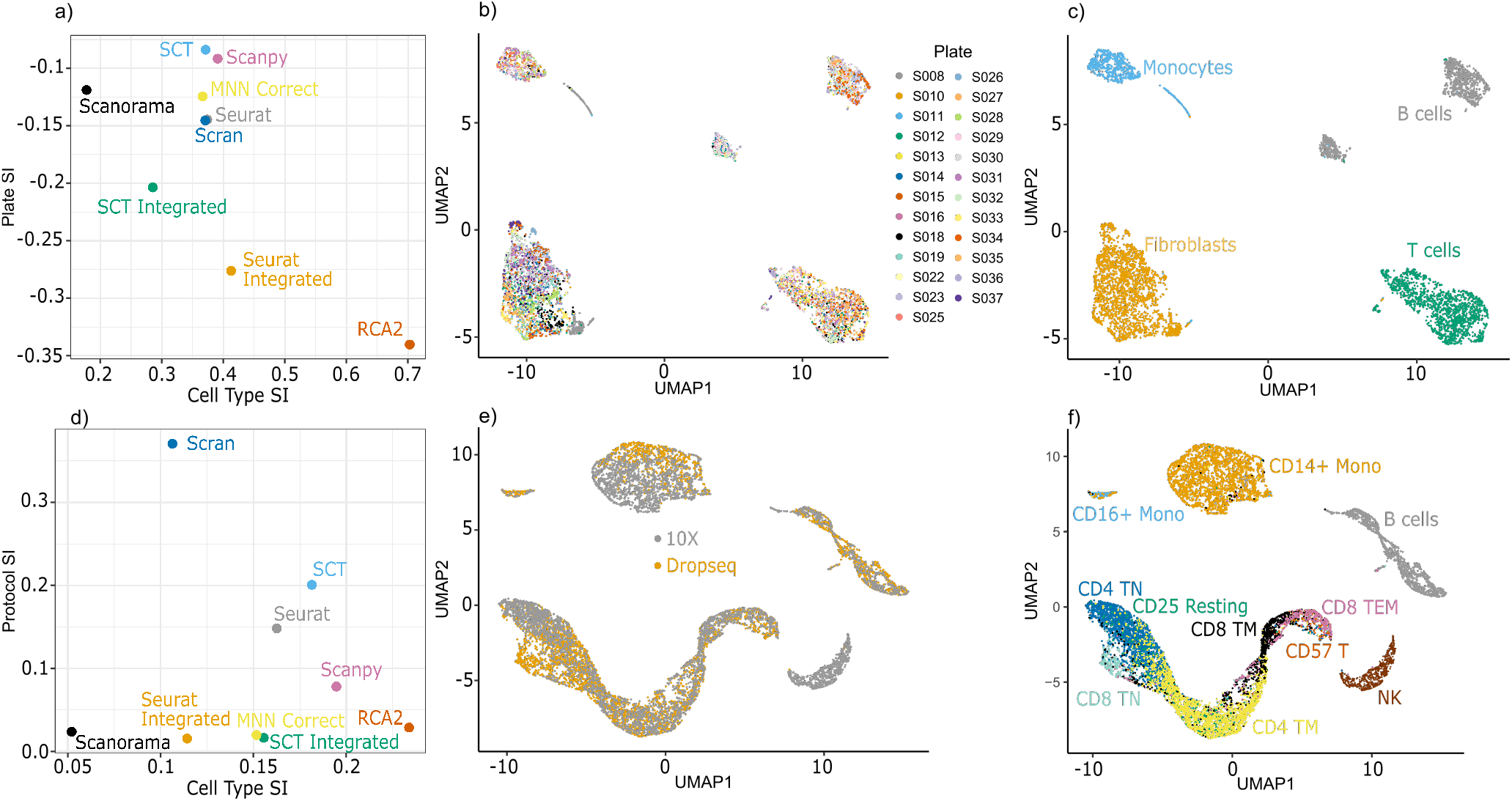
a) Silhouette Index (SI) measuring separation of cells in RA data by plate and cell type. b-c) UMAP visualization of RCA2 clustering of RA data colored by b) plate and c) cell type. d) SI measuring separation of cells in CITE-Seq data by protocol and cell type. e-f) UMAP visualization of RCA2 clustering of CITE-Seq data colored by e) protocol and f) cell type.

**Fig. 4.**
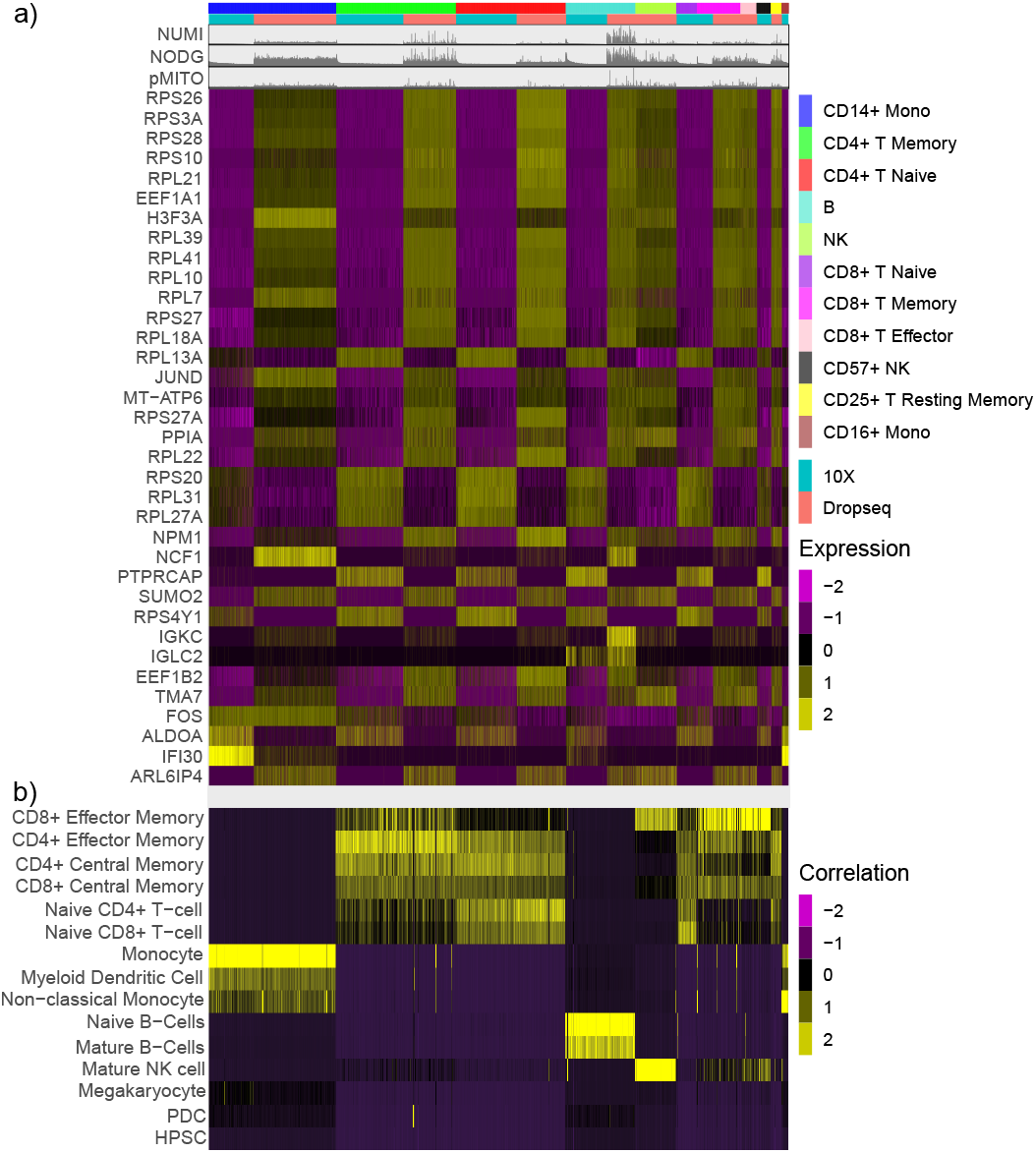
a) Expression of DEG computed for sequencing protocol batch within ADT clusters. b) Reference projection of the CITE-seq data against RCA2’s global panel.

**Fig. 5.**
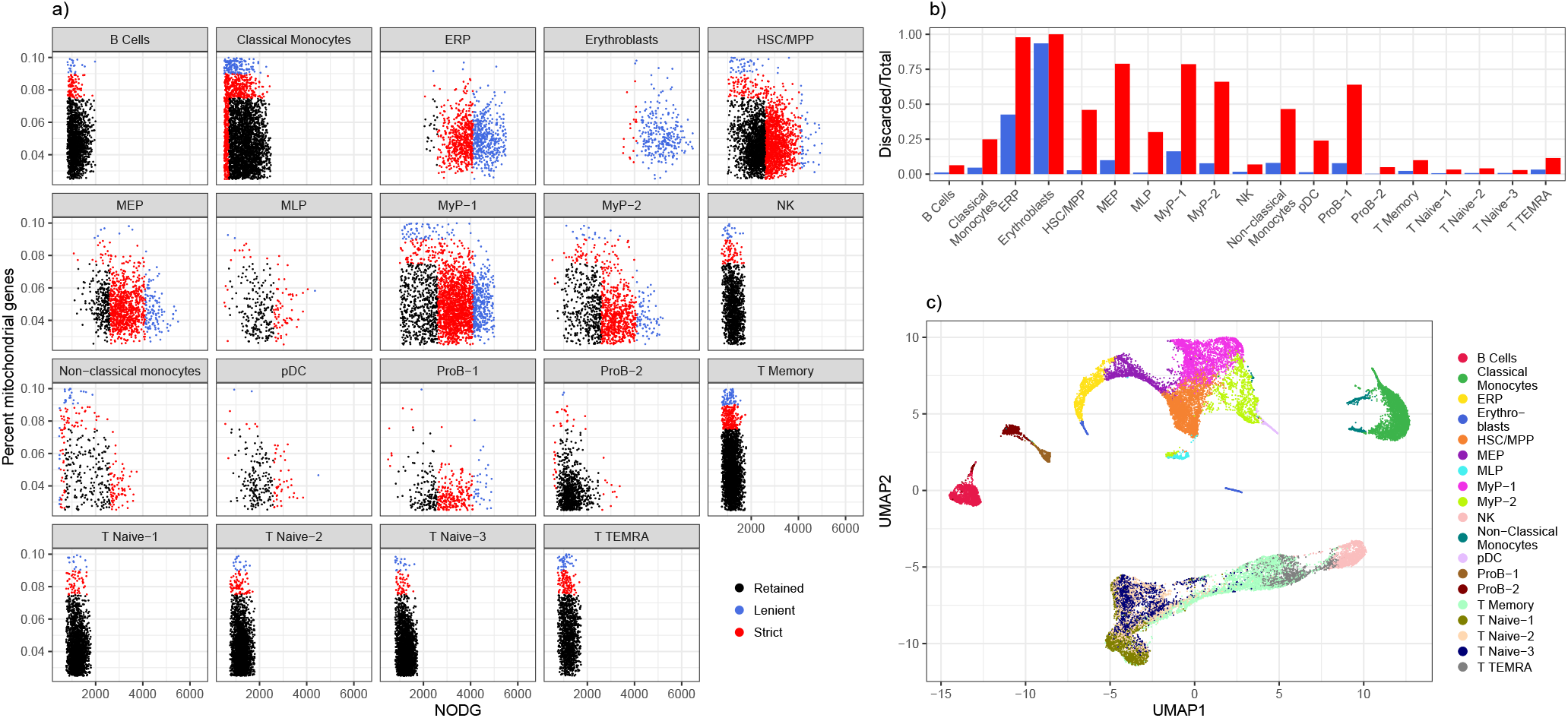
a) Cluster-specific QC based on NODG and pMito. Colors indicates whether cells are discarded (red, blue) or retained (black) if general, cluster-unspecific QC would be used. b) Proportions of cells discarded per cell type using cluster unspecific QC. c) UMAP reduction of a multi panel RCA2 projection coloured by cell type using a resolution of 0.5.

**Fig. 6.**
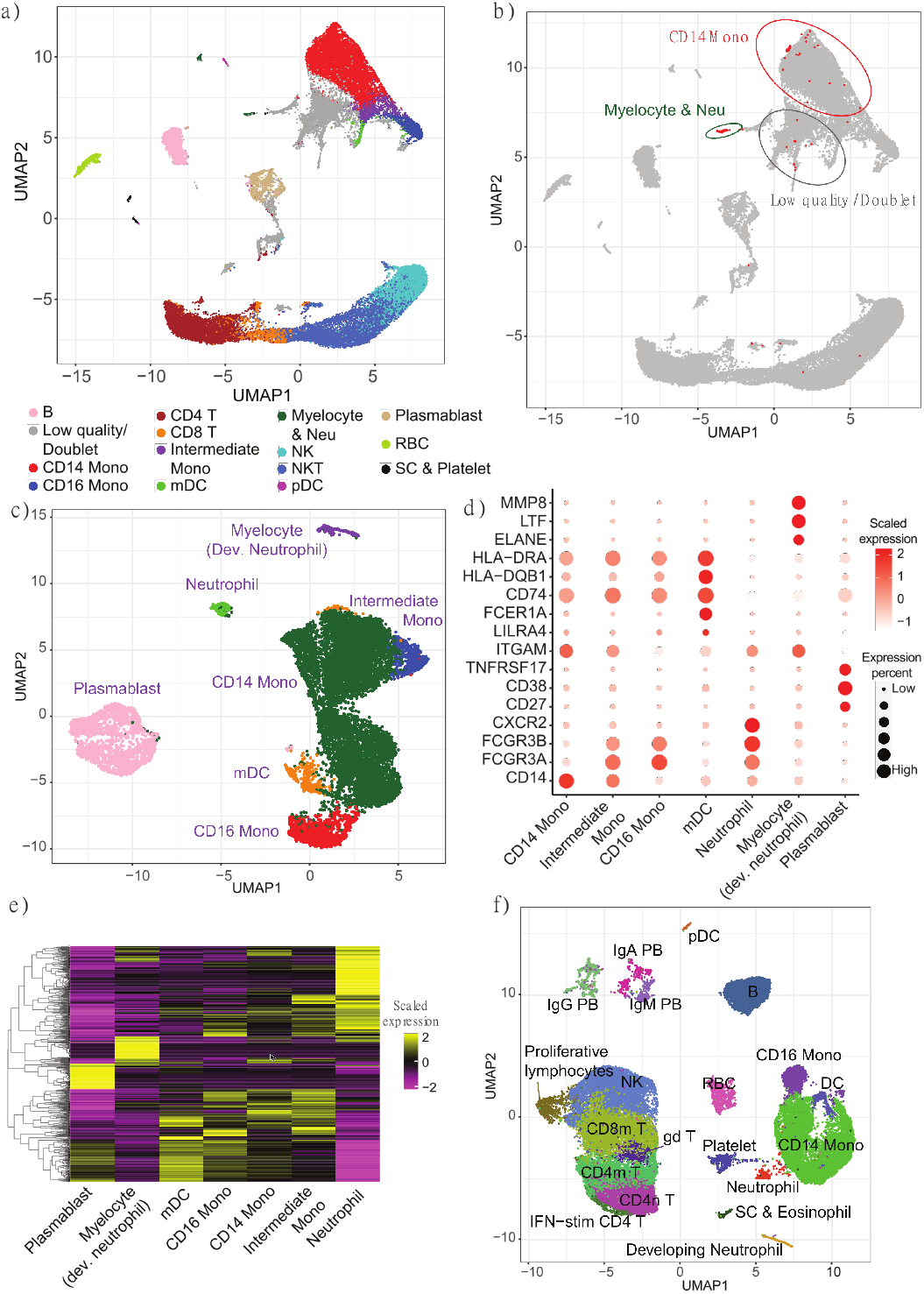
a) UMAP shows the RCA2 clustering of cells from the COVID-19 study by Wilk *et al.* b) The location of developing neutrophils annotated by Wilk *et al.* are marked as red dots in the RCA2 UMAP. c) UMAP represents the sub-clustering of CD14 monocytes, CD16 monocytes, myelocytes, neutrophils, and plasmablasts using an unsupervised approach. d) Bubble plot shows the marker gene expression levels across the identified cell types from c. Bubble size indicates expression percentage for each cell type, while color intensity represents scaled expression levels. (e)Heatmap shows the expression profiles of cell-specific genes across the identified cell types from c. f) UMAP plot shows the cell clustering using the analysis pipeline by Wilk *et al.* after removing low-quality cells and doublets.

To complement the functionality of RCA2, we incorporated SingleR/scMatch like assignment of cell types to individual cells (10, 11). Exploiting RCA2’S clustering algorithms, we also allow annotation on the cluster level (see Methods). For further biological interpretation RCA2 identifies DEG in either a *1 vs all* or a *pairwise* scheme. DEGs are visualized in heatmaps following the established Seurat color scheme (Sup. Fig. 7) and can be used as input for a GO-Term (32) enrichment and KEGG pathway (33) analysis, providing biological insights on clusters beyond lists of marker genes (Sup. Fig. 8).

**Fig. 7.**
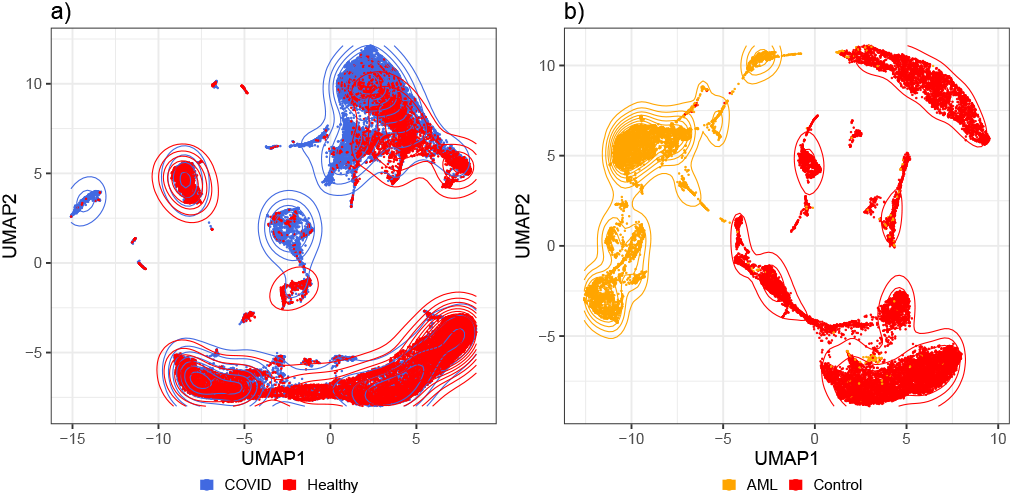
a) UMAP embedding of a reference projection for the Covid-19 PBMC data set from (22). b) UMAP embedding of a reference projection for the AML dataset 809653 from (23). AML and control cells are well separated in reference space.

### Supervised clustering is robust to batch effects

On of the major advantages of supervised clustering is its ability to reduce the contribution of unwanted variation, which manifests in the form of noise or technical variation. This is crucial in the prevention of batch effects. By projecting sc data onto a reference panel of purified transcriptomes, supervised clustering is able to preserve only the cell type-specific variation and ignore variation from other sources. This is based on the concept shown in Sup. Fig. 9, i.e. the batch effect expression signature will be orthogonal to the signature of cell type marker genes, because any two randomly selected vectors are orthogonal in a high dimensional space.

We benchmarked RCA2s robustness to batch effects by comparing its performance against several other commonly used scRNA-seq clustering algorithms: Seurat, Seurat with SCTransform, MultiCCA, MultiCCA with SCTransform, scran, Scanpy, MNN Correct and Scanorama.

The benchmarking relies on SI values to determine clustering quality with respect to batches and cell types (28). A robust method has a low SI in separating batches and a high SI in separating cell types.

### Rheumatoid arthritis (RA) data set benchmarking

We use a RA data sets comprised of 5, 829 sorted cells generated by (19). They detected plate-specific batch effects while clustering cells using the Seurat package, which were more pronounced in some plates as compared to others. RCA2 considerably outperforms the other tested algorithms in terms of cell type separation (Fig. 3a) but also in terms of batch robustness. While data from different plates is readily merged together (Fig. 3b) cells cluster well according to their determined cell type (Fig. 3c). We observed that using SC-Transform in the Seurat Integration workflow worsened both separation by batch and separation by cell type.

### Cite-seq data set benchmarking

Next, we considered PBMC CITE-seq data sequenced using (a) Drop-Seq and (b) 10X Chromium that were analysed together as a single dataset. CITE-Seq data contains both the antibody signature of cells and their transcriptomic profile. Thus it provides the capability of defining a ground truth to an scRNA-seq dataset using antibody-derived tags (ADTs). We used Seurat to cluster SCs in the antibody space in each of the CITE-seq datasets and used the unique ADT signature of each cluster to annotate them (Sup. Fig. 10) (Methods).

Benchmarking on the CITE-Seq data showed that Seurat Integrated, MNN Correct and Scanorama successfully reduced batch effects, similar to RCA2. However, that was achieved at the cost of worsening cell type separation (Fig.3d). Seurat (Sup. Fig. S11), Scran and Scanpy produced clustering results significantly affected by protocol, with Scanpy performing the best of these unsupervised clustering methods without explicit batch correction in terms of cell type separation. RCA2 is among the top method in terms of batch robustness (Fig.3e) and has the best SI value for cell types leading to a crisper separation of cell types (Fig.3f).

Using ADT-tags of the CITE-seq data, we are able to characterize the batch not only from a technical perspective using the SI, but also from a biological point of view: we computed the set of DEGs that characterizes the sc capture protocol batch within each ADT cluster (Fig.4a, Sup. Table 3). While the majority of DEGs are ribosomal genes, we also find several genes that are both cell type and batch specific markers, such as *H3F3A*, *IGKC*, *IFI30*, or *IGLC2*. The latter three are related to immune reaction and gamma-interferon signaling. Another interesting gene that is associated to the batch is *FOS*, which has been associated to several molecular processes and has been linked to cancer progression (34). As it is known that the expression of *FOS* can be easily changed by external stimuli (35), it might be more likely that, in our data set, the observed differences in *FOS* expression are of technical instead of biological nature. Indeed, the RCA2 projection, shown in Fig. 4b, is not affected by any of the DEGs linked to the batch effects and is supporting the antibody based clustering well. The latter is also backed up by the expression of the genes targeted by the antibodies (Sup. Fig. 12).

To characterize the genes defining the observed batch further, we investigated their GO term enrichment separately for genes expressed in the 10X batch (Sup. Fig. 14) and the Drop-seq batch (Sup. Fig. 15). All of the observed GO terms can be exclusively explained by the difference in sequencing protocol and therefore potentially misguide down-stream analysis, if less robust clustering methods are used.

We investigated this hypothesis by computing cluster specific marker genes for clusters identified with various scRNA-seq pipelines, using the same parameters and settings. We compared those method specific DEG to the set of batch specific DEGs, shown in Fig.4a, using an Upset plot (Sup. Fig. 15). Indeed, RCA2 shows the lowest overlap between cluster specific marker genes and the set of batch DEG compared to the other methods, underlining the robustness of RCA2 towards batch effects further.

### Use-case on 10X PBMC data set comprising 5000 cells

We obtained a 10X dataset containing 5025 Peripheral blood mono nuclear cells (PBMCs) from a healthy donor. We imported the CellRanger output directly into RCA, considering cells with an UMI count ≥ 100.

Upon QC using RCA2’s QC functionality (Sup. Fig. 15, Sup. Table 1), the scRNA-seq data is projected against a new, manually curated reference panel of immune cell types, based on purified populations of human hematopoietic cells (16). We utilized the integration of Louvain graph based clustering algorithm into RCA2 to cluster the data using a resolution of 0.1, which leads to a sensible separation of cells in terms of the projection heatmap (Sup. Fig. 16) as well as in the resulting number of clusters (Sup. Fig. 4). We obtained nine clusters forming four disconnected islands in a UMAP based on the cell to cell type correlation space obtained by the reference projection (Sup. Fig. 17a).

RCA’s automated cluster annotation function determines cell types, shown in Sup. Fig. 17b. While the B-cell cluster (red) is very distinct from all remaining clusters, T and natural killer cells form a continuum (blue, brown, yellow, green). Monocytes (turquiose) and non-classical monocytes (black) appear to be well separated within a major myeloid cluster. In close proximity in UMAP space, small populations of myeloid (pink) and plasmacytoid (magenta) dendritic cells were identified. While the automatically determined labels agree with the projection heatmap shown in Sup. Fig. 17a, this can be expected by the design of RCA2 and the projection step. Therefore, we use canonical marker genes, which are used for instance also in the Seurat tutorial, to verify cell types. As shown in Sup. Fig. 17c-j, the abundance of the various marker genes corresponds well to the identified clusters.

To further characterize the clusters, we compute DEGs in a pair-wise manner (Sup. Fig. 7b). Sup. Table 4 lists all identified DEGs. As shown in Sup. Fig. 18, we obtain several significant terms in GO term analysis using these DE genes for the natural killer cell cluster including *cytolysis*, *cell killing* and *cellular defense response*. These are well matching to the expected biological function of natural killer cells. Also, for the naive CD4 T-cell cluster we obtain sensible terms such as *adaptive immune response*, *immune response-activating cell surface receptor signaling pathways*, and *activation of immune response* (Sup. Fig. 19).

This example illustrates that RCA2 allows a hassle-free analysis to characterize clusters with minimal manual efforts. The example can be reproduced by following the online tutorial on our github page.

### Cluster specific quality control is essential to retain high-quality cells in complex data sets

Here, we consider four human bone marrow specimens separated into CD34+ and CD34-fractions (Methods). Clustering the RCA2 reference projection using Louvain clustering with a resolution of 0.1, we find ten clusters representing major cell types (Sup. Fig. 20). With RCA’s cluster specific QC function, we observed that the various cell types included in the dataset do require different QC thresholds (Sup. Fig. 21) For instance, the average NODG for the lymphoid population, e.g. B or T cells, is around 1, 000, while the NODG of progenitor cells can be up to three fold higher. Similarly, the percentage of mitochondrial reads shows different distributions. While it has low standard deviation for Pro-B cells, it values spread out widely e.g. for Classical Monocytes. As indicated by the color code in Fig. 5a, cluster agnostic thresholds (Sup. Fig. 22) would result in a substantial loss of cells, which is quantified for the final clusters in Fig. 5b and clearly illustrates the importance of a (major) cell type specific QC. Final cell type specific QC thresholds are listed in Sup. Tab. 2.

Upon QC on the level of major cell types, we used RCA2 to define cell types on a more detailed level. Using Louvain clustering, we found the most convincing clustering in terms of the projection heatmap using a resolution of 0.5. Doublets have been removed at this stage with DoubletFinder (21) using the 0.97 quantile of all *pANN* values as a threshold, resulting in the identification and removal of 906 doublets (Sup. Fig. 23). Final cell type annotations, based on projection scores (Sup. Fig. 24) and backed up with DEGs (Sup. Fig. 25, Sup. Table 5) as well as canonical markers (Sup. Fig. 26), are indicated in the UMAP representation of the RCA2 projection shown in Fig. 5c.

We separated the bone marrow data into two populations using magnetic bead selection as cells that are either positive or negative for the progenitor marker CD34 (36). As shown in Sup. Fig. 27, the RCA2 reference projection based UMAP of the scRNA-seq data shows distinct levels of CD34 FACS labels. These match well to the identified cell types shown in Fig. 5c.

For example, hematopoietic stem/progenitor clusters (HSPC), i.e. HSC/MPP, LyP-1, ERP, MEP, MyP-1 and MyP-2, representing progenitor populations are almost completely composed of cells with a CD34+ FACS label, while clusters such as B cells or Classical Monocytes, that are composed of differentiated cells are enriched for cells with a CD34-label (Sup. Fig. 28a). However, we note that some clusters like naive T cells and non-classical monocytes also had a small contribution of 10 − 15% from cells labelled as CD34+ cells by our MACS sorting strategy. This is not unexpected because our workflow for purifying HSPCs lacks a prior conventional lineage-depletion (lin-) step.CD34+ populations within the bone marrow are known to be heterogeneous and lin-CD34+ populations were shown to mainly harbour stem cell activity (37). Hence, our MACS sorting strategy is expected to deliver false positives in the form of cells that are retained in the CD34+ magnetic columns but are in reality differentiated cells. However, RCA2 is able to identify such lineage+CD34+ cells and clusters them correctly based on their transcriptome. Thus, RCA2 offers a more precise in-silico alternative to the conventional lineage-depletion step for HSPC studies. Compared to an analysis with Seurat using default parameters (Sup. Fig. 28b), RCA2 achieves a better purity: In 10.05% of RCA2 clusters, the impurity is > 20%, compared to 19.05% using Seurat. The validity of our approach is also supported by the fact that cell type proportions are in agreement with earlier studies (Sup. Fig. 29).

This example illustrates the ability of RCA2 to seamlessly derive meaningful annotations and dimensionality reductions even in highly complex datasets where cells are placed in a continuum and the reference set might not contain exactly matching cell types.

### RCA2 clusters PBMCs from COVID-19 patients more robustly than de-novo approaches

Using scRNA-seq PBMC data obtained from seven COVID-19 patients and six healthy donors, Wilk *et al.* reported a *developing neutrophil* (DN) population that seemingly showed a phenotypic relation with plasmablasts in dimensionality reduction (22). Wilk *et al.* hypothesized *a lymphocyte-to-granulocyte differentiation process* to be associated with severe COVID-19 infection. However, as raised by (38), this UMAP based interpretation contradicts the principle cell lineage fate. By projecting all of 44, 721 cells analyzed by Wilk *et al.* to immune reference panels in RCA2, we identified that a large portion of cells were actually low quality cells or potential doublets (Fig.6a).

We identified 32 clusters using graph-based clustering (resolution 1.3) on the reference projection of all 44, 721 cells (Sup.Fig. 30, 31a). Seven clusters (darkorange, darkred, green, greenyellow, lightcyan and lightgreen) exhibit ambiguous projection profiles showing high correlations with multiple cell types from distinct lineages, suggesting that these cells are either of low quality or doublets (Sup. Fig. 31b). Additional marker gene profiling coupled with a NODG comparison supported this hypothesis (Sup. Fig. 31c,d). For instance, clusters darkred and darkorange had high correlations with monocytes and T cells, respectively (Sup. Fig. 31b). However, both of them also showed high transcriptomic correlations with red blood cells (RBC), implying that these cells could be contaminated by RBC (Sup. Fig. 31b). We observed high expression levels of RBC genes, such as GYPA and HBB, in these two clusters (Sup. Fig. 31c) and significantly lower NODG of these two clusters comparing to monocytes and T cells respectively (Sup. Fig. 31d).

Next, we pinpointed the location of the 206 *developing neutrophils* (DN) annotated by Wilk *et al.* within our RCA2 UMAP, and found these cells to be a mixture of different cell types (Fig. 6b). Similar to the initial SingleR annotation for the *DN* cluster made by Wilk *et al.*, 131 cells were positioned within the *Myelocyte & Neu* group, which consistently showed high expression of premature neutrophil granule genes in our RCA2 analysis (Sup. Fig. 32a). The remaining 75 cells were either low quality/doublets (32 cells) or other cell types (Sup. Fig. 32b). Particularly, 29 *DN* were annotated as CD14 monocytes by RCA. In order to ascertain that the RCA2 approach did correctly cluster and annotate these *DN*, we compared the RCA2 reference projection profiles and feature genes between the 131 and 29 *DN* annotated as myelocytes and CD14 monocytes in the RCA2 analysis respectively (Sup. Fig.32c,d). Indeed, the former 131 *DN* showed high correlation with the transcriptome reference of myelocytes, while the latter 29 *DN* were highly correlated with CD14 monocytes with high expression of the marker genes such as CD14 and CD300E (39)) (Sup. Fig. 32d). Therefore, RCA2 has precisely segregated these *DN* into their real cell types.

Interestingly, we did not observe the phenotypic relation by dimensionality reduction between RCA-annotated myelocytes and plasmablasts as described by Wilk *et al.* (Fig.6a). In order to assess the transcriptomic similarity between myeloid-derived myelocytes and lymphoid-derived plasmablasts, we charted the UMAP embedding and compared the DEGs across these cell types (Fig.6c). As comparison control, we also included myeloid-derived monocytes and DCs in the analysis (Fig.6c). As expected, myelocytes and plasmablasts did not form a *differentiation bridge* between them, and a clear difference at the gene-expression level were characterized between these two cell types (Fig.6c-e). Presumably, the initial findings of the *differentiation bridge* between *DN* and plasmablasts by Wilk *et al.* could be due to the presence of low-quality cells and doublets. Indeed, after removing these sub-optimal cells and applying the same analysis pipeline used by the authors, we observed distinct separation between *DN* and plasmablasts (Fig.6f). Altogether, RCA2 has provided a more precise clustering and UMAP visualization of PBMCs from this COVID-19 study.

### Reference based clustering is able to capture disease states of cells

To address a prevalent misconception that supervised clustering algorithms are unable to identify novel cell types and cell states, we used RCA2 to project and to cluster two publicly available data sets: one Acute Myeloid Leukemia (AML) data set (23) as well as the already introduced Covid-19 dataset (22). Note that no additional QC was performed and data is used as provided by the authors. We refer to the Methods section for further processing details.

As shown in Fig. 7a, we observe that PBMCs from the Covid-19 data occur both in condition specific and in shared neighbourhoods. This is an expected behaviour and corresponds well to the original findings of (22). Upon clustering the data in RCA, we obtained 28 clusters. As shown in Sup. Fig. 33a, several clusters are depleted for cells from Covid-19 patients, whereas five clusters are composed of more than 75% of cells from Covid-19 patients, despite no disease specific reference cell types are included in our panels.

For the AML data set we obtain a clearer picture. According to the authors classification of cells, AML and healthy cells separate almost perfectly in the RCA2 projection although no AML samples are included in the reference panels (Fig.7b.). This separation is also reflected in the cluster composition plot (Sup. Fig. 33b).

## DISCUSSION AND CONCLUSIONS

Although *de novo* clustering is currently the predominant strategy to cluster scRNA-seq data, it does have some disadvantages, the most important of which is vulnerability to batch effects. Consequently, *de novo* clustering necessitates use of explicit batch-effect correction (4). However, one fundamental problem with batch correction is that it cannot distinguish between technical variation and genuine biological differences. Hence, when batch and biology are confounded, there is a risk of erroneously suppressing biological variation. Since reference-based methods can mitigate this problem, mapping of SCs to a reference atlas has recently been identified as one of the grand challenges in the SC field (40). RCA2 directly addresses this challenge.

Indeed, our benchmarking of batch effect robustness supports the above expectation. In two independent benchmarks, RCA2 was the best performer in clustering cells by cell type rather than batch, even without explicit batch correction (Fig. 3,4). Consistently with this finding, DEGs between clusters reflected cell type identity in the case of RCA2, but batch effects in the case of *de novo* clustering. Importantly, in addition to being robust to batch effects, RCA2 is able to detect cell types and states not present in the reference panel (Fig. 5,7). This capability of RCA2 implies that novel cell states could potentially be discriminated even when projected onto reference transcriptomes.

One inter-operability advantage of RCA2 is that count matrices from Seurat can be imported. In return, RCA2 results can be incorporated into a Seurat object. In terms of scalability, RCA2 is a key improvement over the initial release: RCA2 memory usage grows linearly with the number of cells, unlike the quadratic scaling of RCA (Fig. 2). Also, execution time is over ten-fold faster on large datasets. In addition, RCA2 incorporates multiple new reference panels for human and mouse and also supports generation of new panels from user-supplied transcriptome data. RCA2 also provides multiple features for data visualization and interpretation, such as generation of editable (ggplot2) figures, KEGG and GO enrichment analysis (Fig. 1). Lastly, RCA2 simplifies cluster-specific QC, which is essential for discarding low-quality cells and doublets in SC data from heterogeneous samples (Fig. 5).

QC can have a severe effect on clustering, particularly *de novo* clustering. Indeed, our re-analysis of a recently published COVID-19 dataset showed that known cell type markers showed clear cluster-specific expression only after QC using RCA2 (Fig. 6d). In contrast, reference-based clustering using RCA2 was robust to the presence of low-quality cells and doublets (Sup. Fig. 31). Importantly, the “developing neutrophils” identified as plasmablast-derived in the previous study formed a novel cluster after QC that was clearly distinct from plasmablasts.

Overall, we demonstrated that reference-based clustering of scRNA-seq data has unique advantages and provides a complementary strategy to widely-used unsupervised approaches. RCA2, which is freely available on github (https://github.com/prabhakarlab/RCAv2), provides the single-cell community with a robust, scalable and easy-to-use R-package that can be easily integrated into existing workflows to leverage these advantages. We will continue to maintain and enhance RCA2, for example by expanding the set of reference panels and by adding more clustering strategies downstream of reference projection. Given the potential of reference-based methods for SC data analysis, we believe that such methods may in future also prove useful in analyzing multi-modal SC data.

## ACKNOWLEDGEMENTS

The authors thank all members of the Prabhakar lab for testing and providing feed-back on the functionality and user-friendliness of RCA2.

