## Supplementary Material for "Robust clustering and interpretation of scRNA-seq data using reference component analysis"

Supplementary Information

### RCA2 – An improved framework for reference component analysis of scRNA-seq data.

#### 1. Available reference panels in RCA2

In RCA2, we provide the following panels:

a) Global panel from the original RCA (Li et al., 2017) containing both primary cell types and tissues.
b) ColonEpiPanel: 9 colon epithelial samples from from the original RCA (Li et al., 2017)
c) 15 hematopoietic cell types obtained from FACS sorted PBMCs (Novershtern et al., 2011): *Naive B-Cells, Mature B-Cells, Plasmacytoid Dendritic Cell, Myeloid Dendritic Cell, Hematopoietic stem cell, Megakaryocyte, Monocyte, Non-classical Monocyte, Mature NK cell_CD56+ CD16+ CD3, Naive CD8+ T-cell, CD8+ Effector Memory, CD8+ Central Memory, Naive CD4+ T-cell, CD4+ Effector Memory, CD4+ Central Memory*
d) NovershternTCellPanel: Selection of six T cell sub-types from Novershtern et al. (Cell, 2011).
e) 28 immune cell types obtained from blood samples (Monaco et al. 2019) : *Basophils, B-Ex, B-naive, B-NSM, B-SM, CD4 naive, CD8 CM, CD8 EM, CD8 naive, CD8 TE, C-mono, I-mono, MAIT, mDC, NC-mono, Neutrophils, NK, pDC, Plasma blasts, Progenitor, TFH, Th1, Th17, Th1/Th17, Th2, TReg, VD2negative, VD2positive.*
f) MonacoBCellPanel: Selection of six five B cell sub-types from Monaco G., et al. (Cell reports, 2019).
g) MonacoMonoPanel: Selection of five Monocyte sub-types from Monaco G., et al. (Cell reports, 2019).
h) MonacoTCellPanel: Selection of 15 T cell sub-types from Monaco, G., et al. (Cell reports, 2019).
i) Mus Musculus brain panel containing 7 cell types (Zhang et al. 2016): *Astrocytes, Neurons, Microglia, Endothelial Cells, Newly-formed Oligodendrocytes, Myelinating Oligodendrocytes, Oligodendrocyte Precursor Cells.*
j) ENCODE homo sapiens panel containing 97 cell types generated using RCA2 build panel function: *Skin Fibroblast, Endodermal Cell, Neural Cell, T-Cell, Mesendoderm, Neural Stem Progenitor Cell, Endothelial Cell Of Umbilical Vein, Fibroblast Of Skin Of Abdomen, CD4-Positive, Alpha-Beta T Cell, Purkinje Cell, Fibroblast Of Breast, Smooth Muscle Cell, Trophoblast Cell, Common Myeloid Progenitor, CD34-Positive, Mesenchymal Cell, CD8-Positive, Alpha-Beta T Cell, Foreskin Keratinocyte, Mammary Stem Cell, Fibroblast Of Lung, Foreskin Fibroblast, Neurosphere, Myoepithelial Cell Of Mammary Gland, Skeletal Muscle Myoblast, Luminal Epithelial Cell Of Mammary Gland, Mesodermal Cell, Keratinocyte, B Cell, CD14-Positive Monocyte, Peripheral Blood Mononuclear Cell, Ectodermal Cell, Foreskin Melanocyte, Hepatocyte, Cardiac Muscle Cell, Bipolar Neuron, Natural Killer Cell, Astrocyte, Mammary Epithelial Cell, Aortic Smooth Muscle Cell, Epithelial Cell Of Umbilical Artery, Smooth Muscle Cell Of The Umbilical Artery, Tracheal Epithelial Cell, Nasal Cavity Respiratory Epithelium Epithelial Cell Of Viscerocranial Mucosa, Smooth Muscle Cell Of The Coronary Artery, Thoracic Aorta Endothelial Cell, Regular Cardiac Myocyte, Dermis Microvascular Lymphatic Vessel Endothelial*

*Cell, Dermis Lymphatic Vessel Endothelial Cell, Bronchial Smooth Muscle Cell, Endothelial Cell Of Coronary Artery, Mesenchymal Stem Cell Of Whartons Jelly, Smooth Muscle Cell Of The Pulmonary Artery, Bronchus Fibroblast Of Lung, Endometrial Microvascular Endothelial Cells, Uterine Smooth Muscle Cell, Smooth Muscle Cell Of Trachea, Melanocyte Of Skin, Mammary Microvascular Endothelial Cell, Bronchial Epithelial Cell, Hematopoietic Multipotent Progenitor Cell, Bladder Microvascular Endothelial Cell, Fibroblast Of The Aortic Adventitia, Mononuclear Cell, Fibroblast Of Villous Mesenchyme, Placental Epithelial Cell, Osteoblast, Mesenchymal Stem Cell Of Adipose, Skeletal Muscle Satellite Cell, Cardiac Atrium Fibroblast, Epithelial Cell Of Alveolus Of Lung, Kidney Epithelial Cell, Vein Endothelial Cell, Renal Cortical Epithelial Cell, Myotube, Smooth Muscle Cell Of Bladder, Myometrial Cell, Neural Progenitor Cell, Pulmonary Artery Endothelial Cell, Hair Follicle Dermal Papilla Cell, Fibroblast Of Dermis, Lung Microvascular Endothelial Cell, Myocyte, Dermis Blood Vessel Endothelial Cell, Placental Pericyte, Articular Chondrocyte Of Knee Joint, Mesenchymal Stem Cell Of The Bone Marrow, Cardiac Ventricle Fibroblast, Airway Epithelial Cell, Epithelial Cell Of Proximal Tubule, Subcutaneous Preadipocyte, Pericardium Fibroblast, Mesangial Cell, Hair Follicular Keratinocyte, Glomerular Endothelial Cell.*

k) ENCODE Mus Musculus panel containing 15 cell types: *Erythroblast, Megakaryocyte, Erythroid Progenitor Cell, Megakaryocyte Progenitor Cell, Megakaryocyte-Erythroid Progenitor Cell, Granulocyte Monocyte Progenitor Cell, Hematopoietic Multipotent Progenitor Cell, Common Myeloid Progenitor, Monocyte, Neutrophil, Hematopoietic Stem Cell, Bone Marrow Macrophage, Embryonic Fibroblast, Myocyte, Leukemia Stem Cell.*

l) A CITE-seq panel based on Seurat 4.0 containing 34 cell types: *CD14 Mono, CD4 TCM, CD8 Naïve, NK, CD8 TEM, CD16 Mono, B intermediate lambda, CD4 Naïve, CD4 CTL, B naive kappa, MAIT, gdT, CD8 TCM, dnT, B intermediate kappa, B memory kappa, pDC, B naive lambda, CD8 Proliferating, Treg Naïve, Plasma, CD4 TEM, Treg Memory, cDC2, NK Proliferating, ASDC, B memory lambda, HSPC, Platelet, CD4 Proliferating, Eryth, Plasmablast, cDC1, ILC.*

Panel (a) and (b) are identical to the panel from Li et al. (2017), used in the original RCA. Panels (d-i) have been generated in the same way as panels (a) and (b) (Li et al. (2017)). Panels (j-l) have been generated using the methodology described in the Section 6.

Panel (c) was created as follows: FACS sorted microarray of 38 immune cell types was obtained from the Gene Expression Omnibus (GEO) database; accession number: GSE24759. All progenitor cells as well as cells not assigned to a fully differentiated state were excluded as they did not exhibit a distinctive transcriptome. In addition, granulocytes (including neutrophils, basophils and eosinophils) as well as denucleated erythrocytes were excluded, resulting in 15 distinct immune cell types representing PBMCs populations. After quantile normalization and *log10* transformation of the data using the *preprocessCore* R package, two rounds of feature selection were performed. First, 2383 highly variable genes, were selected to generate the distance matrix and dendrogram plot of all cell types. Here, highly variable genes are defined as genes with an expression fold change relative to mean expression across all cell types being higher than 1.8. At this stage, replicates of cell types clustered far apart from the respective cell type clusters were labelled as outliers and consequently removed from the dataset. To further refine the feature genes, a pair-wise differential expression (DE) analysis between nodes in the dendrogram was performed using a two-sided *t-test*, considering a log fold change of 2 and a q-value cut-off of 0.05 as thresholds to determine de genes.

Taking the union of DE genes across all comparisons resulted in a combined set of 966 genes as final feature genes to compile a PBMC reference panel. Importantly, the majority of these genes (955 gene) have been already detected in the first round of feature selection testing for highly variable genes. This indicates that the feature refinement step led to removal of genes which are likely to be common across sup-populations of major cell types. Therefore, removing those improved our ability to detect sub-populations within major cell types.

### 2 Supplementary Methods

#### 2.1 Build Your Own Panel

First, it generates a pseudo bulk for replicates across cell types. Next, RCA2 calculates a PCA of a spearman correlation distance matrix of cell types and clusters the PCA representation of samples considering all PCs that explain at least 1% of the variance in the distance matrix. Based on a user-provided cut-height, similar cell types are identified and differentially expressed genes with a lenient log2 fold-change cut-off (default 2) are computed within the identified clusters using the default edgeR pipeline. This approach ensures that enough DE genes are computed for highly similar cell types without adding unnecessary amounts of noise to easily distinguishable cell types. Next, the default edgeR workflow is applied to call DE genes across all cell types using a more stringent log2 fold-change cut-off (default 6). The union of genes obtained from both steps is selected to be the feature set of the reference panel, which is constructed by selecting the rows corresponding to the features and by merging replicates.

#### 2.2 Cell type annotation

From the projection matrix *P*, we identify, for each cell i, the cell type t associated to the highest score according to

$$t:=\text{index}^{P_i} (\max (P_i)),$$

where *index* returns the column index corresponding to the cell type of the projection matrix *P* holding the highest correlation for cell *i*.

We compute two optional metrics that can be visualized in UMAP plots to estimate the confidence of cell type assignments.

Firstly, we assess the confidence of the annotation for each cell individually by computing a ratio-score r between $\max(P_i)$ and $\max2(P_i)$according to:

$$r=1-\text{abs}(\max(P_i)/\max2(P_i)).$$

The value of *r* is close to 0 if the second highest score for cell *i* in P is similar to the highest score. If the value of *r* is smaller than a user-defined threshold, a cell will be labelled as *Unknown*. The ratio-score is visualized by adding transparency to the UMAP plots: The closer *r* is to 0 the more transparent a cell will be.

Secondly, we calculate a rank based score across all cells that are assigned to cell type *t*. Specifically, we compute for a cell *i* that has been assigned to cell type *t*, the relative rank of the cells score $P_{i,t}$ among all other cells labeled to be of cell identity *t*. The relative rank score is visualized by a color gradient.

#### 2.3 GO and KEGG enrichment

By default, we consider cluster-specific background sets composed of genes expressed within each cluster. Alternatively, all genes expressed across all clusters or simply all genes available in the used annotation can be considered.

Several options are available to distinguish expressed from not expressed genes: a numeric threshold, the 1st quartile, the mean, the median or the 3rd quartile of the distribution of mean gene-expression values within one or across all clusters. For each tested cluster, RCA2 generates barplots, dotplots and GoPlots (if applicable).

#### 2.4 Experimental details bone marrow data

Briefly, bone marrow aspirates were drawn from the posterior iliac crest of donors using standard operating procedures.

Mononuclear cells (MNCs) were isolated using density gradient separation using standard protocols.

Frozen BMMNCs were rapidly thawed at 370-degree C in RPM1 medium supplemented with 10% FBS and containing DNAse1 (50,000 units) (catalog#260913, Calbiochem) to prevent cell clumping.

To isolate viable cells, BMMNCs were subjected to dead cell removal using a dead cell removal kit (catalog no. #130-090-101, Miltenyi biotec, Germany). Next, positive selection for CD34-positive cells was performed using the direct CD34 progenitor cell isolation kit (catalog no. #130-046-703, Miltenyi biotec, Germany).

The “flow-through” from the column was used as the CD34-negative fraction.  Only samples with higher than 80-85 percent viability were used for downstream scRNA seq analysis.

Before loading the samples for scRNA sequencing into the Chromium system (10X Genomics), the single-cell suspension of CD34 positive and CD34 negative BMMNCs were re-counted and assayed for viability by trypan-blue exclusion principle based methodology. Cells were barcoded and UMI resolved single-cell sequencing libraries were generated using Chromium Single Cell 5' Reagent Kit (PN-1000006). We targeted 6,000 and 10,000 cells for CD34 positive and negative cell fractions, respectively.

We followed the instructions as listed in the manufacturer’s product manual to generate an emulsion of cells entrapped along with barcoded beads.

The bead emulsion was transferred into a PCR strip and subjected to 13 cycles of cDNA synthesis. Subsequently, the synthesized cDNA was cleaned up with AMPure XP beads (A63881) and 25ng of cDNA were used for library preparation in compliance with manufacturer’s instructions. Generated libraries were further cleaned up with AMPure XP beads and the final quantity and quality of libraries was assessed on Bioanalyzer using High Sensitivity DNA Kit (5067-4626, 5067-4627; Agilent).

Each library was sequenced on one lane of a HiSeq4000, targeting 70,000-100,000 reads per cell.

#### 2.5 Benchmarking of runtime and memory consumption

We benchmarked RCAv2 against RCAv1 with respect to the runtime of the projection and memory consumption using lovain graph based clustering. We obtained the original RCA implementation from GIS-SP-Group/RCA: R package for robust clustering of single cell RNA sequencing data (github.com). The benchmarking is carried out on a large in-house single cell data set, which is downsampled to 1000, 2000, 5000, 10000, 20000, 50000, 100000 and 200000 cells, respectively. We use a notebook equipped with an Intel(R) Core(TM) i9-9980HK CPU @ 2.40GHz, 2400 Mhz, 8 Core(s), 16 Logical Processor(s) and 64GB memory for the benchmarking experiments.

Upon downsampling we project the scRNA-seq data sets using the dataProject() function with default parameters. We use the system.time() function in R to measure the time used for computation and report the time from the *user* slot. Memory consumption of the dataSClust function is assessed with the profmem() function from the profmem library and the total sum of allocated bytes (excluding NAs) is reported. The same functions, system.time() and profmem(), are used to benchmark RCAv1, which was used using default parameters.

#### 3. Supplementary Tables and Figures

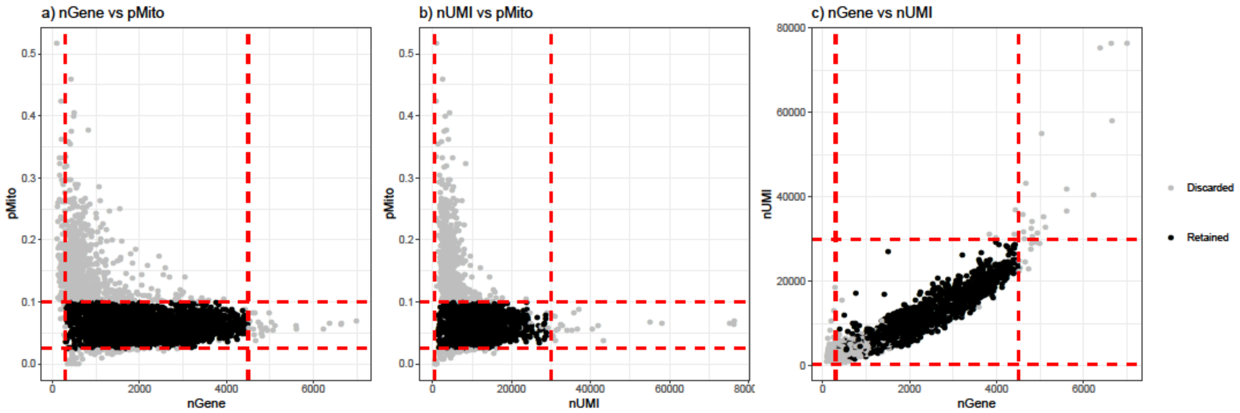


**Supplementary Figure 1:** Illustration of RCA2 quality control metrics and scatter-plots. As a quality metric, the number of detected genes (nGene), the mitochondrial rate (pMito) and the number of unique molecular identifiers (nUMI) are used. Red lines indicate user defined thresholds, cells colored black are retained and cells colored grey are discarded upon quality control.

**a)**

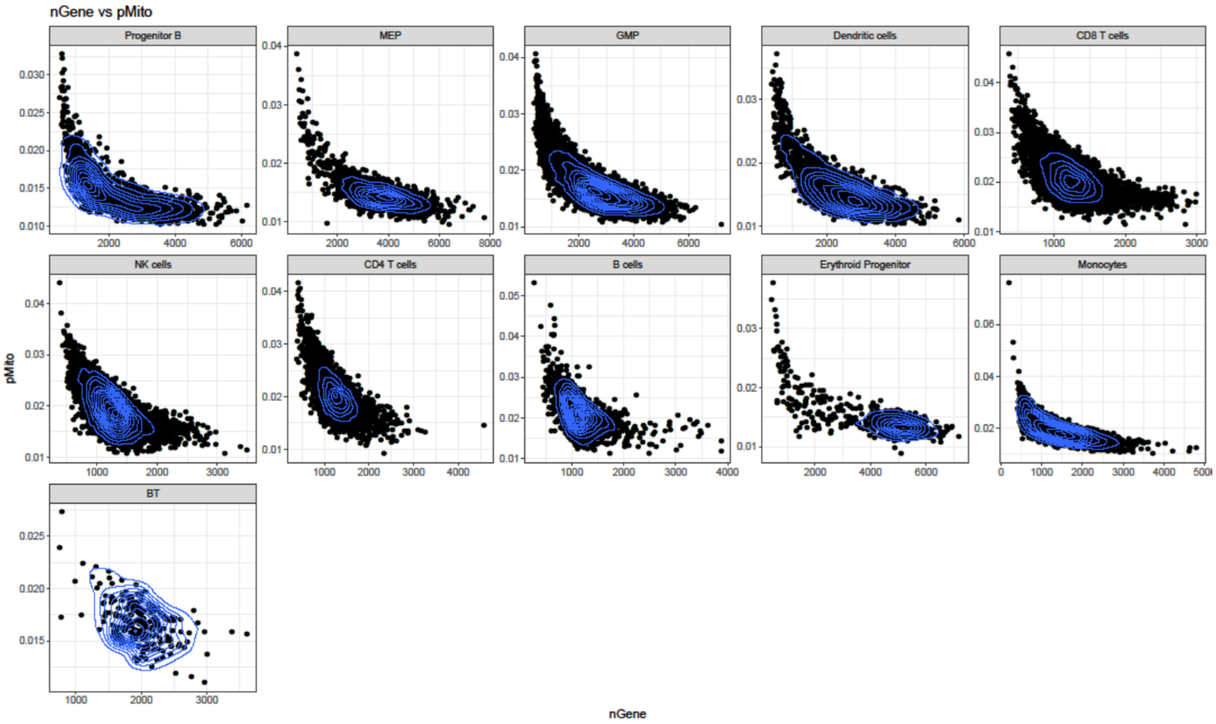


**b)**

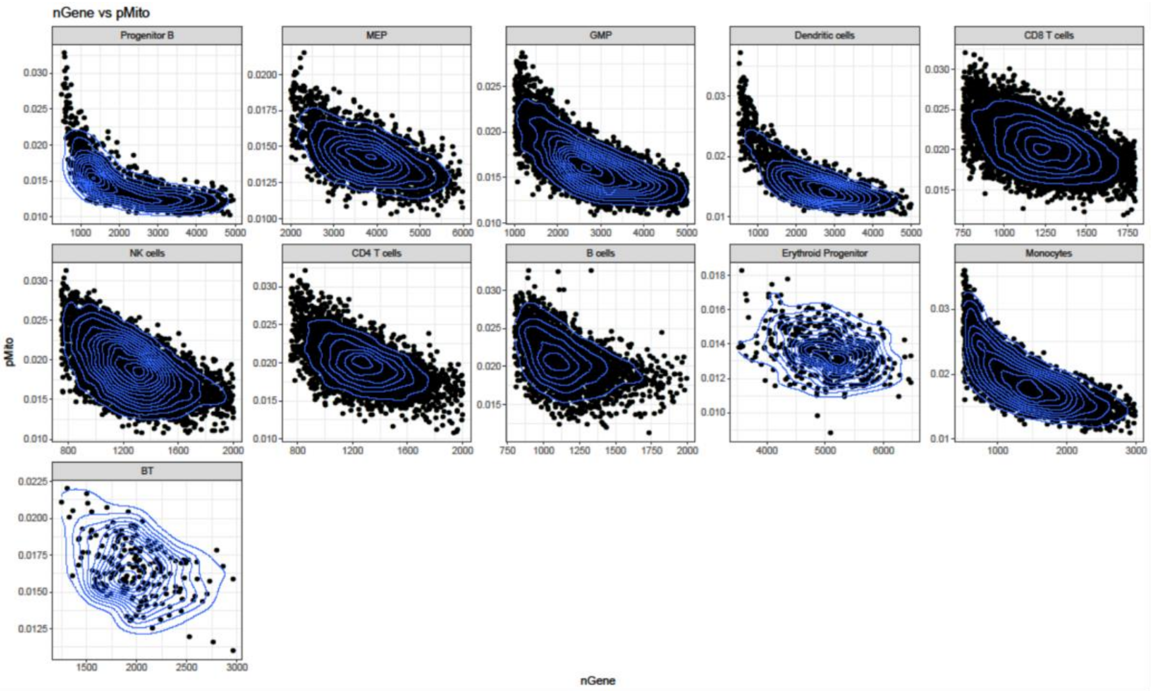


**Supplementary Figure 2**: Illustration of major cell type specific quality control information (number of detected Genes (nGene) versus percent Mitochondrial rates (pMito) calculated **a)** before and **b)** after applying QC-Thresholding using an in-house normal bone marrow data set.

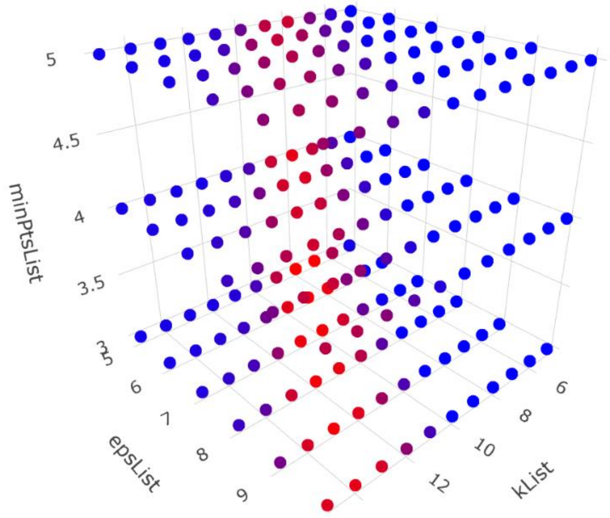


**Supplementary Figure 3**: Three dimensional visualization of the parameter space used for graph based clustering as implemented in *dbscan*. Color code and mouse over events indicate the resulting number of clusters.

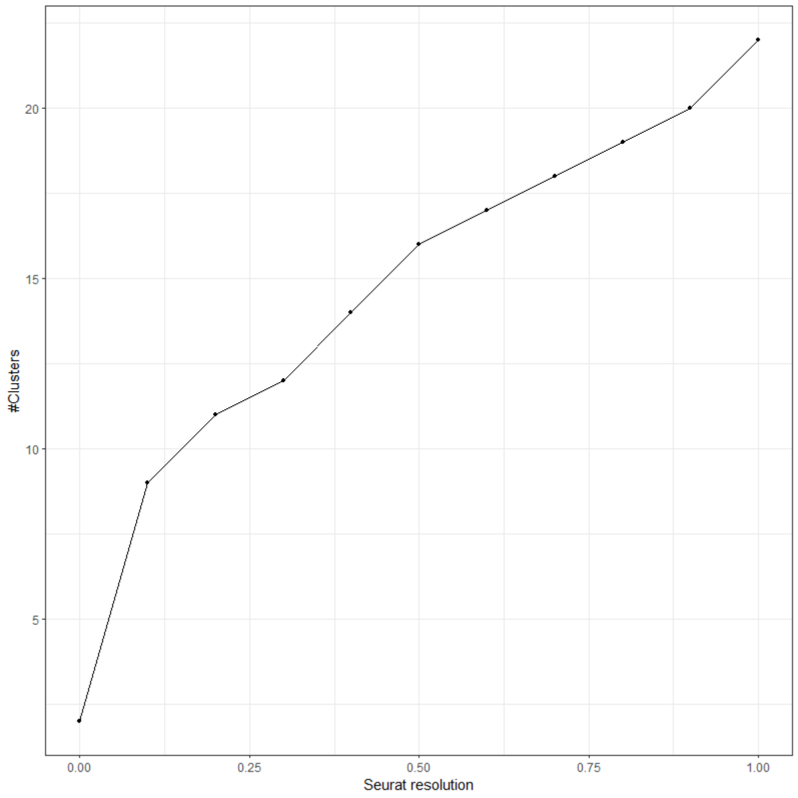


**Supplementary Figure 4:** Overview on #clusters depending on the selected Seurat resolution generated on the 10X PBMC example data set.

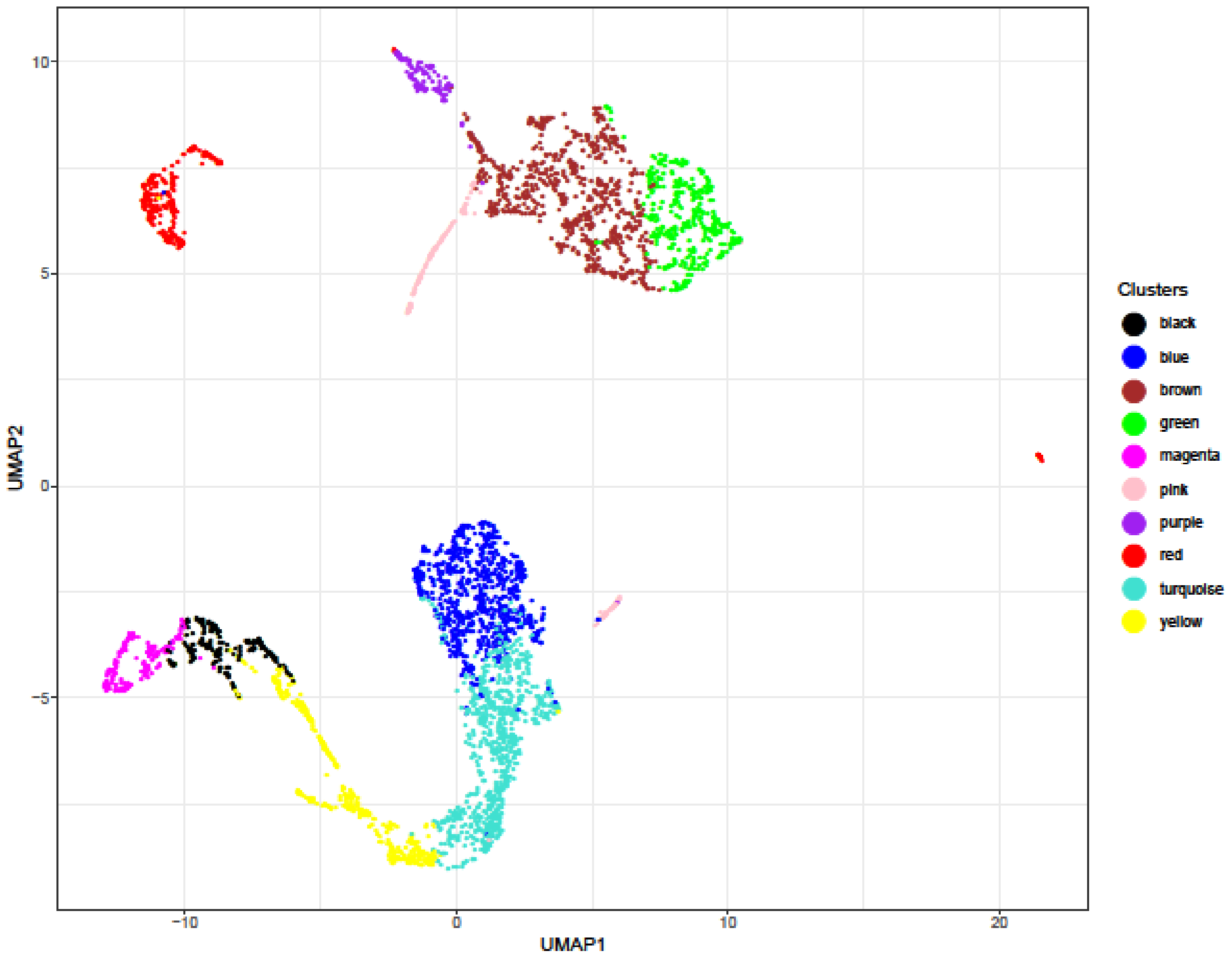


**Supplementary Figure 5:** Example of a 2D UMAP reduction of a RCA projection colored by identified clusters using the 10X PBMC example data set.

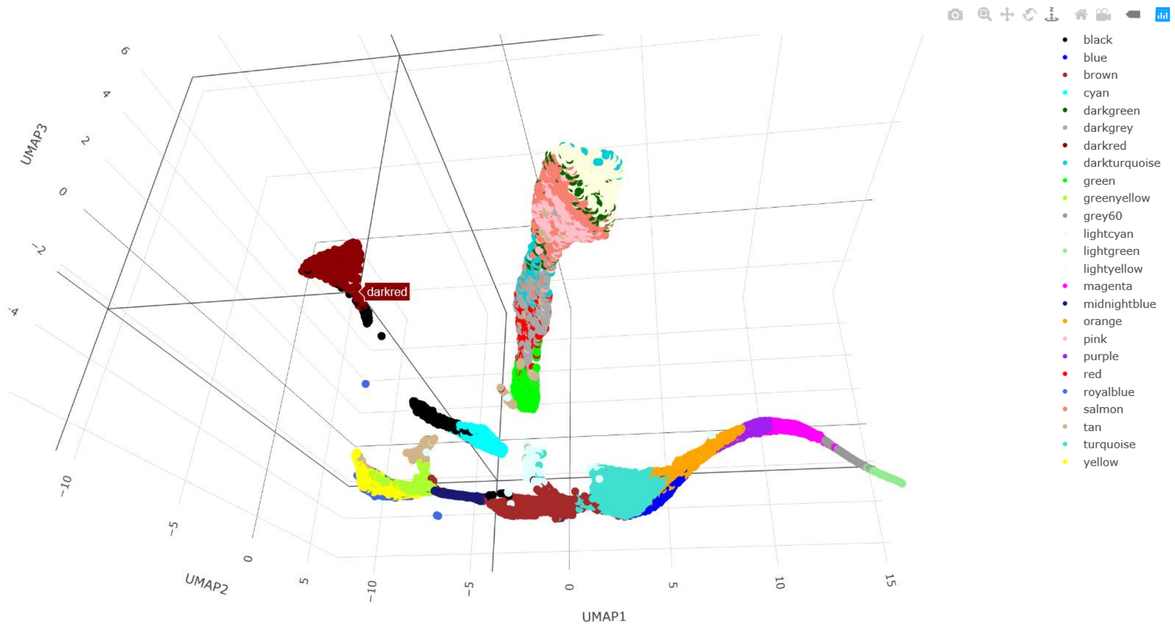


**Supplementary Figure 6:** Reduction of the reference projection to three dimensions using a UMAP on the normal bone marrow data set. Mouse over events as well as the color code allow the user to learn about cell labels and characteristics. The HTML version of the Figure can be interactively viewed and modified.

a)

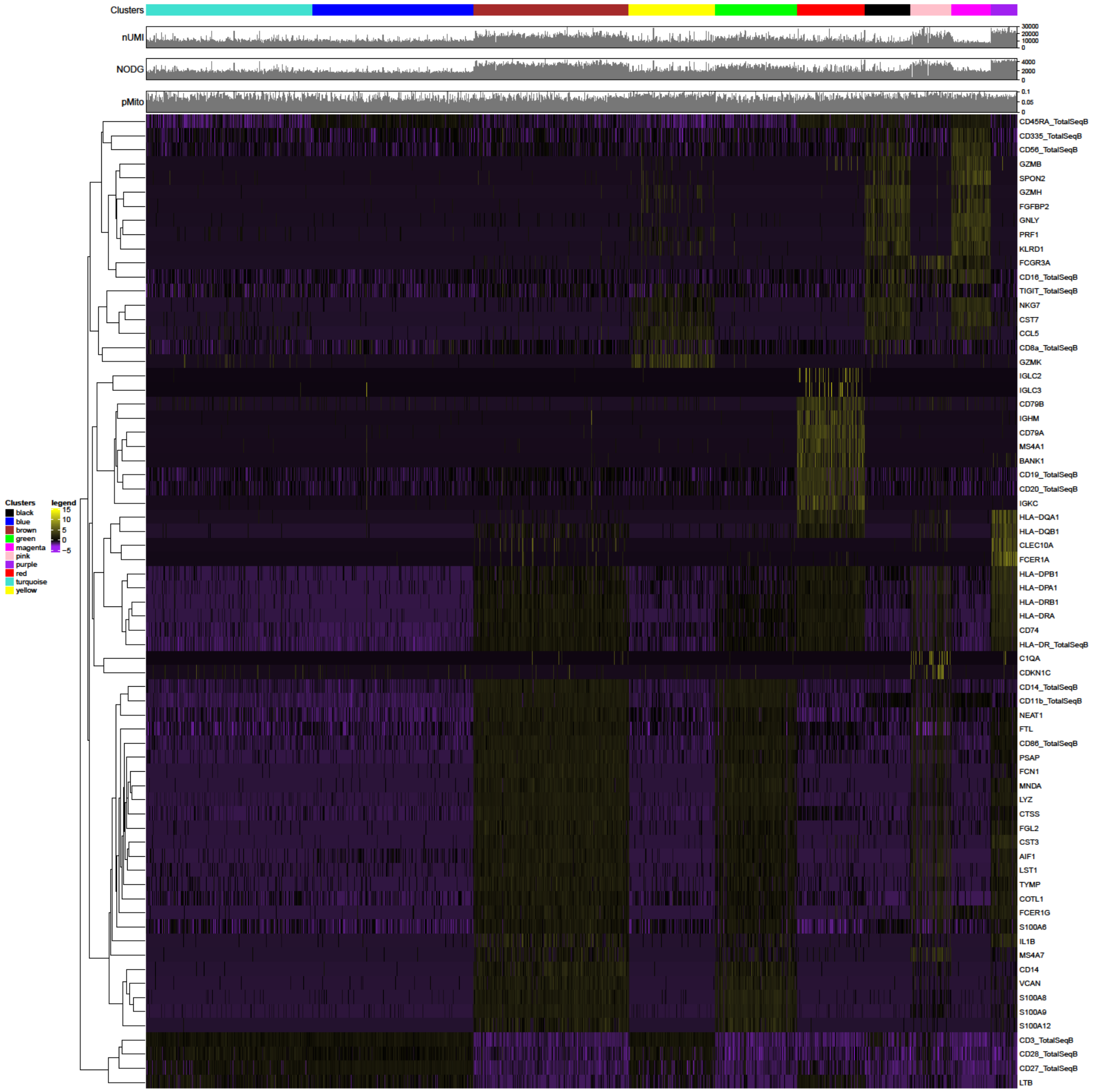

Clusters
nUMI
30000
20000
10000
0
NODG
4000
2000
0
pMito
0.1
0.05
0
CD45RA_TotalSeqB
CD335_TotalSeqB
CD56_TotalSeqB
GZMB
SPON2
GZMH
FGFBP2
GNLY
PRF1
KLRD1
FCGR3A
CD16_TotalSeqB
TIGIT_TotalSeqB
NKG7
CST7
CCL5
CD8a_TotalSeqB
GZMK
IGLC2
IGLC3
CD79B
IGHM
CD79A
MS4A1
BANK1
CD19_TotalSeqB
CD20_TotalSeqB
IGKC
HLA−DQA1
HLA−DQB1
CLEC10A
FCER1A
HLA−DPB1
HLA−DPA1
HLA−DRB1
HLA−DRA
CD74
HLA−DR_TotalSeqB
C1QA
CDKN1C
CD14_TotalSeqB
CD11b_TotalSeqB
NEAT1
FTL
CD86_TotalSeqB
PSAP
FCN1
MNDA
LYZ
CTSS
FGL2
CST3
AIF1
LST1
TYMP
COTL1
FCER1G
S100A6
IL1B
MS4A7
CD14
VCAN
S100A8
S100A9
S100A12
CD3_TotalSeqB
CD28_TotalSeqB
CD27_TotalSeqB
LTB
Clusters
black
blue
brown
green
magenta
pink
purple
red
turquoise
yellow
legend
15
10
5
0
−5

**b)**

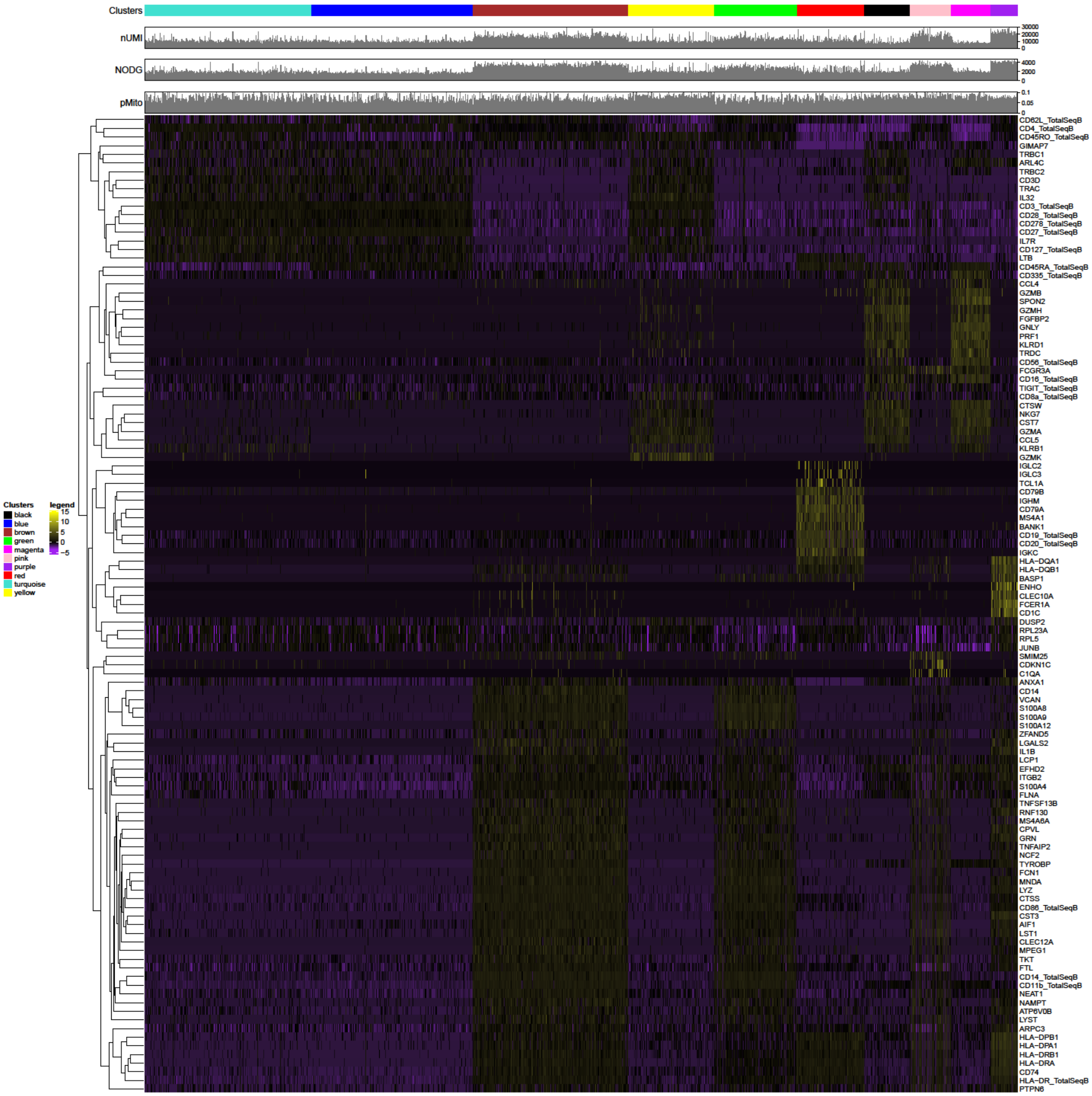


**Supplementary Figure 7:** Heatmap visualizing DE genes computed for the 10X PBMC example clustered using RCA2. DE genes are computed in either **a)** 1 vs all or **b)** pairwise manner.

a.)

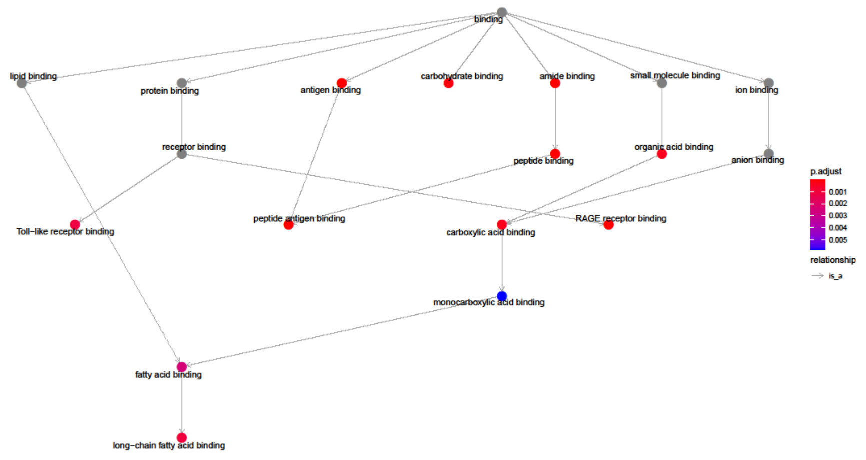


b.)

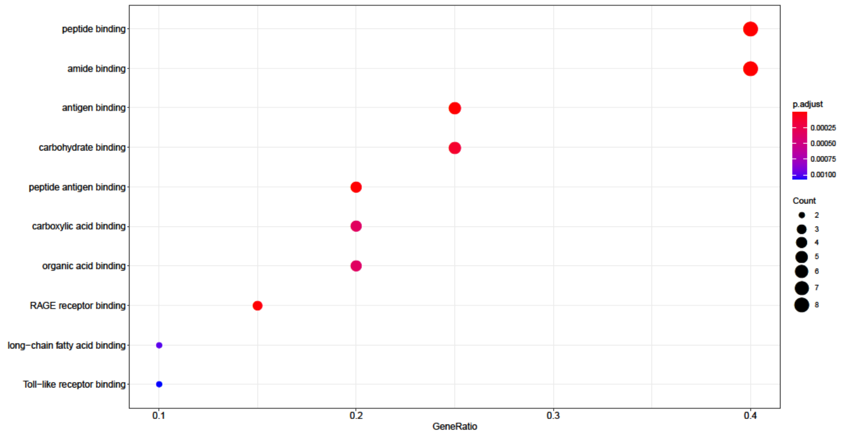


c.)

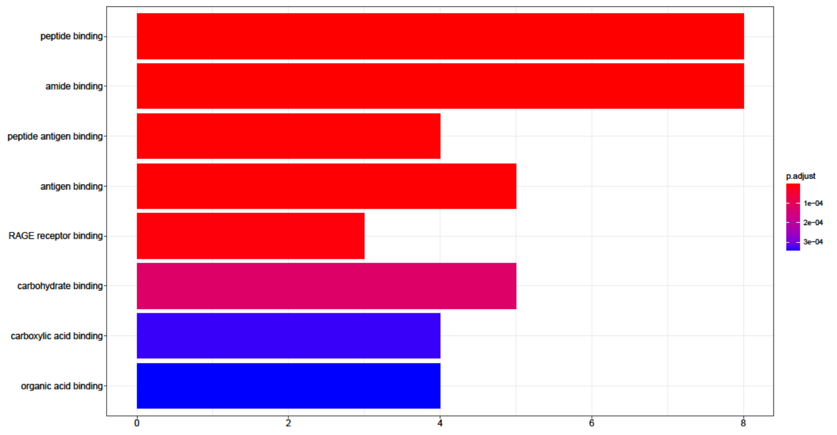


**Supplementary Figure 8:** GO enrichment for Molecular function terms represented as a **a)** Go Plot, **b)** Dot Plot, **c)** Bar plot, generated for one of the clusters using the 10X PBMC data set.

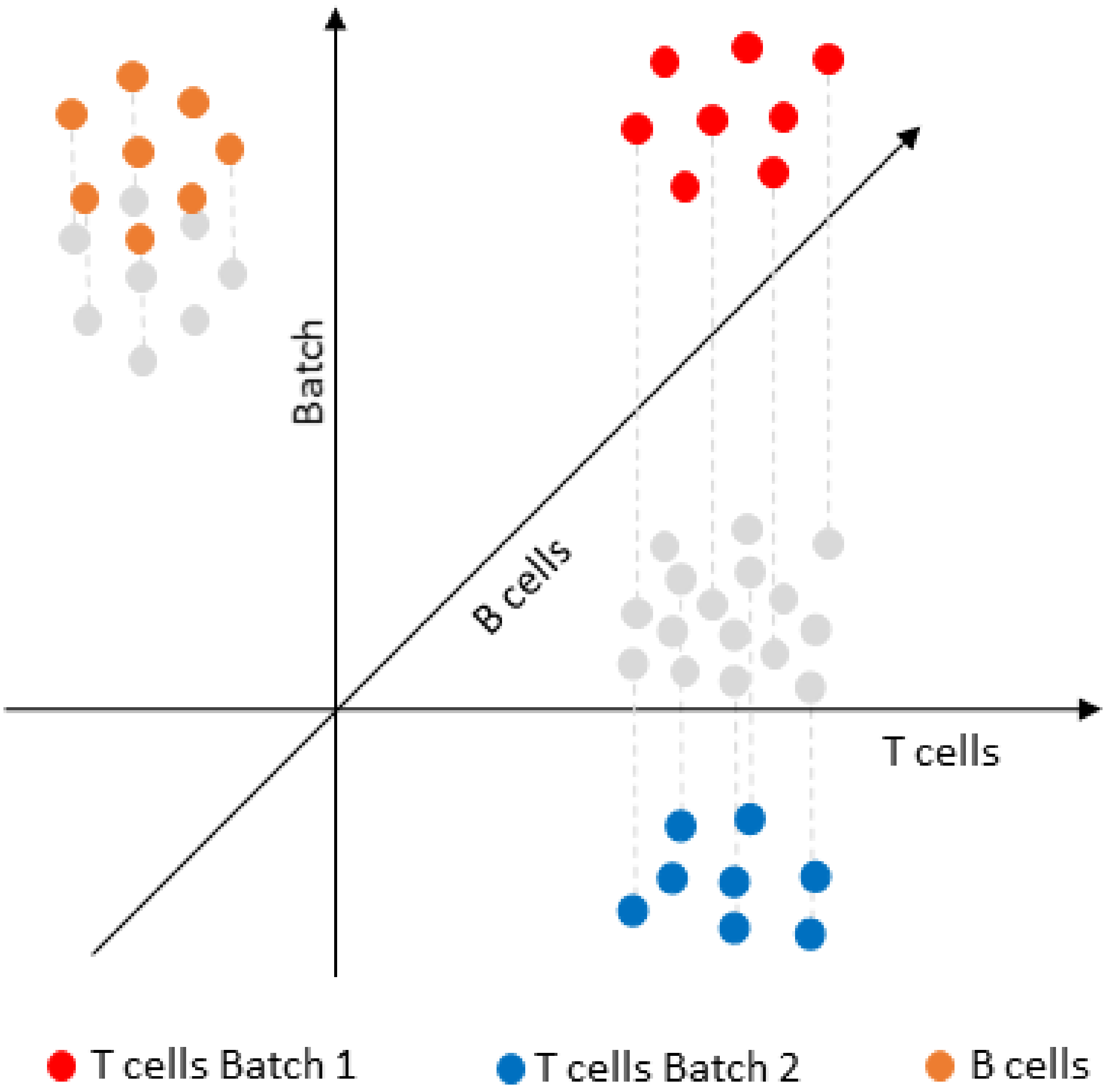


**Supplementary Figure 9:** Illustration of the concept of batch effects. Consider three data, two sets representing T cells (red and blue) and one set representing B cells (orange). In a three dimensional space measuring transcriptomic similarity to T cells (x-axis), B cells (y-axis) and to a Batch variable (z-axis), we find the T cell clusters to be separated by the Batch affect, while in 2D space comprised of only T cell and B cell similarity, they would be very similar to each other and show high similarity to T cells but not to B cells. In contrast to that, the B cell cluster's position stays almost constant neglecting the Batch column, with a high similarity o B cells and a negative score with respect to T cells.

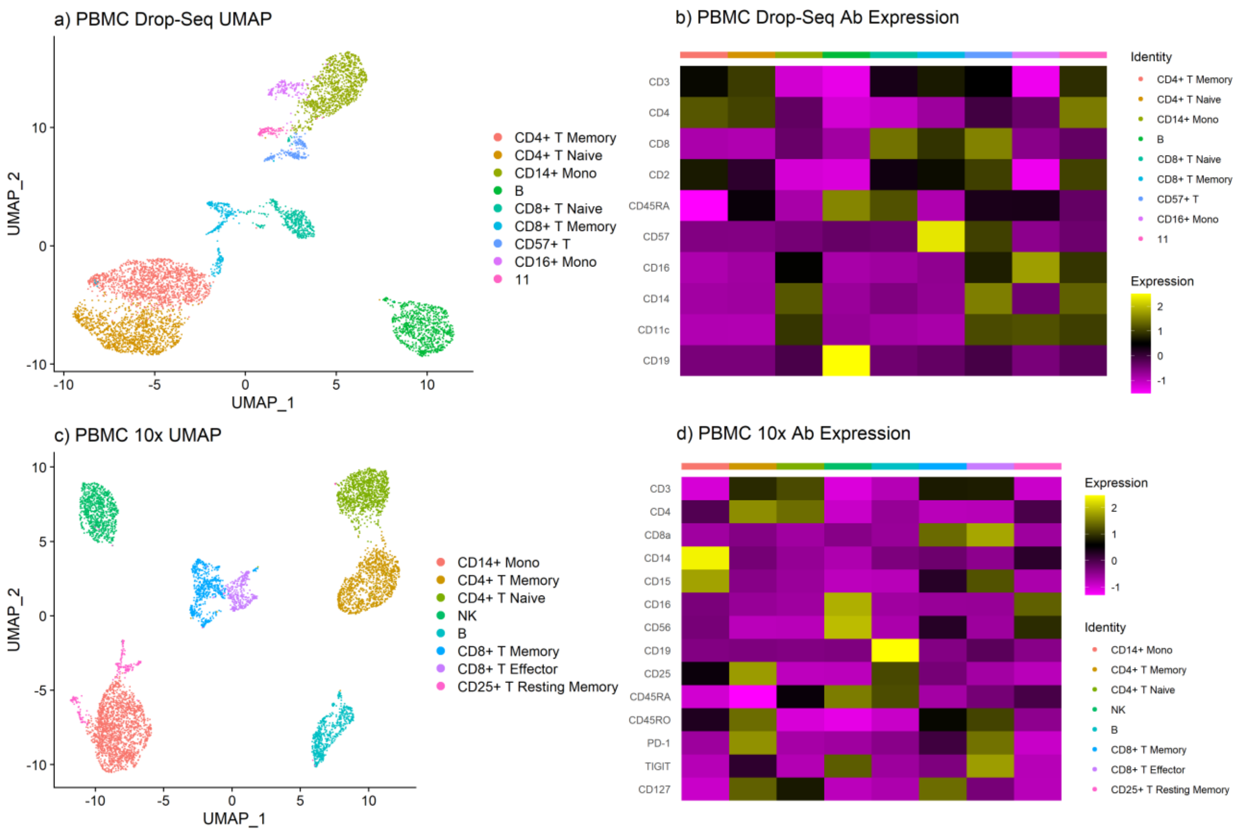


**Supplementary Figure 10:** Antibody-derived clustering of PBMCs. a) UMAP visualization of ADT-based clusters from the Drop-Seq dataset. b) Average antibody-expression of each cluster from the Drop-Seq dataset. c) UMAP visualization of ADT-based clusters from the 10x dataset. d) Average antibody-expression of each cluster from the 10x dataset.

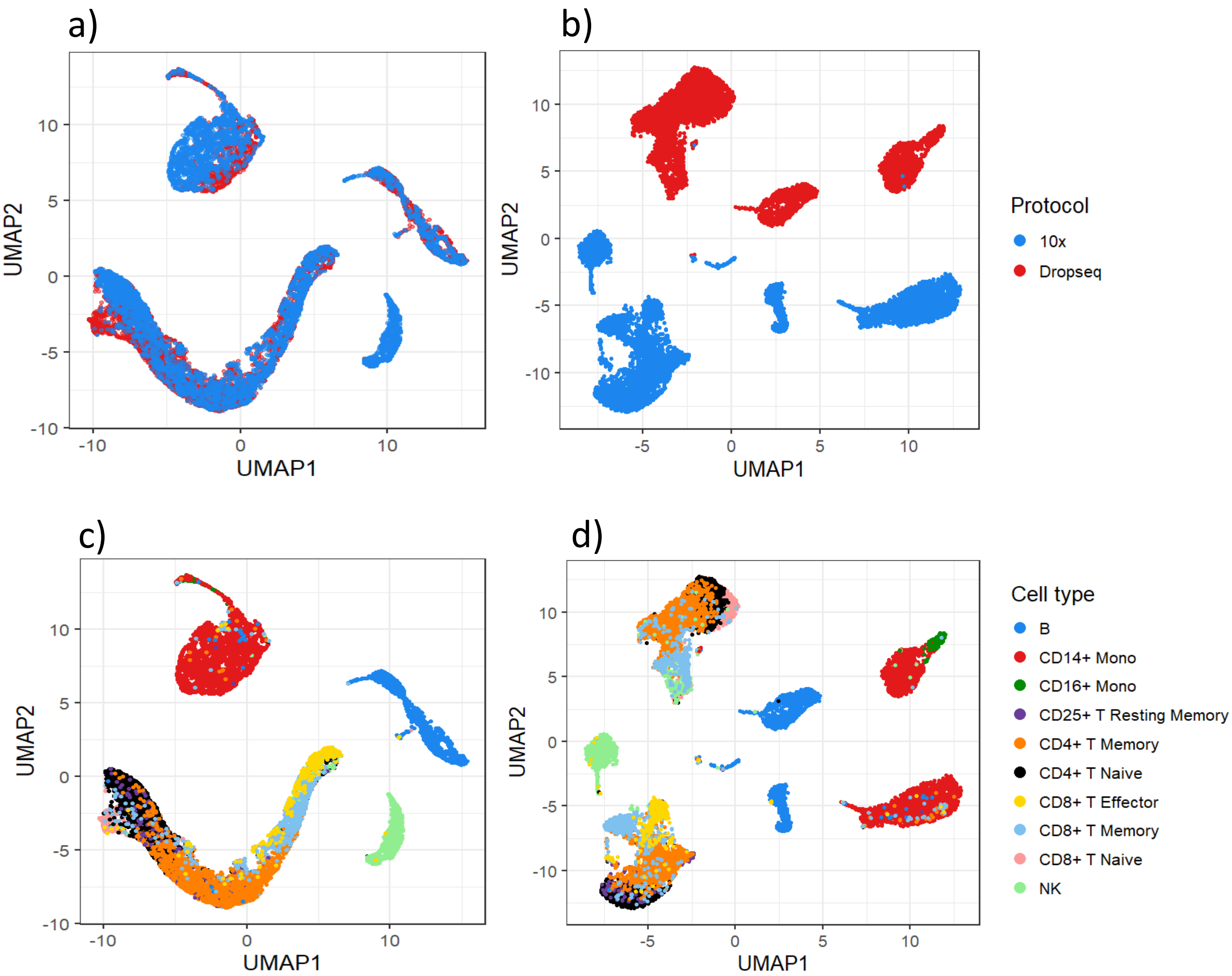


**Supplementary Figure 11: a)** UMAP of RCA reference projection of CITE-seq datasets colored by sequencing protocol. **b)** UMAP of a PCA embedding computed in Seurat using highly variable genes as features on CITE-seq datasets colored by sequencing protocol. **c)** UMAP of RCA reference projection of CITE-seq datasets colored by cell type. **d)** UMAP of a PCA embedding computed in Seurat using highly variable genes as features on CITE-seq datasets colored by cell type.

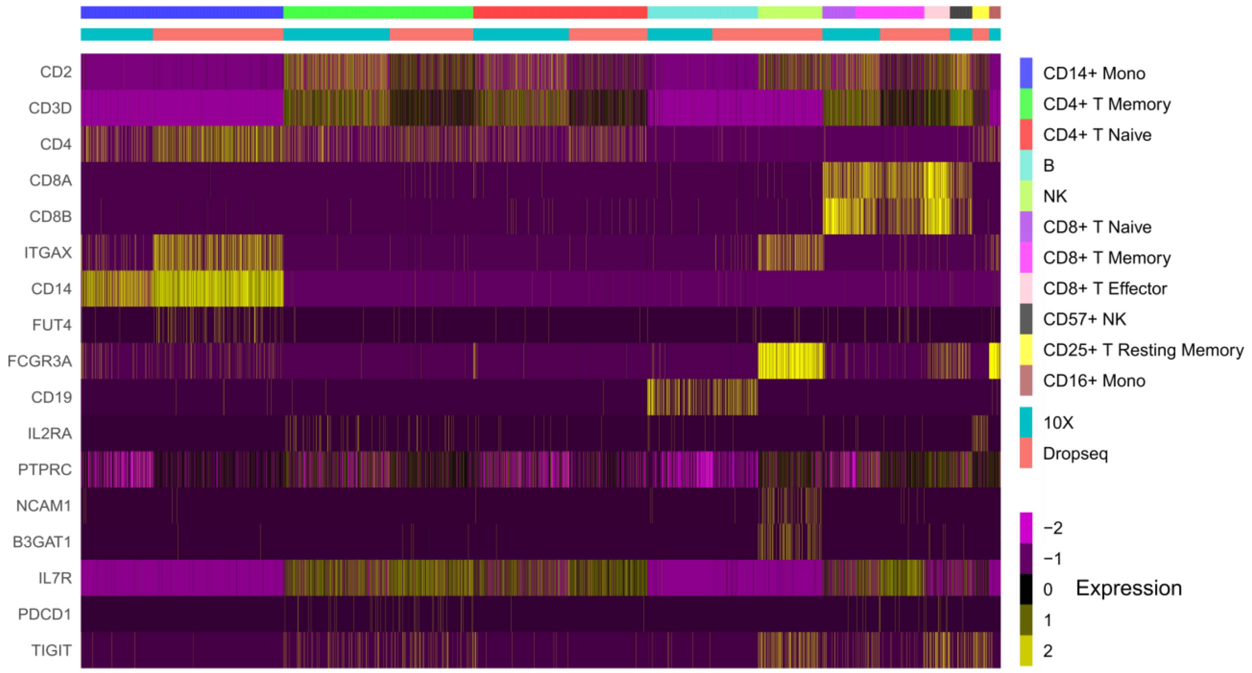


**Supplementary Figure 12:** Expression of genes captured with antibodies in the RNA-seq fraction of the Cite-seq PBMC data sets.

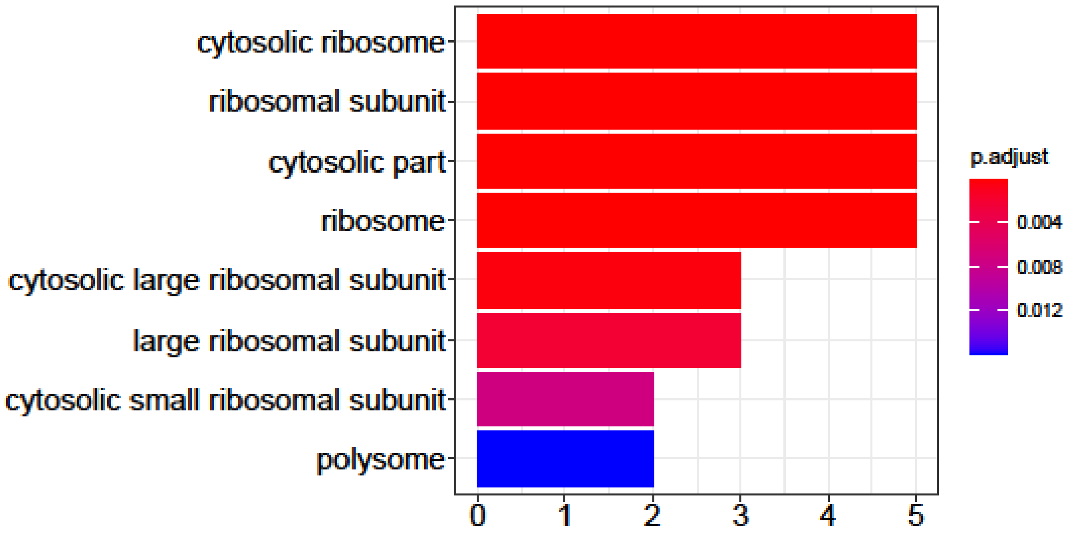


**Supplementary Figure 13:** GO enrichment (Cellular Compartment) for DE genes upregulated in the 10X CITE-seq PBMC data set.

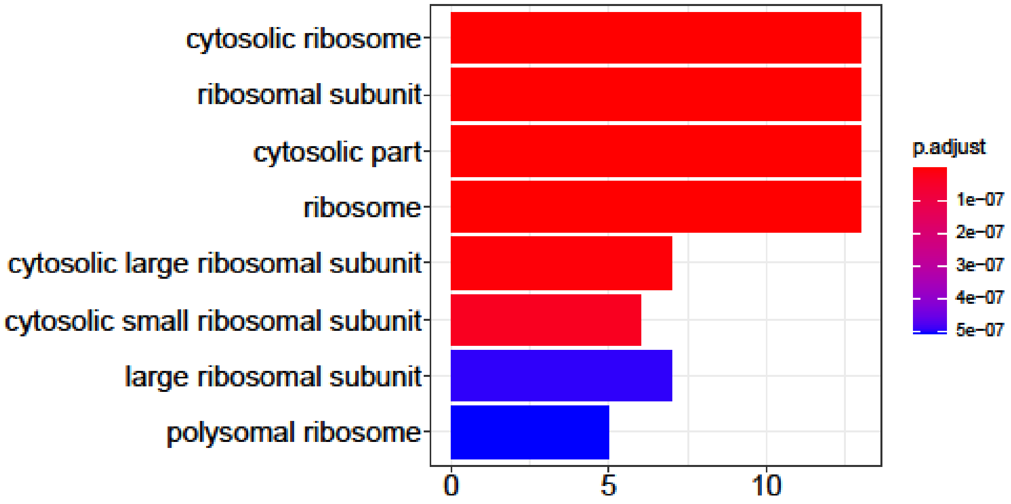


**Supplementary Figure 14:** GO enrichment (Cellular Compartment) for DE genes upregulated in the Dropseq PBMC data set.

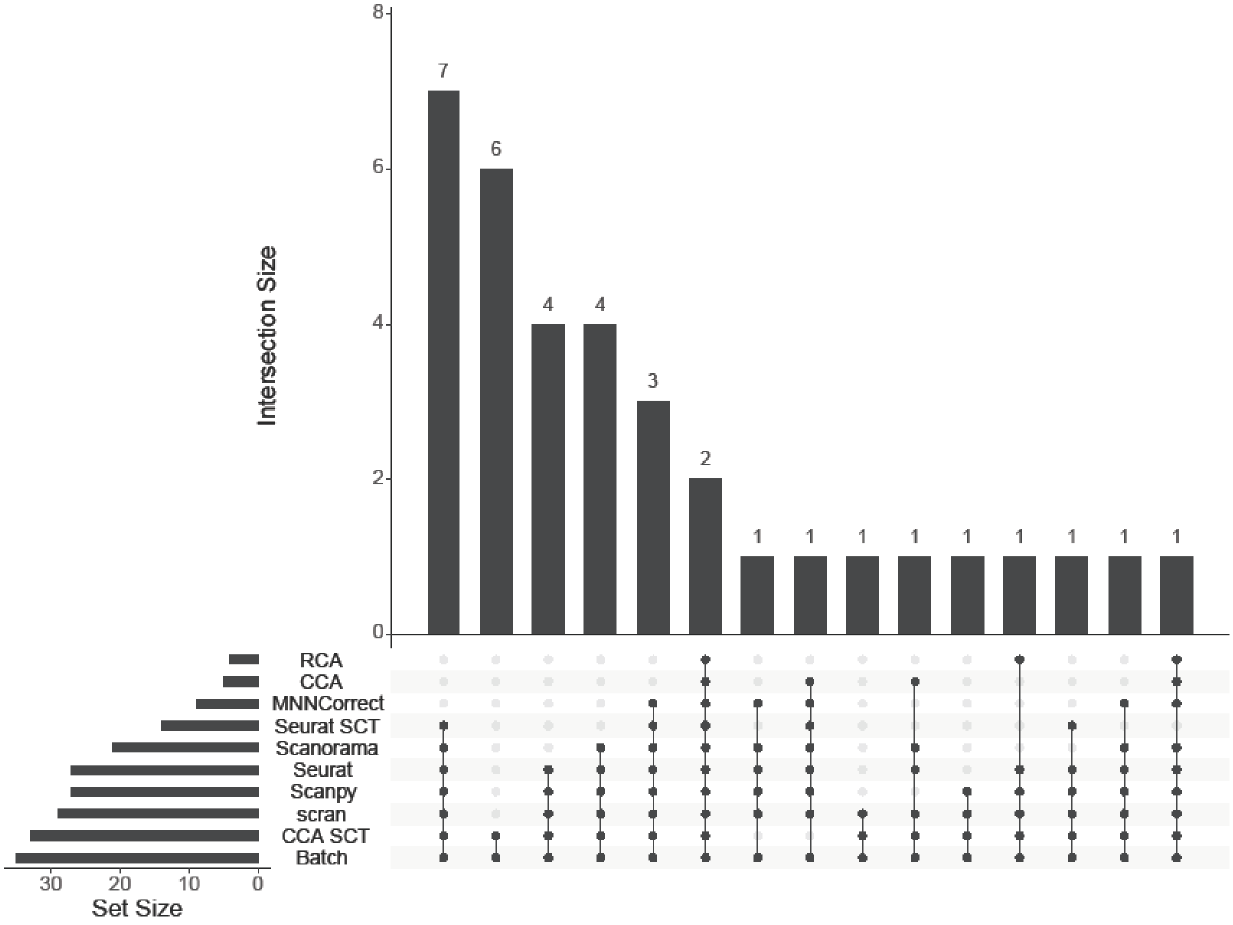


**Supplementary Figure 15:** UpSet plot showing the relationship between DE genes of scRNA-seq clusters and batch specific DE genes with respect to the used clustering method.

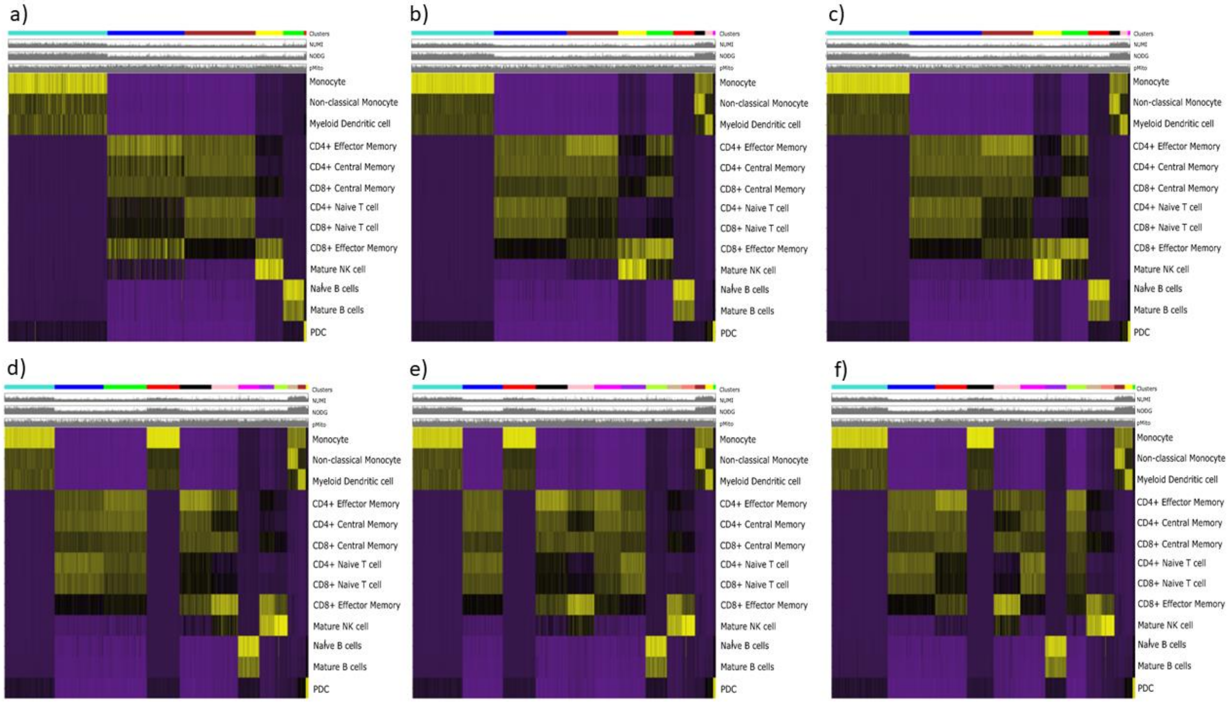


**Supplementary Figure 16:** Projection heatmaps with different resolutions for graph based clustering on a 10X PBMC data set.

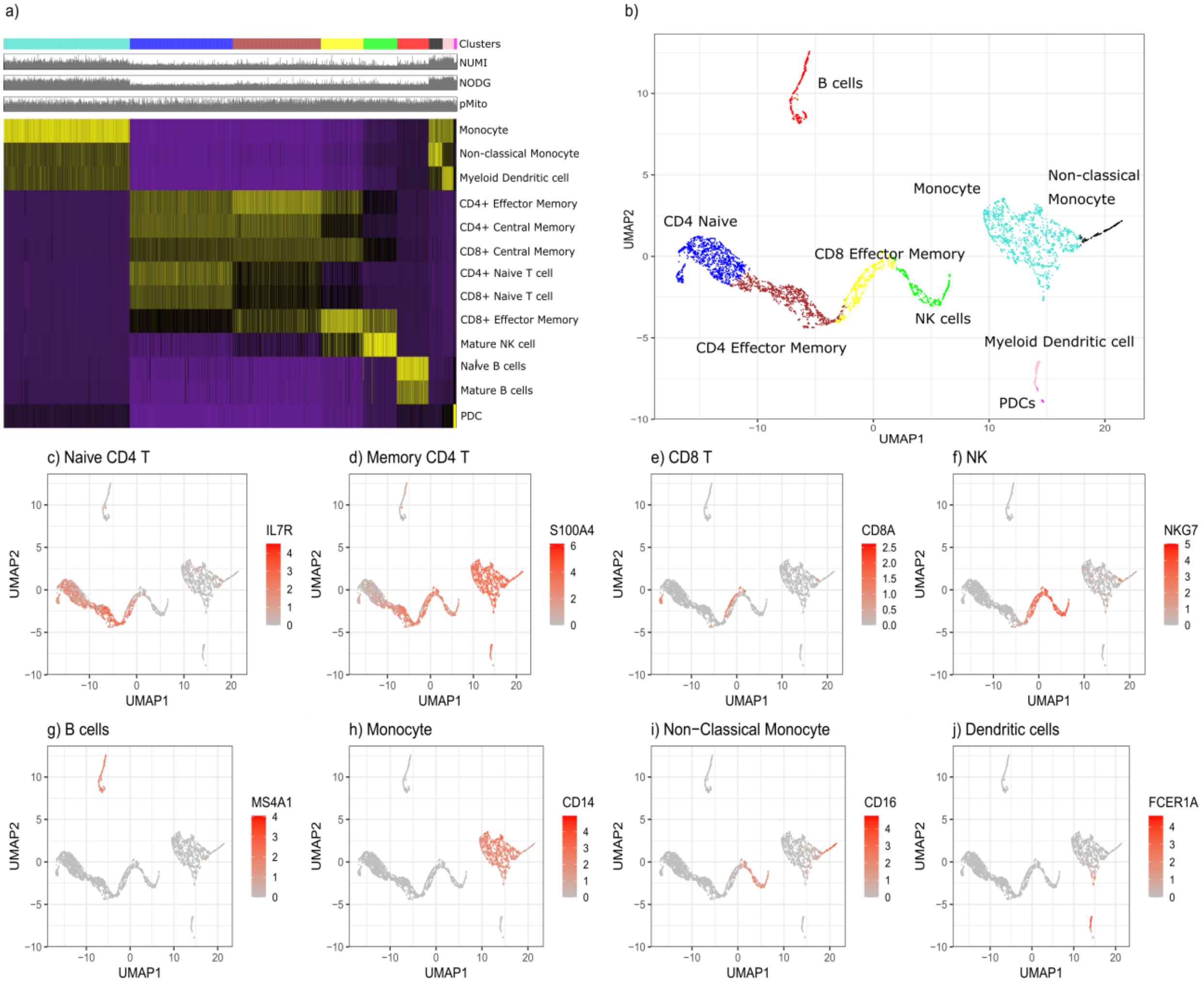


**Supplementary Figure 17:** a) Clusters obtained using graph based clustering on the reference projection of the 10X PBMC data set. b) Per cluster cell type prediction identifies CD4+ Naive T cells, CD4+ Memory cells, CD8+ Memory T cells, NK cells, B Cells, DC cells, as well as Classical and non-classical monocytes c)-j) Cannoical marker genes for all cell types shown in c-j.

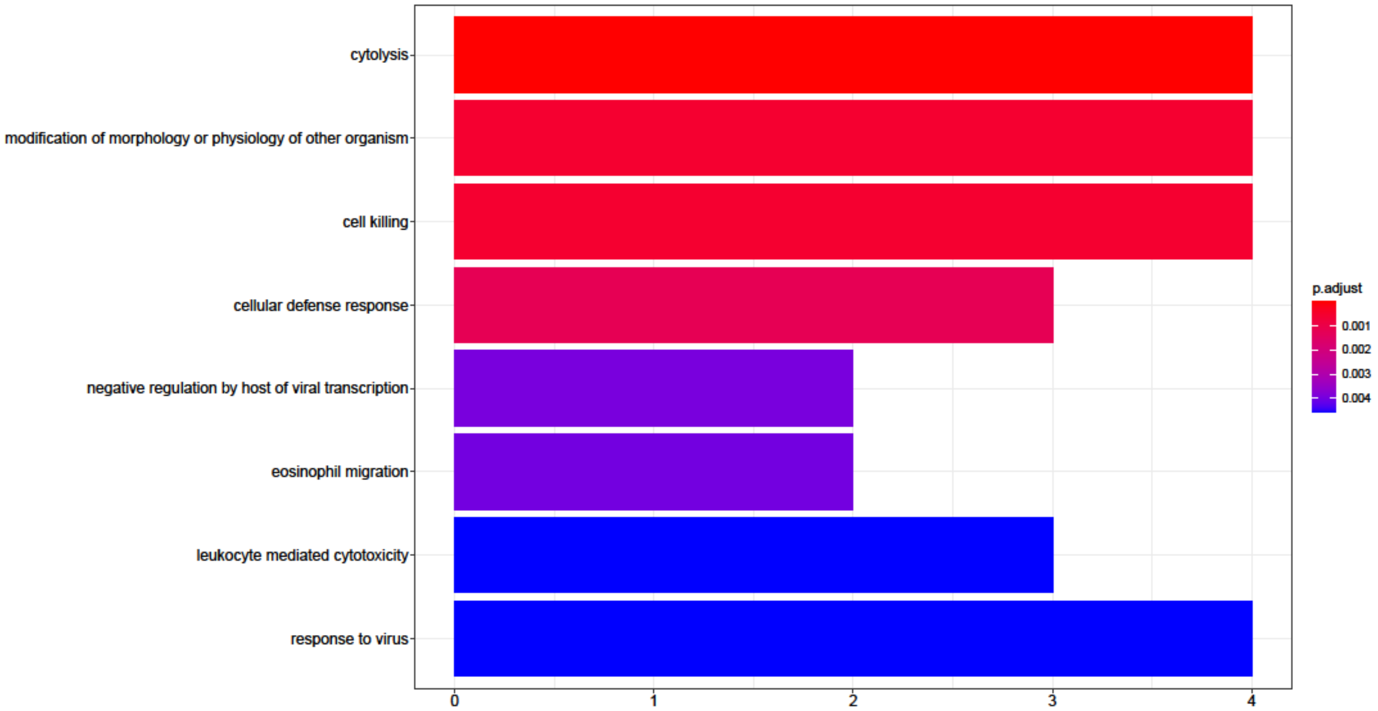


**Supplementary Figure 18:** Biological Process Terms computed for the NK cell cluster form the 10X PBMC data set.

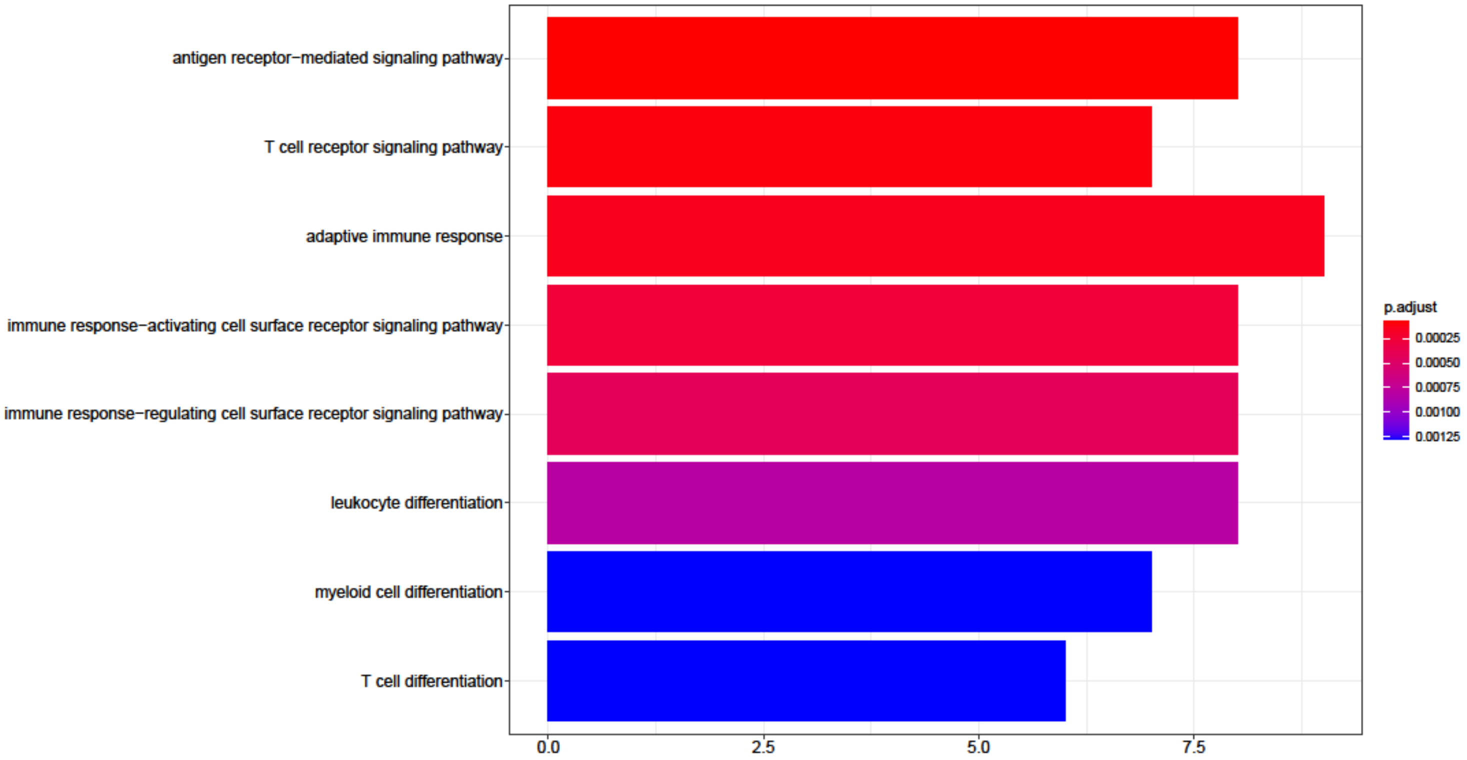


**Supplementary Figure 19:** Biological Process Terms computed for the T cell cluster form the 10X PBMC data set.

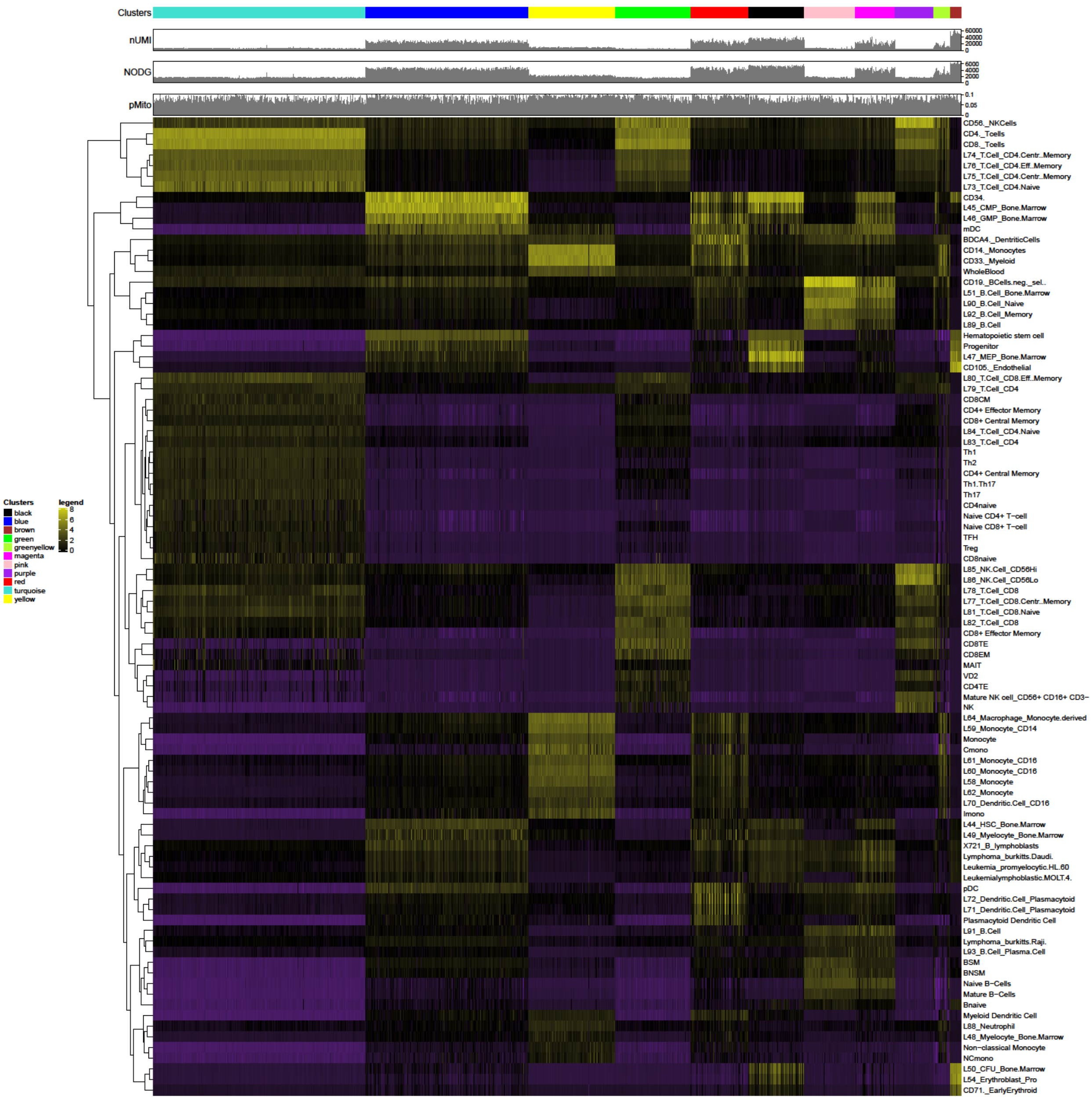


**Supplementary Figure 20:** Clustering of the RCA projection of the in-house bone marrow data set to obtain a classification into major cell types.

a.)

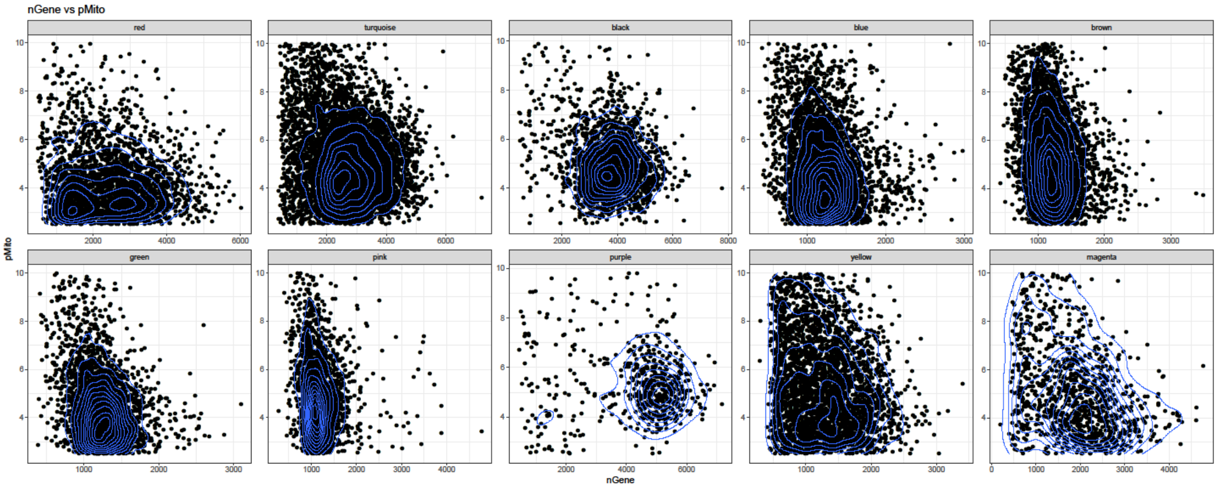


b.)

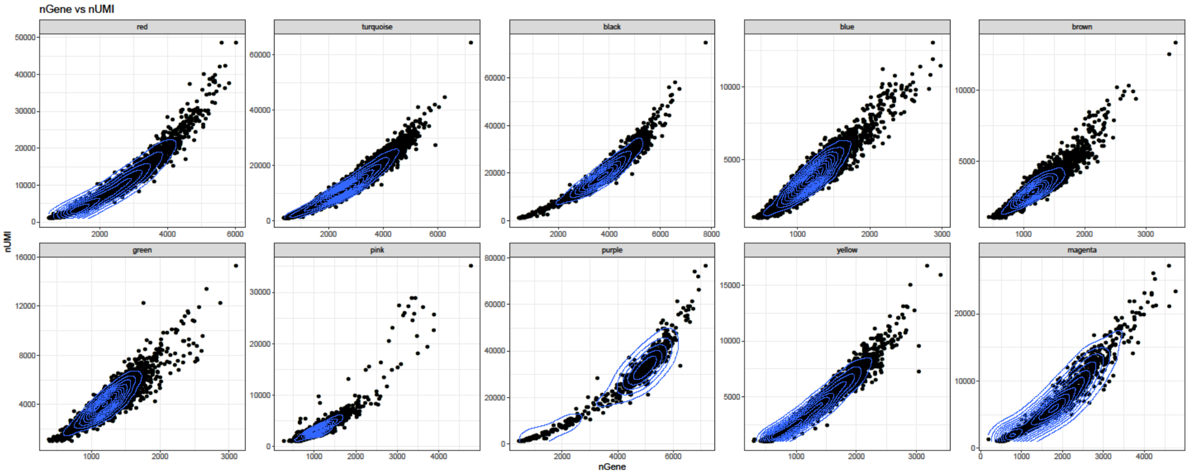


c.)

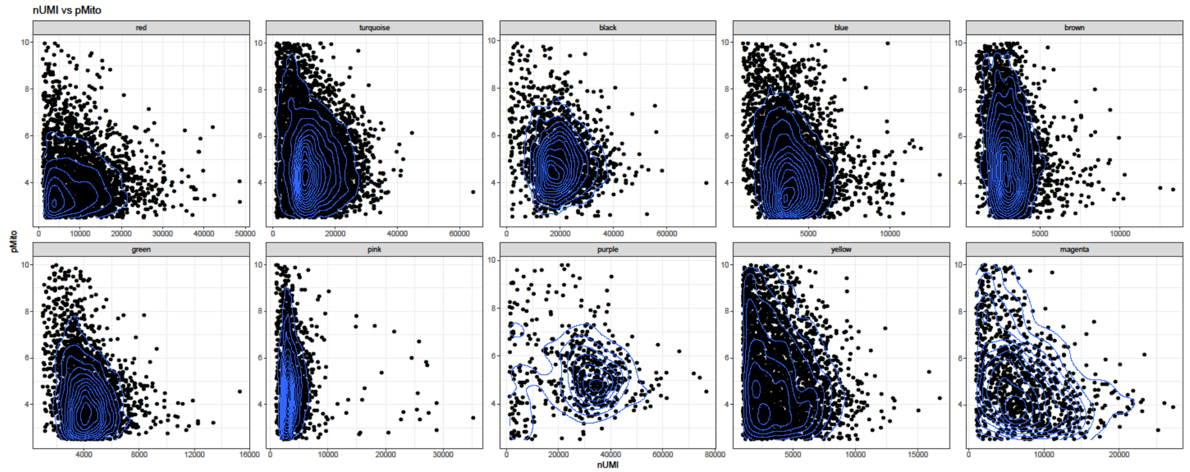


**Supplementary Figure 21:** Cluster specific QC plots for all major cell types identified in the bone marrow data set.

**a.)**

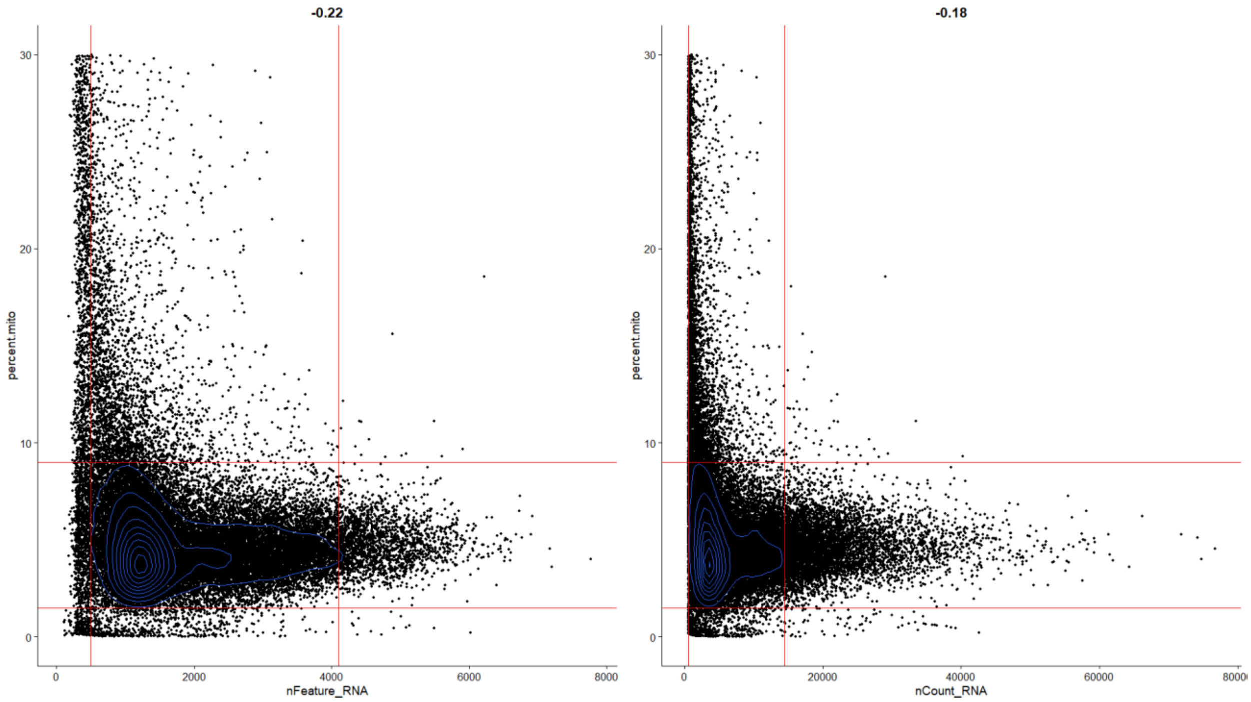


**b.)**

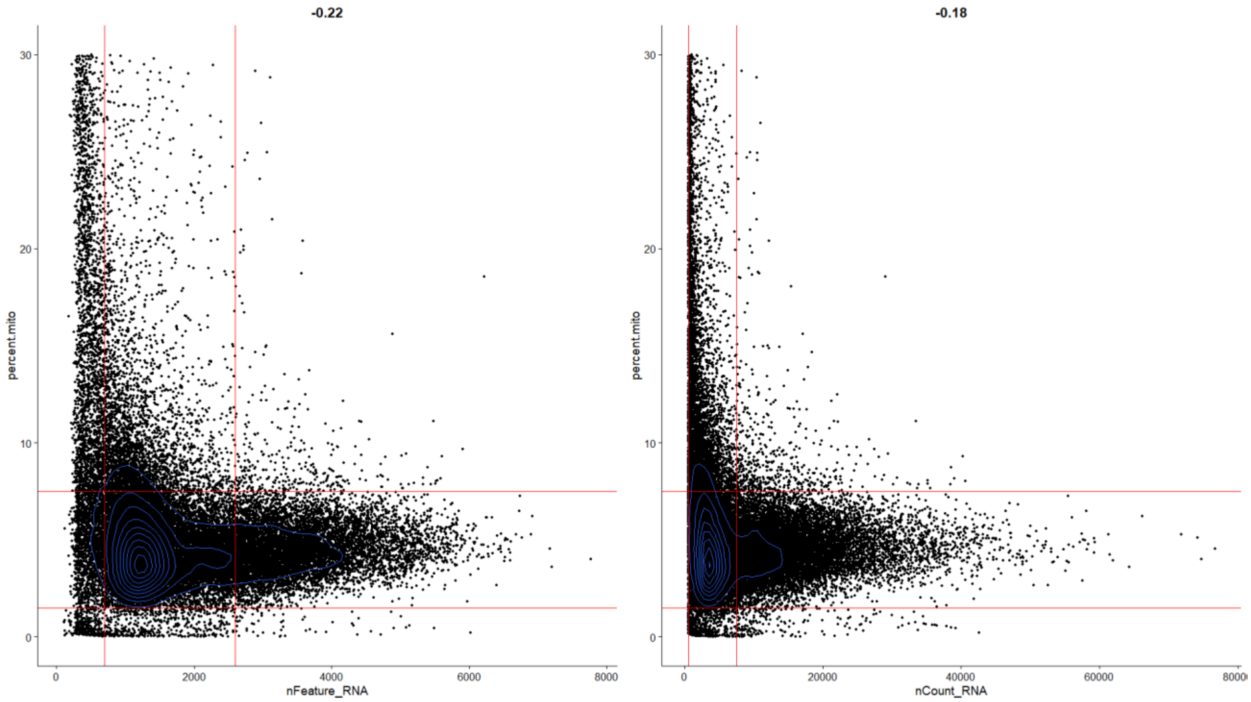


**Supplementary Figure 22:** Illustration of **a)** lenient and **b)** strict quality control cut-offs applied on the entire normal bone marrow data set.

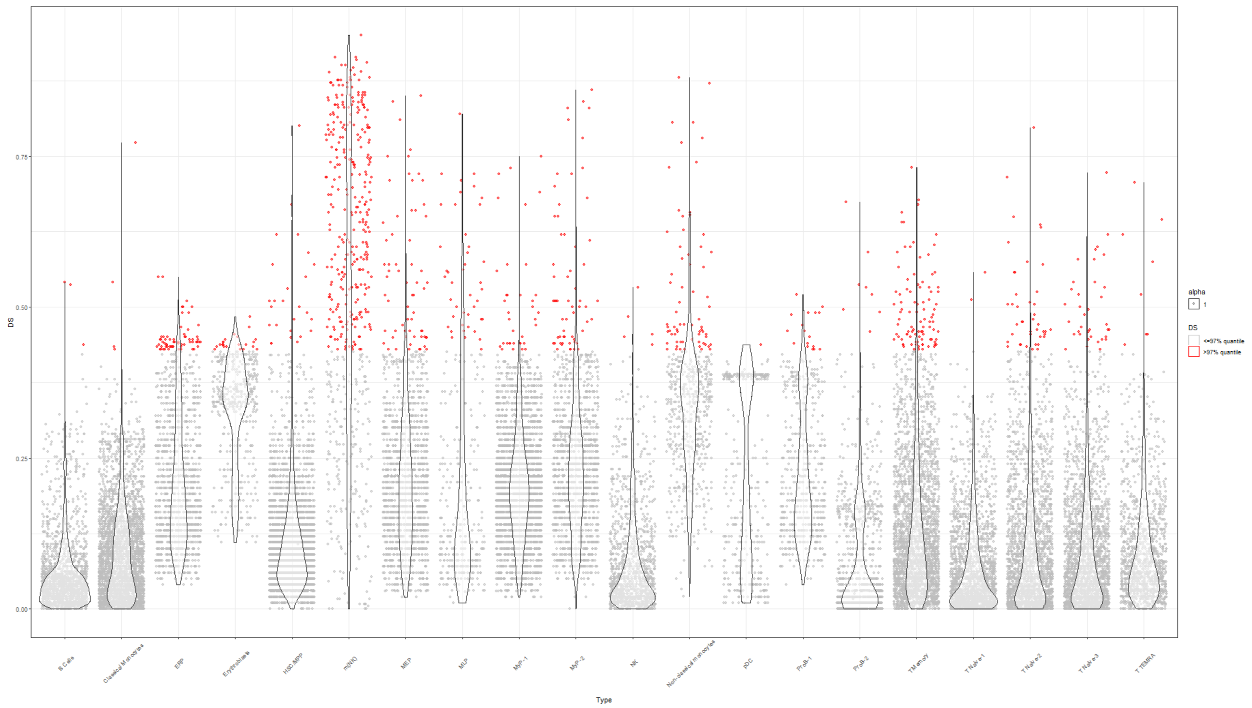


**Supplementary Figure 23:** Violin Plot depicting the PANN score of DoubletFinder individually for each cluster identified in the NBM data set. Cells with a PANN score >=0.97 quantile of the scores distribution are termed doublets and shown in red.

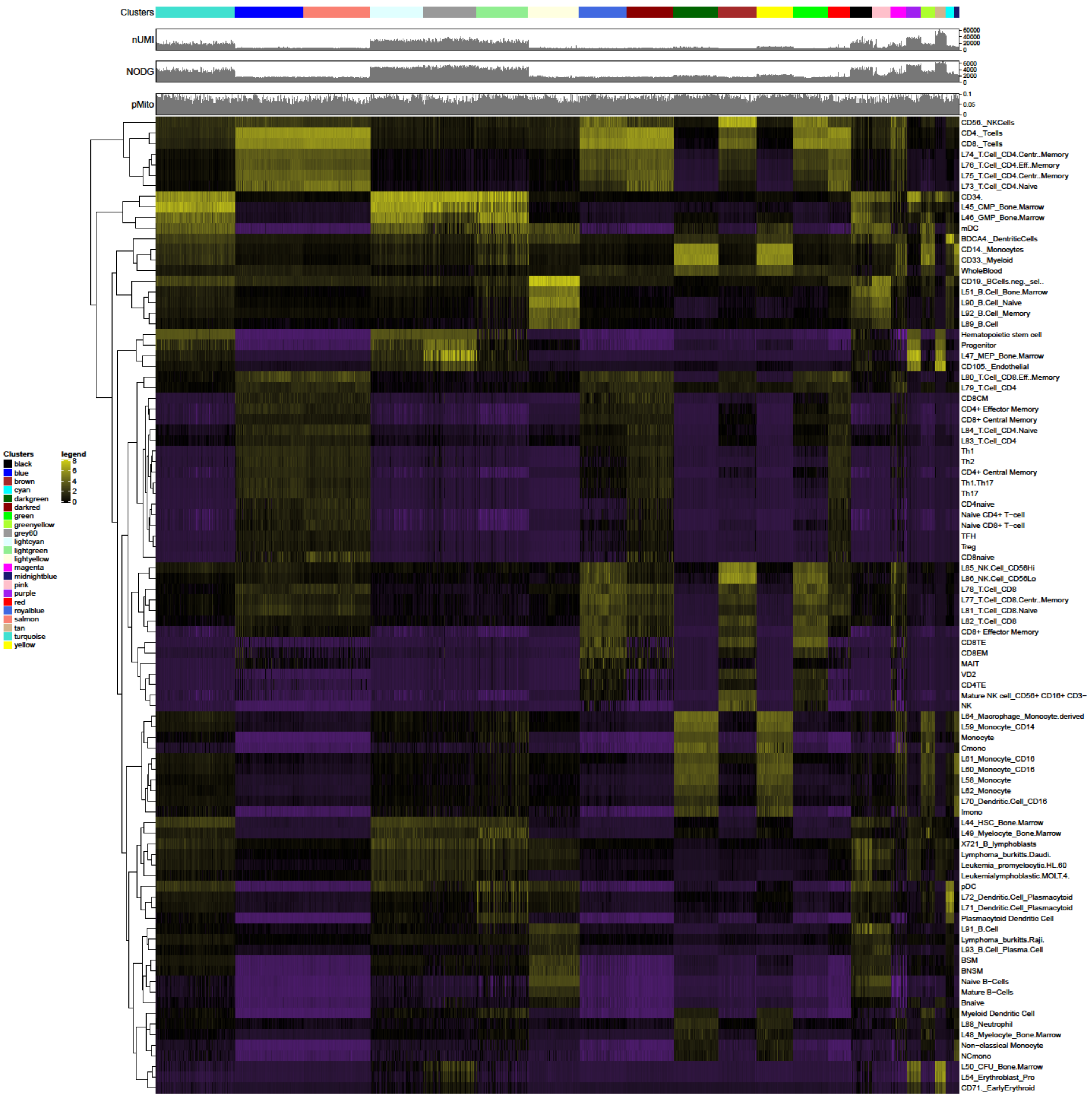


**Supplementary Figure 24:** Clustering of the RCA projection of the in-house bone marrow data set to obtain a classification into subtypes using a resolution of 0.5 for Seurat's graph based clustering.

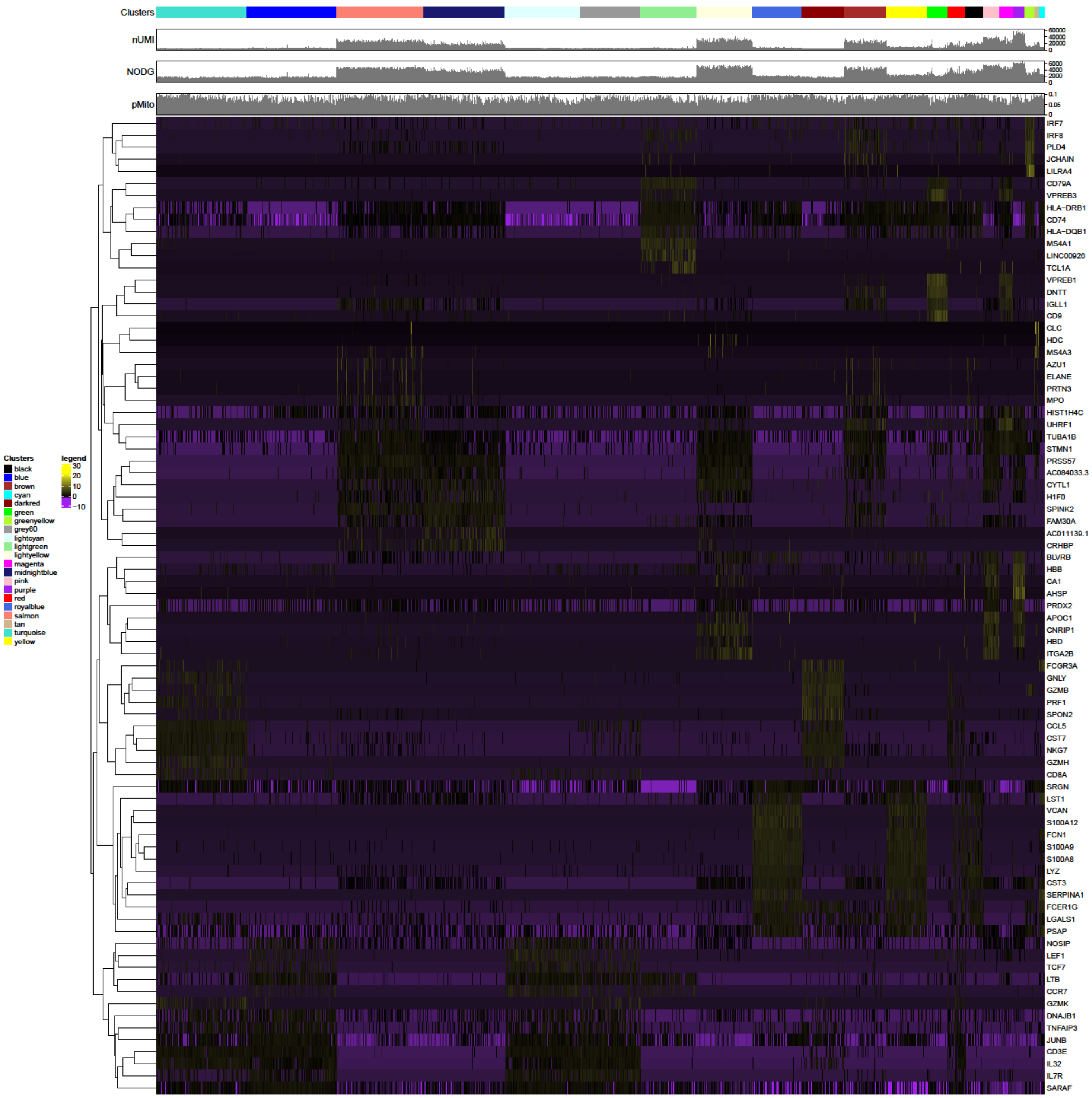


**Supplementary Figure 25:** Heatmap visualizing scaled counts for differentially expressed genes computed in a 1 vs all manner for the clusters shown in Sup. Fig. 20 on the in-house bone marrow data set.

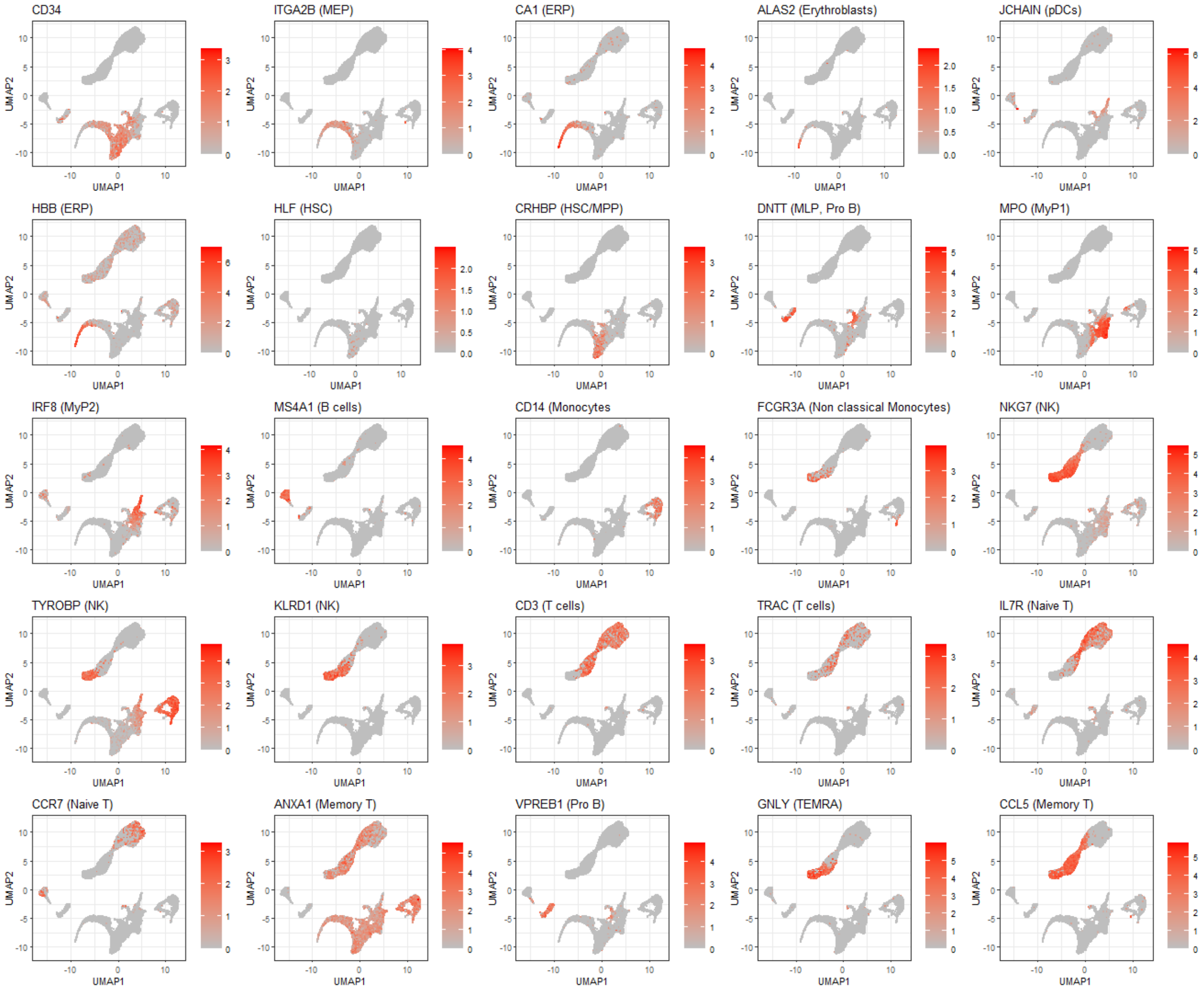


**Supplementary Figure 26:** Overview of the expression of 25 literature curated marker genes supporting the annotation of the normal bone marrow data set in shown in UMAP space.

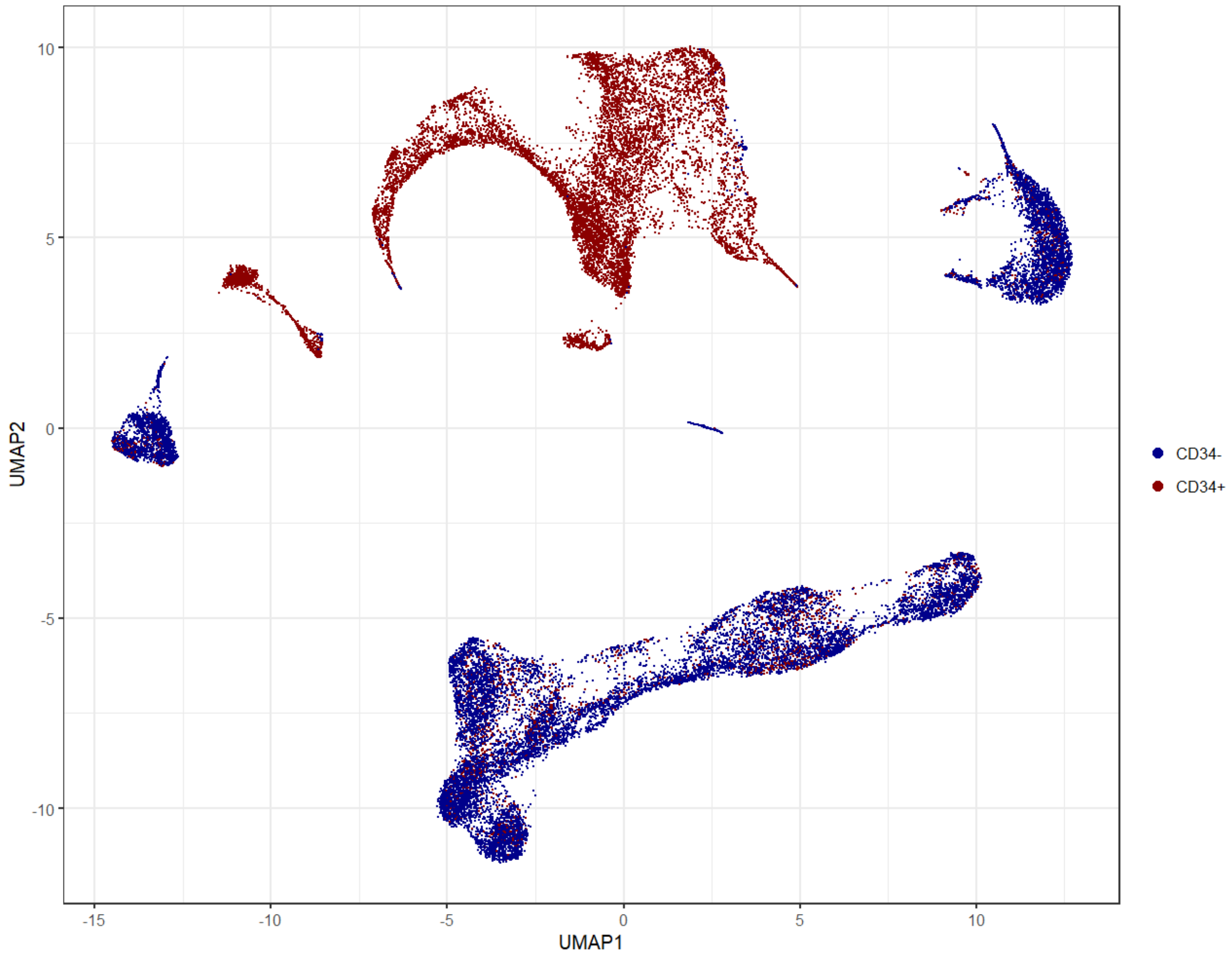


**Supplementary Figure 27:** UMAP visualization of a RCA2 reference projection of human bone marrow data colored by the FACS readout of each cell for the progenitor marker CD34.

**a.)**

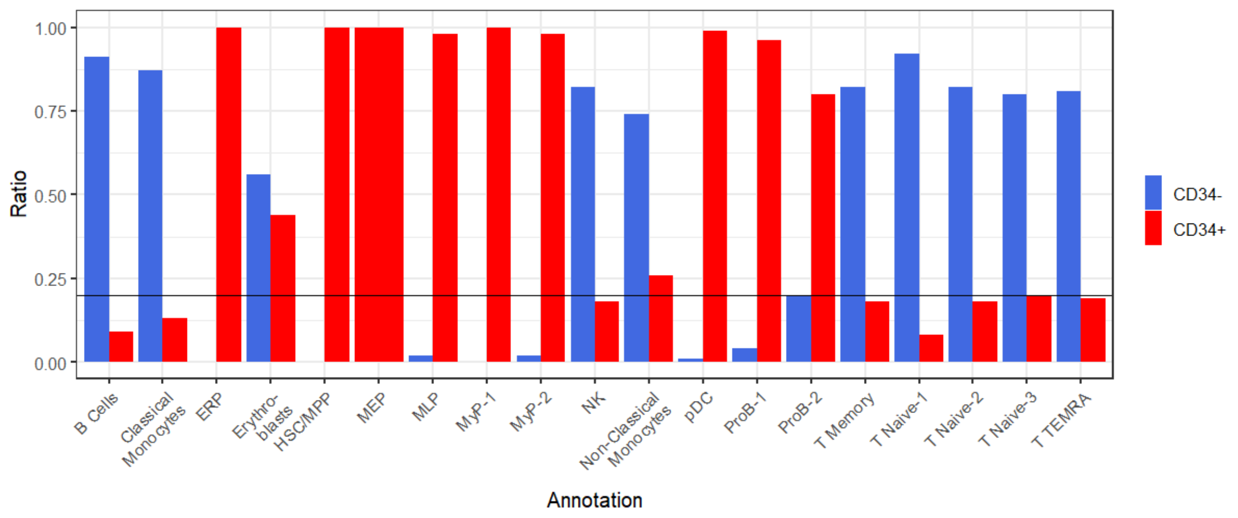


**b.)**

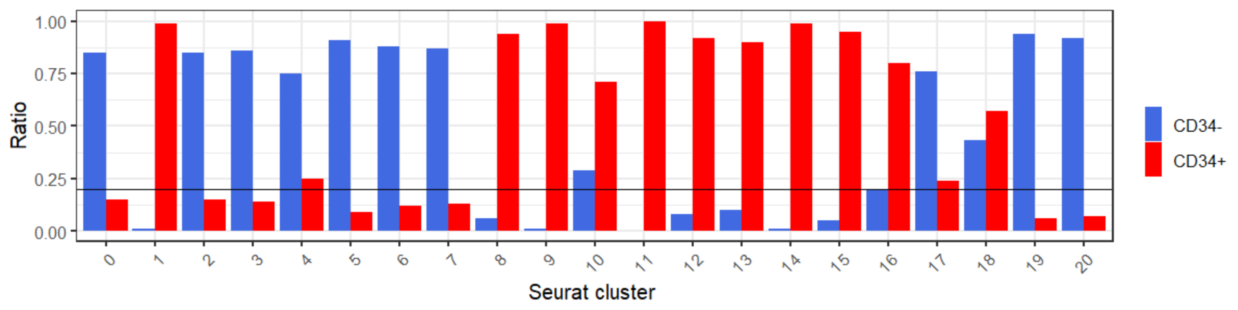


**Supplementary Figure 28:** Proportions of CD34 FACS labelled cells in clusters defined by scRNA-seq data of normal bone marrow samples using **a.)** RCA2 and **b.)** Seruat. In 10.05% of RCA clusters, the impurity is > 20%, compared to 19.05% of clusters in Seurat.

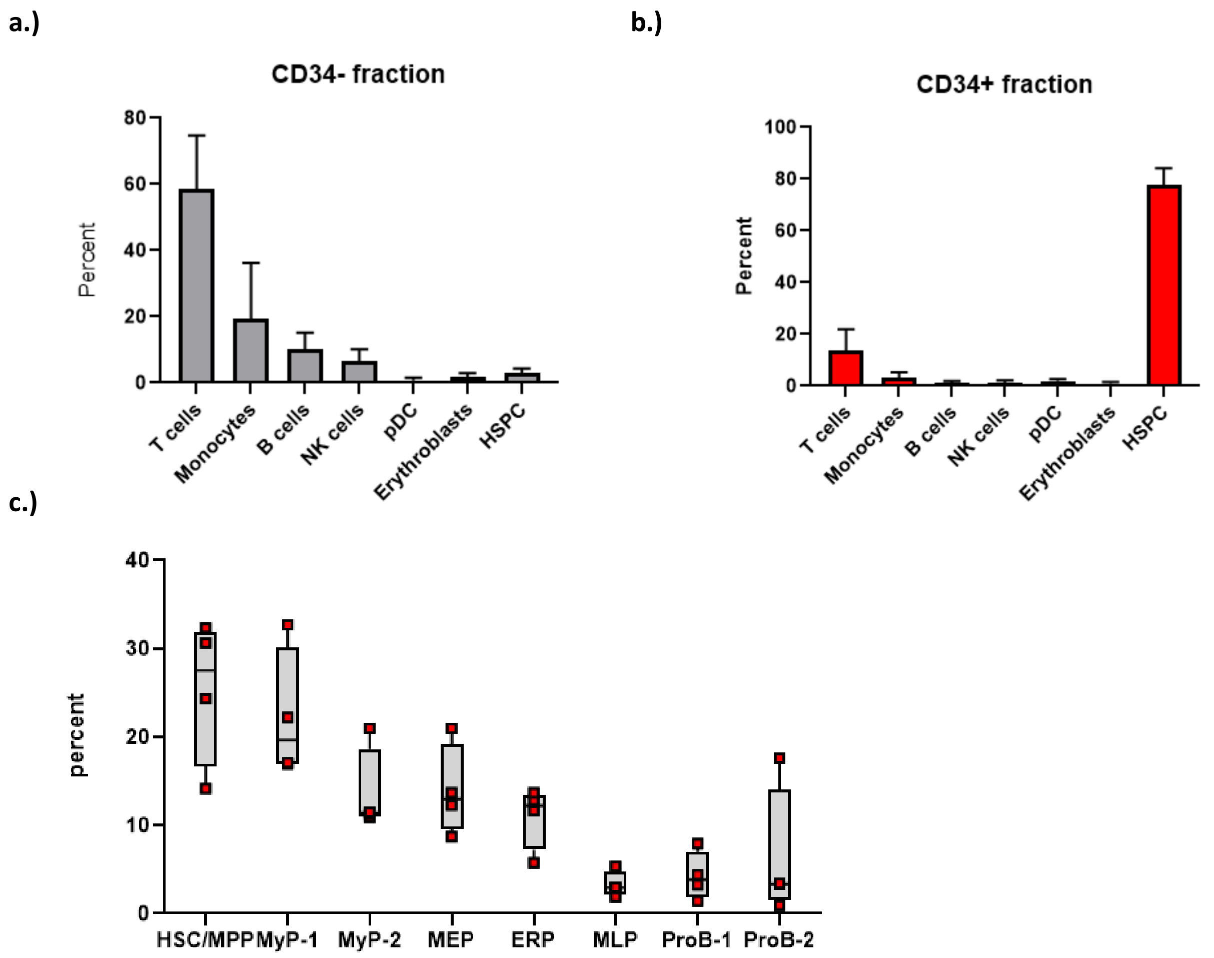


**Supplementary Figure 29:** Cluster frequency calculations for the **a.)** CD34- fraction of cells, **b.)** CD34+ fraction of cells and **c)** the CD34+ fraction cells depleted of differentiated clusters (according to transcriptome).

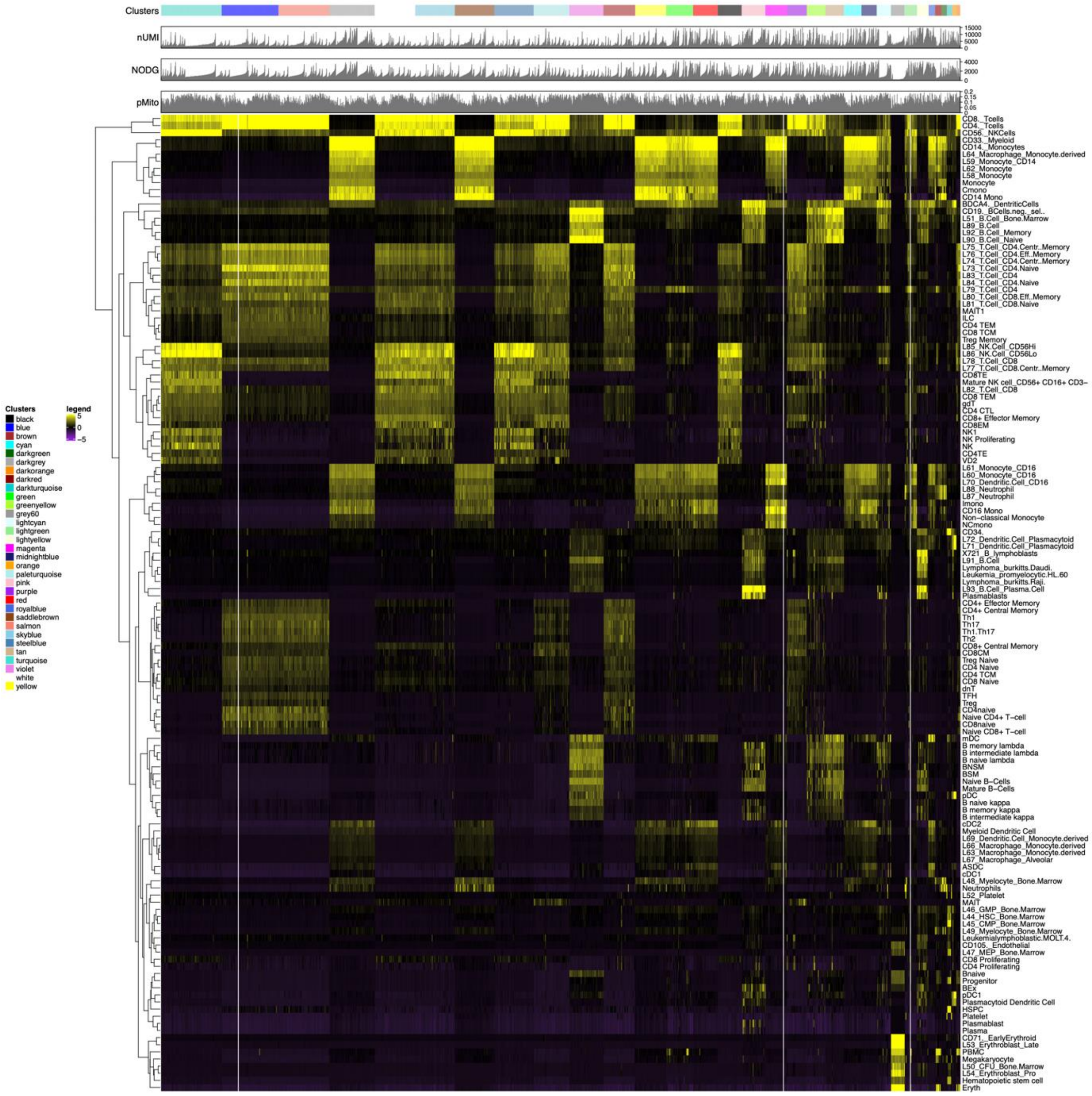

**Supplementary Figure 30**. Heatmap shows RCA reference projection profiles of 44,721 cells reported by Wilk et al. Each row represents a projection reference, while each column corresponds to a single cell.

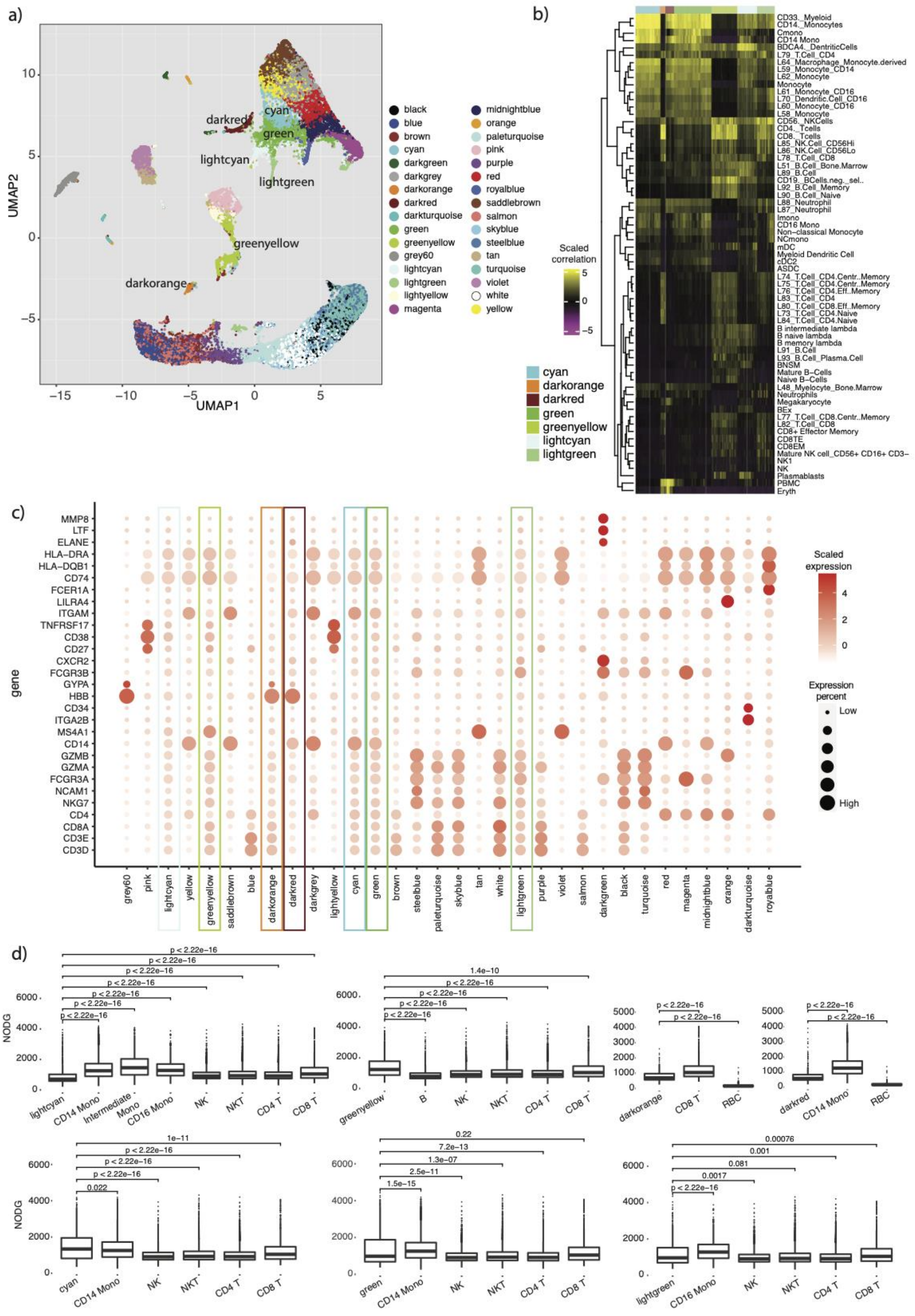

a)
b)
c)
d)
UMAP1
UMAP2
darkred
cyan
green
lightcyan
lightgreen
greenyellow
darkorange
Scaled correlation
Scaled expression
Expression percent
Low
High
gene
NODG

**Supplementary Figure 31**. a) UMAP plot shows the RCA clustering of 44,72 cells reported by Wilk et al. b) Heatmap shows the RCA reference projection profiles of 7 clusters identified as low quality or doublets. Each row represents a projection reference, while each column corresponds to a single cell. c) Bubble plot illustrates the marker gene expression levels across the 32 identified clusters. The bubble size indicates expression percentage for each cell type, while color intensity implies scaled expression level. d) Boxplots show NODG across different clusters or cell types. The value between two boxes is p-value calculated using the Wilcoxon Rank Sum test.

a)

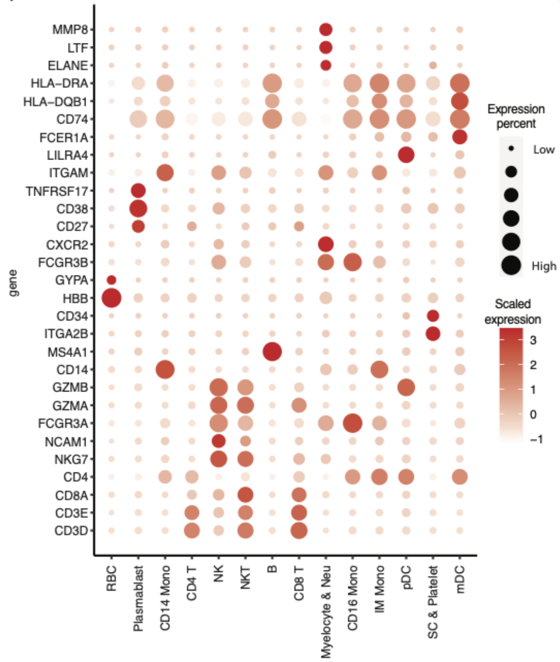


c)

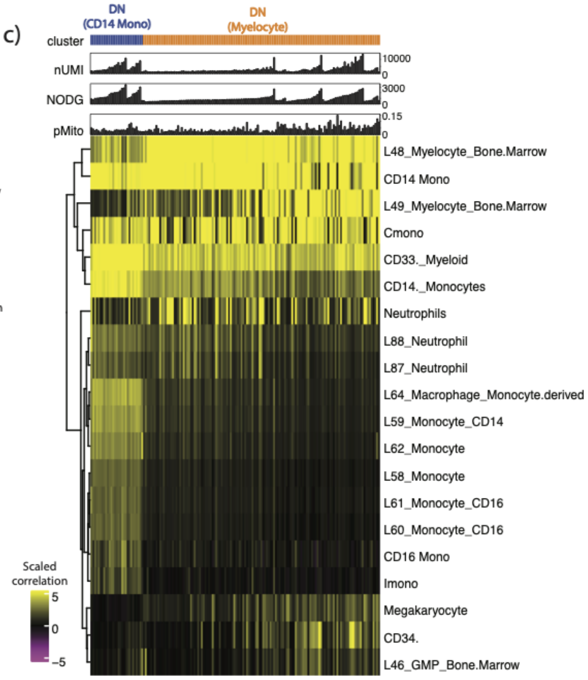


b)

| Cell type | number |
|---|---|
| Myelocyte & Neu | 131 |
| CD14 Mono | 29 |
| Low quality/Doublet | 32 |
| SC & Platelet | 4 |
| mDC | 3 |
| Intermediate Mono | 3 |
| NK | 2 |
| NKT | 1 |
| CD16 Mono | 1 |

d)

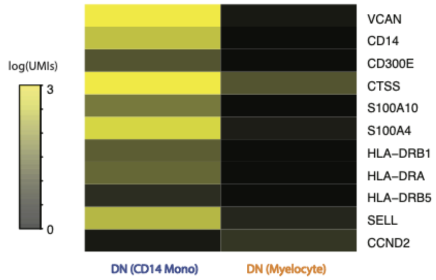


**Supplementary Figure 32**: a) Bubble plot shows the marker gene expression levels across all annotated cell types by RCA. The bubble size indicates expression percentage for each cell type, while color intensity represents scaled expression level. b) Table illustrates RCA annotations of "developing neutrophils" identified by Wilk et al. c) Heatmap shows the RCA reference projection profiles of 131 and 29 "developing neutrophils" annotated as myelocytes (orange) and CD14 monocytes (blue) by RCA respectively. Each row represents a projection reference, while each column corresponds to a single cell. d) Heatmap represents the DEGs identified between 131 and 29 "developing neutrophils" annotated as myelocytes (orange) and CD14 monocytes (blue) by RCA respectively.

a.)

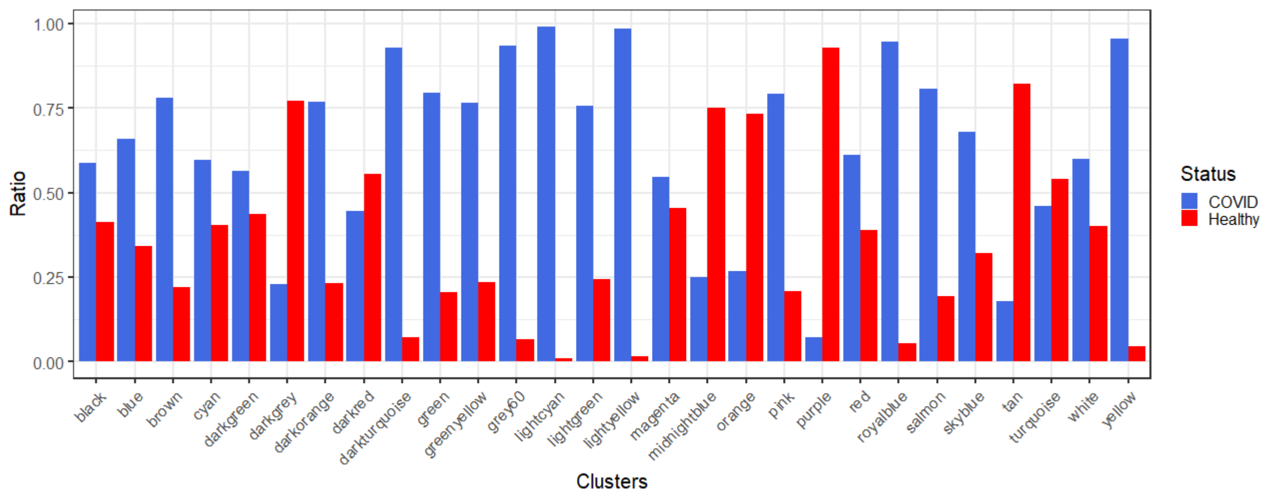


b.)

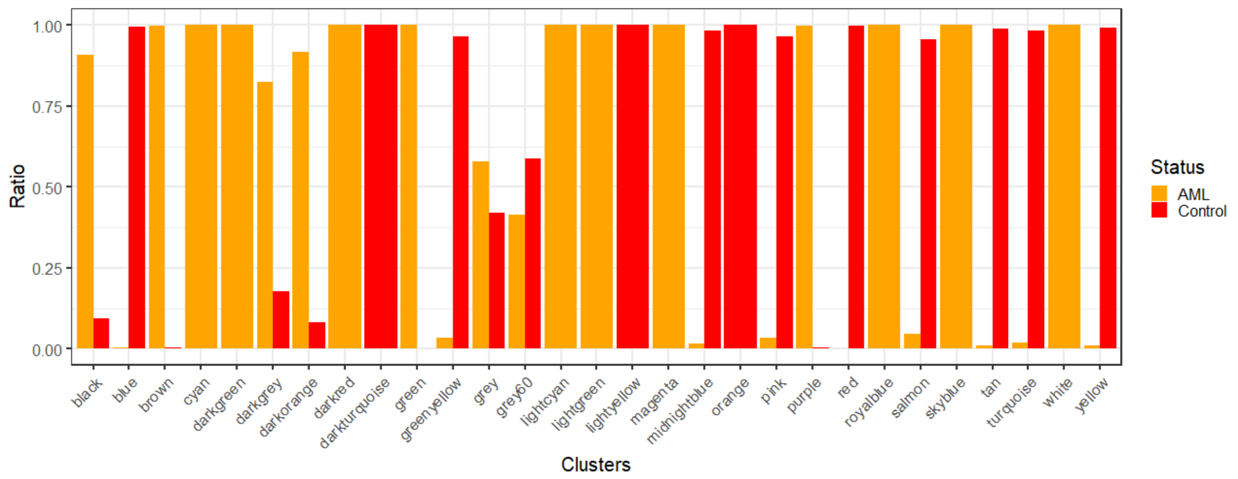


**Supplementary Figure 33:** Ratio of cells within RCA2 clusters determined for a.) Covid-19 disease specific clusters using data from Wilk et al. and for b.) AML Clusters using data from Petti et al.

| Dataset | NODG Lower | NODG Upper | NUMI Lower | NUMI Upper | pMito Lower | pMito Upper |
|---|---|---|---|---|---|---|
| PBMCs 10X | 300 | 4500 | 100 | ∞ | 0.025 | 0.1 |
| CITE-seq 10X (After Cellranger filtering) | 200 | ∞ | - | - | - | - |
| CITE-seq Drop-Seq (After Cellranger filtering) | 200 | ∞ | - | - | - | - |
| RA data set (as in paper) | 1000 | ∞ | - | - | 0 | 0.25 |

**Sup.Tab1:** Quality control parameters for a 10X PBMC data set as well as for two CITE-seq and several Rheumatoid arthritis scRNA-seq data sets. NODG is the number of detected genes, NUMI is the number of unique molecular identifiers, pMito is the percentage of mitochondrial genes.

| Dataset | NODG Lower | NODG Upper | NUMI Lower | NUMI Upper | pMito Lower | pMito Upper |
|---|---|---|---|---|---|---|
| General | NA | NA | 1000 | ∞ | 0.025 | 0.1 |
| red | 500 | 5200 | 1000 | ∞ | 0.025 | 0.1 |
| turquoise | 1000 | 5000 | 1000 | ∞ | 0.025 | 0.1 |
| black | 2000 | 5500 | 1000 | ∞ | 0.025 | 0.1 |
| blue | 750 | 1800 | 1000 | ∞ | 0.025 | 0.1 |
| brown | 750 | 1750 | 1000 | ∞ | 0.025 | 0.1 |
| green | 800 | 1750 | 1000 | ∞ | 0.025 | 0.1 |
| pink | 800 | 2000 | 1000 | ∞ | 0.025 | 0.1 |
| purple | 3500 | 6500 | 1000 | ∞ | 0.025 | 0.1 |
| yellow | 500 | 2500 | 1000 | ∞ | 0.025 | 0.1 |
| magenta | 400 | 3750 | 1000 | ∞ | 0.025 | 0.1 |

**Sup. Tab.2:** Quality control parameters for the bone marrow data set. NODG is the number of detected genes, NUMI is the number of unique molecular identifiers, pMito is the percentage reads overlapping mitochondrial genes.

| p_val | avg_logFC | pct.1 | pct.2 | p_val_adj | Gene |
|---|---|---|---|---|---|
| 8.83E-149 | 2.76087855 | 0.81791045 | 0.00163399 | 2.19E-147 | PTPRCAP |
| 1.51E-148 | 2.70292509 | 0.81492537 | 0 | 2.19E-147 | RPS4Y1 |
| 8.22E-148 | -3.7102365 | 0.04776119 | 0.99673203 | 6.55E-147 | RPS28 |
| 9.03E-148 | -4.7730572 | 0.05074627 | 0.99673203 | 6.55E-147 | EEF1A1 |
| 1.37E-147 | -3.9709835 | 0.0358209 | 0.99509804 | 7.92E-147 | RPS3A |
| 1.61E-146 | -3.9914604 | 0.09253731 | 0.99509804 | 7.79E-146 | RPL41 |
| 2.27E-146 | -3.912639 | 0.10149254 | 0.99509804 | 9.41E-146 | RPL39 |
| 2.60E-146 | -3.5784117 | 0.0358209 | 0.99019608 | 9.44E-146 | RPS26 |
| 7.93E-146 | -3.0633749 | 0.04179104 | 0.99019608 | 2.55E-145 | RPL21 |
| 2.46E-145 | -3.5434198 | 0.28656716 | 1 | 7.13E-145 | RPL10 |
| 5.74E-142 | -2.8568548 | 0.4238806 | 0.99509804 | 1.51E-141 | RPS27 |
| 6.75E-142 | -2.4048975 | 0.43283582 | 0.99509804 | 1.63E-141 | RPL18A |
| 7.89E-141 | -2.6310264 | 0.00597015 | 0.96895425 | 1.76E-140 | RPS10 |
| 2.39E-140 | -2.0721405 | 0.6119403 | 0.99509804 | 4.95E-140 | RPS27A |
| 4.03E-139 | -1.5683971 | 0.93731343 | 1 | 7.79E-139 | MT-ATP6 |
| 3.06E-135 | -2.3208831 | 0.01492537 | 0.95588235 | 5.55E-135 | H3F3A |
| 8.30E-134 | -2.3201426 | 0.09552239 | 0.9869281 | 1.42E-133 | JUND |
| 1.43E-133 | 1.8352117 | 0.99701493 | 0.85457516 | 2.30E-133 | RPL13A |
| 1.18E-126 | -2.1645287 | 0.01492537 | 0.91993464 | 1.80E-126 | RPL7 |
| 2.30E-126 | -1.8926456 | 0.25074627 | 0.98366013 | 3.33E-126 | RPL22 |
| 5.46E-125 | 1.82357426 | 0.9761194 | 0.75980392 | 7.54E-125 | RPL31 |
| 7.37E-124 | 2.26682795 | 0.93731343 | 0.43954248 | 9.72E-124 | RPS20 |
| 1.65E-117 | 1.53695151 | 0.99104478 | 0.88235294 | 2.08E-117 | RPL27A |
| 5.19E-114 | -1.7256419 | 0.22985075 | 0.97222222 | 6.27E-114 | EEF1B2 |
| 1.24E-111 | -1.7591346 | 0.13134328 | 0.94934641 | 1.43E-111 | NPM1 |
| 9.36E-109 | -1.7830268 | 0.08955224 | 0.91176471 | 1.04E-108 | PPIA |
| 2.58E-104 | -1.6304162 | 0.08955224 | 0.91339869 | 2.77E-104 | TMA7 |
| 1.17E-92 | -1.5553256 | 0.02985075 | 0.79411765 | 1.21E-92 | SUMO2 |
| 1.71E-73 | 1.76891989 | 0.61492537 | 0.11111111 | 1.71E-73 | ALDOA |

**Sup. Tab. 3:** DE genes identified for different CITE-seq batches within ADT derived clusters using a minimum log fold change of 1.5 and a q-value cut-off of 0.05.

| p_val | avg_logFC | pct.1 | pct.2 | p_val_adj | group1 | gene |
|---|---|---|---|---|---|---|
| 0 | 1.985549 | 1 | 0.997423 | 0 | brown | CD14 |
| 0 | 1.784014 | 1 | 0.218499 | 0 | brown | FCN1 |
| 0 | 1.77773 | 1 | 0.53866 | 0 | brown | CTSS |
| 0 | 1.739737 | 1 | 0.406071 | 0 | brown | LYZ |
| 0 | 1.659181 | 0.982827 | 0.180985 | 0 | brown | VCAN |
| 0 | 1.641425 | 0.97358 | 0.15063 | 0 | brown | CD14 |
| 0 | 1.585389 | 0.994716 | 0.216781 | 0 | brown | MNDA |
| 0 | 1.579942 | 1 | 0.422108 | 0 | brown | PSAP |
| 0 | 1.570105 | 0.836196 | 0.166667 | 0 | brown | IL1B |
| 0 | 1.567362 | 1 | 0.31701 | 0 | brown | S100A9 |
| 0 | 1.514141 | 0.993395 | 0.254009 | 0 | brown | FGL2 |
| 3.96E-300 | 1.635497 | 1 | 0.793814 | 8.25E-300 | brown | NEAT1 |
| 1.53E-297 | 1.876263 | 1 | 0.997709 | 2.94E-297 | brown | CD11b |
| 6.66E-294 | 1.744686 | 1 | 0.92984 | 1.19E-293 | brown | CD86 |
| 1.05E-144 | -1.78428 | 0.380449 | 0.756014 | 1.75E-144 | brown | LTB |
| 9.44E-136 | -1.74853 | 0.985469 | 0.990836 | 1.48E-135 | brown | CD278 |
| 3.93E-124 | -2.42807 | 0.998679 | 0.998282 | 5.78E-124 | brown | CD3 |
| 5.08E-111 | -2.50527 | 0.982827 | 0.9874 | 7.05E-111 | brown | CD27 |
| 1.12E-103 | -1.52405 | 1 | 0.997423 | 1.48E-103 | brown | CD28 |
| 4.72E-79 | -1.60163 | 0.974901 | 0.978522 | 5.90E-79 | brown | CD127 |
| 6.27E-25 | -1.57057 | 0.985469 | 0.930412 | 7.47E-25 | brown | CD19 |
| 9.21E-18 | -1.51976 | 0.159841 | 0.284078 | 1.05E-17 | brown | NKG7 |
| 1.25E-07 | -1.57728 | 0.993395 | 0.987973 | 1.35E-07 | brown | CD8a |
| 0.002099 | -1.65726 | 0.996037 | 0.988832 | 0.002187 | brown | CD20 |
| 5.92E-148 | -2.6834 | 0.1133 | 0.61158 | 1.84E-146 | turquoise | HLA-DRB1 |
| 1.45E-144 | -2.32317 | 0.034483 | 0.543206 | 2.24E-143 | turquoise | TYROBP |
| 2.83E-140 | -4.3644 | 0.997537 | 0.998254 | 2.93E-139 | turquoise | CD11b |
| 2.41E-136 | -3.03101 | 0.118227 | 0.583649 | 1.87E-135 | turquoise | HLA-DRA |
| 8.97E-135 | -2.34219 | 0.19335 | 0.641257 | 5.56E-134 | turquoise | HLA-DPA1 |
| 4.69E-130 | -2.18084 | 0.25 | 0.675007 | 2.42E-129 | turquoise | HLA-DPB1 |
| 8.10E-127 | -2.70223 | 0.998768 | 0.999418 | 3.59E-126 | turquoise | HLA-DR |
| 2.25E-120 | -1.68951 | 0.03202 | 0.482688 | 8.72E-120 | turquoise | FCER1G |
| 6.35E-109 | -2.3621 | 0.450739 | 0.719232 | 2.19E-108 | turquoise | CD74 |
| 9.89E-107 | -2.79853 | 0.04803 | 0.459703 | 3.07E-106 | turquoise | CST3 |
| 4.18E-106 | -2.00011 | 0.020936 | 0.43439 | 1.18E-105 | turquoise | MNDA |
| 2.68E-102 | -1.52866 | 0.061576 | 0.462322 | 6.93E-102 | turquoise | FGL2 |
| 2.14E-98 | -2.30942 | 0.37931 | 0.677917 | 5.09E-98 | turquoise | CTSS |
| 3.85E-96 | -2.03039 | 0.174877 | 0.536514 | 8.52E-96 | turquoise | AIF1 |
| 9.50E-95 | -2.49692 | 0.051724 | 0.430026 | 1.96E-94 | turquoise | FCN1 |
| 1.71E-94 | -1.94261 | 0.254926 | 0.588886 | 3.32E-94 | turquoise | PSAP |

| | | | | | | |
|---|---|---|---|---|---|---|
| 7.61E-92 | -4.12916 | 0.251232 | 0.573465 | 1.39E-91 | turquoise | LYZ |
| 6.26E-90 | -2.98163 | 0.916256 | 0.948502 | 1.08E-89 | turquoise | CD86 |
| 1.84E-87 | -3.08595 | 0.059113 | 0.418388 | 3.01E-87 | turquoise | S100A8 |
| 4.18E-86 | -1.88255 | 0.934729 | 0.967413 | 6.48E-86 | turquoise | FTL |
| 3.59E-84 | -1.88408 | 0.04064 | 0.390748 | 5.30E-84 | turquoise | VCAN |
| 3.86E-80 | -3.8168 | 0.183498 | 0.498982 | 5.44E-80 | turquoise | S100A9 |
| 9.65E-74 | -1.62341 | 0.20936 | 0.5016 | 1.30E-73 | turquoise | LST1 |
| 1.56E-68 | -1.63704 | 0.04064 | 0.343905 | 2.01E-68 | turquoise | IL1B |
| 1.37E-66 | -3.12355 | 0.998768 | 0.997672 | 1.70E-66 | turquoise | CD14 |
| 1.16E-65 | -1.67921 | 0.763547 | 0.846378 | 1.38E-65 | turquoise | NEAT1 |
| 1.01E-50 | -1.87966 | 0.061576 | 0.309281 | 1.16E-50 | turquoise | NKG7 |
| 1.64E-22 | -2.48891 | 0.921182 | 0.944719 | 1.82E-22 | turquoise | CD19 |
| 3.15E-18 | -2.4766 | 0.868227 | 0.898749 | 3.37E-18 | turquoise | CD16 |
| 1.91E-14 | -2.16742 | 0.992611 | 0.989526 | 1.97E-14 | turquoise | CD20 |
| 5.34E-13 | -1.58249 | 0.950739 | 0.954903 | 5.34E-13 | turquoise | CD56 |
| 0 | 1.930266 | 0.739857 | 0.044909 | 0 | yellow | GZMK |
| 1.83E-276 | 1.644729 | 0.966587 | 0.193734 | 2.48E-275 | yellow | CCL5 |
| 1.79E-60 | -2.05909 | 0.093079 | 0.508355 | 1.61E-59 | yellow | AIF1 |
| 6.84E-59 | -2.07747 | 0.078759 | 0.486162 | 4.62E-58 | yellow | TYROBP |
| 5.26E-56 | -1.57703 | 0.033413 | 0.436292 | 2.84E-55 | yellow | FCER1G |
| 1.71E-46 | -2.41262 | 0.042959 | 0.392167 | 7.69E-46 | yellow | FCN1 |
| 2.08E-46 | -1.8802 | 0.040573 | 0.389817 | 8.02E-46 | yellow | MNDA |
| 6.43E-45 | -2.66194 | 0.078759 | 0.414099 | 2.17E-44 | yellow | CST3 |
| 6.90E-44 | -2.58075 | 0.231504 | 0.523499 | 2.07E-43 | yellow | HLA-DRA |
| 7.93E-42 | -2.97874 | 0.057279 | 0.381723 | 2.14E-41 | yellow | S100A8 |
| 2.08E-41 | -1.80871 | 0.033413 | 0.355614 | 5.10E-41 | yellow | VCAN |
| 2.27E-40 | -3.97364 | 0.260143 | 0.539426 | 5.12E-40 | yellow | LYZ |
| 4.97E-40 | -2.3104 | 0.885442 | 0.948564 | 1.03E-39 | yellow | CD86 |
| 2.50E-38 | -1.56822 | 0.188544 | 0.47389 | 4.82E-38 | yellow | LST1 |
| 1.35E-36 | -3.71205 | 0.193317 | 0.465535 | 2.42E-36 | yellow | S100A9 |
| 4.68E-35 | -1.77506 | 0.319809 | 0.54752 | 7.91E-35 | yellow | PSAP |
| 6.90E-34 | -1.5507 | 0.033413 | 0.313577 | 1.10E-33 | yellow | IL1B |
| 5.15E-33 | -2.07619 | 0.460621 | 0.638381 | 7.73E-33 | yellow | CTSS |
| 1.24E-31 | -3.03568 | 1 | 0.99765 | 1.76E-31 | yellow | CD14 |
| 5.80E-26 | -1.77117 | 0.997613 | 0.999478 | 7.83E-26 | yellow | HLA-DR |
| 3.76E-24 | -1.95667 | 0.386635 | 0.530548 | 4.84E-24 | yellow | HLA-DRB1 |
| 3.11E-23 | -3.39639 | 0.997613 | 0.998172 | 3.82E-23 | yellow | CD11b |
| 3.57E-23 | -1.64707 | 0.983294 | 0.958747 | 4.19E-23 | yellow | FTL |
| 1.29E-20 | 2.010093 | 0.997613 | 0.98799 | 1.46E-20 | yellow | CD8a |
| 2.14E-14 | -1.75334 | 0.501193 | 0.561619 | 2.31E-14 | yellow | HLA-DPA1 |
| 9.74E-12 | -1.68811 | 0.696897 | 0.664752 | 1.01E-11 | yellow | CD74 |

| | | | | | | |
|---|---|---|---|---|---|---|
| 0.000342 | -1.6838 | 0.911695 | 0.943342 | 0.000342 | yellow | CD19 |
| 1.61E-262 | 1.711689 | 0.950249 | 0.197037 | 1.93E-261 | green | S100A12 |
| 5.89E-250 | 2.131024 | 0.987562 | 0.283078 | 3.54E-249 | green | S100A8 |
| 1.81E-209 | 2.013389 | 0.987562 | 0.381336 | 7.25E-209 | green | S100A9 |
| 6.05E-145 | 1.516113 | 1 | 0.99792 | 1.81E-144 | green | CD11b |
| 1.42E-83 | -1.70248 | 0.293532 | 0.730439 | 3.42E-83 | green | LTB |
| 1.76E-71 | -1.96722 | 0.997512 | 0.99844 | 3.53E-71 | green | CD3 |
| 3.76E-68 | -1.50134 | 0.997512 | 0.99792 | 6.44E-68 | green | CD28 |
| 3.15E-51 | -1.67579 | 0.977612 | 0.987523 | 4.72E-51 | green | CD27 |
| 2.12E-50 | -1.82824 | 0.995025 | 0.9987 | 2.82E-50 | green | CD45RA |
| 0.000691 | -2.14912 | 0.893035 | 0.892904 | 0.000829 | green | CD16 |
| 0.012005 | -2.15515 | 0.977612 | 0.936314 | 0.013096 | green | CD19 |
| 0.039519 | -2.0175 | 0.985075 | 0.990642 | 0.039519 | green | CD20 |
| 1.32E-208 | -2.20758 | 0.534943 | 0.869151 | 5.03E-207 | blue | S100A4 |
| 1.04E-179 | -2.67706 | 0.266836 | 0.759099 | 1.98E-178 | blue | CD74 |
| 9.83E-173 | -2.8908 | 0.044473 | 0.623628 | 1.25E-171 | blue | HLA-DRB1 |
| 1.35E-163 | -1.77761 | 0.317662 | 0.762854 | 1.28E-162 | blue | S100A11 |
| 1.56E-157 | -1.82191 | 0.07751 | 0.616407 | 1.18E-156 | blue | LGALS1 |
| 2.82E-152 | -2.33835 | 0.199492 | 0.68342 | 1.78E-151 | blue | HLA-DPB1 |
| 9.39E-150 | -2.42698 | 0.148666 | 0.64818 | 5.10E-149 | blue | HLA-DPA1 |
| 5.60E-149 | -3.12847 | 0.998729 | 0.999422 | 2.66E-148 | blue | HLA-DR |
| 1.76E-148 | -4.45134 | 0.996188 | 0.998556 | 7.45E-148 | blue | CD11b |
| 2.86E-141 | -3.071 | 0.092757 | 0.586077 | 1.09E-140 | blue | HLA-DRA |
| 2.96E-135 | -2.31606 | 0.045743 | 0.536973 | 1.02E-134 | blue | TYROBP |
| 3.74E-132 | -2.16892 | 0.64803 | 0.872039 | 1.19E-131 | blue | NEAT1 |
| 1.87E-119 | -2.40023 | 0.317662 | 0.689775 | 5.48E-119 | blue | CTSS |
| 2.99E-117 | -1.68108 | 0.027954 | 0.480358 | 8.11E-117 | blue | FCER1G |
| 3.99E-115 | -1.52496 | 0.827192 | 0.896303 | 1.01E-114 | blue | S100A6 |
| 8.10E-113 | -1.5542 | 0.11817 | 0.545927 | 1.92E-112 | blue | TYMP |
| 5.01E-106 | -1.53817 | 0.041931 | 0.463894 | 1.12E-105 | blue | FGL2 |
| 9.59E-106 | -1.67676 | 0.43202 | 0.728769 | 2.02E-105 | blue | COTL1 |
| 1.05E-99 | -2.78479 | 0.055909 | 0.454939 | 2.10E-99 | blue | CST3 |
| 1.38E-98 | -1.9416 | 0.232529 | 0.591566 | 2.61E-98 | blue | PSAP |
| 2.13E-97 | -1.97941 | 0.033037 | 0.428654 | 3.85E-97 | blue | MNDA |
| 5.72E-93 | -2.49252 | 0.047014 | 0.428365 | 9.88E-93 | blue | FCN1 |
| 1.49E-86 | -1.6788 | 0.162643 | 0.51011 | 2.47E-86 | blue | LST1 |
| 2.85E-86 | -3.84409 | 0.15756 | 0.5026 | 4.51E-86 | blue | S100A9 |
| 3.56E-85 | -3.07475 | 0.905972 | 0.950607 | 5.41E-85 | blue | CD86 |
| 7.56E-85 | -1.89812 | 0.031766 | 0.390237 | 1.11E-84 | blue | VCAN |
| 1.41E-82 | -4.06033 | 0.263024 | 0.568458 | 1.99E-82 | blue | LYZ |
| 2.19E-81 | -3.0727 | 0.067344 | 0.413923 | 2.97E-81 | blue | S100A8 |

| | | | | | | |
|---|---|---|---|---|---|---|
| 2.27E-78 | -1.84182 | 0.942821 | 0.965338 | 2.97E-78 | blue | FTL |
| 3.87E-71 | -1.75908 | 0.013977 | 0.328134 | 4.91E-71 | blue | CCL5 |
| 2.24E-66 | -1.62925 | 0.03939 | 0.341999 | 2.75E-66 | blue | IL1B |
| 4.31E-66 | -3.10616 | 0.994917 | 0.998556 | 5.12E-66 | blue | CD14 |
| 2.74E-63 | -1.93401 | 0.025413 | 0.315713 | 3.16E-63 | blue | NKG7 |
| 4.42E-29 | -2.52722 | 0.92249 | 0.944252 | 4.93E-29 | blue | CD19 |
| 2.31E-24 | -1.66569 | 0.426938 | 0.476603 | 2.51E-24 | blue | AIF1 |
| 4.41E-16 | -2.17129 | 0.992376 | 0.989601 | 4.65E-16 | blue | CD20 |
| 8.05E-16 | -1.64187 | 0.942821 | 0.956672 | 8.27E-16 | blue | CD56 |
| 1.66E-11 | -2.44895 | 0.869123 | 0.898325 | 1.66E-11 | blue | CD16 |
| 2.72E-150 | 1.850246 | 0.32 | 0.01309 | 2.18E-149 | pink | C1QA |
| 5.24E-111 | 1.804748 | 0.52 | 0.074339 | 2.09E-110 | pink | CDKN1C |
| 1.40E-94 | 2.24548 | 0.63 | 0.142257 | 3.72E-94 | pink | FCGR3A |
| 2.70E-59 | 1.616076 | 0.675 | 0.251914 | 5.40E-59 | pink | MS4A7 |
| 6.89E-40 | 1.764348 | 0.695 | 0.43344 | 1.10E-39 | pink | LST1 |
| 5.48E-37 | 1.534315 | 0.705 | 0.455668 | 7.30E-37 | pink | AIF1 |
| 2.41E-33 | 1.523902 | 1 | 0.999259 | 2.75E-33 | pink | HLA-DR |
| 0 | 3.056948 | 0.875969 | 0.034466 | 0 | purple | FCER1A |
| 0 | 1.520257 | 0.635659 | 0.01068 | 0 | purple | ENHO |
| 2.12E-276 | 2.633534 | 0.844961 | 0.048544 | 2.12E-275 | purple | CLEC10A |
| 1.11E-265 | 2.131767 | 0.782946 | 0.042476 | 8.30E-265 | purple | CD1C |
| 3.84E-149 | 3.181494 | 0.852713 | 0.117233 | 2.30E-148 | purple | HLA-DQA1 |
| 2.44E-110 | 2.862092 | 0.945736 | 0.24733 | 1.22E-109 | purple | HLA-DQB1 |
| 9.55E-98 | 1.682311 | 0.821705 | 0.166505 | 4.09E-97 | purple | BASP1 |
| 1.85E-90 | 2.516439 | 0.976744 | 0.362379 | 6.92E-90 | purple | CST3 |
| 4.46E-89 | 1.534171 | 0.96124 | 0.287621 | 1.49E-88 | purple | CCDC88A |
| 3.53E-86 | 1.901709 | 0.968992 | 0.36068 | 1.06E-85 | purple | HLA-DMA |
| 1.04E-81 | 1.783738 | 0.945736 | 0.265291 | 2.82E-81 | purple | IL1B |
| 5.41E-80 | 2.619512 | 0.984496 | 0.479369 | 1.35E-79 | purple | HLA-DRA |
| 1.19E-78 | 2.624039 | 0.992248 | 0.501456 | 2.75E-78 | purple | HLA-DRB1 |
| 1.18E-75 | 2.394117 | 0.992248 | 0.657767 | 2.53E-75 | purple | CD74 |
| 2.37E-75 | 2.781922 | 0.976744 | 0.542476 | 4.73E-75 | purple | HLA-DPA1 |
| 2.83E-75 | 1.640237 | 0.930233 | 0.287621 | 5.30E-75 | purple | CPVL |
| 4.29E-75 | 1.549795 | 0.945736 | 0.340049 | 7.57E-75 | purple | HLA-DMB |
| 5.47E-74 | 1.698048 | 0.96124 | 0.399515 | 9.11E-74 | purple | CTSZ |
| 2.57E-72 | 2.819087 | 0.976744 | 0.581796 | 4.05E-72 | purple | HLA-DPB1 |
| 1.22E-58 | 1.562241 | 0.945736 | 0.505825 | 1.84E-58 | purple | ANXA2 |
| 3.64E-57 | 1.596383 | 0.96124 | 0.50267 | 5.20E-57 | purple | LGALS1 |
| 2.01E-55 | 1.7448 | 1 | 0.999272 | 2.74E-55 | purple | HLA-DR |
| 2.20E-54 | 1.530723 | 0.992248 | 0.496845 | 2.87E-54 | purple | LYZ |
| 8.38E-52 | 1.582583 | 0.984496 | 0.916019 | 1.05E-51 | purple | VIM |

| | | | | | | |
|---|---|---|---|---|---|---|
| 3.38E-50 | 1.549932 | 0.984496 | 0.804126 | 4.06E-50 | purple | S100A10 |
| 1.92E-28 | -1.72203 | 0.968992 | 0.990534 | 2.21E-28 | purple | CD278 |
| 4.74E-27 | -1.62278 | 0.992248 | 0.998058 | 5.27E-27 | purple | CD28 |
| 1.76E-25 | -1.81855 | 0.930233 | 0.979369 | 1.88E-25 | purple | CD127 |
| 1.82E-21 | -2.28894 | 0.968992 | 0.987136 | 1.88E-21 | purple | CD27 |
| 4.51E-11 | -2.08475 | 1 | 0.998301 | 4.51E-11 | purple | CD3 |
| 0 | 3.878517 | 0.72561 | 0.037745 | 0 | red | IGKC |
| 0 | 2.808865 | 0.783537 | 0.018873 | 0 | red | IGHM |
| 0 | 2.118146 | 0.893293 | 0.019638 | 0 | red | CD79A |
| 0 | 2.062413 | 0.878049 | 0.016832 | 0 | red | MS4A1 |
| 0 | 1.889476 | 0.881098 | 0.120377 | 0 | red | CD79B |
| 0 | 1.552685 | 0.810976 | 0.017853 | 0 | red | BANK1 |
| 0 | 1.514804 | 0.414634 | 0.003315 | 0 | red | TCL1A |
| 4.30E-269 | 2.359806 | 0.341463 | 0.00306 | 2.04E-268 | red | IGLC2 |
| 2.37E-234 | 1.968135 | 0.307927 | 0.003571 | 1.00E-233 | red | IGLC3 |
| 6.84E-151 | 4.04519 | 0.996951 | 0.989543 | 2.60E-150 | red | CD20 |
| 7.20E-150 | 4.183018 | 0.984756 | 0.936496 | 2.49E-149 | red | CD19 |
| 8.71E-107 | -2.15807 | 0.942073 | 0.993624 | 2.76E-106 | red | CD45RO |
| 3.21E-102 | -2.38734 | 0.993902 | 0.99694 | 9.38E-102 | red | CD4 |
| 4.30E-82 | -2.05635 | 0.484756 | 0.834226 | 1.17E-81 | red | S100A4 |
| 7.56E-76 | -2.71168 | 0.996951 | 0.99847 | 1.92E-75 | red | CD3 |
| 2.41E-67 | -1.50215 | 0.615854 | 0.905891 | 5.73E-67 | red | S100A6 |
| 5.53E-67 | -1.54076 | 0.97561 | 0.991074 | 1.24E-66 | red | CD278 |
| 2.49E-62 | -1.84572 | 0.954268 | 0.979852 | 5.26E-62 | red | CD127 |
| 2.43E-61 | -1.54603 | 0.280488 | 0.713849 | 4.86E-61 | red | S100A11 |
| 7.55E-53 | -1.96577 | 0.579268 | 0.851568 | 1.43E-52 | red | NEAT1 |
| 1.32E-44 | -1.8506 | 0.097561 | 0.498342 | 2.39E-44 | red | AIF1 |
| 1.51E-37 | -1.75083 | 0.121951 | 0.473094 | 2.61E-37 | red | TYROBP |
| 1.82E-35 | -2.37079 | 0.045732 | 0.383831 | 3.01E-35 | red | FCN1 |
| 5.83E-32 | -2.9606 | 0.060976 | 0.373884 | 9.23E-32 | red | S100A8 |
| 3.45E-31 | -1.78162 | 0.039634 | 0.347615 | 5.25E-31 | red | VCAN |
| 5.84E-29 | -3.69009 | 0.189024 | 0.459577 | 8.54E-29 | red | S100A9 |
| 1.50E-26 | -3.86553 | 0.304878 | 0.529202 | 2.11E-26 | red | LYZ |
| 2.89E-26 | -1.51187 | 0.216463 | 0.464932 | 3.92E-26 | red | LST1 |
| 3.08E-26 | -1.75346 | 0.112805 | 0.375669 | 4.04E-26 | red | MNDA |
| 4.05E-24 | -2.8654 | 0.993902 | 0.998215 | 5.13E-24 | red | CD14 |
| 1.05E-23 | -1.7458 | 0.134146 | 0.401683 | 1.29E-23 | red | CST3 |
| 1.62E-21 | -1.57259 | 0.051829 | 0.288192 | 1.92E-21 | red | CCL5 |
| 3.55E-21 | -3.53353 | 0.996951 | 0.998215 | 4.09E-21 | red | CD11b |
| 5.12E-19 | -1.75144 | 0.060976 | 0.278755 | 5.72E-19 | red | NKG7 |
| 0.000137 | -2.26421 | 0.859756 | 0.89569 | 0.000149 | red | CD16 |

| | | | | | | |
|---|---|---|---|---|---|---|
| 0.01759 | -1.88254 | 0.978659 | 0.939301 | 0.018567 | red | CD86 |
| 0.018619 | -1.60737 | 0.981707 | 0.989543 | 0.019123 | red | CD8a |
| 4.23E-295 | 1.884026 | 0.90583 | 0.085693 | 1.44E-293 | black | GZMH |
| 6.50E-255 | 1.646382 | 0.829596 | 0.07998 | 1.11E-253 | black | FGFBP2 |
| 3.15E-221 | 1.913751 | 0.843049 | 0.100099 | 3.57E-220 | black | GNLY |
| 1.52E-192 | 1.531231 | 0.950673 | 0.172628 | 1.29E-191 | black | CST7 |
| 5.77E-186 | 2.026789 | 0.991031 | 0.22156 | 3.92E-185 | black | NKG7 |
| 3.06E-167 | 1.770289 | 0.977578 | 0.23075 | 1.73E-166 | black | CCL5 |
| 2.14E-68 | -1.56719 | 0.991031 | 0.997019 | 1.04E-67 | black | CD4 |
| 5.62E-66 | 1.57046 | 0.973094 | 0.856185 | 2.39E-65 | black | TIGIT |
| 5.79E-48 | 2.254625 | 0.950673 | 0.889717 | 2.19E-47 | black | CD16 |
| 7.17E-41 | -1.52264 | 0.309417 | 0.710134 | 2.44E-40 | black | LTB |
| 1.15E-40 | 1.91702 | 0.973094 | 0.953055 | 3.56E-40 | black | CD56 |
| 2.21E-38 | -1.66206 | 0.300448 | 0.694486 | 6.27E-38 | black | COTL1 |
| 2.13E-35 | -2.09411 | 0.058296 | 0.490065 | 5.56E-35 | black | AIF1 |
| 9.70E-28 | -1.64715 | 0.107623 | 0.464481 | 2.36E-27 | black | LST1 |
| 2.86E-26 | -3.75662 | 0.112108 | 0.456781 | 6.47E-26 | black | S100A9 |
| 2.96E-25 | -1.85888 | 0.03139 | 0.373323 | 6.28E-25 | black | MNDA |
| 4.94E-25 | -2.37213 | 0.035874 | 0.375559 | 9.88E-25 | black | FCN1 |
| 1.11E-23 | -1.77807 | 0.013453 | 0.341033 | 2.09E-23 | black | VCAN |
| 4.45E-23 | -2.7633 | 0.923767 | 0.943368 | 7.97E-23 | black | CD86 |
| 1.15E-22 | -2.61629 | 0.089686 | 0.397168 | 1.96E-22 | black | CST3 |
| 2.20E-20 | -3.91465 | 0.278027 | 0.524839 | 3.56E-20 | black | LYZ |
| 6.50E-20 | -2.90843 | 0.080717 | 0.36463 | 1.00E-19 | black | S100A8 |
| 1.99E-19 | -1.79097 | 0.982063 | 0.96001 | 2.94E-19 | black | FTL |
| 1.08E-18 | -2.58214 | 0.295964 | 0.505713 | 1.53E-18 | black | HLA-DRA |
| 8.01E-16 | -2.04827 | 0.484305 | 0.628415 | 1.09E-15 | black | CTSS |
| 1.33E-15 | -1.95579 | 0.58296 | 0.672628 | 1.73E-15 | black | CD74 |
| 4.28E-15 | -2.85405 | 0.995516 | 0.998013 | 5.39E-15 | black | CD14 |
| 1.94E-12 | -1.6129 | 0.399103 | 0.532042 | 2.36E-12 | black | PSAP |
| 6.00E-10 | -2.30533 | 1 | 0.999255 | 7.04E-10 | black | HLA-DR |
| 3.42E-07 | -1.69063 | 0.878924 | 0.943616 | 3.88E-07 | black | CD19 |
| 9.55E-06 | -1.88769 | 1 | 0.998013 | 1.05E-05 | black | CD11b |
| 0.000671 | -1.83667 | 0.591928 | 0.512171 | 0.000713 | black | HLA-DRB1 |
| 0.010728 | -1.52399 | 0.623318 | 0.551913 | 0.011053 | black | HLA-DPA1 |
| 0.022932 | -1.70316 | 0.982063 | 0.990561 | 0.022932 | black | CD20 |
| 0 | 3.075744 | 0.994792 | 0.098595 | 0 | magenta | GNLY |
| 0 | 1.947495 | 0.979167 | 0.071481 | 0 | magenta | GZMB |
| 0 | 1.905935 | 0.932292 | 0.070495 | 0 | magenta | SPON2 |
| 0 | 1.641295 | 0.942708 | 0.061622 | 0 | magenta | KLRF1 |
| 6.58E-291 | 1.706764 | 0.90625 | 0.077644 | 5.77E-290 | magenta | CLIC3 |

| | | | | | | |
|---|---|---|---|---|---|---|
| 6.66E-291 | 1.763303 | 0.984375 | 0.093912 | 5.77E-290 | magenta | KLRD1 |
| 1.84E-274 | 2.024697 | 0.901042 | 0.082327 | 1.36E-273 | magenta | FGFBP2 |
| 8.29E-252 | 1.891258 | 0.994792 | 0.129406 | 5.39E-251 | magenta | PRF1 |
| 3.26E-177 | 2.329661 | 1 | 0.227015 | 1.88E-176 | magenta | NKG7 |
| 1.77E-175 | 1.533617 | 0.963542 | 0.177964 | 9.19E-175 | magenta | CST7 |
| 6.82E-171 | 1.693814 | 0.973958 | 0.188563 | 3.22E-170 | magenta | GZMA |
| 2.35E-170 | 1.542946 | 0.984375 | 0.205571 | 1.02E-169 | magenta | CTSW |
| 6.07E-143 | 1.743508 | 0.848958 | 0.148878 | 2.43E-142 | magenta | CCL4 |
| 6.21E-111 | 2.729956 | 1 | 0.951935 | 2.31E-110 | magenta | CD56 |
| 5.56E-104 | 2.934229 | 0.994792 | 0.888095 | 1.93E-103 | magenta | CD16 |
| 2.67E-96 | 2.162838 | 1 | 0.953414 | 8.69E-96 | magenta | CD335 |
| 8.34E-88 | 1.57542 | 1 | 0.998275 | 2.55E-87 | magenta | CD45RA |
| 9.04E-78 | -3.6456 | 0.984375 | 0.997289 | 2.61E-77 | magenta | CD4 |
| 2.22E-74 | -2.46898 | 0.953125 | 0.991373 | 6.08E-74 | magenta | CD45RO |
| 1.03E-58 | -1.55851 | 0.59375 | 0.9263 | 2.68E-58 | magenta | FOSB |
| 1.37E-49 | -2.62514 | 0.984375 | 0.999014 | 3.39E-49 | magenta | CD3 |
| 3.61E-48 | -1.6522 | 0.729167 | 0.92704 | 8.54E-48 | magenta | VIM |
| 1.07E-45 | -2.71613 | 0.953125 | 0.988169 | 2.42E-45 | magenta | CD27 |
| 1.48E-44 | -1.64503 | 0.963542 | 0.98669 | 3.21E-44 | magenta | CD62L |
| 9.32E-43 | -1.77528 | 0.1875 | 0.69682 | 1.94E-42 | magenta | COTL1 |
| 2.42E-41 | -1.70211 | 0.979167 | 0.964013 | 4.84E-41 | magenta | FTH1 |
| 2.55E-41 | -1.92189 | 0.9375 | 0.979788 | 4.91E-41 | magenta | CD127 |
| 4.18E-38 | -1.75079 | 0.307292 | 0.707173 | 7.75E-38 | magenta | LTB |
| 8.39E-34 | -2.14429 | 0.020833 | 0.488538 | 1.51E-33 | magenta | AIF1 |
| 4.68E-32 | -1.82089 | 0.994792 | 0.998028 | 8.11E-32 | magenta | CD28 |
| 1.88E-23 | -1.88641 | 0.973958 | 0.960562 | 3.15E-23 | magenta | FTL |
| 1.19E-22 | -2.62638 | 1 | 0.999261 | 1.93E-22 | magenta | HLA-DR |
| 2.75E-22 | -3.03097 | 0.880208 | 0.94528 | 4.33E-22 | magenta | CD86 |
| 3.59E-22 | -2.71589 | 0.203125 | 0.508504 | 5.49E-22 | magenta | HLA-DRA |
| 4.38E-22 | -1.58643 | 0.125 | 0.460932 | 6.51E-22 | magenta | LST1 |
| 1.56E-20 | -1.82735 | 0.046875 | 0.369978 | 2.26E-20 | magenta | MNDA |
| 1.60E-19 | -2.33754 | 0.0625 | 0.371703 | 2.24E-19 | magenta | FCN1 |
| 1.12E-18 | -2.94037 | 0.0625 | 0.363323 | 1.53E-18 | magenta | S100A8 |
| 1.29E-18 | -3.70431 | 0.166667 | 0.451565 | 1.71E-18 | magenta | S100A9 |
| 1.46E-18 | -3.8913 | 0.260417 | 0.523786 | 1.89E-18 | magenta | LYZ |
| 1.64E-18 | -1.74601 | 0.036458 | 0.337441 | 2.08E-18 | magenta | VCAN |
| 2.80E-17 | -2.55607 | 0.119792 | 0.393394 | 3.46E-17 | magenta | CST3 |
| 3.43E-16 | -1.51342 | 0.026042 | 0.29825 | 4.14E-16 | magenta | IL1B |
| 6.70E-16 | -2.00865 | 0.53125 | 0.67439 | 7.92E-16 | magenta | CD74 |
| 1.56E-14 | -2.16154 | 0.34375 | 0.524526 | 1.80E-14 | magenta | HLA-DRB1 |
| 3.65E-11 | -2.00991 | 0.557292 | 0.62386 | 4.12E-11 | magenta | CTSS |

| | | | | | | |
|---|---|---|---|---|---|---|
| 5.40E-10 | -2.8677 | 0.994792 | 0.998028 | 5.97E-10 | magenta | CD14 |
| 3.72E-08 | -1.5595 | 0.453125 | 0.528469 | 4.02E-08 | magenta | PSAP |
| 9.01E-08 | -1.76782 | 0.5 | 0.558294 | 9.56E-08 | magenta | HLA-DPA1 |
| 1.56E-06 | -1.60387 | 0.880208 | 0.943061 | 1.62E-06 | magenta | CD19 |
| 0.000133 | -1.51462 | 0.604167 | 0.593296 | 0.000136 | magenta | HLA-DPB1 |
| 0.000813 | -2.03425 | 0.979167 | 0.990633 | 0.000813 | magenta | CD20 |

**Sup. Tab. 4:** DE genes 10X pbmcs a) 1vs all b)

| p_val | avg_logFC | pct.1 | pct.2 | p_val_adj | group1 | gene |
|---|---|---|---|---|---|---|
| 0 | 3.527432 | 0.96181 | 0.054718 | 0 | green | DNTT |
| 0 | 3.145648 | 0.980198 | 0.056134 | 0 | green | VPREB1 |
| 0 | 2.720039 | 0.958982 | 0.066504 | 0 | green | VPREB3 |
| 0 | 2.620183 | 0.879774 | 0.071245 | 0 | green | CD9 |
| 0 | 2.601256 | 0.954738 | 0.239317 | 0 | green | IGLL1 |
| 0 | 2.374099 | 0.967468 | 0.177191 | 0 | green | CD79B |
| 0 | 2.200937 | 0.794908 | 0.183677 | 0 | green | MT1X |
| 0 | 2.15993 | 0.857143 | 0.230888 | 0 | green | SMIM3 |
| 0 | 2.025457 | 0.770863 | 0.014552 | 0 | green | ARPP21 |
| 0 | 1.826118 | 0.885431 | 0.185521 | 0 | green | MZB1 |
| 0 | 1.817024 | 0.745403 | 0.035524 | 0 | green | IGHV5-78 |
| 0 | 1.729646 | 0.553041 | 0.00912 | 0 | green | DEPP1 |
| 0 | 1.663562 | 0.72843 | 0.053829 | 0 | green | CD24 |
| 0 | 1.654976 | 0.678925 | 0.023112 | 0 | green | MME |
| 0 | 1.633568 | 0.824611 | 0.298808 | 0 | green | SSBP2 |
| 0 | 1.605813 | 0.685997 | 0.179397 | 0 | green | LHPP |
| 0 | 1.603649 | 0.937765 | 0.139099 | 0 | green | CD79A |
| 0 | 1.593174 | 0.589816 | 0.010173 | 0 | green | LINC01013 |
| 0 | 1.557855 | 0.81471 | 0.24916 | 0 | green | TCF4 |
| 0 | 1.518797 | 0.712871 | 0.187167 | 0 | green | SOCS2 |
| 0 | 1.458643 | 0.848656 | 0.321426 | 0 | green | RCSD1 |
| 0 | 1.439307 | 0.725601 | 0.246724 | 0 | green | TOP2B |
| 0 | 1.425062 | 0.988685 | 0.925331 | 0 | green | LAPTM5 |
| 0 | 1.409306 | 1 | 0.592875 | 0 | green | HLA-DRA |
| 0 | 1.297089 | 0.514851 | 0.031112 | 0 | green | NEIL1 |
| 0 | 1.295366 | 0.577086 | 0.05159 | 0 | green | LINC00426 |
| 0 | 1.285237 | 0.677511 | 0.197768 | 0 | green | JUP |
| 0 | 1.267424 | 0.565771 | 0.034174 | 0 | green | EBF1 |
| 0 | 1.25004 | 0.480905 | 0.063014 | 0 | green | SMAD1 |
| 0 | 1.247927 | 0.48232 | 0.012642 | 0 | green | AKAP12 |
| 0 | 1.219389 | 0.4314 | 0.016033 | 0 | green | CMTM8 |
| 0 | 1.21015 | 0.353607 | 0.002403 | 0 | green | RPS4Y2 |
| 0 | 1.202214 | 0.585573 | 0.05001 | 0 | green | POU2AF1 |
| 0 | 1.197015 | 0.390382 | 0.009021 | 0 | green | RAG1 |
| 0 | 1.184303 | 0.478076 | 0.013861 | 0 | green | RAG2 |
| 0 | 1.090411 | 0.441301 | 0.053006 | 0 | green | TP53INP1 |
| 0 | 0.980273 | 0.486563 | 0.082702 | 0 | green | BLNK |
| 0 | 0.979866 | 0.356436 | 0.005926 | 0 | green | P4HA2 |
| 0 | 0.910221 | 0.323904 | 0.002173 | 0 | green | DUSP26 |

| | | | | | | |
|---|---|---|---|---|---|---|
| 0 | 0.834822 | 0.295615 | 0.004082 | 0 | green | HMHB1 |
| 0 | 0.784942 | 0.272984 | 0.002403 | 0 | green | SH2D4B |
| 0 | 0.765779 | 0.343706 | 0.014782 | 0 | green | IGHV5-51 |
| 0 | 0.754261 | 0.302687 | 0.011918 | 0 | green | LINC02202 |
| 0 | 0.703976 | 0.288543 | 0.021202 | 0 | green | MYLK |
| 6.02E-308 | 1.286659 | 1 | 0.917001 | 5.31E-307 | green | CD74 |
| 3.20E-291 | 1.365148 | 0.680339 | 0.218114 | 2.76E-290 | green | HPS4 |
| 5.05E-285 | 1.317031 | 0.871287 | 0.381543 | 4.27E-284 | green | SOX4 |
| 2.31E-284 | 1.082835 | 0.544554 | 0.125305 | 1.91E-283 | green | GNG7 |
| 2.18E-279 | 1.24057 | 0.659123 | 0.198262 | 1.77E-278 | green | TSPAN14 |
| 2.50E-278 | 1.032674 | 0.50495 | 0.105386 | 1.98E-277 | green | SH3TC1 |
| 2.30E-267 | 0.815264 | 0.338048 | 0.048002 | 1.79E-266 | green | MILR1 |
| 1.65E-263 | 0.900531 | 0.335219 | 0.048298 | 1.26E-262 | green | GAB1 |
| 3.72E-261 | -0.50076 | 1 | 0.99977 | 2.77E-260 | green | RPL41 |
| 3.77E-261 | -0.60447 | 0.997171 | 0.999407 | 2.77E-260 | green | RPS14 |
| 2.13E-260 | 1.250656 | 0.773692 | 0.320109 | 1.54E-259 | green | TMEM243 |
| 5.22E-260 | -0.72291 | 0.995757 | 0.998815 | 3.70E-259 | green | RPL12 |
| 4.11E-255 | 1.042688 | 0.978784 | 0.645157 | 2.86E-254 | green | HLA-DPB1 |
| 8.02E-252 | 1.238562 | 0.763791 | 0.321229 | 5.49E-251 | green | LAT2 |
| 1.87E-249 | -1.92179 | 0.183876 | 0.84253 | 1.26E-248 | green | SRGN |
| 3.64E-249 | 0.979505 | 0.425743 | 0.084513 | 2.41E-248 | green | ELK3 |
| 9.16E-248 | 1.198705 | 0.967468 | 0.806545 | 5.96E-247 | green | CD99 |
| 1.15E-247 | 1.231398 | 0.855728 | 0.410285 | 7.34E-247 | green | BTG2 |
| 1.68E-242 | 1.074743 | 0.483734 | 0.112695 | 1.06E-241 | green | HIST1H2BK |
| 1.32E-241 | -2.31527 | 0.178218 | 0.826463 | 8.21E-241 | green | S100A4 |
| 1.08E-236 | -2.28263 | 0.192362 | 0.83848 | 6.58E-236 | green | S100A6 |
| 8.38E-227 | 1.150045 | 0.58133 | 0.182788 | 5.04E-226 | green | MLXIP |
| 5.69E-226 | 1.041513 | 0.499293 | 0.127543 | 3.37E-225 | green | QRSL1 |
| 1.25E-225 | 0.951188 | 0.636492 | 0.185224 | 7.29E-225 | green | IGHM |
| 5.18E-224 | -1.02346 | 0.773692 | 0.971192 | 2.98E-223 | green | EEF1B2 |
| 2.21E-221 | -1.32185 | 0.149929 | 0.819912 | 1.26E-220 | green | LSP1 |
| 3.57E-220 | 0.733951 | 0.343706 | 0.056529 | 1.99E-219 | green | RUBCNL |
| 2.17E-219 | 1.126169 | 0.519095 | 0.149536 | 1.20E-218 | green | ERG |
| 6.00E-219 | -0.57229 | 0.998586 | 0.999309 | 3.26E-218 | green | RPL32 |
| 8.69E-219 | -1.49731 | 0.079208 | 0.746823 | 4.66E-218 | green | EMP3 |
| 9.44E-219 | -0.59296 | 0.997171 | 0.999045 | 4.99E-218 | green | RPL13 |
| 1.23E-214 | -0.63999 | 0.998586 | 0.999868 | 6.44E-214 | green | RPLP1 |
| 9.72E-212 | 0.673652 | 0.998586 | 0.992428 | 5.01E-211 | green | H3F3B |
| 1.73E-206 | -1.66953 | 0.053748 | 0.702114 | 8.82E-206 | green | ANXA1 |
| 3.68E-205 | 0.999152 | 0.471004 | 0.11928 | 1.85E-204 | green | LDLRAD4 |
| 3.80E-205 | 1.148673 | 0.770863 | 0.411339 | 1.88E-204 | green | SNX2 |

| | | | | | | |
|---|---|---|---|---|---|---|
| 3.51E-202 | -0.6149 | 0.971711 | 0.994337 | 1.72E-201 | green | RPL36 |
| 4.61E-198 | 1.190428 | 0.705799 | 0.327583 | 2.23E-197 | green | GLRX |
| 3.86E-197 | 1.2039 | 0.680339 | 0.290775 | 1.85E-196 | green | MEF2C |
| 9.76E-196 | -0.89542 | 0.777935 | 0.984427 | 4.61E-195 | green | PFN1 |
| 1.60E-193 | 1.157252 | 0.695898 | 0.326562 | 7.48E-193 | green | MPP1 |
| 9.12E-193 | 1.008416 | 0.850071 | 0.490749 | 4.21E-192 | green | HLA-DMA |
| 1.20E-192 | 0.992977 | 0.363508 | 0.077533 | 5.47E-192 | green | SCHIP1 |
| 1.39E-190 | 0.760898 | 0.960396 | 0.643676 | 6.28E-190 | green | HLA-DPA1 |
| 2.79E-188 | 0.98775 | 0.902405 | 0.633206 | 1.25E-187 | green | TKT |
| 1.70E-185 | 0.706201 | 0.983027 | 0.913116 | 7.48E-185 | green | CCNI |
| 3.06E-183 | -1.30966 | 0.971711 | 0.99707 | 1.34E-182 | green | FTH1 |
| 3.94E-182 | 0.849175 | 0.394625 | 0.093896 | 1.70E-181 | green | DBN1 |
| 5.16E-182 | 0.556125 | 0.261669 | 0.039014 | 2.20E-181 | green | PAX5 |
| 5.65E-181 | -1.15349 | 0.039604 | 0.654474 | 2.39E-180 | green | CNN2 |
| 1.25E-180 | 0.573453 | 0.98727 | 0.969546 | 5.24E-180 | green | ARHGDIB |
| 2.98E-180 | -1.09909 | 0.057992 | 0.666985 | 1.23E-179 | green | CD44 |
| 5.89E-180 | -1.45626 | 0.055163 | 0.657799 | 2.41E-179 | green | ITGB2 |
| 8.91E-180 | -0.53975 | 0.991513 | 0.99572 | 3.61E-179 | green | RPL14 |
| 1.06E-179 | 0.816308 | 0.384724 | 0.088332 | 4.26E-179 | green | HLA-DOA |
| 1.12E-177 | -0.68631 | 0.994342 | 0.996444 | 4.43E-177 | green | RPS18 |
| 5.16E-177 | -1.92133 | 0.236209 | 0.783697 | 2.03E-176 | green | JUNB |
| 1.38E-175 | 1.079391 | 0.486563 | 0.154408 | 5.36E-175 | green | ENG |
| 1.28E-174 | 1.094112 | 0.586987 | 0.230329 | 4.92E-174 | green | CD38 |
| 1.06E-171 | 0.960653 | 0.743989 | 0.381741 | 4.04E-171 | green | CYFIP2 |
| 4.23E-171 | -1.43282 | 0.08628 | 0.670837 | 1.60E-170 | green | HCST |
| 1.47E-170 | 1.12759 | 0.602546 | 0.261474 | 5.51E-170 | green | MAGED1 |
| 6.56E-170 | 0.765165 | 0.994342 | 0.972444 | 2.43E-169 | green | ACTG1 |
| 1.51E-169 | 0.863849 | 0.919378 | 0.761704 | 5.53E-169 | green | PNISR |
| 1.57E-167 | 1.173986 | 0.666195 | 0.328834 | 5.72E-167 | green | UBE2J1 |
| 3.46E-166 | 1.069959 | 0.584158 | 0.242181 | 1.25E-165 | green | ERGIC1 |
| 6.97E-163 | -0.80629 | 0.487977 | 0.915289 | 2.49E-162 | green | PPDPF |
| 7.85E-162 | 0.893707 | 0.411598 | 0.113354 | 2.78E-161 | green | TCF3 |
| 1.43E-161 | 1.01292 | 0.561528 | 0.217291 | 5.03E-161 | green | WASHC4 |
| 5.39E-161 | -1.00392 | 0.964639 | 0.994798 | 1.88E-160 | green | FTL |
| 5.01E-160 | 0.89718 | 0.854314 | 0.575196 | 1.73E-159 | green | 9-Sep |
| 3.79E-159 | 0.559819 | 0.994342 | 0.949134 | 1.30E-158 | green | H3F3A |
| 9.50E-158 | -1.49821 | 0.015559 | 0.576612 | 3.22E-157 | green | CTSW |
| 3.78E-156 | 0.757576 | 0.756719 | 0.359189 | 1.27E-155 | green | HLA-DQB1 |
| 3.83E-156 | 0.935923 | 0.497878 | 0.167742 | 1.28E-155 | green | GBP4 |
| 3.82E-152 | 0.644649 | 0.937765 | 0.573583 | 1.26E-151 | green | HLA-DRB5 |
| 7.93E-152 | 0.705513 | 0.323904 | 0.071541 | 2.60E-151 | green | SYNE3 |

| | | | | | | |
|---|---|---|---|---|---|---|
| 9.76E-151 | 0.619339 | 0.29703 | 0.058932 | 3.17E-150 | green | CD72 |
| 5.28E-147 | -0.90782 | 0.128713 | 0.676269 | 1.70E-146 | green | ICAM3 |
| 1.51E-146 | 1.046218 | 0.710042 | 0.404622 | 4.83E-146 | green | NR3C1 |
| 9.09E-144 | 0.970405 | 0.697313 | 0.375387 | 2.89E-143 | green | PPP1R18 |
| 2.18E-141 | -1.98245 | 0.074965 | 0.584513 | 6.87E-141 | green | IFITM1 |
| 8.51E-141 | 0.850644 | 0.434229 | 0.140252 | 2.66E-140 | green | DOK3 |
| 2.01E-140 | -1.09166 | 0.243281 | 0.759729 | 6.23E-140 | green | PTPRC |
| 3.61E-140 | 1.039156 | 0.605375 | 0.28745 | 1.11E-139 | green | ZCCHC7 |
| 4.57E-140 | 0.595055 | 0.384724 | 0.095509 | 1.39E-139 | green | FAM129C |
| 2.69E-138 | -1.14069 | 0.029703 | 0.549746 | 8.16E-138 | green | FLNA |
| 5.04E-138 | 1.001311 | 0.606789 | 0.291993 | 1.52E-137 | green | MDM4 |
| 7.76E-136 | 0.762717 | 0.282885 | 0.061961 | 2.32E-135 | green | BTBD3 |
| 8.08E-135 | -0.57699 | 0.988685 | 0.993613 | 2.39E-134 | green | RPS6 |
| 5.34E-132 | 0.507741 | 0.263083 | 0.051458 | 1.57E-131 | green | CD22 |
| 6.60E-132 | -1.64348 | 0.123055 | 0.619247 | 1.93E-131 | green | S100A10 |
| 6.44E-131 | -0.51488 | 0.922207 | 0.981662 | 1.87E-130 | green | RPL37A |
| 1.67E-130 | 0.535797 | 0.950495 | 0.915421 | 4.81E-130 | green | SRP14 |
| 5.40E-130 | -0.87605 | 0.043847 | 0.546454 | 1.54E-129 | green | RGS10 |
| 5.02E-129 | 0.672781 | 0.916549 | 0.839567 | 1.42E-128 | green | GMFG |
| 4.40E-128 | 0.659509 | 0.302687 | 0.072825 | 1.24E-127 | green | AEBP1 |
| 9.60E-128 | -0.9069 | 0.582744 | 0.899783 | 2.69E-127 | green | RPS2 |
| 1.23E-127 | -0.93478 | 0.007072 | 0.496115 | 3.41E-127 | green | BIN2 |
| 5.74E-127 | -1.25468 | 0.018388 | 0.507605 | 1.58E-126 | green | FYB1 |
| 2.57E-126 | 0.911006 | 0.462518 | 0.168927 | 7.03E-126 | green | HIST1H2AC |
| 1.70E-125 | 0.619511 | 0.923621 | 0.831138 | 4.62E-125 | green | HNRNPK |
| 3.97E-123 | 1.047945 | 0.588402 | 0.284618 | 1.07E-122 | green | HIST1H1C |
| 2.19E-122 | 0.910389 | 0.612447 | 0.298775 | 5.87E-122 | green | EGFL7 |
| 1.11E-116 | 1.017711 | 0.574257 | 0.272404 | 2.95E-116 | green | CD34 |
| 4.82E-115 | 0.942287 | 0.652051 | 0.407289 | 1.28E-114 | green | ZNF22 |
| 2.87E-113 | -0.77906 | 0.48232 | 0.864588 | 7.55E-113 | green | RAC2 |
| 3.54E-112 | -0.63022 | 0.994342 | 0.994601 | 9.24E-112 | green | RPS27 |
| 6.04E-112 | 0.788774 | 0.359264 | 0.115362 | 1.57E-111 | green | RIMKLB |
| 4.01E-110 | -1.14071 | 0.018388 | 0.464608 | 1.03E-109 | green | AHNAK |
| 1.73E-108 | 0.911531 | 0.650636 | 0.392309 | 4.44E-108 | green | MAP1LC3B |
| 2.35E-106 | 0.938655 | 0.512023 | 0.247942 | 5.98E-106 | green | YBX3 |
| 2.81E-106 | -0.8453 | 0.100424 | 0.555738 | 7.10E-106 | green | PLP2 |
| 7.81E-102 | 0.754124 | 0.545969 | 0.248765 | 1.96E-101 | green | PTPRE |
| 2.73E-100 | -0.54368 | 0.724187 | 0.926516 | 6.83E-100 | green | RPL38 |
| 3.60E-100 | -1.021 | 0.056577 | 0.484658 | 8.93E-100 | green | MT2A |
| 3.85E-100 | -0.76014 | 0.364922 | 0.7487 | 9.49E-100 | green | TPI1 |
| 6.30E-100 | -0.73625 | 0.082037 | 0.523342 | 1.54E-99 | green | PAXX |

| | | | | | | |
|---|---|---|---|---|---|---|
| 2.61E-98 | -1.44722 | 0.009901 | 0.419207 | 6.35E-98 | green | CD7 |
| 5.73E-98 | 0.65696 | 0.793494 | 0.631856 | 1.39E-97 | green | ATP5F1C |
| 2.80E-97 | -0.72699 | 0.203678 | 0.66096 | 6.73E-97 | green | LY6E |
| 2.94E-97 | 0.694096 | 0.270156 | 0.072365 | 7.04E-97 | green | PPP1R14B |
| 1.15E-96 | 0.738575 | 0.635078 | 0.323435 | 2.73E-96 | green | ISG20 |
| 1.32E-96 | -0.57276 | 0.632249 | 0.895371 | 3.11E-96 | green | RPL27A |
| 2.06E-96 | 0.811993 | 0.677511 | 0.46046 | 4.84E-96 | green | TRABD |
| 2.60E-96 | -1.13527 | 0.011315 | 0.419174 | 6.08E-96 | green | TIMP1 |
| 1.11E-95 | 0.727282 | 0.318246 | 0.100415 | 2.57E-95 | green | TIFA |
| 1.86E-95 | 0.634824 | 0.834512 | 0.708435 | 4.29E-95 | green | SRSF3 |
| 3.12E-95 | -1.85266 | 0.011315 | 0.410417 | 7.17E-95 | green | CD3E |
| 5.64E-95 | 0.782208 | 0.704385 | 0.537894 | 1.29E-94 | green | SELENOF |
| 1.30E-94 | 0.866663 | 0.485149 | 0.224468 | 2.95E-94 | green | SPTBN1 |
| 4.00E-94 | -0.6171 | 0.154173 | 0.611411 | 9.02E-94 | green | SELENOW |
| 6.91E-94 | -0.96258 | 0.057992 | 0.456673 | 1.55E-93 | green | CD48 |
| 2.21E-93 | -2.15337 | 0.002829 | 0.397149 | 4.92E-93 | green | IL32 |
| 4.43E-91 | 0.627236 | 0.845827 | 0.5989 | 9.83E-91 | green | CXCR4 |
| 1.02E-90 | 0.685856 | 0.306931 | 0.096267 | 2.25E-90 | green | SUSD3 |
| 2.08E-90 | -0.89432 | 0.178218 | 0.582406 | 4.57E-90 | green | SELL |
| 3.42E-90 | -1.24619 | 0.330976 | 0.727102 | 7.45E-90 | green | FOS |
| 1.79E-89 | -0.81374 | 0.052334 | 0.451142 | 3.89E-89 | green | VSIR |
| 1.45E-88 | 0.605352 | 0.304102 | 0.095312 | 3.12E-88 | green | RIPOR1 |
| 1.65E-88 | 0.769232 | 0.367751 | 0.138342 | 3.55E-88 | green | ARHGEF7 |
| 7.64E-88 | 0.68227 | 0.785007 | 0.663232 | 1.63E-87 | green | ARGLU1 |
| 1.80E-87 | 0.77511 | 0.705799 | 0.553631 | 3.82E-87 | green | WASF2 |
| 4.69E-87 | -1.21055 | 0.352192 | 0.716567 | 9.90E-87 | green | ZFP36 |
| 1.19E-86 | -0.85651 | 0.421499 | 0.78205 | 2.49E-86 | green | IFITM2 |
| 1.53E-86 | 0.921618 | 0.473833 | 0.236617 | 3.21E-86 | green | AP1B1 |
| 2.11E-86 | -0.67017 | 0.026874 | 0.412557 | 4.38E-86 | green | ITM2A |
| 9.16E-84 | -0.99652 | 0.093352 | 0.476822 | 1.89E-83 | green | TRBC2 |
| 1.08E-83 | -1.01151 | 0.043847 | 0.414236 | 2.21E-83 | green | SLC2A3 |
| 5.81E-82 | -0.58493 | 0.742574 | 0.935043 | 1.19E-81 | green | RPS29 |
| 7.41E-82 | 0.669458 | 0.380481 | 0.153388 | 1.51E-81 | green | SYVN1 |
| 2.21E-81 | -0.57419 | 0.46959 | 0.827286 | 4.47E-81 | green | RPS20 |
| 8.77E-81 | -0.59527 | 0.131542 | 0.538158 | 1.77E-80 | green | CSTB |
| 1.72E-80 | -0.64207 | 0.326733 | 0.738098 | 3.45E-80 | green | LCP1 |
| 1.16E-79 | 0.628123 | 0.398868 | 0.160598 | 2.32E-79 | green | PAG1 |
| 1.73E-79 | 0.898241 | 0.635078 | 0.473102 | 3.43E-79 | green | VDAC1 |
| 1.67E-77 | 0.633234 | 0.727016 | 0.566373 | 3.29E-77 | green | YWHAQ |
| 4.76E-77 | -0.54974 | 0.673267 | 0.934615 | 9.35E-77 | green | TOMM7 |
| 2.47E-76 | -1.57375 | 0.039604 | 0.387535 | 4.84E-76 | green | LGALS1 |

| | | | | | | |
|---|---|---|---|---|---|---|
| 7.18E-76 | 0.783642 | 0.520509 | 0.29917 | 1.40E-75 | green | PECAM1 |
| 1.36E-75 | -0.66055 | 0.490806 | 0.828175 | 2.64E-75 | green | CORO1A |
| 7.53E-75 | 0.580482 | 0.277228 | 0.091987 | 1.45E-74 | green | SCMH1 |
| 7.54E-75 | -0.67999 | 0.036775 | 0.389577 | 1.45E-74 | green | C1orf162 |
| 4.24E-74 | 0.688999 | 0.388967 | 0.167643 | 8.09E-74 | green | PHACTR1 |
| 4.27E-74 | -0.75821 | 0.120226 | 0.490485 | 8.12E-74 | green | MCL1 |
| 5.05E-74 | -1.30368 | 0.015559 | 0.354547 | 9.55E-74 | green | S100A11 |
| 1.29E-73 | 0.819921 | 0.478076 | 0.264832 | 2.44E-73 | green | MTF2 |
| 1.71E-73 | 0.763094 | 0.637907 | 0.476263 | 3.20E-73 | green | TRAM1 |
| 1.73E-73 | 0.788185 | 0.560113 | 0.367584 | 3.22E-73 | green | ANKRD10 |
| 3.87E-73 | 0.716338 | 0.540311 | 0.300784 | 7.18E-73 | green | PDE4B |
| 4.54E-73 | -0.55257 | 0.046676 | 0.402384 | 8.38E-73 | green | NDFIP1 |
| 1.05E-72 | -0.67869 | 0.005658 | 0.335879 | 1.93E-72 | green | SELPLG |
| 3.70E-72 | -0.95374 | 0.032532 | 0.367189 | 6.78E-72 | green | NR4A2 |
| 5.27E-72 | 0.632227 | 0.376238 | 0.155363 | 9.60E-72 | green | TMEM131L |
| 7.15E-72 | -1.13382 | 0.001414 | 0.326826 | 1.30E-71 | green | CD3D |
| 1.83E-71 | 0.743008 | 0.45686 | 0.238856 | 3.31E-71 | green | GABPB1-AS1 |
| 2.76E-71 | 0.807802 | 0.465347 | 0.254132 | 4.96E-71 | green | BCL11A |
| 4.20E-71 | 0.58435 | 0.285714 | 0.098341 | 7.51E-71 | green | PIK3AP1 |
| 4.89E-71 | 0.653505 | 0.336634 | 0.133404 | 8.71E-71 | green | CIITA |
| 5.19E-71 | 0.565667 | 0.570014 | 0.287581 | 9.21E-71 | green | FCMR |
| 7.14E-71 | -0.75911 | 0.76662 | 0.910548 | 1.26E-70 | green | VIM |
| 7.31E-71 | 0.889603 | 0.492221 | 0.280799 | 1.28E-70 | green | REXO2 |
| 4.68E-70 | -0.51361 | 0.026874 | 0.356193 | 8.19E-70 | green | DNPH1 |
| 6.63E-70 | 0.584461 | 0.333805 | 0.128761 | 1.15E-69 | green | AFF3 |
| 8.72E-70 | -0.6 | 0.598303 | 0.838579 | 1.51E-69 | green | TAGLN2 |
| 1.24E-69 | 0.642069 | 0.336634 | 0.135445 | 2.13E-69 | green | ERO1B |
| 1.76E-69 | -0.78299 | 0.523338 | 0.742345 | 3.03E-69 | green | GSTP1 |
| 8.06E-69 | -0.76879 | 0.089109 | 0.436854 | 1.38E-68 | green | FOSB |
| 9.79E-69 | -0.70257 | 0.381895 | 0.716731 | 1.67E-68 | green | MT-ATP8 |
| 2.36E-68 | 0.922262 | 0.531825 | 0.331665 | 4.01E-68 | green | KLF10 |
| 3.02E-68 | -0.62342 | 0.093352 | 0.445611 | 5.11E-68 | green | APLP2 |
| 3.16E-68 | 0.614037 | 0.251768 | 0.081517 | 5.31E-68 | green | LMF1 |
| 4.18E-68 | -0.71016 | 0.195191 | 0.563047 | 7.01E-68 | green | LEPROTL1 |
| 8.24E-68 | -1.88447 | 0.015559 | 0.332521 | 1.37E-67 | green | TYROBP |
| 2.62E-67 | -0.56723 | 0.039604 | 0.370843 | 4.36E-67 | green | RNF130 |
| 3.50E-67 | 0.740802 | 0.613861 | 0.426615 | 5.79E-67 | green | ZEB2 |
| 4.07E-67 | 0.689151 | 0.446959 | 0.230131 | 6.70E-67 | green | A1BG |
| 1.48E-66 | -0.71407 | 0.031117 | 0.353394 | 2.43E-66 | green | PLEK |
| 4.49E-66 | 0.730214 | 0.572843 | 0.365312 | 7.34E-66 | green | PBXIP1 |
| 4.61E-66 | -0.62498 | 0.196605 | 0.559097 | 7.50E-66 | green | PRDX2 |

| | | | | | | |
|---|---|---|---|---|---|---|
| 4.83E-66 | 0.726803 | 0.700141 | 0.541417 | 7.83E-66 | green | EZR |
| 1.11E-65 | -0.57911 | 0.164074 | 0.532725 | 1.79E-65 | green | SLC9A3R1 |
| 2.81E-65 | 0.546504 | 0.756719 | 0.645848 | 4.51E-65 | green | FKBP1A |
| 3.24E-65 | 0.750137 | 0.540311 | 0.34332 | 5.19E-65 | green | JMJD1C |
| 2.14E-64 | 0.667572 | 0.312588 | 0.124218 | 3.41E-64 | green | MDM2 |
| 5.66E-64 | 0.753136 | 0.429986 | 0.228419 | 8.98E-64 | green | FHIT |
| 5.82E-64 | 0.501072 | 0.298444 | 0.104333 | 9.20E-64 | green | LILRB2 |
| 6.93E-64 | -0.56079 | 0.09901 | 0.437809 | 1.09E-63 | green | IDS |
| 1.12E-63 | -0.81376 | 0.01273 | 0.311714 | 1.76E-63 | green | ARL4C |
| 1.36E-63 | -0.52508 | 0.121641 | 0.4759 | 2.13E-63 | green | PTPN6 |
| 3.74E-63 | -0.51272 | 0.181047 | 0.54889 | 5.82E-63 | green | SYNGR2 |
| 4.30E-63 | 0.739665 | 0.497878 | 0.3021 | 6.67E-63 | green | SYK |
| 4.87E-63 | -0.51636 | 0.097595 | 0.441562 | 7.52E-63 | green | CTSC |
| 6.10E-63 | 0.560508 | 0.760962 | 0.681701 | 9.39E-63 | green | EPC1 |
| 7.48E-63 | -0.55771 | 0.397454 | 0.7219 | 1.15E-62 | green | LDHA |
| 1.69E-62 | 0.591317 | 0.277228 | 0.102884 | 2.59E-62 | green | CALHM6 |
| 4.44E-62 | -0.80591 | 0.022631 | 0.320603 | 6.75E-62 | green | SAMHD1 |
| 5.47E-62 | 0.579281 | 0.291372 | 0.112893 | 8.29E-62 | green | HMCES |
| 8.85E-62 | 0.533187 | 0.256011 | 0.085995 | 1.34E-61 | green | IGHD |
| 1.03E-61 | -0.88136 | 0.016973 | 0.3102 | 1.55E-61 | green | SPOCK2 |
| 1.95E-61 | -0.53104 | 0.007072 | 0.298644 | 2.92E-61 | green | CYTOR |
| 2.42E-61 | -0.52802 | 0.182461 | 0.530816 | 3.61E-61 | green | HSPB1 |
| 5.70E-61 | -0.70286 | 0.06082 | 0.376012 | 8.48E-61 | green | GRN |
| 6.46E-61 | 0.533403 | 0.267327 | 0.09439 | 9.57E-61 | green | AC116366.3 |
| 7.36E-61 | -0.52602 | 0.422914 | 0.724271 | 1.09E-60 | green | ANP32B |
| 1.43E-60 | -0.51945 | 0.055163 | 0.366695 | 2.11E-60 | green | TMC8 |
| 1.49E-60 | -1.54239 | 0.014144 | 0.303319 | 2.19E-60 | green | CST7 |
| 7.72E-60 | -2.27935 | 0.022631 | 0.311977 | 1.13E-59 | green | NKG7 |
| 1.13E-59 | 0.627572 | 0.391796 | 0.19678 | 1.64E-59 | green | ADA |
| 1.69E-59 | 0.704667 | 0.446959 | 0.259827 | 2.45E-59 | green | CDCA7L |
| 3.39E-59 | 0.771564 | 0.537482 | 0.372819 | 4.89E-59 | green | PFKL |
| 7.13E-59 | 0.616189 | 0.349364 | 0.155627 | 1.03E-58 | green | PPP3CC |
| 2.03E-58 | 0.723325 | 0.574257 | 0.431059 | 2.91E-58 | green | PSMA6 |
| 3.74E-58 | 0.765439 | 0.280057 | 0.112135 | 5.34E-58 | green | POLA2 |
| 5.16E-58 | 0.561146 | 0.338048 | 0.148054 | 7.34E-58 | green | CCM2 |
| 5.42E-58 | 0.573981 | 0.66761 | 0.515573 | 7.68E-58 | green | RBBP4 |
| 1.46E-57 | 0.70986 | 0.480905 | 0.304701 | 2.06E-57 | green | PLIN3 |
| 3.02E-57 | -0.66586 | 0.005658 | 0.281721 | 4.26E-57 | green | CSF3R |
| 5.68E-57 | -0.50806 | 0.141443 | 0.477975 | 7.96E-57 | green | CD63 |
| 7.09E-57 | 0.71988 | 0.462518 | 0.276157 | 9.91E-57 | green | SYNGR1 |
| 2.11E-56 | -0.73406 | 0.005658 | 0.277573 | 2.93E-56 | green | IL10RA |

| | | | | | | |
|---|---|---|---|---|---|---|
| 2.42E-56 | 0.667706 | 0.401697 | 0.209916 | 3.35E-56 | green | EIF1AY |
| 2.57E-56 | -0.54181 | 0.01273 | 0.289721 | 3.56E-56 | green | MGST1 |
| 1.17E-55 | -1.49125 | 0.113154 | 0.414828 | 1.62E-55 | green | CST3 |
| 3.36E-55 | -0.90558 | 0.393211 | 0.668532 | 4.62E-55 | green | TUBA1B |
| 8.59E-55 | 0.585635 | 0.718529 | 0.6203 | 1.18E-54 | green | GDI2 |
| 1.26E-54 | -0.5288 | 0.229137 | 0.562126 | 1.72E-54 | green | STK17B |
| 4.87E-54 | 0.611542 | 0.670438 | 0.556891 | 6.62E-54 | green | PNN |
| 8.00E-54 | -0.6814 | 0.009901 | 0.272733 | 1.08E-53 | green | LAT |
| 9.26E-54 | -0.74917 | 0.005658 | 0.266873 | 1.25E-53 | green | HOPX |
| 1.41E-53 | -0.79597 | 0.584158 | 0.791664 | 1.89E-53 | green | CD52 |
| 2.45E-53 | 0.668362 | 0.353607 | 0.17166 | 3.28E-53 | green | RELL1 |
| 3.46E-53 | -0.88002 | 0.001414 | 0.258379 | 4.63E-53 | green | GZMM |
| 3.83E-53 | -0.60553 | 0.007072 | 0.266083 | 5.10E-53 | green | MYO1F |
| 4.46E-53 | 0.550604 | 0.27157 | 0.106374 | 5.92E-53 | green | ARHGAP27 |
| 5.40E-53 | -0.79541 | 0.001414 | 0.257622 | 7.15E-53 | green | CD3G |
| 1.01E-52 | 0.531715 | 0.683168 | 0.599559 | 1.34E-52 | green | COX7A2L |
| 2.14E-52 | -0.65255 | 0.019802 | 0.284816 | 2.82E-52 | green | BLVRB |
| 4.87E-50 | -0.68613 | 0.933522 | 0.966024 | 6.38E-50 | green | RPS26 |
| 6.67E-50 | 0.537445 | 0.64215 | 0.547705 | 8.71E-50 | green | PPP2R1A |
| 7.22E-50 | 0.550654 | 0.264498 | 0.10779 | 9.40E-50 | green | MYO18A |
| 4.71E-49 | -0.7247 | 0.046676 | 0.306677 | 6.11E-49 | green | ANXA2 |
| 7.04E-49 | 0.688746 | 0.4314 | 0.259729 | 9.10E-49 | green | DCK |
| 1.47E-48 | -0.5581 | 0.646393 | 0.900112 | 1.89E-48 | green | MYL12A |
| 3.34E-48 | 0.515523 | 0.321075 | 0.145025 | 4.29E-48 | green | CSRNP1 |
| 4.68E-48 | 0.587227 | 0.550212 | 0.389511 | 6.00E-48 | green | CLK1 |
| 6.18E-48 | -1.2637 | 0.019802 | 0.263087 | 7.89E-48 | green | FCER1G |
| 6.84E-48 | 0.65504 | 0.329562 | 0.165536 | 8.70E-48 | green | ZMIZ1 |
| 1.05E-47 | -0.50292 | 0.056577 | 0.321624 | 1.34E-47 | green | AGTRAP |
| 1.86E-47 | 0.654641 | 0.611033 | 0.492955 | 2.35E-47 | green | LUC7L3 |
| 1.21E-46 | 0.573231 | 0.339463 | 0.162902 | 1.53E-46 | green | AC243960.1 |
| 2.05E-46 | 0.620887 | 0.442716 | 0.283005 | 2.57E-46 | green | FPGS |
| 2.50E-46 | 0.673528 | 0.44413 | 0.285244 | 3.13E-46 | green | SMARCA4 |
| 8.44E-46 | 0.580579 | 0.632249 | 0.516659 | 1.05E-45 | green | FOXP1 |
| 5.85E-45 | 0.721345 | 0.475248 | 0.315138 | 7.28E-45 | green | SOD2 |
| 1.15E-44 | 0.634388 | 0.404526 | 0.235728 | 1.43E-44 | green | MUC20-OT1 |
| 3.66E-44 | 0.511841 | 0.256011 | 0.108678 | 4.52E-44 | green | BCR |
| 4.04E-44 | -2.49847 | 0.02546 | 0.256206 | 4.98E-44 | green | LYZ |
| 8.42E-44 | -0.74618 | 0.082037 | 0.325245 | 1.04E-43 | green | LCK |
| 5.63E-43 | 0.693282 | 0.441301 | 0.283367 | 6.90E-43 | green | LAIR1 |
| 1.37E-42 | 0.534482 | 0.661952 | 0.524067 | 1.67E-42 | green | CCND3 |
| 2.66E-42 | 0.535845 | 0.487977 | 0.310463 | 3.24E-42 | green | SLC44A2 |

| | | | | | | |
|---|---|---|---|---|---|---|
| 2.95E-42 | -0.89388 | 0.107496 | 0.341246 | 3.58E-42 | green | PRSS57 |
| 3.33E-42 | 0.602992 | 0.586987 | 0.487753 | 4.03E-42 | green | TAF7 |
| 3.80E-42 | 0.516462 | 0.70297 | 0.651478 | 4.58E-42 | green | KMT2E |
| 6.32E-42 | 0.582062 | 0.5686 | 0.450616 | 7.60E-42 | green | PARP1 |
| 1.24E-41 | -0.54179 | 0.053748 | 0.287845 | 1.49E-41 | green | CRIP1 |
| 5.18E-41 | 0.650701 | 0.427157 | 0.282116 | 6.19E-41 | green | NGLY1 |
| 1.94E-40 | -0.63161 | 0.041018 | 0.264996 | 2.31E-40 | green | KLF2 |
| 3.11E-39 | -0.75379 | 0.196605 | 0.471028 | 3.69E-39 | green | CTSS |
| 3.47E-39 | -0.53364 | 0.275813 | 0.562422 | 4.12E-39 | green | DNAJB1 |
| 4.91E-39 | 0.535948 | 0.650636 | 0.554948 | 5.80E-39 | green | YPEL5 |
| 5.02E-39 | -0.71752 | 0.166902 | 0.429117 | 5.91E-39 | green | DUSP1 |
| 6.10E-39 | 0.623785 | 0.510608 | 0.379305 | 7.16E-39 | green | APBB1IP |
| 4.04E-38 | -1.20565 | 0.083451 | 0.29489 | 4.74E-38 | green | IL7R |
| 5.62E-38 | 0.605501 | 0.526167 | 0.425166 | 6.57E-38 | green | NDUFA5 |
| 7.53E-38 | 0.516532 | 0.27157 | 0.13156 | 8.76E-38 | green | NARFL |
| 7.75E-38 | 0.623433 | 0.461103 | 0.334134 | 9.00E-38 | green | CHD9 |
| 1.19E-37 | 0.59969 | 0.462518 | 0.323204 | 1.38E-37 | green | FLI1 |
| 1.49E-37 | 0.546421 | 0.322489 | 0.176829 | 1.72E-37 | green | NT5C2 |
| 3.28E-37 | -0.57578 | 0.048091 | 0.261013 | 3.78E-37 | green | FLT3LG |
| 3.69E-37 | 0.527924 | 0.27157 | 0.130836 | 4.23E-37 | green | CEP135 |
| 4.37E-37 | 0.589494 | 0.485149 | 0.354777 | 5.00E-37 | green | AKIRIN2 |
| 8.86E-37 | 0.611995 | 0.425743 | 0.289063 | 1.01E-36 | green | ATXN7L3B |
| 9.49E-37 | 0.614597 | 0.415842 | 0.287746 | 1.08E-36 | green | MYL6B |
| 1.75E-35 | 0.530346 | 0.550212 | 0.439257 | 1.98E-35 | green | RASGRP2 |
| 2.71E-35 | -0.98263 | 0.113154 | 0.335221 | 3.06E-35 | green | AREG |
| 3.46E-35 | 0.622685 | 0.432815 | 0.306907 | 3.91E-35 | green | SEC63 |
| 2.38E-34 | 0.623549 | 0.421499 | 0.292783 | 2.68E-34 | green | MSI2 |
| 3.99E-34 | 0.537761 | 0.292786 | 0.153849 | 4.47E-34 | green | MEF2A |
| 7.88E-34 | -0.55448 | 0.082037 | 0.300915 | 8.82E-34 | green | NAMPT |
| 1.43E-33 | 0.562235 | 0.439887 | 0.321788 | 1.60E-33 | green | UBE2E1 |
| 5.32E-33 | 0.549624 | 0.304102 | 0.169652 | 5.91E-33 | green | BCL7A |
| 6.66E-33 | -0.72242 | 0.356436 | 0.554059 | 7.38E-33 | green | AIF1 |
| 8.47E-33 | 0.579644 | 0.466761 | 0.352176 | 9.36E-33 | green | CALCOCO2 |
| 9.67E-33 | 0.563572 | 0.415842 | 0.277672 | 1.07E-32 | green | BAZ2A |
| 1.77E-32 | 0.567328 | 0.438472 | 0.31234 | 1.94E-32 | green | PPP4R3B |
| 2.04E-32 | 0.625692 | 0.39604 | 0.267696 | 2.24E-32 | green | ZMYM2 |
| 2.75E-32 | 0.558141 | 0.304102 | 0.171034 | 3.01E-32 | green | MIR181A1HG |
| 4.13E-32 | 0.553322 | 0.424328 | 0.28103 | 4.50E-32 | green | ARID5A |
| 4.76E-32 | 0.540851 | 0.513437 | 0.422829 | 5.18E-32 | green | ESD |
| 7.24E-32 | 0.586597 | 0.326733 | 0.194541 | 7.85E-32 | green | SPIDR |
| 8.88E-32 | 0.505288 | 0.308345 | 0.175249 | 9.61E-32 | green | BCL2L1 |

| | | | | | | |
|---|---|---|---|---|---|---|
| 4.87E-31 | 0.557118 | 0.347949 | 0.21897 | 5.25E-31 | green | STK26 |
| 1.06E-30 | 0.5454 | 0.465347 | 0.330217 | 1.14E-30 | green | SLC38A1 |
| 2.72E-30 | -0.64813 | 0.21075 | 0.426582 | 2.91E-30 | green | TNFAIP3 |
| 8.66E-29 | 0.531766 | 0.398868 | 0.276914 | 9.26E-29 | green | CARHSP1 |
| 1.67E-28 | 0.503479 | 0.526167 | 0.448311 | 1.78E-28 | green | ROCK1 |
| 4.65E-28 | -0.5677 | 0.164074 | 0.385494 | 4.95E-28 | green | LST1 |
| 4.69E-28 | 0.578033 | 0.554455 | 0.494074 | 4.98E-28 | green | WSB1 |
| 6.07E-28 | -0.62767 | 0.079208 | 0.255383 | 6.43E-28 | green | SPINK2 |
| 1.48E-26 | 0.528423 | 0.362093 | 0.246955 | 1.56E-26 | green | TIA1 |
| 1.61E-26 | 0.500046 | 0.49505 | 0.408013 | 1.70E-26 | green | RNASEH2B |
| 2.46E-26 | 0.50034 | 0.292786 | 0.174294 | 2.58E-26 | green | PCMTD2 |
| 3.33E-26 | 0.579268 | 0.360679 | 0.249325 | 3.49E-26 | green | STARD3 |
| 3.75E-26 | 0.509565 | 0.468175 | 0.374366 | 3.92E-26 | green | NIPBL |
| 1.49E-25 | 0.525139 | 0.257426 | 0.146671 | 1.55E-25 | green | GABPB1 |
| 1.18E-24 | 0.52029 | 0.278642 | 0.167578 | 1.22E-24 | green | EIPR1 |
| 2.51E-24 | 0.519715 | 0.349364 | 0.242609 | 2.61E-24 | green | BTK |
| 7.60E-23 | 0.534927 | 0.306931 | 0.200435 | 7.86E-23 | green | AFF1 |
| 8.71E-23 | 0.50496 | 0.448373 | 0.362744 | 8.98E-23 | green | CYBC1 |
| 1.42E-22 | 0.508379 | 0.400283 | 0.302924 | 1.46E-22 | green | TRA2A |
| 4.03E-22 | 0.546949 | 0.312588 | 0.212945 | 4.14E-22 | green | KDM5B |
| 6.44E-22 | -0.55519 | 0.461103 | 0.69092 | 6.59E-22 | green | TSPO |
| 7.14E-22 | 0.530312 | 0.422914 | 0.341114 | 7.28E-22 | green | ATP1B3 |
| 1.28E-21 | 0.51119 | 0.29703 | 0.186903 | 1.30E-21 | green | PMAIP1 |
| 2.69E-21 | 0.544107 | 0.33239 | 0.233753 | 2.73E-21 | green | SMC4 |
| 3.80E-21 | 0.506743 | 0.321075 | 0.219727 | 3.85E-21 | green | PKN2 |
| 2.70E-19 | 0.546713 | 0.415842 | 0.342892 | 2.72E-19 | green | MIS18BP1 |
| 1.88E-18 | 0.508026 | 0.318246 | 0.230757 | 1.89E-18 | green | WDFY2 |
| 4.59E-13 | 0.516942 | 0.315417 | 0.250774 | 4.62E-13 | green | MTIF3 |
| 6.43E-12 | 0.504933 | 0.335219 | 0.284915 | 6.45E-12 | green | ZNF326 |
| 0 | 1.488288 | 0.716923 | 0.055405 | 0 | lightyellow | HBD |
| 0 | 1.159764 | 0.857436 | 0.053483 | 0 | lightyellow | CNRIP1 |
| 0 | 1.123091 | 0.991795 | 0.325598 | 0 | lightyellow | AC084033.3 |
| 0 | 1.077883 | 0.782564 | 0.04188 | 0 | lightyellow | ITGA2B |
| 0 | 1.077299 | 0.936923 | 0.193711 | 0 | lightyellow | CYTL1 |
| 0 | 1.070829 | 0.738462 | 0.060348 | 0 | lightyellow | AL157895.1 |
| 0 | 1.042753 | 0.929231 | 0.183173 | 0 | lightyellow | SLC40A1 |
| 0 | 1.005151 | 0.855897 | 0.05108 | 0 | lightyellow | GATA1 |
| 0 | 1.003803 | 0.736923 | 0.081425 | 0 | lightyellow | FCER1A |
| 0 | 0.979803 | 0.997949 | 0.583845 | 0 | lightyellow | SERPINB1 |
| 0 | 0.909937 | 0.950769 | 0.402904 | 0 | lightyellow | TMEM14C |
| 0 | 0.9008 | 0.974359 | 0.421372 | 0 | lightyellow | CAT |

| | | | | | | |
|---|---|---|---|---|---|---|
| 0 | 0.895378 | 0.992308 | 0.455254 | 0 | lightyellow | IMPDH2 |
| 0 | 0.860733 | 0.944615 | 0.327692 | 0 | lightyellow | TUBA1C |
| 0 | 0.85959 | 0.918462 | 0.316639 | 0 | lightyellow | UROD |
| 0 | 0.825652 | 0.968718 | 0.322783 | 0 | lightyellow | RNF130 |
| 0 | 0.819596 | 0.47641 | 0.044351 | 0 | lightyellow | APOC1 |
| 0 | 0.815728 | 0.9 | 0.251313 | 0 | lightyellow | CYTOR |
| 0 | 0.801816 | 1 | 0.860218 | 0 | lightyellow | HSP90AB1 |
| 0 | 0.785651 | 0.84 | 0.182692 | 0 | lightyellow | PDLIM1 |
| 0 | 0.783918 | 0.977436 | 0.362157 | 0 | lightyellow | CDK6 |
| 0 | 0.77716 | 0.973333 | 0.452405 | 0 | lightyellow | GLUL |
| 0 | 0.732545 | 0.656923 | 0.191617 | 0 | lightyellow | MYC |
| 0 | 0.722505 | 0.94 | 0.294463 | 0 | lightyellow | MPP1 |
| 0 | 0.70882 | 0.728718 | 0.103704 | 0 | lightyellow | GATA2 |
| 0 | 0.707628 | 1 | 0.913254 | 0 | lightyellow | NPM1 |
| 0 | 0.706671 | 0.785128 | 0.095671 | 0 | lightyellow | TPM1 |
| 0 | 0.704481 | 0.848718 | 0.158457 | 0 | lightyellow | PKIG |
| 0 | 0.699048 | 0.922564 | 0.313069 | 0 | lightyellow | CD82 |
| 0 | 0.698464 | 0.789744 | 0.160276 | 0 | lightyellow | NET1 |
| 0 | 0.693096 | 0.989231 | 0.600529 | 0 | lightyellow | TUBB |
| 0 | 0.68716 | 0.941538 | 0.380866 | 0 | lightyellow | FBXO7 |
| 0 | 0.678216 | 0.978462 | 0.50012 | 0 | lightyellow | HSPD1 |
| 0 | 0.6772 | 0.789231 | 0.326182 | 0 | lightyellow | ATF7IP2 |
| 0 | 0.674324 | 0.97641 | 0.293056 | 0 | lightyellow | PRSS57 |
| 0 | 0.670534 | 0.958462 | 0.449899 | 0 | lightyellow | NAA38 |
| 0 | 0.6702 | 0.801538 | 0.165665 | 0 | lightyellow | NFE2 |
| 0 | 0.666564 | 0.703077 | 0.091655 | 0 | lightyellow | MINPP1 |
| 0 | 0.664908 | 0.767179 | 0.161512 | 0 | lightyellow | FSCN1 |
| 0 | 0.664468 | 0.873846 | 0.221929 | 0 | lightyellow | MEST |
| 0 | 0.660169 | 0.911795 | 0.329923 | 0 | lightyellow | CDK4 |
| 0 | 0.659656 | 0.910769 | 0.398888 | 0 | lightyellow | MPC2 |
| 0 | 0.655815 | 0.892821 | 0.235591 | 0 | lightyellow | CTNNBL1 |
| 0 | 0.639593 | 1 | 0.955271 | 0 | lightyellow | HNRNPA1 |
| 0 | 0.629456 | 0.998974 | 0.592942 | 0 | lightyellow | TALDO1 |
| 0 | 0.620955 | 0.808718 | 0.167142 | 0 | lightyellow | ALDH1A1 |
| 0 | 0.618689 | 0.631282 | 0.0702 | 0 | lightyellow | MPIG6B |
| 0 | 0.617025 | 0.987179 | 0.443651 | 0 | lightyellow | STMN1 |
| 0 | 0.612365 | 0.739487 | 0.22392 | 0 | lightyellow | H1F0 |
| 0 | 0.61099 | 0.995897 | 0.560228 | 0 | lightyellow | TXN |
| 0 | 0.599927 | 1 | 0.794617 | 0 | lightyellow | LDHB |
| 0 | 0.588749 | 0.96 | 0.509766 | 0 | lightyellow | DUT |
| 0 | 0.576009 | 0.628205 | 0.044248 | 0 | lightyellow | KLF1 |

| | | | | | | |
|---|---|---|---|---|---|---|
| 0 | 0.573877 | 0.967179 | 0.481274 | 0 | lightyellow | CCT2 |
| 0 | 0.562046 | 0.993333 | 0.657101 | 0 | lightyellow | SH3BGRL |
| 0 | 0.558136 | 0.81641 | 0.241976 | 0 | lightyellow | ISYNA1 |
| 0 | 0.555285 | 0.647179 | 0.13326 | 0 | lightyellow | PHTF1 |
| 0 | 0.552956 | 0.88 | 0.313137 | 0 | lightyellow | DNPH1 |
| 0 | 0.552163 | 0.841026 | 0.292232 | 0 | lightyellow | LIMS1 |
| 0 | 0.545656 | 0.747692 | 0.2474 | 0 | lightyellow | BLVRB |
| 0 | 0.541934 | 0.994872 | 0.722907 | 0 | lightyellow | TPI1 |
| 0 | 0.541254 | 0.858462 | 0.266314 | 0 | lightyellow | PAICS |
| 0 | 0.535095 | 0.994359 | 0.695582 | 0 | lightyellow | RAN |
| 0 | 0.535088 | 0.946154 | 0.383475 | 0 | lightyellow | RTN3 |
| 0 | 0.534384 | 0.875897 | 0.239333 | 0 | lightyellow | CD34 |
| 0 | 0.528494 | 0.747179 | 0.218393 | 0 | lightyellow | HDAC7 |
| 0 | 0.527763 | 1 | 0.995022 | 0 | lightyellow | RPLP0 |
| 0 | 0.526678 | 0.908718 | 0.341629 | 0 | lightyellow | AHCY |
| 0 | 0.52661 | 0.923077 | 0.375339 | 0 | lightyellow | AMD1 |
| 0 | 0.522148 | 0.501026 | 0.107823 | 0 | lightyellow | CPA3 |
| 0 | 0.520506 | 0.917949 | 0.338025 | 0 | lightyellow | BZW2 |
| 0 | 0.520039 | 0.594872 | 0.061069 | 0 | lightyellow | CSF2RB |
| 0 | 0.51989 | 0.919487 | 0.339158 | 0 | lightyellow | DDAH2 |
| 0 | 0.519747 | 0.989744 | 0.680409 | 0 | lightyellow | ATP5F1B |
| 0 | 0.519489 | 0.911795 | 0.347808 | 0 | lightyellow | HDAC2 |
| 0 | 0.509712 | 1 | 0.91473 | 0 | lightyellow | HMGB1 |
| 0 | 0.50953 | 0.997436 | 0.698672 | 0 | lightyellow | ANP32B |
| 0 | 0.509375 | 0.955897 | 0.354983 | 0 | lightyellow | SOX4 |
| 0 | 0.506828 | 0.905641 | 0.334283 | 0 | lightyellow | BEX3 |
| 0 | 0.506597 | 0.891282 | 0.336343 | 0 | lightyellow | FABP5 |
| 0 | 0.503894 | 1 | 0.964471 | 0 | lightyellow | EEF1B2 |
| 0 | 0.503163 | 0.780513 | 0.292541 | 0 | lightyellow | MCM7 |
| 0 | 0.500404 | 0.684615 | 0.126978 | 0 | lightyellow | HACD1 |
| 0 | -0.7418 | 0.995385 | 0.979609 | 0 | lightyellow | UBC |
| 0 | -0.9041 | 1 | 0.998661 | 0 | lightyellow | TMSB10 |
| 0 | -1.0101 | 0.889231 | 0.885037 | 0 | lightyellow | CALM1 |
| 0 | -1.1599 | 0.962051 | 0.945934 | 0 | lightyellow | HLA-E |
| 0 | -1.20394 | 0.715385 | 0.810648 | 0 | lightyellow | LSP1 |
| 0 | -1.22956 | 1 | 0.999657 | 0 | lightyellow | B2M |
| 0 | -1.24429 | 0.982051 | 0.986509 | 0 | lightyellow | HLA-A |
| 0 | -1.27754 | 0.990256 | 0.993924 | 0 | lightyellow | TMSB4X |
| 0 | -1.41478 | 0.575897 | 0.836875 | 0 | lightyellow | CORO1A |
| 0 | -1.41981 | 0.371282 | 0.676702 | 0 | lightyellow | HCST |
| 0 | -1.45055 | 0.998462 | 0.997322 | 0 | lightyellow | HLA-B |

| | | | | | | |
|---|---|---|---|---|---|---|
| 0 | -1.46941 | 0.416923 | 0.727781 | 0 | lightyellow | ZFP36 |
| 0 | -1.49458 | 0.99641 | 0.993031 | 0 | lightyellow | HLA-C |
| 0 | -1.54933 | 0.14359 | 0.635371 | 0 | lightyellow | CXCR4 |
| 0 | -1.87256 | 0.404615 | 0.812536 | 0 | lightyellow | CD52 |
| 0 | -1.98334 | 0.5 | 0.7894 | 0 | lightyellow | JUNB |
| 0 | -2.0345 | 0.205641 | 0.597508 | 0 | lightyellow | IFITM1 |
| 5.85E-306 | -1.09183 | 0.027179 | 0.475747 | 1.48E-305 | lightyellow | CD48 |
| 1.02E-298 | -1.23126 | 0.700513 | 0.826474 | 2.54E-298 | lightyellow | TSC22D3 |
| 1.07E-295 | 0.576575 | 0.975897 | 0.597645 | 2.66E-295 | lightyellow | DAD1 |
| 8.79E-295 | -0.88874 | 0.988205 | 0.96756 | 2.16E-294 | lightyellow | SH3BGRL3 |
| 9.42E-286 | -1.17979 | 0.636923 | 0.755415 | 2.29E-285 | lightyellow | PTPRC |
| 7.35E-282 | 0.542224 | 0.978974 | 0.590402 | 1.77E-281 | lightyellow | H2AFZ |
| 4.63E-278 | 0.535204 | 0.985641 | 0.640623 | 1.10E-277 | lightyellow | TUBA1B |
| 9.10E-270 | -1.31362 | 0.439487 | 0.657787 | 2.15E-269 | lightyellow | ITGB2 |
| 7.17E-266 | -1.89398 | 0.016923 | 0.427071 | 1.68E-265 | lightyellow | CD3E |
| 5.78E-258 | 0.504052 | 0.999487 | 0.885414 | 1.34E-257 | lightyellow | RPS2 |
| 1.58E-255 | -2.17639 | 0.013846 | 0.413237 | 3.62E-255 | lightyellow | IL32 |
| 9.55E-241 | -1.20411 | 0.341026 | 0.60197 | 2.17E-240 | lightyellow | EVL |
| 2.56E-236 | -1.1122 | 0.224615 | 0.5397 | 5.78E-236 | lightyellow | LITAF |
| 4.46E-234 | -1.29167 | 0.078462 | 0.444647 | 9.98E-234 | lightyellow | TNFAIP3 |
| 2.90E-229 | -1.64877 | 0.055385 | 0.419313 | 6.42E-229 | lightyellow | LTB |
| 8.33E-202 | -1.60434 | 0.446154 | 0.618791 | 1.83E-201 | lightyellow | S100A10 |
| 9.93E-202 | -0.69309 | 0.973846 | 0.923621 | 2.16E-201 | lightyellow | LAPTM5 |
| 7.51E-198 | -1.01563 | 0.007692 | 0.340599 | 1.62E-197 | lightyellow | LCK |
| 1.76E-197 | -1.09836 | 0.182564 | 0.487213 | 3.77E-197 | lightyellow | TRBC2 |
| 1.44E-196 | -1.15443 | 0.008718 | 0.340222 | 3.06E-196 | lightyellow | CD3D |
| 2.53E-196 | -0.68314 | 0.853333 | 0.378154 | 5.32E-196 | lightyellow | CST3 |
| 9.71E-191 | -0.91223 | 0.345641 | 0.588445 | 2.03E-190 | lightyellow | SELL |
| 6.28E-186 | -0.73943 | 0.925128 | 0.802513 | 1.30E-185 | lightyellow | CD99 |
| 4.87E-179 | -0.83993 | 0.040513 | 0.349936 | 1.00E-178 | lightyellow | ISG20 |
| 1.22E-178 | -0.5084 | 0.999487 | 0.999176 | 2.48E-178 | lightyellow | MALAT1 |
| 7.02E-177 | -1.00981 | 0.128718 | 0.424359 | 1.42E-176 | lightyellow | SLC2A3 |
| 2.94E-176 | -0.8455 | 0.017949 | 0.324122 | 5.90E-176 | lightyellow | ARL4C |
| 3.04E-175 | -0.89703 | 0.435385 | 0.591329 | 6.06E-175 | lightyellow | 9-Sep |
| 6.92E-175 | -0.92959 | 0.018462 | 0.322612 | 1.37E-174 | lightyellow | SPOCK2 |
| 1.60E-174 | -0.97415 | 0.326154 | 0.541451 | 3.14E-174 | lightyellow | BTG1 |
| 5.18E-173 | -1.46087 | 0.007692 | 0.308984 | 1.01E-172 | lightyellow | IL7R |
| 6.66E-172 | -0.86968 | 0.012821 | 0.312828 | 1.29E-171 | lightyellow | FCMR |
| 1.08E-170 | -0.87799 | 0.149231 | 0.438571 | 2.07E-170 | lightyellow | BTG2 |
| 1.33E-170 | -0.80347 | 0.151282 | 0.432907 | 2.52E-170 | lightyellow | CYTIP |
| 2.37E-168 | -0.80276 | 0.273333 | 0.512581 | 4.48E-168 | lightyellow | ANXA6 |

| | | | | | | |
|---|---|---|---|---|---|---|
| 7.76E-165 | -0.87202 | 0.182564 | 0.445745 | 1.46E-164 | lightyellow | HLA-F |
| 5.40E-159 | -0.65259 | 0.071795 | 0.359514 | 1.01E-158 | lightyellow | GLIPR1 |
| 7.40E-158 | -0.77021 | 0.008205 | 0.289005 | 1.37E-157 | lightyellow | IL10RA |
| 2.44E-155 | -0.85391 | 0.052821 | 0.331297 | 4.48E-155 | lightyellow | SAMHD1 |
| 2.25E-153 | -0.7902 | 0.017436 | 0.293296 | 4.09E-153 | lightyellow | ZAP70 |
| 5.12E-153 | -0.92347 | 0.65641 | 0.717037 | 9.26E-153 | lightyellow | IER2 |
| 1.31E-152 | -0.75024 | 0.520513 | 0.622739 | 2.36E-152 | lightyellow | PSMB9 |
| 5.24E-152 | -1.04268 | 0.686154 | 0.783083 | 9.34E-152 | lightyellow | JUN |
| 2.85E-145 | -0.90143 | 0.006154 | 0.269026 | 5.04E-145 | lightyellow | GZMM |
| 2.24E-143 | -0.81303 | 0.007692 | 0.268134 | 3.94E-143 | lightyellow | CD3G |
| 1.97E-141 | -0.61588 | 0.057949 | 0.322852 | 3.45E-141 | lightyellow | PDE4B |
| 2.15E-138 | -0.77468 | 0.020513 | 0.275926 | 3.72E-138 | lightyellow | KLF2 |
| 1.19E-137 | -2.31989 | 0.062564 | 0.32165 | 2.06E-137 | lightyellow | NKG7 |
| 9.82E-137 | -0.72042 | 0.144103 | 0.392949 | 1.68E-136 | lightyellow | P2RY8 |
| 3.02E-136 | -0.64971 | 0.989744 | 0.901823 | 5.14E-136 | lightyellow | CD37 |
| 6.05E-136 | -0.75678 | 0.421026 | 0.563489 | 1.02E-135 | lightyellow | STK17B |
| 1.68E-135 | -0.6596 | 0.845641 | 0.768219 | 2.82E-135 | lightyellow | GPSM3 |
| 1.99E-135 | -0.78101 | 0.236923 | 0.453606 | 3.32E-135 | lightyellow | 1-Sep |
| 1.84E-132 | -0.93235 | 0.217436 | 0.446672 | 3.05E-132 | lightyellow | GADD45B |
| 7.60E-132 | -0.87038 | 0.010769 | 0.255123 | 1.25E-131 | lightyellow | CD247 |
| 5.41E-130 | -0.71016 | 0.993846 | 0.900244 | 8.84E-130 | lightyellow | SARAF |
| 5.84E-130 | -0.60494 | 0.101026 | 0.347911 | 9.48E-130 | lightyellow | BIN1 |
| 3.59E-128 | -1.51992 | 0.161026 | 0.394254 | 5.79E-128 | lightyellow | LGALS1 |
| 4.75E-127 | -0.93109 | 0.257949 | 0.467612 | 7.61E-127 | lightyellow | AHNAK |
| 8.12E-127 | -0.88536 | 0.197949 | 0.42635 | 1.29E-126 | lightyellow | DDIT4 |
| 3.76E-123 | -0.81398 | 0.392821 | 0.536199 | 5.96E-123 | lightyellow | CCND3 |
| 6.15E-123 | -0.8192 | 0.083077 | 0.321856 | 9.68E-123 | lightyellow | ID2 |
| 1.78E-120 | -0.67509 | 0.66 | 0.652672 | 2.78E-120 | lightyellow | CD44 |
| 2.62E-119 | -1.10706 | 0.101026 | 0.345508 | 4.07E-119 | lightyellow | AREG |
| 1.99E-118 | -1.30386 | 0.138462 | 0.360784 | 3.07E-118 | lightyellow | S100A11 |
| 1.51E-115 | -0.79472 | 0.081026 | 0.305482 | 2.32E-115 | lightyellow | PIK3IP1 |
| 1.83E-113 | -0.62863 | 0.114359 | 0.335862 | 2.79E-113 | lightyellow | CYTH1 |
| 5.25E-113 | -1.02087 | 0.852821 | 0.789674 | 7.97E-113 | lightyellow | TXNIP |
| 2.36E-111 | -0.65057 | 0.151795 | 0.364388 | 3.57E-111 | lightyellow | IL16 |
| 3.70E-111 | -0.97408 | 0.374359 | 0.504651 | 5.56E-111 | lightyellow | FYB1 |
| 3.91E-110 | -0.56944 | 0.945641 | 0.904706 | 5.84E-110 | lightyellow | VIM |
| 1.18E-106 | -0.89336 | 0.614359 | 0.668841 | 1.75E-106 | lightyellow | KLF6 |
| 1.41E-106 | -0.61116 | 0.807179 | 0.72349 | 2.08E-106 | lightyellow | LCP1 |
| 3.19E-106 | -0.66687 | 0.065128 | 0.279256 | 4.68E-106 | lightyellow | LBH |
| 6.16E-103 | -0.62871 | 0.086667 | 0.29563 | 8.97E-103 | lightyellow | CRIP1 |
| 1.00E-102 | -1.45282 | 0.986667 | 0.914352 | 1.45E-102 | lightyellow | CD74 |

| | | | | | | |
|---|---|---|---|---|---|---|
| 8.34E-101 | -0.59508 | 0.061026 | 0.268546 | 1.20E-100 | lightyellow | PTPRE |
| 3.26E-100 | -0.80568 | 0.062051 | 0.266314 | 4.67E-100 | lightyellow | NCF1 |
| 3.41E-100 | -0.68493 | 0.062051 | 0.274244 | 4.86E-100 | lightyellow | HOPX |
| 9.64E-100 | -0.68961 | 0.194359 | 0.381037 | 1.37E-99 | lightyellow | RNF213 |
| 3.56E-94 | -0.56445 | 0.742564 | 0.775943 | 5.01E-94 | lightyellow | IFITM2 |
| 3.77E-90 | -0.55692 | 0.801026 | 0.705091 | 5.28E-90 | lightyellow | HCLS1 |
| 8.68E-89 | -0.53043 | 0.138974 | 0.327074 | 1.21E-88 | lightyellow | TRIM22 |
| 1.57E-88 | -0.6041 | 0.091282 | 0.279805 | 2.18E-88 | lightyellow | PRKCH |
| 1.82E-85 | -0.83078 | 0.346154 | 0.491779 | 2.52E-85 | lightyellow | CD69 |
| 2.75E-84 | -0.54165 | 0.842051 | 0.734063 | 3.77E-84 | lightyellow | ITM2B |
| 3.44E-84 | -0.67657 | 0.087179 | 0.267481 | 4.69E-84 | lightyellow | FLT3LG |
| 2.18E-81 | -0.63954 | 0.610769 | 0.614809 | 2.96E-81 | lightyellow | MYH9 |
| 4.79E-81 | -0.63033 | 0.163077 | 0.333562 | 6.46E-81 | lightyellow | ADGRE5 |
| 7.00E-78 | -0.56611 | 0.976923 | 0.888813 | 9.39E-78 | lightyellow | MYL12A |
| 1.19E-77 | -0.68736 | 0.098462 | 0.271257 | 1.59E-77 | lightyellow | ALOX5AP |
| 1.47E-77 | -1.89493 | 0.172821 | 0.335519 | 1.95E-77 | lightyellow | TYROBP |
| 2.77E-77 | -0.71007 | 0.324615 | 0.435103 | 3.66E-77 | lightyellow | TBC1D10C |
| 1.97E-76 | -0.70498 | 0.337436 | 0.449075 | 2.59E-76 | lightyellow | VSIR |
| 3.47E-75 | -0.56668 | 0.142564 | 0.306306 | 4.54E-75 | lightyellow | AKNA |
| 9.42E-72 | -0.73707 | 0.504615 | 0.533658 | 1.23E-71 | lightyellow | RARRES3 |
| 2.38E-71 | -0.57745 | 0.127692 | 0.287632 | 3.08E-71 | lightyellow | MGAT4A |
| 2.78E-70 | -0.95225 | 0.240513 | 0.376781 | 3.59E-70 | lightyellow | HLA-DQB1 |
| 5.28E-70 | -0.59631 | 0.289744 | 0.411314 | 6.77E-70 | lightyellow | ADD3 |
| 1.16E-68 | -0.51351 | 0.131795 | 0.287117 | 1.47E-68 | lightyellow | ANKRD44 |
| 1.05E-67 | -0.54608 | 0.697436 | 0.624833 | 1.33E-67 | lightyellow | RNASET2 |
| 2.97E-66 | -0.72126 | 0.453846 | 0.498747 | 3.75E-66 | lightyellow | AES |
| 4.39E-66 | -0.5851 | 0.694872 | 0.636847 | 5.51E-66 | lightyellow | CNN2 |
| 3.01E-65 | -0.56126 | 0.198974 | 0.342419 | 3.77E-65 | lightyellow | RCSD1 |
| 1.10E-63 | -0.54894 | 0.182051 | 0.325598 | 1.37E-63 | lightyellow | RNF125 |
| 1.59E-62 | -0.67178 | 0.546154 | 0.544952 | 1.97E-62 | lightyellow | EZR |
| 3.76E-61 | -0.6304 | 0.558462 | 0.551577 | 4.64E-61 | lightyellow | ARHGEF1 |
| 5.47E-61 | -0.53761 | 0.134359 | 0.284886 | 6.71E-61 | lightyellow | CSF3R |
| 3.06E-57 | -0.63469 | 0.301026 | 0.396794 | 3.74E-57 | lightyellow | TRAF3IP3 |
| 2.24E-55 | -0.53305 | 0.223077 | 0.340668 | 2.72E-55 | lightyellow | SLC38A1 |
| 4.69E-53 | -0.61594 | 0.142564 | 0.271189 | 5.67E-53 | lightyellow | TUBA4A |
| 9.70E-51 | -0.66318 | 0.375385 | 0.449212 | 1.17E-50 | lightyellow | IRF1 |
| 6.13E-50 | -0.74251 | 0.194872 | 0.307851 | 7.34E-50 | lightyellow | ANXA2 |
| 3.66E-49 | -0.54118 | 0.573333 | 0.56167 | 4.36E-49 | lightyellow | ZFP36L2 |
| 3.79E-49 | -0.56521 | 0.549744 | 0.529745 | 4.49E-49 | lightyellow | CD53 |
| 1.07E-43 | -0.60438 | 0.405641 | 0.442003 | 1.26E-43 | lightyellow | PIK3R1 |
| 3.96E-43 | -0.86748 | 0.808205 | 0.64234 | 4.66E-43 | lightyellow | HLA-DPB1 |

| | | | | | | |
|---|---|---|---|---|---|---|
| 1.03E-40 | -1.45047 | 0.210769 | 0.302496 | 1.21E-40 | lightyellow | CST7 |
| 3.85E-40 | -0.55443 | 0.554359 | 0.522605 | 4.48E-40 | lightyellow | CYBA |
| 7.46E-39 | -0.71664 | 0.287179 | 0.364423 | 8.64E-39 | lightyellow | NR4A2 |
| 1.68E-35 | -0.54441 | 0.329744 | 0.385019 | 1.93E-35 | lightyellow | C1orf162 |
| 8.86E-35 | -0.69232 | 0.323077 | 0.376403 | 1.02E-34 | lightyellow | MAP3K8 |
| 9.01E-35 | -0.51105 | 0.349744 | 0.392674 | 1.03E-34 | lightyellow | CYFIP2 |
| 1.70E-34 | -0.63371 | 0.330769 | 0.378463 | 1.94E-34 | lightyellow | HERPUD1 |
| 9.38E-31 | -0.65818 | 0.979487 | 0.817377 | 1.06E-30 | lightyellow | SRGN |
| 1.45E-30 | -0.50244 | 0.413846 | 0.425286 | 1.64E-30 | lightyellow | VAMP2 |
| 2.17E-28 | -0.75016 | 0.64 | 0.550273 | 2.44E-28 | lightyellow | DNAJB1 |
| 8.90E-28 | -0.54846 | 0.644615 | 0.551268 | 9.95E-28 | lightyellow | YPEL5 |
| 2.35E-27 | -0.57757 | 0.501026 | 0.48275 | 2.61E-27 | lightyellow | IFITM3 |
| 9.88E-25 | -0.52143 | 0.554872 | 0.495417 | 1.09E-24 | lightyellow | ARRB2 |
| 9.39E-24 | -0.81998 | 0.858462 | 0.636985 | 1.04E-23 | lightyellow | HLA-DPA1 |
| 1.86E-23 | -0.66559 | 0.655385 | 0.547939 | 2.04E-23 | lightyellow | LEPROTL1 |
| 1.04E-22 | -0.51257 | 0.295897 | 0.331262 | 1.14E-22 | lightyellow | EML4 |
| 1.85E-22 | -0.7405 | 0.406667 | 0.424256 | 2.02E-22 | lightyellow | DUSP1 |
| 2.32E-22 | -0.73713 | 0.262564 | 0.310425 | 2.51E-22 | lightyellow | HLA-DQA1 |
| 8.22E-20 | -1.11465 | 0.451282 | 0.407126 | 8.88E-20 | lightyellow | CD7 |
| 2.32E-17 | -0.53968 | 0.449231 | 0.414061 | 2.50E-17 | lightyellow | RIPOR2 |
| 3.47E-17 | -1.04767 | 0.762564 | 0.613745 | 3.71E-17 | lightyellow | SAT1 |
| 5.55E-16 | -0.99301 | 0.862051 | 0.62624 | 5.92E-16 | lightyellow | HLA-DRB1 |
| 1.21E-14 | -0.67488 | 0.973333 | 0.813772 | 1.29E-14 | lightyellow | S100A6 |
| 3.73E-14 | -0.86852 | 0.544615 | 0.459442 | 3.95E-14 | lightyellow | CTSS |
| 5.09E-12 | -1.16147 | 0.240513 | 0.258693 | 5.36E-12 | lightyellow | FCER1G |
| 2.54E-10 | -0.50882 | 0.485128 | 0.41612 | 2.66E-10 | lightyellow | IQGAP1 |
| 9.60E-10 | -0.51866 | 0.571282 | 0.476091 | 1.00E-09 | lightyellow | MCL1 |
| 4.45E-08 | -0.56043 | 0.459487 | 0.408156 | 4.63E-08 | lightyellow | NFKBIA |
| 3.12E-07 | -0.57287 | 0.692821 | 0.527548 | 3.23E-07 | lightyellow | FLNA |
| 3.30E-07 | -0.54045 | 0.550769 | 0.369057 | 3.41E-07 | lightyellow | LST1 |
| 5.96E-07 | -0.94651 | 0.363077 | 0.248395 | 6.13E-07 | lightyellow | IGLL1 |
| 1.84E-06 | -0.81089 | 0.812308 | 0.566441 | 1.89E-06 | lightyellow | HLA-DRB5 |
| 0.001249 | -2.41803 | 0.272308 | 0.249528 | 0.001274 | lightyellow | LYZ |
| 0.030699 | -0.51568 | 0.384615 | 0.29 | 0.031188 | lightyellow | NAMPT |
| 0 | 1.344948 | 0.964261 | 0.179296 | 0 | midnightblue | SPINK2 |
| 0 | 1.301674 | 0.761738 | 0.05477 | 0 | midnightblue | CRHBP |
| 0 | 1.128936 | 0.485284 | 0.024126 | 0 | midnightblue | AC011139.1 |
| 0 | 1.126654 | 0.813595 | 0.199915 | 0 | midnightblue | H1F0 |
| 0 | 1.101975 | 0.870007 | 0.223297 | 0 | midnightblue | FAM30A |
| 0 | 1.053817 | 0.921514 | 0.214334 | 0 | midnightblue | CD34 |
| 0 | 1.020416 | 0.892081 | 0.196408 | 0 | midnightblue | SMIM24 |

| | | | | | | |
|---|---|---|---|---|---|---|
| 0 | 1.017594 | 0.937281 | 0.242073 | 0 | midnightblue | EGFL7 |
| 0 | 1.013516 | 0.804835 | 0.193396 | 0 | midnightblue | TSC22D1 |
| 0 | 0.982326 | 0.71794 | 0.290963 | 0 | midnightblue | AREG |
| 0 | 0.956578 | 0.975123 | 0.537889 | 0 | midnightblue | PRDX1 |
| 0 | 0.941173 | 0.998949 | 0.760903 | 0 | midnightblue | ZFAS1 |
| 0 | 0.854932 | 0.82761 | 0.270769 | 0 | midnightblue | GNA15 |
| 0 | 0.835832 | 0.989488 | 0.571439 | 0 | midnightblue | SERPINB1 |
| 0 | 0.775537 | 0.999299 | 0.855812 | 0 | midnightblue | HSP90AB1 |
| 0 | 0.761121 | 0.664331 | 0.109505 | 0 | midnightblue | MDK |
| 0 | 0.754147 | 0.934478 | 0.488291 | 0 | midnightblue | CD164 |
| 0 | 0.750552 | 0.791521 | 0.241648 | 0 | midnightblue | ANKRD28 |
| 0 | 0.738369 | 0.797828 | 0.183973 | 0 | midnightblue | CYTL1 |
| 0 | 0.734197 | 0.788367 | 0.1929 | 0 | midnightblue | IL18 |
| 0 | 0.723275 | 0.970217 | 0.504198 | 0 | midnightblue | LAPTM4A |
| 0 | 0.706198 | 0.431324 | 0.059517 | 0 | midnightblue | BEX1 |
| 0 | 0.688984 | 0.949194 | 0.442413 | 0 | midnightblue | IMPDH2 |
| 0 | 0.688943 | 0.886826 | 0.323874 | 0 | midnightblue | DDAH2 |
| 0 | 0.684723 | 0.914506 | 0.420555 | 0 | midnightblue | SYPL1 |
| 0 | 0.681761 | 0.873861 | 0.352925 | 0 | midnightblue | CDK6 |
| 0 | 0.681341 | 1 | 0.906614 | 0 | midnightblue | HINT1 |
| 0 | 0.677963 | 1 | 0.910476 | 0 | midnightblue | NPM1 |
| 0 | 0.670005 | 0.940785 | 0.274772 | 0 | midnightblue | PRSS57 |
| 0 | 0.652227 | 0.322355 | 0.01495 | 0 | midnightblue | AVP |
| 0 | 0.645439 | 0.99124 | 0.764481 | 0 | midnightblue | EIF3E |
| 0 | 0.63581 | 0.664681 | 0.129592 | 0 | midnightblue | NPR3 |
| 0 | 0.621806 | 0.942537 | 0.511425 | 0 | midnightblue | BST2 |
| 0 | 0.61828 | 0.977575 | 0.631027 | 0 | midnightblue | SNHG8 |
| 0 | 0.607469 | 0.937631 | 0.611932 | 0 | midnightblue | TUBA1A |
| 0 | 0.602518 | 0.867554 | 0.308216 | 0 | midnightblue | ITM2C |
| 0 | 0.594794 | 1 | 0.997626 | 0 | midnightblue | RPS4X |
| 0 | 0.587836 | 0.840925 | 0.319517 | 0 | midnightblue | AC084033.3 |
| 0 | 0.586408 | 0.953399 | 0.537818 | 0 | midnightblue | APEX1 |
| 0 | 0.579606 | 0.659425 | 0.161689 | 0 | midnightblue | ALDH1A1 |
| 0 | 0.576221 | 0.584793 | 0.103624 | 0 | midnightblue | TFPI |
| 0 | 0.564625 | 0.867204 | 0.319871 | 0 | midnightblue | BEX3 |
| 0 | 0.55425 | 0.942887 | 0.535834 | 0 | midnightblue | SELL |
| 0 | 0.553147 | 1 | 0.991887 | 0 | midnightblue | RACK1 |
| 0 | 0.546628 | 0.759986 | 0.248769 | 0 | midnightblue | MSI2 |
| 0 | 0.539638 | 0.743518 | 0.225316 | 0 | midnightblue | BEX2 |
| 0 | 0.537711 | 0.603714 | 0.132533 | 0 | midnightblue | MSRB3 |
| 0 | 0.537643 | 1 | 0.99915 | 0 | midnightblue | RPS24 |

| | | | | | | |
|---|---|---|---|---|---|---|
| 0 | 0.536571 | 0.756482 | 0.349736 | 0 | midnightblue | TPM4 |
| 0 | 0.53594 | 0.948493 | 0.452723 | 0 | midnightblue | H2AFY |
| 0 | 0.533052 | 0.544499 | 0.084423 | 0 | midnightblue | PROM1 |
| 0 | 0.532558 | 0.480378 | 0.139193 | 0 | midnightblue | AC020916.1 |
| 0 | 0.52948 | 0.707428 | 0.217877 | 0 | midnightblue | MEST |
| 0 | 0.528164 | 0.99965 | 0.963333 | 0 | midnightblue | EEF2 |
| 0 | 0.526563 | 0.994394 | 0.882276 | 0 | midnightblue | RPS2 |
| 0 | 0.525124 | 1 | 0.969887 | 0 | midnightblue | BTF3 |
| 0 | 0.517712 | 0.597057 | 0.141035 | 0 | midnightblue | SERPING1 |
| 0 | 0.516326 | 0.655571 | 0.188472 | 0 | midnightblue | RAB13 |
| 0 | 0.5099 | 0.974422 | 0.62837 | 0 | midnightblue | RSL1D1 |
| 0 | 0.505095 | 0.802383 | 0.331137 | 0 | midnightblue | BZW2 |
| 0 | 0.504164 | 0.680799 | 0.160945 | 0 | midnightblue | JUP |
| 0 | 0.500967 | 0.882621 | 0.407943 | 0 | midnightblue | SPINT2 |
| 0 | -0.57109 | 0.946041 | 0.921671 | 0 | midnightblue | TMA7 |
| 0 | -0.59789 | 0.992292 | 0.976264 | 0 | midnightblue | CFL1 |
| 0 | -0.62129 | 0.970568 | 0.947072 | 0 | midnightblue | ATP5F1E |
| 0 | -0.62395 | 0.979678 | 0.951075 | 0 | midnightblue | MYL6 |
| 0 | -0.66867 | 1 | 0.99961 | 0 | midnightblue | ACTB |
| 0 | -0.73603 | 0.988437 | 0.97885 | 0 | midnightblue | PFN1 |
| 0 | -0.86206 | 0.878416 | 0.895951 | 0 | midnightblue | MYL12A |
| 0 | -0.96371 | 0.983181 | 0.986538 | 0 | midnightblue | HLA-A |
| 0 | -1.02127 | 0.103364 | 0.48241 | 0 | midnightblue | CD48 |
| 0 | -1.06583 | 0.688858 | 0.833812 | 0 | midnightblue | CORO1A |
| 0 | -1.09373 | 1 | 0.999646 | 0 | midnightblue | B2M |
| 0 | -1.16087 | 0.47267 | 0.676232 | 0 | midnightblue | HCST |
| 0 | -1.18019 | 0.998248 | 0.997308 | 0 | midnightblue | HLA-B |
| 0 | -1.21347 | 0.426419 | 0.6661 | 0 | midnightblue | ITGB2 |
| 0 | -1.21987 | 0.723196 | 0.90169 | 0 | midnightblue | CALM1 |
| 0 | -1.23139 | 0.994043 | 0.993163 | 0 | midnightblue | HLA-C |
| 0 | -1.28051 | 0.083392 | 0.538244 | 0 | midnightblue | FYB1 |
| 0 | -1.28253 | 0.660827 | 0.807064 | 0 | midnightblue | TXNIP |
| 0 | -1.31213 | 0.99965 | 0.998654 | 0 | midnightblue | TMSB10 |
| 0 | -1.32418 | 0.992992 | 0.993765 | 0 | midnightblue | TMSB4X |
| 0 | -1.36526 | 0.29082 | 0.636235 | 0 | midnightblue | CXCR4 |
| 0 | -1.45279 | 0.614576 | 0.787083 | 0 | midnightblue | JUNB |
| 0 | -1.63464 | 0.228802 | 0.646296 | 0 | midnightblue | S100A10 |
| 0 | -1.85378 | 0.748073 | 0.831438 | 0 | midnightblue | S100A6 |
| 0 | -1.92079 | 0.022074 | 0.439685 | 0 | midnightblue | CD3E |
| 0 | -2.21172 | 0.006657 | 0.426755 | 0 | midnightblue | IL32 |
| 5.09E-300 | -1.17905 | 0.005606 | 0.351153 | 1.05E-299 | midnightblue | CD3D |

| | | | | | | |
|---|---|---|---|---|---|---|
| 9.86E-296 | -0.59325 | 0.969166 | 0.917349 | 2.02E-295 | midnightblue | OAZ1 |
| 2.49E-291 | -1.03033 | 0.012614 | 0.350763 | 5.03E-291 | midnightblue | LCK |
| 1.07E-288 | -1.59731 | 0.101962 | 0.426259 | 2.13E-288 | midnightblue | LTB |
| 7.46E-288 | -0.84935 | 0.761037 | 0.809084 | 1.48E-287 | midnightblue | LSP1 |
| 2.08E-279 | -0.77625 | 0.988087 | 0.966911 | 4.07E-279 | midnightblue | SH3BGRL3 |
| 1.50E-270 | -0.88542 | 0.022074 | 0.343324 | 2.90E-270 | midnightblue | SAMHD1 |
| 3.82E-269 | -0.89858 | 0.664681 | 0.738406 | 7.33E-269 | midnightblue | EMP3 |
| 4.72E-268 | -1.38731 | 0.12754 | 0.438445 | 8.94E-268 | midnightblue | CD7 |
| 9.95E-262 | -1.48706 | 0.006657 | 0.318737 | 1.87E-261 | midnightblue | IL7R |
| 1.06E-252 | -0.55906 | 0.988087 | 0.979842 | 1.97E-252 | midnightblue | UBC |
| 5.55E-248 | -0.82611 | 0.392782 | 0.600631 | 1.02E-247 | midnightblue | 9-Sep |
| 1.58E-238 | -0.83593 | 0.037842 | 0.331916 | 2.88E-238 | midnightblue | ARL4C |
| 6.28E-236 | -0.90639 | 0.3185 | 0.54912 | 1.13E-235 | midnightblue | BTG1 |
| 1.09E-233 | -0.78239 | 0.010862 | 0.297729 | 1.94E-233 | midnightblue | IL10RA |
| 1.15E-232 | -0.91523 | 0.321654 | 0.559783 | 2.03E-232 | midnightblue | FLNA |
| 2.45E-226 | -1.45891 | 0.896636 | 0.803132 | 4.30E-226 | midnightblue | S100A4 |
| 7.51E-222 | -0.92134 | 0.003504 | 0.277713 | 1.30E-221 | midnightblue | GZMM |
| 1.77E-221 | -0.83465 | 0.003153 | 0.276933 | 3.04E-221 | midnightblue | CD3G |
| 2.23E-218 | -0.71848 | 0.017519 | 0.291955 | 3.80E-218 | midnightblue | LAT |
| 9.45E-217 | 0.832142 | 0.939033 | 0.695752 | 1.60E-216 | midnightblue | FOS |
| 1.07E-205 | -0.8898 | 0.004905 | 0.263542 | 1.79E-205 | midnightblue | CD247 |
| 5.37E-202 | 0.523793 | 0.383322 | 0.157296 | 8.91E-202 | midnightblue | HSPA1B |
| 2.88E-201 | -0.77896 | 0.024527 | 0.284267 | 4.74E-201 | midnightblue | ALOX5AP |
| 2.84E-200 | -0.78207 | 0.289068 | 0.516846 | 4.63E-200 | midnightblue | AES |
| 2.68E-199 | -0.71258 | 0.187106 | 0.444043 | 4.33E-199 | midnightblue | IQGAP1 |
| 1.66E-195 | -0.97908 | 0.462509 | 0.598044 | 2.66E-195 | midnightblue | EVL |
| 1.01E-194 | -0.92939 | 0.351086 | 0.537004 | 1.61E-194 | midnightblue | LITAF |
| 1.56E-194 | -0.89218 | 0.074982 | 0.326638 | 2.46E-194 | midnightblue | SPOCK2 |
| 4.33E-194 | -0.83124 | 0.064471 | 0.317214 | 6.75E-194 | midnightblue | FCMR |
| 2.91E-189 | -1.56649 | 0.066924 | 0.319977 | 4.50E-189 | midnightblue | CST7 |
| 8.52E-188 | -0.78858 | 0.225648 | 0.449818 | 1.31E-187 | midnightblue | HLA-F |
| 2.74E-184 | -0.8849 | 0.237211 | 0.45832 | 4.16E-184 | midnightblue | ZFP36L1 |
| 4.05E-183 | -0.76732 | 0.055361 | 0.298296 | 6.10E-183 | midnightblue | ZAP70 |
| 3.65E-180 | -0.81863 | 0.077435 | 0.323343 | 5.45E-180 | midnightblue | ANXA2 |
| 2.07E-172 | -0.75976 | 0.123336 | 0.351472 | 3.07E-172 | midnightblue | ISG20 |
| 2.60E-163 | -0.88417 | 0.291871 | 0.493428 | 3.83E-163 | midnightblue | MT2A |
| 1.95E-161 | -1.42017 | 0.18185 | 0.399617 | 2.85E-161 | midnightblue | LGALS1 |
| 3.10E-156 | -0.52478 | 0.11843 | 0.347079 | 4.49E-156 | midnightblue | REEP5 |
| 9.99E-156 | -0.55958 | 0.677996 | 0.714458 | 1.44E-155 | midnightblue | HCLS1 |
| 3.37E-154 | -0.62143 | 0.083742 | 0.302618 | 4.80E-154 | midnightblue | CRIP1 |
| 3.24E-151 | -0.65742 | 0.070778 | 0.285542 | 4.59E-151 | midnightblue | LBH |

| | | | | | | |
|---|---|---|---|---|---|---|
| 1.36E-144 | -0.65872 | 0.071128 | 0.282531 | 1.91E-144 | midnightblue | TUBA4A |
| 6.27E-144 | -1.52728 | 0.522775 | 0.577993 | 8.74E-144 | midnightblue | IFITM1 |
| 1.13E-141 | -0.78677 | 0.755781 | 0.747192 | 1.56E-141 | midnightblue | PTPRC |
| 1.80E-138 | -0.55791 | 0.794674 | 0.770893 | 2.48E-138 | midnightblue | GPSM3 |
| 2.95E-137 | -2.28429 | 0.122285 | 0.32391 | 4.02E-137 | midnightblue | NKG7 |
| 1.66E-136 | -0.73035 | 0.421163 | 0.537925 | 2.25E-136 | midnightblue | CCND3 |
| 2.48E-136 | -0.74809 | 0.113875 | 0.309349 | 3.34E-136 | midnightblue | PIK3IP1 |
| 8.28E-134 | -0.54978 | 0.71829 | 0.729798 | 1.10E-133 | midnightblue | LCP1 |
| 2.06E-133 | -1.29923 | 0.078136 | 0.275693 | 2.73E-133 | midnightblue | FCER1G |
| 3.76E-131 | -0.5182 | 0.902943 | 0.832501 | 4.94E-131 | midnightblue | LIMD2 |
| 6.02E-129 | -0.57476 | 0.123686 | 0.323059 | 7.85E-129 | midnightblue | GNG2 |
| 1.78E-127 | -0.68366 | 0.205326 | 0.392674 | 2.31E-127 | midnightblue | HERPUD1 |
| 1.24E-126 | -0.77706 | 0.081289 | 0.270911 | 1.60E-126 | midnightblue | NCF1 |
| 2.66E-120 | -0.55156 | 0.071128 | 0.256669 | 3.40E-120 | midnightblue | CDC25B |
| 5.33E-118 | -0.59801 | 0.688858 | 0.701243 | 6.77E-118 | midnightblue | UCP2 |
| 2.74E-117 | -0.65596 | 0.093203 | 0.272647 | 3.45E-117 | midnightblue | FLT3LG |
| 2.37E-113 | -0.61994 | 0.094954 | 0.276579 | 2.97E-113 | midnightblue | KLF2 |
| 6.30E-110 | -0.76828 | 0.269096 | 0.419634 | 7.84E-110 | midnightblue | SLC2A3 |
| 4.71E-109 | -0.55524 | 0.405746 | 0.524179 | 5.82E-109 | midnightblue | PAXX |
| 1.01E-104 | -0.52936 | 0.983181 | 0.943281 | 1.24E-104 | midnightblue | HLA-E |
| 1.49E-98 | -0.57588 | 0.564471 | 0.61962 | 1.81E-98 | midnightblue | MYH9 |
| 1.36E-95 | -0.54284 | 0.18185 | 0.343182 | 1.64E-95 | midnightblue | SELPLG |
| 1.74E-95 | -0.54467 | 0.184303 | 0.343678 | 2.09E-95 | midnightblue | EML4 |
| 2.36E-95 | -0.5373 | 0.139103 | 0.30198 | 2.81E-95 | midnightblue | PPP2R5C |
| 3.29E-90 | -0.58222 | 0.244919 | 0.381904 | 3.91E-90 | midnightblue | RNF213 |
| 1.32E-89 | -0.75908 | 0.374562 | 0.477557 | 1.55E-89 | midnightblue | TRBC2 |
| 1.81E-89 | -0.57911 | 0.470217 | 0.530095 | 2.13E-89 | midnightblue | CYBA |
| 1.34E-85 | -0.87453 | 0.322355 | 0.431714 | 1.56E-85 | midnightblue | TNFAIP3 |
| 7.92E-81 | -0.60627 | 0.340575 | 0.437028 | 9.17E-81 | midnightblue | TBC1D10C |
| 3.41E-78 | -2.51385 | 0.123686 | 0.263825 | 3.92E-78 | midnightblue | LYZ |
| 3.68E-78 | -0.51323 | 0.204275 | 0.333865 | 4.20E-78 | midnightblue | CYTH1 |
| 3.42E-75 | -0.57909 | 0.359145 | 0.448188 | 3.88E-75 | midnightblue | 1-Sep |
| 1.48E-69 | -0.51432 | 0.16398 | 0.289085 | 1.67E-69 | midnightblue | MGAT4A |
| 2.63E-68 | -0.50009 | 0.482481 | 0.551352 | 2.95E-68 | midnightblue | EZR |
| 3.35E-68 | -1.81837 | 0.225298 | 0.335424 | 3.74E-68 | midnightblue | TYROBP |
| 2.61E-67 | -1.12861 | 0.251927 | 0.356432 | 2.90E-67 | midnightblue | S100A11 |
| 2.85E-67 | -0.7277 | 0.934128 | 0.772062 | 3.15E-67 | midnightblue | CD52 |
| 2.45E-65 | -0.52377 | 0.520673 | 0.555178 | 2.68E-65 | midnightblue | ARHGEF1 |
| 9.82E-65 | -0.54141 | 0.344078 | 0.422432 | 1.07E-64 | midnightblue | CYTIP |
| 1.86E-63 | -0.55701 | 0.323756 | 0.443264 | 2.01E-63 | midnightblue | GADD45B |
| 2.88E-63 | -0.54109 | 0.971969 | 0.899458 | 3.09E-63 | midnightblue | SARAF |

| | | | | | | |
|---|---|---|---|---|---|---|
| 6.41E-63 | -0.55309 | 0.30939 | 0.399015 | 6.86E-63 | midnightblue | TRAF3IP3 |
| 3.12E-60 | -0.53396 | 0.295025 | 0.385659 | 3.32E-60 | midnightblue | P2RY8 |
| 1.01E-54 | -0.62654 | 0.476524 | 0.512275 | 1.07E-54 | midnightblue | NOSIP |
| 5.60E-48 | -0.66426 | 0.548353 | 0.556666 | 5.88E-48 | midnightblue | DNAJB1 |
| 6.11E-43 | -0.64367 | 0.426069 | 0.457328 | 6.39E-43 | midnightblue | AHNAK |
| 1.47E-35 | -0.60707 | 0.580238 | 0.552096 | 1.52E-35 | midnightblue | LEPROTL1 |
| 1.18E-29 | -0.93625 | 0.660827 | 0.619265 | 1.22E-29 | midnightblue | SAT1 |
| 8.11E-27 | -1.11311 | 0.62719 | 0.385801 | 8.34E-27 | midnightblue | CST3 |
| 6.84E-21 | -0.61768 | 0.813945 | 0.681015 | 6.99E-21 | midnightblue | PSAP |
| 2.06E-15 | -0.52133 | 0.808339 | 0.539129 | 2.09E-15 | midnightblue | CTSW |
| 4.86E-15 | -0.73521 | 0.420813 | 0.408793 | 4.92E-15 | midnightblue | TIMP1 |
| 7.04E-05 | -0.50324 | 0.87281 | 0.666773 | 7.08E-05 | midnightblue | TSPO |
| 0.001881 | -0.70496 | 0.544849 | 0.45669 | 0.001881 | midnightblue | CTSS |
| 0 | 1.52706 | 0.522064 | 0.108315 | 0 | brown | MPO |
| 0 | 1.410424 | 0.974881 | 0.453019 | 0 | brown | STMN1 |
| 0 | 1.409407 | 0.727088 | 0.10818 | 0 | brown | IRF8 |
| 0 | 1.346738 | 0.477936 | 0.046339 | 0 | brown | JCHAIN |
| 0 | 1.315912 | 0.756959 | 0.144083 | 0 | brown | PLD4 |
| 0 | 1.205831 | 0.938221 | 0.330789 | 0 | brown | ITM2C |
| 0 | 1.199952 | 0.761032 | 0.230444 | 0 | brown | IGLL1 |
| 0 | 1.193864 | 0.968092 | 0.647055 | 0 | brown | TUBA1B |
| 0 | 1.193147 | 0.851324 | 0.222947 | 0 | brown | NREP |
| 0 | 1.167738 | 0.925322 | 0.366185 | 0 | brown | SOX4 |
| 0 | 1.139997 | 0.724372 | 0.146683 | 0 | brown | UHRF1 |
| 0 | 1.091778 | 0.947047 | 0.608889 | 0 | brown | TUBB |
| 0 | 1.073716 | 0.458248 | 0.056302 | 0 | brown | DNTT |
| 0 | 1.058845 | 0.864902 | 0.47933 | 0 | brown | PLAC8 |
| 0 | 1.041943 | 0.989138 | 0.849433 | 0 | brown | HMGN2 |
| 0 | 1.019154 | 0.949762 | 0.598115 | 0 | brown | H2AFZ |
| 0 | 1.006836 | 0.611677 | 0.174784 | 0 | brown | IGHM |
| 0 | 0.982843 | 0.999321 | 0.971629 | 0 | brown | ACTG1 |
| 0 | 0.857858 | 0.830278 | 0.306471 | 0 | brown | LAT2 |
| 0 | 0.846646 | 0.727088 | 0.227709 | 0 | brown | SPINK2 |
| 0 | 0.834255 | 0.998642 | 0.916171 | 0 | brown | HMGB1 |
| 0 | 0.828419 | 0.964019 | 0.654012 | 0 | brown | VAMP8 |
| 0 | 0.82731 | 0.873727 | 0.346156 | 0 | brown | FABP5 |
| 0 | 0.794568 | 0.961982 | 0.53749 | 0 | brown | NUCB2 |
| 0 | 0.762002 | 0.75017 | 0.152324 | 0 | brown | SPNS3 |
| 0 | 0.74774 | 0.708079 | 0.239834 | 0 | brown | TCF4 |
| 0 | 0.746586 | 0.997284 | 0.724433 | 0 | brown | GSTP1 |
| 0 | 0.733867 | 0.704684 | 0.176169 | 0 | brown | ADA |

| | | | | | | |
|---|---|---|---|---|---|---|
| 0 | 0.732466 | 0.446707 | 0.025128 | 0 | brown | KCNE5 |
| 0 | 0.724355 | 0.933469 | 0.476594 | 0 | brown | H2AFY |
| 0 | 0.690378 | 0.76239 | 0.235308 | 0 | brown | SMIM24 |
| 0 | 0.684057 | 0.57909 | 0.067651 | 0 | brown | BLNK |
| 0 | 0.671054 | 0.334691 | 0.019049 | 0 | brown | ACY3 |
| 0 | 0.668961 | 0.549898 | 0.097913 | 0 | brown | RNASE6 |
| 0 | 0.66414 | 0.875085 | 0.39263 | 0 | brown | NASP |
| 0 | 0.641349 | 0.98167 | 0.688463 | 0 | brown | NAP1L1 |
| 0 | 0.635303 | 0.976918 | 0.739429 | 0 | brown | SLC25A5 |
| 0 | 0.627416 | 0.999321 | 0.96913 | 0 | brown | GAPDH |
| 0 | 0.621989 | 0.776646 | 0.293873 | 0 | brown | CAPG |
| 0 | 0.620563 | 0.631365 | 0.180053 | 0 | brown | MZB1 |
| 0 | 0.616691 | 0.863544 | 0.387429 | 0 | brown | RNASEH2B |
| 0 | 0.613369 | 0.608282 | 0.099027 | 0 | brown | FLT3 |
| 0 | 0.571428 | 0.826205 | 0.343218 | 0 | brown | NUDT1 |
| 0 | 0.566073 | 0.404616 | 0.056775 | 0 | brown | IL3RA |
| 0 | 0.562774 | 0.500339 | 0.075892 | 0 | brown | RUNX2 |
| 0 | 0.562354 | 0.607604 | 0.187821 | 0 | brown | C12orf75 |
| 0 | 0.557984 | 0.602172 | 0.163604 | 0 | brown | GSN |
| 0 | 0.551904 | 0.419552 | 0.039111 | 0 | brown | LGMN |
| 0 | 0.533812 | 0.720978 | 0.265367 | 0 | brown | LAIR1 |
| 0 | 0.52693 | 1 | 0.914854 | 0 | brown | CD74 |
| 0 | 0.523832 | 1 | 0.999189 | 0 | brown | PTMA |
| 0 | 0.512767 | 0.997963 | 0.973386 | 0 | brown | PPIA |
| 0 | -1.13097 | 0.895451 | 0.949507 | 0 | brown | HLA-E |
| 5.18E-297 | 0.618224 | 0.81127 | 0.353891 | 1.62E-296 | brown | DDAH2 |
| 2.15E-296 | 1.135489 | 0.73184 | 0.310186 | 6.61E-296 | brown | AREG |
| 9.79E-286 | 0.645746 | 0.967413 | 0.684714 | 2.95E-285 | brown | SEC61B |
| 4.92E-283 | 0.600574 | 0.95112 | 0.630944 | 1.46E-282 | brown | HMGN1 |
| 2.22E-281 | 0.58743 | 0.978955 | 0.56809 | 6.47E-281 | brown | TXN |
| 5.32E-277 | -0.67731 | 0.997963 | 0.994427 | 1.52E-276 | brown | RPS27 |
| 2.61E-269 | 0.598073 | 0.835709 | 0.415766 | 7.34E-269 | brown | IDH2 |
| 5.16E-267 | -1.327 | 0.513917 | 0.800527 | 1.43E-266 | brown | CD52 |
| 3.25E-266 | 0.54813 | 0.745418 | 0.302148 | 8.85E-266 | brown | MCM7 |
| 1.29E-265 | 0.532787 | 0.63408 | 0.235409 | 3.45E-265 | brown | APP |
| 9.26E-264 | 0.790607 | 0.777325 | 0.359835 | 2.45E-263 | brown | LGALS1 |
| 4.70E-261 | 0.51353 | 0.356415 | 0.087443 | 1.22E-260 | brown | MKI67 |
| 5.56E-253 | 0.600178 | 0.59131 | 0.220346 | 1.42E-252 | brown | PCNA |
| 2.17E-242 | 0.592906 | 0.986422 | 0.842272 | 5.48E-242 | brown | PPIB |
| 1.38E-238 | 0.586559 | 0.383571 | 0.107201 | 3.44E-238 | brown | NUSAP1 |
| 4.43E-238 | 0.63482 | 0.900204 | 0.594569 | 1.09E-237 | brown | DBI |

| | | | | | | |
|---|---|---|---|---|---|---|
| 2.60E-232 | 0.54292 | 0.787508 | 0.313463 | 6.27E-232 | brown | PRSS57 |
| 1.61E-221 | -1.72348 | 0.577733 | 0.78087 | 3.83E-221 | brown | JUNB |
| 3.88E-221 | -1.82166 | 0.203666 | 0.591293 | 9.11E-221 | brown | IFITM1 |
| 9.33E-208 | 0.583118 | 0.898167 | 0.556539 | 2.16E-207 | brown | NUCKS1 |
| 5.11E-205 | -0.83007 | 0.976239 | 0.986727 | 1.17E-204 | brown | HLA-A |
| 2.46E-196 | 0.59149 | 0.855397 | 0.529958 | 5.55E-196 | brown | PLP2 |
| 3.96E-195 | 0.750258 | 0.314325 | 0.086092 | 8.80E-195 | brown | UBE2C |
| 1.93E-194 | 0.505049 | 0.505771 | 0.195961 | 4.23E-194 | brown | CKS1B |
| 2.12E-192 | 0.501079 | 0.405974 | 0.13395 | 4.59E-192 | brown | CENPF |
| 3.85E-190 | 0.556994 | 0.89002 | 0.520501 | 8.24E-190 | brown | DUT |
| 7.09E-189 | -2.14144 | 0.015614 | 0.406714 | 1.50E-188 | brown | IL32 |
| 1.55E-185 | 0.562582 | 0.305499 | 0.084234 | 3.22E-185 | brown | TOP2A |
| 8.03E-178 | -0.56003 | 0.947726 | 0.927722 | 1.66E-177 | brown | TOMM7 |
| 3.55E-171 | 0.517812 | 0.849287 | 0.514895 | 7.24E-171 | brown | SIVA1 |
| 2.41E-170 | -0.51932 | 0.995927 | 0.999358 | 4.86E-170 | brown | MALAT1 |
| 6.99E-163 | -0.67685 | 0.911066 | 0.927553 | 1.39E-162 | brown | LAPTM5 |
| 1.78E-159 | 0.503466 | 0.942295 | 0.695893 | 3.50E-159 | brown | HSP90B1 |
| 1.57E-152 | -1.13162 | 0.709437 | 0.797825 | 3.05E-152 | brown | TXNIP |
| 2.77E-152 | 0.594669 | 0.588595 | 0.280195 | 5.33E-152 | brown | CLEC11A |
| 2.37E-151 | -0.94676 | 0.626612 | 0.754019 | 4.50E-151 | brown | PTPRC |
| 1.19E-148 | -0.82672 | 0.989817 | 0.993414 | 2.24E-148 | brown | HLA-C |
| 5.66E-148 | -0.70809 | 1 | 0.999662 | 1.05E-147 | brown | B2M |
| 8.66E-146 | -1.14326 | 0.010183 | 0.334808 | 1.59E-145 | brown | CD3D |
| 1.05E-142 | 0.749633 | 0.922607 | 0.670224 | 1.91E-142 | brown | HMGB2 |
| 1.76E-142 | -1.46917 | 0.74406 | 0.827749 | 3.16E-142 | brown | S100A6 |
| 1.64E-141 | -0.70783 | 0.995927 | 0.997467 | 2.92E-141 | brown | HLA-B |
| 1.30E-137 | 0.542596 | 0.721656 | 0.389489 | 2.29E-137 | brown | TUBB4B |
| 3.78E-137 | 0.515621 | 0.691785 | 0.357201 | 6.59E-137 | brown | MAP3K8 |
| 1.44E-133 | -0.64624 | 0.798371 | 0.840989 | 2.48E-133 | brown | LIMD2 |
| 1.62E-115 | -0.84281 | 0.176511 | 0.451027 | 2.77E-115 | brown | ZFP36L1 |
| 1.21E-113 | -0.80496 | 0.031908 | 0.30431 | 2.04E-113 | brown | PIK3IP1 |
| 4.67E-113 | -0.77565 | 0.769857 | 0.774048 | 7.82E-113 | brown | IFITM2 |
| 1.12E-112 | -1.1612 | 0.151392 | 0.422757 | 1.86E-112 | brown | CD7 |
| 2.18E-112 | -1.1213 | 0.180584 | 0.433667 | 3.57E-112 | brown | TNFAIP3 |
| 3.56E-112 | -0.86045 | 0.044807 | 0.316401 | 5.79E-112 | brown | SPOCK2 |
| 4.05E-111 | -0.9412 | 0.394433 | 0.618583 | 6.52E-111 | brown | S100A10 |
| 6.59E-109 | -1.55376 | 0.174474 | 0.412625 | 1.05E-108 | brown | CD3E |
| 4.86E-108 | -0.88867 | 0.006789 | 0.26476 | 7.68E-108 | brown | GZMM |
| 2.23E-107 | -0.79843 | 0.006789 | 0.263983 | 3.49E-107 | brown | CD3G |
| 2.33E-106 | 0.579789 | 0.518669 | 0.27094 | 3.62E-106 | brown | FAM30A |
| 6.45E-105 | -0.86293 | 0.695859 | 0.306877 | 9.90E-105 | brown | TYROBP |

| | | | | | | |
|---|---|---|---|---|---|---|
| 9.46E-105 | -0.69459 | 0.023761 | 0.278844 | 1.44E-104 | brown | LAT |
| 4.18E-97 | -0.85594 | 0.012899 | 0.251081 | 6.31E-97 | brown | CD247 |
| 4.71E-95 | -0.88613 | 0.094365 | 0.330924 | 7.05E-95 | brown | LCK |
| 1.05E-92 | -0.59404 | 0.021724 | 0.254289 | 1.56E-92 | brown | SKAP1 |
| 7.00E-92 | -0.86633 | 0.267481 | 0.493346 | 1.03E-91 | brown | CD69 |
| 8.45E-92 | -0.86928 | 0.687712 | 0.781444 | 1.23E-91 | brown | JUN |
| 2.24E-89 | -0.59348 | 0.867617 | 0.909315 | 3.24E-89 | brown | CD37 |
| 9.97E-88 | -0.77661 | 0.371351 | 0.535734 | 1.43E-87 | brown | BTG1 |
| 1.29E-87 | -0.71606 | 0.115411 | 0.341225 | 1.83E-87 | brown | ISG20 |
| 3.54E-86 | -0.57053 | 0.201629 | 0.421373 | 4.99E-86 | brown | TNFRSF14 |
| 9.38E-86 | -0.62804 | 0.921928 | 0.90533 | 1.31E-85 | brown | SARAF |
| 3.79E-78 | -0.85806 | 0.421589 | 0.562584 | 5.25E-78 | brown | DNAJB1 |
| 7.64E-78 | -0.83793 | 0.224711 | 0.41482 | 1.05E-77 | brown | SLC2A3 |
| 1.97E-77 | -0.6336 | 0.066531 | 0.275736 | 2.68E-77 | brown | LBH |
| 1.16E-69 | -0.65742 | 0.389681 | 0.538908 | 1.56E-69 | brown | RARRES3 |
| 1.73E-68 | -0.58621 | 0.328581 | 0.492772 | 2.32E-68 | brown | BIN2 |
| 5.95E-66 | -0.67224 | 0.082145 | 0.268745 | 7.92E-66 | brown | KLF2 |
| 4.17E-65 | -0.66921 | 0.101154 | 0.284687 | 5.51E-65 | brown | ZAP70 |
| 5.60E-64 | -0.64291 | 0.082824 | 0.264793 | 7.34E-64 | brown | FLT3LG |
| 5.52E-63 | -0.69831 | 0.297352 | 0.435794 | 7.17E-63 | brown | HLA-F |
| 1.10E-57 | 0.840844 | 0.820095 | 0.584133 | 1.41E-57 | brown | HIST1H4C |
| 3.95E-56 | -0.7351 | 0.528853 | 0.588422 | 5.06E-56 | brown | EVL |
| 8.60E-56 | -0.60146 | 0.293958 | 0.434849 | 1.09E-55 | brown | TBC1D10C |
| 9.83E-56 | -0.69731 | 0.3537 | 0.452276 | 1.24E-55 | brown | CD48 |
| 1.29E-54 | -0.55398 | 0.111337 | 0.27577 | 1.62E-54 | brown | PRKCH |
| 7.76E-54 | -0.72471 | 0.205703 | 0.367232 | 9.64E-54 | brown | NR4A2 |
| 3.82E-51 | -0.55104 | 0.470468 | 0.551574 | 4.71E-51 | brown | STK4 |
| 2.61E-50 | -0.54716 | 0.996606 | 0.993988 | 3.19E-50 | brown | FTL |
| 2.08E-48 | -0.69907 | 0.497624 | 0.557518 | 2.53E-48 | brown | LEPROTL1 |
| 2.14E-48 | -1.0755 | 0.145961 | 0.297251 | 2.59E-48 | brown | IL7R |
| 5.13E-48 | -0.7527 | 0.467753 | 0.522528 | 6.15E-48 | brown | LITAF |
| 1.27E-47 | -0.56809 | 0.116768 | 0.2704 | 1.51E-47 | brown | TUBA4A |
| 3.72E-45 | -0.58306 | 0.369314 | 0.489597 | 4.40E-45 | brown | IFITM3 |
| 1.69E-44 | -0.55534 | 0.757637 | 0.730343 | 1.99E-44 | brown | EMP3 |
| 4.28E-44 | -0.73844 | 0.441276 | 0.499223 | 4.99E-44 | brown | FYB1 |
| 1.16E-43 | -0.62215 | 0.14664 | 0.277594 | 1.34E-43 | brown | IL10RA |
| 2.75E-39 | -1.62018 | 0.55465 | 0.292995 | 3.16E-39 | brown | NKG7 |
| 4.01E-38 | -0.5662 | 0.339443 | 0.419008 | 4.58E-38 | brown | CYTIP |
| 8.72E-37 | -0.97801 | 0.899525 | 0.807349 | 9.90E-37 | brown | S100A4 |
| 1.73E-33 | -1.17295 | 0.182621 | 0.302418 | 1.95E-33 | brown | CST7 |
| 4.35E-32 | -0.68199 | 0.750849 | 0.690354 | 4.87E-32 | brown | PSAP |

| | | | | | | |
|---|---|---|---|---|---|---|
| 1.08E-31 | -0.63654 | 0.290563 | 0.40175 | 1.20E-31 | brown | LTB |
| 1.23E-31 | -0.56961 | 0.460964 | 0.49767 | 1.36E-31 | brown | AES |
| 6.95E-31 | -0.59683 | 0.201629 | 0.298602 | 7.62E-31 | brown | FCMR |
| 1.57E-29 | -0.54475 | 0.351663 | 0.423838 | 1.71E-29 | brown | BTG2 |
| 1.92E-26 | -0.53883 | 0.863544 | 0.816333 | 2.09E-26 | brown | TSC22D3 |
| 8.36E-23 | -0.66606 | 0.704684 | 0.641077 | 9.00E-23 | brown | ITGB2 |
| 1.94E-20 | -0.58071 | 0.442634 | 0.476527 | 2.07E-20 | brown | MT2A |
| 7.59E-20 | -0.71129 | 0.632722 | 0.605748 | 8.07E-20 | brown | NEAT1 |
| 4.64E-18 | -0.51482 | 0.534963 | 0.538064 | 4.91E-18 | brown | FLNA |
| 1.39E-16 | -0.75731 | 0.801765 | 0.703627 | 1.46E-16 | brown | ZFP36 |
| 2.27E-13 | -0.53394 | 0.392396 | 0.412321 | 2.36E-13 | brown | NFKBIA |
| 1.58E-11 | -0.96352 | 0.334012 | 0.347474 | 1.64E-11 | brown | S100A11 |
| 4.29E-09 | -0.53485 | 0.260692 | 0.29769 | 4.42E-09 | brown | NAMPT |
| 2.55E-06 | -0.58808 | 0.441955 | 0.422217 | 2.62E-06 | brown | DUSP1 |
| 1.70E-05 | -0.85609 | 0.724372 | 0.618042 | 1.73E-05 | brown | SAT1 |
| 0 | 2.417921 | 0.67374 | 0.069268 | 0 | salmon | MPO |
| 0 | 2.090525 | 0.30305 | 0.022448 | 0 | salmon | PRTN3 |
| 0 | 1.77073 | 0.378647 | 0.040834 | 0 | salmon | AZU1 |
| 0 | 1.700717 | 0.285146 | 0.0217 | 0 | salmon | ELANE |
| 0 | 1.567446 | 0.993037 | 0.265313 | 0 | salmon | PRSS57 |
| 0 | 1.250656 | 0.957891 | 0.18532 | 0 | salmon | SMIM24 |
| 0 | 1.225052 | 0.788462 | 0.24176 | 0 | salmon | CLEC11A |
| 0 | 1.181894 | 0.929377 | 0.178514 | 0 | salmon | SPINK2 |
| 0 | 1.174234 | 0.770889 | 0.282808 | 0 | salmon | AREG |
| 0 | 1.159933 | 0.9937 | 0.688758 | 0 | salmon | CALR |
| 0 | 1.084294 | 0.918435 | 0.215179 | 0 | salmon | MGST1 |
| 0 | 0.997581 | 0.98508 | 0.627579 | 0 | salmon | TUBA1B |
| 0 | 0.971677 | 0.988064 | 0.511349 | 0 | salmon | NUCB2 |
| 0 | 0.938782 | 0.95126 | 0.308819 | 0 | salmon | FABP5 |
| 0 | 0.898749 | 0.988395 | 0.422875 | 0 | salmon | STMN1 |
| 0 | 0.894635 | 0.552056 | 0.038803 | 0 | salmon | CTSG |
| 0 | 0.888148 | 0.996021 | 0.568324 | 0 | salmon | SERPINB1 |
| 0 | 0.868508 | 0.983753 | 0.575129 | 0 | salmon | H2AFZ |
| 0 | 0.865723 | 0.850464 | 0.191662 | 0 | salmon | IGLL1 |
| 0 | 0.858168 | 1 | 0.967397 | 0 | salmon | GAPDH |
| 0 | 0.818626 | 0.697281 | 0.155532 | 0 | salmon | EREG |
| 0 | 0.818394 | 0.995027 | 0.697773 | 0 | salmon | ENO1 |
| 0 | 0.806023 | 0.568966 | 0.084661 | 0 | salmon | RNASE2 |
| 0 | 0.802762 | 0.950597 | 0.402922 | 0 | salmon | CAT |
| 0 | 0.796795 | 0.691645 | 0.072403 | 0 | salmon | CPA3 |
| 0 | 0.784106 | 0.966844 | 0.491929 | 0 | salmon | DUT |

| | | | | | | |
|---|---|---|---|---|---|---|
| 0 | 0.783593 | 0.971817 | 0.437057 | 0 | salmon | IMPDH2 |
| 0 | 0.76333 | 0.966844 | 0.302975 | 0 | salmon | AC084033.3 |
| 0 | 0.763078 | 0.975133 | 0.649742 | 0 | salmon | GYPC |
| 0 | 0.749455 | 0.963528 | 0.475539 | 0 | salmon | HSPB1 |
| 0 | 0.748953 | 0.967838 | 0.588063 | 0 | salmon | TUBB |
| 0 | 0.74319 | 0.999668 | 0.881062 | 0 | salmon | RPS2 |
| 0 | 0.728118 | 0.936008 | 0.309318 | 0 | salmon | BEX3 |
| 0 | 0.722189 | 0.998674 | 0.709282 | 0 | salmon | GSTP1 |
| 0 | 0.718256 | 0.975464 | 0.482273 | 0 | salmon | HSPD1 |
| 0 | 0.707129 | 0.934682 | 0.343381 | 0 | salmon | CDK6 |
| 0 | 0.704697 | 0.992374 | 0.544058 | 0 | salmon | TXN |
| 0 | 0.698652 | 0.988727 | 0.671584 | 0 | salmon | NAP1L1 |
| 0 | 0.675755 | 0.878316 | 0.244397 | 0 | salmon | EGFL7 |
| 0 | 0.653964 | 0.847812 | 0.266774 | 0 | salmon | MCM7 |
| 0 | 0.650239 | 0.767905 | 0.177944 | 0 | salmon | CDCA7 |
| 0 | 0.648962 | 0.846817 | 0.265491 | 0 | salmon | GNA15 |
| 0 | 0.648308 | 1 | 0.854908 | 0 | salmon | HSP90AB1 |
| 0 | 0.622533 | 0.997679 | 0.808587 | 0 | salmon | MIF |
| 0 | 0.621949 | 0.900531 | 0.309033 | 0 | salmon | CDK4 |
| 0 | 0.621649 | 0.915119 | 0.361803 | 0 | salmon | NASP |
| 0 | 0.620657 | 0.956897 | 0.537324 | 0 | salmon | PRDX1 |
| 0 | 0.617374 | 0.997347 | 0.840869 | 0 | salmon | HMGN2 |
| 0 | 0.602325 | 0.854775 | 0.213184 | 0 | salmon | CSF3R |
| 0 | 0.601139 | 0.820955 | 0.221058 | 0 | salmon | CD34 |
| 0 | 0.594831 | 0.987069 | 0.634491 | 0 | salmon | VAMP8 |
| 0 | 0.58971 | 0.854111 | 0.258186 | 0 | salmon | ARMH1 |
| 0 | 0.582899 | 0.999668 | 0.906111 | 0 | salmon | HINT1 |
| 0 | 0.578796 | 0.907493 | 0.300695 | 0 | salmon | ITM2C |
| 0 | 0.571093 | 1 | 0.994833 | 0 | salmon | RPLP0 |
| 0 | 0.567246 | 0.989721 | 0.684732 | 0 | salmon | RAN |
| 0 | 0.562402 | 0.991379 | 0.578336 | 0 | salmon | TALDO1 |
| 0 | 0.557447 | 0.602454 | 0.102975 | 0 | salmon | AC002454.1 |
| 0 | 0.551211 | 0.987401 | 0.629396 | 0 | salmon | RPL22L1 |
| 0 | 0.550506 | 0.999005 | 0.910066 | 0 | salmon | NPM1 |
| 0 | 0.546791 | 0.866048 | 0.292143 | 0 | salmon | HMGA1 |
| 0 | 0.536094 | 0.967175 | 0.501158 | 0 | salmon | SNRPD1 |
| 0 | 0.530658 | 0.849801 | 0.3199 | 0 | salmon | RANBP1 |
| 0 | 0.524332 | 0.644231 | 0.085765 | 0 | salmon | SPARC |
| 0 | 0.520702 | 0.791777 | 0.238446 | 0 | salmon | ANKRD28 |
| 0 | 0.519292 | 0.687003 | 0.123285 | 0 | salmon | PHGDH |
| 0 | 0.515778 | 0.873011 | 0.323927 | 0 | salmon | AHCY |

| | | | | | | |
|---|---|---|---|---|---|---|
| 0 | 0.514398 | 0.772215 | 0.224906 | 0 | salmon | ISYNA1 |
| 0 | 0.509261 | 0.913462 | 0.418671 | 0 | salmon | PDIA6 |
| 0 | 0.508314 | 0.947613 | 0.417887 | 0 | salmon | COMT |
| 0 | 0.501165 | 0.958554 | 0.470479 | 0 | salmon | OSTC |
| 0 | 0.500393 | 0.972812 | 0.533333 | 0 | salmon | APEX1 |
| 0 | -0.52953 | 0.999337 | 0.996615 | 0 | salmon | EIF1 |
| 0 | -0.60019 | 0.999337 | 0.994085 | 0 | salmon | RPS27 |
| 0 | -0.60363 | 0.924403 | 0.873223 | 0 | salmon | YWHAZ |
| 0 | -0.61628 | 0.918435 | 0.902369 | 0 | salmon | DAZAP2 |
| 0 | -0.75482 | 0.985743 | 0.980046 | 0 | salmon | UBC |
| 0 | -0.78386 | 0.854775 | 0.837271 | 0 | salmon | LIMD2 |
| 0 | -0.81619 | 0.931034 | 0.904792 | 0 | salmon | CD37 |
| 0 | -0.88009 | 0.706897 | 0.780189 | 0 | salmon | GPSM3 |
| 0 | -0.88358 | 0.930371 | 0.90351 | 0 | salmon | SARAF |
| 0 | -0.89683 | 0.988064 | 0.966791 | 0 | salmon | SH3BGRL3 |
| 0 | -0.90269 | 0.998342 | 0.999287 | 0 | salmon | MALAT1 |
| 0 | -0.93731 | 0.920424 | 0.891538 | 0 | salmon | MYL12A |
| 0 | -0.93827 | 0.988395 | 0.985997 | 0 | salmon | HLA-A |
| 0 | -0.9502 | 0.892905 | 0.930412 | 0 | salmon | LAPTM5 |
| 0 | -0.96228 | 0.831897 | 0.819277 | 0 | salmon | CORO1A |
| 0 | -1.01862 | 0.877984 | 0.886086 | 0 | salmon | CALM1 |
| 0 | -1.03122 | 1 | 0.99861 | 0 | salmon | TMSB10 |
| 0 | -1.03866 | 0.01061 | 0.352931 | 0 | salmon | LCK |
| 0 | -1.11039 | 0.999337 | 0.993087 | 0 | salmon | TMSB4X |
| 0 | -1.13003 | 0.765584 | 0.824265 | 0 | salmon | TSC22D3 |
| 0 | -1.14168 | 1 | 0.999644 | 0 | salmon | B2M |
| 0 | -1.16015 | 0.597812 | 0.746018 | 0 | salmon | EMP3 |
| 0 | -1.18678 | 0.636605 | 0.75995 | 0 | salmon | PTPRC |
| 0 | -1.18884 | 0.001658 | 0.353572 | 0 | salmon | CD3D |
| 0 | -1.19667 | 0.366048 | 0.609193 | 0 | salmon | EVL |
| 0 | -1.21291 | 0.996353 | 0.997506 | 0 | salmon | HLA-B |
| 0 | -1.24944 | 0.993369 | 0.99323 | 0 | salmon | HLA-C |
| 0 | -1.2688 | 0.905836 | 0.951363 | 0 | salmon | HLA-E |
| 0 | -1.34616 | 0.751989 | 0.7907 | 0 | salmon | CD52 |
| 0 | -1.56261 | 0.628979 | 0.811331 | 0 | salmon | TXNIP |
| 0 | -1.64108 | 0.871021 | 0.805345 | 0 | salmon | S100A4 |
| 0 | -1.69728 | 0.038793 | 0.434919 | 0 | salmon | LTB |
| 0 | -1.78625 | 0.313329 | 0.639622 | 0 | salmon | S100A10 |
| 0 | -1.86217 | 0.814655 | 0.824764 | 0 | salmon | S100A6 |
| 0 | -1.86864 | 0.639589 | 0.785391 | 0 | salmon | JUNB |
| 0 | -1.93585 | 0.023541 | 0.441938 | 0 | salmon | CD3E |

| | | | | | | |
|---|---|---|---|---|---|---|
| 0 | -2.22731 | 0.002984 | 0.429574 | 0 | salmon | IL32 |
| 3.00E-280 | -1.49107 | 0.004973 | 0.32072 | 6.05E-280 | salmon | IL7R |
| 2.58E-278 | -1.22526 | 0.600133 | 0.7199 | 5.17E-278 | salmon | ZFP36 |
| 1.25E-275 | -0.93885 | 0.327255 | 0.548691 | 2.47E-275 | salmon | CCND3 |
| 1.39E-271 | -1.1267 | 0.283156 | 0.519401 | 2.73E-271 | salmon | FYB1 |
| 4.71E-263 | -0.99852 | 0.190318 | 0.464636 | 9.17E-263 | salmon | ZFP36L1 |
| 2.03E-262 | -0.89416 | 0.445955 | 0.596116 | 3.92E-262 | salmon | 9-Sep |
| 5.06E-257 | -1.17478 | 0.496353 | 0.616141 | 9.68E-257 | salmon | CXCR4 |
| 6.84E-250 | -1.39619 | 0.165119 | 0.436202 | 1.30E-249 | salmon | CD7 |
| 1.16E-248 | -0.87899 | 0.032162 | 0.322145 | 2.18E-248 | salmon | FCMR |
| 1.15E-241 | -0.84359 | 0.048077 | 0.332514 | 2.14E-241 | salmon | ARL4C |
| 1.16E-241 | -0.85003 | 0.034483 | 0.319009 | 2.14E-241 | salmon | PIK3IP1 |
| 1.03E-239 | -0.84001 | 0 | 0.278853 | 1.88E-239 | salmon | CD3G |
| 2.92E-238 | -0.92629 | 0.001658 | 0.279494 | 5.31E-238 | salmon | GZMM |
| 5.70E-238 | -0.83386 | 0.079244 | 0.357527 | 1.03E-237 | salmon | ISG20 |
| 5.62E-235 | -1.20918 | 0.19695 | 0.445822 | 1.01E-234 | salmon | TNFAIP3 |
| 2.64E-223 | -0.7768 | 0.029509 | 0.297381 | 4.69E-223 | salmon | IL10RA |
| 1.76E-222 | -0.71874 | 0.024536 | 0.292785 | 3.10E-222 | salmon | LAT |
| 3.01E-220 | -0.92115 | 0.064655 | 0.3292 | 5.26E-220 | salmon | SPOCK2 |
| 2.96E-219 | -0.82052 | 0.07195 | 0.339818 | 5.13E-219 | salmon | SAMHD1 |
| 1.25E-212 | -1.71802 | 0.489058 | 0.581935 | 2.15E-212 | salmon | IFITM1 |
| 1.80E-212 | -0.89277 | 0.009284 | 0.264564 | 3.08E-212 | salmon | CD247 |
| 5.49E-208 | -0.91001 | 0.270889 | 0.466595 | 9.30E-208 | salmon | CD48 |
| 2.00E-207 | -0.75259 | 0.880968 | 0.796472 | 3.34E-207 | salmon | LSP1 |
| 2.00E-207 | -0.92 | 0.394231 | 0.542312 | 3.34E-207 | salmon | BTG1 |
| 4.44E-198 | -1.07578 | 0.636273 | 0.644931 | 7.35E-198 | salmon | ITGB2 |
| 3.87E-194 | -0.8448 | 0.034151 | 0.277071 | 6.36E-194 | salmon | NCF1 |
| 2.24E-188 | -0.78193 | 0.061671 | 0.29902 | 3.65E-188 | salmon | ZAP70 |
| 7.03E-181 | -1.00966 | 0.679708 | 0.655158 | 1.14E-180 | salmon | HCST |
| 7.83E-180 | -0.7578 | 0.051061 | 0.282345 | 1.26E-179 | salmon | KLF2 |
| 7.00E-168 | -0.80592 | 0.274536 | 0.445858 | 1.12E-167 | salmon | HLA-F |
| 1.62E-165 | -0.59316 | 0.759947 | 0.705861 | 2.57E-165 | salmon | HCLS1 |
| 2.27E-161 | -0.68732 | 0.072613 | 0.286585 | 3.58E-161 | salmon | LBH |
| 2.42E-159 | -0.78026 | 0.375 | 0.508926 | 3.79E-159 | salmon | AES |
| 3.89E-153 | -0.9263 | 0.255637 | 0.421949 | 6.04E-153 | salmon | SLC2A3 |
| 4.96E-152 | -0.69492 | 0.765252 | 0.693105 | 7.64E-152 | salmon | UCP2 |
| 7.73E-151 | -0.71793 | 0.072281 | 0.280634 | 1.18E-150 | salmon | ALOX5AP |
| 1.06E-140 | -0.73692 | 0.299403 | 0.44201 | 1.61E-140 | salmon | TBC1D10C |
| 2.03E-140 | -0.56395 | 0.138594 | 0.333405 | 3.07E-140 | salmon | SLC44A2 |
| 1.97E-131 | -0.6925 | 0.089523 | 0.274078 | 2.96E-131 | salmon | FLT3LG |
| 5.08E-130 | -0.57934 | 0.07626 | 0.261215 | 7.56E-130 | salmon | SKAP1 |

| | | | | | | |
|---|---|---|---|---|---|---|
| 4.28E-128 | -0.61582 | 0.119363 | 0.300053 | 6.34E-128 | salmon | CRIP1 |
| 5.39E-127 | -0.89463 | 0.478448 | 0.52439 | 7.92E-127 | salmon | LITAF |
| 1.50E-125 | -0.6807 | 0.514589 | 0.556031 | 2.19E-125 | salmon | ARHGEF1 |
| 2.06E-125 | -0.85484 | 0.783488 | 0.68352 | 2.99E-125 | salmon | PSAP |
| 4.20E-122 | -0.65184 | 0.503647 | 0.552467 | 6.05E-122 | salmon | STK4 |
| 3.98E-121 | -0.66757 | 0.104111 | 0.280207 | 5.70E-121 | salmon | TUBA4A |
| 3.63E-111 | -0.64224 | 0.628979 | 0.613006 | 5.17E-111 | salmon | MYH9 |
| 2.92E-110 | -0.59492 | 0.160146 | 0.320292 | 4.12E-110 | salmon | GNG2 |
| 1.93E-108 | -0.75304 | 0.319297 | 0.431285 | 2.72E-108 | salmon | BTG2 |
| 8.01E-103 | -0.77689 | 0.324934 | 0.443827 | 1.12E-102 | salmon | GADD45B |
| 4.84E-100 | -0.50469 | 0.844164 | 0.729735 | 6.71E-100 | salmon | ITM2B |
| 1.40E-98 | -0.64766 | 0.320955 | 0.42651 | 1.92E-98 | salmon | RIPOR2 |
| 4.37E-96 | -0.68233 | 0.373342 | 0.447176 | 5.98E-96 | salmon | 1-Sep |
| 6.37E-93 | -0.59769 | 0.245358 | 0.362409 | 8.67E-93 | salmon | IL16 |
| 1.16E-92 | -0.57252 | 0.204244 | 0.334616 | 1.57E-92 | salmon | CYTH1 |
| 1.47E-90 | -0.65738 | 0.344828 | 0.422804 | 1.98E-90 | salmon | CYTIP |
| 2.28E-86 | -1.26973 | 0.134615 | 0.270764 | 3.05E-86 | salmon | FCER1G |
| 3.98E-84 | -0.79865 | 0.553714 | 0.556138 | 5.29E-84 | salmon | DNAJB1 |
| 2.62E-83 | -0.51854 | 0.980769 | 0.561447 | 3.46E-83 | salmon | HLA-DRA |
| 3.48E-82 | -0.52842 | 0.171088 | 0.300695 | 4.58E-82 | salmon | ATM |
| 1.99E-81 | -0.55836 | 0.528846 | 0.542811 | 2.59E-81 | salmon | RSRP1 |
| 6.33E-81 | -0.63955 | 0.314324 | 0.399002 | 8.22E-81 | salmon | TRAF3IP3 |
| 1.20E-80 | -0.58162 | 0.154841 | 0.280135 | 1.54E-80 | salmon | PRKCH |
| 1.08E-76 | -0.57508 | 0.572281 | 0.552646 | 1.39E-76 | salmon | STK17B |
| 1.34E-75 | -0.82079 | 0.41943 | 0.458222 | 1.72E-75 | salmon | AHNAK |
| 1.47E-75 | -0.55565 | 0.522546 | 0.524853 | 1.87E-75 | salmon | ANKRD12 |
| 9.58E-72 | -0.54097 | 0.241048 | 0.343666 | 1.21E-71 | salmon | RBL2 |
| 1.45E-67 | -0.73238 | 0.589854 | 0.5509 | 1.82E-67 | salmon | LEPROTL1 |
| 1.73E-65 | -0.50874 | 0.252984 | 0.34641 | 2.16E-65 | salmon | CDC42SE1 |
| 2.08E-64 | -0.5214 | 0.196618 | 0.301514 | 2.58E-64 | salmon | KIAA1551 |
| 6.79E-64 | -0.66461 | 0.345159 | 0.419205 | 8.37E-64 | salmon | DDIT4 |
| 4.43E-61 | -0.59652 | 0.379642 | 0.424835 | 5.43E-61 | salmon | IQGAP1 |
| 5.97E-58 | -0.56832 | 0.596154 | 0.552931 | 7.28E-58 | salmon | YPEL5 |
| 1.14E-56 | -0.6562 | 0.58057 | 0.526599 | 1.39E-56 | salmon | RARRES3 |
| 3.93E-56 | -0.51573 | 0.258621 | 0.339426 | 4.75E-56 | salmon | CDKN1B |
| 5.91E-56 | -0.54432 | 0.257626 | 0.336718 | 7.09E-56 | salmon | EML4 |
| 1.86E-52 | -0.59132 | 0.328581 | 0.373704 | 2.23E-52 | salmon | RNF213 |
| 1.05E-50 | -1.1834 | 0.301393 | 0.351719 | 1.24E-50 | salmon | S100A11 |
| 8.48E-49 | -0.61863 | 0.443966 | 0.441867 | 1.00E-48 | salmon | VSIR |
| 6.91E-46 | -0.77381 | 0.499668 | 0.464707 | 8.12E-46 | salmon | TRBC2 |
| 1.78E-45 | -0.52659 | 0.508952 | 0.483592 | 2.08E-45 | salmon | WIPF1 |

| | | | | | | |
|---|---|---|---|---|---|---|
| 3.51E-45 | -0.54217 | 0.511605 | 0.487048 | 4.08E-45 | salmon | PRMT2 |
| 3.57E-43 | -0.6672 | 0.597812 | 0.53148 | 4.13E-43 | salmon | FLNA |
| 5.00E-43 | -0.52762 | 0.216844 | 0.284126 | 5.76E-43 | salmon | MGAT4A |
| 2.83E-42 | -0.72998 | 0.322944 | 0.363513 | 3.24E-42 | salmon | NR4A2 |
| 1.51E-40 | -0.50812 | 0.225796 | 0.293604 | 1.72E-40 | salmon | PPP2R5C |
| 4.65E-40 | -0.59283 | 0.346154 | 0.378621 | 5.26E-40 | salmon | HERPUD1 |
| 1.01E-39 | -0.5144 | 0.395889 | 0.404525 | 1.14E-39 | salmon | ADD3 |
| 1.70E-37 | -0.70358 | 0.471485 | 0.277962 | 1.90E-37 | salmon | CST7 |
| 1.09E-36 | -0.54196 | 0.6187 | 0.53711 | 1.22E-36 | salmon | EZR |
| 3.79E-34 | -0.90477 | 0.504973 | 0.460467 | 4.21E-34 | salmon | CTSS |
| 2.70E-31 | -0.51116 | 0.537467 | 0.479352 | 2.98E-31 | salmon | BIN2 |
| 1.09E-25 | -0.53014 | 0.47679 | 0.289221 | 1.20E-25 | salmon | HLA-DQA1 |
| 4.69E-23 | -1.94549 | 0.382294 | 0.236843 | 5.12E-23 | salmon | LYZ |
| 1.02E-21 | -0.5062 | 0.604443 | 0.486086 | 1.11E-21 | salmon | ANXA6 |
| 1.21E-20 | -0.64045 | 0.287135 | 0.302227 | 1.31E-20 | salmon | ANXA2 |
| 1.38E-19 | -0.59515 | 0.567308 | 0.346838 | 1.49E-19 | salmon | HLA-DQB1 |
| 3.65E-19 | -0.94286 | 0.736074 | 0.610939 | 3.91E-19 | salmon | SAT1 |
| 1.84E-18 | -1.20433 | 0.633621 | 0.383716 | 1.97E-18 | salmon | CST3 |
| 1.25E-17 | -0.73825 | 0.999668 | 0.910208 | 1.33E-17 | salmon | CD74 |
| 4.32E-17 | -0.6177 | 0.946618 | 0.608195 | 4.57E-17 | salmon | HLA-DRB1 |
| 6.13E-16 | -0.54545 | 0.296751 | 0.307964 | 6.45E-16 | salmon | ID2 |
| 8.68E-16 | -0.58274 | 0.621353 | 0.496918 | 9.09E-16 | salmon | NOSIP |
| 2.78E-13 | -0.52983 | 0.503979 | 0.438197 | 2.90E-13 | salmon | IRF1 |
| 1.76E-06 | -0.5229 | 0.901857 | 0.662462 | 1.83E-06 | salmon | TSPO |
| 8.48E-06 | -0.73685 | 0.261605 | 0.188028 | 8.75E-06 | salmon | CD79B |
| 1.46E-05 | -0.75865 | 0.495027 | 0.400748 | 1.51E-05 | salmon | TIMP1 |
| 4.36E-05 | -0.8643 | 0.538793 | 0.362516 | 4.46E-05 | salmon | LGALS1 |
| 4.56E-05 | -0.52317 | 0.579907 | 0.47219 | 4.65E-05 | salmon | CD69 |
| 0 | 2.199973 | 0.971223 | 0.200655 | 0 | pink | HBB |
| 0 | 1.967983 | 0.978417 | 0.080852 | 0 | pink | HBD |
| 0 | 1.85724 | 0.992806 | 0.054676 | 0 | pink | APOC1 |
| 0 | 1.747325 | 0.841727 | 0.056609 | 0 | pink | CA1 |
| 0 | 1.462421 | 0.996403 | 0.265717 | 0 | pink | BLVRB |
| 0 | 1.336166 | 0.93705 | 0.207764 | 0 | pink | MYC |
| 0 | 1.290576 | 1 | 0.427027 | 0 | pink | TMEM14C |
| 0 | 1.262015 | 0.985612 | 0.087862 | 0 | pink | CNRIP1 |
| 0 | 1.178294 | 0.985612 | 0.085471 | 0 | pink | GATA1 |
| 0 | 1.153428 | 0.996403 | 0.342703 | 0 | pink | UROD |
| 0 | 1.065775 | 0.969424 | 0.072301 | 0 | pink | ITGA2B |
| 0 | 1.005609 | 0.953237 | 0.064996 | 0 | pink | KLF1 |
| 0 | 1.004481 | 0.953237 | 0.086355 | 0 | pink | PVT1 |

| | | | | | | |
|---|---|---|---|---|---|---|
| 0 | 0.944128 | 0.991007 | 0.27276 | 0 | pink | REXO2 |
| 0 | 0.922365 | 0.989209 | 0.226699 | 0 | pink | CYTL1 |
| 0 | 0.853948 | 0.964029 | 0.185586 | 0 | pink | FSCN1 |
| 0 | 0.83494 | 0.785971 | 0.033579 | 0 | pink | FAM178B |
| 0 | 0.819356 | 0.994604 | 0.279214 | 0 | pink | CYTOR |
| 0 | 0.787893 | 0.926259 | 0.101556 | 0 | pink | SMIM10 |
| 0 | 0.784786 | 0.830935 | 0.047011 | 0 | pink | EPCAM |
| 0 | 0.777306 | 0.904676 | 0.067912 | 0 | pink | FN3K |
| 0 | 0.77137 | 0.951439 | 0.20285 | 0 | pink | PLIN2 |
| 0 | 0.767711 | 0.96223 | 0.193645 | 0 | pink | ALDH1A1 |
| 0 | 0.737353 | 0.76259 | 0.033481 | 0 | pink | AHSP |
| 0 | 0.67342 | 0.834532 | 0.037772 | 0 | pink | KCNH2 |
| 0 | 0.667636 | 0.956835 | 0.20095 | 0 | pink | NME4 |
| 0 | 0.665217 | 0.877698 | 0.070238 | 0 | pink | SMIM1 |
| 0 | 0.658974 | 0.928058 | 0.186503 | 0 | pink | NET1 |
| 0 | 0.654135 | 0.877698 | 0.065684 | 0 | pink | TFR2 |
| 0 | 0.64738 | 0.897482 | 0.182473 | 0 | pink | CENPU |
| 0 | 0.64647 | 0.868705 | 0.071974 | 0 | pink | FHL2 |
| 0 | 0.644029 | 0.839928 | 0.047076 | 0 | pink | ANK1 |
| 0 | 0.636719 | 0.895683 | 0.176708 | 0 | pink | LMNA |
| 0 | 0.636042 | 0.94964 | 0.213792 | 0 | pink | ISOC2 |
| 0 | 0.635402 | 0.96223 | 0.238624 | 0 | pink | HDAC7 |
| 0 | 0.633805 | 0.94964 | 0.19027 | 0 | pink | MIR4435-2HG |
| 0 | 0.633242 | 0.870504 | 0.137592 | 0 | pink | CASP3 |
| 0 | 0.629206 | 0.915468 | 0.159148 | 0 | pink | MAP7 |
| 0 | 0.613198 | 0.911871 | 0.192695 | 0 | pink | NFE2 |
| 0 | 0.6093 | 0.915468 | 0.146929 | 0 | pink | HMBS |
| 0 | 0.601029 | 0.902878 | 0.125012 | 0 | pink | TPM1 |
| 0 | 0.593119 | 0.965827 | 0.216577 | 0 | pink | SLC40A1 |
| 0 | 0.590209 | 0.958633 | 0.210549 | 0 | pink | PDLIM1 |
| 0 | 0.581365 | 0.920863 | 0.217101 | 0 | pink | MPST |
| 0 | 0.574936 | 0.859712 | 0.102604 | 0 | pink | TST |
| 0 | 0.567837 | 0.870504 | 0.132973 | 0 | pink | FADS2 |
| 0 | 0.562991 | 0.935252 | 0.236462 | 0 | pink | PSMG1 |
| 0 | 0.561863 | 0.877698 | 0.144767 | 0 | pink | CMBL |
| 0 | 0.558971 | 0.901079 | 0.191417 | 0 | pink | HNRNPAB |
| 0 | 0.557741 | 0.913669 | 0.177854 | 0 | pink | PCCB |
| 0 | 0.552811 | 0.766187 | 0.135561 | 0 | pink | CENPF |
| 0 | 0.550562 | 0.859712 | 0.095692 | 0 | pink | NFIA |
| 0 | 0.547916 | 0.832734 | 0.05294 | 0 | pink | PPP1R14A |
| 0 | 0.536922 | 0.883094 | 0.187158 | 0 | pink | FKBP4 |

| | | | | | | |
|---|---|---|---|---|---|---|
| 0 | 0.534365 | 0.926259 | 0.188567 | 0 | pink | PKIG |
| 0 | 0.532492 | 0.884892 | 0.188075 | 0 | pink | MCM2 |
| 0 | 0.531277 | 0.908273 | 0.208976 | 0 | pink | AKR1C3 |
| 0 | 0.528753 | 0.834532 | 0.117183 | 0 | pink | MINPP1 |
| 0 | 0.528441 | 0.931655 | 0.21448 | 0 | pink | MTHFD1 |
| 0 | 0.527814 | 0.911871 | 0.170025 | 0 | pink | LPCAT3 |
| 0 | 0.516632 | 0.915468 | 0.204324 | 0 | pink | STAT5A |
| 1.41E-304 | -2.27667 | 0.964029 | 0.994234 | 8.51E-304 | pink | TMSB4X |
| 5.13E-301 | 0.990745 | 1 | 0.355872 | 3.04E-300 | pink | AC084033.3 |
| 5.59E-299 | 0.605391 | 0.951439 | 0.250287 | 3.26E-298 | pink | MEST |
| 1.12E-295 | 0.939386 | 0.994604 | 0.420737 | 6.40E-295 | pink | MPC2 |
| 9.93E-294 | 0.599585 | 0.953237 | 0.259099 | 5.61E-293 | pink | MGST2 |
| 3.08E-292 | 0.699512 | 0.93705 | 0.268436 | 1.71E-291 | pink | SYNGR1 |
| 7.93E-289 | 0.775065 | 0.985612 | 0.337101 | 4.35E-288 | pink | DNPH1 |
| 1.74E-288 | 0.842149 | 0.992806 | 0.35502 | 9.43E-288 | pink | CDK4 |
| 3.40E-286 | 0.669544 | 0.971223 | 0.296118 | 1.81E-285 | pink | TPGS2 |
| 4.64E-280 | 0.655004 | 0.980216 | 0.291138 | 2.44E-279 | pink | PAICS |
| 1.16E-279 | 0.726527 | 0.902878 | 0.240721 | 5.99E-279 | pink | CKS2 |
| 5.87E-278 | 0.858014 | 0.996403 | 0.405471 | 3.00E-277 | pink | FBXO7 |
| 3.19E-274 | 0.779422 | 0.991007 | 0.339656 | 1.61E-273 | pink | CD82 |
| 6.87E-274 | 0.635679 | 0.967626 | 0.289599 | 3.42E-273 | pink | DKC1 |
| 1.37E-273 | 0.511301 | 0.931655 | 0.244717 | 6.70E-273 | pink | SLC39A8 |
| 3.24E-268 | 0.561649 | 0.967626 | 0.275381 | 1.57E-267 | pink | HADH |
| 3.06E-264 | 0.616188 | 0.93705 | 0.266011 | 1.47E-263 | pink | ISYNA1 |
| 6.72E-264 | 0.878848 | 0.998201 | 0.323866 | 3.17E-263 | pink | PRSS57 |
| 7.49E-264 | 0.584364 | 0.940647 | 0.253104 | 3.49E-263 | pink | MCM3 |
| 3.42E-261 | 0.519439 | 0.971223 | 0.279574 | 1.57E-260 | pink | MRPS2 |
| 4.40E-252 | 0.893525 | 0.998201 | 0.542703 | 2.00E-251 | pink | PRDX2 |
| 1.50E-250 | -1.59801 | 0.973022 | 0.96878 | 6.73E-250 | pink | SH3BGRL3 |
| 1.61E-250 | 0.548113 | 0.776978 | 0.187813 | 7.14E-250 | pink | UNG |
| 3.62E-250 | 0.934037 | 0.998201 | 0.521605 | 1.59E-249 | pink | HSPD1 |
| 9.56E-250 | 0.579689 | 0.983813 | 0.306601 | 4.15E-249 | pink | ATIC |
| 5.46E-248 | 0.860609 | 1 | 0.823784 | 2.34E-247 | pink | MIF |
| 2.03E-241 | 0.592211 | 0.978417 | 0.32036 | 8.59E-241 | pink | RUVBL2 |
| 1.27E-239 | 0.603193 | 0.964029 | 0.299787 | 5.31E-239 | pink | DNAJC9 |
| 1.09E-238 | 0.596271 | 1 | 0.966093 | 4.53E-238 | pink | EEF1B2 |
| 5.74E-238 | -1.6346 | 0.998201 | 0.997379 | 2.35E-237 | pink | HLA-B |
| 3.84E-236 | -2.19486 | 0.589928 | 0.913055 | 1.56E-235 | pink | VIM |
| 1.98E-235 | -1.64855 | 1 | 0.99312 | 7.93E-235 | pink | HLA-C |
| 1.12E-233 | 0.752859 | 0.989209 | 0.355053 | 4.45E-233 | pink | TUBA1C |
| 4.60E-230 | 0.824792 | 0.998201 | 0.446192 | 1.80E-229 | pink | CAT |

| | | | | | | |
|---|---|---|---|---|---|---|
| 6.59E-229 | 0.518857 | 0.785971 | 0.200164 | 2.56E-228 | pink | CKS1B |
| 5.82E-228 | 0.533975 | 0.969424 | 0.301753 | 2.24E-227 | pink | SCCPDH |
| 1.71E-225 | -1.70064 | 0.897482 | 0.947846 | 6.49E-225 | pink | HLA-E |
| 1.04E-224 | -1.52543 | 0.980216 | 0.986339 | 3.92E-224 | pink | HLA-A |
| 4.82E-223 | 0.694236 | 0.998201 | 0.425717 | 1.80E-222 | pink | TIMM13 |
| 5.23E-223 | 0.510427 | 0.98741 | 0.323079 | 1.93E-222 | pink | MPP1 |
| 1.89E-220 | 0.83669 | 0.998201 | 0.479672 | 6.91E-220 | pink | IMPDH2 |
| 1.29E-219 | 0.732559 | 0.996403 | 0.51145 | 4.67E-219 | pink | TMEM14B |
| 2.63E-218 | 0.738211 | 1 | 0.917215 | 9.42E-218 | pink | NPM1 |
| 2.55E-217 | 0.645806 | 0.991007 | 0.366028 | 9.07E-217 | pink | AHCY |
| 5.92E-214 | 0.58319 | 0.985612 | 0.336216 | 2.08E-213 | pink | HMGA1 |
| 1.13E-213 | 0.511935 | 0.973022 | 0.321278 | 3.94E-213 | pink | PAFAH1B3 |
| 2.69E-213 | 0.633695 | 0.958633 | 0.311581 | 9.29E-213 | pink | MCM7 |
| 7.52E-212 | -1.76694 | 0.305755 | 0.829877 | 2.57E-211 | pink | CORO1A |
| 6.69E-208 | 0.581535 | 0.98741 | 0.35679 | 2.27E-207 | pink | HPRT1 |
| 3.60E-207 | 0.714124 | 1 | 0.476921 | 1.21E-206 | pink | CCT6A |
| 8.02E-207 | 0.75204 | 1 | 0.709124 | 2.67E-206 | pink | RAN |
| 1.34E-205 | -2.18157 | 0.192446 | 0.797772 | 4.41E-205 | pink | CD52 |
| 2.84E-205 | 0.61302 | 0.983813 | 0.370778 | 9.29E-205 | pink | PRDX3 |
| 7.95E-205 | 0.711435 | 1 | 0.490975 | 2.58E-204 | pink | ATP5MC1 |
| 5.27E-203 | 0.734927 | 1 | 0.696314 | 1.70E-202 | pink | ATP5IF1 |
| 2.86E-202 | 0.604394 | 0.902878 | 0.291007 | 9.12E-202 | pink | MCM5 |
| 3.85E-201 | 0.621574 | 0.989209 | 0.393251 | 1.22E-200 | pink | CCT5 |
| 8.24E-201 | 0.728597 | 0.994604 | 0.472465 | 2.58E-200 | pink | NAA38 |
| 1.17E-198 | 0.55474 | 1 | 0.965635 | 3.64E-198 | pink | RPL4 |
| 7.53E-198 | -1.25908 | 1 | 0.999672 | 2.32E-197 | pink | B2M |
| 3.70E-196 | 0.754055 | 1 | 0.468239 | 1.13E-195 | pink | STMN1 |
| 9.58E-196 | 0.510393 | 0.967626 | 0.339328 | 2.90E-195 | pink | MRPL37 |
| 5.07E-195 | 0.72465 | 1 | 0.580049 | 1.52E-194 | pink | TXN |
| 3.61E-192 | 0.539682 | 1 | 0.99525 | 1.08E-191 | pink | RPLP0 |
| 7.10E-192 | 0.684096 | 0.996403 | 0.502932 | 2.10E-191 | pink | CCT2 |
| 4.03E-191 | 0.952809 | 0.998201 | 0.618116 | 1.18E-190 | pink | TUBB |
| 6.46E-189 | 0.793559 | 1 | 0.658018 | 1.88E-188 | pink | RPL22L1 |
| 1.41E-188 | 0.537169 | 0.985612 | 0.371204 | 4.07E-188 | pink | STOML2 |
| 4.05E-188 | 0.547589 | 0.985612 | 0.372219 | 1.16E-187 | pink | HDAC2 |
| 4.38E-188 | 0.51072 | 1 | 0.974087 | 1.25E-187 | pink | PPIA |
| 1.19E-184 | 0.656922 | 1 | 0.918624 | 3.38E-184 | pink | HMGB1 |
| 1.92E-182 | 0.550078 | 0.980216 | 0.360229 | 5.39E-182 | pink | RANBP1 |
| 2.46E-182 | 0.668006 | 0.998201 | 0.55181 | 6.84E-182 | pink | PA2G4 |
| 3.15E-180 | 0.710087 | 1 | 0.866601 | 8.71E-180 | pink | HSP90AB1 |
| 9.10E-178 | 0.519252 | 0.985612 | 0.388534 | 2.50E-177 | pink | TAF9 |

| | | | | | | |
|---|---|---|---|---|---|---|
| 3.22E-176 | 0.55555 | 1 | 0.957314 | 8.75E-176 | pink | HNRNPA1 |
| 3.14E-172 | 0.632925 | 0.998201 | 0.486159 | 8.48E-172 | pink | EIF3A |
| 3.77E-172 | 0.766631 | 1 | 0.607764 | 1.01E-171 | pink | H2AFZ |
| 1.02E-171 | 0.560771 | 0.98741 | 0.429025 | 2.73E-171 | pink | NDUFB6 |
| 1.63E-170 | 0.571516 | 0.994604 | 0.439214 | 4.31E-170 | pink | PHB |
| 5.45E-170 | 0.614481 | 1 | 0.651564 | 1.43E-169 | pink | SERBP1 |
| 4.23E-167 | 0.610785 | 1 | 0.694349 | 1.10E-166 | pink | ATP5F1B |
| 1.14E-162 | 0.582627 | 1 | 0.70552 | 2.97E-162 | pink | ATP5MC3 |
| 3.04E-162 | -1.82746 | 0.521583 | 0.82398 | 7.82E-162 | pink | TSC22D3 |
| 1.03E-160 | -1.49565 | 0.548561 | 0.809337 | 2.62E-160 | pink | LSP1 |
| 3.77E-159 | 0.571157 | 0.991007 | 0.390008 | 9.58E-159 | pink | CDK6 |
| 9.57E-159 | 0.559029 | 0.983813 | 0.405143 | 2.41E-158 | pink | NASP |
| 1.22E-158 | 0.50642 | 0.989209 | 0.416871 | 3.06E-158 | pink | MRPS34 |
| 1.06E-157 | 0.618318 | 0.994604 | 0.5486 | 2.63E-157 | pink | HSPE1 |
| 2.62E-157 | -1.32645 | 0.104317 | 0.693759 | 6.47E-157 | pink | PKM |
| 9.84E-157 | 0.772569 | 0.998201 | 0.656151 | 2.42E-156 | pink | TUBA1B |
| 1.19E-155 | 0.585049 | 1 | 0.538116 | 2.91E-155 | pink | SNRPD1 |
| 1.23E-155 | 0.597727 | 1 | 0.807568 | 2.98E-155 | pink | NCL |
| 2.67E-155 | -1.19399 | 0.958633 | 0.906405 | 6.42E-155 | pink | CD37 |
| 1.95E-153 | 0.53112 | 1 | 0.780835 | 4.67E-153 | pink | UQCRH |
| 1.30E-150 | 0.622852 | 1 | 0.85982 | 3.08E-150 | pink | HSP90AA1 |
| 2.48E-148 | 0.613753 | 1 | 0.607797 | 5.85E-148 | pink | YBX1 |
| 4.52E-146 | -2.0365 | 0.251799 | 0.716593 | 1.06E-145 | pink | ZFP36 |
| 5.06E-146 | 0.522781 | 0.994604 | 0.469189 | 1.18E-145 | pink | PRMT1 |
| 6.31E-145 | 0.623639 | 1 | 0.649631 | 1.46E-144 | pink | EIF5A |
| 7.46E-145 | 0.563391 | 0.992806 | 0.536937 | 1.72E-144 | pink | ILF2 |
| 5.69E-141 | 0.533525 | 0.998201 | 0.634038 | 1.30E-140 | pink | TUFM |
| 9.81E-140 | 0.518174 | 0.998201 | 0.460475 | 2.23E-139 | pink | SNRPE |
| 4.69E-139 | 0.500022 | 0.98741 | 0.455201 | 1.06E-138 | pink | CACYBP |
| 3.51E-138 | 0.543487 | 0.996403 | 0.541785 | 7.90E-138 | pink | NHP2 |
| 4.33E-138 | 0.533409 | 0.996403 | 0.62955 | 9.67E-138 | pink | COX5A |
| 9.91E-138 | -0.57819 | 1 | 0.981491 | 2.20E-137 | pink | PFDN5 |
| 1.06E-137 | 0.673654 | 0.998201 | 0.529631 | 2.34E-137 | pink | DUT |
| 2.30E-136 | 0.537999 | 1 | 0.677281 | 5.04E-136 | pink | PEBP1 |
| 2.46E-136 | 0.623025 | 0.931655 | 0.395643 | 5.37E-136 | pink | TUBB4B |
| 6.06E-136 | 0.524876 | 0.998201 | 0.509877 | 1.31E-135 | pink | SSB |
| 6.08E-136 | -1.59557 | 0.17446 | 0.666339 | 1.31E-135 | pink | HCST |
| 2.67E-134 | -0.55271 | 1 | 0.989091 | 5.73E-134 | pink | EEF1D |
| 3.07E-132 | -1.64109 | 0.070144 | 0.614251 | 6.55E-132 | pink | CXCR4 |
| 9.61E-132 | 0.521829 | 1 | 0.803997 | 2.04E-131 | pink | LDHB |
| 2.58E-131 | 0.53069 | 0.994604 | 0.475807 | 5.44E-131 | pink | GLUL |

| | | | | | | |
|---|---|---|---|---|---|---|
| 2.41E-130 | 0.553831 | 1 | 0.735233 | 5.06E-130 | pink | TPI1 |
| 4.55E-130 | -1.31074 | 0.44964 | 0.779754 | 9.49E-130 | pink | IFITM2 |
| 2.05E-126 | 0.501595 | 1 | 0.526421 | 4.24E-126 | pink | CCT8 |
| 4.69E-126 | -0.57854 | 0.998201 | 0.969435 | 9.67E-126 | pink | ARHGDIB |
| 9.47E-126 | -2.18469 | 0.510791 | 0.775987 | 1.94E-125 | pink | JUNB |
| 1.96E-121 | -0.80605 | 0.998201 | 0.999214 | 4.00E-121 | pink | MALAT1 |
| 1.03E-117 | -0.804 | 0.996403 | 0.980311 | 2.09E-117 | pink | UBC |
| 5.57E-114 | -1.35597 | 0.097122 | 0.581884 | 1.12E-113 | pink | SELL |
| 2.14E-105 | -1.07849 | 0.242806 | 0.623129 | 4.29E-105 | pink | PSMB9 |
| 3.59E-105 | -0.66476 | 0.794964 | 0.245766 | 7.16E-105 | pink | IGLL1 |
| 1.36E-104 | -1.34393 | 0.636691 | 0.750008 | 2.71E-104 | pink | PTPRC |
| 1.50E-104 | -0.5929 | 1 | 0.996822 | 2.96E-104 | pink | EIF1 |
| 3.58E-103 | -0.68025 | 1 | 0.992432 | 7.02E-103 | pink | H3F3B |
| 4.08E-100 | -0.81722 | 1 | 0.998722 | 7.96E-100 | pink | TMSB10 |
| 1.58E-99 | -1.67767 | 0.535971 | 0.781392 | 3.07E-99 | pink | JUN |
| 3.84E-98 | -0.9795 | 0.996403 | 0.904472 | 7.42E-98 | pink | SARAF |
| 1.15E-95 | -0.67978 | 0.982014 | 0.866929 | 2.21E-95 | pink | ARPC2 |
| 2.59E-92 | -0.96691 | 0.066547 | 0.505422 | 4.96E-92 | pink | ANXA6 |
| 5.30E-91 | -1.37817 | 0.379496 | 0.648911 | 1.01E-90 | pink | ITGB2 |
| 1.85E-90 | -2.01895 | 0.208633 | 0.579558 | 3.50E-90 | pink | IFITM1 |
| 4.89E-89 | -0.97298 | 0.789568 | 0.772776 | 9.21E-89 | pink | GPSM3 |
| 1.54E-88 | -1.07306 | 0.010791 | 0.455561 | 2.89E-88 | pink | CD48 |
| 1.18E-87 | -0.54434 | 0.996403 | 0.904013 | 2.20E-87 | pink | COMMD6 |
| 1.70E-87 | 0.73036 | 0.978417 | 0.588337 | 3.16E-87 | pink | HIST1H4C |
| 1.87E-85 | -1.29868 | 0.071942 | 0.490123 | 3.45E-85 | pink | CD69 |
| 3.85E-85 | -1.16901 | 0.138489 | 0.52688 | 7.07E-85 | pink | LITAF |
| 1.73E-82 | -1.23622 | 0.068345 | 0.475381 | 3.17E-82 | pink | TRBC2 |
| 2.22E-76 | -0.76766 | 0.996403 | 0.925504 | 4.03E-76 | pink | LAPTM5 |
| 3.49E-76 | -1.89403 | 0.978417 | 0.917805 | 6.32E-76 | pink | CD74 |
| 2.27E-74 | -1.85773 | 0.014388 | 0.408387 | 4.09E-74 | pink | CD3E |
| 5.11E-72 | -1.63794 | 0.017986 | 0.403374 | 9.17E-72 | pink | LTB |
| 2.77E-71 | -2.1428 | 0.01259 | 0.39502 | 4.95E-71 | pink | IL32 |
| 2.00E-69 | -1.07979 | 0.041367 | 0.412449 | 3.55E-69 | pink | SLC2A3 |
| 1.12E-68 | -1.28916 | 0.064748 | 0.428174 | 1.98E-68 | pink | TNFAIP3 |
| 1.63E-68 | -1.21311 | 0.663669 | 0.714136 | 2.87E-68 | pink | IER2 |
| 3.65E-68 | -1.20068 | 0.183453 | 0.502179 | 6.39E-68 | pink | FYB1 |
| 3.76E-68 | -0.96385 | 0.06295 | 0.426929 | 6.55E-68 | pink | BTG2 |
| 6.45E-68 | -0.87322 | 0.960432 | 0.883931 | 1.12E-67 | pink | CALM1 |
| 1.14E-67 | -1.28951 | 0.526978 | 0.667944 | 1.97E-67 | pink | KLF6 |
| 2.64E-66 | -1.22688 | 0.386691 | 0.589222 | 4.53E-66 | pink | EVL |
| 2.36E-62 | -1.64845 | 0.039568 | 0.385815 | 4.05E-62 | pink | LGALS1 |

| | | | | | | |
|---|---|---|---|---|---|---|
| 3.69E-62 | -0.85009 | 0.089928 | 0.421163 | 6.28E-62 | pink | CYTIP |
| 4.95E-62 | -0.85006 | 0.814748 | 0.727174 | 8.39E-62 | pink | LCP1 |
| 5.55E-62 | -1.04526 | 0.276978 | 0.532514 | 9.37E-62 | pink | BTG1 |
| 3.94E-58 | -1.03951 | 0.102518 | 0.417658 | 6.62E-58 | pink | DDIT4 |
| 1.53E-57 | -0.91207 | 0.133094 | 0.434627 | 2.56E-57 | pink | HLA-F |
| 1.00E-56 | -0.72252 | 0.034173 | 0.356822 | 1.67E-56 | pink | IL16 |
| 1.07E-56 | -0.65626 | 0.014388 | 0.338215 | 1.78E-56 | pink | BIN1 |
| 2.37E-55 | -0.98709 | 0.007194 | 0.325405 | 3.91E-55 | pink | LCK |
| 3.90E-55 | -0.68241 | 0.030576 | 0.347125 | 6.41E-55 | pink | GLIPR1 |
| 4.82E-55 | -1.14917 | 0.982014 | 0.824734 | 7.89E-55 | pink | SRGN |
| 1.13E-54 | -1.07547 | 0.181655 | 0.459427 | 1.85E-54 | pink | AHNAK |
| 3.11E-54 | -0.54498 | 1 | 0.972449 | 5.04E-54 | pink | ACTG1 |
| 3.76E-54 | -0.83183 | 0.023381 | 0.336118 | 6.07E-54 | pink | ISG20 |
| 4.76E-54 | -1.1251 | 0.01259 | 0.325012 | 7.66E-54 | pink | CD3D |
| 3.08E-53 | -1.41839 | 0.026978 | 0.335692 | 4.93E-53 | pink | AREG |
| 1.61E-52 | -0.74764 | 0.089928 | 0.386863 | 2.57E-52 | pink | C1orf162 |
| 6.78E-52 | -0.82686 | 0.007194 | 0.310336 | 1.08E-51 | pink | ARL4C |
| 8.16E-52 | -0.63865 | 0.008993 | 0.311646 | 1.29E-51 | pink | PDE4B |
| 2.46E-51 | -0.84462 | 0.181655 | 0.444717 | 3.87E-51 | pink | 1-Sep |
| 3.99E-51 | -2.33244 | 0.010791 | 0.310762 | 6.25E-51 | pink | NKG7 |
| 9.12E-51 | -0.90827 | 0.010791 | 0.308862 | 1.42E-50 | pink | SPOCK2 |
| 6.32E-50 | -0.94209 | 0.345324 | 0.530516 | 9.82E-50 | pink | CCND3 |
| 2.14E-49 | -0.64977 | 0.97482 | 0.803997 | 3.31E-49 | pink | GABARAP |
| 5.09E-49 | -0.64261 | 0.057554 | 0.351319 | 7.85E-49 | pink | PLEK |
| 5.58E-49 | -0.61764 | 0.071942 | 0.359083 | 8.57E-49 | pink | SORL1 |
| 2.78E-48 | -0.76308 | 0.106115 | 0.382277 | 4.25E-48 | pink | P2RY8 |
| 8.28E-48 | -0.84384 | 0.014388 | 0.299099 | 1.26E-47 | pink | FCMR |
| 1.31E-47 | -0.62285 | 0.089928 | 0.367207 | 1.98E-47 | pink | VAMP5 |
| 2.37E-47 | -1.42529 | 0.01259 | 0.295135 | 3.58E-47 | pink | IL7R |
| 9.01E-46 | -0.91673 | 0.508993 | 0.582867 | 1.36E-45 | pink | 9-Sep |
| 1.06E-45 | -0.84087 | 0.215827 | 0.446192 | 1.59E-45 | pink | VSIR |
| 2.33E-45 | -0.7031 | 0.809353 | 0.70932 | 3.48E-45 | pink | HCLS1 |
| 2.35E-45 | -1.95322 | 0.055755 | 0.330221 | 3.50E-45 | pink | TYROBP |
| 2.85E-45 | -0.60001 | 0.046763 | 0.320164 | 4.22E-45 | pink | TRIM22 |
| 1.44E-44 | -0.74727 | 0.007194 | 0.2762 | 2.12E-44 | pink | IL10RA |
| 2.72E-44 | -0.77127 | 0.01259 | 0.280786 | 3.99E-44 | pink | ZAP70 |
| 5.14E-44 | -0.6936 | 0.874101 | 0.738411 | 7.52E-44 | pink | ITM2B |
| 8.80E-44 | -1.33122 | 0.467626 | 0.551057 | 1.28E-43 | pink | AIF1 |
| 4.70E-43 | -1.02362 | 0.210432 | 0.436331 | 6.82E-43 | pink | GADD45B |
| 9.88E-43 | -0.75335 | 0.005396 | 0.265586 | 1.43E-42 | pink | HOPX |
| 1.54E-42 | -0.66379 | 0.017986 | 0.280131 | 2.22E-42 | pink | CSF3R |

| | | | | | | |
|---|---|---|---|---|---|---|
| 3.09E-42 | -1.60081 | 0.600719 | 0.608092 | 4.43E-42 | pink | S100A10 |
| 3.13E-42 | -0.90827 | 0.30036 | 0.48724 | 4.47E-42 | pink | IFITM3 |
| 2.49E-41 | -0.79081 | 0.003597 | 0.256314 | 3.54E-41 | pink | CD3G |
| 4.49E-41 | -0.82635 | 0.467626 | 0.556134 | 6.36E-41 | pink | STK17B |
| 4.70E-41 | -0.87425 | 0.005396 | 0.257035 | 6.64E-41 | pink | GZMM |
| 6.03E-41 | -1.28095 | 0.758993 | 0.650811 | 8.49E-41 | pink | HLA-DPB1 |
| 1.32E-40 | -0.83245 | 0.068345 | 0.318296 | 1.85E-40 | pink | SAMHD1 |
| 4.55E-40 | -0.64123 | 0.023381 | 0.272432 | 6.36E-40 | pink | PRKCH |
| 1.06E-39 | -0.64626 | 0.091727 | 0.337821 | 1.47E-39 | pink | RCSD1 |
| 1.33E-39 | -0.7532 | 0.017986 | 0.264308 | 1.85E-39 | pink | KLF2 |
| 1.74E-39 | -0.73724 | 0.142086 | 0.373464 | 2.41E-39 | pink | RNF213 |
| 2.09E-39 | -1.04736 | 0.138489 | 0.372416 | 2.87E-39 | pink | HLA-DQB1 |
| 5.30E-39 | -0.6155 | 0.077338 | 0.321114 | 7.27E-39 | pink | CAPG |
| 1.05E-37 | -0.75878 | 0.026978 | 0.264668 | 1.44E-37 | pink | ALOX5AP |
| 6.78E-37 | -0.73269 | 0.030576 | 0.264472 | 9.22E-37 | pink | SMIM24 |
| 4.24E-36 | -0.89598 | 0.02518 | 0.255495 | 5.75E-36 | pink | SPINK2 |
| 2.46E-35 | -0.64509 | 0.059353 | 0.286585 | 3.33E-35 | pink | CRIP1 |
| 1.18E-34 | -0.6287 | 0.998201 | 0.892449 | 1.58E-34 | pink | MYL12A |
| 1.63E-34 | -0.67843 | 0.260791 | 0.433186 | 2.18E-34 | pink | IDS |
| 2.62E-34 | -0.61967 | 0.118705 | 0.335168 | 3.50E-34 | pink | LAT2 |
| 5.92E-34 | -0.85991 | 0.464029 | 0.533071 | 7.89E-34 | pink | RARRES3 |
| 1.15E-33 | -0.73931 | 0.205036 | 0.394169 | 1.53E-33 | pink | TRAF3IP3 |
| 1.46E-33 | -0.65583 | 0.203237 | 0.393382 | 1.94E-33 | pink | CYFIP2 |
| 2.60E-32 | -0.93719 | 0.192446 | 0.383882 | 3.43E-32 | pink | LST1 |
| 5.15E-32 | -0.80742 | 0.046763 | 0.257265 | 6.77E-32 | pink | NCF1 |
| 5.39E-32 | -0.57289 | 0.134892 | 0.337592 | 7.06E-32 | pink | ARHGAP30 |
| 2.75E-31 | -0.6117 | 0.203237 | 0.390434 | 3.58E-31 | pink | TPM4 |
| 2.80E-31 | -0.70289 | 0.784173 | 0.650745 | 3.64E-31 | pink | CD44 |
| 3.15E-30 | -0.58818 | 0.055755 | 0.259165 | 4.08E-30 | pink | PTPRE |
| 7.87E-30 | -0.76677 | 0.291367 | 0.430663 | 1.02E-29 | pink | TBC1D10C |
| 2.49E-29 | -0.78951 | 0.782374 | 0.639279 | 3.21E-29 | pink | TUBA1A |
| 1.34E-28 | -0.58095 | 0.836331 | 0.660672 | 1.71E-28 | pink | ICAM3 |
| 2.19E-28 | -0.77769 | 0.106115 | 0.294775 | 2.80E-28 | pink | PIK3IP1 |
| 9.75E-28 | -0.87977 | 0.18705 | 0.362719 | 1.24E-27 | pink | NR4A2 |
| 1.06E-27 | -0.66439 | 0.26259 | 0.406257 | 1.35E-27 | pink | ADD3 |
| 1.46E-27 | -1.17057 | 0.839928 | 0.647437 | 1.85E-27 | pink | HLA-DPA1 |
| 1.99E-26 | -0.62175 | 0.577338 | 0.558755 | 2.51E-26 | pink | WAS |
| 6.64E-26 | -1.25208 | 0.18705 | 0.349746 | 8.33E-26 | pink | S100A11 |
| 7.21E-26 | -0.72967 | 0.703237 | 0.61294 | 9.03E-26 | pink | MYH9 |
| 7.65E-26 | -0.62585 | 0.241007 | 0.385291 | 9.54E-26 | pink | PPP1R18 |
| 1.26E-25 | -0.62529 | 0.303957 | 0.422277 | 1.56E-25 | pink | AKAP13 |

| | | | | | | |
|---|---|---|---|---|---|---|
| 1.36E-25 | -0.61111 | 0.158273 | 0.324947 | 1.68E-25 | pink | CYTH1 |
| 1.55E-25 | -0.78403 | 0.136691 | 0.309975 | 1.91E-25 | pink | ID2 |
| 2.79E-25 | -0.53816 | 0.106115 | 0.280491 | 3.43E-25 | pink | ANKRD44 |
| 4.07E-24 | -0.71602 | 0.516187 | 0.52475 | 5.00E-24 | pink | CYBA |
| 5.92E-24 | -0.59289 | 0.113309 | 0.28059 | 7.25E-24 | pink | MGAT4A |
| 5.11E-23 | -0.5789 | 0.519784 | 0.524324 | 6.23E-23 | pink | CELF2 |
| 6.89E-23 | -0.69409 | 0.47482 | 0.49959 | 8.37E-23 | pink | ARRB2 |
| 2.49E-22 | -0.53515 | 0.258993 | 0.379656 | 3.01E-22 | pink | RABAC1 |
| 7.92E-22 | -0.57539 | 0.149281 | 0.298706 | 9.56E-22 | pink | AKNA |
| 2.04E-21 | -0.56327 | 0.107914 | 0.262965 | 2.45E-21 | pink | MYO1F |
| 6.24E-21 | -0.66533 | 0.107914 | 0.25887 | 7.48E-21 | pink | FLT3LG |
| 8.67E-21 | -0.58562 | 0.751799 | 0.617625 | 1.04E-20 | pink | TAPBP |
| 1.16E-20 | -0.52235 | 0.992806 | 0.80688 | 1.39E-20 | pink | CD99 |
| 1.34E-20 | -0.6964 | 0.602518 | 0.551089 | 1.59E-20 | pink | ARHGEF1 |
| 1.03E-19 | -0.54255 | 0.210432 | 0.333726 | 1.22E-19 | pink | SASH3 |
| 2.27E-19 | -0.69336 | 0.388489 | 0.440655 | 2.68E-19 | pink | PIK3R1 |
| 7.48E-19 | -0.66488 | 0.600719 | 0.546765 | 8.80E-19 | pink | STK4 |
| 3.88E-18 | -0.62371 | 0.134892 | 0.268206 | 4.55E-18 | pink | LBH |
| 8.16E-18 | -0.52338 | 0.845324 | 0.64914 | 9.55E-18 | pink | KMT2E |
| 8.28E-18 | -0.82258 | 0.638489 | 0.553153 | 9.65E-18 | pink | LEPROTL1 |
| 3.55E-17 | -0.68518 | 0.638489 | 0.555643 | 4.13E-17 | pink | YPEL5 |
| 4.13E-17 | -0.78294 | 0.739209 | 0.717707 | 4.78E-17 | pink | FOS |
| 5.83E-17 | -1.28784 | 0.861511 | 0.637019 | 6.73E-17 | pink | HLA-DRB1 |
| 1.17E-16 | -1.24702 | 0.134892 | 0.259787 | 1.34E-16 | pink | FCER1G |
| 1.22E-16 | -0.565 | 0.210432 | 0.318526 | 1.40E-16 | pink | RNF125 |
| 4.80E-16 | -0.55479 | 0.821942 | 0.62588 | 5.49E-16 | pink | RNASET2 |
| 5.63E-16 | -0.57961 | 0.366906 | 0.413268 | 6.42E-16 | pink | ARHGAP9 |
| 9.95E-16 | -0.86976 | 0.956835 | 0.790663 | 1.13E-15 | pink | TXNIP |
| 1.32E-15 | -0.56516 | 0.366906 | 0.415659 | 1.50E-15 | pink | CLEC2B |
| 1.58E-15 | -0.60575 | 0.839928 | 0.636855 | 1.79E-15 | pink | CNN2 |
| 2.79E-15 | -0.61285 | 0.230216 | 0.324554 | 3.14E-15 | pink | ADGRE5 |
| 9.00E-15 | -0.80892 | 0.210432 | 0.309189 | 1.01E-14 | pink | HLA-DQA1 |
| 3.96E-13 | -0.57351 | 0.413669 | 0.424767 | 4.44E-13 | pink | VAMP2 |
| 4.93E-13 | -0.5597 | 0.361511 | 0.394169 | 5.50E-13 | pink | ARHGAP45 |
| 7.09E-13 | -0.99108 | 0.496403 | 0.46421 | 7.90E-13 | pink | CTSS |
| 1.27E-12 | -0.55023 | 0.303957 | 0.358952 | 1.41E-12 | pink | MYO1G |
| 1.56E-12 | -0.53044 | 0.26259 | 0.331663 | 1.73E-12 | pink | TMEM243 |
| 2.29E-12 | -1.12653 | 0.77518 | 0.578346 | 2.53E-12 | pink | HLA-DRB5 |
| 9.01E-12 | -0.72461 | 0.321942 | 0.360262 | 9.91E-12 | pink | ITM2C |
| 3.11E-11 | -0.78666 | 0.348921 | 0.373497 | 3.41E-11 | pink | MAP3K8 |
| 4.08E-11 | -0.65588 | 0.922662 | 0.683079 | 4.46E-11 | pink | ANXA1 |

| | | | | | | |
|---|---|---|---|---|---|---|
| 6.03E-11 | -0.69254 | 0.451439 | 0.444455 | 6.57E-11 | pink | IRF1 |
| 9.26E-11 | -0.53806 | 0.640288 | 0.533104 | 1.01E-10 | pink | RGS10 |
| 9.77E-11 | -0.61635 | 0.555755 | 0.483702 | 1.06E-10 | pink | BIN2 |
| 2.22E-10 | -0.59519 | 0.672662 | 0.528419 | 2.40E-10 | pink | CD53 |
| 6.81E-10 | -0.54196 | 0.42446 | 0.410713 | 7.34E-10 | pink | TNFRSF14 |
| 3.99E-09 | -0.50231 | 0.530576 | 0.291663 | 4.29E-09 | pink | NAMPT |
| 9.37E-09 | -0.7289 | 0.580935 | 0.480262 | 1.00E-08 | pink | MCL1 |
| 2.55E-08 | -0.70709 | 0.257194 | 0.301556 | 2.72E-08 | pink | ANXA2 |
| 2.60E-07 | -0.5937 | 0.555755 | 0.461982 | 2.77E-07 | pink | HIF1A |
| 6.98E-07 | -1.42011 | 0.895683 | 0.59679 | 7.42E-07 | pink | HLA-DRA |
| 2.05E-06 | -0.51399 | 0.604317 | 0.479476 | 2.17E-06 | pink | PPP1R2 |
| 2.61E-06 | -0.53288 | 0.280576 | 0.302932 | 2.76E-06 | pink | TAGAP |
| 2.92E-06 | -0.62136 | 0.399281 | 0.375037 | 3.08E-06 | pink | HERPUD1 |
| 5.31E-06 | -0.64258 | 0.640288 | 0.493301 | 5.58E-06 | pink | AES |
| 5.67E-06 | -0.50742 | 0.221223 | 0.255987 | 5.95E-06 | pink | ABLIM1 |
| 1.23E-05 | -0.57822 | 0.235612 | 0.26362 | 1.28E-05 | pink | TUBA4A |
| 3.01E-05 | -0.70741 | 0.73741 | 0.552596 | 3.14E-05 | pink | DNAJB1 |
| 4.63E-05 | -0.79854 | 0.478417 | 0.422146 | 4.81E-05 | pink | DUSP1 |
| 5.14E-05 | -0.50734 | 0.805755 | 0.468894 | 5.33E-05 | pink | MT2A |
| 7.36E-05 | -0.57744 | 0.498201 | 0.414775 | 7.60E-05 | pink | RIPOR2 |
| 0.000237 | -0.59309 | 1 | 0.994005 | 0.000245 | pink | FTL |
| 0.000269 | -0.57931 | 0.776978 | 0.540803 | 0.000277 | pink | EZR |
| 0.000717 | -0.53801 | 0.742806 | 0.52036 | 0.000735 | pink | SLC9A3R1 |
| 0.001229 | -0.53253 | 0.663669 | 0.486257 | 0.001256 | pink | PRMT2 |
| 0.001657 | -0.76259 | 0.843525 | 0.602719 | 0.001689 | pink | NEAT1 |
| 0.007897 | -0.50817 | 0.381295 | 0.328092 | 0.008026 | pink | EML4 |
| 0.010979 | -1.10883 | 0.543165 | 0.407469 | 0.011129 | pink | CD7 |
| 0.014623 | -2.3629 | 0.264388 | 0.250713 | 0.014782 | pink | LYZ |
| 0 | 1.770557 | 0.962417 | 0.241313 | 0 | grey60 | IL7R |
| 0 | 1.376293 | 0.443863 | 0.069568 | 0 | grey60 | GZMK |
| 0 | 1.242189 | 0.987155 | 0.344732 | 0 | grey60 | IL32 |
| 0 | 1.0236 | 0.99001 | 0.358639 | 0 | grey60 | CD3E |
| 0 | 0.89606 | 0.864891 | 0.362504 | 0 | grey60 | LTB |
| 0 | 0.875424 | 0.936727 | 0.759239 | 0 | grey60 | JUNB |
| 0 | 0.858506 | 0.860609 | 0.280168 | 0 | grey60 | CD3D |
| 0 | 0.842507 | 0.803996 | 0.284585 | 0 | grey60 | LCK |
| 0 | 0.835317 | 0.52236 | 0.125505 | 0 | grey60 | TRAC |
| 0 | 0.824512 | 0.627022 | 0.191656 | 0 | grey60 | CD2 |
| 0 | 0.795645 | 0.433397 | 0.096518 | 0 | grey60 | AQP3 |
| 0 | 0.792271 | 0.758801 | 0.397219 | 0 | grey60 | TNFAIP3 |
| 0 | 0.77493 | 0.59039 | 0.176473 | 0 | grey60 | TCF7 |

| | | | | | | |
|---|---|---|---|---|---|---|
| 0 | 0.770605 | 0.595623 | 0.194037 | 0 | grey60 | GIMAP7 |
| 0 | 0.769998 | 0.837298 | 0.471755 | 0 | grey60 | FYB1 |
| 0 | 0.752107 | 0.527593 | 0.149453 | 0 | grey60 | GIMAP4 |
| 0 | 0.729748 | 0.517602 | 0.143932 | 0 | grey60 | CD27 |
| 0 | 0.704945 | 0.689343 | 0.275544 | 0 | grey60 | SPOCK2 |
| 0 | 0.704266 | 0.671265 | 0.221367 | 0 | grey60 | CD3G |
| 0 | 0.703031 | 0.876308 | 0.564512 | 0 | grey60 | EVL |
| 0 | 0.682124 | 0.939106 | 0.783084 | 0 | grey60 | TXNIP |
| 0 | 0.680724 | 0.374405 | 0.092722 | 0 | grey60 | NELL2 |
| 0 | 0.667129 | 0.436727 | 0.116498 | 0 | grey60 | ITK |
| 0 | 0.636061 | 0.962417 | 0.54467 | 0 | grey60 | IFITM1 |
| 0 | 0.61574 | 0.457184 | 0.141965 | 0 | grey60 | CAMK4 |
| 0 | 0.595112 | 0.62607 | 0.250595 | 0 | grey60 | ZAP70 |
| 0 | 0.588324 | 0.600856 | 0.188619 | 0 | grey60 | CCL5 |
| 0 | 0.552827 | 0.977165 | 0.900963 | 0 | grey60 | SARAF |
| 0 | 0.547766 | 0.294957 | 0.060458 | 0 | grey60 | TRAT1 |
| 0 | 0.514143 | 0.273073 | 0.05535 | 0 | grey60 | CD28 |
| 0 | 0.506443 | 1 | 0.999655 | 0 | grey60 | B2M |
| 0 | -0.79697 | 0.863463 | 0.956451 | 0 | grey60 | H3F3A |
| 0 | -1.02821 | 0.052807 | 0.531281 | 0 | grey60 | HLA-DMA |
| 0 | -1.09235 | 0.463368 | 0.757238 | 0 | grey60 | GSTP1 |
| 0 | -1.28396 | 0.141294 | 0.579178 | 0 | grey60 | AIF1 |
| 0 | -1.34453 | 0.23882 | 0.680769 | 0 | grey60 | HLA-DPA1 |
| 0 | -1.78926 | 0.046147 | 0.620725 | 0 | grey60 | HLA-DRB5 |
| 0 | -1.89898 | 0.124167 | 0.678526 | 0 | grey60 | HLA-DRB1 |
| 0 | -1.96991 | 0.805899 | 0.927085 | 0 | grey60 | CD74 |
| 0 | -2.71375 | 0.039962 | 0.642914 | 0 | grey60 | HLA-DRA |
| 4.27E-307 | 0.647795 | 0.476213 | 0.163774 | 1.71E-306 | grey60 | TC2N |
| 3.61E-300 | -1.21919 | 0.262607 | 0.681045 | 1.41E-299 | grey60 | HLA-DPB1 |
| 4.62E-298 | -1.82581 | 0.009039 | 0.436903 | 1.76E-297 | grey60 | CST3 |
| 7.09E-298 | -0.98095 | 0.073264 | 0.514959 | 2.64E-297 | grey60 | GLUL |
| 1.12E-297 | -0.94214 | 0.96099 | 0.971255 | 4.08E-297 | grey60 | GAPDH |
| 9.60E-293 | -0.93325 | 0.282112 | 0.66524 | 3.42E-292 | grey60 | TKT |
| 9.33E-292 | 0.548246 | 0.964795 | 0.774043 | 3.26E-291 | grey60 | CD52 |
| 6.44E-290 | 0.655153 | 0.621313 | 0.26747 | 2.20E-289 | grey60 | PIK3IP1 |
| 2.15E-287 | 0.571859 | 0.914843 | 0.735878 | 7.20E-287 | grey60 | PTPRC |
| 7.93E-281 | -1.131 | 0.009515 | 0.420477 | 2.60E-280 | grey60 | SOX4 |
| 8.46E-279 | 0.607276 | 0.56803 | 0.233548 | 2.72E-278 | grey60 | FLT3LG |
| 3.25E-269 | 0.600386 | 0.469553 | 0.171607 | 1.03E-268 | grey60 | CD96 |
| 1.23E-267 | 0.659878 | 0.781161 | 0.538252 | 3.82E-267 | grey60 | LEPROTL1 |
| 8.39E-266 | -0.7403 | 0.477165 | 0.770523 | 2.55E-265 | grey60 | SLC25A5 |

| | | | | | | |
|---|---|---|---|---|---|---|
| 5.61E-260 | -0.84514 | 0.305423 | 0.641119 | 1.67E-259 | grey60 | TALDO1 |
| 3.24E-258 | -1.03251 | 0.275928 | 0.634045 | 9.48E-258 | grey60 | SERPINB1 |
| 2.21E-256 | -0.71343 | 0.048525 | 0.458608 | 6.37E-256 | grey60 | ZEB2 |
| 2.28E-252 | 0.554514 | 0.40961 | 0.134615 | 6.45E-252 | grey60 | LINC00861 |
| 7.36E-244 | 0.631199 | 0.558516 | 0.250561 | 2.04E-243 | grey60 | IL10RA |
| 7.50E-243 | 0.648661 | 0.405328 | 0.150764 | 2.05E-242 | grey60 | INPP4B |
| 1.55E-239 | 0.733312 | 0.701713 | 0.495014 | 4.18E-239 | grey60 | NOSIP |
| 6.98E-233 | -1.21923 | 0.00333 | 0.360054 | 1.85E-232 | grey60 | PRSS57 |
| 2.08E-231 | -1.4229 | 0.153663 | 0.50126 | 5.41E-231 | grey60 | STMN1 |
| 1.90E-229 | -0.74754 | 0.139391 | 0.524276 | 4.86E-229 | grey60 | H2AFY |
| 4.95E-229 | 0.527806 | 0.355852 | 0.114635 | 1.25E-228 | grey60 | RORA |
| 9.03E-229 | 0.584703 | 0.411989 | 0.158184 | 2.24E-228 | grey60 | LDLRAP1 |
| 3.08E-228 | -0.87921 | 0.031874 | 0.391732 | 7.55E-228 | grey60 | AC084033.3 |
| 3.74E-227 | 0.632432 | 0.766889 | 0.540598 | 9.02E-227 | grey60 | DNAJB1 |
| 4.45E-227 | 0.599234 | 0.740247 | 0.448359 | 1.06E-226 | grey60 | TRBC2 |
| 2.82E-225 | 0.566763 | 0.545671 | 0.246523 | 6.62E-225 | grey60 | LAT |
| 7.74E-223 | -0.68525 | 0.006185 | 0.354878 | 1.79E-222 | grey60 | LAT2 |
| 2.03E-216 | -0.7339 | 0.039962 | 0.395183 | 4.62E-216 | grey60 | FABP5 |
| 9.03E-214 | 0.583326 | 0.465271 | 0.207392 | 2.03E-213 | grey60 | DGKA |
| 4.27E-212 | -0.60467 | 0.007612 | 0.344698 | 9.47E-212 | grey60 | GNA15 |
| 1.75E-211 | -0.73523 | 0.293054 | 0.608924 | 3.83E-211 | grey60 | TXN |
| 2.31E-211 | -0.59832 | 0.034253 | 0.387177 | 4.98E-211 | grey60 | RNF130 |
| 9.99E-211 | -0.59429 | 0.096575 | 0.475068 | 2.13E-210 | grey60 | PGD |
| 2.05E-209 | -0.65473 | 0.089439 | 0.462852 | 4.31E-209 | grey60 | APLP2 |
| 9.92E-208 | 0.527333 | 0.431018 | 0.173332 | 2.06E-207 | grey60 | LIME1 |
| 1.50E-207 | -1.02705 | 0.045195 | 0.391663 | 3.07E-207 | grey60 | HLA-DQB1 |
| 7.68E-207 | -0.66801 | 0.001427 | 0.327996 | 1.56E-206 | grey60 | EGFL7 |
| 2.05E-206 | -0.52539 | 0.001903 | 0.328652 | 4.10E-206 | grey60 | SYK |
| 1.98E-204 | -0.58091 | 0.048525 | 0.399289 | 3.90E-204 | grey60 | DDAH2 |
| 3.99E-204 | 0.532181 | 0.314462 | 0.100694 | 7.80E-204 | grey60 | GATA3 |
| 1.80E-200 | -0.59998 | 0.001903 | 0.321233 | 3.46E-200 | grey60 | MEF2C |
| 3.82E-200 | -0.7389 | 0.132731 | 0.47952 | 7.29E-200 | grey60 | CAT |
| 9.69E-200 | -0.55149 | 0.119886 | 0.494634 | 1.83E-199 | grey60 | COMT |
| 1.80E-199 | 0.603775 | 0.620362 | 0.374133 | 3.36E-199 | grey60 | TRAF3IP3 |
| 2.08E-196 | 0.50088 | 0.573264 | 0.27375 | 3.83E-196 | grey60 | FCMR |
| 3.60E-196 | -1.88051 | 0.024738 | 0.347113 | 6.57E-196 | grey60 | TYROBP |
| 1.79E-195 | 0.545883 | 0.783539 | 0.630111 | 3.22E-195 | grey60 | CNN2 |
| 4.10E-195 | 0.689355 | 0.377735 | 0.134235 | 7.31E-195 | grey60 | CD8B |
| 6.00E-195 | 0.518345 | 0.714082 | 0.428276 | 1.06E-194 | grey60 | CD48 |
| 1.64E-194 | -0.52409 | 0.501903 | 0.76583 | 2.87E-194 | grey60 | SLC25A3 |
| 1.67E-190 | 0.511504 | 0.417697 | 0.164982 | 2.88E-190 | grey60 | SYNE2 |

| | | | | | | |
|---|---|---|---|---|---|---|
| 2.27E-190 | -0.51819 | 0.029971 | 0.357086 | 3.88E-190 | grey60 | MPP1 |
| 3.43E-190 | -0.77279 | 0.7255 | 0.865523 | 5.79E-190 | grey60 | HMGN2 |
| 5.57E-190 | 0.547244 | 0.486204 | 0.225646 | 9.33E-190 | grey60 | SKAP1 |
| 5.57E-187 | 0.512207 | 0.823977 | 0.588599 | 9.22E-187 | grey60 | CXCR4 |
| 6.79E-187 | -0.59721 | 0.001427 | 0.303875 | 1.11E-186 | grey60 | MGST1 |
| 2.27E-185 | -0.64639 | 0 | 0.299527 | 3.68E-185 | grey60 | CD34 |
| 5.75E-185 | -0.63422 | 0.002379 | 0.303013 | 9.25E-185 | grey60 | FAM30A |
| 7.08E-184 | -0.88191 | 0.019029 | 0.328341 | 1.13E-183 | grey60 | HLA-DQA1 |
| 1.24E-180 | -0.70283 | 0.000951 | 0.295352 | 1.95E-180 | grey60 | CSF3R |
| 2.83E-177 | 0.520295 | 0.408183 | 0.170537 | 4.42E-177 | grey60 | FYN |
| 7.91E-177 | -0.73556 | 0.20647 | 0.509438 | 1.22E-176 | grey60 | IMPDH2 |
| 8.28E-177 | -0.60686 | 0.152236 | 0.507505 | 1.27E-176 | grey60 | PYCARD |
| 1.59E-176 | 0.56523 | 0.647954 | 0.424928 | 2.42E-176 | grey60 | 1-Sep |
| 1.20E-175 | -0.6362 | 0.833968 | 0.896822 | 1.81E-175 | grey60 | RPS2 |
| 1.23E-172 | 0.524704 | 0.707897 | 0.466303 | 1.83E-172 | grey60 | CD69 |
| 9.28E-170 | -0.68755 | 0.071361 | 0.390421 | 1.37E-169 | grey60 | GRN |
| 1.63E-169 | -0.56823 | 0.31922 | 0.636254 | 2.39E-169 | grey60 | YBX1 |
| 7.68E-169 | -0.5733 | 0.001427 | 0.280928 | 1.12E-168 | grey60 | TCF4 |
| 3.09E-168 | -0.66868 | 0.867269 | 0.92391 | 4.45E-168 | grey60 | HMGB1 |
| 8.65E-168 | -0.69235 | 0.023311 | 0.314504 | 1.23E-167 | grey60 | CLEC11A |
| 1.61E-165 | 0.718229 | 0.294006 | 0.098106 | 2.28E-165 | grey60 | KLRB1 |
| 2.47E-165 | -0.76143 | 0.00333 | 0.278926 | 3.46E-165 | grey60 | SMIM24 |
| 2.83E-163 | -0.72577 | 0.084206 | 0.394009 | 3.94E-163 | grey60 | MAP3K8 |
| 7.45E-162 | 0.506911 | 0.418649 | 0.189586 | 1.03E-161 | grey60 | OXNAD1 |
| 1.45E-160 | -1.42202 | 0.004282 | 0.273819 | 1.98E-160 | grey60 | IGLL1 |
| 2.68E-159 | -0.82304 | 0.09039 | 0.401498 | 3.63E-159 | grey60 | LST1 |
| 3.29E-158 | 0.530307 | 0.388677 | 0.154491 | 4.43E-158 | grey60 | CD8A |
| 3.84E-157 | -0.6234 | 0.118934 | 0.421202 | 5.12E-157 | grey60 | CDK6 |
| 1.03E-156 | -1.17675 | 0.464795 | 0.676593 | 1.36E-156 | grey60 | TUBA1B |
| 1.45E-154 | 0.524056 | 0.671741 | 0.526933 | 1.90E-154 | grey60 | ACAP1 |
| 1.99E-152 | -0.57623 | 0.219315 | 0.571 | 2.59E-152 | grey60 | NPC2 |
| 1.03E-150 | -0.59607 | 0.011418 | 0.274026 | 1.33E-150 | grey60 | H1F0 |
| 3.37E-148 | -0.523 | 0.05138 | 0.336002 | 4.31E-148 | grey60 | CAPG |
| 8.66E-146 | 0.570014 | 0.489058 | 0.276994 | 1.10E-145 | grey60 | KIAA1551 |
| 6.82E-142 | 0.533798 | 0.523311 | 0.319956 | 8.61E-142 | grey60 | RBL2 |
| 4.73E-138 | 0.562387 | 0.515699 | 0.31823 | 5.93E-138 | grey60 | CDKN1B |
| 2.44E-137 | -0.53383 | 0.557088 | 0.738914 | 3.03E-137 | grey60 | ENO1 |
| 9.25E-137 | 0.505251 | 0.408658 | 0.202353 | 1.14E-136 | grey60 | PARP8 |
| 5.82E-136 | -0.62973 | 0.03901 | 0.29618 | 7.13E-136 | grey60 | BLVRB |
| 5.89E-135 | -1.25912 | 0.029496 | 0.274095 | 7.15E-135 | grey60 | FCER1G |
| 2.60E-132 | -0.52001 | 0.286394 | 0.608406 | 3.14E-132 | grey60 | ATP6V0B |

| | | | | | | |
|---|---|---|---|---|---|---|
| 2.29E-129 | -0.51517 | 0.132731 | 0.424997 | 2.74E-129 | grey60 | TUBB4B |
| 4.92E-128 | -0.85462 | 0.031874 | 0.267297 | 5.85E-128 | grey60 | SPINK2 |
| 6.90E-128 | -0.54173 | 0.090866 | 0.364574 | 8.14E-128 | grey60 | PLEK |
| 1.27E-126 | -0.7793 | 0.995243 | 0.996584 | 1.49E-126 | grey60 | FTH1 |
| 1.04E-124 | -0.50345 | 0.081351 | 0.340695 | 1.22E-124 | grey60 | MCM7 |
| 7.90E-124 | -2.51556 | 0.034729 | 0.266641 | 9.12E-124 | grey60 | LYZ |
| 6.56E-121 | 0.551891 | 0.566127 | 0.43359 | 7.52E-121 | grey60 | ARHGAP15 |
| 9.93E-121 | -0.8041 | 0.401047 | 0.630284 | 1.13E-120 | grey60 | H2AFZ |
| 7.57E-120 | -0.61955 | 0.353473 | 0.594327 | 8.56E-120 | grey60 | PRDX1 |
| 3.84E-112 | -0.54539 | 0.175071 | 0.447358 | 4.31E-112 | grey60 | FOSB |
| 3.22E-106 | 0.507846 | 0.514272 | 0.359571 | 3.59E-106 | grey60 | PBXIP1 |
| 1.35E-105 | -0.53428 | 0.845385 | 0.870699 | 1.49E-105 | grey60 | HSP90AB1 |
| 8.34E-105 | -0.57943 | 0.077545 | 0.311777 | 9.18E-105 | grey60 | NAMPT |
| 2.18E-96 | -0.54183 | 0.138915 | 0.375582 | 2.39E-96 | grey60 | ITM2C |
| 1.82E-95 | 0.641347 | 0.43863 | 0.296249 | 1.98E-95 | grey60 | LYAR |
| 9.08E-93 | -0.8586 | 0.459087 | 0.636944 | 9.79E-93 | grey60 | TUBB |
| 8.52E-87 | -1.08231 | 0.189819 | 0.393388 | 9.13E-87 | grey60 | LGALS1 |
| 1.63E-84 | -0.72476 | 0.992388 | 0.994237 | 1.74E-84 | grey60 | FTL |
| 2.22E-83 | -0.57316 | 0.528069 | 0.731875 | 2.35E-83 | grey60 | FOS |
| 2.76E-71 | -0.9765 | 0.162226 | 0.342351 | 2.90E-71 | grey60 | AREG |
| 9.25E-67 | -0.67399 | 0.260228 | 0.479623 | 9.66E-67 | grey60 | CTSS |
| 4.44E-62 | -0.68501 | 0.395338 | 0.609821 | 4.61E-62 | grey60 | HIST1H4C |
| 2.48E-55 | -0.58515 | 0.491912 | 0.699748 | 2.56E-55 | grey60 | TSPO |
| 3.28E-53 | -0.59681 | 0.404377 | 0.621726 | 3.37E-53 | grey60 | NEAT1 |
| 2.46E-52 | -0.58827 | 0.486679 | 0.708202 | 2.51E-52 | grey60 | PSAP |
| 2.74E-51 | -0.84867 | 0.429591 | 0.637117 | 2.77E-51 | grey60 | SAT1 |
| 4.72E-11 | -0.64222 | 0.886299 | 0.819248 | 4.74E-11 | grey60 | S100A6 |
| 0 | 2.418617 | 0.87037 | 0.062723 | 0 | magenta | DNTT |
| 0 | 2.216036 | 0.995885 | 0.243831 | 0 | magenta | IGLL1 |
| 0 | 2.076435 | 0.95679 | 0.161628 | 0 | magenta | UHRF1 |
| 0 | 2.021737 | 1 | 0.469456 | 0 | magenta | STMN1 |
| 0 | 2.007693 | 0.993827 | 0.062592 | 0 | magenta | VPREB1 |
| 0 | 1.840944 | 0.897119 | 0.056022 | 0 | magenta | CD24 |
| 0 | 1.711525 | 0.971193 | 0.072757 | 0 | magenta | VPREB3 |
| 0 | 1.443577 | 0.936214 | 0.313417 | 0 | magenta | MCM7 |
| 0 | 1.36784 | 0.761317 | 0.110116 | 0 | magenta | NUSAP1 |
| 0 | 1.344803 | 0.895062 | 0.227488 | 0 | magenta | PCNA |
| 0 | 1.344576 | 0.971193 | 0.18284 | 0 | magenta | CD79B |
| 0 | 1.302493 | 0.621399 | 0.086354 | 0 | magenta | TOP2A |
| 0 | 1.277464 | 0.884774 | 0.22569 | 0 | magenta | SMC4 |
| 0 | 1.196855 | 0.728395 | 0.090211 | 0 | magenta | MKI67 |

| | | | | | | |
|---|---|---|---|---|---|---|
| 0 | 1.121895 | 0.769547 | 0.040268 | 0 | magenta | IGHV5-78 |
| 0 | 1.11133 | 0.932099 | 0.269325 | 0 | magenta | CARHSP1 |
| 0 | 1.075501 | 0.518519 | 0.063638 | 0 | magenta | HIST1H3B |
| 0 | 1.0735 | 0.829218 | 0.13901 | 0 | magenta | DHFR |
| 0 | 1.023214 | 0.860082 | 0.184703 | 0 | magenta | CENPU |
| 0 | 1.019743 | 0.748971 | 0.142311 | 0 | magenta | GINS2 |
| 0 | 0.969782 | 0.73251 | 0.086387 | 0 | magenta | PKMYT1 |
| 0 | 0.965454 | 0.94856 | 0.144697 | 0 | magenta | CD79A |
| 0 | 0.961199 | 0.77572 | 0.034679 | 0 | magenta | EBF1 |
| 0 | 0.932596 | 0.895062 | 0.184377 | 0 | magenta | IGHM |
| 0 | 0.929732 | 0.623457 | 0.081157 | 0 | magenta | CD9 |
| 0 | 0.893185 | 0.648148 | 0.085373 | 0 | magenta | BIRC5 |
| 0 | 0.892413 | 0.713992 | 0.107338 | 0 | magenta | TK1 |
| 0 | 0.887092 | 0.520576 | 0.017617 | 0 | magenta | CMTM8 |
| 0 | 0.882957 | 0.736626 | 0.127014 | 0 | magenta | CENPM |
| 0 | 0.880948 | 0.792181 | 0.128322 | 0 | magenta | ZWINT |
| 0 | 0.877015 | 0.67284 | 0.088642 | 0 | magenta | ASF1B |
| 0 | 0.84324 | 0.450617 | 0.010328 | 0 | magenta | NPY |
| 0 | 0.837564 | 0.748971 | 0.051283 | 0 | magenta | POU2AF1 |
| 0 | 0.827944 | 0.687243 | 0.109201 | 0 | magenta | CLSPN |
| 0 | 0.827286 | 0.765432 | 0.08119 | 0 | magenta | BLNK |
| 0 | 0.807205 | 0.641975 | 0.079131 | 0 | magenta | CDCA5 |
| 0 | 0.782061 | 0.58642 | 0.066122 | 0 | magenta | KIFC1 |
| 0 | 0.752272 | 0.522634 | 0.015395 | 0 | magenta | AKAP12 |
| 0 | 0.750612 | 0.662551 | 0.028109 | 0 | magenta | MME |
| 0 | 0.695338 | 0.683128 | 0.098905 | 0 | magenta | RAD51AP1 |
| 0 | 0.693417 | 0.633745 | 0.085373 | 0 | magenta | CDC45 |
| 0 | 0.678184 | 0.49177 | 0.057885 | 0 | magenta | AURKB |
| 0 | 0.674867 | 0.559671 | 0.061415 | 0 | magenta | CCNA2 |
| 0 | 0.668548 | 0.668724 | 0.034156 | 0 | magenta | PAX5 |
| 0 | 0.661616 | 0.541152 | 0.020886 | 0 | magenta | E2F2 |
| 0 | 0.655407 | 0.506173 | 0.042589 | 0 | magenta | RRM2 |
| 0 | 0.651288 | 0.553498 | 0.035169 | 0 | magenta | FAM111B |
| 0 | 0.646292 | 0.384774 | 0.017617 | 0 | magenta | LINC01013 |
| 0 | 0.642236 | 0.590535 | 0.083706 | 0 | magenta | CDC6 |
| 0 | 0.632488 | 0.516461 | 0.067037 | 0 | magenta | CDK1 |
| 0 | 0.602104 | 0.440329 | 0.017813 | 0 | magenta | RAG2 |
| 0 | 0.599561 | 0.547325 | 0.068312 | 0 | magenta | NCAPH |
| 0 | 0.594355 | 0.487654 | 0.048635 | 0 | magenta | CKAP2L |
| 0 | 0.577497 | 0.502058 | 0.062265 | 0 | magenta | FBXO5 |
| 0 | 0.570442 | 0.522634 | 0.063834 | 0 | magenta | KNL1 |

| | | | | | | |
|---|---|---|---|---|---|---|
| 0 | 0.566041 | 0.430041 | 0.025429 | 0 | magenta | ARPP21 |
| 0 | 0.562418 | 0.502058 | 0.056284 | 0 | magenta | NCAPG |
| 0 | 0.526967 | 0.425926 | 0.035921 | 0 | magenta | CCNE2 |
| 0 | 0.504868 | 0.557613 | 0.053963 | 0 | magenta | CD19 |
| 4.82E-307 | 1.064634 | 0.864198 | 0.2252 | 3.63E-306 | magenta | CDCA7 |
| 1.28E-306 | 0.849429 | 0.800412 | 0.186893 | 9.51E-306 | magenta | CENPW |
| 2.63E-305 | 0.932289 | 0.769547 | 0.16859 | 1.92E-304 | magenta | PHGDH |
| 2.08E-304 | 0.664349 | 0.701646 | 0.125838 | 1.49E-303 | magenta | GNG7 |
| 2.35E-303 | 1.129225 | 0.903292 | 0.276385 | 1.66E-302 | magenta | MAD2L2 |
| 1.60E-302 | 0.64642 | 0.654321 | 0.117078 | 1.11E-301 | magenta | CENPN |
| 1.69E-301 | 1.789463 | 1 | 0.91881 | 1.16E-300 | magenta | HMGB1 |
| 3.26E-301 | 1.070549 | 0.911523 | 0.251708 | 2.20E-300 | magenta | TCF4 |
| 5.18E-301 | 1.167118 | 1 | 0.999216 | 3.44E-300 | magenta | PTMA |
| 1.08E-300 | 0.60866 | 0.580247 | 0.092303 | 7.05E-300 | magenta | UBE2T |
| 7.49E-300 | 0.783821 | 0.868313 | 0.190848 | 4.84E-299 | magenta | MZB1 |
| 1.11E-298 | 0.786999 | 0.716049 | 0.143422 | 7.05E-298 | magenta | ATAD2 |
| 1.83E-296 | 0.839353 | 0.82716 | 0.198856 | 1.15E-295 | magenta | JUP |
| 5.81E-296 | 0.787342 | 0.788066 | 0.156137 | 3.60E-295 | magenta | PAG1 |
| 2.56E-292 | 0.543686 | 0.54321 | 0.082301 | 1.57E-291 | magenta | TCF19 |
| 1.46E-290 | 0.578421 | 0.460905 | 0.059356 | 8.78E-290 | magenta | CDCA8 |
| 5.58E-289 | 1.014436 | 0.567901 | 0.089426 | 3.32E-288 | magenta | UBE2C |
| 1.04E-286 | 0.90843 | 0.878601 | 0.239974 | 6.11E-286 | magenta | ERGIC1 |
| 5.87E-285 | 0.672676 | 0.697531 | 0.140252 | 3.40E-284 | magenta | HELLS |
| 1.36E-283 | 0.528902 | 0.487654 | 0.068018 | 7.76E-283 | magenta | RACGAP1 |
| 4.33E-283 | 1.114166 | 0.917695 | 0.29665 | 2.45E-282 | magenta | USP1 |
| 6.17E-283 | 0.569864 | 0.545267 | 0.086779 | 3.44E-282 | magenta | DTL |
| 3.66E-278 | 1.353101 | 0.981481 | 0.49057 | 2.02E-277 | magenta | H2AFY |
| 6.19E-278 | 2.030732 | 0.995885 | 0.65697 | 3.37E-277 | magenta | TUBA1B |
| 8.09E-278 | 0.572802 | 0.604938 | 0.107534 | 4.35E-277 | magenta | CTNNAL1 |
| 2.43E-276 | 0.605464 | 0.617284 | 0.113548 | 1.29E-275 | magenta | TIMELESS |
| 1.93E-275 | 1.955207 | 0.997942 | 0.61899 | 1.01E-274 | magenta | TUBB |
| 2.60E-275 | 0.792358 | 0.788066 | 0.18771 | 1.35E-274 | magenta | NRM |
| 1.37E-274 | 1.665707 | 1 | 0.853767 | 7.02E-274 | magenta | HMGN2 |
| 1.22E-272 | 0.756326 | 0.773663 | 0.185455 | 6.18E-272 | magenta | MAD2L1 |
| 5.40E-272 | 0.774617 | 0.730453 | 0.16555 | 2.71E-271 | magenta | MCM4 |
| 1.97E-268 | 1.592461 | 0.99177 | 0.608792 | 9.79E-268 | magenta | H2AFZ |
| 3.32E-267 | 0.805595 | 0.753086 | 0.17882 | 1.63E-266 | magenta | SMC2 |
| 7.13E-267 | 0.662754 | 0.687243 | 0.145906 | 3.47E-266 | magenta | GMNN |
| 1.06E-265 | 0.841724 | 0.792181 | 0.204543 | 5.08E-265 | magenta | RRM1 |
| 1.55E-265 | 1.068939 | 0.936214 | 0.321687 | 7.39E-265 | magenta | LAT2 |
| 2.41E-265 | 0.520902 | 0.561728 | 0.095342 | 1.13E-264 | magenta | WDR76 |

| | | | | | | |
|---|---|---|---|---|---|---|
| 1.33E-263 | 0.552589 | 0.567901 | 0.098317 | 6.18E-263 | magenta | MND1 |
| 7.69E-262 | 1.618579 | 1 | 0.677137 | 3.55E-261 | magenta | HMGB2 |
| 1.13E-260 | 1.02752 | 0.54321 | 0.092401 | 5.18E-260 | magenta | HIST1H1B |
| 1.23E-259 | 0.837041 | 0.87037 | 0.254453 | 5.55E-259 | magenta | CDCA7L |
| 5.08E-258 | 1.159492 | 0.954733 | 0.406929 | 2.27E-257 | magenta | NASP |
| 6.45E-258 | 0.877323 | 0.563786 | 0.101585 | 2.86E-257 | magenta | TPX2 |
| 8.78E-258 | 0.818504 | 0.730453 | 0.173787 | 3.85E-257 | magenta | FEN1 |
| 4.06E-257 | 1.055744 | 0.843621 | 0.256218 | 1.76E-256 | magenta | MCM3 |
| 1.14E-256 | 0.890576 | 0.997942 | 0.974179 | 4.92E-256 | magenta | PPIA |
| 2.55E-256 | 0.943751 | 0.858025 | 0.24808 | 1.09E-255 | magenta | TOP2B |
| 2.89E-255 | 0.980587 | 0.915638 | 0.30116 | 1.22E-254 | magenta | SSBP2 |
| 1.06E-253 | 0.744136 | 0.769547 | 0.191502 | 4.43E-253 | magenta | MCM2 |
| 2.05E-252 | 0.585019 | 0.534979 | 0.089361 | 8.51E-252 | magenta | CDKN3 |
| 3.66E-242 | 0.601456 | 0.631687 | 0.134009 | 1.50E-241 | magenta | RFC5 |
| 5.41E-242 | 0.993663 | 0.967078 | 0.445138 | 2.20E-241 | magenta | PARP1 |
| 4.17E-241 | 0.706794 | 0.742798 | 0.185651 | 1.68E-240 | magenta | LMNB1 |
| 2.21E-240 | 1.122768 | 1 | 0.970093 | 8.86E-240 | magenta | GAPDH |
| 2.76E-240 | 0.582768 | 0.604938 | 0.123027 | 1.09E-239 | magenta | EZH2 |
| 9.25E-238 | 0.589559 | 0.399177 | 0.053277 | 3.63E-237 | magenta | ASPM |
| 5.35E-236 | 0.92663 | 0.995885 | 0.913221 | 2.08E-235 | magenta | HNRNPA2B1 |
| 6.72E-236 | 0.922723 | 0.76749 | 0.210296 | 2.60E-235 | magenta | H2AFX |
| 1.10E-235 | 0.709043 | 0.534979 | 0.096127 | 4.22E-235 | magenta | CCNB2 |
| 1.65E-235 | 0.534784 | 0.54321 | 0.100474 | 6.25E-235 | magenta | FANCI |
| 8.84E-235 | 0.669165 | 0.722222 | 0.18065 | 3.33E-234 | magenta | CENPH |
| 5.75E-234 | -1.61048 | 0.94856 | 0.994836 | 2.15E-233 | magenta | FTL |
| 6.17E-234 | 0.511064 | 0.532922 | 0.095048 | 2.29E-233 | magenta | BARD1 |
| 9.23E-234 | 0.601907 | 0.415638 | 0.05831 | 3.39E-233 | magenta | HIST1H2AL |
| 1.58E-232 | 1.111351 | 0.635802 | 0.139075 | 5.75E-232 | magenta | CENPF |
| 1.95E-231 | 0.562697 | 0.588477 | 0.118124 | 7.06E-231 | magenta | ATAD5 |
| 1.04E-229 | 0.821944 | 0.436214 | 0.065207 | 3.75E-229 | magenta | CDC20 |
| 5.13E-227 | 1.021946 | 0.755144 | 0.206733 | 1.83E-226 | magenta | PTTG1 |
| 1.55E-226 | 0.600437 | 0.615226 | 0.130315 | 5.46E-226 | magenta | PCLAF |
| 4.45E-223 | 0.573502 | 0.619342 | 0.136885 | 1.56E-222 | magenta | RBBP8 |
| 6.34E-222 | 0.650645 | 0.734568 | 0.194901 | 2.20E-221 | magenta | RFC2 |
| 6.38E-222 | 0.607784 | 0.368313 | 0.048243 | 2.20E-221 | magenta | HIST1H3C |
| 2.58E-221 | 0.845144 | 0.938272 | 0.323811 | 8.82E-221 | magenta | RCSD1 |
| 8.55E-220 | 0.74976 | 0.82716 | 0.25782 | 2.91E-219 | magenta | SAE1 |
| 1.09E-219 | 0.902996 | 0.997942 | 0.972545 | 3.69E-219 | magenta | ACTG1 |
| 1.93E-218 | 0.970491 | 0.989712 | 0.746887 | 6.47E-218 | magenta | SLC25A5 |
| 8.77E-218 | 0.506117 | 0.576132 | 0.120314 | 2.91E-217 | magenta | CDK2 |
| 1.03E-217 | 0.574582 | 0.617284 | 0.128354 | 3.41E-217 | magenta | QRSL1 |

| | | | | | | |
|---|---|---|---|---|---|---|
| 1.19E-213 | 0.804642 | 0.769547 | 0.223435 | 3.90E-213 | magenta | TMEM106C |
| 3.36E-210 | 1.204228 | 0.973251 | 0.5311 | 1.09E-209 | magenta | DUT |
| 8.78E-209 | 0.736626 | 0.849794 | 0.279948 | 2.83E-208 | magenta | SMARCA4 |
| 9.41E-208 | 0.870102 | 0.866255 | 0.30286 | 3.02E-207 | magenta | DNAJC9 |
| 5.25E-204 | 0.897785 | 0.868313 | 0.311685 | 1.67E-203 | magenta | TMPO |
| 3.17E-202 | 0.703128 | 0.742798 | 0.20219 | 1.00E-201 | magenta | CKS1B |
| 7.91E-202 | 0.587679 | 0.73251 | 0.198954 | 2.48E-201 | magenta | DTYMK |
| 9.58E-202 | 0.901623 | 0.685185 | 0.191404 | 2.99E-201 | magenta | SOCS2 |
| 2.62E-201 | 0.701055 | 0.711934 | 0.193692 | 8.10E-201 | magenta | LIG1 |
| 1.46E-200 | 0.547653 | 0.639918 | 0.153816 | 4.49E-200 | magenta | WDR34 |
| 1.67E-200 | -0.65514 | 0.995885 | 0.999281 | 5.10E-200 | magenta | RPL34 |
| 9.61E-199 | 1.120452 | 0.969136 | 0.571825 | 2.92E-198 | magenta | PRDX1 |
| 1.84E-197 | 0.825504 | 0.923868 | 0.365158 | 5.56E-197 | magenta | H2AFV |
| 1.22E-195 | 0.746444 | 1 | 0.949371 | 3.65E-195 | magenta | H3F3A |
| 3.82E-194 | 0.854701 | 0.971193 | 0.563654 | 1.14E-193 | magenta | YWHAQ |
| 7.78E-194 | 0.89289 | 0.930041 | 0.426442 | 2.30E-193 | magenta | PSMA6 |
| 3.61E-193 | 0.587285 | 0.613169 | 0.146821 | 1.06E-192 | magenta | CDCA4 |
| 1.02E-191 | 0.923565 | 0.977366 | 0.562118 | 2.97E-191 | magenta | SRSF10 |
| 1.19E-191 | 0.92493 | 0.835391 | 0.293479 | 3.44E-191 | magenta | MCM5 |
| 2.60E-191 | 0.837848 | 0.977366 | 0.707076 | 7.47E-191 | magenta | SRSF3 |
| 1.17E-190 | 0.820259 | 0.76749 | 0.244386 | 3.34E-190 | magenta | CKS2 |
| 3.84E-189 | 0.920364 | 0.975309 | 0.710181 | 1.09E-188 | magenta | RAN |
| 2.16E-185 | 0.599052 | 0.707819 | 0.193169 | 6.11E-185 | magenta | ADA |
| 1.08E-184 | 0.57811 | 0.306584 | 0.039614 | 3.03E-184 | magenta | HIST1H2AI |
| 8.76E-184 | 0.678974 | 0.695473 | 0.200981 | 2.45E-183 | magenta | MCM6 |
| 9.40E-183 | -1.67588 | 0.162551 | 0.814872 | 2.61E-182 | magenta | LSP1 |
| 1.70E-181 | 0.897834 | 0.950617 | 0.486027 | 4.69E-181 | magenta | DEK |
| 2.08E-180 | 0.626345 | 0.763374 | 0.230103 | 5.69E-180 | magenta | CD38 |
| 5.04E-179 | 0.765515 | 0.878601 | 0.32757 | 1.37E-178 | magenta | GLRX |
| 4.09E-176 | 0.58859 | 0.726337 | 0.21899 | 1.11E-175 | magenta | CBX5 |
| 7.12E-176 | -2.44846 | 0.298354 | 0.832129 | 1.92E-175 | magenta | S100A6 |
| 1.02E-173 | 0.559212 | 0.695473 | 0.202026 | 2.74E-173 | magenta | YEATS4 |
| 4.18E-173 | -0.55262 | 1 | 0.999346 | 1.11E-172 | magenta | RPS14 |
| 5.02E-173 | 0.798096 | 0.934156 | 0.458604 | 1.33E-172 | magenta | DDX39A |
| 2.49E-172 | 0.533789 | 0.728395 | 0.20049 | 6.55E-172 | magenta | TSPAN14 |
| 3.02E-172 | 0.7714 | 0.923868 | 0.40183 | 7.89E-172 | magenta | RNASEH2B |
| 2.69E-171 | 0.85911 | 0.942387 | 0.524203 | 6.98E-171 | magenta | SIVA1 |
| 3.18E-171 | 0.737925 | 0.835391 | 0.317013 | 8.22E-171 | magenta | RPA3 |
| 2.50E-170 | 0.732885 | 0.872428 | 0.369603 | 6.42E-170 | magenta | SSRP1 |
| 5.10E-170 | 0.763697 | 0.792181 | 0.295702 | 1.30E-169 | magenta | PAICS |
| 2.54E-169 | -0.56749 | 0.997942 | 0.999575 | 6.44E-169 | magenta | RPL30 |

| | | | | | | |
|---|---|---|---|---|---|---|
| 1.63E-168 | 0.821711 | 0.960905 | 0.512012 | 4.10E-168 | magenta | RBBP4 |
| 2.01E-168 | 0.727551 | 0.726337 | 0.229319 | 5.05E-168 | magenta | HMGB3 |
| 3.09E-168 | 0.829825 | 0.973251 | 0.678608 | 7.71E-168 | magenta | PKM |
| 3.44E-168 | 0.741199 | 0.711934 | 0.220951 | 8.55E-168 | magenta | HPS4 |
| 4.21E-168 | 0.819789 | 0.997942 | 0.860173 | 1.04E-167 | magenta | HSP90AA1 |
| 3.86E-167 | 0.762239 | 0.814815 | 0.291453 | 9.47E-167 | magenta | MEF2C |
| 6.46E-167 | 0.629049 | 0.786008 | 0.255336 | 1.58E-166 | magenta | DCK |
| 5.39E-164 | 0.774481 | 0.981481 | 0.727635 | 1.31E-163 | magenta | HNRNPA3 |
| 1.62E-162 | -1.36223 | 0.520576 | 0.913483 | 3.92E-162 | magenta | CD37 |
| 2.04E-161 | -2.41839 | 0.360082 | 0.818892 | 4.91E-161 | magenta | S100A4 |
| 1.35E-160 | -0.74021 | 0.975309 | 0.981925 | 3.23E-160 | magenta | PFDN5 |
| 9.93E-160 | 0.649203 | 0.995885 | 0.838797 | 2.36E-159 | magenta | SUMO2 |
| 1.47E-159 | 0.50697 | 0.646091 | 0.186043 | 3.47E-159 | magenta | RNASEH2A |
| 5.28E-159 | 0.744731 | 0.967078 | 0.625887 | 1.24E-158 | magenta | ERH |
| 3.27E-158 | 0.788744 | 0.853909 | 0.361726 | 7.67E-158 | magenta | DNMT1 |
| 1.99E-157 | -1.90252 | 0.469136 | 0.833241 | 4.64E-157 | magenta | SRGN |
| 1.01E-156 | 0.837676 | 0.927984 | 0.469619 | 2.33E-156 | magenta | VDAC1 |
| 2.07E-155 | 2.022531 | 0.94856 | 0.589704 | 4.78E-155 | magenta | HIST1H4C |
| 6.81E-155 | 0.68181 | 0.839506 | 0.323484 | 1.56E-154 | magenta | C19orf48 |
| 9.83E-154 | 0.853178 | 0.971193 | 0.614741 | 2.24E-153 | magenta | SRSF2 |
| 9.26E-152 | 0.608125 | 0.773663 | 0.280046 | 2.10E-151 | magenta | HADH |
| 3.35E-150 | 0.604495 | 0.757202 | 0.261481 | 7.56E-150 | magenta | MAGED1 |
| 4.91E-150 | 0.794666 | 0.841564 | 0.36385 | 1.10E-149 | magenta | RANBP1 |
| 1.12E-149 | -0.85056 | 0.993827 | 0.994607 | 2.51E-149 | magenta | RPS27 |
| 1.63E-147 | -0.55467 | 1 | 0.998987 | 3.62E-147 | magenta | RPL13 |
| 3.61E-147 | 0.615612 | 0.987654 | 0.830789 | 8.01E-147 | magenta | HNRNPK |
| 8.67E-147 | 0.654434 | 0.987654 | 0.744174 | 1.91E-146 | magenta | SLC25A3 |
| 4.78E-145 | 0.681142 | 0.841564 | 0.336656 | 1.05E-144 | magenta | MIS18BP1 |
| 6.82E-144 | -1.52426 | 0.18107 | 0.740382 | 1.49E-143 | magenta | EMP3 |
| 1.09E-143 | 0.526001 | 0.722222 | 0.241052 | 2.38E-143 | magenta | ARL6IP6 |
| 2.77E-143 | 0.649154 | 0.835391 | 0.336199 | 5.99E-143 | magenta | PRKDC |
| 4.62E-143 | 0.632556 | 0.839506 | 0.349142 | 9.95E-143 | magenta | COPS3 |
| 4.48E-142 | 0.644783 | 0.755144 | 0.272855 | 9.60E-142 | magenta | SYNGR1 |
| 7.36E-142 | 0.699275 | 0.740741 | 0.263899 | 1.57E-141 | magenta | KIF22 |
| 1.29E-141 | 0.694736 | 0.944444 | 0.509822 | 2.74E-141 | magenta | HNRNPR |
| 2.96E-141 | 0.616454 | 0.798354 | 0.30505 | 6.26E-141 | magenta | SMC3 |
| 3.05E-141 | -0.57834 | 0.99177 | 0.993855 | 6.41E-141 | magenta | RPL36 |
| 1.47E-140 | 0.832859 | 0.942387 | 0.552966 | 3.08E-140 | magenta | HNRNPF |
| 7.35E-140 | 0.55416 | 0.666667 | 0.22033 | 1.53E-139 | magenta | MTHFD1 |
| 2.88E-139 | 0.595578 | 0.777778 | 0.288054 | 5.97E-139 | magenta | MSI2 |
| 3.53E-138 | 0.615057 | 0.831276 | 0.345677 | 7.29E-138 | magenta | SMS |

| | | | | | | |
|---|---|---|---|---|---|---|
| 9.91E-137 | 0.525042 | 0.771605 | 0.28302 | 2.04E-136 | magenta | MYL6B |
| 2.17E-136 | 0.714884 | 0.995885 | 0.808073 | 4.44E-136 | magenta | NCL |
| 3.21E-136 | -1.33439 | 0.993827 | 0.996535 | 6.55E-136 | magenta | FTH1 |
| 5.85E-136 | 0.816144 | 0.950617 | 0.566727 | 1.19E-135 | magenta | NUCKS1 |
| 8.92E-136 | 0.664499 | 0.946502 | 0.570093 | 1.80E-135 | magenta | SRP9 |
| 4.29E-135 | 0.681345 | 0.942387 | 0.538781 | 8.61E-135 | magenta | ILF2 |
| 1.80E-134 | 0.707666 | 0.886831 | 0.428501 | 3.61E-134 | magenta | IDH2 |
| 2.06E-134 | 0.54922 | 0.786008 | 0.286877 | 4.10E-134 | magenta | ZCCHC7 |
| 2.24E-134 | -1.67353 | 0.135802 | 0.696127 | 4.43E-134 | magenta | ANXA1 |
| 2.66E-134 | 0.541977 | 1 | 0.957411 | 5.25E-134 | magenta | HNRNPA1 |
| 8.64E-133 | -1.22678 | 0.034979 | 0.650106 | 1.70E-132 | magenta | CNN2 |
| 1.48E-132 | 0.744612 | 0.932099 | 0.553881 | 2.89E-132 | magenta | PA2G4 |
| 1.53E-132 | 0.584847 | 0.763374 | 0.283739 | 2.99E-132 | magenta | SMC1A |
| 1.11E-131 | 0.518126 | 0.572016 | 0.171564 | 2.16E-131 | magenta | MSH6 |
| 5.52E-131 | 0.549625 | 0.534979 | 0.151953 | 1.07E-130 | magenta | ERG |
| 5.62E-131 | 0.701225 | 0.796296 | 0.325674 | 1.08E-130 | magenta | ANP32E |
| 1.40E-129 | -1.61016 | 0.092593 | 0.666514 | 2.67E-129 | magenta | HCST |
| 2.90E-128 | 0.63141 | 0.917695 | 0.457526 | 5.53E-128 | magenta | CACYBP |
| 7.73E-128 | 0.576407 | 0.796296 | 0.324596 | 1.47E-127 | magenta | HSPB11 |
| 4.60E-126 | 0.542688 | 0.705761 | 0.251839 | 8.70E-126 | magenta | BCL11A |
| 1.08E-125 | 0.618959 | 0.919753 | 0.474064 | 2.03E-125 | magenta | CBX3 |
| 2.11E-125 | 0.529686 | 0.989712 | 0.905083 | 3.96E-125 | magenta | RHOA |
| 2.79E-125 | 0.590029 | 0.919753 | 0.443373 | 5.21E-125 | magenta | RAB11A |
| 3.09E-125 | 0.712645 | 0.683128 | 0.238176 | 5.76E-125 | magenta | SMIM3 |
| 2.51E-124 | 0.748845 | 0.845679 | 0.363622 | 4.65E-124 | magenta | FABP5 |
| 3.81E-124 | 0.511547 | 0.847737 | 0.32221 | 7.04E-124 | magenta | TMEM243 |
| 3.99E-123 | 0.689192 | 0.965021 | 0.672299 | 7.34E-123 | magenta | PPP1CA |
| 1.34E-122 | 0.631574 | 0.975309 | 0.698023 | 2.45E-122 | magenta | NAP1L1 |
| 4.10E-122 | -0.70407 | 0.958848 | 0.948717 | 7.46E-122 | magenta | FXYD5 |
| 4.23E-121 | 0.616182 | 0.884774 | 0.421638 | 7.68E-121 | magenta | LSM4 |
| 7.08E-121 | 0.586179 | 0.880658 | 0.418827 | 1.28E-120 | magenta | SMARCB1 |
| 1.37E-119 | 0.505524 | 0.985597 | 0.886746 | 2.47E-119 | magenta | CHCHD2 |
| 1.89E-119 | 0.582771 | 0.853909 | 0.375715 | 3.40E-119 | magenta | HDAC2 |
| 2.00E-119 | -0.63788 | 0.995885 | 0.994542 | 3.57E-119 | magenta | MT-ND3 |
| 2.13E-118 | 0.611367 | 0.950617 | 0.604118 | 3.79E-118 | magenta | UBE2I |
| 4.36E-118 | 0.619788 | 0.911523 | 0.454846 | 7.73E-118 | magenta | YWHAE |
| 1.74E-117 | 0.592666 | 0.87037 | 0.404184 | 3.07E-117 | magenta | MRPL51 |
| 2.93E-117 | 0.503756 | 0.773663 | 0.300801 | 5.14E-117 | magenta | TPGS2 |
| 4.42E-117 | 0.560789 | 0.81893 | 0.327276 | 7.74E-117 | magenta | MPP1 |
| 1.37E-116 | 0.728336 | 0.958848 | 0.710639 | 2.39E-116 | magenta | LDHA |
| 5.29E-116 | 0.527179 | 0.740741 | 0.287335 | 9.20E-116 | magenta | SUPT16H |

| | | | | | | |
|---|---|---|---|---|---|---|
| 5.41E-116 | 0.602284 | 0.866255 | 0.41242 | 9.36E-116 | magenta | SNX2 |
| 5.11E-115 | 0.89093 | 0.91358 | 0.521066 | 8.81E-115 | magenta | CCND3 |
| 5.92E-115 | 0.526608 | 0.662551 | 0.230757 | 1.02E-114 | magenta | FAM111A |
| 9.51E-115 | 0.61417 | 0.839506 | 0.396993 | 1.63E-114 | magenta | CCT5 |
| 1.07E-114 | 1.001268 | 0.814815 | 0.398725 | 1.83E-114 | magenta | TUBB4B |
| 1.44E-114 | 0.588311 | 0.884774 | 0.417585 | 2.44E-114 | magenta | ESD |
| 1.55E-113 | 0.623182 | 0.944444 | 0.565746 | 2.63E-113 | magenta | NONO |
| 2.11E-113 | 0.573884 | 0.658436 | 0.223599 | 3.56E-113 | magenta | SPTBN1 |
| 7.11E-113 | 0.589546 | 0.944444 | 0.602255 | 1.20E-112 | magenta | SNRPB |
| 8.46E-113 | 0.553837 | 0.851852 | 0.374277 | 1.42E-112 | magenta | PRDX3 |
| 2.35E-112 | 0.511415 | 0.995885 | 0.595882 | 3.92E-112 | magenta | HLA-DRA |
| 2.31E-111 | -0.6948 | 0.911523 | 0.939761 | 3.84E-111 | magenta | RPL36AL |
| 3.65E-111 | 0.603904 | 0.977366 | 0.73767 | 6.04E-111 | magenta | HNRNPA0 |
| 5.86E-111 | 0.611978 | 0.930041 | 0.540284 | 9.66E-111 | magenta | SNRPD1 |
| 6.72E-111 | 0.583295 | 0.969136 | 0.706684 | 1.10E-110 | magenta | ATP5MC3 |
| 9.63E-111 | 0.595709 | 0.965021 | 0.637326 | 1.58E-110 | magenta | HNRNPM |
| 4.45E-109 | 0.604217 | 0.907407 | 0.494558 | 7.26E-109 | magenta | KPNB1 |
| 4.10E-108 | 0.502619 | 0.641975 | 0.226508 | 6.67E-108 | magenta | FHIT |
| 4.38E-107 | -1.51157 | 0.032922 | 0.572283 | 7.10E-107 | magenta | CTSW |
| 1.56E-106 | 0.518267 | 0.895062 | 0.43579 | 2.51E-106 | magenta | PSMB2 |
| 1.81E-106 | 0.5041 | 0.769547 | 0.319464 | 2.92E-106 | magenta | POLD2 |
| 4.22E-106 | 0.517957 | 0.644033 | 0.237196 | 6.77E-106 | magenta | CKAP2 |
| 4.74E-106 | 0.639266 | 0.911523 | 0.506684 | 7.57E-106 | magenta | RAD21 |
| 7.66E-106 | 0.579424 | 0.604938 | 0.215231 | 1.22E-105 | magenta | KPNA2 |
| 1.22E-105 | 0.62308 | 0.983539 | 0.763295 | 1.93E-105 | magenta | FUS |
| 1.86E-105 | 0.536519 | 0.792181 | 0.353881 | 2.93E-105 | magenta | POLE3 |
| 2.09E-105 | 0.553607 | 0.823045 | 0.369472 | 3.29E-105 | magenta | PFKL |
| 3.66E-105 | 0.574267 | 0.874486 | 0.44151 | 5.75E-105 | magenta | PTBP1 |
| 4.29E-104 | 0.570711 | 0.839506 | 0.375454 | 6.70E-104 | magenta | PPP1R18 |
| 6.21E-102 | 0.557476 | 0.95679 | 0.600882 | 9.67E-102 | magenta | PSMB8 |
| 1.14E-101 | 0.564597 | 0.930041 | 0.503121 | 1.77E-101 | magenta | SNRPA |
| 1.40E-101 | 0.590659 | 0.907407 | 0.505475 | 2.17E-101 | magenta | CCT2 |
| 6.33E-101 | 0.551482 | 0.954733 | 0.593823 | 9.77E-101 | magenta | EIF4H |
| 7.41E-101 | 0.536953 | 0.761317 | 0.335284 | 1.14E-100 | magenta | CENPX |
| 1.36E-100 | 0.580061 | 0.606996 | 0.217846 | 2.08E-100 | magenta | PDLIM1 |
| 8.90E-100 | -0.9766 | 0.962963 | 0.986599 | 1.36E-99 | magenta | HLA-A |
| 1.13E-99 | 0.551459 | 0.923868 | 0.545056 | 1.72E-99 | magenta | XRCC5 |
| 7.46E-99 | 0.51305 | 0.804527 | 0.361137 | 1.13E-98 | magenta | HPRT1 |
| 2.17E-98 | 0.596301 | 0.969136 | 0.640987 | 3.29E-98 | magenta | HMGN1 |
| 2.38E-98 | 0.571535 | 0.569959 | 0.191665 | 3.58E-98 | magenta | MT1X |
| 4.97E-98 | -1.07584 | 0.911523 | 0.947508 | 7.46E-98 | magenta | HLA-E |

| | | | | | | |
|---|---|---|---|---|---|---|
| 1.28E-97 | 0.547473 | 0.975309 | 0.69544 | 1.92E-97 | magenta | ATP5F1B |
| 1.45E-96 | -0.96527 | 0.273663 | 0.670011 | 2.16E-96 | magenta | ICAM3 |
| 2.16E-96 | 0.542863 | 0.95679 | 0.683085 | 3.22E-96 | magenta | PTGES3 |
| 2.71E-96 | 0.537359 | 0.95679 | 0.604903 | 4.01E-96 | magenta | EWSR1 |
| 5.33E-95 | 0.556501 | 0.967078 | 0.641772 | 7.87E-95 | magenta | HNRNPU |
| 5.95E-95 | 0.531113 | 0.930041 | 0.55679 | 8.76E-95 | magenta | SNRPD3 |
| 2.38E-94 | 0.561024 | 0.701646 | 0.285014 | 3.49E-94 | magenta | HIST1H1C |
| 3.29E-94 | 0.509673 | 0.860082 | 0.405753 | 4.82E-94 | magenta | ZNF22 |
| 3.37E-94 | 0.563613 | 0.936214 | 0.617552 | 4.92E-94 | magenta | GDI2 |
| 4.41E-94 | 0.500526 | 0.855967 | 0.424089 | 6.42E-94 | magenta | PIN1 |
| 1.04E-92 | 0.517863 | 0.806584 | 0.389835 | 1.50E-92 | magenta | RBBP7 |
| 1.51E-92 | 0.552939 | 0.921811 | 0.558948 | 2.19E-92 | magenta | ANP32A |
| 1.54E-92 | 0.53246 | 0.954733 | 0.575617 | 2.22E-92 | magenta | 9-Sep |
| 1.72E-92 | 0.504092 | 0.954733 | 0.655728 | 2.46E-92 | magenta | XRCC6 |
| 1.26E-91 | -0.99764 | 0.269547 | 0.659225 | 1.80E-91 | magenta | CD44 |
| 1.77E-91 | 0.502113 | 0.909465 | 0.536395 | 2.52E-91 | magenta | PGAM1 |
| 6.39E-91 | -1.36856 | 0.259259 | 0.650204 | 9.08E-91 | magenta | ITGB2 |
| 3.70E-90 | -0.68597 | 0.91358 | 0.92891 | 5.24E-90 | magenta | TOMM7 |
| 4.08E-90 | 0.664547 | 0.736626 | 0.329858 | 5.76E-90 | magenta | KLF10 |
| 9.32E-90 | -0.99706 | 0.91358 | 0.905998 | 1.31E-89 | magenta | SARAF |
| 1.24E-89 | 0.505473 | 0.962963 | 0.72355 | 1.74E-89 | magenta | COX8A |
| 3.51E-89 | 0.503668 | 0.835391 | 0.434221 | 4.91E-89 | magenta | NUTF2 |
| 3.58E-89 | 0.50121 | 0.932099 | 0.554404 | 5.00E-89 | magenta | SNRPF |
| 3.65E-89 | 0.535595 | 0.919753 | 0.543357 | 5.08E-89 | magenta | RALY |
| 1.27E-88 | 0.527031 | 0.915638 | 0.528714 | 1.76E-88 | magenta | BANF1 |
| 4.33E-88 | -1.1599 | 0.434156 | 0.752966 | 5.98E-88 | magenta | PTPRC |
| 4.97E-88 | 0.548533 | 0.954733 | 0.653081 | 6.85E-88 | magenta | SERBP1 |
| 5.14E-88 | 0.592856 | 0.987654 | 0.858081 | 7.07E-88 | magenta | HSPA8 |
| 1.04E-87 | -1.01838 | 0.985597 | 0.993365 | 1.43E-87 | magenta | HLA-C |
| 4.60E-87 | 0.514479 | 0.792181 | 0.387743 | 6.28E-87 | magenta | SYNCRIP |
| 2.01E-86 | -0.92948 | 0.014403 | 0.492466 | 2.73E-86 | magenta | BIN2 |
| 4.11E-85 | 0.681127 | 0.633745 | 0.262821 | 5.58E-85 | magenta | HIST1H1D |
| 8.92E-84 | 0.512709 | 0.907407 | 0.528975 | 1.21E-83 | magenta | CCT8 |
| 1.19E-83 | 0.516734 | 0.897119 | 0.536101 | 1.61E-83 | magenta | SNRPG |
| 1.37E-83 | -0.91233 | 0.995885 | 0.997418 | 1.84E-83 | magenta | HLA-B |
| 1.99E-83 | 0.511563 | 0.866255 | 0.480242 | 2.67E-83 | magenta | CCT6A |
| 1.58E-81 | 0.550272 | 0.950617 | 0.678804 | 2.11E-81 | magenta | PEBP1 |
| 3.71E-81 | 0.545722 | 0.81893 | 0.424154 | 4.94E-81 | magenta | BUB3 |
| 3.44E-79 | -1.23303 | 0.061728 | 0.503383 | 4.57E-79 | magenta | FYB1 |
| 4.21E-78 | 0.681295 | 0.872428 | 0.524694 | 5.58E-78 | magenta | HSPD1 |
| 5.00E-78 | 0.512834 | 0.878601 | 0.460141 | 6.61E-78 | magenta | PDIA6 |

| | | | | | | |
|---|---|---|---|---|---|---|
| 3.44E-77 | 0.570629 | 0.989712 | 0.925772 | 4.54E-77 | magenta | LAPTM5 |
| 8.39E-76 | -1.15973 | 0.024691 | 0.461285 | 1.10E-75 | magenta | AHNAK |
| 4.13E-69 | -0.55958 | 0.886831 | 0.905867 | 5.41E-69 | magenta | PPDPF |
| 6.03E-67 | -1.44765 | 0.014403 | 0.416179 | 7.87E-67 | magenta | CD7 |
| 1.78E-66 | -0.52603 | 0.930041 | 0.905279 | 2.32E-66 | magenta | COMMD6 |
| 1.06E-65 | -1.53096 | 0.555556 | 0.774669 | 1.38E-65 | magenta | JUNB |
| 1.84E-64 | 0.577616 | 0.946502 | 0.736297 | 2.38E-64 | magenta | CALM2 |
| 4.59E-64 | -2.14371 | 0.004115 | 0.39428 | 5.93E-64 | magenta | IL32 |
| 8.12E-63 | -1.87224 | 0.283951 | 0.577513 | 1.05E-62 | magenta | IFITM1 |
| 5.67E-61 | -0.92985 | 0.679012 | 0.775355 | 7.28E-61 | magenta | IFITM2 |
| 2.31E-60 | -1.84264 | 0.039095 | 0.407093 | 2.96E-60 | magenta | CD3E |
| 4.95E-60 | -0.78278 | 0.631687 | 0.775323 | 6.32E-60 | magenta | GPSM3 |
| 9.63E-60 | -0.75555 | 1 | 0.999673 | 1.23E-59 | magenta | B2M |
| 3.30E-59 | -1.54974 | 0.343621 | 0.612159 | 4.19E-59 | magenta | S100A10 |
| 2.32E-57 | -0.98359 | 0.22428 | 0.542899 | 2.94E-57 | magenta | FLNA |
| 1.36E-55 | -1.10816 | 0.065844 | 0.415362 | 1.72E-55 | magenta | TIMP1 |
| 7.94E-55 | -0.84626 | 0.106996 | 0.447393 | 1.00E-54 | magenta | VSIR |
| 6.56E-52 | -1.15942 | 0.676955 | 0.795489 | 8.24E-52 | magenta | TXNIP |
| 2.50E-51 | -1.34869 | 0.020576 | 0.352018 | 3.14E-51 | magenta | S100A11 |
| 7.67E-51 | -1.71878 | 0.08642 | 0.413074 | 9.58E-51 | magenta | CST3 |
| 2.24E-50 | -1.60256 | 0.057613 | 0.384736 | 2.79E-50 | magenta | LGALS1 |
| 7.27E-50 | -0.67883 | 0.00823 | 0.333453 | 9.03E-50 | magenta | SELPLG |
| 1.69E-49 | -1.12951 | 0.002058 | 0.324465 | 2.10E-49 | magenta | CD3D |
| 3.69E-48 | -1.97008 | 0.012346 | 0.330283 | 4.55E-48 | magenta | TYROBP |
| 2.78E-46 | -0.65431 | 0.199588 | 0.493447 | 3.43E-46 | magenta | STAT3 |
| 3.67E-46 | -0.93722 | 0.557613 | 0.687727 | 4.50E-46 | magenta | TSPO |
| 8.84E-46 | -0.60303 | 0.685185 | 0.74254 | 1.08E-45 | magenta | GSTK1 |
| 5.27E-44 | -1.29242 | 0.613169 | 0.709789 | 6.44E-44 | magenta | ZFP36 |
| 9.83E-44 | -0.99473 | 0.12963 | 0.410198 | 1.20E-43 | magenta | SLC2A3 |
| 3.29E-41 | -0.65227 | 0.693416 | 0.741592 | 4.00E-41 | magenta | ITM2B |
| 1.76E-40 | -0.6701 | 0.308642 | 0.54035 | 2.13E-40 | magenta | ACAP1 |
| 2.44E-40 | -0.88657 | 0.030864 | 0.307861 | 2.95E-40 | magenta | SPOCK2 |
| 1.20E-39 | -0.89184 | 0.218107 | 0.45125 | 1.44E-39 | magenta | CD48 |
| 3.20E-38 | -0.56116 | 0.020576 | 0.287596 | 3.85E-38 | magenta | MGST1 |
| 5.92E-38 | -0.69223 | 0.00823 | 0.270861 | 7.10E-38 | magenta | LAT |
| 5.21E-37 | -0.61496 | 0.061728 | 0.333845 | 6.23E-37 | magenta | FCGRT |
| 1.06E-36 | -0.55922 | 0.13786 | 0.398366 | 1.27E-36 | magenta | NDFIP1 |
| 1.38E-36 | -0.79047 | 0.002058 | 0.255761 | 1.64E-36 | magenta | CD3G |
| 1.74E-36 | -0.65212 | 0.020576 | 0.27949 | 2.07E-36 | magenta | CSF3R |
| 2.46E-36 | -0.89081 | 0.631687 | 0.694198 | 2.91E-36 | magenta | PSAP |
| 2.73E-36 | -0.87378 | 0.004115 | 0.25648 | 3.22E-36 | magenta | GZMM |

| | | | | | | |
|---|---|---|---|---|---|---|
| 3.81E-36 | -0.72625 | 0.020576 | 0.275372 | 4.48E-36 | magenta | IL10RA |
| 6.56E-36 | -0.67334 | 0.131687 | 0.385521 | 7.71E-36 | magenta | C1orf162 |
| 8.58E-36 | -1.08021 | 0.728395 | 0.777774 | 1.00E-35 | magenta | JUN |
| 1.53E-35 | -0.77247 | 0.055556 | 0.308874 | 1.79E-35 | magenta | ARL4C |
| 2.43E-35 | -0.65585 | 0.347737 | 0.537996 | 2.84E-35 | magenta | RGS10 |
| 4.15E-35 | -0.72617 | 0.100823 | 0.349959 | 4.82E-35 | magenta | PLEK |
| 2.24E-34 | -1.29756 | 0.016461 | 0.261383 | 2.59E-34 | magenta | FCER1G |
| 4.63E-34 | -0.81022 | 0.076132 | 0.317601 | 5.35E-34 | magenta | SAMHD1 |
| 1.21E-33 | -0.65165 | 0.372428 | 0.525282 | 1.39E-33 | magenta | HSPB1 |
| 1.85E-32 | -0.60173 | 0.921811 | 0.893904 | 2.13E-32 | magenta | MYL12A |
| 2.27E-32 | -0.83345 | 0.399177 | 0.55715 | 2.60E-32 | magenta | LEPROTL1 |
| 5.40E-32 | -0.74446 | 0.059671 | 0.295081 | 6.18E-32 | magenta | PIK3IP1 |
| 1.25E-31 | -0.58205 | 0.030864 | 0.263834 | 1.42E-31 | magenta | MYO1F |
| 3.14E-31 | -0.58085 | 0.50823 | 0.631312 | 3.58E-31 | magenta | RNASET2 |
| 4.46E-31 | -0.55482 | 0.13786 | 0.36689 | 5.07E-31 | magenta | RNF130 |
| 7.57E-31 | -1.52141 | 0.072016 | 0.300311 | 8.57E-31 | magenta | CST7 |
| 2.28E-30 | -0.61045 | 0.228395 | 0.433306 | 2.58E-30 | magenta | IDS |
| 2.99E-30 | -1.2162 | 0.617284 | 0.719693 | 3.37E-30 | magenta | FOS |
| 3.36E-30 | -0.56924 | 0.1893 | 0.407191 | 3.78E-30 | magenta | ITM2A |
| 9.15E-30 | -0.57402 | 0.403292 | 0.542703 | 1.03E-29 | magenta | SYNGR2 |
| 5.90E-29 | -0.77357 | 0.306584 | 0.48485 | 6.60E-29 | magenta | MCL1 |
| 6.84E-29 | -0.72 | 0.047325 | 0.264324 | 7.63E-29 | magenta | HOPX |
| 1.20E-28 | -0.56562 | 0.450617 | 0.579931 | 1.33E-28 | magenta | ATP6V0E1 |
| 1.56E-28 | -0.82006 | 0.144033 | 0.363 | 1.74E-28 | magenta | NR4A2 |
| 1.87E-28 | -0.88955 | 0.294239 | 0.470861 | 2.07E-28 | magenta | TRBC2 |
| 2.16E-28 | -0.51384 | 0.851852 | 0.838765 | 2.38E-28 | magenta | LIMD2 |
| 7.13E-28 | -2.50101 | 0.041152 | 0.25429 | 7.86E-28 | magenta | LYZ |
| 1.49E-26 | -0.66112 | 0.201646 | 0.39379 | 1.64E-26 | magenta | TRAF3IP3 |
| 1.05E-25 | -0.68221 | 0.479424 | 0.574702 | 1.15E-25 | magenta | SELL |
| 4.30E-25 | -0.52045 | 0.209877 | 0.409283 | 4.70E-25 | magenta | LCP2 |
| 2.60E-24 | -0.54901 | 0.154321 | 0.344435 | 2.84E-24 | magenta | GLIPR1 |
| 4.61E-24 | -0.63265 | 0.08642 | 0.281843 | 5.03E-24 | magenta | BLVRB |
| 4.86E-24 | -0.68497 | 0.403292 | 0.533878 | 5.28E-24 | magenta | RARRES3 |
| 4.56E-23 | -0.87238 | 0.154321 | 0.32234 | 4.94E-23 | magenta | LCK |
| 1.81E-21 | 0.559413 | 0.695473 | 0.472528 | 1.96E-21 | magenta | HIST1H1E |
| 2.00E-21 | -0.54665 | 0.337449 | 0.47243 | 2.16E-21 | magenta | CD63 |
| 9.55E-21 | -0.59047 | 0.462963 | 0.549077 | 1.03E-20 | magenta | STK4 |
| 1.47E-19 | -0.79586 | 0.685185 | 0.608629 | 1.57E-19 | magenta | SERPINB1 |
| 7.40E-19 | -0.622 | 0.100823 | 0.258637 | 7.93E-19 | magenta | FLT3LG |
| 4.30E-18 | -0.6518 | 0.294239 | 0.431084 | 4.59E-18 | magenta | FOSB |
| 5.84E-18 | -0.69436 | 0.23251 | 0.37529 | 6.23E-18 | magenta | MAP3K8 |

| | | | | | | |
|---|---|---|---|---|---|---|
| 8.91E-18 | -0.98533 | 0.211934 | 0.337898 | 9.48E-18 | magenta | PRSS57 |
| 1.18E-17 | -0.91669 | 0.36214 | 0.466416 | 1.25E-17 | magenta | CTSS |
| 1.24E-17 | -0.6565 | 0.22428 | 0.371139 | 1.31E-17 | magenta | GRN |
| 4.86E-17 | -0.91753 | 0.320988 | 0.423272 | 5.13E-17 | magenta | TNFAIP3 |
| 5.53E-17 | -0.56479 | 0.333333 | 0.439255 | 5.83E-17 | magenta | APLP2 |
| 8.42E-17 | -0.53404 | 0.434156 | 0.525772 | 8.85E-17 | magenta | SLC9A3R1 |
| 1.70E-16 | -0.51438 | 0.36214 | 0.466318 | 1.79E-16 | magenta | ASAH1 |
| 3.87E-15 | -0.63036 | 0.131687 | 0.261938 | 4.05E-15 | magenta | KLF2 |
| 1.16E-14 | -0.51569 | 0.524691 | 0.555025 | 1.21E-14 | magenta | STK17B |
| 1.92E-14 | -0.6187 | 0.353909 | 0.43043 | 2.00E-14 | magenta | HLA-F |
| 4.58E-14 | -0.51297 | 0.979424 | 0.968688 | 4.75E-14 | magenta | SH3BGRL3 |
| 3.28E-13 | -0.66904 | 0.895062 | 0.785226 | 3.40E-13 | magenta | CD52 |
| 1.08E-12 | -0.70266 | 0.407407 | 0.475993 | 1.12E-12 | magenta | MT2A |
| 3.96E-12 | -0.80786 | 0.347737 | 0.42435 | 4.09E-12 | magenta | DUSP1 |
| 3.06E-11 | -2.15899 | 0.213992 | 0.306848 | 3.15E-11 | magenta | NKG7 |
| 3.72E-11 | -0.51351 | 0.355967 | 0.416179 | 3.82E-11 | magenta | CYTIP |
| 5.11E-11 | -1.04457 | 0.658436 | 0.62252 | 5.24E-11 | magenta | SAT1 |
| 8.70E-11 | -0.9451 | 0.23251 | 0.331721 | 8.90E-11 | magenta | AREG |
| 4.01E-08 | -0.64007 | 0.666667 | 0.584311 | 4.09E-08 | magenta | EVL |
| 1.24E-07 | -0.71999 | 0.644033 | 0.606439 | 1.27E-07 | magenta | NEAT1 |
| 1.08E-06 | -0.64312 | 0.584362 | 0.55545 | 1.09E-06 | magenta | DNAJB1 |
| 1.09E-06 | -0.51326 | 0.37037 | 0.412028 | 1.10E-06 | magenta | NFKBIA |
| 1.97E-05 | -0.53303 | 0.220165 | 0.261056 | 2.00E-05 | magenta | ALOX5AP |
| 6.91E-05 | -0.68405 | 0.693416 | 0.547279 | 6.97E-05 | magenta | AIF1 |
| 6.92E-05 | -0.52072 | 0.226337 | 0.260827 | 6.97E-05 | magenta | SMIM24 |
| 9.94E-05 | -0.59295 | 0.27572 | 0.30116 | 9.99E-05 | magenta | ANXA2 |
| 0.021478 | -0.62903 | 0.259259 | 0.25125 | 0.021525 | magenta | SPINK2 |
| 0 | 2.320061 | 0.98737 | 0.129039 | 0 | turquoise | CCL5 |
| 0 | 2.204233 | 0.701926 | 0.049545 | 0 | turquoise | GZMH |
| 0 | 1.985344 | 0.986422 | 0.228129 | 0 | turquoise | NKG7 |
| 0 | 1.782964 | 0.700347 | 0.110196 | 0 | turquoise | CD8A |
| 0 | 1.752967 | 0.975371 | 0.219746 | 0 | turquoise | CST7 |
| 0 | 1.693613 | 0.90401 | 0.101598 | 0 | turquoise | GZMA |
| 0 | 1.643559 | 0.616356 | 0.056996 | 0 | turquoise | CCL4 |
| 0 | 1.631938 | 0.568361 | 0.074515 | 0 | turquoise | GNLY |
| 0 | 1.567066 | 0.972845 | 0.321846 | 0 | turquoise | IL32 |
| 0 | 1.544832 | 0.55731 | 0.056244 | 0 | turquoise | FGFBP2 |
| 0 | 1.539687 | 0.443638 | 0.055313 | 0 | turquoise | GZMK |
| 0 | 1.368495 | 0.587938 | 0.101096 | 0 | turquoise | CD8B |
| 0 | 1.256639 | 0.918219 | 0.523644 | 0 | turquoise | CTSW |
| 0 | 1.194844 | 0.792232 | 0.191302 | 0 | turquoise | GZMM |

| | | | | | | |
|---|---|---|---|---|---|---|
| 0 | 1.178867 | 0.973161 | 0.336462 | 0 | turquoise | CD3E |
| 0 | 1.151292 | 0.485317 | 0.059003 | 0 | turquoise | KLRD1 |
| 0 | 1.126506 | 0.520051 | 0.103926 | 0 | turquoise | KLRG1 |
| 0 | 1.09386 | 0.826334 | 0.261912 | 0 | turquoise | CD3D |
| 0 | 1.036055 | 0.941585 | 0.625313 | 0 | turquoise | HCST |
| 0 | 1.016365 | 0.603094 | 0.08125 | 0 | turquoise | PRF1 |
| 0 | 0.998119 | 0.666877 | 0.2047 | 0 | turquoise | CD3G |
| 0 | 0.98493 | 1 | 0.992477 | 0 | turquoise | HLA-C |
| 0 | 0.980357 | 0.562046 | 0.229204 | 0 | turquoise | TUBA4A |
| 0 | 0.949057 | 0.473003 | 0.065988 | 0 | turquoise | GZMB |
| 0 | 0.943202 | 1 | 0.999642 | 0 | turquoise | B2M |
| 0 | 0.932265 | 0.574676 | 0.180984 | 0 | turquoise | CD2 |
| 0 | 0.921598 | 0.81623 | 0.486315 | 0 | turquoise | LITAF |
| 0 | 0.905669 | 0.56994 | 0.275919 | 0 | turquoise | LYAR |
| 0 | 0.904112 | 0.395011 | 0.033675 | 0 | turquoise | FCRL6 |
| 0 | 0.88607 | 0.999368 | 0.984739 | 0 | turquoise | HLA-A |
| 0 | 0.884689 | 1 | 0.997098 | 0 | turquoise | HLA-B |
| 0 | 0.862816 | 0.501105 | 0.150928 | 0 | turquoise | FYN |
| 0 | 0.860805 | 0.513735 | 0.144444 | 0 | turquoise | SYNE2 |
| 0 | 0.853421 | 0.323019 | 0.08734 | 0 | turquoise | KLRB1 |
| 0 | 0.83704 | 0.966846 | 0.52823 | 0 | turquoise | IFITM1 |
| 0 | 0.836897 | 0.710452 | 0.275382 | 0 | turquoise | LCK |
| 0 | 0.833196 | 0.976634 | 0.874937 | 0 | turquoise | CALM1 |
| 0 | 0.824191 | 0.922008 | 0.728237 | 0 | turquoise | PTPRC |
| 0 | 0.820689 | 0.754026 | 0.506627 | 0 | turquoise | RARRES3 |
| 0 | 0.819788 | 0.744869 | 0.436698 | 0 | turquoise | TRBC2 |
| 0 | 0.794251 | 0.45532 | 0.126496 | 0 | turquoise | SH2D2A |
| 0 | 0.793652 | 0.406063 | 0.070753 | 0 | turquoise | SAMD3 |
| 0 | 0.786754 | 0.877487 | 0.617611 | 0 | turquoise | ITGB2 |
| 0 | 0.786212 | 0.526997 | 0.230744 | 0 | turquoise | HOPX |
| 0 | 0.778319 | 0.532049 | 0.259225 | 0 | turquoise | PPP2R5C |
| 0 | 0.776732 | 0.716451 | 0.388228 | 0 | turquoise | TNFAIP3 |
| 0 | 0.772848 | 0.616988 | 0.267966 | 0 | turquoise | SPOCK2 |
| 0 | 0.757109 | 0.83928 | 0.577882 | 0 | turquoise | CXCR4 |
| 0 | 0.749786 | 0.696558 | 0.42699 | 0 | turquoise | AHNAK |
| 0 | 0.745042 | 0.993369 | 0.941678 | 0 | turquoise | HLA-E |
| 0 | 0.736672 | 0.598358 | 0.239414 | 0 | turquoise | ZAP70 |
| 0 | 0.736127 | 0.454373 | 0.179731 | 0 | turquoise | C12orf75 |
| 0 | 0.723138 | 0.402273 | 0.138354 | 0 | turquoise | TRBC1 |
| 0 | 0.715663 | 0.518787 | 0.239521 | 0 | turquoise | PRKCH |
| 0 | 0.713253 | 0.877171 | 0.689117 | 0 | turquoise | ZFP36 |

| | | | | | | |
|---|---|---|---|---|---|---|
| 0 | 0.707043 | 0.323019 | 0.052089 | 0 | turquoise | TGFBR3 |
| 0 | 0.696872 | 0.909694 | 0.773017 | 0 | turquoise | CD52 |
| 0 | 0.692366 | 0.942848 | 0.888837 | 0 | turquoise | MYL12A |
| 0 | 0.688474 | 0.938428 | 0.795658 | 0 | turquoise | CD99 |
| 0 | 0.685593 | 0.54468 | 0.235222 | 0 | turquoise | LAT |
| 0 | 0.679566 | 0.324913 | 0.052733 | 0 | turquoise | PYHIN1 |
| 0 | 0.67941 | 0.52605 | 0.242495 | 0 | turquoise | IL10RA |
| 0 | 0.679023 | 0.415535 | 0.16515 | 0 | turquoise | ITGAL |
| 0 | 0.677866 | 0.984212 | 0.967113 | 0 | turquoise | SH3BGRL3 |
| 0 | 0.676069 | 0.415851 | 0.154725 | 0 | turquoise | SYTL3 |
| 0 | 0.657507 | 0.32365 | 0.101992 | 0 | turquoise | GBP5 |
| 0 | 0.628879 | 0.380486 | 0.12646 | 0 | turquoise | TRAC |
| 0 | 0.621067 | 1 | 0.992978 | 0 | turquoise | TMSB4X |
| 0 | 0.599409 | 0.55131 | 0.204449 | 0 | turquoise | CD247 |
| 0 | 0.585921 | 0.335965 | 0.107688 | 0 | turquoise | RORA |
| 0 | 0.572055 | 0.359331 | 0.091245 | 0 | turquoise | IL2RB |
| 0 | 0.554638 | 0.2545 | 0.032528 | 0 | turquoise | S1PR5 |
| 0 | 0.503743 | 0.994001 | 0.978111 | 0 | turquoise | PFN1 |
| 0 | -0.52731 | 0.912536 | 0.977395 | 0 | turquoise | RPL7 |
| 0 | -0.52819 | 0.011051 | 0.350613 | 0 | turquoise | ARMH1 |
| 0 | -0.5431 | 0.900221 | 0.955829 | 0 | turquoise | H3F3A |
| 0 | -0.60545 | 0.997474 | 0.995092 | 0 | turquoise | RPLP0 |
| 0 | -0.61846 | 0.000947 | 0.333524 | 0 | turquoise | MEF2C |
| 0 | -0.62533 | 0.004421 | 0.357921 | 0 | turquoise | GNA15 |
| 0 | -0.64175 | 0.374171 | 0.739593 | 0 | turquoise | NAP1L1 |
| 0 | -0.64395 | 0.016104 | 0.410296 | 0 | turquoise | BEX3 |
| 0 | -0.66759 | 0.209978 | 0.614423 | 0 | turquoise | SELL |
| 0 | -0.68004 | 0.002842 | 0.340295 | 0 | turquoise | EGFL7 |
| 0 | -0.685 | 0.883486 | 0.922691 | 0 | turquoise | NPM1 |
| 0 | -0.7373 | 0.63467 | 0.827112 | 0 | turquoise | LDHB |
| 0 | -0.73831 | 0.118724 | 0.49434 | 0 | turquoise | CAT |
| 0 | -0.7513 | 0.131039 | 0.539908 | 0 | turquoise | H2AFY |
| 0 | -0.76375 | 0.017998 | 0.500716 | 0 | turquoise | SPINT2 |
| 0 | -0.87514 | 0.265235 | 0.658487 | 0 | turquoise | TALDO1 |
| 0 | -0.88372 | 0.128829 | 0.529806 | 0 | turquoise | IMPDH2 |
| 0 | -0.88404 | 0.032207 | 0.405424 | 0 | turquoise | AC084033.3 |
| 0 | -0.95944 | 0.04168 | 0.418894 | 0 | turquoise | LST1 |
| 0 | -0.97824 | 0.233344 | 0.685391 | 0 | turquoise | TKT |
| 0 | -1.16032 | 0.00821 | 0.436304 | 0 | turquoise | SOX4 |
| 0 | -1.21986 | 0.009788 | 0.372931 | 0 | turquoise | PRSS57 |
| 0 | -1.32805 | 0.178087 | 0.51175 | 0 | turquoise | STMN1 |

| | | | | | | |
|---|---|---|---|---|---|---|
| 0 | -1.59063 | 0.037891 | 0.607616 | 0 | turquoise | AIF1 |
| 0 | -1.72763 | 0.049574 | 0.448628 | 0 | turquoise | CST3 |
| 0 | -2.02275 | 0.224187 | 0.645017 | 0 | turquoise | HLA-DRA |
| 2.91E-305 | -0.55882 | 0.099779 | 0.489145 | 6.39E-305 | turquoise | PGD |
| 1.70E-303 | 0.756986 | 0.656457 | 0.454324 | 3.70E-303 | turquoise | MT2A |
| 2.51E-302 | 0.659987 | 0.369119 | 0.140216 | 5.42E-302 | turquoise | RUNX3 |
| 2.72E-301 | -0.61236 | 0.096306 | 0.475962 | 5.81E-301 | turquoise | TMEM14C |
| 2.93E-301 | 0.55822 | 0.79318 | 0.562048 | 6.19E-301 | turquoise | EVL |
| 4.69E-301 | -0.52512 | 0.026524 | 0.369958 | 9.84E-301 | turquoise | MPP1 |
| 1.36E-300 | -0.50916 | 0.01042 | 0.340152 | 2.83E-300 | turquoise | SYK |
| 1.06E-298 | -0.62122 | 0.026524 | 0.365874 | 2.18E-298 | turquoise | LAT2 |
| 5.39E-298 | -0.56616 | 0.047048 | 0.39919 | 1.10E-297 | turquoise | RNF130 |
| 6.59E-291 | -0.75786 | 0.051153 | 0.394569 | 1.33E-290 | turquoise | ITM2C |
| 2.22E-290 | 0.595776 | 0.395327 | 0.150032 | 4.45E-290 | turquoise | GIMAP4 |
| 3.20E-289 | -0.61056 | 0.00221 | 0.315326 | 6.35E-289 | turquoise | MGST1 |
| 4.91E-288 | -0.6632 | 0.000316 | 0.310919 | 9.65E-288 | turquoise | CD34 |
| 1.37E-287 | -0.64933 | 0.002526 | 0.314466 | 2.68E-287 | turquoise | FAM30A |
| 5.18E-287 | -0.60733 | 0.02305 | 0.350075 | 1.00E-286 | turquoise | CAPG |
| 5.44E-287 | 0.688653 | 0.397221 | 0.166691 | 1.04E-286 | turquoise | SYNE1 |
| 4.16E-285 | 0.6088 | 0.453426 | 0.194848 | 7.89E-285 | turquoise | GIMAP7 |
| 9.69E-285 | -0.73818 | 0.011367 | 0.326969 | 1.82E-284 | turquoise | CLEC11A |
| 4.81E-281 | 0.671932 | 0.648879 | 0.497922 | 8.98E-281 | turquoise | PAXX |
| 4.13E-280 | 0.655661 | 0.425008 | 0.192627 | 7.64E-280 | turquoise | PARP8 |
| 5.62E-280 | 0.699322 | 0.693401 | 0.520277 | 1.03E-279 | turquoise | FLNA |
| 1.64E-278 | 0.717716 | 0.418693 | 0.190155 | 2.98E-278 | turquoise | TENT5C |
| 4.60E-275 | -0.70891 | 0.003789 | 0.306262 | 8.29E-275 | turquoise | CSF3R |
| 3.12E-271 | 0.586627 | 0.539312 | 0.278319 | 5.58E-271 | turquoise | ARL4C |
| 2.23E-268 | -0.66488 | 0.218503 | 0.565487 | 3.96E-268 | turquoise | HSPD1 |
| 6.41E-268 | 0.60721 | 0.459425 | 0.218743 | 1.13E-267 | turquoise | SKAP1 |
| 2.01E-267 | 0.571714 | 0.383328 | 0.150319 | 3.51E-267 | turquoise | CD5 |
| 7.07E-267 | -0.51318 | 0.544995 | 0.812567 | 1.22E-266 | turquoise | EIF3E |
| 1.49E-266 | -0.68609 | 0.168298 | 0.536433 | 2.56E-266 | turquoise | HLA-DMA |
| 5.10E-266 | -0.57274 | 0.532681 | 0.811134 | 8.69E-266 | turquoise | ZFAS1 |
| 5.55E-266 | 0.617906 | 0.628039 | 0.406678 | 9.39E-266 | turquoise | HLA-F |
| 8.30E-266 | -0.588 | 0.527629 | 0.775991 | 1.39E-265 | turquoise | SLC25A5 |
| 3.82E-265 | -0.51513 | 0.070729 | 0.410153 | 6.37E-265 | turquoise | DDAH2 |
| 1.55E-264 | -0.5072 | 0.131986 | 0.499427 | 2.57E-264 | turquoise | YWHAE |
| 1.47E-261 | -0.63205 | 0.781497 | 0.905173 | 2.41E-261 | turquoise | RPS2 |
| 8.15E-260 | -0.53149 | 0.439848 | 0.748907 | 1.33E-259 | turquoise | ANP32B |
| 4.71E-258 | -0.70271 | 0.171456 | 0.520671 | 7.62E-258 | turquoise | GLUL |
| 6.29E-258 | -0.80597 | 0.33186 | 0.641363 | 1.01E-257 | turquoise | SERPINB1 |

| | | | | | | |
|---|---|---|---|---|---|---|
| 7.62E-256 | -0.77826 | 0.004105 | 0.289353 | 1.21E-255 | turquoise | SMIM24 |
| 8.39E-256 | 0.552935 | 0.788443 | 0.721967 | 1.33E-255 | turquoise | LCP1 |
| 5.02E-255 | -0.54934 | 0.257973 | 0.612058 | 7.89E-255 | turquoise | APEX1 |
| 8.05E-255 | -0.57689 | 0.005052 | 0.29118 | 1.26E-254 | turquoise | TCF4 |
| 1.24E-254 | -0.58688 | 0.198295 | 0.586802 | 1.93E-254 | turquoise | NPC2 |
| 1.52E-254 | -0.50729 | 0.010736 | 0.301748 | 2.34E-254 | turquoise | TNFSF13B |
| 8.69E-253 | -0.66999 | 0.07515 | 0.402164 | 1.33E-252 | turquoise | GRN |
| 1.03E-249 | 0.556853 | 0.382381 | 0.158917 | 1.56E-249 | turquoise | DOK2 |
| 1.47E-249 | -0.6253 | 0.107673 | 0.434012 | 2.21E-249 | turquoise | CDK6 |
| 3.84E-247 | -1.4423 | 0.005684 | 0.283944 | 5.75E-247 | turquoise | IGLL1 |
| 1.69E-243 | -0.6316 | 0.006947 | 0.284553 | 2.51E-243 | turquoise | H1F0 |
| 2.06E-243 | -0.50627 | 0.00221 | 0.275919 | 3.05E-243 | turquoise | SPI1 |
| 3.45E-238 | -0.62174 | 0.770761 | 0.880132 | 5.05E-238 | turquoise | HSP90AB1 |
| 5.68E-238 | 0.652505 | 0.501737 | 0.311457 | 8.28E-238 | turquoise | APMAP |
| 3.58E-237 | -0.50712 | 0.000632 | 0.267536 | 5.18E-237 | turquoise | CYTL1 |
| 3.42E-236 | -0.93636 | 0.008841 | 0.278892 | 4.91E-236 | turquoise | SPINK2 |
| 1.19E-229 | 0.973674 | 0.439848 | 0.24991 | 1.70E-229 | turquoise | CMC1 |
| 5.79E-227 | 0.64848 | 0.636249 | 0.511643 | 8.22E-227 | turquoise | SLC9A3R1 |
| 1.95E-226 | 0.620307 | 0.487212 | 0.284051 | 2.75E-226 | turquoise | GNG2 |
| 5.47E-225 | 0.632058 | 0.612567 | 0.471699 | 7.67E-225 | turquoise | WIPF1 |
| 2.48E-222 | 0.672048 | 0.373855 | 0.180268 | 3.45E-222 | turquoise | APOBEC3G |
| 2.98E-222 | -0.54638 | 0.095674 | 0.402415 | 4.13E-222 | turquoise | FABP5 |
| 3.95E-222 | 0.531368 | 0.383328 | 0.170022 | 5.43E-222 | turquoise | CD96 |
| 4.25E-222 | -0.54498 | 0.063783 | 0.359748 | 5.80E-222 | turquoise | FCGRT |
| 2.63E-218 | 0.545553 | 0.370382 | 0.16386 | 3.57E-218 | turquoise | TC2N |
| 2.43E-212 | 0.600549 | 0.611304 | 0.47066 | 3.28E-212 | turquoise | BIN2 |
| 1.33E-207 | 0.580049 | 0.403536 | 0.198395 | 1.78E-207 | turquoise | CLEC2D |
| 1.62E-205 | 0.583809 | 0.620145 | 0.483664 | 2.15E-205 | turquoise | ANXA6 |
| 5.16E-204 | -0.6445 | 0.308178 | 0.608655 | 6.84E-204 | turquoise | PRDX1 |
| 2.40E-203 | -0.53801 | 0.430692 | 0.750699 | 3.16E-203 | turquoise | FOS |
| 3.36E-201 | -0.80364 | 0.991159 | 0.997098 | 4.39E-201 | turquoise | FTH1 |
| 3.33E-200 | 0.514448 | 0.710452 | 0.643763 | 4.34E-200 | turquoise | LY6E |
| 3.85E-198 | -0.67017 | 0.703821 | 0.873325 | 4.99E-198 | turquoise | HMGN2 |
| 1.15E-197 | 0.618563 | 0.464477 | 0.276922 | 1.48E-197 | turquoise | AKNA |
| 1.48E-195 | -1.10488 | 0.034417 | 0.282869 | 1.90E-195 | turquoise | FCER1G |
| 4.23E-192 | 0.52213 | 0.68898 | 0.608082 | 5.38E-192 | turquoise | PSMB9 |
| 7.77E-192 | 0.543785 | 0.461636 | 0.262198 | 9.82E-192 | turquoise | CRIP1 |
| 9.71E-192 | 0.510927 | 0.364383 | 0.171061 | 1.22E-191 | turquoise | LIME1 |
| 1.03E-188 | 0.634366 | 0.586044 | 0.42853 | 1.28E-188 | turquoise | IRF1 |
| 2.67E-187 | 0.52161 | 0.452479 | 0.244644 | 3.32E-187 | turquoise | LBH |
| 7.04E-180 | -2.50722 | 0.042311 | 0.274629 | 8.70E-180 | turquoise | LYZ |

| | | | | | | |
|---|---|---|---|---|---|---|
| 4.95E-177 | 0.532757 | 0.671298 | 0.54281 | 6.08E-177 | turquoise | DNAJB1 |
| 3.68E-174 | 0.53588 | 0.293338 | 0.130651 | 4.49E-174 | turquoise | ZBTB38 |
| 9.24E-171 | -0.51338 | 0.377644 | 0.611378 | 1.12E-170 | turquoise | TXN |
| 1.12E-170 | -0.6406 | 0.982318 | 0.969227 | 1.35E-170 | turquoise | GAPDH |
| 2.92E-170 | -0.56283 | 0.061257 | 0.303468 | 3.51E-170 | turquoise | BLVRB |
| 3.94E-170 | -0.53703 | 0.322703 | 0.584259 | 4.71E-170 | turquoise | NUCB2 |
| 5.60E-170 | 0.549891 | 0.519735 | 0.352261 | 6.66E-170 | turquoise | RNF213 |
| 7.55E-169 | -0.63029 | 0.640354 | 0.74837 | 8.93E-169 | turquoise | GSTP1 |
| 2.88E-165 | 0.582367 | 0.516261 | 0.361575 | 3.39E-165 | turquoise | P2RY8 |
| 1.22E-164 | -0.72949 | 0.157562 | 0.438525 | 1.43E-164 | turquoise | TIMP1 |
| 1.30E-162 | -0.75116 | 0.386801 | 0.640646 | 1.51E-162 | turquoise | H2AFZ |
| 5.65E-158 | -0.99251 | 0.11462 | 0.354625 | 6.55E-158 | turquoise | AREG |
| 6.96E-157 | 0.53935 | 0.563309 | 0.394856 | 8.02E-157 | turquoise | DDIT4 |
| 2.20E-155 | -1.0301 | 0.493527 | 0.681414 | 2.52E-155 | turquoise | TUBA1B |
| 1.79E-153 | 0.539256 | 0.33186 | 0.170488 | 2.04E-153 | turquoise | CBLB |
| 2.94E-151 | 0.581097 | 0.622987 | 0.466719 | 3.34E-151 | turquoise | CD69 |
| 9.43E-151 | 0.590239 | 0.517524 | 0.403131 | 1.06E-150 | turquoise | CLEC2B |
| 1.12E-149 | -0.87931 | 0.425639 | 0.647525 | 1.26E-149 | turquoise | TUBB |
| 4.11E-148 | -0.80194 | 0.373224 | 0.605538 | 4.59E-148 | turquoise | HLA-DRB5 |
| 1.63E-145 | 0.50459 | 0.564256 | 0.425915 | 1.82E-145 | turquoise | 1-Sep |
| 1.72E-143 | 0.504511 | 0.729713 | 0.703625 | 1.90E-143 | turquoise | SRSF7 |
| 3.00E-140 | 0.550864 | 0.460688 | 0.314108 | 3.31E-140 | turquoise | EML4 |
| 9.82E-136 | 0.517981 | 0.325229 | 0.171706 | 1.08E-135 | turquoise | AOAH |
| 5.65E-135 | 0.500293 | 0.346069 | 0.195744 | 6.15E-135 | turquoise | OPTN |
| 9.99E-132 | 0.655229 | 0.524155 | 0.430143 | 1.08E-131 | turquoise | PIK3R1 |
| 4.10E-130 | -0.65096 | 0.154405 | 0.392491 | 4.43E-130 | turquoise | HLA-DQB1 |
| 1.54E-124 | 0.661358 | 0.340385 | 0.208605 | 1.65E-124 | turquoise | ADGRG1 |
| 4.37E-121 | 0.519685 | 0.341648 | 0.200437 | 4.66E-121 | turquoise | BHLHE40 |
| 2.32E-119 | -0.51253 | 0.06694 | 0.260013 | 2.47E-119 | turquoise | ATP2B1-AS1 |
| 5.53E-117 | -0.67509 | 0.346069 | 0.623594 | 5.85E-117 | turquoise | HIST1H4C |
| 1.91E-115 | 0.6515 | 0.403221 | 0.283263 | 2.01E-115 | turquoise | ITGB1 |
| 8.10E-108 | 0.595988 | 0.467635 | 0.381386 | 8.48E-108 | turquoise | CCNH |
| 2.57E-105 | -0.57645 | 0.092201 | 0.271799 | 2.68E-105 | turquoise | NCF1 |
| 4.35E-99 | 0.573329 | 0.473319 | 0.346672 | 4.51E-99 | turquoise | NR4A2 |
| 2.35E-96 | -0.62549 | 0.376066 | 0.633231 | 2.42E-96 | turquoise | NEAT1 |
| 2.78E-85 | -0.86805 | 0.168614 | 0.343089 | 2.86E-85 | turquoise | TYROBP |
| 4.50E-79 | -1.10322 | 0.919166 | 0.918858 | 4.61E-79 | turquoise | CD74 |
| 7.37E-65 | -0.6305 | 0.284812 | 0.40915 | 7.51E-65 | turquoise | LTB |
| 2.53E-49 | -0.52881 | 0.302179 | 0.483234 | 2.56E-49 | turquoise | CTSS |
| 1.11E-38 | -0.6246 | 0.988633 | 0.994734 | 1.12E-38 | turquoise | FTL |
| 3.37E-34 | -0.66908 | 0.581939 | 0.647739 | 3.39E-34 | turquoise | HLA-DRB1 |

| | | | | | | |
|---|---|---|---|---|---|---|
| 1.15E-28 | -0.65191 | 0.462583 | 0.641291 | 1.15E-28 | turquoise | SAT1 |
| 0 | 1.177564 | 0.975942 | 0.748084 | 0 | blue | JUNB |
| 0 | 1.149569 | 0.938272 | 0.335184 | 0 | blue | LTB |
| 0 | 0.986686 | 0.857233 | 0.521811 | 0 | blue | DNAJB1 |
| 0 | 0.951254 | 0.992403 | 0.896354 | 0 | blue | SARAF |
| 0 | 0.93492 | 0.802469 | 0.37859 | 0 | blue | TNFAIP3 |
| 0 | 0.926229 | 0.701804 | 0.205752 | 0 | blue | FLT3LG |
| 0 | 0.912157 | 0.871162 | 0.518874 | 0 | blue | LEPROTL1 |
| 0 | 0.853967 | 0.984805 | 0.335327 | 0 | blue | CD3E |
| 0 | 0.844063 | 0.980057 | 0.526861 | 0 | blue | IFITM1 |
| 0 | 0.831061 | 0.741374 | 0.367846 | 0 | blue | SLC2A3 |
| 0 | 0.816688 | 0.507439 | 0.204212 | 0 | blue | ZNF331 |
| 0 | 0.801293 | 0.531497 | 0.224124 | 0 | blue | ABLIM1 |
| 0 | 0.763994 | 0.628997 | 0.156436 | 0 | blue | TCF7 |
| 0 | 0.740477 | 0.54321 | 0.132297 | 0 | blue | CD5 |
| 0 | 0.717897 | 0.471668 | 0.087386 | 0 | blue | MAL |
| 0 | 0.70821 | 0.67648 | 0.247833 | 0 | blue | PIK3IP1 |
| 0 | 0.697838 | 0.719848 | 0.256429 | 0 | blue | SPOCK2 |
| 0 | 0.690333 | 0.706553 | 0.486641 | 0 | blue | NOSIP |
| 0 | 0.682581 | 0.810383 | 0.263878 | 0 | blue | CD3D |
| 0 | 0.678679 | 0.522634 | 0.122627 | 0 | blue | CAMK4 |
| 0 | 0.65118 | 0.747072 | 0.238378 | 0 | blue | IL7R |
| 0 | 0.644014 | 0.846787 | 0.456844 | 0 | blue | FYB1 |
| 0 | 0.640209 | 0.513137 | 0.170224 | 0 | blue | OXNAD1 |
| 0 | 0.630574 | 0.971193 | 0.964616 | 0 | blue | RPS26 |
| 0 | 0.613546 | 0.545426 | 0.247296 | 0 | blue | MGAT4A |
| 0 | 0.600946 | 1 | 0.993983 | 0 | blue | RPS27 |
| 0 | 0.600522 | 0.783159 | 0.367667 | 0 | blue | CD7 |
| 0 | 0.597868 | 0.35106 | 0.093081 | 0 | blue | AQP3 |
| 0 | 0.586863 | 0.443178 | 0.118724 | 0 | blue | CD6 |
| 0 | 0.563601 | 0.918645 | 0.328164 | 0 | blue | IL32 |
| 0 | 0.56334 | 0.465021 | 0.135735 | 0 | blue | CD27 |
| 0 | 0.554876 | 0.560304 | 0.225915 | 0 | blue | KLF2 |
| 0 | 0.540761 | 0.871478 | 0.553255 | 0 | blue | EVL |
| 0 | 0.540414 | 0.432732 | 0.134625 | 0 | blue | LEF1 |
| 0 | 0.52266 | 0.559671 | 0.233615 | 0 | blue | LAT |
| 0 | 0.521433 | 0.616967 | 0.210479 | 0 | blue | CD3G |
| 0 | 0.514469 | 0.747705 | 0.413652 | 0 | blue | CD48 |
| 0 | 0.503238 | 0.978474 | 0.925256 | 0 | blue | RPS29 |
| 0 | -0.53944 | 0.002532 | 0.34095 | 0 | blue | SYK |
| 0 | -0.58207 | 0.806584 | 0.935177 | 0 | blue | OAZ1 |

| | | | | | | |
|---|---|---|---|---|---|---|
| 0 | -0.5871 | 0.127255 | 0.507987 | 0 | blue | COMT |
| 0 | -0.61271 | 0.002849 | 0.333214 | 0 | blue | MEF2C |
| 0 | -0.63528 | 0.510605 | 0.765919 | 0 | blue | TPI1 |
| 0 | -0.64494 | 0.035771 | 0.41401 | 0 | blue | DDAH2 |
| 0 | -0.66484 | 0.233935 | 0.626531 | 0 | blue | ATP6V0B |
| 0 | -0.686 | 0.001899 | 0.340305 | 0 | blue | EGFL7 |
| 0 | -0.70116 | 0.006964 | 0.367989 | 0 | blue | LAT2 |
| 0 | -0.70774 | 0.08452 | 0.477545 | 0 | blue | APLP2 |
| 0 | -0.71858 | 0.128522 | 0.523637 | 0 | blue | PYCARD |
| 0 | -0.72356 | 0.054448 | 0.406991 | 0 | blue | FABP5 |
| 0 | -0.72544 | 0.918645 | 0.953728 | 0 | blue | H3F3A |
| 0 | -0.76124 | 0.164926 | 0.535957 | 0 | blue | H2AFY |
| 0 | -0.77053 | 0.137069 | 0.492157 | 0 | blue | CAT |
| 0 | -0.80066 | 0.29408 | 0.620765 | 0 | blue | TXN |
| 0 | -0.80177 | 0.016777 | 0.383318 | 0 | blue | PLEK |
| 0 | -0.80515 | 0.864198 | 0.926402 | 0 | blue | HMGB1 |
| 0 | -0.81017 | 0.393796 | 0.667108 | 0 | blue | TKT |
| 0 | -0.85785 | 0.014562 | 0.477974 | 0 | blue | ZEB2 |
| 0 | -0.87359 | 0.039886 | 0.410751 | 0 | blue | MAP3K8 |
| 0 | -0.88532 | 0.041785 | 0.404233 | 0 | blue | AC084033.3 |
| 0 | -0.88725 | 0.110795 | 0.527434 | 0 | blue | GLUL |
| 0 | -0.89484 | 0.461222 | 0.834002 | 0 | blue | CLIC1 |
| 0 | -0.91556 | 0.049383 | 0.431524 | 0 | blue | SOX4 |
| 0 | -0.93057 | 0.297879 | 0.654681 | 0 | blue | TALDO1 |
| 0 | -0.99024 | 0.336499 | 0.640749 | 0 | blue | SERPINB1 |
| 0 | -1.04161 | 0.041152 | 0.405236 | 0 | blue | HLA-DQB1 |
| 0 | -1.05 | 0.059196 | 0.548671 | 0 | blue | HLA-DMA |
| 0 | -1.05165 | 0.966762 | 0.970991 | 0 | blue | GAPDH |
| 0 | -1.23784 | 0.006331 | 0.373218 | 0 | blue | PRSS57 |
| 0 | -1.36282 | 0.377018 | 0.778132 | 0 | blue | GSTP1 |
| 0 | -1.48274 | 0.173473 | 0.706969 | 0 | blue | HLA-DPB1 |
| 0 | -1.4878 | 0.150681 | 0.514755 | 0 | blue | STMN1 |
| 0 | -1.56068 | 0.169991 | 0.705286 | 0 | blue | HLA-DPA1 |
| 0 | -1.74516 | 0.055397 | 0.64143 | 0 | blue | HLA-DRB5 |
| 0 | -1.83757 | 0.01836 | 0.452045 | 0 | blue | CST3 |
| 0 | -2.01114 | 0.087053 | 0.70371 | 0 | blue | HLA-DRB1 |
| 0 | -2.0123 | 0.014562 | 0.360468 | 0 | blue | TYROBP |
| 0 | -2.25501 | 0.775245 | 0.935141 | 0 | blue | CD74 |
| 0 | -2.6306 | 0.067426 | 0.662632 | 0 | blue | HLA-DRA |
| 2.61E-306 | -0.60823 | 0.048116 | 0.398969 | 5.09E-306 | blue | RNF130 |
| 7.26E-306 | -0.59349 | 0.112694 | 0.487573 | 1.40E-305 | blue | PGD |

| | | | | | | |
|---|---|---|---|---|---|---|
| 1.11E-303 | -0.61055 | 0.126939 | 0.509168 | 2.10E-303 | blue | CD63 |
| 3.31E-302 | -0.5777 | 0.019626 | 0.356099 | 6.23E-302 | blue | GNA15 |
| 4.92E-302 | -0.66123 | 0.538145 | 0.747941 | 9.14E-302 | blue | ENO1 |
| 5.26E-301 | -0.93178 | 0.012029 | 0.340842 | 9.65E-301 | blue | HLA-DQA1 |
| 1.62E-296 | -0.64042 | 0.404242 | 0.698625 | 2.93E-296 | blue | VAMP8 |
| 2.86E-292 | -1.26467 | 0.446027 | 0.686734 | 5.13E-292 | blue | TUBA1B |
| 1.65E-289 | -0.61378 | 0.001899 | 0.315271 | 2.93E-289 | blue | MGST1 |
| 8.47E-288 | -0.65272 | 0.002216 | 0.314412 | 1.48E-287 | blue | FAM30A |
| 1.18E-287 | -0.57713 | 0.377335 | 0.665389 | 2.05E-287 | blue | COX5A |
| 2.83E-287 | -0.66331 | 0.000317 | 0.31083 | 4.85E-287 | blue | CD34 |
| 1.85E-280 | -0.50617 | 0.070909 | 0.416911 | 3.13E-280 | blue | PRDX3 |
| 9.52E-280 | -0.56997 | 0.433048 | 0.720829 | 1.60E-279 | blue | GNG5 |
| 2.72E-274 | -0.7137 | 0.004115 | 0.306139 | 4.51E-274 | blue | CSF3R |
| 3.91E-272 | 0.507213 | 0.376701 | 0.144366 | 6.42E-272 | blue | INPP4B |
| 3.39E-260 | -0.78835 | 0.001583 | 0.289557 | 5.50E-260 | blue | SMIM24 |
| 9.00E-258 | -0.58356 | 0.003482 | 0.291276 | 1.45E-257 | blue | TCF4 |
| 2.20E-253 | -0.56971 | 0.606204 | 0.76703 | 3.50E-253 | blue | SLC25A5 |
| 2.63E-253 | -0.73619 | 0.320671 | 0.607156 | 4.15E-253 | blue | PRDX1 |
| 1.69E-250 | -0.52446 | 0.162077 | 0.495917 | 2.63E-250 | blue | YWHAE |
| 7.55E-250 | -0.50532 | 0.216841 | 0.558914 | 1.17E-249 | blue | SET |
| 1.04E-242 | -0.506 | 0.002216 | 0.27584 | 1.58E-242 | blue | SPI1 |
| 2.82E-242 | -1.44054 | 0.00823 | 0.283576 | 4.27E-242 | blue | IGLL1 |
| 1.44E-236 | -0.69157 | 0.088636 | 0.400544 | 2.15E-236 | blue | GRN |
| 5.37E-236 | -0.50721 | 0.00095 | 0.267424 | 7.98E-236 | blue | CYTL1 |
| 2.21E-234 | -2.11454 | 0.049066 | 0.334396 | 3.25E-234 | blue | NKG7 |
| 8.75E-234 | -0.90328 | 0.391263 | 0.640069 | 1.28E-233 | blue | H2AFZ |
| 2.11E-232 | -0.52717 | 0.191516 | 0.514648 | 3.05E-232 | blue | NAA38 |
| 1.44E-226 | -0.5135 | 0.090852 | 0.397572 | 2.06E-226 | blue | TUBA1C |
| 3.77E-226 | -0.54934 | 0.139285 | 0.446745 | 5.34E-226 | blue | NASP |
| 1.37E-218 | -0.81525 | 0.992086 | 0.994341 | 1.93E-218 | blue | FTL |
| 9.54E-216 | 0.550341 | 0.510605 | 0.294642 | 1.33E-215 | blue | RNF125 |
| 5.91E-215 | -1.30064 | 0.026274 | 0.283719 | 8.15E-215 | blue | FCER1G |
| 2.32E-214 | -0.83669 | 0.114593 | 0.410536 | 3.17E-214 | blue | LST1 |
| 1.47E-211 | -0.53254 | 0.173156 | 0.497672 | 2.00E-211 | blue | ASAH1 |
| 3.91E-210 | -0.88648 | 0.331117 | 0.638242 | 5.26E-210 | blue | NEAT1 |
| 1.49E-209 | -0.58986 | 0.024375 | 0.282501 | 1.98E-209 | blue | H1F0 |
| 6.94E-208 | -0.91473 | 0.3327 | 0.574099 | 9.17E-208 | blue | AIF1 |
| 1.13E-206 | -0.51448 | 0.065527 | 0.345176 | 1.48E-206 | blue | CAPG |
| 2.75E-205 | -0.54715 | 0.076607 | 0.351049 | 3.58E-205 | blue | MCM7 |
| 9.85E-204 | -0.60189 | 0.010446 | 0.254137 | 1.27E-203 | blue | HLA-DQA2 |
| 1.59E-203 | -0.89879 | 0.029756 | 0.276449 | 2.03E-203 | blue | SPINK2 |

| | | | | | | |
|---|---|---|---|---|---|---|
| 4.77E-203 | -0.58732 | 0.150364 | 0.429088 | 6.05E-203 | blue | CDK6 |
| 1.59E-200 | 0.684124 | 0.550491 | 0.337977 | 2.00E-200 | blue | NR4A2 |
| 1.26E-194 | -0.63392 | 0.049383 | 0.304742 | 1.58E-194 | blue | BLVRB |
| 2.54E-192 | -0.63385 | 0.063628 | 0.320966 | 3.14E-192 | blue | CLEC11A |
| 1.21E-189 | -0.51125 | 0.472618 | 0.685803 | 1.49E-189 | blue | RPL22L1 |
| 2.78E-188 | -0.95823 | 0.452991 | 0.644366 | 3.39E-188 | blue | TUBB |
| 2.52E-178 | 0.500462 | 0.54321 | 0.377444 | 3.05E-178 | blue | NDFIP1 |
| 3.67E-172 | -0.61945 | 0.129471 | 0.38561 | 4.41E-172 | blue | ITM2C |
| 1.18E-166 | -0.54362 | 0.055714 | 0.293425 | 1.41E-166 | blue | CMC1 |
| 1.54E-159 | 0.612009 | 0.565369 | 0.445778 | 1.82E-159 | blue | PTP4A1 |
| 3.39E-157 | -2.47231 | 0.058246 | 0.27276 | 3.98E-157 | blue | LYZ |
| 1.82E-156 | -0.60043 | 0.059829 | 0.283683 | 2.12E-156 | blue | HOPX |
| 2.99E-151 | 0.518723 | 0.501425 | 0.350978 | 3.46E-151 | blue | FAM177A1 |
| 2.77E-150 | 0.508263 | 0.671098 | 0.550104 | 3.18E-150 | blue | ZFP36L2 |
| 3.65E-149 | -0.55123 | 0.396645 | 0.575818 | 4.16E-149 | blue | NUCB2 |
| 5.09E-147 | -0.56569 | 0.320671 | 0.507987 | 5.75E-147 | blue | IMPDH2 |
| 3.64E-142 | -0.51436 | 0.356758 | 0.549746 | 4.08E-142 | blue | HSPD1 |
| 5.33E-140 | -1.04544 | 0.181387 | 0.402049 | 5.93E-140 | blue | LGALS1 |
| 1.44E-134 | -0.60606 | 0.81798 | 0.860361 | 1.59E-134 | blue | HMGN2 |
| 1.64E-129 | -1.10519 | 0.114277 | 0.317384 | 1.81E-129 | blue | CST7 |
| 3.26E-121 | -0.50972 | 0.358974 | 0.558269 | 3.56E-121 | blue | DUT |
| 1.09E-106 | -0.73253 | 0.266223 | 0.48725 | 1.18E-106 | blue | CTSS |
| 4.08E-104 | -0.51008 | 0.119658 | 0.31588 | 4.39E-104 | blue | NAMPT |
| 1.17E-99 | -0.65223 | 0.358974 | 0.587028 | 1.25E-99 | blue | CTSW |
| 1.15E-97 | -0.66495 | 0.505856 | 0.714419 | 1.22E-97 | blue | PSAP |
| 1.40E-67 | -0.58575 | 0.550491 | 0.700988 | 1.48E-67 | blue | TSPO |
| 9.50E-52 | -0.78147 | 0.471668 | 0.640212 | 9.95E-52 | blue | SAT1 |
| 3.80E-51 | -0.51301 | 0.289016 | 0.438328 | 3.95E-51 | blue | DUSP1 |
| 9.47E-49 | -0.67388 | 0.232985 | 0.359716 | 9.79E-49 | blue | S100A11 |
| 2.52E-46 | -0.52639 | 0.712884 | 0.8229 | 2.58E-46 | blue | S100A4 |
| 7.96E-45 | -0.5406 | 0.277303 | 0.424898 | 8.12E-45 | blue | TIMP1 |
| 3.62E-09 | -0.68295 | 0.310225 | 0.332426 | 3.66E-09 | blue | AREG |
| 5.27E-07 | -1.33578 | 0.258626 | 0.209441 | 5.30E-07 | blue | HBB |
| 1.98E-116 | 0.785628 | 0.483465 | 0.145963 | 1.19E-115 | red | S100A8 |
| 4.36E-106 | 0.600704 | 0.480315 | 0.149445 | 1.31E-105 | red | S100A9 |
| 1.17E-84 | 0.505256 | 0.370079 | 0.108389 | 2.35E-84 | red | VCAN |
| 2.10E-77 | 0.522451 | 0.340157 | 0.101557 | 3.15E-77 | red | S100A12 |
| 2.54E-22 | -0.66389 | 0.171654 | 0.339355 | 3.04E-22 | red | PRSS57 |
| 7.58E-17 | -0.66787 | 0.111811 | 0.258589 | 7.58E-17 | red | IGLL1 |
| 0 | 3.074676 | 0.946694 | 0.083685 | 0 | darkred | GNLY |
| 0 | 2.607935 | 0.929825 | 0.066286 | 0 | darkred | GZMB |

| | | | | | | |
|---|---|---|---|---|---|---|
| 0 | 2.561113 | 1 | 0.270617 | 0 | darkred | NKG7 |
| 0 | 2.439064 | 0.920378 | 0.095071 | 0 | darkred | PRF1 |
| 0 | 2.299723 | 0.775978 | 0.077604 | 0 | darkred | SPON2 |
| 0 | 2.132962 | 0.835358 | 0.070847 | 0 | darkred | FGFBP2 |
| 0 | 2.103063 | 0.994602 | 0.261799 | 0 | darkred | CST7 |
| 0 | 2.073091 | 0.747638 | 0.037603 | 0 | darkred | CLIC3 |
| 0 | 1.941482 | 0.714575 | 0.083922 | 0 | darkred | CCL4 |
| 0 | 1.937056 | 0.851552 | 0.064935 | 0 | darkred | KLRD1 |
| 0 | 1.916512 | 0.803644 | 0.076692 | 0 | darkred | KLRB1 |
| 0 | 1.91369 | 0.769906 | 0.042062 | 0 | darkred | FCGR3A |
| 0 | 1.908545 | 0.950067 | 0.144971 | 0 | darkred | GZMA |
| 0 | 1.875194 | 0.919703 | 0.20575 | 0 | darkred | CD247 |
| 0 | 1.837485 | 0.788124 | 0.085037 | 0 | darkred | IL2RB |
| 0 | 1.828883 | 0.981107 | 0.542958 | 0 | darkred | CTSW |
| 0 | 1.765148 | 0.952767 | 0.382716 | 0 | darkred | CD7 |
| 0 | 1.706626 | 0.822537 | 0.186155 | 0 | darkred | CCL5 |
| 0 | 1.617661 | 0.65857 | 0.015237 | 0 | darkred | KLRF1 |
| 0 | 1.565511 | 0.603914 | 0.005135 | 0 | darkred | SH2D1B |
| 0 | 1.554328 | 0.802294 | 0.233825 | 0 | darkred | HOPX |
| 0 | 1.492838 | 0.692982 | 0.248049 | 0 | darkred | CMC1 |
| 0 | 1.489554 | 0.991228 | 0.551978 | 0 | darkred | IFITM1 |
| 0 | 1.430575 | 0.762483 | 0.309233 | 0 | darkred | APMAP |
| 0 | 1.429188 | 0.371795 | 0.00652 | 0 | darkred | KIR3DL1 |
| 0 | 1.38155 | 0.497301 | 0.02446 | 0 | darkred | TRDC |
| 0 | 1.374153 | 0.651822 | 0.089192 | 0 | darkred | GZMH |
| 0 | 1.357559 | 0.969636 | 0.764046 | 0 | darkred | IFITM2 |
| 0 | 1.34622 | 0.825911 | 0.223825 | 0 | darkred | GZMM |
| 0 | 1.34422 | 0.591768 | 0.096929 | 0 | darkred | TTC38 |
| 0 | 1.320395 | 0.962213 | 0.628163 | 0 | darkred | ITGB2 |
| 0 | 1.309437 | 0.423077 | 0.028008 | 0 | darkred | XCL2 |
| 0 | 1.296667 | 0.896086 | 0.501098 | 0 | darkred | LITAF |
| 0 | 1.273772 | 0.641026 | 0.135917 | 0 | darkred | SH2D2A |
| 0 | 1.273067 | 0.816464 | 0.229569 | 0 | darkred | FCER1G |
| 0 | 1.26779 | 0.525641 | 0.031589 | 0 | darkred | S1PR5 |
| 0 | 1.192947 | 0.925101 | 0.697422 | 0 | darkred | ZFP36 |
| 0 | 1.179591 | 0.545209 | 0.146221 | 0 | darkred | TRBC1 |
| 0 | 1.178555 | 0.557355 | 0.205243 | 0 | darkred | ADGRG1 |
| 0 | 1.178152 | 0.311741 | 0.009798 | 0 | darkred | KLRC1 |
| 0 | 1.149351 | 0.589744 | 0.167202 | 0 | darkred | AOAH |
| 0 | 1.139857 | 0.972335 | 0.292915 | 0 | darkred | TYROBP |
| 0 | 1.124858 | 0.646424 | 0.241089 | 0 | darkred | ALOX5AP |

| | | | | | | |
|---|---|---|---|---|---|---|
| 0 | 1.107143 | 0.280027 | 0.002534 | 0 | darkred | KIR2DL1 |
| 0 | 1.103172 | 0.446019 | 0.036961 | 0 | darkred | PRSS23 |
| 0 | 1.092414 | 0.701754 | 0.254671 | 0 | darkred | ZAP70 |
| 0 | 1.088725 | 0.387314 | 0.026014 | 0 | darkred | CX3CR1 |
| 0 | 1.085174 | 0.646424 | 0.287645 | 0 | darkred | GNG2 |
| 0 | 1.04977 | 0.48448 | 0.069428 | 0 | darkred | NCR3 |
| 0 | 1.020606 | 0.312416 | 0.017399 | 0 | darkred | LAIR2 |
| 0 | 1.019658 | 0.449393 | 0.051522 | 0 | darkred | FCRL6 |
| 0 | 0.999198 | 0.410931 | 0.031521 | 0 | darkred | PTGDR |
| 0 | 0.991488 | 0.562078 | 0.138417 | 0 | darkred | FGR |
| 0 | 0.979909 | 0.659919 | 0.287138 | 0 | darkred | ARL4C |
| 0 | 0.964081 | 0.794197 | 0.5187 | 0 | darkred | RARRES3 |
| 0 | 0.951522 | 0.429825 | 0.062975 | 0 | darkred | PYHIN1 |
| 0 | 0.950849 | 0.931849 | 0.643806 | 0 | darkred | HCST |
| 0 | 0.947687 | 1 | 0.992905 | 0 | darkred | HLA-C |
| 0 | 0.94643 | 0.349528 | 0.010068 | 0 | darkred | NCR1 |
| 0 | 0.916242 | 0.458165 | 0.087233 | 0 | darkred | SAMD3 |
| 0 | 0.90955 | 0.99865 | 0.985608 | 0 | darkred | HLA-A |
| 0 | 0.894692 | 1 | 0.999662 | 0 | darkred | B2M |
| 0 | 0.889321 | 0.995277 | 0.944525 | 0 | darkred | HLA-E |
| 0 | 0.887209 | 0.520918 | 0.164702 | 0 | darkred | DOK2 |
| 0 | 0.881846 | 0.31444 | 0.022805 | 0 | darkred | CD160 |
| 0 | 0.86456 | 0.964912 | 0.890807 | 0 | darkred | MYL12A |
| 0 | 0.857267 | 0.977733 | 0.880672 | 0 | darkred | CALM1 |
| 0 | 0.844737 | 0.917004 | 0.739518 | 0 | darkred | PTPRC |
| 0 | 0.815826 | 1 | 0.997263 | 0 | darkred | HLA-B |
| 0 | 0.805771 | 0.997976 | 0.978817 | 0 | darkred | PFN1 |
| 0 | 0.804206 | 0.397436 | 0.090814 | 0 | darkred | ITGAM |
| 0 | 0.804175 | 0.364372 | 0.071793 | 0 | darkred | TXK |
| 0 | 0.802664 | 0.259109 | 0.013379 | 0 | darkred | KLRC3 |
| 0 | 0.783208 | 0.331309 | 0.065408 | 0 | darkred | ABI3 |
| 0 | 0.777996 | 0.94197 | 0.851583 | 0 | darkred | RAC2 |
| 0 | 0.766335 | 0.355601 | 0.065881 | 0 | darkred | TGFBR3 |
| 0 | 0.737632 | 0.27193 | 0.010879 | 0 | darkred | IL18RAP |
| 0 | 0.729419 | 0.930499 | 0.865908 | 0 | darkred | ARPC2 |
| 0 | 0.669098 | 0.280702 | 0.039528 | 0 | darkred | TIGIT |
| 0 | 0.584848 | 1 | 0.993378 | 0 | darkred | TMSB4X |
| 0 | -0.50245 | 0.996626 | 0.998716 | 0 | darkred | RPL35A |
| 0 | -0.50267 | 0.999325 | 0.999223 | 0 | darkred | RPL34 |
| 0 | -0.50574 | 0.99865 | 0.99902 | 0 | darkred | RPL18 |
| 0 | -0.51637 | 0.997301 | 0.999324 | 0 | darkred | RPS24 |

| | | | | | | |
|---|---|---|---|---|---|---|
| 0 | -0.52813 | 0.982456 | 0.995878 | 0 | darkred | RPS21 |
| 0 | -0.56748 | 0.997976 | 0.998851 | 0 | darkred | RPS3A |
| 0 | -0.57039 | 0.999325 | 0.998986 | 0 | darkred | RPL13 |
| 0 | -0.57216 | 0.999325 | 0.998716 | 0 | darkred | RPL12 |
| 0 | -0.5815 | 0.977058 | 0.994662 | 0 | darkred | RPL36 |
| 0 | -0.58556 | 0.979757 | 0.993277 | 0 | darkred | RACK1 |
| 0 | -0.59197 | 0.980432 | 0.994561 | 0 | darkred | RPL9 |
| 0 | -0.60038 | 0.999325 | 0.999358 | 0 | darkred | RPL11 |
| 0 | -0.60369 | 0.956815 | 0.980168 | 0 | darkred | RPL10A |
| 0 | -0.60449 | 0.962213 | 0.994865 | 0 | darkred | RPL22 |
| 0 | -0.60657 | 1 | 0.999831 | 0 | darkred | RPLP1 |
| 0 | -0.60832 | 0.995277 | 0.999122 | 0 | darkred | RPL8 |
| 0 | -0.62453 | 0.987854 | 0.993108 | 0 | darkred | RPL5 |
| 0 | -0.64924 | 0.969636 | 0.993682 | 0 | darkred | RPS9 |
| 0 | -0.65062 | 1 | 0.999358 | 0 | darkred | RPS12 |
| 0 | -0.65545 | 0.99865 | 0.997804 | 0 | darkred | RPS4X |
| 0 | -0.66014 | 0.997976 | 0.999257 | 0 | darkred | RPL39 |
| 0 | -0.66579 | 0.867072 | 0.975979 | 0 | darkred | RPL7 |
| 0 | -0.67238 | 0.999325 | 0.999054 | 0 | darkred | RPS23 |
| 0 | -0.69432 | 0.99865 | 0.998649 | 0 | darkred | RPL18A |
| 0 | -0.72239 | 0.999325 | 0.999291 | 0 | darkred | RPL32 |
| 0 | -0.72388 | 0.993252 | 0.993513 | 0 | darkred | RPS6 |
| 0 | -0.74054 | 0.975034 | 0.983108 | 0 | darkred | RPS5 |
| 0 | -0.76101 | 0.99865 | 0.999054 | 0 | darkred | RPS13 |
| 0 | -0.79357 | 0.997301 | 0.996351 | 0 | darkred | RPS18 |
| 0 | -0.83701 | 0.997976 | 0.999561 | 0 | darkred | RPS8 |
| 0 | -0.97314 | 0.982456 | 0.99598 | 0 | darkred | RPLP0 |
| 0 | -1.0208 | 0.475709 | 0.824116 | 0 | darkred | LDHB |
| 0 | -1.31334 | 0.568826 | 0.92422 | 0 | darkred | VIM |
| 0 | -1.61995 | 0.012146 | 0.576472 | 0 | darkred | AIF1 |
| 1.27E-307 | -0.70665 | 0.936572 | 0.968208 | 3.89E-307 | darkred | EEF1B2 |
| 9.35E-307 | -1.03848 | 0.589744 | 0.907733 | 2.83E-306 | darkred | RPS2 |
| 8.98E-302 | 0.93623 | 0.502024 | 0.173013 | 2.69E-301 | darkred | ADAM8 |
| 4.31E-301 | 0.878758 | 0.25776 | 0.041285 | 1.28E-300 | darkred | CCL3 |
| 4.80E-299 | 0.699504 | 0.310391 | 0.062029 | 1.42E-298 | darkred | ZBP1 |
| 1.51E-291 | 0.837598 | 0.508097 | 0.165749 | 4.40E-291 | darkred | SYNE2 |
| 3.36E-290 | 0.890632 | 0.420378 | 0.123416 | 9.73E-290 | darkred | SLA2 |
| 1.22E-287 | 0.940526 | 0.784076 | 0.452279 | 3.51E-287 | darkred | TRBC2 |
| 1.33E-287 | -0.60888 | 0.879217 | 0.970607 | 3.79E-287 | darkred | RPL4 |
| 4.05E-283 | -2.2238 | 0.1417 | 0.62519 | 1.15E-282 | darkred | HLA-DRA |
| 5.71E-281 | 0.914222 | 0.590418 | 0.251833 | 1.60E-280 | darkred | PRKCH |

| | | | | | | |
|---|---|---|---|---|---|---|
| 1.70E-280 | 0.840377 | 0.491228 | 0.165816 | 4.74E-280 | darkred | SYTL3 |
| 1.75E-272 | 0.516302 | 0.989204 | 0.977161 | 4.85E-272 | darkred | CFL1 |
| 6.09E-269 | -0.52627 | 0.912281 | 0.988141 | 1.67E-268 | darkred | PABPC1 |
| 6.06E-266 | -0.8678 | 0.792173 | 0.925031 | 1.65E-265 | darkred | NPM1 |
| 3.90E-258 | 0.937025 | 0.719973 | 0.473225 | 1.05E-257 | darkred | BIN2 |
| 3.47E-254 | 0.77755 | 0.845479 | 0.572587 | 9.29E-254 | darkred | EVL |
| 1.63E-249 | 0.636022 | 0.932524 | 0.79827 | 4.33E-249 | darkred | LSP1 |
| 7.45E-239 | -0.76327 | 0.008097 | 0.473732 | 1.97E-238 | darkred | SPINT2 |
| 1.48E-238 | 0.822932 | 0.497976 | 0.193182 | 3.87E-238 | darkred | C12orf75 |
| 3.93E-237 | 0.952549 | 0.701754 | 0.431704 | 1.02E-236 | darkred | IRF1 |
| 6.66E-231 | 0.933693 | 0.512821 | 0.232778 | 1.72E-230 | darkred | TPST2 |
| 1.15E-227 | 0.824119 | 0.4278 | 0.150309 | 2.94E-227 | darkred | RUNX3 |
| 8.95E-219 | -0.63478 | 0.734818 | 0.92422 | 2.28E-218 | darkred | HINT1 |
| 8.96E-218 | 0.799921 | 0.456815 | 0.17683 | 2.26E-217 | darkred | SYNE1 |
| 4.33E-215 | 0.890501 | 0.439271 | 0.174465 | 1.09E-214 | darkred | CD300A |
| 4.59E-215 | 0.929017 | 0.676788 | 0.398763 | 1.14E-214 | darkred | DDIT4 |
| 5.41E-214 | 0.933679 | 0.840081 | 0.706882 | 1.34E-213 | darkred | IER2 |
| 2.35E-213 | 0.671518 | 0.280702 | 0.069698 | 5.76E-213 | darkred | RHOC |
| 8.41E-213 | 0.9246 | 0.633603 | 0.401365 | 2.05E-212 | darkred | ARHGAP9 |
| 6.93E-210 | -0.77317 | 0.402834 | 0.801784 | 1.68E-209 | darkred | ZFAS1 |
| 1.40E-209 | 0.807476 | 0.757085 | 0.632893 | 3.37E-209 | darkred | RAP1B |
| 7.18E-209 | 0.706966 | 0.379217 | 0.125241 | 1.71E-208 | darkred | RAB29 |
| 7.91E-206 | -1.27807 | 0.168016 | 0.602588 | 1.88E-205 | darkred | HLA-DRB5 |
| 9.33E-205 | 0.709719 | 0.362348 | 0.112673 | 2.20E-204 | darkred | GBP5 |
| 2.50E-204 | 0.687965 | 0.334008 | 0.098888 | 5.85E-204 | darkred | GPR65 |
| 4.48E-204 | 0.556766 | 0.908907 | 0.805264 | 1.04E-203 | darkred | CD99 |
| 9.18E-201 | -0.51017 | 0.931849 | 0.968411 | 2.12E-200 | darkred | EEF2 |
| 1.39E-198 | -0.89211 | 0.239541 | 0.659347 | 3.20E-198 | darkred | TKT |
| 3.42E-198 | 0.556498 | 0.923077 | 0.868273 | 7.79E-198 | darkred | UBB |
| 9.98E-198 | -0.68593 | 0.434548 | 0.802865 | 2.26E-197 | darkred | EIF3E |
| 3.04E-197 | -0.74393 | 0.290823 | 0.722964 | 6.85E-197 | darkred | NAP1L1 |
| 1.82E-195 | -0.85939 | 0.239541 | 0.637386 | 4.08E-195 | darkred | TALDO1 |
| 4.42E-194 | 0.767312 | 0.877868 | 0.789419 | 9.83E-194 | darkred | TXNIP |
| 2.03E-192 | -0.80732 | 0.656545 | 0.879624 | 4.48E-192 | darkred | HSP90AB1 |
| 4.82E-191 | 0.666156 | 0.635628 | 0.303895 | 1.06E-190 | darkred | LCK |
| 1.04E-189 | 0.533097 | 0.908232 | 0.876685 | 2.26E-189 | darkred | YWHAZ |
| 1.14E-186 | -0.64798 | 0.107962 | 0.556404 | 2.48E-186 | darkred | RGS10 |
| 4.95E-186 | 0.761198 | 0.68556 | 0.504679 | 1.07E-185 | darkred | PAXX |
| 1.66E-184 | -0.8624 | 0.095142 | 0.519139 | 3.56E-184 | darkred | HLA-DMA |
| 1.92E-184 | -0.71441 | 0.044534 | 0.456941 | 4.08E-184 | darkred | TMEM14C |
| 1.33E-182 | 0.750184 | 0.732119 | 0.528194 | 2.81E-182 | darkred | FLNA |

| | | | | | | |
|---|---|---|---|---|---|---|
| 4.28E-182 | 0.739182 | 0.746964 | 0.650258 | 9.00E-182 | darkred | MSN |
| 1.85E-180 | -0.86636 | 0.178138 | 0.598061 | 3.87E-180 | darkred | PRDX1 |
| 2.72E-180 | 0.683179 | 0.810391 | 0.724653 | 5.64E-180 | darkred | LCP1 |
| 6.61E-180 | 0.840213 | 0.425101 | 0.187574 | 1.36E-179 | darkred | C1orf21 |
| 2.25E-178 | -0.62202 | 0.068151 | 0.485827 | 4.61E-178 | darkred | SYPL1 |
| 1.40E-175 | 0.86788 | 0.55803 | 0.316869 | 2.85E-175 | darkred | SELPLG |
| 8.42E-175 | -0.62294 | 0.010796 | 0.388121 | 1.71E-174 | darkred | BEX3 |
| 1.39E-174 | 0.745838 | 0.42915 | 0.178722 | 2.81E-174 | darkred | ITGAL |
| 2.40E-174 | 0.698194 | 0.298246 | 0.093415 | 4.82E-174 | darkred | CD244 |
| 3.43E-174 | 0.745596 | 0.657895 | 0.417784 | 6.84E-174 | darkred | HLA-F |
| 2.89E-172 | -0.8863 | 0.007422 | 0.377209 | 5.73E-172 | darkred | ITM2C |
| 5.59E-172 | -0.65032 | 0.345479 | 0.736038 | 1.10E-171 | darkred | ANP32B |
| 1.86E-171 | -0.98738 | 0.021592 | 0.398426 | 3.65E-171 | darkred | LST1 |
| 1.23E-170 | -1.3668 | 0.122807 | 0.495523 | 2.41E-170 | darkred | STMN1 |
| 1.77E-170 | -0.84804 | 0.120108 | 0.507416 | 3.44E-170 | darkred | IMPDH2 |
| 3.42E-170 | 0.781168 | 0.682186 | 0.516436 | 6.60E-170 | darkred | SLC9A3R1 |
| 5.62E-170 | 0.655312 | 0.748988 | 0.656982 | 1.08E-169 | darkred | 7-Sep |
| 2.67E-168 | -0.67755 | 0.136977 | 0.56583 | 5.09E-168 | darkred | PLP2 |
| 1.11E-167 | 0.824745 | 0.568151 | 0.353863 | 2.10E-167 | darkred | OSTF1 |
| 2.91E-166 | 0.809449 | 0.377868 | 0.147539 | 5.48E-166 | darkred | SERTAD1 |
| 2.92E-166 | 0.684064 | 0.428475 | 0.174499 | 5.48E-166 | darkred | FYN |
| 1.36E-165 | -1.29039 | 0.291498 | 0.658536 | 2.54E-165 | darkred | HLA-DRB1 |
| 2.43E-162 | 0.682182 | 0.487179 | 0.231055 | 4.51E-162 | darkred | SKAP1 |
| 7.95E-161 | 0.749172 | 0.497301 | 0.248319 | 1.47E-160 | darkred | MYO1F |
| 5.85E-159 | -0.98041 | 0.044534 | 0.410115 | 1.08E-158 | darkred | SOX4 |
| 1.92E-157 | 0.702172 | 0.369096 | 0.146863 | 3.51E-157 | darkred | PTPN4 |
| 2.52E-156 | 0.824026 | 0.597841 | 0.425521 | 4.58E-156 | darkred | CTSC |
| 8.22E-155 | 0.71777 | 0.321188 | 0.117605 | 1.49E-154 | darkred | CEP78 |
| 1.20E-154 | 0.727354 | 0.551957 | 0.294605 | 2.17E-154 | darkred | ID2 |
| 8.23E-154 | -0.74134 | 0.165992 | 0.548363 | 1.47E-153 | darkred | HSPD1 |
| 2.76E-153 | 0.731099 | 0.643725 | 0.478158 | 4.91E-153 | darkred | WIPF1 |
| 4.77E-152 | -0.5926 | 0.020243 | 0.367884 | 8.45E-152 | darkred | CD82 |
| 5.83E-152 | 0.763788 | 0.415655 | 0.189196 | 1.03E-151 | darkred | APOBEC3G |
| 2.14E-151 | -0.50902 | 0.138327 | 0.558465 | 3.75E-151 | darkred | TMEM123 |
| 7.19E-151 | 0.520718 | 0.904858 | 0.816278 | 1.26E-150 | darkred | CORO1A |
| 2.12E-150 | -0.53832 | 0.000675 | 0.331802 | 3.69E-150 | darkred | ARMH1 |
| 1.08E-149 | -0.59855 | 0.004723 | 0.337782 | 1.87E-149 | darkred | GNA15 |
| 4.04E-149 | 0.675137 | 0.740891 | 0.646035 | 6.94E-149 | darkred | LY6E |
| 3.68E-148 | -0.9852 | 0.037112 | 0.38481 | 6.30E-148 | darkred | HLA-DQB1 |
| 1.34E-146 | -0.55867 | 0.076248 | 0.45211 | 2.28E-146 | darkred | RILPL2 |
| 3.76E-146 | -0.52973 | 0.111336 | 0.504814 | 6.38E-146 | darkred | PPA1 |

| | | | | | | |
|---|---|---|---|---|---|---|
| 2.89E-145 | 0.706086 | 0.409582 | 0.187506 | 4.88E-145 | darkred | PXN |
| 3.72E-145 | -0.66527 | 0.137652 | 0.516301 | 6.25E-145 | darkred | H2AFY |
| 2.10E-144 | 0.662079 | 0.458165 | 0.209331 | 3.52E-144 | darkred | GIMAP7 |
| 4.48E-144 | -0.53881 | 0.271255 | 0.679618 | 7.46E-144 | darkred | RSL1D1 |
| 2.07E-143 | -0.65714 | 0.00135 | 0.321159 | 3.43E-143 | darkred | EGFL7 |
| 2.24E-141 | -1.38171 | 0.087045 | 0.411973 | 3.70E-141 | darkred | LTB |
| 4.26E-141 | 0.799654 | 0.382591 | 0.17683 | 6.99E-141 | darkred | MATK |
| 7.31E-141 | 0.76615 | 0.593792 | 0.362005 | 1.19E-140 | darkred | MAP3K8 |
| 9.24E-140 | -0.58561 | 0.152497 | 0.541437 | 1.50E-139 | darkred | HSPB1 |
| 3.67E-139 | 0.713686 | 0.553306 | 0.335687 | 5.93E-139 | darkred | PLEK |
| 3.76E-139 | 0.721043 | 0.570175 | 0.359269 | 6.06E-139 | darkred | RNF213 |
| 8.57E-139 | 0.652863 | 0.325236 | 0.123754 | 1.37E-138 | darkred | EFHD2 |
| 1.29E-138 | -0.77556 | 0.049933 | 0.38329 | 2.06E-138 | darkred | AC084033.3 |
| 5.53E-138 | -0.54839 | 0.334008 | 0.70053 | 8.79E-138 | darkred | PEBP1 |
| 8.24E-138 | -0.65562 | 0.048583 | 0.38731 | 1.30E-137 | darkred | FABP5 |
| 5.90E-137 | 0.815072 | 0.418354 | 0.204635 | 9.28E-137 | darkred | BHLHE40 |
| 1.47E-135 | -1.56269 | 0.08502 | 0.424136 | 2.30E-135 | darkred | CST3 |
| 4.79E-135 | 0.807879 | 0.462213 | 0.266327 | 7.48E-135 | darkred | CHST12 |
| 5.03E-135 | -0.53742 | 0.043185 | 0.379337 | 7.81E-135 | darkred | RNF130 |
| 1.84E-132 | 0.697347 | 0.526991 | 0.312646 | 2.85E-132 | darkred | ADGRE5 |
| 7.14E-132 | 0.61183 | 0.692308 | 0.518936 | 1.10E-131 | darkred | CCND3 |
| 5.44E-130 | -0.57192 | 0.020243 | 0.331599 | 8.33E-130 | darkred | CAPG |
| 8.57E-130 | 0.581773 | 0.334683 | 0.133214 | 1.31E-129 | darkred | STAT4 |
| 1.18E-129 | -0.58706 | 0.00135 | 0.297544 | 1.79E-129 | darkred | MGST1 |
| 4.97E-129 | -0.65362 | 0.124831 | 0.472651 | 7.51E-129 | darkred | CAT |
| 4.40E-128 | 0.680187 | 0.6417 | 0.490354 | 6.63E-128 | darkred | ANXA6 |
| 2.58E-127 | -0.62113 | 0.003374 | 0.296665 | 3.86E-127 | darkred | FAM30A |
| 4.14E-127 | 0.649688 | 0.607962 | 0.41917 | 6.18E-127 | darkred | TBC1D10C |
| 6.98E-127 | -0.5995 | 0.031714 | 0.344505 | 1.04E-126 | darkred | FCGRT |
| 1.69E-126 | -0.62787 | 0.002024 | 0.293152 | 2.50E-126 | darkred | CD34 |
| 1.15E-125 | -0.50375 | 0.291498 | 0.681442 | 1.69E-125 | darkred | SNHG8 |
| 3.67E-125 | 0.597941 | 0.730094 | 0.675631 | 5.39E-125 | darkred | PDIA3 |
| 5.09E-121 | -0.67555 | 0.005398 | 0.288962 | 7.45E-121 | darkred | CSF3R |
| 3.46E-119 | -0.51774 | 0.020918 | 0.313592 | 5.04E-119 | darkred | MEF2C |
| 6.50E-119 | 0.539554 | 0.721997 | 0.674617 | 9.42E-119 | darkred | PPP1CA |
| 6.66E-119 | 0.659748 | 0.352227 | 0.153721 | 9.63E-119 | darkred | CEMIP2 |
| 7.43E-117 | -0.61293 | 0.2861 | 0.602655 | 1.07E-116 | darkred | TXN |
| 8.02E-116 | 0.537591 | 0.311741 | 0.121896 | 1.15E-115 | darkred | RORA |
| 1.78E-113 | 0.631795 | 0.585695 | 0.412176 | 2.54E-113 | darkred | IQGAP1 |
| 1.84E-113 | 0.529302 | 0.34413 | 0.143654 | 2.61E-113 | darkred | LINC00861 |
| 1.52E-111 | 0.582673 | 0.695007 | 0.612386 | 2.16E-111 | darkred | PSMB9 |

| | | | | | | |
|---|---|---|---|---|---|---|
| 2.41E-111 | -0.73757 | 0.007422 | 0.272948 | 3.40E-111 | darkred | SMIM24 |
| 1.99E-110 | -1.39872 | 0.005398 | 0.268117 | 2.80E-110 | darkred | IGLL1 |
| 1.26E-109 | -0.91374 | 0.065452 | 0.349471 | 1.75E-109 | darkred | PRSS57 |
| 1.26E-109 | -1.29926 | 0.033063 | 0.302949 | 1.75E-109 | darkred | IL7R |
| 4.88E-109 | 0.623818 | 0.398785 | 0.207068 | 6.78E-109 | darkred | SUN2 |
| 2.72E-107 | -0.59531 | 0.008097 | 0.268692 | 3.76E-107 | darkred | H1F0 |
| 6.76E-107 | 0.5993 | 0.255061 | 0.095882 | 9.30E-107 | darkred | GK5 |
| 1.35E-106 | 0.656567 | 0.630904 | 0.525997 | 1.86E-106 | darkred | CD53 |
| 1.69E-106 | 0.689391 | 0.5 | 0.3292 | 2.31E-106 | darkred | CDC42SE1 |
| 7.85E-106 | 0.732104 | 0.402834 | 0.223926 | 1.07E-105 | darkred | JAK1 |
| 2.22E-105 | 0.659632 | 0.604588 | 0.483665 | 3.01E-105 | darkred | PRMT2 |
| 2.86E-105 | 0.632991 | 0.364372 | 0.176459 | 3.86E-105 | darkred | DENND2D |
| 9.86E-105 | -0.62301 | 0.175439 | 0.500591 | 1.33E-104 | darkred | GLUL |
| 1.06E-104 | -0.82222 | 0.988529 | 0.996892 | 1.42E-104 | darkred | FTH1 |
| 1.70E-104 | 0.656672 | 0.47166 | 0.287239 | 2.27E-104 | darkred | AKNA |
| 1.78E-104 | -0.59416 | 0.082996 | 0.383155 | 2.36E-104 | darkred | GRN |
| 7.51E-104 | 0.656862 | 0.455466 | 0.27359 | 9.94E-104 | darkred | RASSF5 |
| 4.68E-103 | -0.88473 | 0.011471 | 0.263387 | 6.18E-103 | darkred | SPINK2 |
| 1.50E-101 | -0.50198 | 0.017544 | 0.274266 | 1.98E-101 | darkred | TCF4 |
| 9.29E-101 | 0.66659 | 0.602564 | 0.477955 | 1.22E-100 | darkred | IFITM3 |
| 5.54E-100 | -0.75441 | 0.049258 | 0.320349 | 7.23E-100 | darkred | HLA-DQA1 |
| 9.73E-100 | -0.58617 | 0.134278 | 0.443697 | 1.26E-99 | darkred | FOSB |
| 9.16E-99 | 0.669587 | 0.504049 | 0.356769 | 1.19E-98 | darkred | SYTL1 |
| 9.67E-99 | -0.77357 | 0.061404 | 0.332342 | 1.25E-98 | darkred | CD3D |
| 3.95E-98 | 0.756591 | 0.482456 | 0.337647 | 5.08E-98 | darkred | HSH2D |
| 1.06E-97 | 0.654653 | 0.525641 | 0.375553 | 1.36E-97 | darkred | PPP1R18 |
| 6.56E-97 | 0.532423 | 0.458165 | 0.256191 | 8.37E-97 | darkred | LBH |
| 7.03E-96 | 0.650843 | 0.473684 | 0.306531 | 8.94E-96 | darkred | SLC44A2 |
| 1.68E-94 | 0.594393 | 0.624831 | 0.571438 | 2.14E-94 | darkred | BZW1 |
| 2.12E-94 | -0.56141 | 0.232119 | 0.55333 | 2.68E-94 | darkred | DUT |
| 3.57E-94 | 0.592831 | 0.603239 | 0.492314 | 4.49E-94 | darkred | PLAC8 |
| 3.19E-92 | 0.608056 | 0.475709 | 0.3167 | 4.00E-92 | darkred | RHOF |
| 1.69E-91 | -0.51959 | 0.130229 | 0.414305 | 2.11E-91 | darkred | CDK6 |
| 4.32E-91 | 0.620804 | 0.552632 | 0.454272 | 5.37E-91 | darkred | DHRS7 |
| 2.39E-90 | 0.557914 | 0.628205 | 0.519409 | 2.97E-90 | darkred | CYBA |
| 1.07E-87 | 0.653543 | 0.472335 | 0.32447 | 1.32E-87 | darkred | SASH3 |
| 9.46E-86 | 0.642607 | 0.443995 | 0.279165 | 1.16E-85 | darkred | PPP2R5C |
| 1.28E-82 | -0.6926 | 0.037112 | 0.264333 | 1.57E-82 | darkred | NCF1 |
| 4.78E-82 | 0.636649 | 0.417679 | 0.263218 | 5.85E-82 | darkred | UTRN |
| 1.46E-81 | 0.51488 | 0.661943 | 0.629312 | 1.77E-81 | darkred | CAP1 |
| 1.52E-81 | 0.624346 | 0.356275 | 0.198486 | 1.85E-81 | darkred | BTN3A2 |

| | | | | | | |
|---|---|---|---|---|---|---|
| 2.70E-81 | 0.539736 | 0.645749 | 0.618737 | 3.26E-81 | darkred | TAPBP |
| 7.15E-81 | 0.511899 | 0.623482 | 0.590189 | 8.62E-81 | darkred | BSG |
| 2.75E-80 | 0.591059 | 0.558704 | 0.465894 | 3.30E-80 | darkred | CD63 |
| 3.86E-80 | 0.601791 | 0.612011 | 0.583331 | 4.63E-80 | darkred | PSMB10 |
| 6.79E-78 | -0.55434 | 0.068151 | 0.307341 | 8.11E-78 | darkred | NAMPT |
| 7.21E-78 | -0.51406 | 0.067476 | 0.306193 | 8.58E-78 | darkred | CLEC11A |
| 7.42E-78 | 0.54112 | 0.267881 | 0.123349 | 8.80E-78 | darkred | AC012615.1 |
| 1.08E-77 | -0.60252 | 0.976383 | 0.970269 | 1.27E-77 | darkred | GAPDH |
| 2.36E-77 | -0.59384 | 0.04251 | 0.262272 | 2.77E-77 | darkred | CD3G |
| 7.62E-77 | -0.50234 | 0.304318 | 0.57029 | 8.94E-77 | darkred | NUCB2 |
| 2.68E-76 | 0.686801 | 0.509447 | 0.352073 | 3.13E-76 | darkred | NR4A2 |
| 1.83E-74 | 0.532714 | 0.261134 | 0.121187 | 2.13E-74 | darkred | DIP2A |
| 2.17E-74 | 0.634581 | 0.394737 | 0.244299 | 2.52E-74 | darkred | PDE4D |
| 1.31E-73 | 0.536218 | 0.709177 | 0.671779 | 1.52E-73 | darkred | HSPA5 |
| 1.09E-72 | 0.652271 | 0.576923 | 0.469813 | 1.26E-72 | darkred | MT2A |
| 5.51E-72 | 0.514288 | 0.620108 | 0.5486 | 6.34E-72 | darkred | ARHGEF1 |
| 3.37E-70 | 0.609496 | 0.523617 | 0.457347 | 3.87E-70 | darkred | FAM49B |
| 4.20E-69 | -0.80198 | 0.388664 | 0.636744 | 4.80E-69 | darkred | TUBB |
| 7.41E-69 | -2.37006 | 0.059379 | 0.260549 | 8.45E-69 | darkred | LYZ |
| 5.96E-62 | 0.5614 | 0.381242 | 0.251191 | 6.77E-62 | darkred | STK10 |
| 2.28E-61 | 0.532516 | 0.340081 | 0.204602 | 2.58E-61 | darkred | OPTN |
| 1.14E-60 | 0.563841 | 0.398111 | 0.275888 | 1.28E-60 | darkred | TAP1 |
| 5.77E-60 | 0.553403 | 0.579622 | 0.539545 | 6.49E-60 | darkred | RSRP1 |
| 3.10E-59 | 0.551718 | 0.311741 | 0.184905 | 3.48E-59 | darkred | TSEN54 |
| 1.04E-58 | 0.532645 | 0.252362 | 0.131288 | 1.17E-58 | darkred | UBE2F |
| 1.49E-58 | 0.626477 | 0.337382 | 0.219568 | 1.66E-58 | darkred | MCTP2 |
| 1.73E-58 | 0.59069 | 0.262483 | 0.140613 | 1.92E-58 | darkred | GTF3C1 |
| 2.06E-58 | 0.595304 | 0.437922 | 0.333491 | 2.28E-58 | darkred | CDK2AP2 |
| 2.51E-58 | 0.521207 | 0.35695 | 0.224197 | 2.78E-58 | darkred | GLIPR2 |
| 4.49E-57 | -0.71261 | 0.448043 | 0.661036 | 4.95E-57 | darkred | HLA-DPA1 |
| 4.78E-56 | 0.637438 | 0.381916 | 0.275617 | 5.24E-56 | darkred | STOM |
| 8.58E-56 | 0.627393 | 0.390688 | 0.289469 | 9.38E-56 | darkred | G6PD |
| 9.40E-56 | 0.56363 | 0.445344 | 0.34937 | 1.02E-55 | darkred | SORL1 |
| 3.02E-55 | 0.61041 | 0.406208 | 0.30849 | 3.29E-55 | darkred | ARPC5L |
| 1.36E-54 | -0.5356 | 0.417004 | 0.61948 | 1.47E-54 | darkred | SERPINB1 |
| 1.76E-54 | -0.57975 | 0.182186 | 0.421298 | 1.90E-54 | darkred | TIMP1 |
| 1.96E-53 | -1.23069 | 0.91498 | 0.919085 | 2.11E-53 | darkred | CD74 |
| 2.37E-53 | 0.545096 | 0.487854 | 0.411129 | 2.54E-53 | darkred | CLEC2B |
| 5.55E-52 | 0.537124 | 0.502024 | 0.453292 | 5.94E-52 | darkred | SPSB3 |
| 2.70E-51 | -0.62875 | 0.407557 | 0.625156 | 2.88E-51 | darkred | H2AFZ |
| 7.37E-50 | 0.575859 | 0.426451 | 0.348289 | 7.84E-50 | darkred | ACTN4 |

| | | | | | | |
|---|---|---|---|---|---|---|
| 3.96E-49 | 0.543701 | 0.374494 | 0.269029 | 4.20E-49 | darkred | MBP |
| 3.64E-48 | 0.551654 | 0.405533 | 0.300889 | 3.85E-48 | darkred | LYAR |
| 7.41E-48 | 0.583405 | 0.429825 | 0.358897 | 7.82E-48 | darkred | MAPRE2 |
| 1.32E-47 | 0.517293 | 0.326586 | 0.219433 | 1.39E-47 | darkred | RPS6KA1 |
| 2.24E-45 | 0.509421 | 0.532389 | 0.427278 | 2.35E-45 | darkred | GADD45B |
| 1.19E-43 | 0.530094 | 0.320513 | 0.208081 | 1.24E-43 | darkred | TENT5C |
| 2.42E-43 | 0.548008 | 0.397436 | 0.311497 | 2.52E-43 | darkred | CFLAR |
| 1.88E-42 | 0.504826 | 0.410256 | 0.311902 | 1.96E-42 | darkred | RNF125 |
| 5.14E-40 | 0.529428 | 0.323887 | 0.231866 | 5.32E-40 | darkred | LASP1 |
| 1.79E-39 | -0.54131 | 0.432524 | 0.698368 | 1.85E-39 | darkred | TSPO |
| 6.42E-38 | -0.82566 | 0.539811 | 0.668401 | 6.60E-38 | darkred | TUBA1B |
| 2.86E-37 | -0.57693 | 0.367072 | 0.606743 | 2.93E-37 | darkred | HIST1H4C |
| 3.52E-35 | 0.605323 | 0.297571 | 0.217676 | 3.60E-35 | darkred | AKR1C3 |
| 1.70E-33 | 0.525218 | 0.312416 | 0.22494 | 1.73E-33 | darkred | IFNGR1 |
| 9.34E-32 | -0.51125 | 0.279352 | 0.407446 | 9.50E-32 | darkred | CD3E |
| 1.85E-30 | -0.63038 | 0.991228 | 0.994257 | 1.87E-30 | darkred | FTL |
| 1.77E-22 | 0.500164 | 0.292173 | 0.235751 | 1.79E-22 | darkred | CD38 |
| 2.33E-16 | 0.500423 | 0.367746 | 0.345451 | 2.35E-16 | darkred | AC007952.4 |
| 2.01E-14 | -0.67099 | 0.454116 | 0.631542 | 2.02E-14 | darkred | SAT1 |
| 2.02E-07 | -1.11334 | 0.261134 | 0.212102 | 2.03E-07 | darkred | HBB |
| 0.000285 | -0.56197 | 0.807018 | 0.824622 | 0.000285 | darkred | S100A6 |
| 0 | 1.244311 | 0.790847 | 0.149945 | 0 | lightcyan | TCF7 |
| 0 | 1.163169 | 0.713691 | 0.110068 | 0 | lightcyan | CCR7 |
| 0 | 1.100863 | 0.695159 | 0.115624 | 0 | lightcyan | LEF1 |
| 0 | 1.070133 | 0.853253 | 0.476984 | 0 | lightcyan | NOSIP |
| 0 | 1.060064 | 0.877458 | 0.235468 | 0 | lightcyan | IL7R |
| 0 | 1.028872 | 0.995083 | 0.346134 | 0 | lightcyan | CD3E |
| 0 | 1.003597 | 0.33056 | 0.042621 | 0 | lightcyan | LINC02446 |
| 0 | 0.997138 | 0.441755 | 0.123642 | 0 | lightcyan | CD8B |
| 0 | 0.950281 | 0.784418 | 0.24556 | 0 | lightcyan | PIK3IP1 |
| 0 | 0.907634 | 0.713313 | 0.213665 | 0 | lightcyan | FLT3LG |
| 0 | 0.891548 | 0.558245 | 0.086296 | 0 | lightcyan | MAL |
| 0 | 0.877628 | 0.88767 | 0.266589 | 0 | lightcyan | CD3D |
| 0 | 0.875578 | 0.619894 | 0.166508 | 0 | lightcyan | OXNAD1 |
| 0 | 0.872268 | 0.613843 | 0.127862 | 0 | lightcyan | CD27 |
| 0 | 0.862842 | 0.609304 | 0.121813 | 0 | lightcyan | CAMK4 |
| 0 | 0.855803 | 0.996218 | 0.897739 | 0 | lightcyan | SARAF |
| 0 | 0.849646 | 0.549168 | 0.140591 | 0 | lightcyan | LDLRAP1 |
| 0 | 0.843637 | 0.501513 | 0.075535 | 0 | lightcyan | NELL2 |
| 0 | 0.835631 | 0.923979 | 0.347435 | 0 | lightcyan | LTB |
| 0 | 0.817384 | 0.857035 | 0.526568 | 0 | lightcyan | LEPROTL1 |

| | | | | | | |
|---|---|---|---|---|---|---|
| 0 | 0.811596 | 0.899395 | 0.364384 | 0 | lightcyan | CD7 |
| 0 | 0.780221 | 0.879349 | 0.460878 | 0 | lightcyan | FYB1 |
| 0 | 0.744576 | 0.577912 | 0.225375 | 0 | lightcyan | ABLIM1 |
| 0 | 0.738697 | 0.798411 | 0.510532 | 0 | lightcyan | RGS10 |
| 0 | 0.736115 | 0.613086 | 0.246404 | 0 | lightcyan | MGAT4A |
| 0 | 0.731382 | 0.720877 | 0.208179 | 0 | lightcyan | CD3G |
| 0 | 0.729713 | 0.50416 | 0.120582 | 0 | lightcyan | LINC00861 |
| 0 | 0.720899 | 0.783283 | 0.276611 | 0 | lightcyan | LCK |
| 0 | 0.71813 | 0.916793 | 0.554805 | 0 | lightcyan | EVL |
| 0 | 0.703076 | 0.947806 | 0.754826 | 0 | lightcyan | JUNB |
| 0 | 0.698167 | 0.405825 | 0.059254 | 0 | lightcyan | TRABD2A |
| 0 | 0.659078 | 0.512103 | 0.198122 | 0 | lightcyan | DGKA |
| 0 | 0.646591 | 0.677005 | 0.364173 | 0 | lightcyan | TRAF3IP3 |
| 0 | 0.642883 | 0.721256 | 0.467876 | 0 | lightcyan | PRMT2 |
| 0 | 0.641679 | 0.975794 | 0.535464 | 0 | lightcyan | IFITM1 |
| 0 | 0.625872 | 0.783661 | 0.469177 | 0 | lightcyan | AES |
| 0 | 0.613213 | 0.700076 | 0.26666 | 0 | lightcyan | SPOCK2 |
| 0 | 0.594786 | 0.442133 | 0.109892 | 0 | lightcyan | ITK |
| 0 | 0.592904 | 0.581694 | 0.236453 | 0 | lightcyan | LBH |
| 0 | 0.587585 | 0.594554 | 0.236277 | 0 | lightcyan | LAT |
| 0 | 0.574413 | 0.928896 | 0.768893 | 0 | lightcyan | C12orf57 |
| 0 | 0.570432 | 0.999622 | 0.994127 | 0 | lightcyan | RPS27 |
| 0 | 0.538076 | 0.365734 | 0.07768 | 0 | lightcyan | BCL11B |
| 0 | 0.505251 | 0.283283 | 0.069733 | 0 | lightcyan | EPHX2 |
| 0 | 0.503297 | 0.361952 | 0.09674 | 0 | lightcyan | AQP3 |
| 0 | 0.502879 | 0.274962 | 0.036783 | 0 | lightcyan | LINC01550 |
| 0 | 0.50003 | 0.462935 | 0.147203 | 0 | lightcyan | CD5 |
| 0 | -0.69695 | 0.638427 | 0.851215 | 0 | lightcyan | TAGLN2 |
| 0 | -0.75355 | 0.509077 | 0.838204 | 0 | lightcyan | CD99 |
| 0 | -0.76151 | 0.056732 | 0.473011 | 0 | lightcyan | APLP2 |
| 0 | -0.86943 | 0.871785 | 0.95745 | 0 | lightcyan | H3F3A |
| 0 | -0.87125 | 0.00643 | 0.470338 | 0 | lightcyan | ZEB2 |
| 0 | -0.90594 | 0.84531 | 0.927032 | 0 | lightcyan | HMGB1 |
| 0 | -0.94668 | 0.286309 | 0.649295 | 0 | lightcyan | TALDO1 |
| 0 | -0.98311 | 0.009077 | 0.406899 | 0 | lightcyan | MAP3K8 |
| 0 | -1.06643 | 0.296142 | 0.638991 | 0 | lightcyan | SERPINB1 |
| 0 | -1.07835 | 0.041982 | 0.526286 | 0 | lightcyan | GLUL |
| 0 | -1.08858 | 0.163011 | 0.627035 | 0 | lightcyan | TXN |
| 0 | -1.08966 | 0.041982 | 0.541407 | 0 | lightcyan | HLA-DMA |
| 0 | -1.1389 | 0.01059 | 0.401484 | 0 | lightcyan | HLA-DQB1 |
| 0 | -1.1659 | 0.380484 | 0.834758 | 0 | lightcyan | CLIC1 |

| | | | | | | |
|---|---|---|---|---|---|---|
| 0 | -1.18944 | 0.594175 | 0.849246 | 0 | lightcyan | SRGN |
| 0 | -1.31181 | 0.94289 | 0.973134 | 0 | lightcyan | GAPDH |
| 0 | -1.32453 | 0.396747 | 0.769033 | 0 | lightcyan | GSTP1 |
| 0 | -1.59614 | 0.151664 | 0.699335 | 0 | lightcyan | HLA-DPB1 |
| 0 | -1.81629 | 0.087368 | 0.703274 | 0 | lightcyan | HLA-DPA1 |
| 0 | -1.84183 | 0.010212 | 0.444948 | 0 | lightcyan | CST3 |
| 0 | -1.93545 | 0.009455 | 0.635088 | 0 | lightcyan | HLA-DRB5 |
| 0 | -2.31041 | 0.019667 | 0.698808 | 0 | lightcyan | HLA-DRB1 |
| 0 | -2.69179 | 0.678517 | 0.941239 | 0 | lightcyan | CD74 |
| 0 | -2.78464 | 0.025719 | 0.65573 | 0 | lightcyan | HLA-DRA |
| 6.15E-307 | -0.78993 | 0.142209 | 0.53135 | 1.74E-306 | lightcyan | H2AFY |
| 6.22E-307 | 0.554458 | 0.757564 | 0.418785 | 1.74E-306 | lightcyan | CD48 |
| 1.35E-304 | -0.83069 | 0.006051 | 0.377677 | 3.73E-304 | lightcyan | PLEK |
| 1.60E-304 | -0.73174 | 0.207262 | 0.621901 | 4.35E-304 | lightcyan | ATP6V0B |
| 2.12E-302 | -0.65944 | 0.746218 | 0.873123 | 5.68E-302 | lightcyan | HSP90AA1 |
| 4.48E-302 | -0.93573 | 0.996974 | 0.996448 | 1.19E-301 | lightcyan | FTH1 |
| 1.52E-300 | 0.519327 | 0.366112 | 0.116327 | 3.98E-300 | lightcyan | PDE3B |
| 6.34E-300 | 0.51294 | 0.934191 | 0.780568 | 1.64E-299 | lightcyan | TXNIP |
| 1.73E-298 | -1.23467 | 0.003026 | 0.366881 | 4.42E-298 | lightcyan | PRSS57 |
| 1.82E-298 | 0.512115 | 0.610439 | 0.264585 | 4.59E-298 | lightcyan | FCMR |
| 1.97E-298 | -0.67513 | 0.386914 | 0.727046 | 4.89E-298 | lightcyan | SEC61B |
| 1.07E-294 | -0.83542 | 0.366112 | 0.664733 | 2.63E-294 | lightcyan | TKT |
| 2.50E-294 | -0.71208 | 0.001513 | 0.361958 | 6.06E-294 | lightcyan | LAT2 |
| 2.56E-293 | -0.96957 | 0.045008 | 0.42501 | 6.14E-293 | lightcyan | SOX4 |
| 1.53E-289 | -0.63708 | 0.033661 | 0.407357 | 3.63E-289 | lightcyan | DDAH2 |
| 4.50E-285 | -0.63393 | 0.000378 | 0.351795 | 1.06E-284 | lightcyan | GNA15 |
| 1.90E-284 | -0.53205 | 0.800681 | 0.924887 | 4.40E-284 | lightcyan | ARPC3 |
| 7.94E-284 | -0.66103 | 0.108548 | 0.503956 | 1.82E-283 | lightcyan | CD63 |
| 1.88E-275 | -0.64811 | 0.529879 | 0.759503 | 4.25E-275 | lightcyan | TPI1 |
| 2.92E-275 | -0.71301 | 0.128971 | 0.51644 | 6.54E-275 | lightcyan | PYCARD |
| 2.37E-273 | -2.02412 | 0.010968 | 0.354538 | 5.25E-273 | lightcyan | TYROBP |
| 1.75E-271 | 0.506983 | 0.496974 | 0.195555 | 3.84E-271 | lightcyan | GIMAP7 |
| 5.83E-271 | -0.74556 | 0.04652 | 0.401343 | 1.26E-270 | lightcyan | FABP5 |
| 1.01E-270 | -0.72403 | 0.068835 | 0.436509 | 2.18E-270 | lightcyan | TUBB4B |
| 1.69E-270 | -0.62013 | 0.354387 | 0.662306 | 3.59E-270 | lightcyan | COX5A |
| 8.12E-268 | -0.6197 | 0.403933 | 0.718325 | 1.71E-267 | lightcyan | GNG5 |
| 9.50E-265 | -0.53425 | 0.001513 | 0.334916 | 1.98E-264 | lightcyan | SYK |
| 3.01E-264 | -0.67624 | 0.001513 | 0.334212 | 6.19E-264 | lightcyan | EGFL7 |
| 1.22E-263 | -1.56183 | 0.065809 | 0.408798 | 2.49E-263 | lightcyan | LGALS1 |
| 3.22E-262 | -0.9463 | 0.003782 | 0.335654 | 6.50E-262 | lightcyan | HLA-DQA1 |
| 2.82E-260 | -0.5782 | 0.125567 | 0.501248 | 5.65E-260 | lightcyan | COMT |

| | | | | | | |
|---|---|---|---|---|---|---|
| 1.05E-258 | -0.60519 | 0.043116 | 0.393079 | 2.08E-258 | lightcyan | RNF130 |
| 1.25E-258 | -1.30203 | 0.439486 | 0.682983 | 2.44E-258 | lightcyan | TUBA1B |
| 2.70E-258 | -0.60292 | 0.093419 | 0.465872 | 5.25E-258 | lightcyan | RILPL2 |
| 6.55E-257 | -0.6081 | 0.001513 | 0.327355 | 1.26E-256 | lightcyan | MEF2C |
| 1.21E-256 | -0.70408 | 0.297277 | 0.678342 | 2.31E-256 | lightcyan | PPP1R15A |
| 2.49E-256 | -0.58809 | 0.126324 | 0.49833 | 4.70E-256 | lightcyan | PDIA6 |
| 4.35E-254 | -0.66021 | 0.540469 | 0.743925 | 8.13E-254 | lightcyan | ENO1 |
| 1.03E-253 | -0.63506 | 0.589637 | 0.765657 | 1.91E-253 | lightcyan | SLC25A5 |
| 2.15E-253 | -0.83253 | 0.060893 | 0.395893 | 3.95E-253 | lightcyan | AC084033.3 |
| 2.12E-248 | -0.80901 | 0.293495 | 0.604494 | 3.85E-248 | lightcyan | PRDX1 |
| 6.55E-247 | 0.551479 | 0.368003 | 0.141998 | 1.18E-246 | lightcyan | SH3YL1 |
| 2.41E-244 | -1.27876 | 0.614599 | 0.830045 | 4.31E-244 | lightcyan | S100A4 |
| 1.60E-243 | -0.59889 | 0.054463 | 0.3954 | 2.85E-243 | lightcyan | TUBA1C |
| 4.32E-243 | 0.554763 | 0.531014 | 0.265534 | 7.60E-243 | lightcyan | ATM |
| 2.53E-241 | -0.63617 | 0.410741 | 0.692689 | 4.41E-241 | lightcyan | VAMP8 |
| 2.63E-240 | -0.73181 | 0.152042 | 0.484334 | 4.54E-240 | lightcyan | CAT |
| 3.06E-240 | -0.60806 | 0.000756 | 0.309702 | 5.24E-240 | lightcyan | MGST1 |
| 9.62E-240 | -0.55701 | 0.00643 | 0.318564 | 1.63E-239 | lightcyan | CYTOR |
| 1.22E-236 | -0.64315 | 0.002269 | 0.308753 | 2.05E-236 | lightcyan | FAM30A |
| 5.35E-236 | -0.57902 | 0.217474 | 0.577628 | 8.95E-236 | lightcyan | LAPTM4A |
| 1.01E-235 | -0.6533 | 0.000756 | 0.305166 | 1.68E-235 | lightcyan | CD34 |
| 5.82E-234 | -0.55522 | 0.83056 | 0.930619 | 9.58E-234 | lightcyan | OAZ1 |
| 2.02E-233 | -0.56778 | 0.0382 | 0.363927 | 3.30E-233 | lightcyan | KLF10 |
| 1.11E-232 | -0.61679 | 0.026475 | 0.343742 | 1.80E-232 | lightcyan | CAPG |
| 1.50E-232 | -0.73507 | 0.06354 | 0.397229 | 2.42E-232 | lightcyan | GRN |
| 4.56E-229 | -0.71161 | 0.002269 | 0.30084 | 7.27E-229 | lightcyan | CSF3R |
| 3.77E-227 | -0.60737 | 0.344175 | 0.639976 | 5.96E-227 | lightcyan | YBX1 |
| 3.91E-227 | -1.5423 | 0.019667 | 0.322502 | 6.14E-227 | lightcyan | CST7 |
| 8.70E-225 | 0.521412 | 0.313918 | 0.109927 | 1.35E-224 | lightcyan | PASK |
| 2.01E-224 | -0.88556 | 0.079047 | 0.408482 | 3.10E-224 | lightcyan | LST1 |
| 2.49E-224 | -1.27263 | 0.479576 | 0.740268 | 3.82E-224 | lightcyan | FOS |
| 7.68E-224 | -0.60921 | 0.086611 | 0.415023 | 1.17E-223 | lightcyan | TPM4 |
| 1.23E-222 | -0.54261 | 0.15053 | 0.490945 | 1.85E-222 | lightcyan | YWHAE |
| 1.12E-221 | 0.5084 | 0.668684 | 0.41875 | 1.68E-221 | lightcyan | 1-Sep |
| 2.53E-221 | -0.54327 | 0.132375 | 0.478953 | 3.77E-221 | lightcyan | PGD |
| 9.07E-219 | -0.60141 | 0.416036 | 0.687203 | 1.34E-218 | lightcyan | RPL22L1 |
| 1.89E-217 | -0.77274 | 0.360817 | 0.693744 | 2.77E-217 | lightcyan | KLF6 |
| 4.24E-216 | -0.5259 | 0.128215 | 0.464254 | 6.18E-216 | lightcyan | IDH2 |
| 8.58E-215 | -0.71501 | 0.409985 | 0.713155 | 1.24E-214 | lightcyan | ANXA1 |
| 1.77E-214 | -0.58061 | 0.002269 | 0.286176 | 2.55E-214 | lightcyan | TCF4 |
| 3.02E-214 | -0.77799 | 0.001513 | 0.284348 | 4.31E-214 | lightcyan | SMIM24 |

| | | | | | | |
|---|---|---|---|---|---|---|
| 3.33E-214 | -0.81216 | 0.030635 | 0.325878 | 4.71E-214 | lightcyan | ANXA2 |
| 1.22E-208 | -2.25959 | 0.039334 | 0.330133 | 1.72E-208 | lightcyan | NKG7 |
| 2.78E-208 | -0.93613 | 0.381619 | 0.63646 | 3.88E-208 | lightcyan | H2AFZ |
| 3.94E-206 | -0.51017 | 0.013238 | 0.296163 | 5.47E-206 | lightcyan | TNFSF13B |
| 9.00E-205 | -1.43429 | 0.004539 | 0.278932 | 1.24E-204 | lightcyan | IGLL1 |
| 8.68E-204 | -0.50531 | 0 | 0.27109 | 1.19E-203 | lightcyan | SPI1 |
| 1.91E-203 | -0.64257 | 0.120651 | 0.426803 | 2.60E-203 | lightcyan | CDK6 |
| 5.77E-203 | -0.76677 | 0.009077 | 0.284348 | 7.80E-203 | lightcyan | HOPX |
| 1.96E-201 | -0.5786 | 0.135023 | 0.464817 | 2.62E-201 | lightcyan | MYADM |
| 6.41E-196 | -0.50254 | 0 | 0.262686 | 8.55E-196 | lightcyan | CYTL1 |
| 2.54E-194 | -1.2577 | 0.257943 | 0.498189 | 3.36E-194 | lightcyan | STMN1 |
| 1.39E-191 | -0.94174 | 0.319213 | 0.633787 | 1.83E-191 | lightcyan | NEAT1 |
| 4.13E-189 | 0.519504 | 0.6059 | 0.394943 | 5.40E-189 | lightcyan | RHOH |
| 3.44E-188 | -0.52764 | 0.714448 | 0.861624 | 4.47E-188 | lightcyan | PPIB |
| 7.22E-186 | -0.59979 | 0.240166 | 0.575764 | 9.32E-186 | lightcyan | NPC2 |
| 1.06E-185 | -0.59957 | 0.015885 | 0.278616 | 1.37E-185 | lightcyan | H1F0 |
| 2.60E-185 | -1.19223 | 0.339259 | 0.649471 | 3.32E-185 | lightcyan | SAT1 |
| 3.23E-183 | -0.5902 | 0.02118 | 0.282414 | 4.09E-183 | lightcyan | MYO1F |
| 4.82E-183 | -0.62509 | 0.001135 | 0.250589 | 6.07E-183 | lightcyan | HLA-DQA2 |
| 5.89E-182 | -0.53047 | 0.259077 | 0.572388 | 7.37E-182 | lightcyan | P4HB |
| 3.45E-181 | -0.50553 | 0.1823 | 0.490664 | 4.29E-181 | lightcyan | DYNLL1 |
| 8.39E-180 | -0.5491 | 0.234493 | 0.55157 | 1.04E-179 | lightcyan | CYBA |
| 8.45E-180 | -0.71151 | 0.181165 | 0.479868 | 1.04E-179 | lightcyan | AHNAK |
| 2.25E-177 | -0.52808 | 0.147126 | 0.440447 | 2.75E-177 | lightcyan | NASP |
| 3.91E-176 | -0.6526 | 0.0382 | 0.301157 | 4.74E-176 | lightcyan | BLVRB |
| 1.20E-160 | -0.5194 | 0.109304 | 0.392271 | 1.45E-160 | lightcyan | JUND |
| 2.66E-157 | -0.51656 | 0.08472 | 0.345325 | 3.19E-157 | lightcyan | MCM7 |
| 5.23E-157 | -0.67122 | 0.797655 | 0.861483 | 6.24E-157 | lightcyan | HMGN2 |
| 4.70E-156 | -0.82256 | 0.17171 | 0.446531 | 5.58E-156 | lightcyan | DUSP1 |
| 2.49E-153 | -0.60482 | 0.219743 | 0.506453 | 2.94E-153 | lightcyan | MCL1 |
| 1.20E-150 | -0.58412 | 0.453101 | 0.703485 | 1.41E-150 | lightcyan | HMGB2 |
| 2.77E-148 | -2.52097 | 0.042738 | 0.270317 | 3.23E-148 | lightcyan | LYZ |
| 2.78E-147 | -0.64156 | 0.069592 | 0.316981 | 3.22E-147 | lightcyan | NAMPT |
| 2.48E-145 | -0.59856 | 0.549546 | 0.734044 | 2.85E-145 | lightcyan | CALR |
| 1.43E-140 | -0.76738 | 0.990923 | 0.994409 | 1.64E-140 | lightcyan | FTL |
| 3.71E-137 | -0.80761 | 0.169062 | 0.432289 | 4.23E-137 | lightcyan | TIMP1 |
| 2.74E-134 | -0.53965 | 0.146747 | 0.396737 | 3.11E-134 | lightcyan | HERPUD1 |
| 3.78E-134 | -0.50954 | 0.052194 | 0.274431 | 4.26E-134 | lightcyan | PTPRE |
| 1.76E-133 | -0.82048 | 0.058245 | 0.269332 | 1.98E-133 | lightcyan | SPINK2 |
| 2.27E-133 | -0.5434 | 0.059002 | 0.288814 | 2.53E-133 | lightcyan | CMC1 |
| 8.93E-130 | -0.59909 | 0.140696 | 0.379928 | 9.90E-130 | lightcyan | ITM2C |

| | | | | | | |
|---|---|---|---|---|---|---|
| 1.38E-124 | -0.50205 | 0.452345 | 0.694131 | 1.52E-124 | lightcyan | HSPA5 |
| 4.77E-124 | -1.04341 | 0.144856 | 0.365615 | 5.23E-124 | lightcyan | S100A11 |
| 6.42E-124 | -1.17999 | 0.070348 | 0.274959 | 7.01E-124 | lightcyan | FCER1G |
| 5.46E-117 | -0.69899 | 0.481089 | 0.704716 | 5.93E-117 | lightcyan | TSPO |
| 7.82E-111 | -0.87333 | 0.500378 | 0.636495 | 8.44E-111 | lightcyan | TUBB |
| 4.15E-109 | -0.52641 | 0.349849 | 0.555509 | 4.45E-109 | lightcyan | DUT |
| 2.58E-108 | -0.5239 | 0.213691 | 0.448957 | 2.76E-108 | lightcyan | FOSB |
| 2.28E-107 | -0.5174 | 0.103253 | 0.312621 | 2.42E-107 | lightcyan | CLEC11A |
| 1.05E-106 | -0.52289 | 0.332829 | 0.503464 | 1.11E-106 | lightcyan | IMPDH2 |
| 1.29E-105 | -0.56648 | 0.929652 | 0.972501 | 1.35E-105 | lightcyan | SH3BGRL3 |
| 4.78E-103 | -0.74994 | 0.250378 | 0.484721 | 5.00E-103 | lightcyan | CTSS |
| 4.79E-102 | -0.53381 | 0.51475 | 0.731688 | 4.98E-102 | lightcyan | IER2 |
| 3.57E-99 | -1.06161 | 0.15885 | 0.346098 | 3.70E-99 | lightcyan | AREG |
| 5.62E-90 | -0.75698 | 0.579425 | 0.720259 | 5.79E-90 | lightcyan | ZFP36 |
| 3.70E-83 | -0.66381 | 0.498109 | 0.711362 | 3.79E-83 | lightcyan | PSAP |
| 5.74E-75 | -0.52012 | 0.153177 | 0.321166 | 5.85E-75 | lightcyan | ID2 |
| 2.24E-68 | -0.50394 | 0.09947 | 0.253437 | 2.28E-68 | lightcyan | ATP2B1-AS1 |
| 1.85E-67 | -0.6747 | 0.454992 | 0.558357 | 1.87E-67 | lightcyan | AIF1 |
| 3.18E-16 | -1.01162 | 0.842284 | 0.822063 | 3.20E-16 | lightcyan | S100A6 |
| 0 | 2.648862 | 0.884264 | 0.026725 | 0 | lightgreen | MS4A1 |
| 0 | 2.563511 | 0.981726 | 0.101474 | 0 | lightgreen | CD79A |
| 0 | 2.246158 | 0.585279 | 0.010168 | 0 | lightgreen | TCL1A |
| 0 | 1.960052 | 1 | 0.9134 | 0 | lightgreen | CD74 |
| 0 | 1.92268 | 0.756853 | 0.021195 | 0 | lightgreen | LINC00926 |
| 0 | 1.889977 | 0.317766 | 0.011439 | 0 | lightgreen | IGLC2 |
| 0 | 1.87332 | 0.887817 | 0.268146 | 0 | lightgreen | HLA-DQA1 |
| 0 | 1.779783 | 0.840102 | 0.151523 | 0 | lightgreen | CD79B |
| 0 | 1.761657 | 0.927411 | 0.330391 | 0 | lightgreen | HLA-DQB1 |
| 0 | 1.751992 | 0.611168 | 0.012435 | 0 | lightgreen | FCER2 |
| 0 | 1.713054 | 0.784772 | 0.155611 | 0 | lightgreen | IGHM |
| 0 | 1.693614 | 0.754315 | 0.057951 | 0 | lightgreen | FAM129C |
| 0 | 1.685691 | 0.996447 | 0.901309 | 0 | lightgreen | CD37 |
| 0 | 1.675119 | 0.725888 | 0.064993 | 0 | lightgreen | BANK1 |
| 0 | 1.674308 | 0.487817 | 0.040809 | 0 | lightgreen | IGKC |
| 0 | 1.632287 | 0.994924 | 0.575556 | 0 | lightgreen | HLA-DRA |
| 0 | 1.548466 | 0.990355 | 0.617395 | 0 | lightgreen | HLA-DRB1 |
| 0 | 1.491627 | 0.585787 | 0.124077 | 0 | lightgreen | CD83 |
| 0 | 1.482605 | 0.293909 | 0.009309 | 0 | lightgreen | IGLL5 |
| 0 | 1.472333 | 0.268528 | 0.006733 | 0 | lightgreen | IGLC3 |
| 0 | 1.472145 | 0.980203 | 0.628594 | 0 | lightgreen | HLA-DPA1 |
| 0 | 1.454851 | 0.86802 | 0.211913 | 0 | lightgreen | NCF1 |

| | | | | | | |
|---|---|---|---|---|---|---|
| 0 | 1.404514 | 0.980711 | 0.630552 | 0 | lightgreen | HLA-DPB1 |
| 0 | 1.34292 | 0.607614 | 0.087424 | 0 | lightgreen | RALGPS2 |
| 0 | 1.34217 | 0.562944 | 0.018893 | 0 | lightgreen | FCRLA |
| 0 | 1.336126 | 0.585787 | 0.070145 | 0 | lightgreen | HVCN1 |
| 0 | 1.335482 | 0.602538 | 0.025248 | 0 | lightgreen | CD19 |
| 0 | 1.247826 | 0.54264 | 0.192779 | 0 | lightgreen | PLPP5 |
| 0 | 1.173165 | 0.953299 | 0.556731 | 0 | lightgreen | HLA-DRB5 |
| 0 | 1.158809 | 0.478173 | 0.006355 | 0 | lightgreen | FCRL1 |
| 0 | 1.153174 | 0.481726 | 0.041256 | 0 | lightgreen | CD24 |
| 0 | 1.126809 | 0.420812 | 0.096115 | 0 | lightgreen | IL4R |
| 0 | 1.122455 | 0.753807 | 0.26289 | 0 | lightgreen | FCMR |
| 0 | 1.112556 | 0.473096 | 0.035313 | 0 | lightgreen | RUBCNL |
| 0 | 1.109472 | 0.447716 | 0.065645 | 0 | lightgreen | IGHD |
| 0 | 1.096207 | 0.470558 | 0.028237 | 0 | lightgreen | CD22 |
| 0 | 1.085767 | 0.52132 | 0.105184 | 0 | lightgreen | BIRC3 |
| 0 | 1.082844 | 0.615228 | 0.224692 | 0 | lightgreen | CYB561A3 |
| 0 | 1.079049 | 0.611675 | 0.051286 | 0 | lightgreen | VPREB3 |
| 0 | 1.075657 | 0.494924 | 0.108962 | 0 | lightgreen | AFF3 |
| 0 | 1.06688 | 0.471066 | 0.014702 | 0 | lightgreen | BLK |
| 0 | 1.054782 | 0.759391 | 0.416269 | 0 | lightgreen | ZFP36L1 |
| 0 | 1.053256 | 0.932487 | 0.810862 | 0 | lightgreen | TSC22D3 |
| 0 | 1.052744 | 0.634518 | 0.276974 | 0 | lightgreen | MEF2C |
| 0 | 1.051695 | 0.861929 | 0.364982 | 0 | lightgreen | LTB |
| 0 | 1.045354 | 0.459391 | 0.110268 | 0 | lightgreen | ADAM28 |
| 0 | 1.022668 | 0.932487 | 0.78424 | 0 | lightgreen | TXNIP |
| 0 | 1.010166 | 0.464467 | 0.03497 | 0 | lightgreen | POU2AF1 |
| 0 | 1.005208 | 0.58934 | 0.168046 | 0 | lightgreen | HLA-DMB |
| 0 | 0.991687 | 0.308122 | 0.050462 | 0 | lightgreen | JCHAIN |
| 0 | 0.986815 | 0.504061 | 0.149222 | 0 | lightgreen | SWAP70 |
| 0 | 0.98512 | 0.856853 | 0.4747 | 0 | lightgreen | HLA-DMA |
| 0 | 0.951675 | 0.374112 | 0.043386 | 0 | lightgreen | CD72 |
| 0 | 0.951502 | 0.909645 | 0.583869 | 0 | lightgreen | CXCR4 |
| 0 | 0.946454 | 0.430457 | 0.044004 | 0 | lightgreen | CD40 |
| 0 | 0.941074 | 0.945178 | 0.776236 | 0 | lightgreen | CD52 |
| 0 | 0.920009 | 0.421827 | 0.031054 | 0 | lightgreen | 1-Mar |
| 0 | 0.916854 | 0.984772 | 0.922847 | 0 | lightgreen | LAPTM5 |
| 0 | 0.903765 | 0.689848 | 0.401223 | 0 | lightgreen | SNX2 |
| 0 | 0.876692 | 0.374112 | 0.005737 | 0 | lightgreen | HLA-DOB |
| 0 | 0.840406 | 0.435533 | 0.113909 | 0 | lightgreen | SMIM14 |
| 0 | 0.831226 | 0.361421 | 0.022603 | 0 | lightgreen | PAX5 |
| 0 | 0.809584 | 0.430457 | 0.114836 | 0 | lightgreen | GNG7 |

| | | | | | | |
|---|---|---|---|---|---|---|
| 0 | 0.770559 | 0.45533 | 0.147367 | 0 | lightgreen | POU2F2 |
| 0 | 0.757004 | 0.402538 | 0.070867 | 0 | lightgreen | BLNK |
| 0 | 0.756757 | 0.426396 | 0.127546 | 0 | lightgreen | SNX29 |
| 0 | 0.748798 | 0.664467 | 0.307925 | 0 | lightgreen | ISG20 |
| 0 | 0.737837 | 0.346701 | 0.045412 | 0 | lightgreen | NAPSA |
| 0 | 0.731847 | 0.371574 | 0.094947 | 0 | lightgreen | MTSS1 |
| 0 | 0.725254 | 0.939086 | 0.832194 | 0 | lightgreen | LIMD2 |
| 0 | 0.70042 | 0.346193 | 0.054035 | 0 | lightgreen | ALOX5 |
| 0 | 0.698647 | 0.484264 | 0.114046 | 0 | lightgreen | IRF8 |
| 0 | 0.693991 | 0.279188 | 0.008519 | 0 | lightgreen | CXCR5 |
| 0 | 0.693518 | 0.336041 | 0.078767 | 0 | lightgreen | HLA-DOA |
| 0 | 0.651832 | 0.279188 | 0.015767 | 0 | lightgreen | FCGR2B |
| 0 | 0.650555 | 0.266497 | 0.025592 | 0 | lightgreen | SNX22 |
| 0 | 0.643575 | 0.275127 | 0.03655 | 0 | lightgreen | IGHV5-78 |
| 0 | 0.631529 | 0.418274 | 0.119371 | 0 | lightgreen | LY86 |
| 0 | 0.625917 | 0.288832 | 0.029508 | 0 | lightgreen | FGD2 |
| 0 | 0.615853 | 0.264467 | 0.011233 | 0 | lightgreen | ARHGAP24 |
| 0 | 0.589713 | 0.326904 | 0.0675 | 0 | lightgreen | SP140 |
| 0 | -0.88299 | 0.840609 | 0.981897 | 0 | lightgreen | ACTG1 |
| 0 | -0.96515 | 0.092893 | 0.58593 | 0 | lightgreen | LEPROTL1 |
| 0 | -1.07564 | 0.147208 | 0.617361 | 0 | lightgreen | TXN |
| 0 | -1.09088 | 0.035533 | 0.504655 | 0 | lightgreen | MT2A |
| 0 | -1.13679 | 0.690863 | 0.92192 | 0 | lightgreen | VIM |
| 0 | -1.16381 | 0.452284 | 0.831541 | 0 | lightgreen | LDHB |
| 0 | -1.17378 | 0.059898 | 0.591289 | 0 | lightgreen | NUCB2 |
| 0 | -1.19295 | 0.372081 | 0.762083 | 0 | lightgreen | GSTP1 |
| 0 | -1.31331 | 0.012183 | 0.52925 | 0 | lightgreen | FYB1 |
| 0 | -1.44715 | 0.086802 | 0.696163 | 0 | lightgreen | HCST |
| 0 | -1.54912 | 0.009645 | 0.601353 | 0 | lightgreen | CTSW |
| 0 | -1.59508 | 0.031472 | 0.584624 | 0 | lightgreen | AIF1 |
| 0 | -1.59936 | 0.783756 | 0.983202 | 0 | lightgreen | GAPDH |
| 0 | -1.74161 | 0.423858 | 0.850847 | 0 | lightgreen | S100A6 |
| 0 | -1.79839 | 0.024873 | 0.732197 | 0 | lightgreen | ANXA1 |
| 0 | -2.07913 | 0.148223 | 0.873519 | 0 | lightgreen | SRGN |
| 0 | -2.13934 | 0.234518 | 0.850778 | 0 | lightgreen | S100A4 |
| 2.91E-307 | -0.88177 | 0.427411 | 0.83611 | 7.80E-307 | lightgreen | CD99 |
| 1.60E-302 | -0.83832 | 0.339594 | 0.739892 | 4.25E-302 | lightgreen | LDHA |
| 1.30E-299 | 0.815377 | 0.736548 | 0.527258 | 3.42E-299 | lightgreen | SYNGR2 |
| 9.60E-298 | 0.858946 | 0.651269 | 0.383395 | 2.49E-297 | lightgreen | POLD4 |
| 3.87E-287 | 0.571629 | 0.356345 | 0.094706 | 9.96E-287 | lightgreen | IKZF3 |
| 1.40E-285 | -1.48062 | 0.005584 | 0.437257 | 3.58E-285 | lightgreen | CD7 |

| | | | | | | |
|---|---|---|---|---|---|---|
| 2.33E-285 | -0.7207 | 0.016244 | 0.457078 | 5.87E-285 | lightgreen | SPN |
| 1.03E-277 | -0.67973 | 0.307614 | 0.736182 | 2.57E-277 | lightgreen | NDUFS5 |
| 1.99E-274 | 0.704712 | 0.372589 | 0.122153 | 4.92E-274 | lightgreen | CIITA |
| 2.51E-272 | -0.73411 | 0.108629 | 0.563876 | 6.15E-272 | lightgreen | RGS10 |
| 7.53E-267 | -1.86404 | 0.018274 | 0.427261 | 1.83E-266 | lightgreen | CD3E |
| 3.24E-266 | -1.13848 | 0.018274 | 0.436399 | 7.80E-266 | lightgreen | TIMP1 |
| 3.21E-265 | 0.857974 | 0.574112 | 0.284154 | 7.65E-265 | lightgreen | TAGAP |
| 1.38E-264 | -1.78301 | 0.018782 | 0.434303 | 3.27E-264 | lightgreen | CST3 |
| 1.29E-262 | 0.556182 | 0.394416 | 0.120229 | 3.03E-262 | lightgreen | CTSH |
| 1.65E-260 | -0.87652 | 0.279188 | 0.641373 | 3.83E-260 | lightgreen | TALDO1 |
| 5.23E-260 | 0.769422 | 0.647208 | 0.404005 | 1.20E-259 | lightgreen | SP100 |
| 3.08E-258 | 0.544145 | 0.289848 | 0.074611 | 7.03E-258 | lightgreen | TMEM154 |
| 1.55E-255 | 0.758668 | 0.750761 | 0.531105 | 3.52E-255 | lightgreen | EZR |
| 1.08E-254 | -0.50584 | 0.879188 | 0.956236 | 2.44E-254 | lightgreen | ATP5MC2 |
| 1.04E-249 | -0.77526 | 0.426904 | 0.7469 | 2.32E-249 | lightgreen | ENO1 |
| 3.41E-244 | -2.11008 | 0.028934 | 0.41249 | 7.53E-244 | lightgreen | IL32 |
| 5.47E-243 | -1.03015 | 0.269036 | 0.632888 | 1.20E-242 | lightgreen | SERPINB1 |
| 1.46E-241 | -0.64523 | 0.007107 | 0.394696 | 3.18E-241 | lightgreen | BEX3 |
| 2.26E-240 | -0.68198 | 0.032995 | 0.431418 | 4.87E-240 | lightgreen | LCP2 |
| 1.34E-235 | 0.762316 | 0.414213 | 0.171344 | 2.86E-235 | lightgreen | STX7 |
| 1.01E-234 | 0.752323 | 0.816244 | 0.692247 | 2.14E-234 | lightgreen | UCP2 |
| 1.10E-232 | -1.55681 | 0.245685 | 0.595067 | 2.31E-232 | lightgreen | IFITM1 |
| 7.73E-227 | -0.82288 | 0.311675 | 0.661503 | 1.61E-226 | lightgreen | TKT |
| 5.64E-226 | 0.683825 | 0.45533 | 0.208272 | 1.17E-225 | lightgreen | PDLIM1 |
| 6.85E-223 | -0.66839 | 0.131472 | 0.550926 | 1.41E-222 | lightgreen | SLC9A3R1 |
| 8.83E-218 | -0.52976 | 0.005584 | 0.364879 | 1.80E-217 | lightgreen | TMEM173 |
| 5.24E-217 | -1.21265 | 0.003553 | 0.358421 | 1.06E-216 | lightgreen | PRSS57 |
| 1.72E-214 | -0.5623 | 0.16599 | 0.588609 | 3.45E-214 | lightgreen | ABRACL |
| 1.77E-212 | -0.62301 | 0.688325 | 0.874103 | 3.52E-212 | lightgreen | HSP90AA1 |
| 9.04E-212 | -0.64484 | 0.101015 | 0.495311 | 1.79E-211 | lightgreen | CD63 |
| 1.62E-211 | -0.78333 | 0.114213 | 0.506956 | 3.19E-211 | lightgreen | MCL1 |
| 3.06E-210 | -0.65619 | 0.125381 | 0.524441 | 5.96E-210 | lightgreen | ARRB2 |
| 1.15E-209 | -0.58414 | 0.101015 | 0.494212 | 2.23E-209 | lightgreen | COMT |
| 2.69E-209 | -0.71045 | 0.175635 | 0.581636 | 5.17E-209 | lightgreen | DNAJB1 |
| 6.96E-205 | -0.61562 | 0.002538 | 0.343513 | 1.33E-204 | lightgreen | GNA15 |
| 1.04E-203 | -0.98966 | 0.287817 | 0.636907 | 1.96E-203 | lightgreen | H2AFZ |
| 2.33E-202 | -1.15306 | 0.004061 | 0.340765 | 4.38E-202 | lightgreen | CD3D |
| 8.89E-201 | -0.6076 | 0.03198 | 0.38573 | 1.66E-200 | lightgreen | RNF130 |
| 9.15E-198 | -0.59865 | 0.060406 | 0.427021 | 1.70E-197 | lightgreen | ITM2A |
| 6.54E-197 | -0.5861 | 0.082741 | 0.45749 | 1.20E-196 | lightgreen | CTSC |
| 4.38E-196 | -0.68857 | 0.779695 | 0.90021 | 8.01E-196 | lightgreen | RPS2 |

| | | | | | | |
|---|---|---|---|---|---|---|
| 2.23E-194 | -0.70681 | 0.173096 | 0.556113 | 4.04E-194 | lightgreen | RARRES3 |
| 3.58E-194 | -0.66721 | 0.000508 | 0.326578 | 6.45E-194 | lightgreen | EGFL7 |
| 3.61E-194 | -0.96765 | 0.244162 | 0.631583 | 6.46E-194 | lightgreen | NEAT1 |
| 5.52E-192 | -0.6588 | 0.092386 | 0.463227 | 9.82E-192 | lightgreen | PIK3R1 |
| 1.28E-188 | -1.9619 | 0.020812 | 0.345917 | 2.27E-188 | lightgreen | TYROBP |
| 2.14E-185 | 0.807107 | 0.588325 | 0.361066 | 3.76E-185 | lightgreen | HERPUD1 |
| 2.78E-185 | -0.8384 | 0.313198 | 0.7109 | 4.86E-185 | lightgreen | TSPO |
| 9.65E-184 | 0.782381 | 0.545178 | 0.319089 | 1.67E-183 | lightgreen | RCSD1 |
| 5.19E-183 | -0.6533 | 0.160406 | 0.521109 | 8.93E-183 | lightgreen | H2AFY |
| 2.76E-182 | -0.57637 | 0.336041 | 0.656419 | 4.72E-182 | lightgreen | COX5A |
| 8.37E-182 | -0.84816 | 0.01066 | 0.326921 | 1.42E-181 | lightgreen | ID2 |
| 8.24E-181 | -0.59811 | 0.490863 | 0.756827 | 1.39E-180 | lightgreen | TPI1 |
| 9.40E-178 | -0.59529 | 0.197462 | 0.569304 | 1.58E-177 | lightgreen | P4HB |
| 1.92E-177 | 0.653587 | 0.618782 | 0.402563 | 3.19E-177 | lightgreen | RIPOR2 |
| 1.88E-176 | -0.66352 | 0.393401 | 0.7008 | 3.11E-176 | lightgreen | GYPC |
| 1.11E-175 | -0.59726 | 0.000508 | 0.302566 | 1.84E-175 | lightgreen | MGST1 |
| 8.62E-172 | -0.6412 | 0.001015 | 0.2981 | 1.41E-171 | lightgreen | CD34 |
| 1.21E-171 | -0.61193 | 0.032487 | 0.348391 | 1.96E-171 | lightgreen | SELPLG |
| 2.19E-171 | -1.56498 | 0.014721 | 0.315826 | 3.54E-171 | lightgreen | CST7 |
| 1.86E-170 | -0.72296 | 0.063452 | 0.387963 | 2.99E-170 | lightgreen | AC084033.3 |
| 1.34E-169 | -0.64074 | 0.118782 | 0.463948 | 2.14E-169 | lightgreen | VSIR |
| 9.59E-169 | -0.6409 | 0.088832 | 0.421868 | 1.52E-168 | lightgreen | CDK6 |
| 1.49E-168 | -2.30001 | 0.02335 | 0.324482 | 2.35E-168 | lightgreen | NKG7 |
| 2.29E-168 | -0.5152 | 0.116244 | 0.472021 | 3.59E-168 | lightgreen | PGD |
| 3.47E-168 | -0.70687 | 0.013198 | 0.313868 | 5.41E-168 | lightgreen | CLEC11A |
| 4.46E-168 | -0.79951 | 0.029442 | 0.33307 | 6.91E-168 | lightgreen | SAMHD1 |
| 7.76E-167 | -0.57397 | 0.144162 | 0.508158 | 1.20E-166 | lightgreen | GLUL |
| 1.86E-166 | -0.52088 | 0.01269 | 0.310913 | 2.85E-166 | lightgreen | CYTOR |
| 5.32E-166 | 0.517772 | 0.320812 | 0.119955 | 8.10E-166 | lightgreen | AC119396.1 |
| 7.33E-163 | -0.68513 | 0.005584 | 0.293703 | 1.11E-162 | lightgreen | CSF3R |
| 5.97E-161 | -0.72665 | 0.513706 | 0.791453 | 8.99E-161 | lightgreen | IFITM2 |
| 4.77E-160 | -0.52381 | 0.357868 | 0.689739 | 7.14E-160 | lightgreen | VAMP8 |
| 2.23E-159 | 0.674038 | 0.505076 | 0.292776 | 3.32E-159 | lightgreen | PDE4B |
| 1.15E-157 | -0.52561 | 0.419289 | 0.748136 | 1.70E-157 | lightgreen | COX8A |
| 5.58E-157 | -0.65621 | 0.006091 | 0.2857 | 8.22E-157 | lightgreen | PRKCH |
| 8.64E-157 | -1.17083 | 0.435533 | 0.677613 | 1.26E-156 | lightgreen | TUBA1B |
| 8.91E-157 | -0.70631 | 0.005584 | 0.284429 | 1.30E-156 | lightgreen | LAT |
| 2.22E-156 | -0.50782 | 0.200508 | 0.544708 | 3.21E-156 | lightgreen | HSPB1 |
| 3.64E-155 | -0.61643 | 0.081726 | 0.390746 | 5.24E-155 | lightgreen | FABP5 |
| 5.23E-154 | -1.13329 | 0.335533 | 0.626396 | 7.49E-154 | lightgreen | S100A10 |
| 1.96E-153 | -0.75611 | 0.015228 | 0.293635 | 2.80E-153 | lightgreen | ZAP70 |

| | | | | | | |
|---|---|---|---|---|---|---|
| 2.43E-151 | 0.637885 | 0.531472 | 0.343822 | 3.45E-151 | lightgreen | SP110 |
| 3.73E-151 | -1.35722 | 0.030964 | 0.307616 | 5.26E-151 | lightgreen | IL7R |
| 4.66E-151 | -1.18113 | 0.073096 | 0.36536 | 6.54E-151 | lightgreen | S100A11 |
| 4.54E-150 | -0.8989 | 0.00203 | 0.269486 | 6.34E-150 | lightgreen | GZMM |
| 2.28E-148 | -0.50997 | 0.490863 | 0.732747 | 3.16E-148 | lightgreen | ANP32B |
| 2.53E-148 | -0.77521 | 0.346701 | 0.664216 | 3.49E-148 | lightgreen | ITGB2 |
| 3.12E-147 | 0.73489 | 0.471574 | 0.245577 | 4.28E-147 | lightgreen | KLF2 |
| 1.07E-146 | -0.80439 | 0.004569 | 0.268524 | 1.47E-146 | lightgreen | CD3G |
| 1.18E-145 | -1.31032 | 0.009645 | 0.274329 | 1.60E-145 | lightgreen | FCER1G |
| 3.60E-145 | 0.509147 | 0.304061 | 0.122084 | 4.87E-145 | lightgreen | NCOA3 |
| 2.78E-144 | -0.55378 | 0.170558 | 0.504655 | 3.74E-144 | lightgreen | PYCARD |
| 2.81E-144 | -0.50142 | 0.633503 | 0.863694 | 3.76E-144 | lightgreen | PPIB |
| 1.84E-141 | -1.32539 | 0.116244 | 0.397444 | 2.45E-141 | lightgreen | LGALS1 |
| 3.37E-140 | 0.561232 | 0.395431 | 0.187283 | 4.46E-140 | lightgreen | ARID5B |
| 5.87E-140 | 0.652916 | 0.607107 | 0.459414 | 7.74E-140 | lightgreen | REL |
| 1.37E-139 | -0.56743 | 0.011675 | 0.272818 | 1.80E-139 | lightgreen | H1F0 |
| 1.55E-139 | -0.52838 | 0.928426 | 0.918038 | 2.02E-139 | lightgreen | NPM1 |
| 7.69E-139 | 0.575009 | 0.298985 | 0.12497 | 9.99E-139 | lightgreen | QRSL1 |
| 5.81E-137 | -0.91988 | 0.21269 | 0.495689 | 7.51E-137 | lightgreen | STMN1 |
| 8.43E-136 | -0.87312 | 0.152792 | 0.439868 | 1.08E-135 | lightgreen | TNFAIP3 |
| 9.59E-136 | -0.51899 | 0.054315 | 0.334341 | 1.23E-135 | lightgreen | RNF125 |
| 3.90E-135 | -0.54864 | 0.119797 | 0.424547 | 4.96E-135 | lightgreen | TUBB4B |
| 1.20E-134 | 0.77326 | 0.419797 | 0.25444 | 1.52E-134 | lightgreen | EAF2 |
| 1.16E-133 | -0.85015 | 0.008629 | 0.255436 | 1.46E-133 | lightgreen | CD247 |
| 1.21E-132 | -0.81494 | 0.075635 | 0.33623 | 1.52E-132 | lightgreen | LCK |
| 1.50E-132 | -0.50616 | 0.056853 | 0.333104 | 1.88E-132 | lightgreen | AGTRAP |
| 1.74E-132 | -0.63831 | 0.241117 | 0.558002 | 2.16E-132 | lightgreen | FLNA |
| 2.35E-132 | 0.616186 | 0.593401 | 0.459345 | 2.92E-132 | lightgreen | PTPN6 |
| 3.36E-131 | -0.56558 | 0.201015 | 0.529834 | 4.14E-131 | lightgreen | NOSIP |
| 1.21E-130 | -0.88444 | 0.020305 | 0.267012 | 1.48E-130 | lightgreen | SPINK2 |
| 3.36E-130 | -0.52239 | 0.052284 | 0.323039 | 4.10E-130 | lightgreen | LYAR |
| 9.96E-129 | -0.65967 | 0.024873 | 0.276219 | 1.21E-128 | lightgreen | SMIM24 |
| 1.53E-128 | -0.76583 | 0.108629 | 0.398853 | 1.85E-128 | lightgreen | LST1 |
| 2.65E-128 | -0.55937 | 0.038071 | 0.293806 | 3.19E-128 | lightgreen | MGAT4A |
| 1.70E-125 | 0.622942 | 0.45736 | 0.283673 | 2.04E-125 | lightgreen | ZCCHC7 |
| 2.63E-124 | -0.57462 | 0.277665 | 0.547216 | 3.14E-124 | lightgreen | HSPD1 |
| 4.15E-124 | 0.629303 | 0.500508 | 0.318917 | 4.93E-124 | lightgreen | TMEM243 |
| 7.26E-122 | 0.848848 | 0.635025 | 0.47233 | 8.59E-122 | lightgreen | CD69 |
| 5.27E-118 | 0.510375 | 0.685787 | 0.517262 | 6.20E-118 | lightgreen | BTG1 |
| 2.05E-114 | 0.585412 | 0.43198 | 0.26478 | 2.41E-114 | lightgreen | EBLN3P |
| 2.23E-111 | -0.55979 | 0.045178 | 0.284429 | 2.60E-111 | lightgreen | CMC1 |

| | | | | | | |
|---|---|---|---|---|---|---|
| 1.37E-109 | -0.90646 | 0.418274 | 0.638899 | 1.60E-109 | lightgreen | TUBB |
| 5.50E-109 | 0.523128 | 0.315736 | 0.149875 | 6.36E-109 | lightgreen | TMEM131L |
| 2.19E-108 | -0.56802 | 0.056345 | 0.293841 | 2.52E-108 | lightgreen | BLVRB |
| 6.22E-102 | -0.58496 | 0.073096 | 0.311016 | 7.14E-102 | lightgreen | NAMPT |
| 1.13E-101 | 0.57187 | 0.37868 | 0.235993 | 1.29E-101 | lightgreen | BTK |
| 2.53E-101 | -0.5621 | 0.281218 | 0.503006 | 2.88E-101 | lightgreen | IMPDH2 |
| 2.76E-101 | 0.566265 | 0.34467 | 0.19326 | 3.13E-101 | lightgreen | ARHGAP17 |
| 1.70E-100 | 0.531301 | 0.353299 | 0.194119 | 1.91E-100 | lightgreen | PPM1K |
| 3.04E-99 | -0.5415 | 0.298985 | 0.554189 | 3.41E-99 | lightgreen | DUT |
| 3.10E-99 | -2.39091 | 0.051777 | 0.264436 | 3.46E-99 | lightgreen | LYZ |
| 8.09E-99 | 0.702559 | 0.434518 | 0.297894 | 9.01E-99 | lightgreen | SYK |
| 9.01E-98 | -0.58413 | 0.093401 | 0.319226 | 9.99E-98 | lightgreen | ARL4C |
| 1.45E-95 | 0.52504 | 0.27665 | 0.132733 | 1.61E-95 | lightgreen | PLEKHA2 |
| 2.68E-95 | -0.85568 | 0.052284 | 0.269348 | 2.95E-95 | lightgreen | IGLL1 |
| 3.18E-94 | 0.544184 | 0.601523 | 0.513723 | 3.48E-94 | lightgreen | FOXP1 |
| 4.54E-92 | 0.513912 | 0.373096 | 0.217684 | 4.95E-92 | lightgreen | PIM2 |
| 5.89E-92 | 0.525106 | 0.540102 | 0.40428 | 6.40E-92 | lightgreen | RHOH |
| 5.74E-89 | 0.748669 | 0.296954 | 0.164302 | 6.21E-89 | lightgreen | BCL7A |
| 1.19E-88 | -0.53749 | 0.885279 | 0.922435 | 1.28E-88 | lightgreen | HMGB1 |
| 1.09E-87 | 0.504351 | 0.385787 | 0.241764 | 1.17E-87 | lightgreen | LMBRD1 |
| 2.01E-84 | -0.52881 | 0.247208 | 0.499914 | 2.15E-84 | lightgreen | IFITM3 |
| 1.53E-83 | 0.518329 | 0.325381 | 0.193398 | 1.62E-83 | lightgreen | PKIG |
| 4.48E-83 | -0.55692 | 0.293909 | 0.535227 | 4.75E-83 | lightgreen | LITAF |
| 4.23E-82 | -0.52489 | 0.193401 | 0.426815 | 4.46E-82 | lightgreen | DDIT4 |
| 4.34E-81 | -0.51376 | 0.073604 | 0.273608 | 4.56E-81 | lightgreen | HOPX |
| 8.46E-79 | 0.572574 | 0.280203 | 0.150321 | 8.86E-79 | lightgreen | SYVN1 |
| 1.33E-77 | -0.51469 | 0.079188 | 0.268146 | 1.39E-77 | lightgreen | FLT3LG |
| 1.57E-75 | 0.560628 | 0.356345 | 0.236921 | 1.63E-75 | lightgreen | IFT57 |
| 1.38E-71 | 0.539352 | 0.306091 | 0.17715 | 1.43E-71 | lightgreen | PELI1 |
| 4.83E-63 | 0.500177 | 0.311675 | 0.201573 | 4.98E-63 | lightgreen | ORAI2 |
| 7.00E-59 | -0.70249 | 0.99797 | 0.996393 | 7.19E-59 | lightgreen | FTH1 |
| 4.49E-57 | 0.701971 | 0.292893 | 0.18402 | 4.59E-57 | lightgreen | LHPP |
| 7.34E-55 | 0.576836 | 0.370558 | 0.288276 | 7.48E-55 | lightgreen | ADK |
| 2.32E-53 | -0.86049 | 0.17665 | 0.340559 | 2.36E-53 | lightgreen | AREG |
| 4.16E-45 | 0.618743 | 0.301523 | 0.194669 | 4.21E-45 | lightgreen | MZB1 |
| 6.77E-28 | -0.54837 | 0.441117 | 0.60575 | 6.82E-28 | lightgreen | HIST1H4C |
| 3.20E-05 | -0.60969 | 0.523858 | 0.629796 | 3.22E-05 | lightgreen | SAT1 |
| 0.00026 | 0.949922 | 0.280203 | 0.292295 | 0.00026 | lightgreen | HIST1H1C |
| 0 | 2.625101 | 0.990798 | 0.164461 | 0 | greenyellow | PLD4 |
| 0 | 2.418006 | 0.981595 | 0.197269 | 0 | greenyellow | IRF7 |
| 0 | 2.408831 | 0.957055 | 0.057357 | 0 | greenyellow | JCHAIN |

| | | | | | | |
|---|---|---|---|---|---|---|
| 0 | 2.403595 | 0.929448 | 0.012746 | 0 | greenyellow | LILRA4 |
| 0 | 2.29759 | 0.981595 | 0.128564 | 0 | greenyellow | IRF8 |
| 0 | 2.206829 | 0.699387 | 0.034466 | 0 | greenyellow | RASD1 |
| 0 | 2.199401 | 0.978528 | 0.246627 | 0 | greenyellow | APP |
| 0 | 1.890482 | 0.917178 | 0.064315 | 0 | greenyellow | IL3RA |
| 0 | 1.79477 | 0.619632 | 0.021167 | 0 | greenyellow | SERPINF1 |
| 0 | 1.761572 | 0.730061 | 0.040676 | 0 | greenyellow | IRF4 |
| 0 | 1.562164 | 0.871166 | 0.058169 | 0 | greenyellow | LILRB4 |
| 0 | 1.537107 | 0.644172 | 0.010145 | 0 | greenyellow | SMPD3 |
| 0 | 1.534822 | 0.865031 | 0.111429 | 0 | greenyellow | RNASE6 |
| 0 | 1.507087 | 0.809816 | 0.064445 | 0 | greenyellow | UGCG |
| 0 | 1.486089 | 0.868098 | 0.048545 | 0 | greenyellow | LGMN |
| 0 | 1.462333 | 0.855828 | 0.063827 | 0 | greenyellow | CCDC50 |
| 0 | 1.455133 | 0.812883 | 0.061258 | 0 | greenyellow | IGKC |
| 0 | 1.441855 | 0.779141 | 0.120371 | 0 | greenyellow | LDLRAD4 |
| 0 | 1.363692 | 0.539877 | 0.035376 | 0 | greenyellow | TPM2 |
| 0 | 1.350765 | 0.754601 | 0.10278 | 0 | greenyellow | DERL3 |
| 0 | 1.287074 | 0.680982 | 0.006015 | 0 | greenyellow | CLEC4C |
| 0 | 1.279965 | 0.828221 | 0.067176 | 0 | greenyellow | TGFBI |
| 0 | 1.23708 | 0.815951 | 0.094521 | 0 | greenyellow | FAM129C |
| 0 | 1.224695 | 0.662577 | 0.070655 | 0 | greenyellow | PPP1R14B |
| 0 | 1.210738 | 0.542945 | 0.005918 | 0 | greenyellow | SCT |
| 0 | 1.12943 | 0.708589 | 0.091595 | 0 | greenyellow | TNFRSF21 |
| 0 | 0.995216 | 0.567485 | 0.047927 | 0 | greenyellow | FCRLA |
| 0 | 0.982964 | 0.57362 | 0.043212 | 0 | greenyellow | SLC15A4 |
| 0 | 0.890025 | 0.592025 | 0.058917 | 0 | greenyellow | NAPSA |
| 0 | 0.879265 | 0.56135 | 0.035994 | 0 | greenyellow | TNNI2 |
| 0 | 0.856316 | 0.533742 | 0.028158 | 0 | greenyellow | SPIB |
| 0 | 0.826106 | 0.328221 | 0.002634 | 0 | greenyellow | PTCRA |
| 0 | 0.77901 | 0.398773 | 0.029524 | 0 | greenyellow | ZFAT |
| 0 | 0.752329 | 0.469325 | 0.018143 | 0 | greenyellow | SHD |
| 0 | 0.677762 | 0.41411 | 0.004422 | 0 | greenyellow | PACSIN1 |
| 0 | 0.661178 | 0.404908 | 0.025622 | 0 | greenyellow | ADAM19 |
| 0 | 0.64357 | 0.392638 | 0.004357 | 0 | greenyellow | PROC |
| 0 | 0.626444 | 0.383436 | 0.005528 | 0 | greenyellow | PTPRS |
| 0 | 0.606491 | 0.352761 | 0.006405 | 0 | greenyellow | AC119428.2 |
| 0 | 0.578553 | 0.294479 | 0.01125 | 0 | greenyellow | PLVAP |
| 0 | 0.578147 | 0.355828 | 0.008161 | 0 | greenyellow | SCAMP5 |
| 0 | 0.545839 | 0.315951 | 0.002764 | 0 | greenyellow | AC023590.1 |
| 0 | 0.51577 | 0.266871 | 0.003869 | 0 | greenyellow | FAM160A1 |
| 5.92E-307 | 1.300827 | 0.907975 | 0.156983 | 4.74E-306 | greenyellow | CD68 |

| | | | | | | |
|---|---|---|---|---|---|---|
| 8.28E-301 | 1.580942 | 0.929448 | 0.200065 | 6.48E-300 | greenyellow | C12orf75 |
| 5.19E-292 | 1.681052 | 0.966258 | 0.25456 | 3.97E-291 | greenyellow | TCF4 |
| 1.88E-281 | 1.591546 | 0.895706 | 0.194082 | 1.40E-280 | greenyellow | MZB1 |
| 2.57E-279 | 1.236196 | 0.858896 | 0.177239 | 1.89E-278 | greenyellow | GSN |
| 3.39E-269 | 2.117101 | 0.990798 | 0.352886 | 2.43E-268 | greenyellow | ITM2C |
| 5.99E-253 | 0.844147 | 0.631902 | 0.086165 | 4.22E-252 | greenyellow | BLNK |
| 1.20E-246 | 0.996191 | 0.714724 | 0.116274 | 8.27E-246 | greenyellow | FCER1A |
| 4.01E-242 | 0.913872 | 0.696319 | 0.122061 | 2.71E-241 | greenyellow | ST6GALNAC4 |
| 7.95E-241 | 0.890412 | 0.733129 | 0.116534 | 5.28E-240 | greenyellow | CYBB |
| 1.18E-231 | 1.485594 | 0.92638 | 0.253357 | 7.69E-231 | greenyellow | ALOX5AP |
| 2.51E-228 | 0.920477 | 0.567485 | 0.08558 | 1.61E-227 | greenyellow | MAP1A |
| 8.36E-228 | 0.551968 | 0.263804 | 0.017233 | 5.25E-227 | greenyellow | CRYM |
| 2.10E-227 | 0.615399 | 0.43865 | 0.046952 | 1.30E-226 | greenyellow | SEL1L3 |
| 4.98E-227 | 0.827119 | 0.592025 | 0.090749 | 3.02E-226 | greenyellow | RUNX2 |
| 1.79E-226 | 0.605303 | 0.273006 | 0.018566 | 1.07E-225 | greenyellow | EGLN3 |
| 5.26E-225 | 0.584106 | 0.5 | 0.05843 | 3.09E-224 | greenyellow | RUBCNL |
| 5.69E-217 | 1.005085 | 0.733129 | 0.14596 | 3.28E-216 | greenyellow | IGFLR1 |
| 4.10E-212 | 0.997194 | 0.708589 | 0.143847 | 2.33E-211 | greenyellow | RPS6KA4 |
| 1.12E-209 | 0.644011 | 0.404908 | 0.045651 | 6.28E-209 | greenyellow | SCN9A |
| 2.98E-199 | 0.557195 | 0.41411 | 0.046236 | 1.64E-198 | greenyellow | SULF2 |
| 1.65E-192 | 1.506013 | 0.993865 | 0.694976 | 8.94E-192 | greenyellow | SEC61B |
| 2.09E-191 | 1.325805 | 0.932515 | 0.35178 | 1.11E-190 | greenyellow | CTSB |
| 5.55E-191 | 0.891981 | 0.662577 | 0.131979 | 2.92E-190 | greenyellow | MAPKAPK2 |
| 1.89E-190 | 1.510959 | 0.97546 | 0.540823 | 9.80E-190 | greenyellow | PLP2 |
| 4.92E-190 | 0.615285 | 0.411043 | 0.051016 | 2.51E-189 | greenyellow | DAB2 |
| 1.10E-188 | 0.517574 | 0.365031 | 0.040286 | 5.52E-188 | greenyellow | NUDT17 |
| 4.23E-188 | 0.852599 | 0.699387 | 0.138969 | 2.10E-187 | greenyellow | JAML |
| 2.62E-187 | 0.53039 | 0.349693 | 0.037587 | 1.28E-186 | greenyellow | KCNK17 |
| 1.52E-176 | 0.557697 | 0.43865 | 0.057877 | 7.32E-176 | greenyellow | CARD11 |
| 5.24E-176 | 1.179356 | 0.815951 | 0.24344 | 2.49E-175 | greenyellow | CYB561A3 |
| 1.26E-173 | 1.507721 | 0.960123 | 0.4927 | 5.94E-173 | greenyellow | PLAC8 |
| 4.50E-173 | 0.916289 | 0.59816 | 0.120631 | 2.08E-172 | greenyellow | FCHSD2 |
| 7.07E-171 | 1.209545 | 0.684049 | 0.149212 | 3.23E-170 | greenyellow | GPR183 |
| 2.26E-170 | 1.348813 | 0.98773 | 0.847635 | 1.02E-169 | greenyellow | PPIB |
| 6.84E-169 | 1.363779 | 0.92638 | 0.369631 | 3.05E-168 | greenyellow | HERPUD1 |
| 8.72E-164 | 0.776176 | 0.453988 | 0.064998 | 3.84E-163 | greenyellow | CXCR3 |
| 6.00E-162 | 0.51207 | 0.352761 | 0.042335 | 2.61E-161 | greenyellow | UNC93B1 |
| 1.52E-161 | 1.485497 | 1 | 0.91803 | 6.53E-161 | greenyellow | CD74 |
| 1.20E-159 | 0.783284 | 0.521472 | 0.095822 | 5.07E-159 | greenyellow | RHEX |
| 4.30E-157 | 0.73728 | 0.481595 | 0.075955 | 1.80E-156 | greenyellow | MPEG1 |
| 9.20E-157 | 1.616974 | 0.923313 | 0.387059 | 3.81E-156 | greenyellow | SOX4 |

| | | | | | | |
|---|---|---|---|---|---|---|
| 1.03E-155 | 0.893644 | 0.880368 | 0.246854 | 4.20E-155 | greenyellow | NCF1 |
| 1.06E-153 | 1.180963 | 0.972393 | 0.682166 | 4.28E-153 | greenyellow | SPCS1 |
| 1.01E-152 | 1.140505 | 0.993865 | 0.83141 | 4.04E-152 | greenyellow | TAGLN2 |
| 8.88E-152 | 0.94359 | 0.791411 | 0.253292 | 3.51E-151 | greenyellow | BCL11A |
| 7.31E-150 | 0.815114 | 0.907975 | 0.250658 | 2.86E-149 | greenyellow | FCER1G |
| 3.84E-148 | 0.817414 | 0.674847 | 0.167355 | 1.49E-147 | greenyellow | PHACTR1 |
| 1.21E-147 | 0.594754 | 0.481595 | 0.084799 | 4.61E-147 | greenyellow | ZDHHC17 |
| 3.19E-147 | 1.233866 | 1 | 0.597919 | 1.21E-146 | greenyellow | HLA-DRA |
| 4.89E-147 | 0.605227 | 0.404908 | 0.060933 | 1.83E-146 | greenyellow | WDFY4 |
| 3.43E-145 | 1.393754 | 0.969325 | 0.542741 | 1.27E-144 | greenyellow | NPC2 |
| 4.79E-145 | 1.092977 | 0.978528 | 0.493838 | 1.76E-144 | greenyellow | HLA-DMA |
| 9.08E-145 | 0.815023 | 0.625767 | 0.155552 | 3.29E-144 | greenyellow | CD2AP |
| 8.89E-144 | 0.621745 | 0.613497 | 0.129215 | 3.19E-143 | greenyellow | SMIM14 |
| 1.17E-142 | 0.720372 | 0.407975 | 0.063632 | 4.16E-142 | greenyellow | GRAMD1B |
| 1.62E-139 | 0.973613 | 0.969325 | 0.402016 | 5.70E-139 | greenyellow | CST3 |
| 6.12E-139 | 0.623561 | 0.481595 | 0.095139 | 2.13E-138 | greenyellow | CCDC189 |
| 1.22E-138 | 1.015481 | 0.843558 | 0.311169 | 4.21E-138 | greenyellow | CAPG |
| 1.40E-137 | 1.098738 | 0.996933 | 0.577467 | 4.79E-137 | greenyellow | HLA-DRB5 |
| 1.13E-136 | 0.992949 | 0.90184 | 0.363193 | 3.82E-136 | greenyellow | GRN |
| 1.81E-134 | 1.010746 | 0.990798 | 0.647277 | 6.06E-134 | greenyellow | HLA-DPA1 |
| 9.88E-132 | 1.038591 | 0.984663 | 0.802764 | 3.28E-131 | greenyellow | LSP1 |
| 3.92E-131 | 0.941944 | 0.788344 | 0.249878 | 1.29E-130 | greenyellow | PTPRE |
| 4.64E-131 | 0.681741 | 0.558282 | 0.129312 | 1.51E-130 | greenyellow | ATG101 |
| 5.94E-130 | 0.890768 | 0.644172 | 0.187092 | 1.92E-129 | greenyellow | C12orf45 |
| 2.85E-127 | 0.928284 | 0.97546 | 0.78345 | 9.11E-127 | greenyellow | ERP29 |
| 1.91E-125 | 1.120341 | 0.978528 | 0.704698 | 6.07E-125 | greenyellow | HSP90B1 |
| 1.21E-124 | 0.972829 | 0.993865 | 0.637295 | 3.80E-124 | greenyellow | HLA-DRB1 |
| 2.15E-122 | 0.739598 | 0.555215 | 0.142806 | 6.69E-122 | greenyellow | CORO1C |
| 7.13E-122 | 0.646025 | 0.41411 | 0.067826 | 2.20E-121 | greenyellow | CLIC3 |
| 1.21E-121 | 0.987926 | 0.929448 | 0.555422 | 3.71E-121 | greenyellow | CNPY3 |
| 7.16E-121 | 0.877778 | 0.641104 | 0.184881 | 2.17E-120 | greenyellow | SLC20A1 |
| 2.91E-120 | 0.97744 | 0.97546 | 0.625719 | 8.75E-120 | greenyellow | RNASET2 |
| 5.23E-118 | 0.702378 | 0.641104 | 0.175776 | 1.56E-117 | greenyellow | SPNS3 |
| 6.42E-118 | 0.932013 | 0.969325 | 0.792554 | 1.90E-117 | greenyellow | SSR4 |
| 2.61E-115 | 0.667833 | 0.677914 | 0.189628 | 7.65E-115 | greenyellow | HLA-DMB |
| 7.35E-115 | 0.728652 | 0.641104 | 0.17893 | 2.14E-114 | greenyellow | RAB11FIP1 |
| 1.98E-114 | 0.906175 | 0.935583 | 0.535165 | 5.71E-114 | greenyellow | TMED10 |
| 2.91E-114 | 0.672507 | 0.693252 | 0.185433 | 8.32E-114 | greenyellow | LIME1 |
| 1.39E-113 | 0.765318 | 0.98773 | 0.915461 | 3.95E-113 | greenyellow | SRP14 |
| 2.45E-113 | 0.731112 | 0.723926 | 0.229881 | 6.89E-113 | greenyellow | A1BG |
| 2.92E-113 | 0.613218 | 0.546012 | 0.125053 | 8.17E-113 | greenyellow | CD36 |

| | | | | | | |
|---|---|---|---|---|---|---|
| 1.47E-112 | 0.752759 | 0.57362 | 0.159941 | 4.08E-112 | greenyellow | IDH3A |
| 6.23E-111 | 0.847817 | 0.98773 | 0.649195 | 1.71E-110 | greenyellow | HLA-DPB1 |
| 1.11E-108 | -0.66994 | 0.969325 | 0.994082 | 3.03E-108 | greenyellow | RPL36 |
| 2.54E-108 | 1.016725 | 0.794479 | 0.333637 | 6.87E-108 | greenyellow | CDK2AP2 |
| 9.56E-108 | 0.981897 | 0.812883 | 0.308535 | 2.57E-107 | greenyellow | SAMHD1 |
| 2.49E-107 | 0.651536 | 0.429448 | 0.09101 | 6.63E-107 | greenyellow | CSF2RB |
| 1.24E-106 | 0.505475 | 0.328221 | 0.051861 | 3.29E-106 | greenyellow | ICAM1 |
| 4.83E-106 | 1.010084 | 0.325153 | 0.055113 | 1.27E-105 | greenyellow | MYBL2 |
| 9.26E-103 | 0.855965 | 0.742331 | 0.290782 | 2.41E-102 | greenyellow | MAN2B1 |
| 7.76E-102 | 1.103283 | 0.647239 | 0.224939 | 2.01E-101 | greenyellow | HLA-DQA2 |
| 4.73E-100 | 0.819885 | 0.935583 | 0.520241 | 1.21E-99 | greenyellow | CYBA |
| 1.23E-99 | 0.633231 | 0.46319 | 0.110974 | 3.13E-99 | greenyellow | FLNB |
| 1.44E-99 | 0.992327 | 0.828221 | 0.398472 | 3.64E-99 | greenyellow | DSTN |
| 8.35E-99 | 0.722157 | 0.5 | 0.128532 | 2.10E-98 | greenyellow | SLC7A5 |
| 3.67E-98 | 0.901342 | 0.693252 | 0.25986 | 9.16E-98 | greenyellow | SELENOS |
| 4.17E-97 | 0.817219 | 0.730061 | 0.297578 | 1.03E-96 | greenyellow | CCDC88A |
| 9.88E-96 | 0.823522 | 0.656442 | 0.211705 | 2.43E-95 | greenyellow | B4GALT1 |
| 2.27E-93 | 0.586412 | 0.466258 | 0.104959 | 5.56E-93 | greenyellow | LILRB2 |
| 3.97E-93 | 1.328589 | 0.453988 | 0.103788 | 9.64E-93 | greenyellow | GZMB |
| 6.58E-93 | 0.65232 | 0.631902 | 0.199675 | 1.59E-92 | greenyellow | PPM1K |
| 3.14E-92 | 0.821659 | 0.730061 | 0.309771 | 7.52E-92 | greenyellow | PDIA4 |
| 4.66E-92 | 0.558813 | 0.785276 | 0.302357 | 1.11E-91 | greenyellow | HLA-DQA1 |
| 1.68E-90 | 0.819418 | 0.91411 | 0.541831 | 3.96E-90 | greenyellow | P4HB |
| 2.53E-90 | 0.503544 | 0.352761 | 0.067469 | 5.93E-90 | greenyellow | SIDT1 |
| 8.66E-90 | -0.50808 | 0.98773 | 0.999415 | 2.02E-89 | greenyellow | RPL32 |
| 6.28E-89 | 0.767013 | 0.92638 | 0.667404 | 1.46E-88 | greenyellow | ATP6V0C |
| 8.04E-89 | 0.51922 | 0.377301 | 0.078361 | 1.85E-88 | greenyellow | RUBCN |
| 9.53E-89 | 0.642807 | 0.588957 | 0.187482 | 2.18E-88 | greenyellow | SFT2D2 |
| 5.35E-87 | -0.52429 | 0.984663 | 0.998894 | 1.21E-86 | greenyellow | RPL12 |
| 1.70E-86 | 0.75811 | 0.858896 | 0.447212 | 3.83E-86 | greenyellow | SPINT2 |
| 5.05E-86 | 0.589852 | 0.97546 | 0.953471 | 1.13E-85 | greenyellow | MYL6 |
| 4.15E-85 | 0.701714 | 0.634969 | 0.229946 | 9.25E-85 | greenyellow | NUDT22 |
| 1.54E-84 | 0.776657 | 0.892638 | 0.513868 | 3.41E-84 | greenyellow | OSTC |
| 3.31E-84 | 0.740013 | 0.552147 | 0.178052 | 7.28E-84 | greenyellow | RRBP1 |
| 4.67E-84 | 0.617138 | 0.481595 | 0.138514 | 1.02E-83 | greenyellow | GAPT |
| 3.77E-83 | 0.754495 | 0.806748 | 0.34622 | 8.19E-83 | greenyellow | IL16 |
| 4.20E-83 | 0.82235 | 0.911043 | 0.628093 | 9.07E-83 | greenyellow | SEC61G |
| 8.28E-83 | 0.635164 | 0.484663 | 0.140595 | 1.78E-82 | greenyellow | POLB |
| 3.95E-82 | 0.756651 | 0.717791 | 0.310551 | 8.44E-82 | greenyellow | BID |
| 4.13E-82 | 0.708616 | 0.644172 | 0.240904 | 8.75E-82 | greenyellow | SMIM3 |
| 2.71E-81 | 0.7835 | 0.91411 | 0.555259 | 5.72E-81 | greenyellow | CANX |

| | | | | | | |
|---|---|---|---|---|---|---|
| 7.93E-81 | -1.54699 | 0.331288 | 0.791774 | 1.66E-80 | greenyellow | CD52 |
| 8.19E-80 | 0.668716 | 0.610429 | 0.209039 | 1.71E-79 | greenyellow | THEMIS2 |
| 1.20E-79 | 0.692867 | 0.733129 | 0.332304 | 2.48E-79 | greenyellow | UBE2J1 |
| 2.96E-79 | 0.590309 | 0.54908 | 0.181076 | 6.10E-79 | greenyellow | C15orf39 |
| 1.71E-78 | 0.791661 | 0.95092 | 0.697448 | 3.49E-78 | greenyellow | UCP2 |
| 2.52E-78 | 0.519117 | 0.53681 | 0.149732 | 5.12E-78 | greenyellow | CYTH4 |
| 6.00E-77 | 0.6959 | 0.779141 | 0.381076 | 1.21E-76 | greenyellow | TMEM109 |
| 1.17E-76 | 0.734976 | 0.757669 | 0.3682 | 2.36E-76 | greenyellow | FKBP2 |
| 5.17E-75 | -2.13406 | 0.021472 | 0.578768 | 1.03E-74 | greenyellow | IFITM1 |
| 7.54E-75 | 0.69308 | 0.892638 | 0.545667 | 1.50E-74 | greenyellow | RPN2 |
| 2.71E-74 | 0.848431 | 0.613497 | 0.214989 | 5.36E-74 | greenyellow | RGS2 |
| 4.60E-73 | 0.543868 | 0.340491 | 0.077516 | 9.05E-73 | greenyellow | C3orf58 |
| 1.44E-72 | 0.625414 | 0.972393 | 0.893513 | 2.81E-72 | greenyellow | MYL12A |
| 2.26E-72 | 0.699745 | 0.92638 | 0.665973 | 4.40E-72 | greenyellow | VAMP8 |
| 8.76E-72 | 0.66349 | 0.932515 | 0.677841 | 1.69E-71 | greenyellow | PARK7 |
| 1.17E-71 | 0.664213 | 0.831288 | 0.455015 | 2.24E-71 | greenyellow | DHRS7 |
| 2.41E-71 | 0.649688 | 0.993865 | 0.906357 | 4.60E-71 | greenyellow | VIM |
| 2.53E-71 | 0.746921 | 0.920245 | 0.473061 | 4.81E-71 | greenyellow | STMN1 |
| 1.05E-70 | 0.631748 | 0.595092 | 0.210795 | 1.98E-70 | greenyellow | BHLHE40 |
| 1.43E-70 | 0.650472 | 0.677914 | 0.295627 | 2.69E-70 | greenyellow | MEF2C |
| 7.98E-70 | 0.593318 | 0.47546 | 0.152528 | 1.49E-69 | greenyellow | HYOU1 |
| 8.03E-69 | 0.505061 | 0.481595 | 0.147716 | 1.50E-68 | greenyellow | RASSF2 |
| 5.26E-67 | 0.602068 | 0.935583 | 0.773338 | 9.74E-67 | greenyellow | ARF1 |
| 7.31E-66 | 0.526612 | 0.539877 | 0.180036 | 1.35E-65 | greenyellow | UNC119 |
| 8.05E-66 | 0.584166 | 0.662577 | 0.268347 | 1.48E-65 | greenyellow | CCDC69 |
| 1.14E-65 | 0.72055 | 0.898773 | 0.605983 | 2.08E-65 | greenyellow | DBI |
| 1.83E-65 | 0.806814 | 0.696319 | 0.347976 | 3.31E-65 | greenyellow | TM9SF2 |
| 2.43E-63 | 0.644393 | 0.920245 | 0.675662 | 4.39E-63 | greenyellow | PDIA3 |
| 1.16E-62 | -1.05583 | 0.039877 | 0.537051 | 2.08E-62 | greenyellow | RARRES3 |
| 1.59E-62 | 0.704612 | 0.929448 | 0.67085 | 2.84E-62 | greenyellow | HSPA5 |
| 3.17E-62 | -1.0672 | 0.309816 | 0.685254 | 5.64E-62 | greenyellow | GYPC |
| 4.60E-62 | 0.65172 | 0.809816 | 0.459568 | 8.14E-62 | greenyellow | MYDGF |
| 2.11E-61 | -1.242 | 0.116564 | 0.590571 | 3.72E-61 | greenyellow | EVL |
| 3.18E-61 | 0.522516 | 0.990798 | 0.977597 | 5.56E-61 | greenyellow | CFL1 |
| 3.37E-61 | 0.575189 | 0.960123 | 0.806178 | 5.87E-61 | greenyellow | SUB1 |
| 4.10E-61 | 0.711933 | 0.634969 | 0.283271 | 7.11E-61 | greenyellow | LAIR1 |
| 1.25E-60 | 0.524567 | 0.493865 | 0.172427 | 2.15E-60 | greenyellow | RELL1 |
| 1.83E-60 | 0.680454 | 0.355828 | 0.091822 | 3.15E-60 | greenyellow | RGS1 |
| 4.56E-60 | 0.631715 | 0.831288 | 0.509673 | 7.78E-60 | greenyellow | RPN1 |
| 5.78E-59 | 0.613737 | 0.935583 | 0.569371 | 9.82E-59 | greenyellow | SELL |
| 7.24E-59 | 0.566479 | 0.886503 | 0.58345 | 1.22E-58 | greenyellow | ATP6V0B |

| | | | | | | |
|---|---|---|---|---|---|---|
| 7.47E-59 | 0.531651 | 0.653374 | 0.302878 | 1.26E-58 | greenyellow | SYK |
| 1.07E-58 | 0.516042 | 0.506135 | 0.177922 | 1.80E-58 | greenyellow | PARP10 |
| 1.61E-58 | 0.518043 | 0.97546 | 0.877158 | 2.69E-58 | greenyellow | YWHAZ |
| 1.74E-58 | 0.534806 | 0.429448 | 0.142091 | 2.88E-58 | greenyellow | PRMT7 |
| 1.87E-58 | 0.652249 | 0.644172 | 0.310096 | 3.09E-58 | greenyellow | SEC11C |
| 2.73E-58 | 0.587677 | 0.733129 | 0.300374 | 4.50E-58 | greenyellow | ARL4C |
| 3.51E-58 | 0.995096 | 0.766871 | 0.375516 | 5.75E-58 | greenyellow | LGALS1 |
| 5.77E-58 | 0.651178 | 0.472393 | 0.177077 | 9.40E-58 | greenyellow | EIF4A3 |
| 8.53E-58 | 0.692498 | 0.96319 | 0.58358 | 1.38E-57 | greenyellow | TXN |
| 4.79E-57 | -1.70744 | 0.453988 | 0.774606 | 7.73E-57 | greenyellow | JUNB |
| 6.36E-57 | 0.672191 | 0.674847 | 0.349894 | 1.02E-56 | greenyellow | AP3S1 |
| 1.39E-56 | 0.51752 | 0.91411 | 0.69088 | 2.22E-56 | greenyellow | PSAP |
| 3.99E-56 | 0.582788 | 0.920245 | 0.643017 | 6.36E-56 | greenyellow | PPP1R15A |
| 7.21E-56 | 0.595058 | 0.503067 | 0.201691 | 1.14E-55 | greenyellow | DCPS |
| 9.43E-55 | 0.508393 | 0.650307 | 0.276508 | 1.49E-54 | greenyellow | PRKCB |
| 1.07E-53 | 0.605544 | 0.766871 | 0.435084 | 1.69E-53 | greenyellow | SSR3 |
| 3.42E-53 | 0.665166 | 0.751534 | 0.407966 | 5.36E-53 | greenyellow | NR3C1 |
| 4.73E-53 | -1.28003 | 0.042945 | 0.487303 | 7.37E-53 | greenyellow | CD69 |
| 7.86E-53 | 0.678865 | 0.687117 | 0.342904 | 1.22E-52 | greenyellow | AC007952.4 |
| 8.84E-53 | 0.600223 | 0.754601 | 0.441652 | 1.36E-52 | greenyellow | DBNL |
| 1.30E-52 | 0.534819 | 0.803681 | 0.461811 | 2.00E-52 | greenyellow | TRABD |
| 3.79E-52 | 0.624768 | 0.650307 | 0.31842 | 5.80E-52 | greenyellow | GNA15 |
| 9.89E-52 | -0.79447 | 0.555215 | 0.841977 | 1.51E-51 | greenyellow | LIMD2 |
| 1.01E-51 | -1.09251 | 0.303681 | 0.691432 | 1.53E-51 | greenyellow | ANXA1 |
| 1.50E-51 | 0.598362 | 0.346626 | 0.100016 | 2.26E-51 | greenyellow | Z93241.1 |
| 3.18E-51 | 0.544978 | 0.684049 | 0.354284 | 4.79E-51 | greenyellow | AKIRIN2 |
| 3.84E-51 | -0.83214 | 0.248466 | 0.65482 | 5.74E-51 | greenyellow | LY6E |
| 4.48E-51 | 0.516211 | 0.883436 | 0.578345 | 6.68E-51 | greenyellow | 9-Sep |
| 1.99E-50 | 0.541296 | 0.837423 | 0.532629 | 2.96E-50 | greenyellow | MLF2 |
| 5.25E-50 | 0.623103 | 0.855828 | 0.543716 | 7.76E-50 | greenyellow | LAPTM4A |
| 1.12E-49 | -1.18041 | 0.046012 | 0.472574 | 1.65E-49 | greenyellow | TRBC2 |
| 1.52E-49 | 0.521852 | 0.957055 | 0.852707 | 2.22E-49 | greenyellow | RBM3 |
| 2.81E-49 | 0.825007 | 0.97546 | 0.854788 | 4.11E-49 | greenyellow | HMGN2 |
| 1.59E-48 | 0.587588 | 0.763804 | 0.445814 | 2.32E-48 | greenyellow | PHB |
| 3.96E-48 | 0.616065 | 0.773006 | 0.476833 | 5.74E-48 | greenyellow | TRAM1 |
| 4.71E-47 | -1.22793 | 0.521472 | 0.796521 | 6.79E-47 | greenyellow | TXNIP |
| 6.62E-47 | -1.18804 | 0.214724 | 0.611185 | 9.51E-47 | greenyellow | NEAT1 |
| 1.42E-46 | -1.02557 | 0.144172 | 0.560267 | 2.03E-46 | greenyellow | DNAJB1 |
| 1.17E-45 | 0.515365 | 0.530675 | 0.21073 | 1.66E-45 | greenyellow | AC016831.7 |
| 1.73E-45 | 0.553437 | 0.883436 | 0.641749 | 2.46E-45 | greenyellow | MTDH |
| 2.59E-45 | 0.689445 | 0.996933 | 0.972687 | 3.66E-45 | greenyellow | ACTG1 |

| | | | | | | |
|---|---|---|---|---|---|---|
| 3.47E-45 | 0.560772 | 0.812883 | 0.537636 | 4.88E-45 | greenyellow | DDOST |
| 5.62E-45 | 0.518413 | 0.57362 | 0.281027 | 7.88E-45 | greenyellow | TSPAN3 |
| 8.14E-45 | 0.527198 | 0.803681 | 0.532141 | 1.14E-44 | greenyellow | COPE |
| 1.96E-44 | 0.561545 | 0.800613 | 0.536336 | 2.73E-44 | greenyellow | TMED2 |
| 2.52E-44 | 0.501221 | 0.809816 | 0.506714 | 3.49E-44 | greenyellow | NUCB1 |
| 5.89E-44 | 0.529615 | 0.441718 | 0.182377 | 8.13E-44 | greenyellow | EIF2AK4 |
| 6.40E-44 | 0.595638 | 0.730061 | 0.453682 | 8.80E-44 | greenyellow | HM13 |
| 6.48E-44 | -2.14831 | 0 | 0.392294 | 8.88E-44 | greenyellow | IL32 |
| 5.13E-43 | 0.523565 | 0.59816 | 0.308633 | 7.00E-43 | greenyellow | SEC61A1 |
| 3.73E-42 | -1.00337 | 0.058282 | 0.442042 | 5.07E-42 | greenyellow | ZFP36L1 |
| 7.67E-42 | -1.05171 | 0.030675 | 0.409787 | 1.04E-41 | greenyellow | SLC2A3 |
| 3.75E-41 | 0.602755 | 0.702454 | 0.430889 | 5.06E-41 | greenyellow | CTSC |
| 3.86E-41 | -0.90572 | 0.595092 | 0.775744 | 5.18E-41 | greenyellow | IFITM2 |
| 8.33E-41 | 0.503285 | 0.935583 | 0.76121 | 1.12E-40 | greenyellow | TMBIM6 |
| 5.26E-40 | 0.507872 | 0.708589 | 0.420484 | 7.01E-40 | greenyellow | DNAJC4 |
| 5.90E-40 | 0.516149 | 0.794479 | 0.539002 | 7.84E-40 | greenyellow | SELENOF |
| 1.13E-39 | 0.526379 | 0.886503 | 0.669225 | 1.49E-39 | greenyellow | EIF4G2 |
| 4.03E-39 | 0.553033 | 0.634969 | 0.336108 | 5.32E-39 | greenyellow | OGT |
| 8.84E-39 | 0.517727 | 0.736196 | 0.470558 | 1.16E-38 | greenyellow | OS9 |
| 1.14E-38 | -0.96839 | 0.407975 | 0.688636 | 1.49E-38 | greenyellow | TSPO |
| 1.88E-38 | -0.95227 | 0.113497 | 0.478751 | 2.45E-38 | greenyellow | MT2A |
| 5.81E-38 | 0.513628 | 0.788344 | 0.533182 | 7.55E-38 | greenyellow | BRD2 |
| 1.66E-37 | 0.504864 | 0.628834 | 0.362705 | 2.15E-37 | greenyellow | ERCC1 |
| 1.87E-37 | 0.555702 | 0.684049 | 0.434694 | 2.41E-37 | greenyellow | KIF5B |
| 1.34E-36 | -0.6515 | 0.570552 | 0.741766 | 1.73E-36 | greenyellow | TPI1 |
| 2.68E-36 | 0.544644 | 0.496933 | 0.22728 | 3.43E-36 | greenyellow | AC245014.3 |
| 3.88E-36 | 0.580902 | 0.546012 | 0.293383 | 4.95E-36 | greenyellow | FYTTD1 |
| 1.79E-35 | 0.545113 | 0.481595 | 0.234336 | 2.27E-35 | greenyellow | TP53I13 |
| 3.49E-35 | 0.518219 | 0.779141 | 0.510486 | 4.42E-35 | greenyellow | PAXX |
| 4.29E-35 | -0.84807 | 0.196319 | 0.541538 | 5.41E-35 | greenyellow | FLNA |
| 5.66E-35 | -0.85445 | 0.150307 | 0.487433 | 7.12E-35 | greenyellow | IFITM3 |
| 7.23E-35 | -1.12205 | 0.254601 | 0.567127 | 9.05E-35 | greenyellow | CTSW |
| 6.87E-34 | -1.12738 | 0 | 0.322809 | 8.58E-34 | greenyellow | CD3D |
| 1.36E-33 | -0.60145 | 0.392638 | 0.666688 | 1.69E-33 | greenyellow | ICAM3 |
| 2.82E-33 | -1.02092 | 0.079755 | 0.413396 | 3.49E-33 | greenyellow | TIMP1 |
| 5.96E-33 | -0.94691 | 0.039877 | 0.362965 | 7.36E-33 | greenyellow | NR4A2 |
| 8.02E-32 | -0.62301 | 0.08589 | 0.409559 | 9.87E-32 | greenyellow | LCP2 |
| 1.03E-31 | -0.96188 | 0.015337 | 0.322939 | 1.27E-31 | greenyellow | LCK |
| 1.05E-31 | 0.617934 | 0.601227 | 0.343359 | 1.28E-31 | greenyellow | PLEK |
| 1.62E-31 | -0.74426 | 0.907975 | 0.947358 | 1.97E-31 | greenyellow | HLA-E |
| 2.50E-30 | -0.83793 | 0.165644 | 0.450593 | 3.04E-30 | greenyellow | CD48 |

| | | | | | | |
|---|---|---|---|---|---|---|
| 3.02E-30 | -0.56026 | 0.08589 | 0.407153 | 3.65E-30 | greenyellow | ITM2A |
| 3.41E-30 | -0.88452 | 0.009202 | 0.306649 | 4.11E-30 | greenyellow | SPOCK2 |
| 4.16E-29 | -1.41545 | 0.006135 | 0.293091 | 5.00E-29 | greenyellow | IL7R |
| 8.70E-29 | -0.80804 | 0.009202 | 0.294391 | 1.04E-28 | greenyellow | PIK3IP1 |
| 1.58E-28 | -1.55358 | 0.015337 | 0.299724 | 1.89E-28 | greenyellow | CST7 |
| 3.44E-28 | -0.65292 | 0.282209 | 0.525443 | 4.09E-28 | greenyellow | HSPB1 |
| 3.97E-28 | -0.61457 | 0.003067 | 0.282198 | 4.71E-28 | greenyellow | CD34 |
| 7.26E-28 | -1.09059 | 0.058282 | 0.338872 | 8.58E-28 | greenyellow | PRSS57 |
| 2.42E-27 | -0.66081 | 0.138037 | 0.431247 | 2.85E-27 | greenyellow | TBC1D10C |
| 4.53E-27 | -0.69515 | 0 | 0.269582 | 5.31E-27 | greenyellow | LAT |
| 6.04E-27 | -0.52615 | 0.208589 | 0.488994 | 7.06E-27 | greenyellow | PPA1 |
| 3.75E-26 | -0.62373 | 0.202454 | 0.487986 | 4.37E-26 | greenyellow | BIN2 |
| 7.03E-26 | -0.74941 | 0.003067 | 0.263664 | 8.17E-26 | greenyellow | HOPX |
| 8.94E-26 | -1.52486 | 0.162577 | 0.403869 | 1.04E-25 | greenyellow | CD3E |
| 1.23E-25 | 0.748532 | 0.650307 | 0.429979 | 1.42E-25 | greenyellow | GADD45B |
| 1.45E-25 | -0.74963 | 0.018405 | 0.278719 | 1.66E-25 | greenyellow | ZAP70 |
| 2.94E-25 | -0.78903 | 0 | 0.254463 | 3.37E-25 | greenyellow | CD3G |
| 6.51E-25 | -0.86878 | 0.003067 | 0.255178 | 7.44E-25 | greenyellow | GZMM |
| 7.47E-25 | -0.56269 | 0.04908 | 0.31842 | 8.51E-25 | greenyellow | AGTRAP |
| 9.52E-25 | -0.64757 | 0.021472 | 0.278134 | 1.08E-24 | greenyellow | CSF3R |
| 6.35E-24 | -0.6902 | 0.01227 | 0.258755 | 7.19E-24 | greenyellow | FLT3LG |
| 1.21E-23 | -0.56493 | 0.052147 | 0.3086 | 1.37E-23 | greenyellow | EGFL7 |
| 1.90E-23 | -0.52696 | 0.444785 | 0.657096 | 2.14E-23 | greenyellow | MSN |
| 5.21E-23 | -0.72332 | 0.021472 | 0.262429 | 5.84E-23 | greenyellow | KLF2 |
| 2.68E-22 | -0.57975 | 0.021472 | 0.257844 | 2.99E-22 | greenyellow | ABLIM1 |
| 5.36E-22 | -0.81205 | 0.690184 | 0.819932 | 5.97E-22 | greenyellow | TSC22D3 |
| 1.34E-21 | -0.58376 | 0.251534 | 0.500179 | 1.48E-21 | greenyellow | ANXA6 |
| 1.46E-21 | -0.84899 | 0.168712 | 0.425849 | 1.61E-21 | greenyellow | DUSP1 |
| 2.79E-21 | -0.61992 | 0.033742 | 0.26555 | 3.08E-21 | greenyellow | TUBA4A |
| 5.31E-21 | -0.94861 | 0.165644 | 0.412486 | 5.84E-21 | greenyellow | CD7 |
| 6.00E-21 | -0.93757 | 0.196319 | 0.424061 | 6.58E-21 | greenyellow | TNFAIP3 |
| 1.08E-20 | -0.58247 | 0.042945 | 0.270363 | 1.18E-20 | greenyellow | PRKCH |
| 8.72E-19 | -0.58517 | 0.055215 | 0.268054 | 9.50E-19 | greenyellow | LBH |
| 1.76E-18 | -0.51943 | 0.082822 | 0.308243 | 1.91E-18 | greenyellow | LYAR |
| 2.93E-18 | -0.54095 | 0.981595 | 0.986279 | 3.17E-18 | greenyellow | HLA-A |
| 1.04E-17 | -0.58572 | 0.223926 | 0.43141 | 1.13E-17 | greenyellow | HLA-F |
| 2.50E-17 | -0.54928 | 0.165644 | 0.379581 | 2.69E-17 | greenyellow | P2RY8 |
| 2.79E-17 | -0.52847 | 0.469325 | 0.643668 | 3.00E-17 | greenyellow | TUBA1A |
| 2.89E-17 | -0.61649 | 0.343558 | 0.490489 | 3.10E-17 | greenyellow | IMPDH2 |
| 6.32E-17 | -0.64489 | 0.358896 | 0.556755 | 6.74E-17 | greenyellow | LEPROTL1 |
| 2.96E-16 | -0.62051 | 0.659509 | 0.748919 | 3.15E-16 | greenyellow | PTPRC |

| | | | | | | |
|---|---|---|---|---|---|---|
| 6.62E-16 | -0.62023 | 0.365031 | 0.52967 | 7.02E-16 | greenyellow | BTG1 |
| 2.39E-15 | -0.56665 | 0.144172 | 0.332499 | 2.52E-15 | greenyellow | ISG20 |
| 6.70E-15 | -0.62447 | 0.214724 | 0.413461 | 7.06E-15 | greenyellow | NFKBIA |
| 7.89E-15 | -0.66195 | 0.122699 | 0.295822 | 8.29E-15 | greenyellow | FCMR |
| 6.41E-14 | -0.77092 | 0.453988 | 0.550577 | 6.71E-14 | greenyellow | AIF1 |
| 7.89E-13 | -2.2654 | 0.08589 | 0.252707 | 8.24E-13 | greenyellow | LYZ |
| 9.95E-13 | -0.67469 | 0.595092 | 0.609982 | 1.04E-12 | greenyellow | SERPINB1 |
| 8.99E-12 | -0.54259 | 0.107362 | 0.261909 | 9.34E-12 | greenyellow | SMIM24 |
| 9.58E-12 | -0.86989 | 0.254601 | 0.397984 | 9.92E-12 | greenyellow | LTB |
| 1.60E-10 | -0.53706 | 0.355828 | 0.510616 | 1.65E-10 | greenyellow | NOSIP |
| 2.19E-10 | -0.60074 | 0.110429 | 0.252869 | 2.26E-10 | greenyellow | SPINK2 |
| 7.06E-10 | -0.55147 | 0.144172 | 0.297545 | 7.24E-10 | greenyellow | NAMPT |
| 9.28E-10 | -0.60735 | 0.423313 | 0.520956 | 9.50E-10 | greenyellow | LITAF |
| 8.43E-09 | -0.50061 | 0.147239 | 0.280182 | 8.60E-09 | greenyellow | BLVRB |
| 1.37E-08 | -0.62682 | 0.242331 | 0.381922 | 1.39E-08 | greenyellow | LST1 |
| 5.37E-08 | -0.73932 | 0.809816 | 0.811738 | 5.45E-08 | greenyellow | S100A4 |
| 1.13E-05 | -0.53537 | 0.996933 | 0.994082 | 1.15E-05 | greenyellow | FTL |
| 0.016644 | -0.72323 | 0.322086 | 0.347098 | 0.016787 | greenyellow | S100A11 |
| 0 | 4.308029 | 0.708571 | 0.003527 | 0 | tan | CLC |
| 0 | 2.054001 | 0.72 | 0.008639 | 0 | tan | HDC |
| 0 | 2.004146 | 0.857143 | 0.041739 | 0 | tan | MS4A3 |
| 0 | 1.466738 | 0.56 | 0.007054 | 0 | tan | MS4A2 |
| 0 | 1.153442 | 0.485714 | 0.003462 | 0 | tan | PRG2 |
| 0 | 0.546662 | 0.302857 | 0.004303 | 0 | tan | ANGPT2 |
| 2.12E-262 | 1.161114 | 0.411429 | 0.021258 | 6.99E-261 | tan | TPSAB1 |
| 1.09E-196 | 1.146121 | 0.462857 | 0.035818 | 3.16E-195 | tan | HPGD |
| 1.52E-183 | 1.292346 | 0.817143 | 0.128616 | 3.90E-182 | tan | CPA3 |
| 3.09E-178 | 0.502862 | 0.257143 | 0.011939 | 7.14E-177 | tan | TPSD1 |
| 8.28E-158 | 1.530365 | 0.771429 | 0.128033 | 1.74E-156 | tan | RNASE2 |
| 3.94E-150 | 0.533419 | 0.428571 | 0.039604 | 7.58E-149 | tan | SLC45A3 |
| 2.32E-147 | 1.558008 | 0.72 | 0.119168 | 4.11E-146 | tan | FCER1A |
| 4.16E-143 | 1.254655 | 0.702857 | 0.11865 | 6.86E-142 | tan | LMO4 |
| 2.17E-138 | 0.736384 | 0.257143 | 0.015563 | 3.34E-137 | tan | TPSB2 |
| 1.75E-136 | 0.944014 | 0.611429 | 0.091633 | 2.53E-135 | tan | CSF2RB |
| 9.53E-129 | 1.355326 | 0.714286 | 0.139682 | 1.29E-127 | tan | GATA2 |
| 8.79E-124 | 0.735565 | 0.582857 | 0.089012 | 1.13E-122 | tan | EXD3 |
| 1.50E-101 | 0.790603 | 0.542857 | 0.094512 | 1.82E-100 | tan | HPGDS |
| 7.18E-100 | 1.237727 | 0.971429 | 0.33233 | 8.29E-99 | tan | PRSS57 |
| 8.84E-97 | 1.165964 | 0.788571 | 0.204815 | 9.73E-96 | tan | EREG |
| 2.51E-90 | 0.727832 | 0.571429 | 0.101275 | 2.63E-89 | tan | CNRIP1 |
| 1.80E-88 | 0.633559 | 0.537143 | 0.097813 | 1.81E-87 | tan | RHEX |

| | | | | | | |
|---|---|---|---|---|---|---|
| 7.82E-87 | 1.601784 | 0.994286 | 0.826603 | 7.53E-86 | tan | SRGN |
| 5.31E-79 | -1.46559 | 1 | 0.999676 | 4.90E-78 | tan | B2M |
| 1.92E-72 | 0.899118 | 0.942857 | 0.364136 | 1.71E-71 | tan | AC084033.3 |
| 7.97E-72 | 1.281606 | 0.457143 | 0.088074 | 6.82E-71 | tan | OSM |
| 1.21E-69 | 1.013605 | 0.971429 | 0.543325 | 1.00E-68 | tan | P4HB |
| 1.99E-64 | 0.877302 | 0.982857 | 0.848347 | 1.59E-63 | tan | PPIB |
| 1.84E-62 | 0.500771 | 0.994286 | 0.999256 | 1.42E-61 | tan | RPS24 |
| 1.05E-61 | 0.579381 | 0.554286 | 0.138776 | 7.81E-61 | tan | NAALADL1 |
| 1.76E-61 | 0.760304 | 0.6 | 0.166926 | 1.27E-60 | tan | TENT5A |
| 5.88E-60 | 0.787193 | 0.977143 | 0.696531 | 4.11E-59 | tan | SEC61B |
| 7.07E-59 | 0.846545 | 0.994286 | 0.891995 | 4.80E-58 | tan | RPS2 |
| 5.50E-57 | -1.40068 | 0.931429 | 0.993593 | 3.63E-56 | tan | HLA-C |
| 3.59E-56 | 1.016249 | 0.72 | 0.257814 | 2.30E-55 | tan | ALOX5AP |
| 3.88E-56 | 1.060593 | 0.651429 | 0.213777 | 2.42E-55 | tan | PLIN2 |
| 6.27E-56 | -1.32161 | 0.977143 | 0.997509 | 3.81E-55 | tan | HLA-B |
| 1.61E-55 | 0.76203 | 0.84 | 0.369637 | 9.53E-55 | tan | FKBP2 |
| 4.36E-55 | 0.970766 | 0.891429 | 0.467935 | 2.52E-54 | tan | CD63 |
| 4.69E-54 | 0.696864 | 0.502857 | 0.137125 | 2.64E-53 | tan | KIT |
| 6.38E-52 | 0.557626 | 0.514286 | 0.134375 | 3.51E-51 | tan | RFLNB |
| 7.28E-52 | 0.881787 | 0.977143 | 0.716883 | 3.91E-51 | tan | CALR |
| 3.96E-50 | -2.4243 | 0.8 | 0.919563 | 2.08E-49 | tan | CD74 |
| 8.35E-49 | 0.782641 | 0.965714 | 0.629166 | 4.29E-48 | tan | SEC61G |
| 1.35E-48 | 0.6376 | 0.668571 | 0.24128 | 6.76E-48 | tan | TRIM28 |
| 3.39E-48 | 0.672421 | 0.994286 | 0.793276 | 1.66E-47 | tan | SSR4 |
| 3.46E-48 | 0.577645 | 0.371429 | 0.078431 | 1.66E-47 | tan | NR4A3 |
| 4.65E-48 | -1.27658 | 0.794286 | 0.947809 | 2.19E-47 | tan | HLA-E |
| 9.67E-48 | 0.647111 | 0.748571 | 0.280787 | 4.47E-47 | tan | MGST1 |
| 2.26E-47 | -0.75776 | 1 | 0.999644 | 1.02E-46 | tan | ACTB |
| 8.18E-47 | 0.681157 | 0.794286 | 0.345305 | 3.63E-46 | tan | HMGA1 |
| 2.30E-46 | -1.71977 | 0.422857 | 0.789005 | 1.00E-45 | tan | CD52 |
| 3.10E-46 | 0.732185 | 0.845714 | 0.461076 | 1.33E-45 | tan | MYDGF |
| 4.24E-46 | 0.823279 | 0.828571 | 0.365592 | 1.78E-45 | tan | JUND |
| 1.46E-45 | 0.857342 | 0.382857 | 0.086941 | 6.01E-45 | tan | CTSG |
| 5.75E-45 | 0.568674 | 0.651429 | 0.2395 | 2.33E-44 | tan | VAT1 |
| 6.97E-45 | -1.60025 | 0.342857 | 0.796188 | 2.77E-44 | tan | TXNIP |
| 1.85E-44 | 0.635977 | 0.662857 | 0.252022 | 7.26E-44 | tan | RUNX1 |
| 2.97E-44 | -0.65894 | 0.954286 | 0.970038 | 1.14E-43 | tan | ARHGDIB |
| 5.63E-44 | 0.898545 | 0.725714 | 0.29237 | 2.13E-43 | tan | CLEC11A |
| 6.56E-44 | 1.063622 | 0.868571 | 0.482916 | 2.44E-43 | tan | GLUL |
| 7.67E-43 | 0.702891 | 0.817143 | 0.43642 | 2.81E-42 | tan | SSR3 |
| 6.18E-42 | 0.615349 | 0.805714 | 0.371449 | 2.23E-41 | tan | H2AFV |

| | | | | | | |
|---|---|---|---|---|---|---|
| 8.22E-42 | -1.48183 | 0.462857 | 0.820585 | 2.92E-41 | tan | TSC22D3 |
| 6.01E-41 | 0.502095 | 0.434286 | 0.11577 | 2.10E-40 | tan | CITED2 |
| 8.53E-41 | 0.608493 | 0.96 | 0.659937 | 2.94E-40 | tan | TMEM258 |
| 4.49E-40 | -0.8653 | 0.994286 | 0.99877 | 1.53E-39 | tan | TMSB10 |
| 3.68E-39 | 0.663779 | 0.845714 | 0.390118 | 1.23E-38 | tan | SOX4 |
| 2.21E-38 | -1.24877 | 0.354286 | 0.75021 | 7.31E-38 | tan | PTPRC |
| 1.44E-37 | 0.557733 | 0.782857 | 0.360933 | 4.69E-37 | tan | RNF130 |
| 2.46E-37 | -1.95744 | 0.44 | 0.773118 | 7.89E-37 | tan | JUNB |
| 2.74E-37 | -0.58303 | 0.988571 | 0.990358 | 8.66E-37 | tan | MT-ND4L |
| 7.12E-37 | 0.652042 | 0.874286 | 0.515822 | 2.22E-36 | tan | OSTC |
| 1.08E-36 | 0.864174 | 0.771429 | 0.401702 | 3.32E-36 | tan | ITM2A |
| 1.52E-36 | 0.513761 | 0.657143 | 0.274283 | 4.63E-36 | tan | ZNF593 |
| 2.31E-36 | 0.672397 | 0.914286 | 0.555588 | 6.94E-36 | tan | NUCB2 |
| 3.78E-36 | 0.510614 | 1 | 0.919627 | 1.12E-35 | tan | HMGB1 |
| 1.37E-34 | 0.540551 | 0.645714 | 0.26655 | 4.00E-34 | tan | FKBP11 |
| 1.51E-34 | 0.770242 | 0.954286 | 0.685854 | 4.35E-34 | tan | ANXA1 |
| 3.59E-34 | -0.70724 | 1 | 0.999191 | 1.02E-33 | tan | MALAT1 |
| 4.26E-34 | -1.29066 | 0.131429 | 0.588171 | 1.20E-33 | tan | EVL |
| 5.80E-34 | 0.530968 | 0.942857 | 0.642594 | 1.61E-33 | tan | MTDH |
| 1.43E-33 | 0.616866 | 0.788571 | 0.368796 | 3.95E-33 | tan | FABP5 |
| 2.48E-33 | -1.09184 | 0.594286 | 0.805863 | 6.75E-33 | tan | LSP1 |
| 1.18E-32 | 1.664451 | 0.302857 | 0.072316 | 3.16E-32 | tan | AZU1 |
| 6.39E-32 | -0.68534 | 0.754286 | 0.904776 | 1.70E-31 | tan | DAZAP2 |
| 2.12E-31 | -1.02402 | 0.651429 | 0.821459 | 5.57E-31 | tan | CORO1A |
| 5.33E-31 | -0.90039 | 0.942857 | 0.986475 | 1.38E-30 | tan | HLA-A |
| 5.89E-31 | -1.55002 | 0.337143 | 0.652656 | 1.51E-30 | tan | HLA-DPA1 |
| 1.30E-30 | 0.562192 | 0.565714 | 0.205656 | 3.29E-30 | tan | PLAUR |
| 7.49E-30 | 0.56161 | 0.445714 | 0.151556 | 1.88E-29 | tan | GADD45A |
| 1.09E-28 | 0.505601 | 0.634286 | 0.289232 | 2.72E-28 | tan | SNX5 |
| 2.23E-28 | -0.94841 | 0.177143 | 0.583835 | 5.48E-28 | tan | 9-Sep |
| 4.07E-28 | 0.558997 | 0.474286 | 0.176147 | 9.89E-28 | tan | H1FX |
| 7.47E-28 | -0.56409 | 0.817143 | 0.886818 | 1.80E-27 | tan | PSME1 |
| 8.24E-28 | 0.84432 | 0.685714 | 0.379764 | 1.96E-27 | tan | AKR1B1 |
| 8.85E-28 | -0.83098 | 0.268571 | 0.65531 | 2.09E-27 | tan | CD44 |
| 1.48E-27 | 0.544652 | 0.925714 | 0.691646 | 3.46E-27 | tan | NDUFB9 |
| 3.67E-27 | -0.92301 | 0.754286 | 0.886042 | 8.49E-27 | tan | CALM1 |
| 1.69E-26 | 0.617072 | 0.731429 | 0.422248 | 3.87E-26 | tan | CAMLG |
| 2.35E-25 | -1.86937 | 0 | 0.403611 | 5.31E-25 | tan | CD3E |
| 4.28E-25 | -0.60309 | 0.954286 | 0.980748 | 9.59E-25 | tan | UBC |
| 5.79E-25 | -1.20742 | 0.365714 | 0.645668 | 1.29E-24 | tan | ITGB2 |
| 6.10E-25 | -1.65275 | 0 | 0.398725 | 1.34E-24 | tan | LTB |

| | | | | | | |
|---|---|---|---|---|---|---|
| 7.03E-25 | -0.81393 | 0.582857 | 0.774154 | 1.53E-24 | tan | GPSM3 |
| 1.18E-24 | 0.573164 | 0.411429 | 0.145765 | 2.55E-24 | tan | XIST |
| 1.31E-24 | 0.581923 | 0.971429 | 0.826118 | 2.79E-24 | tan | MIF |
| 5.00E-24 | -0.8656 | 0.811429 | 0.906652 | 1.06E-23 | tan | SARAF |
| 7.57E-24 | -1.3155 | 0.428571 | 0.654015 | 1.59E-23 | tan | HLA-DPB1 |
| 7.94E-24 | -1.84781 | 0.268571 | 0.574646 | 1.65E-23 | tan | IFITM1 |
| 8.00E-24 | -2.13226 | 0.005714 | 0.390345 | 1.65E-23 | tan | IL32 |
| 1.28E-23 | -0.79192 | 0.754286 | 0.895134 | 2.61E-23 | tan | MYL12A |
| 1.39E-23 | -1.67049 | 0.422857 | 0.64227 | 2.82E-23 | tan | HLA-DRB1 |
| 3.16E-22 | -0.94216 | 0.091429 | 0.449621 | 6.34E-22 | tan | CD48 |
| 3.87E-22 | -0.56959 | 0.771429 | 0.86954 | 7.70E-22 | tan | ARPC2 |
| 7.64E-22 | -2.09919 | 0.4 | 0.603281 | 1.51E-21 | tan | HLA-DRA |
| 9.33E-22 | -0.6799 | 0.811429 | 0.927425 | 1.83E-21 | tan | LAPTM5 |
| 1.50E-21 | -1.0972 | 0.102857 | 0.466835 | 2.91E-21 | tan | CTSS |
| 1.06E-20 | -0.89097 | 0.217143 | 0.528959 | 2.03E-20 | tan | CCND3 |
| 1.94E-20 | -0.68529 | 0.994286 | 0.993691 | 3.71E-20 | tan | TMSB4X |
| 2.48E-20 | -0.72515 | 0.217143 | 0.536821 | 4.70E-20 | tan | RGS10 |
| 3.33E-20 | -0.76701 | 0.171429 | 0.499418 | 6.26E-20 | tan | ANXA6 |
| 4.81E-20 | -0.83204 | 0.52 | 0.701126 | 8.96E-20 | tan | UCP2 |
| 1.21E-19 | -0.68371 | 0.514286 | 0.712224 | 2.24E-19 | tan | HCLS1 |
| 6.23E-19 | -1.43029 | 0.36 | 0.609364 | 1.14E-18 | tan | S100A10 |
| 7.00E-19 | -0.98337 | 0 | 0.321523 | 1.27E-18 | tan | LCK |
| 7.35E-19 | -1.12407 | 0 | 0.321232 | 1.33E-18 | tan | CD3D |
| 1.53E-18 | -0.97787 | 0.44 | 0.709797 | 2.74E-18 | tan | ZFP36 |
| 3.65E-18 | -0.77105 | 0.108571 | 0.416974 | 6.49E-18 | tan | CYTIP |
| 4.37E-18 | -0.65814 | 0.028571 | 0.343234 | 7.70E-18 | tan | GLIPR1 |
| 5.17E-18 | -0.63879 | 0.022857 | 0.334175 | 9.06E-18 | tan | BIN1 |
| 5.85E-18 | -0.81868 | 0.245714 | 0.533456 | 1.02E-17 | tan | RARRES3 |
| 6.00E-18 | -1.36257 | 0.388571 | 0.582961 | 1.03E-17 | tan | HLA-DRB5 |
| 1.03E-17 | -0.9069 | 0 | 0.305248 | 1.76E-17 | tan | SPOCK2 |
| 4.78E-17 | -1.03673 | 0.068571 | 0.369928 | 8.11E-17 | tan | HLA-DQB1 |
| 9.00E-17 | -1.42397 | 0 | 0.291723 | 1.52E-16 | tan | IL7R |
| 1.04E-16 | -0.71412 | 0.12 | 0.417945 | 1.75E-16 | tan | RIPOR2 |
| 1.14E-16 | 0.558747 | 0.508571 | 0.254837 | 1.90E-16 | tan | H1F0 |
| 1.31E-16 | -0.79935 | 0.148571 | 0.430822 | 2.17E-16 | tan | HLA-F |
| 1.33E-16 | -0.8371 | 0.005714 | 0.295638 | 2.18E-16 | tan | FCMR |
| 1.36E-16 | -0.92949 | 0.268571 | 0.521355 | 2.21E-16 | tan | LITAF |
| 1.81E-16 | -0.68735 | 0.868571 | 0.907558 | 2.93E-16 | tan | CD37 |
| 2.27E-16 | -0.50449 | 0.828571 | 0.87847 | 3.65E-16 | tan | YWHAZ |
| 3.02E-16 | -0.66877 | 0.062857 | 0.352682 | 4.80E-16 | tan | IL16 |
| 3.03E-16 | -0.9662 | 0.485714 | 0.658513 | 4.80E-16 | tan | HCST |

| | | | | | | |
|---|---|---|---|---|---|---|
| 3.32E-16 | -0.71505 | 0.971429 | 0.968841 | 5.22E-16 | tan | SH3BGRL3 |
| 4.61E-16 | -0.80352 | 0.028571 | 0.31544 | 7.20E-16 | tan | SAMHD1 |
| 7.50E-16 | -0.54077 | 0.84 | 0.855983 | 1.16E-15 | tan | RAC2 |
| 8.38E-16 | -0.83565 | 0.148571 | 0.439656 | 1.29E-15 | tan | ZFP36L1 |
| 1.39E-15 | -0.81047 | 0.017143 | 0.292953 | 2.12E-15 | tan | PIK3IP1 |
| 3.75E-15 | -0.95548 | 0.217143 | 0.46952 | 5.70E-15 | tan | TRBC2 |
| 5.48E-15 | -0.67671 | 0.308571 | 0.553388 | 8.28E-15 | tan | ARHGEF1 |
| 6.29E-15 | -0.76205 | 0.034286 | 0.306445 | 9.44E-15 | tan | ARL4C |
| 1.42E-14 | -0.64146 | 0.788571 | 0.810328 | 2.10E-14 | tan | CD99 |
| 1.42E-14 | -0.72754 | 0.011429 | 0.27286 | 2.10E-14 | tan | IL10RA |
| 2.11E-14 | -0.74376 | 0.017143 | 0.277454 | 3.10E-14 | tan | ZAP70 |
| 2.27E-14 | -0.63576 | 0.011429 | 0.26943 | 3.31E-14 | tan | PRKCH |
| 3.03E-14 | -0.86178 | 0.045714 | 0.308904 | 4.41E-14 | tan | HLA-DQA1 |
| 3.22E-14 | -0.78636 | 0 | 0.253219 | 4.64E-14 | tan | CD3G |
| 4.17E-14 | -0.57028 | 0.754286 | 0.839449 | 5.98E-14 | tan | LIMD2 |
| 4.93E-14 | -0.56965 | 0.331429 | 0.560377 | 7.03E-14 | tan | WAS |
| 5.11E-14 | -0.66388 | 0.337143 | 0.555782 | 7.24E-14 | tan | STK17B |
| 5.24E-14 | -0.54184 | 0.634286 | 0.685757 | 7.38E-14 | tan | 6-Sep |
| 5.78E-14 | -0.52324 | 0.04 | 0.301107 | 8.10E-14 | tan | MEF2C |
| 7.78E-14 | -0.86444 | 0.005714 | 0.253931 | 1.08E-13 | tan | GZMM |
| 1.66E-13 | -0.59648 | 0.48 | 0.651524 | 2.30E-13 | tan | LY6E |
| 5.82E-13 | -0.62726 | 0.474286 | 0.641429 | 8.00E-13 | tan | CNN2 |
| 1.13E-12 | -0.55096 | 0.177143 | 0.420792 | 1.55E-12 | tan | SP100 |
| 1.40E-12 | -0.6286 | 0.474286 | 0.617129 | 1.91E-12 | tan | PSMB9 |
| 1.50E-12 | -1.21453 | 0.188571 | 0.41115 | 2.02E-12 | tan | CD7 |
| 1.51E-12 | -0.54499 | 0.074286 | 0.315861 | 2.03E-12 | tan | SLC44A2 |
| 1.56E-12 | -0.65804 | 0.034286 | 0.267133 | 2.09E-12 | tan | LBH |
| 1.61E-12 | -0.71877 | 0.028571 | 0.261211 | 2.14E-12 | tan | KLF2 |
| 2.63E-12 | -1.0187 | 0.205714 | 0.422895 | 3.48E-12 | tan | TNFAIP3 |
| 3.19E-12 | -0.55392 | 0.457143 | 0.630363 | 4.18E-12 | tan | RNASET2 |
| 3.83E-12 | -0.62172 | 0.194286 | 0.429496 | 5.00E-12 | tan | TBC1D10C |
| 4.68E-12 | -0.51803 | 0.08 | 0.316605 | 6.08E-12 | tan | TRIM22 |
| 4.76E-12 | -0.5962 | 0.068571 | 0.303857 | 6.15E-12 | tan | TAGAP |
| 5.15E-12 | -0.81274 | 0.571429 | 0.693911 | 6.61E-12 | tan | PSAP |
| 6.81E-12 | -0.90255 | 0.331429 | 0.497412 | 8.70E-12 | tan | FYB1 |
| 7.36E-12 | -0.66045 | 0.04 | 0.264382 | 9.35E-12 | tan | TUBA4A |
| 2.54E-11 | -0.55063 | 0.188571 | 0.407397 | 3.21E-11 | tan | LCP2 |
| 2.75E-11 | -0.58724 | 0.445714 | 0.615512 | 3.46E-11 | tan | MYH9 |
| 3.62E-11 | -0.52219 | 0.445714 | 0.621012 | 4.52E-11 | tan | TAPBP |
| 5.14E-11 | -0.71419 | 0.462857 | 0.573837 | 6.38E-11 | tan | SELL |
| 7.83E-11 | -0.6894 | 0.125714 | 0.331683 | 9.67E-11 | tan | ISG20 |

| | | | | | | |
|---|---|---|---|---|---|---|
| 8.85E-11 | -0.57995 | 0.28 | 0.486152 | 1.09E-10 | tan | BIN2 |
| 9.33E-11 | -0.56331 | 0.091429 | 0.30596 | 1.14E-10 | tan | GNG2 |
| 1.06E-10 | -0.61211 | 0.377143 | 0.558144 | 1.29E-10 | tan | YPEL5 |
| 1.18E-10 | -0.65109 | 0.051429 | 0.257329 | 1.43E-10 | tan | FLT3LG |
| 1.20E-10 | -0.7191 | 0.24 | 0.433379 | 1.44E-10 | tan | GADD45B |
| 1.60E-10 | -0.65899 | 0.051429 | 0.262117 | 1.91E-10 | tan | HOPX |
| 2.34E-10 | -0.6191 | 0.245714 | 0.440821 | 2.79E-10 | tan | PIK3R1 |
| 5.50E-10 | -0.57682 | 0.691429 | 0.777487 | 6.51E-10 | tan | JUN |
| 6.96E-10 | -0.6836 | 0.205714 | 0.413188 | 8.20E-10 | tan | DDIT4 |
| 8.30E-10 | -0.79368 | 0.228571 | 0.406814 | 9.74E-10 | tan | SLC2A3 |
| 1.06E-09 | -0.50907 | 0.4 | 0.622145 | 1.24E-09 | tan | RPS4Y1 |
| 1.50E-09 | -0.50831 | 0.348571 | 0.525626 | 1.74E-09 | tan | ANKRD12 |
| 3.09E-09 | -0.5537 | 0.325714 | 0.490358 | 3.57E-09 | tan | PRMT2 |
| 5.06E-09 | -0.55827 | 0.222857 | 0.421536 | 5.81E-09 | tan | BTG2 |
| 7.21E-09 | -0.56524 | 0.348571 | 0.5 | 8.25E-09 | tan | ARRB2 |
| 1.10E-08 | -0.50728 | 0.2 | 0.383744 | 1.25E-08 | tan | PPP1R18 |
| 1.34E-08 | 1.079773 | 0.262857 | 0.12716 | 1.51E-08 | tan | MPO |
| 2.22E-08 | -0.50294 | 0.137143 | 0.317608 | 2.50E-08 | tan | RNF125 |
| 2.57E-08 | -0.56079 | 0.28 | 0.440918 | 2.88E-08 | tan | 1-Sep |
| 3.93E-08 | -0.52873 | 0.297143 | 0.464602 | 4.38E-08 | tan | HIF1A |
| 5.15E-08 | -0.53422 | 0.308571 | 0.483628 | 5.72E-08 | tan | CD69 |
| 9.06E-08 | -0.56802 | 0.274286 | 0.445545 | 1.00E-07 | tan | IRF1 |
| 3.79E-07 | -0.68574 | 0.102857 | 0.254352 | 4.17E-07 | tan | NCF1 |
| 8.10E-07 | -0.54577 | 0.451429 | 0.545557 | 8.87E-07 | tan | EZR |
| 9.41E-07 | -0.55838 | 0.314286 | 0.442794 | 1.03E-06 | tan | VSIR |
| 2.73E-06 | -0.51221 | 0.382857 | 0.49657 | 2.96E-06 | tan | AES |
| 3.13E-06 | -0.62702 | 0.417143 | 0.499385 | 3.37E-06 | tan | HLA-DMA |
| 4.87E-06 | -0.50298 | 0.245714 | 0.370025 | 5.23E-06 | tan | RNF213 |
| 6.73E-06 | -0.58573 | 0.171429 | 0.307643 | 7.19E-06 | tan | ID2 |
| 9.52E-06 | -0.51358 | 0.28 | 0.391413 | 1.01E-05 | tan | TRAF3IP3 |
| 0.000103 | -0.86234 | 0.834286 | 0.81159 | 0.000109 | tan | S100A4 |
| 0.000104 | -0.59096 | 0.262857 | 0.360124 | 0.000109 | tan | NR4A2 |
| 0.000969 | -0.97092 | 0.28 | 0.347214 | 0.001017 | tan | S100A11 |
| 0.001073 | -0.54837 | 0.417143 | 0.475248 | 0.001122 | tan | MT2A |
| 0.003246 | 0.841873 | 0.28 | 0.199185 | 0.003377 | tan | HSPA1A |
| 0.003353 | -0.56622 | 0.24 | 0.301107 | 0.003474 | tan | ANXA2 |
| 0 | 3.268949 | 0.987903 | 0.103459 | 0 | royalblue | S100A8 |
| 0 | 3.123391 | 0.985023 | 0.107173 | 0 | royalblue | S100A9 |
| 0 | 2.895284 | 0.948733 | 0.056602 | 0 | royalblue | S100A12 |
| 0 | 2.436901 | 0.971198 | 0.080831 | 0 | royalblue | FCN1 |
| 0 | 2.392616 | 0.987327 | 0.207395 | 0 | royalblue | LYZ |

| | | | | | | |
|---|---|---|---|---|---|---|
| 0 | 2.34613 | 0.926267 | 0.065667 | 0 | royalblue | VCAN |
| 0 | 2.130508 | 0.986175 | 0.286216 | 0 | royalblue | TYROBP |
| 0 | 2.103125 | 0.689516 | 0.049583 | 0 | royalblue | G0S2 |
| 0 | 2.049205 | 0.990783 | 0.813904 | 0 | royalblue | S100A6 |
| 0 | 2.024363 | 0.858871 | 0.16916 | 0 | royalblue | PLAUR |
| 0 | 1.944188 | 0.821429 | 0.052343 | 0 | royalblue | CD14 |
| 0 | 1.936872 | 0.972926 | 0.602385 | 0 | royalblue | SAT1 |
| 0 | 1.921458 | 0.982719 | 0.373965 | 0 | royalblue | CST3 |
| 0 | 1.891669 | 0.861751 | 0.0641 | 0 | royalblue | SERPINA1 |
| 0 | 1.876247 | 0.957949 | 0.310683 | 0 | royalblue | S100A11 |
| 0 | 1.745141 | 0.918203 | 0.437962 | 0 | royalblue | CTSS |
| 0 | 1.731017 | 0.850806 | 0.107003 | 0 | royalblue | CSTA |
| 0 | 1.728331 | 0.849654 | 0.263179 | 0 | royalblue | NAMPT |
| 0 | 1.714892 | 0.445853 | 0.041983 | 0 | royalblue | CXCL8 |
| 0 | 1.700738 | 0.898041 | 0.395059 | 0 | royalblue | DUSP1 |
| 0 | 1.694307 | 0.959677 | 0.586165 | 0 | royalblue | NEAT1 |
| 0 | 1.676815 | 0.808756 | 0.104549 | 0 | royalblue | LGALS3 |
| 0 | 1.661845 | 0.999424 | 0.993798 | 0 | royalblue | FTL |
| 0 | 1.661757 | 0.964286 | 0.669211 | 0 | royalblue | TSPO |
| 0 | 1.641747 | 0.930876 | 0.21772 | 0 | royalblue | FCER1G |
| 0 | 1.63467 | 0.785714 | 0.208076 | 0 | royalblue | ATP2B1-AS1 |
| 0 | 1.627475 | 0.979263 | 0.585994 | 0 | royalblue | S100A10 |
| 0 | 1.592885 | 0.773618 | 0.113273 | 0 | royalblue | TYMP |
| 0 | 1.58461 | 0.988479 | 0.801261 | 0 | royalblue | S100A4 |
| 0 | 1.562929 | 0.867512 | 0.382825 | 0 | royalblue | TIMP1 |
| 0 | 1.553924 | 0.796083 | 0.127517 | 0 | royalblue | CD68 |
| 0 | 1.54128 | 0.711982 | 0.082535 | 0 | royalblue | SLC11A1 |
| 0 | 1.540802 | 0.698157 | 0.050775 | 0 | royalblue | TMEM176B |
| 0 | 1.517766 | 0.799539 | 0.160436 | 0 | royalblue | COTL1 |
| 0 | 1.476087 | 0.751152 | 0.137604 | 0 | royalblue | CFD |
| 0 | 1.456643 | 0.892857 | 0.350145 | 0 | royalblue | LST1 |
| 0 | 1.456179 | 0.614055 | 0.039189 | 0 | royalblue | PLBD1 |
| 0 | 1.45074 | 0.960829 | 0.525234 | 0 | royalblue | AIF1 |
| 0 | 1.442775 | 0.618664 | 0.038201 | 0 | royalblue | C5AR1 |
| 0 | 1.424761 | 0.943548 | 0.34626 | 0 | royalblue | LGALS1 |
| 0 | 1.41368 | 0.710253 | 0.190118 | 0 | royalblue | RGS2 |
| 0 | 1.407692 | 0.453341 | 0.03418 | 0 | royalblue | THBS1 |
| 0 | 1.402134 | 0.669355 | 0.067473 | 0 | royalblue | MS4A6A |
| 0 | 1.401013 | 0.557604 | 0.021537 | 0 | royalblue | RBP7 |
| 0 | 1.399964 | 0.718318 | 0.128642 | 0 | royalblue | CFP |
| 0 | 1.397821 | 1 | 0.996286 | 0 | royalblue | FTH1 |

| | | | | | | |
|---|---|---|---|---|---|---|
| 0 | 1.392322 | 0.658986 | 0.054013 | 0 | royalblue | CLEC7A |
| 0 | 1.378613 | 0.934332 | 0.678957 | 0 | royalblue | PSAP |
| 0 | 1.368363 | 0.789747 | 0.343943 | 0 | royalblue | GRN |
| 0 | 1.367793 | 0.529378 | 0.037519 | 0 | royalblue | MAFB |
| 0 | 1.316251 | 0.477535 | 0.032919 | 0 | royalblue | APOBEC3A |
| 0 | 1.304193 | 0.641129 | 0.133924 | 0 | royalblue | CTSD |
| 0 | 1.287368 | 0.577765 | 0.0351 | 0 | royalblue | CDA |
| 0 | 1.286002 | 0.578341 | 0.040211 | 0 | royalblue | FPR1 |
| 0 | 1.285941 | 0.835253 | 0.461169 | 0 | royalblue | MCL1 |
| 0 | 1.2701 | 0.631912 | 0.167661 | 0 | royalblue | STXBP2 |
| 0 | 1.266115 | 0.574885 | 0.077526 | 0 | royalblue | KLF4 |
| 0 | 1.256664 | 0.50288 | 0.069927 | 0 | royalblue | MNDA |
| 0 | 1.240489 | 0.56106 | 0.046482 | 0 | royalblue | LILRA5 |
| 0 | 1.23654 | 0.728111 | 0.248662 | 0 | royalblue | CSF3R |
| 0 | 1.235288 | 0.554147 | 0.055137 | 0 | royalblue | TREM1 |
| 0 | 1.21422 | 0.981567 | 0.818436 | 0 | royalblue | SRGN |
| 0 | 1.209708 | 0.50288 | 0.065156 | 0 | royalblue | ANPEP |
| 0 | 1.205418 | 0.572005 | 0.058068 | 0 | royalblue | NCF2 |
| 0 | 1.187261 | 0.589862 | 0.16732 | 0 | royalblue | BCL2A1 |
| 0 | 1.182937 | 0.698157 | 0.394411 | 0 | royalblue | NFKBIA |
| 0 | 1.179126 | 0.58871 | 0.095451 | 0 | royalblue | CYBB |
| 0 | 1.171857 | 0.718318 | 0.306594 | 0 | royalblue | FCGRT |
| 0 | 1.154544 | 0.492512 | 0.037076 | 0 | royalblue | TMEM176A |
| 0 | 1.138847 | 0.764401 | 0.418265 | 0 | royalblue | APLP2 |
| 0 | 1.137565 | 0.987903 | 0.70213 | 0 | royalblue | FOS |
| 0 | 1.119331 | 0.74712 | 0.447981 | 0 | royalblue | ASAH1 |
| 0 | 1.116868 | 0.502304 | 0.06093 | 0 | royalblue | CEBPD |
| 0 | 1.099707 | 0.940668 | 0.799148 | 0 | royalblue | GABARAP |
| 0 | 1.091936 | 0.514401 | 0.106696 | 0 | royalblue | CD36 |
| 0 | 1.089506 | 0.555876 | 0.082297 | 0 | royalblue | LILRB2 |
| 0 | 1.08681 | 0.355415 | 0.033873 | 0 | royalblue | RETN |
| 0 | 1.085826 | 0.660138 | 0.229443 | 0 | royalblue | NCF1 |
| 0 | 1.072969 | 0.500576 | 0.060862 | 0 | royalblue | IGSF6 |
| 0 | 1.066703 | 0.691244 | 0.293372 | 0 | royalblue | AGTRAP |
| 0 | 1.065654 | 0.84735 | 0.571205 | 0 | royalblue | ATP6V0B |
| 0 | 1.055186 | 0.488479 | 0.051525 | 0 | royalblue | TNFAIP2 |
| 0 | 1.054438 | 0.642281 | 0.232646 | 0 | royalblue | PTPRE |
| 0 | 1.05438 | 0.788018 | 0.465463 | 0 | royalblue | PYCARD |
| 0 | 1.047497 | 0.663594 | 0.350724 | 0 | royalblue | JUND |
| 0 | 1.046522 | 0.501728 | 0.048151 | 0 | royalblue | PILRA |
| 0 | 1.045603 | 0.589286 | 0.215301 | 0 | royalblue | NINJ1 |

| | | | | | | |
|---|---|---|---|---|---|---|
| 0 | 1.043221 | 0.474078 | 0.059976 | 0 | royalblue | FGL2 |
| 0 | 1.039223 | 0.436636 | 0.033703 | 0 | royalblue | CD300E |
| 0 | 1.031173 | 0.702189 | 0.277015 | 0 | royalblue | ANXA2 |
| 0 | 1.016829 | 0.653802 | 0.303357 | 0 | royalblue | GNAI2 |
| 0 | 1.013493 | 0.580645 | 0.203953 | 0 | royalblue | FMNL1 |
| 0 | 1.01202 | 0.47235 | 0.050026 | 0 | royalblue | IFI30 |
| 0 | 1.011876 | 0.56394 | 0.112387 | 0 | royalblue | CTSH |
| 0 | 1.009208 | 0.440092 | 0.073164 | 0 | royalblue | SGK1 |
| 0 | 0.996958 | 0.50576 | 0.120463 | 0 | royalblue | C19orf38 |
| 0 | 0.992854 | 0.875576 | 0.65299 | 0 | royalblue | KLF6 |
| 0 | 0.987557 | 0.9303 | 0.719884 | 0 | royalblue | EMP3 |
| 0 | 0.98746 | 0.550691 | 0.12946 | 0 | royalblue | TNFRSF1B |
| 0 | 0.984543 | 0.734447 | 0.424774 | 0 | royalblue | VSIR |
| 0 | 0.981322 | 0.979263 | 0.918726 | 0 | royalblue | OAZ1 |
| 0 | 0.979485 | 0.567972 | 0.229102 | 0 | royalblue | SPI1 |
| 0 | 0.979085 | 0.551267 | 0.144352 | 0 | royalblue | UPP1 |
| 0 | 0.968219 | 0.665899 | 0.306969 | 0 | royalblue | ANXA5 |
| 0 | 0.964744 | 0.989631 | 0.902402 | 0 | royalblue | VIM |
| 0 | 0.960025 | 0.881336 | 0.625013 | 0 | royalblue | TKT |
| 0 | 0.947102 | 0.581797 | 0.133583 | 0 | royalblue | FGR |
| 0 | 0.940677 | 0.438364 | 0.040007 | 0 | royalblue | LRP1 |
| 0 | 0.938884 | 0.764977 | 0.534333 | 0 | royalblue | NPC2 |
| 0 | 0.919259 | 0.991359 | 0.967524 | 0 | royalblue | SH3BGRL3 |
| 0 | 0.899251 | 0.362327 | 0.055035 | 0 | royalblue | IER3 |
| 0 | 0.894411 | 0.38076 | 0.041813 | 0 | royalblue | AC020656.1 |
| 0 | 0.883108 | 0.415323 | 0.042903 | 0 | royalblue | SLC7A7 |
| 0 | 0.877915 | 0.52765 | 0.201806 | 0 | royalblue | FOSL2 |
| 0 | 0.867867 | 0.406106 | 0.058681 | 0 | royalblue | RAB31 |
| 0 | 0.860884 | 0.455069 | 0.120225 | 0 | royalblue | PLEC |
| 0 | 0.83538 | 0.396889 | 0.056125 | 0 | royalblue | TGFBI |
| 0 | 0.83183 | 0.417051 | 0.08642 | 0 | royalblue | LILRA2 |
| 0 | 0.828527 | 0.339862 | 0.041949 | 0 | royalblue | BASP1 |
| 0 | 0.824272 | 0.262097 | 0.032578 | 0 | royalblue | LGALS2 |
| 0 | 0.824145 | 0.377304 | 0.062634 | 0 | royalblue | MPEG1 |
| 0 | 0.822074 | 0.867512 | 0.60368 | 0 | royalblue | TALDO1 |
| 0 | 0.819248 | 0.335253 | 0.029715 | 0 | royalblue | MCEMP1 |
| 0 | 0.811524 | 0.417627 | 0.091498 | 0 | royalblue | STX11 |
| 0 | 0.805025 | 0.982143 | 0.947282 | 0 | royalblue | ATP5F1E |
| 0 | 0.797481 | 0.375576 | 0.076061 | 0 | royalblue | SLC43A2 |
| 0 | 0.79482 | 0.334677 | 0.046447 | 0 | royalblue | DMXL2 |
| 0 | 0.794122 | 0.870968 | 0.788209 | 0 | royalblue | ELOB |

| | | | | | | |
|---|---|---|---|---|---|---|
| 0 | 0.790161 | 0.364631 | 0.047504 | 0 | royalblue | CXCL16 |
| 0 | 0.790041 | 0.463134 | 0.1191 | 0 | royalblue | LY86 |
| 0 | 0.789315 | 0.351382 | 0.029102 | 0 | royalblue | SMIM25 |
| 0 | 0.786075 | 0.321429 | 0.035781 | 0 | royalblue | TRIB1 |
| 0 | 0.780854 | 0.315092 | 0.015846 | 0 | royalblue | CLEC4E |
| 0 | 0.761534 | 0.982719 | 0.951985 | 0 | royalblue | MYL6 |
| 0 | 0.757749 | 0.335253 | 0.046856 | 0 | royalblue | ITGAX |
| 0 | 0.756592 | 0.360599 | 0.063793 | 0 | royalblue | MIR22HG |
| 0 | 0.745618 | 0.392281 | 0.088465 | 0 | royalblue | ITGAM |
| 0 | 0.740325 | 0.289747 | 0.059261 | 0 | royalblue | PADI4 |
| 0 | 0.724998 | 0.363479 | 0.093133 | 0 | royalblue | GRINA |
| 0 | 0.718127 | 0.994816 | 0.998978 | 0 | royalblue | TMSB10 |
| 0 | 0.71606 | 0.303571 | 0.019969 | 0 | royalblue | HK3 |
| 0 | 0.70807 | 0.288594 | 0.02992 | 0 | royalblue | STAB1 |
| 0 | 0.691501 | 0.332373 | 0.056194 | 0 | royalblue | HCK |
| 0 | 0.67851 | 0.331221 | 0.0748 | 0 | royalblue | CTSZ |
| 0 | 0.675277 | 0.993664 | 0.981462 | 0 | royalblue | SERF2 |
| 0 | 0.67262 | 0.282258 | 0.026069 | 0 | royalblue | LILRB3 |
| 0 | 0.668778 | 0.271889 | 0.029307 | 0 | royalblue | CKAP4 |
| 0 | 0.663879 | 0.284562 | 0.036258 | 0 | royalblue | TMEM170B |
| 0 | 0.658897 | 0.934908 | 0.913103 | 0 | royalblue | ARPC3 |
| 0 | 0.653055 | 0.256912 | 0.034997 | 0 | royalblue | CES1 |
| 0 | 0.651718 | 0.3053 | 0.063691 | 0 | royalblue | BRI3 |
| 0 | 0.647766 | 0.260945 | 0.022218 | 0 | royalblue | OSCAR |
| 0 | 0.63436 | 0.25288 | 0.025626 | 0 | royalblue | LRRK2 |
| 0 | 0.630231 | 0.292627 | 0.049105 | 0 | royalblue | CD86 |
| 0 | 0.603521 | 0.282258 | 0.036361 | 0 | royalblue | SULF2 |
| 0 | 0.601863 | 0.259217 | 0.025967 | 0 | royalblue | QPCT |
| 0 | 0.597673 | 0.263825 | 0.037178 | 0 | royalblue | HMOX1 |
| 0 | 0.590405 | 0.268433 | 0.043415 | 0 | royalblue | CLEC4A |
| 0 | 0.556846 | 0.261521 | 0.043278 | 0 | royalblue | LRRC25 |
| 0 | -0.50403 | 0.913594 | 0.987834 | 0 | royalblue | RPL35 |
| 0 | -0.50564 | 0.995392 | 0.999523 | 0 | royalblue | RPL32 |
| 0 | -0.53546 | 0.925691 | 0.99022 | 0 | royalblue | RPL21 |
| 0 | -0.54014 | 0.991359 | 0.999455 | 0 | royalblue | RPL13 |
| 0 | -0.54227 | 0.990207 | 0.999148 | 0 | royalblue | RPL18A |
| 0 | -0.56916 | 0.960253 | 0.997444 | 0 | royalblue | RPL15 |
| 0 | -0.60247 | 0.948157 | 0.995263 | 0 | royalblue | RACK1 |
| 0 | -0.60898 | 0.99712 | 0.999864 | 0 | royalblue | RPL10 |
| 0 | -0.61842 | 0.985023 | 0.99891 | 0 | royalblue | RPL7A |
| 0 | -0.62416 | 0.993088 | 0.999148 | 0 | royalblue | RPS15A |

| | | | | | | |
|---|---|---|---|---|---|---|
| 0 | -0.62595 | 0.841014 | 0.973658 | 0 | royalblue | RPL4 |
| 0 | -0.64386 | 0.993664 | 0.999625 | 0 | royalblue | RPS27A |
| 0 | -0.67131 | 0.993088 | 0.999421 | 0 | royalblue | RPS23 |
| 0 | -0.69222 | 0.975806 | 0.998875 | 0 | royalblue | RPS19 |
| 0 | -0.69698 | 0.583525 | 0.936003 | 0 | royalblue | CIRBP |
| 0 | -0.7147 | 0.99424 | 0.999693 | 0 | royalblue | RPS12 |
| 0 | -0.76178 | 0.959677 | 0.997751 | 0 | royalblue | RPL14 |
| 0 | -0.78264 | 0.949309 | 0.997581 | 0 | royalblue | RPS25 |
| 0 | -0.78555 | 0.885945 | 0.994445 | 0 | royalblue | RPL23A |
| 0 | -0.79067 | 0.700461 | 0.927892 | 0 | royalblue | HINT1 |
| 0 | -0.79214 | 0.90841 | 0.995195 | 0 | royalblue | MT-ND4L |
| 0 | -0.80674 | 0.4303 | 0.833532 | 0 | royalblue | NCL |
| 0 | -0.83182 | 0.8053 | 0.976214 | 0 | royalblue | EEF2 |
| 0 | -0.85272 | 0.987327 | 0.999489 | 0 | royalblue | RPS3A |
| 0 | -0.92051 | 0.968318 | 0.998058 | 0 | royalblue | RPS18 |
| 0 | -0.92259 | 0.940668 | 0.995945 | 0 | royalblue | RPL5 |
| 0 | -0.92356 | 0.960829 | 0.997376 | 0 | royalblue | RPLP0 |
| 0 | -0.93113 | 0.235599 | 0.814858 | 0 | royalblue | C12orf57 |
| 0 | -0.94249 | 0.156682 | 0.716749 | 0 | royalblue | 6-Sep |
| 0 | -0.95222 | 0.952189 | 0.997103 | 0 | royalblue | RPS27 |
| 0 | -0.96237 | 0.979839 | 0.99908 | 0 | royalblue | RPS3 |
| 0 | -0.96359 | 0.830069 | 0.987868 | 0 | royalblue | RPL10A |
| 0 | -0.98169 | 0.980415 | 0.998875 | 0 | royalblue | RPS4X |
| 0 | -1.00989 | 0.995392 | 0.999864 | 0 | royalblue | EEF1A1 |
| 0 | -1.02229 | 0.733871 | 0.971341 | 0 | royalblue | HNRNPA1 |
| 0 | -1.02334 | 0.163018 | 0.713818 | 0 | royalblue | PEBP1 |
| 0 | -1.02898 | 0.927419 | 0.99741 | 0 | royalblue | RPS6 |
| 0 | -1.06756 | 0.414747 | 0.886454 | 0 | royalblue | HSPA8 |
| 0 | -1.09015 | 0.171083 | 0.711501 | 0 | royalblue | GYPC |
| 0 | -1.17165 | 0.796083 | 0.976793 | 0 | royalblue | EEF1B2 |
| 0 | -1.18038 | 0.826613 | 0.99022 | 0 | royalblue | RPSA |
| 0 | -1.18815 | 0.849078 | 0.990629 | 0 | royalblue | RPS5 |
| 0 | -1.25598 | 0.895737 | 0.994241 | 0 | royalblue | RPL3 |
| 0 | -1.26498 | 0.507488 | 0.890373 | 0 | royalblue | HSP90AB1 |
| 0 | -1.31804 | 0.047235 | 0.617448 | 0 | royalblue | EVL |
| 0 | -1.33478 | 0.014401 | 0.510342 | 0 | royalblue | CD69 |
| 0 | -1.3793 | 0.338134 | 0.83527 | 0 | royalblue | LDHB |
| 0 | -1.54835 | 0.006912 | 0.596797 | 0 | royalblue | CTSW |
| 0 | -1.57833 | 0.524194 | 0.942034 | 0 | royalblue | NPM1 |
| 0 | -1.85465 | 0.059908 | 0.603271 | 0 | royalblue | IFITM1 |
| 4.11E-306 | 0.804905 | 0.604839 | 0.259499 | 9.74E-306 | royalblue | BLVRB |

| | | | | | | |
|---|---|---|---|---|---|---|
| 2.64E-304 | 0.95478 | 0.641705 | 0.34108 | 6.23E-304 | royalblue | CTSB |
| 9.96E-304 | -1.27722 | 0.006336 | 0.495417 | 2.34E-303 | royalblue | TRBC2 |
| 1.94E-301 | 0.667744 | 0.293779 | 0.063793 | 4.54E-301 | royalblue | ASGR1 |
| 1.93E-300 | -1.73913 | 0.020161 | 0.504822 | 4.48E-300 | royalblue | STMN1 |
| 1.97E-299 | 0.581333 | 0.262673 | 0.050264 | 4.56E-299 | royalblue | EMILIN2 |
| 8.14E-298 | 0.7556 | 0.425115 | 0.128267 | 1.88E-297 | royalblue | JAML |
| 6.34E-295 | 1.058357 | 0.702765 | 0.449514 | 1.46E-294 | royalblue | HIF1A |
| 1.20E-294 | -0.73263 | 0.036866 | 0.533379 | 2.75E-294 | royalblue | TSTD1 |
| 2.86E-293 | 0.855885 | 0.737327 | 0.386199 | 6.49E-293 | royalblue | SLC2A3 |
| 3.14E-291 | 0.799447 | 0.815668 | 0.688363 | 7.11E-291 | royalblue | LAMTOR4 |
| 1.59E-290 | 0.994481 | 0.382488 | 0.111092 | 3.58E-290 | royalblue | DUSP6 |
| 7.91E-290 | -0.94877 | 0.568548 | 0.926086 | 1.77E-289 | royalblue | SARAF |
| 5.45E-289 | -1.05642 | 0.114055 | 0.583847 | 1.22E-288 | royalblue | NUCB2 |
| 1.30E-284 | 0.631509 | 0.291475 | 0.063759 | 2.90E-284 | royalblue | ODF3B |
| 1.25E-283 | 0.822906 | 0.507488 | 0.195843 | 2.76E-283 | royalblue | THEMIS2 |
| 4.52E-283 | 0.799213 | 0.410714 | 0.132322 | 9.95E-283 | royalblue | ZNF385A |
| 8.97E-283 | 0.842201 | 0.570853 | 0.241813 | 1.97E-282 | royalblue | MYO1F |
| 8.56E-280 | 0.669015 | 0.895737 | 0.799284 | 1.87E-279 | royalblue | LSP1 |
| 3.96E-279 | 0.947077 | 0.551843 | 0.255546 | 8.62E-279 | royalblue | TNFSF13B |
| 6.59E-277 | 0.546041 | 0.285138 | 0.059976 | 1.43E-276 | royalblue | ALOX5 |
| 3.64E-276 | 0.588621 | 0.272465 | 0.058136 | 7.84E-276 | royalblue | RXRA |
| 4.29E-275 | 0.504392 | 0.994816 | 0.993219 | 9.20E-275 | royalblue | MT-ND4 |
| 1.53E-273 | 0.761369 | 0.361175 | 0.104277 | 3.27E-273 | royalblue | TLE3 |
| 3.76E-272 | 0.87125 | 0.721774 | 0.485977 | 8.00E-272 | royalblue | ARRB2 |
| 4.42E-272 | -0.76989 | 0.224078 | 0.688806 | 9.38E-272 | royalblue | SNHG8 |
| 4.62E-271 | 0.819289 | 0.474078 | 0.181837 | 9.74E-271 | royalblue | ZYX |
| 8.35E-271 | 0.816053 | 0.743664 | 0.527517 | 1.76E-270 | royalblue | AP2S1 |
| 3.64E-269 | 0.755368 | 0.362903 | 0.100665 | 7.62E-269 | royalblue | CDKN1A |
| 4.83E-262 | -0.94005 | 0.322005 | 0.741796 | 1.01E-261 | royalblue | CALR |
| 8.13E-260 | 0.587254 | 0.259217 | 0.05558 | 1.69E-259 | royalblue | FCGR2A |
| 1.10E-259 | -0.65912 | 0.245968 | 0.72687 | 2.27E-259 | royalblue | SOD1 |
| 3.94E-259 | -1.20841 | 0.167051 | 0.620651 | 8.10E-259 | royalblue | HIST1H4C |
| 4.91E-259 | -0.66773 | 0.008065 | 0.451457 | 1.01E-258 | royalblue | OCIAD2 |
| 8.03E-258 | 0.743508 | 0.318548 | 0.08533 | 1.64E-257 | royalblue | NLRP3 |
| 2.18E-257 | 1.001056 | 0.6947 | 0.413222 | 4.44E-257 | royalblue | FOSB |
| 1.42E-256 | -1.29902 | 0.236175 | 0.647913 | 2.88E-256 | royalblue | TUBB |
| 3.25E-255 | 0.619931 | 0.263825 | 0.057591 | 6.56E-255 | royalblue | MS4A7 |
| 1.34E-253 | -0.7226 | 0.269585 | 0.732118 | 2.70E-253 | royalblue | SRSF7 |
| 2.34E-253 | 0.870586 | 0.669931 | 0.436429 | 4.68E-253 | royalblue | PGD |
| 7.24E-253 | 0.919537 | 0.560484 | 0.277287 | 1.44E-252 | royalblue | ATP2B1 |
| 7.27E-253 | 0.839007 | 0.764977 | 0.647061 | 1.44E-252 | royalblue | ALDOA |

| | | | | | | |
|---|---|---|---|---|---|---|
| 2.52E-250 | 0.68478 | 0.831797 | 0.632987 | 4.99E-250 | royalblue | ITGB2 |
| 7.49E-250 | -0.81311 | 0.006336 | 0.43694 | 1.47E-249 | royalblue | RHOH |
| 2.29E-248 | -0.70914 | 0.456221 | 0.804771 | 4.49E-248 | royalblue | EIF3E |
| 4.08E-248 | 0.527819 | 0.972926 | 0.981053 | 7.98E-248 | royalblue | UBC |
| 1.31E-246 | -0.56697 | 0.5697 | 0.912114 | 2.55E-246 | royalblue | HNRNPDL |
| 1.63E-246 | 0.703674 | 0.816244 | 0.684205 | 3.15E-246 | royalblue | GNG5 |
| 3.75E-246 | -1.12811 | 0.202189 | 0.628318 | 7.24E-246 | royalblue | CXCR4 |
| 1.58E-245 | -0.65054 | 0.266705 | 0.712251 | 3.04E-245 | royalblue | PTGES3 |
| 3.79E-245 | 0.714868 | 0.391129 | 0.128676 | 7.25E-245 | royalblue | EVI2B |
| 1.27E-244 | -0.98257 | 0.088134 | 0.51266 | 2.42E-244 | royalblue | IMPDH2 |
| 2.79E-244 | 0.854411 | 0.583525 | 0.300971 | 5.30E-244 | royalblue | CAPG |
| 2.59E-242 | -0.74399 | 0.005184 | 0.427364 | 4.90E-242 | royalblue | ITM2A |
| 5.98E-240 | -1.87613 | 0.009793 | 0.424502 | 1.13E-239 | royalblue | CD3E |
| 1.66E-239 | 0.767285 | 0.364055 | 0.11116 | 3.11E-239 | royalblue | RHOB |
| 4.04E-239 | -1.4397 | 0.016129 | 0.433191 | 7.55E-239 | royalblue | CD7 |
| 5.82E-237 | 0.723272 | 0.386521 | 0.132936 | 1.09E-236 | royalblue | AP1S2 |
| 6.62E-234 | -0.6751 | 0.108871 | 0.562038 | 1.23E-233 | royalblue | ACAP1 |
| 5.02E-232 | 0.549108 | 0.297235 | 0.075175 | 9.29E-232 | royalblue | SH3BP5 |
| 7.75E-232 | 0.587092 | 0.921659 | 0.726461 | 1.43E-231 | royalblue | GSTP1 |
| 1.85E-231 | 0.873335 | 0.585253 | 0.290407 | 3.39E-231 | royalblue | ID2 |
| 4.71E-230 | 0.847167 | 0.717166 | 0.471358 | 8.62E-230 | royalblue | GLUL |
| 8.07E-230 | -0.88577 | 0.016129 | 0.423513 | 1.47E-229 | royalblue | CDK6 |
| 1.86E-229 | 0.777682 | 0.748848 | 0.525439 | 3.38E-229 | royalblue | FLNA |
| 6.44E-228 | -0.58915 | 0.13076 | 0.582109 | 1.17E-227 | royalblue | TECR |
| 6.98E-223 | -0.62962 | 0.081797 | 0.512626 | 1.26E-222 | royalblue | NOP58 |
| 2.44E-222 | -0.61742 | 0.027074 | 0.435679 | 4.39E-222 | royalblue | ZNF22 |
| 3.96E-222 | 0.841937 | 0.452189 | 0.191685 | 7.09E-222 | royalblue | CPVL |
| 3.05E-220 | -2.09945 | 0.019009 | 0.410019 | 5.44E-220 | royalblue | IL32 |
| 6.18E-220 | 0.701936 | 0.80818 | 0.675822 | 1.10E-219 | royalblue | PKM |
| 4.33E-218 | -0.61109 | 0.322005 | 0.732391 | 7.68E-218 | royalblue | ST13 |
| 7.49E-217 | 0.849062 | 0.637673 | 0.42215 | 1.32E-216 | royalblue | RILPL2 |
| 1.44E-216 | 0.782875 | 0.661866 | 0.406168 | 2.53E-216 | royalblue | IQGAP1 |
| 7.80E-215 | -1.58242 | 0.031682 | 0.418061 | 1.37E-214 | royalblue | LTB |
| 8.47E-215 | -0.70861 | 0.368088 | 0.73481 | 1.48E-214 | royalblue | RAN |
| 8.67E-215 | -0.6823 | 0.126728 | 0.546328 | 1.51E-214 | royalblue | HSPB1 |
| 5.99E-213 | 0.709889 | 0.345622 | 0.119169 | 1.04E-212 | royalblue | BST1 |
| 2.76E-212 | -0.58257 | 0.388825 | 0.796524 | 4.77E-212 | royalblue | EIF4B |
| 9.37E-212 | -0.60904 | 0.147465 | 0.565173 | 1.61E-211 | royalblue | CCT7 |
| 2.59E-211 | -0.63045 | 0.065668 | 0.476231 | 4.44E-211 | royalblue | PARP1 |
| 7.14E-210 | -0.67517 | 0.124424 | 0.534674 | 1.22E-209 | royalblue | CCT2 |
| 1.47E-208 | -0.58864 | 0.365207 | 0.763674 | 2.52E-208 | royalblue | HNRNPA0 |

| | | | | | | |
|---|---|---|---|---|---|---|
| 2.57E-208 | 0.653507 | 0.259217 | 0.06594 | 4.37E-208 | royalblue | RGCC |
| 2.46E-207 | -0.81527 | 0.16129 | 0.551951 | 4.16E-207 | royalblue | HSPD1 |
| 1.45E-205 | -0.58646 | 0.059908 | 0.46519 | 2.44E-205 | royalblue | ARHGAP15 |
| 1.76E-205 | -0.58304 | 0.285714 | 0.719918 | 2.97E-205 | royalblue | EIF4A2 |
| 9.09E-205 | -0.68497 | 0.209677 | 0.597649 | 1.52E-204 | royalblue | APEX1 |
| 3.16E-204 | -0.58735 | 0.108295 | 0.516238 | 5.28E-204 | royalblue | CCT3 |
| 2.54E-199 | 0.592381 | 0.833525 | 0.767047 | 4.22E-199 | royalblue | SERP1 |
| 1.80E-197 | 0.672382 | 0.406682 | 0.161663 | 2.99E-197 | royalblue | SCPEP1 |
| 3.03E-197 | -0.88013 | 0.013825 | 0.380031 | 5.02E-197 | royalblue | ITM2C |
| 9.47E-197 | -0.5669 | 0.099654 | 0.501005 | 1.56E-196 | royalblue | PRMT1 |
| 8.15E-196 | -0.58091 | 0.184908 | 0.595093 | 1.34E-195 | royalblue | RBMX |
| 1.18E-195 | -0.59492 | 0.198733 | 0.598296 | 1.94E-195 | royalblue | SRP9 |
| 1.21E-195 | -0.6275 | 0.531682 | 0.875072 | 1.97E-195 | royalblue | RAC2 |
| 2.99E-195 | -0.53735 | 0.019585 | 0.390867 | 4.87E-195 | royalblue | BEX3 |
| 1.58E-192 | 0.706913 | 0.359447 | 0.135935 | 2.57E-192 | royalblue | ALDH2 |
| 1.63E-190 | -0.50753 | 0.687212 | 0.92796 | 2.63E-190 | royalblue | HNRNPA2B1 |
| 6.89E-190 | 0.659071 | 0.845046 | 0.678037 | 1.11E-189 | royalblue | ANXA1 |
| 4.07E-189 | 0.733292 | 0.543779 | 0.285023 | 6.54E-189 | royalblue | TCIRG1 |
| 8.70E-189 | 0.806088 | 0.553571 | 0.306696 | 1.39E-188 | royalblue | CARD16 |
| 1.30E-187 | 0.656856 | 0.28629 | 0.088635 | 2.07E-187 | royalblue | PPIF |
| 3.21E-187 | -0.60696 | 0.1947 | 0.595093 | 5.11E-187 | royalblue | NUCKS1 |
| 1.12E-186 | -1.20036 | 0.007488 | 0.355359 | 1.78E-186 | royalblue | PRSS57 |
| 4.61E-185 | -0.66008 | 0.005184 | 0.352701 | 7.29E-185 | royalblue | SLC38A1 |
| 6.04E-184 | -0.69526 | 0.084101 | 0.461067 | 9.52E-184 | royalblue | 1-Sep |
| 7.39E-184 | -0.65938 | 0.174539 | 0.573113 | 1.16E-183 | royalblue | PRDX2 |
| 6.22E-183 | 0.699277 | 0.387673 | 0.157028 | 9.75E-183 | royalblue | MYO9B |
| 1.20E-182 | -0.55411 | 0.705069 | 0.944011 | 1.87E-182 | royalblue | RPS29 |
| 1.95E-182 | -0.57852 | 0.138825 | 0.541796 | 3.04E-182 | royalblue | FOXP1 |
| 3.12E-182 | -0.57322 | 0.198733 | 0.591992 | 4.84E-182 | royalblue | YWHAQ |
| 7.49E-182 | -0.51316 | 0.115207 | 0.512183 | 1.16E-181 | royalblue | TAF7 |
| 4.58E-180 | -0.51688 | 0.072005 | 0.44563 | 7.07E-180 | royalblue | CCNG1 |
| 8.58E-180 | -0.60389 | 0.150922 | 0.545578 | 1.32E-179 | royalblue | ITGA4 |
| 8.86E-180 | 0.716806 | 0.414747 | 0.177884 | 1.36E-179 | royalblue | MIDN |
| 5.09E-179 | -0.54885 | 0.406106 | 0.786539 | 7.77E-179 | royalblue | PNISR |
| 5.40E-179 | -0.52941 | 0.265553 | 0.684784 | 8.23E-179 | royalblue | 7-Sep |
| 1.65E-176 | 0.694275 | 0.428571 | 0.191344 | 2.50E-176 | royalblue | TOM1 |
| 1.65E-176 | -1.14348 | 0.004608 | 0.338047 | 2.50E-176 | royalblue | CD3D |
| 2.72E-176 | -0.52724 | 0.520737 | 0.814585 | 4.11E-176 | royalblue | EIF3L |
| 7.93E-176 | -0.99848 | 0.005184 | 0.33832 | 1.19E-175 | royalblue | LCK |
| 2.05E-175 | -0.57971 | 0.03629 | 0.38596 | 3.08E-175 | royalblue | CDK4 |
| 3.39E-175 | -0.69166 | 0.086406 | 0.44839 | 5.07E-175 | royalblue | TBC1D10C |

| | | | | | | |
|---|---|---|---|---|---|---|
| 4.04E-175 | 0.62969 | 0.843894 | 0.705503 | 6.03E-175 | royalblue | IER2 |
| 8.30E-174 | -0.58998 | 0.127304 | 0.510854 | 1.23E-173 | royalblue | PRMT2 |
| 1.26E-173 | -0.50736 | 0.077765 | 0.451184 | 1.87E-173 | royalblue | BUB3 |
| 1.57E-173 | -0.81917 | 0.044355 | 0.386505 | 2.31E-173 | royalblue | AC084033.3 |
| 9.46E-173 | -0.67385 | 0.18606 | 0.576487 | 1.39E-172 | royalblue | LEPROTL1 |
| 3.07E-172 | -0.52951 | 0.308756 | 0.679025 | 4.51E-172 | royalblue | PHB2 |
| 8.72E-171 | -0.60947 | 0.056452 | 0.409712 | 1.28E-170 | royalblue | CYFIP2 |
| 3.38E-170 | 0.654366 | 0.746544 | 0.665599 | 4.93E-170 | royalblue | ATP6V0C |
| 5.39E-170 | 0.774729 | 0.437788 | 0.21404 | 7.85E-170 | royalblue | NUP214 |
| 2.36E-169 | -0.56006 | 0.167051 | 0.539853 | 3.42E-169 | royalblue | RBBP4 |
| 1.85E-167 | -0.72112 | 0.198733 | 0.558085 | 2.68E-167 | royalblue | DUT |
| 2.49E-167 | 0.670898 | 0.456221 | 0.217175 | 3.60E-167 | royalblue | GLIPR2 |
| 2.11E-166 | -0.66358 | 0.148041 | 0.530346 | 3.03E-166 | royalblue | NOSIP |
| 4.21E-166 | -0.53671 | 0.303571 | 0.666383 | 6.04E-166 | royalblue | HMGN1 |
| 1.30E-165 | -0.9269 | 0.003456 | 0.321281 | 1.86E-165 | royalblue | SPOCK2 |
| 1.76E-164 | 0.525494 | 0.272465 | 0.084035 | 2.51E-164 | royalblue | HOTAIRM1 |
| 1.78E-164 | -0.58216 | 0.047811 | 0.389095 | 2.54E-164 | royalblue | PBXIP1 |
| 3.43E-164 | -0.56529 | 0.582373 | 0.841404 | 4.86E-164 | royalblue | MIF |
| 6.67E-164 | 0.672908 | 0.389401 | 0.17267 | 9.43E-164 | royalblue | PRAM1 |
| 1.11E-163 | 0.592758 | 0.305876 | 0.104617 | 1.57E-163 | royalblue | METRNL |
| 1.71E-163 | -0.52934 | 0.327189 | 0.677356 | 2.41E-163 | royalblue | SERBP1 |
| 5.56E-163 | 0.619446 | 0.782834 | 0.738354 | 7.80E-163 | royalblue | ITM2B |
| 1.60E-162 | -1.21872 | 0.400346 | 0.677765 | 2.24E-162 | royalblue | TUBA1B |
| 1.58E-161 | -0.5066 | 0.218318 | 0.586267 | 2.20E-161 | royalblue | CCT4 |
| 9.44E-161 | 0.715406 | 0.672235 | 0.441574 | 1.31E-160 | royalblue | AHNAK |
| 1.87E-160 | -0.52512 | 0.204493 | 0.577407 | 2.59E-160 | royalblue | HSPE1 |
| 3.08E-160 | -0.6088 | 0.1803 | 0.547725 | 4.26E-160 | royalblue | CCND3 |
| 2.65E-159 | 0.702656 | 0.586982 | 0.369399 | 3.66E-159 | royalblue | C1orf162 |
| 3.07E-159 | 0.554338 | 0.794355 | 0.755393 | 4.22E-159 | royalblue | HIGD2A |
| 3.44E-159 | -0.60194 | 0.009217 | 0.323462 | 4.72E-159 | royalblue | EGFL7 |
| 5.21E-159 | 0.677804 | 0.563364 | 0.351474 | 7.12E-159 | royalblue | RNF130 |
| 1.14E-157 | 0.681519 | 0.413594 | 0.19281 | 1.55E-157 | royalblue | UBXN11 |
| 1.72E-157 | -0.85452 | 0.004032 | 0.31116 | 2.34E-157 | royalblue | FCMR |
| 2.69E-157 | 0.628689 | 0.383641 | 0.166672 | 3.64E-157 | royalblue | SAMSN1 |
| 2.21E-156 | -0.60229 | 0.09447 | 0.434486 | 2.99E-156 | royalblue | NASP |
| 2.05E-155 | 0.546566 | 0.790323 | 0.723564 | 2.77E-155 | royalblue | COX8A |
| 1.98E-154 | 0.717442 | 0.386521 | 0.171784 | 2.66E-154 | royalblue | RAB11FIP1 |
| 5.12E-154 | -0.50224 | 0.267857 | 0.626887 | 6.88E-154 | royalblue | FBL |
| 6.29E-154 | 0.657697 | 0.368088 | 0.158392 | 8.43E-154 | royalblue | GNB2 |
| 1.02E-153 | -1.43625 | 0.00576 | 0.306901 | 1.36E-153 | royalblue | IL7R |
| 1.06E-153 | -0.57816 | 0.420507 | 0.731913 | 1.41E-153 | royalblue | LDHA |

| | | | | | | |
|---|---|---|---|---|---|---|
| 1.62E-153 | -0.51623 | 0.221774 | 0.579008 | 2.15E-153 | royalblue | HNRNPF |
| 1.14E-151 | 0.550544 | 0.254032 | 0.08124 | 1.51E-151 | royalblue | CARS2 |
| 1.28E-151 | -0.50566 | 0.103687 | 0.44839 | 1.69E-151 | royalblue | SPN |
| 3.17E-151 | -0.62614 | 0.002304 | 0.299267 | 4.18E-151 | royalblue | FAM30A |
| 6.59E-150 | 0.645284 | 0.677995 | 0.537536 | 8.66E-150 | royalblue | PLP2 |
| 1.71E-149 | 0.568014 | 0.264977 | 0.09068 | 2.25E-149 | royalblue | CD302 |
| 5.97E-149 | 0.518101 | 0.309908 | 0.112523 | 7.80E-149 | royalblue | SH3BP2 |
| 2.71E-148 | -0.63001 | 0.00288 | 0.295621 | 3.53E-148 | royalblue | CD34 |
| 1.80E-146 | 0.785885 | 0.505184 | 0.307753 | 2.34E-146 | royalblue | SOD2 |
| 1.90E-146 | -0.78911 | 0.00288 | 0.292145 | 2.47E-146 | royalblue | ZAP70 |
| 1.05E-145 | -0.57231 | 0.042051 | 0.3496 | 1.36E-145 | royalblue | BIN1 |
| 1.67E-145 | 0.555485 | 0.779954 | 0.736991 | 2.15E-145 | royalblue | UQCR11 |
| 1.12E-144 | -0.5937 | 0.195276 | 0.551746 | 1.44E-144 | royalblue | RARRES3 |
| 1.51E-144 | -0.58292 | 0.029954 | 0.333549 | 1.93E-144 | royalblue | RNF125 |
| 6.51E-144 | 0.713903 | 0.505184 | 0.302777 | 8.34E-144 | royalblue | SAT2 |
| 7.64E-144 | 0.727068 | 0.516705 | 0.319748 | 9.76E-144 | royalblue | GRK2 |
| 3.31E-141 | 0.713515 | 0.525922 | 0.331027 | 4.21E-141 | royalblue | PTP4A2 |
| 3.87E-141 | -0.58312 | 0.038594 | 0.33999 | 4.92E-141 | royalblue | MCM7 |
| 1.92E-140 | 0.650249 | 0.346774 | 0.154132 | 2.44E-140 | royalblue | TNFSF13 |
| 5.53E-139 | -0.77793 | 0 | 0.276367 | 6.99E-139 | royalblue | HOPX |
| 5.12E-138 | -0.76432 | 0.000576 | 0.275652 | 6.46E-138 | royalblue | SMIM24 |
| 7.12E-138 | -0.67487 | 0.22523 | 0.575464 | 8.96E-138 | royalblue | DNAJB1 |
| 1.60E-137 | -0.69818 | 0.004608 | 0.281274 | 2.01E-137 | royalblue | LBH |
| 2.99E-137 | -0.77675 | 0.024194 | 0.307207 | 3.74E-137 | royalblue | PIK3IP1 |
| 8.16E-136 | -0.58925 | 0.085253 | 0.394616 | 1.02E-135 | royalblue | P2RY8 |
| 8.30E-136 | 0.535067 | 0.315092 | 0.12261 | 1.03E-135 | royalblue | EFHD2 |
| 1.96E-135 | 0.69802 | 0.512097 | 0.318589 | 2.44E-135 | royalblue | ARPC1B |
| 4.04E-135 | 0.70652 | 0.493664 | 0.304515 | 5.00E-135 | royalblue | ATP6V0D1 |
| 3.11E-134 | 0.640782 | 0.362327 | 0.165139 | 3.84E-134 | royalblue | GPCPD1 |
| 7.34E-134 | 0.637757 | 0.387673 | 0.191208 | 9.05E-134 | royalblue | CAPNS1 |
| 1.59E-133 | 0.60926 | 0.661866 | 0.516476 | 1.95E-133 | royalblue | CYBA |
| 1.88E-133 | 0.546317 | 0.395161 | 0.176657 | 2.30E-133 | royalblue | GABARAPL1 |
| 1.19E-132 | 0.624715 | 0.332373 | 0.145885 | 1.46E-132 | royalblue | PLXDC2 |
| 1.52E-132 | -1.48547 | 0.033986 | 0.312285 | 1.86E-132 | royalblue | CST7 |
| 1.76E-132 | -0.61215 | 0.002304 | 0.270336 | 2.15E-132 | royalblue | ABLIM1 |
| 2.74E-132 | 0.599512 | 0.717742 | 0.64931 | 3.33E-132 | royalblue | CD44 |
| 1.02E-131 | 0.598274 | 0.866935 | 0.771682 | 1.23E-131 | royalblue | JUN |
| 1.78E-131 | -1.40888 | 0.004032 | 0.270472 | 2.16E-131 | royalblue | IGLL1 |
| 2.46E-131 | 0.642693 | 0.614631 | 0.461782 | 2.96E-131 | royalblue | CD63 |
| 4.01E-131 | -0.93756 | 0.001728 | 0.266144 | 4.83E-131 | royalblue | SPINK2 |
| 4.83E-131 | -0.89094 | 0.002304 | 0.267337 | 5.79E-131 | royalblue | GZMM |

| | | | | | | |
|---|---|---|---|---|---|---|
| 2.00E-130 | -0.62303 | 0.012673 | 0.283081 | 2.40E-130 | royalblue | PRKCH |
| 3.88E-129 | -0.79561 | 0.003456 | 0.266485 | 4.63E-129 | royalblue | CD3G |
| 3.38E-128 | -0.63045 | 0.803571 | 0.926972 | 4.02E-128 | royalblue | HMGB1 |
| 2.03E-127 | 0.674182 | 0.384793 | 0.198024 | 2.41E-127 | royalblue | MXD1 |
| 8.69E-127 | -0.61608 | 0.000576 | 0.257625 | 1.03E-126 | royalblue | SKAP1 |
| 1.58E-126 | 0.722124 | 0.440092 | 0.259772 | 1.87E-126 | royalblue | PLSCR1 |
| 2.87E-125 | 0.562942 | 0.341014 | 0.148339 | 3.38E-125 | royalblue | ADA2 |
| 4.10E-125 | -0.50518 | 0.319124 | 0.659499 | 4.83E-125 | royalblue | CNN2 |
| 7.68E-125 | 0.704989 | 0.402074 | 0.216255 | 9.03E-125 | royalblue | SLC25A37 |
| 1.38E-124 | -0.56217 | 0.096198 | 0.387425 | 1.61E-124 | royalblue | FABP5 |
| 1.27E-123 | 0.544243 | 0.256336 | 0.094156 | 1.49E-123 | royalblue | CBX6 |
| 8.03E-123 | -0.86246 | 0.002304 | 0.253842 | 9.36E-123 | royalblue | CD247 |
| 1.41E-122 | -0.61535 | 0.020161 | 0.281343 | 1.64E-122 | royalblue | LAT |
| 2.27E-122 | -0.746 | 0.574885 | 0.832987 | 2.63E-122 | royalblue | TSC22D3 |
| 1.10E-119 | 0.658367 | 0.354839 | 0.177543 | 1.27E-119 | royalblue | METTL9 |
| 2.34E-119 | 0.518346 | 0.273618 | 0.107276 | 2.70E-119 | royalblue | C9orf72 |
| 2.13E-116 | -0.50916 | 0.375 | 0.719339 | 2.46E-116 | royalblue | UCP2 |
| 2.40E-115 | 0.636549 | 0.633065 | 0.531539 | 2.76E-115 | royalblue | PGLS |
| 7.16E-115 | 0.603143 | 0.65265 | 0.589981 | 8.21E-115 | royalblue | ATP6V1F |
| 1.10E-114 | 0.665804 | 0.371544 | 0.194514 | 1.26E-114 | royalblue | RAC1 |
| 7.16E-114 | 0.63248 | 0.31682 | 0.148816 | 8.17E-114 | royalblue | SULT1A1 |
| 7.62E-114 | 0.58198 | 0.30818 | 0.136343 | 8.68E-114 | royalblue | IRS2 |
| 8.06E-113 | 0.516897 | 0.27765 | 0.112421 | 9.16E-113 | royalblue | IL17RA |
| 4.03E-112 | -0.62928 | 0.082949 | 0.34517 | 4.57E-112 | royalblue | ISG20 |
| 5.22E-112 | 0.586529 | 0.46659 | 0.278753 | 5.91E-112 | royalblue | CAST |
| 5.79E-112 | 0.660637 | 0.400346 | 0.223309 | 6.54E-112 | royalblue | ZFAND5 |
| 2.15E-111 | -0.64369 | 0.025346 | 0.269825 | 2.42E-111 | royalblue | FLT3LG |
| 3.77E-108 | 0.519949 | 0.709677 | 0.657591 | 4.24E-108 | royalblue | ATP5MPL |
| 9.15E-108 | -0.53418 | 0.194124 | 0.471562 | 1.03E-107 | royalblue | CAT |
| 3.75E-107 | -0.55565 | 0.018433 | 0.252718 | 4.20E-107 | royalblue | CDC25B |
| 4.69E-106 | 0.578437 | 0.320276 | 0.153689 | 5.23E-106 | royalblue | LYN |
| 8.58E-106 | 0.566239 | 0.645737 | 0.554234 | 9.56E-106 | royalblue | CNPY3 |
| 1.13E-105 | 0.668342 | 0.391705 | 0.225183 | 1.26E-105 | royalblue | AMPD2 |
| 4.04E-105 | 0.528489 | 0.718894 | 0.692554 | 4.48E-105 | royalblue | NDUFA1 |
| 3.02E-104 | 0.544362 | 0.285714 | 0.125848 | 3.35E-104 | royalblue | MAP2K3 |
| 5.90E-96 | 0.549362 | 0.654378 | 0.605486 | 6.52E-96 | royalblue | RHOG |
| 3.53E-95 | 0.519699 | 0.304724 | 0.148714 | 3.89E-95 | royalblue | CHP1 |
| 1.25E-94 | 0.52563 | 0.27765 | 0.129324 | 1.37E-94 | royalblue | CD33 |
| 1.98E-94 | -2.11585 | 0.087558 | 0.318283 | 2.17E-94 | royalblue | NKG7 |
| 9.51E-94 | 0.619444 | 0.565668 | 0.455989 | 1.04E-93 | royalblue | GRB2 |
| 2.77E-92 | 0.607512 | 0.343894 | 0.182859 | 3.02E-92 | royalblue | ANKRD13D |

| | | | | | | |
|---|---|---|---|---|---|---|
| 1.45E-91 | 0.50023 | 0.270161 | 0.120327 | 1.58E-91 | royalblue | ARAP1 |
| 1.85E-91 | 0.503269 | 0.271889 | 0.12227 | 2.01E-91 | royalblue | PFKFB3 |
| 5.69E-91 | -0.52756 | 0.052419 | 0.282092 | 6.17E-91 | royalblue | CMC1 |
| 1.82E-89 | 0.656839 | 0.437212 | 0.289794 | 1.97E-89 | royalblue | IVNS1ABP |
| 1.73E-88 | 0.596414 | 0.272465 | 0.128472 | 1.86E-88 | royalblue | RFLNB |
| 3.24E-88 | 0.585207 | 0.326037 | 0.176146 | 3.49E-88 | royalblue | QSOX1 |
| 8.58E-88 | 0.530608 | 0.641129 | 0.580814 | 9.23E-88 | royalblue | LAMTOR1 |
| 2.28E-84 | 0.524563 | 0.50288 | 0.336787 | 2.45E-84 | royalblue | PLEK |
| 2.20E-83 | 0.549038 | 0.285138 | 0.144488 | 2.36E-83 | royalblue | CRTAP |
| 2.27E-83 | 0.528269 | 0.660714 | 0.622321 | 2.43E-83 | royalblue | TCF25 |
| 1.35E-81 | 0.534704 | 0.484447 | 0.333004 | 1.44E-81 | royalblue | GLIPR1 |
| 1.29E-80 | -0.57009 | 0.18318 | 0.425558 | 1.37E-80 | royalblue | DDIT4 |
| 8.51E-80 | 0.559202 | 0.583525 | 0.499642 | 9.04E-80 | royalblue | LRRFIP1 |
| 1.71E-79 | 0.590483 | 0.527074 | 0.413904 | 1.82E-79 | royalblue | CD55 |
| 3.81E-79 | 0.508452 | 0.376152 | 0.220412 | 4.02E-79 | royalblue | IFNGR1 |
| 5.19E-77 | 0.647634 | 0.446429 | 0.322508 | 5.48E-77 | royalblue | CTSA |
| 1.34E-76 | 0.559416 | 0.549539 | 0.43009 | 1.41E-76 | royalblue | MYADM |
| 2.81E-76 | -0.56456 | 0.700461 | 0.865258 | 2.96E-76 | royalblue | HMGN2 |
| 5.52E-75 | 0.568509 | 0.357143 | 0.21939 | 5.79E-75 | royalblue | UBR4 |
| 2.50E-74 | 0.58741 | 0.457949 | 0.34016 | 2.61E-74 | royalblue | PCBP1 |
| 4.56E-74 | 0.516921 | 0.285714 | 0.149225 | 4.76E-74 | royalblue | MAP4K4 |
| 1.16E-71 | -0.83904 | 0.131912 | 0.341898 | 1.21E-71 | royalblue | AREG |
| 8.92E-69 | 0.529606 | 0.323733 | 0.192844 | 9.28E-69 | royalblue | CARD19 |
| 1.09E-67 | 0.541276 | 0.360023 | 0.228148 | 1.14E-67 | royalblue | PNPLA2 |
| 1.33E-65 | 0.576995 | 0.357143 | 0.236906 | 1.38E-65 | royalblue | TBXAS1 |
| 1.41E-64 | 0.86934 | 0.273041 | 0.151303 | 1.46E-64 | royalblue | IL1B |
| 4.65E-64 | 0.510797 | 0.325461 | 0.198671 | 4.79E-64 | royalblue | PRELID1 |
| 2.13E-63 | 0.527507 | 0.522465 | 0.436838 | 2.19E-63 | royalblue | RNH1 |
| 4.20E-63 | -0.58228 | 0.227535 | 0.433157 | 4.31E-63 | royalblue | TNFAIP3 |
| 1.16E-59 | 0.653626 | 0.288594 | 0.163537 | 1.19E-59 | royalblue | AC020916.1 |
| 1.59E-59 | 0.620798 | 0.384217 | 0.267269 | 1.63E-59 | royalblue | NFKBIZ |
| 1.51E-51 | 0.533635 | 0.495968 | 0.43452 | 1.54E-51 | royalblue | GSTO1 |
| 4.78E-47 | -0.54922 | 0.464862 | 0.62365 | 4.87E-47 | royalblue | H2AFZ |
| 5.20E-47 | 0.518152 | 0.415323 | 0.318487 | 5.29E-47 | royalblue | PER1 |
| 7.20E-46 | 0.51985 | 0.381912 | 0.289249 | 7.30E-46 | royalblue | POLE4 |
| 5.81E-44 | 0.508348 | 0.360023 | 0.269143 | 5.88E-44 | royalblue | SRA1 |
| 6.21E-43 | 0.534252 | 0.379032 | 0.296712 | 6.27E-43 | royalblue | CTNNB1 |
| 2.18E-41 | 0.516553 | 0.427419 | 0.365071 | 2.20E-41 | royalblue | C20orf27 |
| 3.57E-34 | 0.513768 | 0.279378 | 0.200954 | 3.59E-34 | royalblue | GCA |
| 1.11E-31 | 0.502789 | 0.41129 | 0.359891 | 1.11E-31 | royalblue | H2AFJ |
| 1.56E-27 | 0.571081 | 0.279954 | 0.203851 | 1.56E-27 | royalblue | EREG |

| | | | | | | |
|---|---|---|---|---|---|---|
| 2.13E-09 | -1.09095 | 0.273618 | 0.210939 | 2.13E-09 | royalblue | HBB |
| 0 | 2.087709 | 0.987138 | 0.235924 | 0 | black | LYZ |
| 0 | 1.74077 | 0.966238 | 0.396566 | 0 | black | CST3 |
| 0 | 1.702069 | 0.961415 | 0.36774 | 0 | black | LGALS1 |
| 0 | 1.126773 | 0.728296 | 0.141108 | 0 | black | S100A8 |
| 0 | 1.058361 | 0.810289 | 0.135034 | 0 | black | CSTA |
| 0 | 1.049571 | 0.631833 | 0.083128 | 0 | black | MNDA |
| 0 | 0.910902 | 0.326367 | 0.006369 | 0 | black | CLEC10A |
| 0 | 0.908476 | 0.493569 | 0.051906 | 0 | black | AC020656.1 |
| 0 | 0.853644 | 0.625402 | 0.074395 | 0 | black | IGSF6 |
| 0 | 0.83892 | 0.308682 | 0.01067 | 0 | black | CD1C |
| 0 | 0.828777 | 0.453376 | 0.056568 | 0 | black | NAPSA |
| 0 | 0.820437 | 0.77492 | 0.14357 | 0 | black | S100A9 |
| 0 | 0.789379 | 0.663987 | 0.126859 | 0 | black | CTSH |
| 0 | 0.781603 | 0.689711 | 0.127056 | 0 | black | LY86 |
| 0 | 0.712375 | 0.413183 | 0.039923 | 0 | black | CES1 |
| 0 | 0.528875 | 0.461415 | 0.054565 | 0 | black | CD86 |
| 7.42E-300 | 0.512653 | 0.472669 | 0.065465 | 8.38E-299 | black | IFI30 |
| 1.08E-299 | 0.735088 | 0.792605 | 0.183952 | 1.16E-298 | black | COTL1 |
| 5.92E-298 | 1.230486 | 0.601286 | 0.122066 | 5.98E-297 | black | RNASE2 |
| 2.06E-291 | 1.2452 | 0.371383 | 0.045307 | 1.98E-290 | black | RETN |
| 6.18E-291 | 1.377618 | 0.836013 | 0.289832 | 5.65E-290 | black | ANXA2 |
| 4.54E-288 | 0.926222 | 0.885852 | 0.244722 | 3.96E-287 | black | FCER1G |
| 8.67E-285 | 0.861163 | 0.681672 | 0.150957 | 7.24E-284 | black | CFP |
| 2.32E-265 | 0.594284 | 0.475884 | 0.075085 | 1.86E-264 | black | FGL2 |
| 2.96E-265 | 0.602983 | 0.670418 | 0.139532 | 2.27E-264 | black | TYMP |
| 1.16E-261 | 0.613147 | 0.567524 | 0.110181 | 8.56E-261 | black | RNASE6 |
| 6.80E-255 | 1.080537 | 0.874598 | 0.358515 | 4.84E-254 | black | GRN |
| 1.02E-242 | 0.741094 | 0.956592 | 0.31242 | 7.02E-242 | black | TYROBP |
| 1.54E-240 | 0.748418 | 0.643087 | 0.163531 | 1.02E-239 | black | PLD4 |
| 5.00E-240 | 0.583475 | 0.323151 | 0.039726 | 3.20E-239 | black | LGALS2 |
| 6.80E-233 | 0.537341 | 0.456592 | 0.078007 | 4.21E-232 | black | CEBPD |
| 1.75E-229 | 1.375075 | 0.511254 | 0.114613 | 1.05E-228 | black | FCER1A |
| 5.08E-223 | 0.728594 | 1 | 0.99872 | 2.96E-222 | black | TMSB10 |
| 3.72E-219 | 1.218702 | 0.948553 | 0.601005 | 2.10E-218 | black | S100A10 |
| 3.07E-216 | 0.516147 | 0.498392 | 0.097278 | 1.68E-215 | black | KLF4 |
| 2.22E-214 | 0.778004 | 0.889068 | 0.370071 | 1.19E-213 | black | LST1 |
| 1.70E-207 | 0.823568 | 0.692926 | 0.197774 | 8.84E-207 | black | PLAUR |
| 2.74E-207 | 0.735793 | 0.881029 | 0.335927 | 1.38E-206 | black | S100A11 |
| 4.34E-204 | 0.947279 | 0.776527 | 0.307364 | 2.14E-203 | black | CAPG |
| 1.48E-201 | 0.544747 | 0.525723 | 0.105322 | 7.09E-201 | black | VCAN |

| | | | | | | |
|---|---|---|---|---|---|---|
| 4.93E-199 | -0.68689 | 0.998392 | 0.999639 | 2.31E-198 | black | EEF1A1 |
| 6.64E-198 | 0.90204 | 0.990354 | 0.820381 | 3.03E-197 | black | S100A6 |
| 7.30E-197 | 1.085861 | 0.773312 | 0.304442 | 3.26E-196 | black | SAMHD1 |
| 4.89E-196 | 0.582829 | 0.57074 | 0.121573 | 2.13E-195 | black | FCN1 |
| 7.92E-196 | 1.616311 | 0.934084 | 0.63505 | 3.38E-195 | black | HLA-DRB1 |
| 6.59E-193 | 0.707075 | 0.707395 | 0.238649 | 2.75E-192 | black | SPI1 |
| 8.45E-188 | 0.899311 | 0.770096 | 0.317968 | 3.45E-187 | black | ANXA5 |
| 1.58E-180 | 0.602099 | 0.55627 | 0.156473 | 6.34E-180 | black | PAK1 |
| 8.31E-179 | 0.816616 | 0.898714 | 0.532256 | 3.26E-178 | black | AP2S1 |
| 1.11E-178 | 0.875651 | 0.440514 | 0.094159 | 4.25E-178 | black | MS4A6A |
| 2.61E-175 | 0.512992 | 0.459807 | 0.102564 | 9.82E-175 | black | STX11 |
| 1.06E-174 | 0.965915 | 0.980707 | 0.905775 | 3.92E-174 | black | VIM |
| 4.67E-172 | 0.84187 | 0.972669 | 0.732559 | 1.69E-171 | black | GSTP1 |
| 1.96E-170 | 0.631161 | 0.998392 | 0.981812 | 6.98E-170 | black | SERF2 |
| 3.63E-170 | 0.883582 | 0.985531 | 0.824321 | 1.27E-169 | black | SRGN |
| 4.92E-170 | 0.830365 | 0.983923 | 0.808201 | 1.69E-169 | black | S100A4 |
| 2.41E-169 | 0.539883 | 0.524116 | 0.140188 | 8.13E-169 | black | ZNF385A |
| 2.71E-169 | 1.233065 | 0.991961 | 0.917397 | 8.97E-169 | black | CD74 |
| 9.39E-169 | 1.574156 | 0.88746 | 0.575626 | 3.06E-168 | black | HLA-DRB5 |
| 1.71E-166 | 0.578129 | 0.514469 | 0.140944 | 5.46E-166 | black | ALDH2 |
| 1.61E-164 | 0.622561 | 0.525723 | 0.13707 | 5.07E-164 | black | JAML |
| 1.32E-160 | 1.5532 | 0.942122 | 0.595194 | 4.10E-160 | black | HLA-DRA |
| 5.98E-159 | 0.620167 | 0.583601 | 0.175613 | 1.82E-158 | black | RAB11FIP1 |
| 2.27E-158 | 0.746886 | 0.826367 | 0.401392 | 6.81E-158 | black | TIMP1 |
| 7.63E-154 | 0.621254 | 0.561093 | 0.176697 | 2.25E-153 | black | GSN |
| 5.47E-153 | -0.64788 | 0.974277 | 0.989034 | 1.59E-152 | black | RPL3 |
| 6.69E-153 | 0.645898 | 0.590032 | 0.186677 | 1.92E-152 | black | HLA-DMB |
| 4.76E-152 | 0.780252 | 0.768489 | 0.360386 | 1.34E-151 | black | C20orf27 |
| 9.30E-152 | 0.596273 | 0.985531 | 0.953052 | 2.59E-151 | black | MYL6 |
| 1.75E-151 | 0.818009 | 0.946945 | 0.680357 | 4.80E-151 | black | TSPO |
| 4.61E-151 | 1.58358 | 0.731511 | 0.360813 | 1.25E-150 | black | HLA-DQB1 |
| 2.41E-148 | 0.79235 | 0.863344 | 0.475721 | 6.42E-148 | black | PYCARD |
| 9.05E-148 | 0.744033 | 0.522508 | 0.147313 | 2.38E-147 | black | GPR183 |
| 6.14E-144 | 0.697059 | 0.73955 | 0.321219 | 1.59E-143 | black | FCGRT |
| 3.89E-141 | 0.546067 | 0.977492 | 0.920976 | 9.95E-141 | black | OAZ1 |
| 1.79E-139 | 0.944485 | 0.559486 | 0.199022 | 4.53E-139 | black | CPVL |
| 2.83E-139 | 0.623153 | 0.938907 | 0.80436 | 7.05E-139 | black | GABARAP |
| 4.17E-139 | 0.538496 | 0.586817 | 0.185561 | 1.03E-138 | black | STXBP2 |
| 7.28E-131 | 0.700748 | 0.948553 | 0.722775 | 1.77E-130 | black | COX8A |
| 1.41E-130 | 0.596375 | 0.932476 | 0.541745 | 3.37E-130 | black | AIF1 |
| 4.45E-130 | 1.042109 | 0.78135 | 0.493155 | 1.05E-129 | black | HLA-DMA |

| | | | | | | |
|---|---|---|---|---|---|---|
| 2.23E-126 | 0.511041 | 0.475884 | 0.137857 | 5.23E-126 | black | NR4A1 |
| 7.39E-126 | 0.554936 | 0.672026 | 0.247021 | 1.71E-125 | black | PTPRE |
| 1.95E-123 | -0.71195 | 0.985531 | 0.99478 | 4.47E-123 | black | RPS27 |
| 3.56E-121 | 1.460085 | 0.639871 | 0.300634 | 8.03E-121 | black | HLA-DQA1 |
| 1.06E-119 | 0.637722 | 0.720257 | 0.314817 | 2.37E-119 | black | GNAI2 |
| 3.45E-118 | 0.668385 | 0.848875 | 0.517975 | 7.61E-118 | black | CYBA |
| 1.24E-117 | 1.537151 | 0.847267 | 0.64687 | 2.70E-117 | black | HLA-DPA1 |
| 1.88E-115 | 0.717457 | 0.754019 | 0.435241 | 4.06E-115 | black | RNH1 |
| 2.15E-115 | 0.6263 | 0.868167 | 0.580879 | 4.59E-115 | black | ATP6V0B |
| 1.86E-114 | 0.624741 | 0.998392 | 0.999672 | 3.93E-114 | black | ACTB |
| 1.66E-113 | 1.44305 | 0.839228 | 0.648938 | 3.46E-113 | black | HLA-DPB1 |
| 1.81E-109 | 0.985897 | 0.300643 | 0.068978 | 3.74E-109 | black | AZU1 |
| 2.88E-108 | 0.542445 | 0.638264 | 0.264618 | 5.88E-108 | black | TNFSF13B |
| 1.07E-107 | 0.587273 | 0.827974 | 0.474999 | 2.16E-107 | black | MCL1 |
| 9.36E-107 | -0.50403 | 0.998392 | 0.997833 | 1.87E-106 | black | RPS4X |
| 9.92E-106 | 0.502273 | 0.990354 | 0.94839 | 1.96E-105 | black | ATP5F1E |
| 5.62E-104 | 0.578406 | 0.919614 | 0.686923 | 1.10E-103 | black | GNG5 |
| 2.31E-103 | 0.59353 | 0.901929 | 0.678748 | 4.47E-103 | black | PKM |
| 3.44E-102 | 0.723686 | 0.702572 | 0.374996 | 6.60E-102 | black | C1orf162 |
| 6.59E-102 | 0.613633 | 0.871383 | 0.618011 | 1.25E-101 | black | SAT1 |
| 3.21E-98 | -1.7466 | 0.165595 | 0.58124 | 6.05E-98 | black | IFITM1 |
| 3.73E-97 | 0.657694 | 0.832797 | 0.604485 | 6.95E-97 | black | DBI |
| 3.01E-95 | 0.579358 | 0.914791 | 0.727897 | 5.55E-95 | black | EMP3 |
| 4.08E-95 | -0.55882 | 0.877814 | 0.968482 | 7.46E-95 | black | EEF2 |
| 1.84E-94 | 0.587285 | 0.618971 | 0.295643 | 3.33E-94 | black | CCDC88A |
| 4.98E-91 | 0.507959 | 0.673633 | 0.322368 | 8.93E-91 | black | ARPC1B |
| 6.52E-90 | -0.62383 | 0.998392 | 0.999212 | 1.16E-89 | black | MALAT1 |
| 2.11E-89 | -1.15237 | 0.057878 | 0.476477 | 3.71E-89 | black | TRBC2 |
| 9.04E-89 | 0.66771 | 0.890675 | 0.683214 | 1.58E-88 | black | ANXA1 |
| 1.18E-88 | 0.516774 | 0.940514 | 0.856102 | 2.04E-88 | black | TPM3 |
| 9.53E-87 | -0.86102 | 0.824759 | 0.94944 | 1.63E-86 | black | HLA-E |
| 1.50E-86 | 0.587352 | 0.471061 | 0.165764 | 2.56E-86 | black | CFD |
| 1.60E-83 | 0.557319 | 0.893891 | 0.643061 | 2.70E-83 | black | ATP5MF |
| 2.86E-81 | 0.574701 | 0.734727 | 0.428051 | 4.77E-81 | black | RILPL2 |
| 7.43E-80 | 0.828097 | 0.646302 | 0.323714 | 1.23E-79 | black | AREG |
| 1.06E-78 | 0.584935 | 0.596463 | 0.309071 | 1.74E-78 | black | BID |
| 5.03E-78 | -0.68669 | 0.868167 | 0.919728 | 8.19E-78 | black | NPM1 |
| 3.39E-77 | -0.52565 | 0.81672 | 0.918349 | 5.47E-77 | black | CIRBP |
| 8.14E-76 | -0.85994 | 0.172026 | 0.539184 | 1.30E-75 | black | RARRES3 |
| 1.55E-73 | -1.73343 | 0.049839 | 0.408516 | 2.45E-73 | black | CD3E |
| 4.56E-73 | -0.65212 | 0.57717 | 0.786697 | 7.18E-73 | black | C12orf57 |

| | | | | | | |
|---|---|---|---|---|---|---|
| 5.34E-73 | -0.68486 | 0.045016 | 0.41111 | 8.34E-73 | black | ITM2A |
| 3.63E-72 | 0.51608 | 0.823151 | 0.53137 | 5.61E-72 | black | PGLS |
| 1.31E-69 | -0.71134 | 0.059486 | 0.420106 | 2.01E-69 | black | RHOH |
| 1.13E-68 | -1.9993 | 0.053055 | 0.395023 | 1.72E-68 | black | IL32 |
| 2.82E-67 | -0.58588 | 0.707395 | 0.863226 | 4.27E-67 | black | HSPA8 |
| 3.11E-66 | 0.510336 | 0.78135 | 0.540563 | 4.66E-66 | black | PLP2 |
| 2.11E-64 | 0.618312 | 0.442122 | 0.186743 | 3.15E-64 | black | ANKRD13D |
| 4.76E-64 | -0.72328 | 0.493569 | 0.685151 | 7.03E-64 | black | GYPC |
| 1.21E-63 | -1.33581 | 0.09164 | 0.416396 | 1.78E-63 | black | CD7 |
| 4.65E-63 | -1.01153 | 0.16881 | 0.489051 | 6.76E-63 | black | CD69 |
| 1.63E-61 | -0.76835 | 0.731511 | 0.909682 | 2.35E-61 | black | SARAF |
| 4.00E-61 | 0.500341 | 0.737942 | 0.52464 | 5.73E-61 | black | CSTB |
| 1.19E-60 | 0.544718 | 0.654341 | 0.369776 | 1.69E-60 | black | HERPUD1 |
| 8.85E-59 | 0.587698 | 0.639871 | 0.367609 | 1.25E-58 | black | MAP3K8 |
| 1.14E-57 | 0.514963 | 0.847267 | 0.623297 | 1.60E-57 | black | COX7B |
| 2.63E-56 | -0.95487 | 0.027331 | 0.325684 | 3.65E-56 | black | LCK |
| 3.61E-56 | -1.08058 | 0.027331 | 0.325388 | 4.98E-56 | black | CD3D |
| 5.57E-56 | -1.10021 | 0.310289 | 0.569027 | 7.63E-56 | black | CTSW |
| 2.83E-55 | -0.5683 | 0.786174 | 0.807939 | 3.86E-55 | black | LDHB |
| 5.22E-55 | 1.196443 | 0.435691 | 0.225155 | 7.06E-55 | black | HLA-DQA2 |
| 1.11E-50 | 0.802809 | 0.323151 | 0.123937 | 1.50E-50 | black | MPO |
| 4.03E-50 | 0.539277 | 0.636656 | 0.362717 | 5.38E-50 | black | JUND |
| 9.42E-50 | -0.86186 | 0.03537 | 0.309006 | 1.25E-49 | black | SPOCK2 |
| 3.90E-49 | -0.89822 | 0.382637 | 0.589744 | 5.12E-49 | black | EVL |
| 1.12E-47 | 0.588919 | 0.858521 | 0.609803 | 1.46E-47 | black | H2AFZ |
| 1.47E-47 | -0.74675 | 0.017685 | 0.281263 | 1.90E-47 | black | ZAP70 |
| 1.95E-46 | -0.83837 | 0.323151 | 0.560655 | 2.52E-46 | black | DNAJB1 |
| 8.70E-46 | -0.51684 | 0.734727 | 0.858367 | 1.11E-45 | black | RAC2 |
| 3.76E-45 | 0.526953 | 0.567524 | 0.325914 | 4.78E-45 | black | GRK2 |
| 3.83E-45 | -0.55446 | 0.046624 | 0.311205 | 4.84E-45 | black | EGFL7 |
| 4.56E-45 | -0.77369 | 0.329582 | 0.559276 | 5.72E-45 | black | LEPROTL1 |
| 1.76E-44 | -1.33536 | 0.041801 | 0.295151 | 2.20E-44 | black | IL7R |
| 3.97E-43 | -0.71616 | 0.019293 | 0.265866 | 4.92E-43 | black | HOPX |
| 4.94E-43 | -0.56678 | 0.033762 | 0.284284 | 6.08E-43 | black | CD34 |
| 1.17E-42 | -0.84651 | 0.016077 | 0.257362 | 1.43E-42 | black | GZMM |
| 3.93E-42 | -1.27421 | 0.167203 | 0.401162 | 4.78E-42 | black | LTB |
| 2.60E-40 | -0.70406 | 0.054662 | 0.296234 | 3.14E-40 | black | PIK3IP1 |
| 5.97E-40 | -0.68577 | 0.05627 | 0.298861 | 7.16E-40 | black | FCMR |
| 7.30E-40 | 0.528103 | 0.364952 | 0.174004 | 8.71E-40 | black | H1FX |
| 1.51E-39 | -0.62645 | 0.036977 | 0.271447 | 1.78E-39 | black | LAT |
| 7.14E-39 | -0.68132 | 0.633441 | 0.790078 | 8.42E-39 | black | CD52 |

| | | | | | | |
|---|---|---|---|---|---|---|
| 7.42E-39 | -0.74994 | 0.030547 | 0.256312 | 8.69E-39 | black | CD3G |
| 3.40E-38 | -0.59379 | 0.212219 | 0.444663 | 3.96E-38 | black | 1-Sep |
| 2.63E-36 | -0.79779 | 0.678457 | 0.795988 | 3.04E-36 | black | TXNIP |
| 5.39E-36 | -0.53761 | 0.996785 | 0.999737 | 6.20E-36 | black | B2M |
| 1.13E-35 | -0.50258 | 0.141479 | 0.3747 | 1.29E-35 | black | PBXIP1 |
| 2.91E-35 | -0.59545 | 0.051447 | 0.270199 | 3.31E-35 | black | LBH |
| 7.43E-35 | -0.79927 | 0.041801 | 0.255655 | 8.40E-35 | black | SPINK2 |
| 2.16E-33 | -0.60512 | 0.228296 | 0.432253 | 2.43E-33 | black | TBC1D10C |
| 5.14E-33 | -0.5776 | 0.065916 | 0.272103 | 5.74E-33 | black | PRKCH |
| 7.27E-33 | -0.50129 | 0.345659 | 0.540628 | 8.07E-33 | black | ACAP1 |
| 8.95E-33 | -0.61386 | 0.215434 | 0.404544 | 9.88E-33 | black | CDK6 |
| 1.11E-32 | -0.54399 | 0.053055 | 0.259496 | 1.22E-32 | black | ABLIM1 |
| 1.41E-32 | -0.64198 | 0.057878 | 0.260219 | 1.53E-32 | black | FLT3LG |
| 3.02E-32 | -0.55006 | 0.292605 | 0.48892 | 3.27E-32 | black | BIN2 |
| 2.30E-29 | -0.92287 | 0.257235 | 0.42503 | 2.48E-29 | black | TNFAIP3 |
| 2.53E-28 | -0.59408 | 0.073955 | 0.264093 | 2.71E-28 | black | SMIM24 |
| 3.86E-28 | -0.93044 | 0.064309 | 0.259496 | 4.11E-28 | black | IGLL1 |
| 8.69E-28 | -0.50118 | 0.209003 | 0.393677 | 9.22E-28 | black | CYFIP2 |
| 6.04E-27 | -0.52071 | 0.258842 | 0.443416 | 6.37E-27 | black | PIK3R1 |
| 2.69E-26 | -0.55696 | 0.207395 | 0.380807 | 2.83E-26 | black | P2RY8 |
| 3.07E-26 | -0.57807 | 0.157556 | 0.334056 | 3.20E-26 | black | ISG20 |
| 7.28E-25 | -0.5882 | 0.350482 | 0.51223 | 7.56E-25 | black | NOSIP |
| 4.62E-24 | -0.63211 | 0.635048 | 0.779901 | 4.77E-24 | black | JUN |
| 7.43E-23 | -0.66923 | 0.167203 | 0.339374 | 7.63E-23 | black | PRSS57 |
| 2.04E-20 | -0.58448 | 0.755627 | 0.819856 | 2.08E-20 | black | TSC22D3 |
| 6.43E-18 | -0.51867 | 0.678457 | 0.749401 | 6.53E-18 | black | PTPRC |
| 4.56E-11 | -1.32392 | 0.487138 | 0.301684 | 4.61E-11 | black | NKG7 |
| 0.022167 | -0.80856 | 0.400322 | 0.294626 | 0.022283 | black | CST7 |
| 0 | 4.601167 | 1 | 0.203757 | 0 | purple | HBB |
| 0 | 3.13059 | 0.935252 | 0.058896 | 0 | purple | CA1 |
| 0 | 2.981082 | 0.997602 | 0.03359 | 0 | purple | AHSP |
| 0 | 2.207559 | 0.98801 | 0.08479 | 0 | purple | HBD |
| 0 | 2.128665 | 1 | 0.26898 | 0 | purple | BLVRB |
| 0 | 1.800372 | 0.995204 | 0.034144 | 0 | purple | FAM178B |
| 0 | 1.705627 | 1 | 0.058831 | 0 | purple | APOC1 |
| 0 | 1.547588 | 0.997602 | 0.039166 | 0 | purple | KCNH2 |
| 0 | 1.477336 | 0.997602 | 0.068419 | 0 | purple | KLF1 |
| 0 | 1.406327 | 1 | 0.27061 | 0 | purple | SYNGR1 |
| 0 | 1.400413 | 0.971223 | 0.069169 | 0 | purple | CA2 |
| 0 | 1.392821 | 1 | 0.275894 | 0 | purple | REXO2 |
| 0 | 1.359163 | 1 | 0.089356 | 0 | purple | GATA1 |

| | | | | | | |
|---|---|---|---|---|---|---|
| 0 | 1.299182 | 1 | 0.345617 | 0 | purple | UROD |
| 0 | 1.245377 | 0.781775 | 0.009392 | 0 | purple | HBA1 |
| 0 | 1.226333 | 0.990408 | 0.149393 | 0 | purple | HMBS |
| 0 | 1.163795 | 0.990408 | 0.019371 | 0 | purple | PKLR |
| 0 | 1.159606 | 1 | 0.203789 | 0 | purple | NME4 |
| 0 | 1.159356 | 0.985612 | 0.070604 | 0 | purple | FN3K |
| 0 | 1.152697 | 0.966427 | 0.048722 | 0 | purple | EPCAM |
| 0 | 1.145978 | 0.980815 | 0.018491 | 0 | purple | RHAG |
| 0 | 1.13826 | 0.892086 | 0.011316 | 0 | purple | NMU |
| 0 | 1.120595 | 1 | 0.178548 | 0 | purple | LMNA |
| 0 | 1.081356 | 0.990408 | 0.176298 | 0 | purple | HEBP1 |
| 0 | 1.050725 | 0.992806 | 0.117728 | 0 | purple | CD36 |
| 0 | 1.007185 | 0.98801 | 0.211062 | 0 | purple | AKR1C3 |
| 0 | 1.001947 | 0.980815 | 0.242662 | 0 | purple | CKS2 |
| 0 | 1.000629 | 0.992806 | 0.091834 | 0 | purple | CNRIP1 |
| 0 | 0.981827 | 0.98801 | 0.107031 | 0 | purple | FECH |
| 0 | 0.976242 | 0.980815 | 0.109183 | 0 | purple | UBAC1 |
| 0 | 0.974796 | 0.990408 | 0.048624 | 0 | purple | ANK1 |
| 0 | 0.973992 | 0.779376 | 0.004794 | 0 | purple | ALAS2 |
| 0 | 0.952457 | 0.985612 | 0.07243 | 0 | purple | SMIM1 |
| 0 | 0.93103 | 0.990408 | 0.196745 | 0 | purple | ALDH1A1 |
| 0 | 0.902101 | 0.971223 | 0.09004 | 0 | purple | PVT1 |
| 0 | 0.868276 | 0.935252 | 0.037145 | 0 | purple | HBQ1 |
| 0 | 0.863813 | 0.966427 | 0.144893 | 0 | purple | SLC29A1 |
| 0 | 0.86188 | 0.935252 | 0.072756 | 0 | purple | GFI1B |
| 0 | 0.851093 | 0.983213 | 0.172417 | 0 | purple | LPCAT3 |
| 0 | 0.842562 | 0.976019 | 0.068028 | 0 | purple | TFR2 |
| 0 | 0.836004 | 0.98801 | 0.15073 | 0 | purple | ENG |
| 0 | 0.835846 | 0.952038 | 0.118836 | 0 | purple | MINPP1 |
| 0 | 0.833599 | 0.973621 | 0.229553 | 0 | purple | OAT |
| 0 | 0.82829 | 0.769784 | 0.090334 | 0 | purple | HIST1H1B |
| 0 | 0.815436 | 0.901679 | 0.017415 | 0 | purple | MYL4 |
| 0 | 0.805776 | 0.964029 | 0.135729 | 0 | purple | CENPF |
| 0 | 0.805376 | 0.976019 | 0.104455 | 0 | purple | TST |
| 0 | 0.800551 | 0.870504 | 0.079703 | 0 | purple | TUBB2A |
| 0 | 0.800325 | 0.966427 | 0.176167 | 0 | purple | TFRC |
| 0 | 0.784036 | 0.983213 | 0.184549 | 0 | purple | CENPU |
| 0 | 0.783719 | 0.781775 | 0.061342 | 0 | purple | CDC20 |
| 0 | 0.783699 | 0.964029 | 0.138729 | 0 | purple | DHFR |
| 0 | 0.783551 | 0.992806 | 0.216541 | 0 | purple | ISOC2 |
| 0 | 0.780173 | 0.995204 | 0.244913 | 0 | purple | BOLA3 |

| | | | | | | |
|---|---|---|---|---|---|---|
| 0 | 0.768791 | 0.940048 | 0.091606 | 0 | purple | CCNB2 |
| 0 | 0.765087 | 0.968825 | 0.104716 | 0 | purple | SMIM10 |
| 0 | 0.753462 | 0.964029 | 0.097737 | 0 | purple | NFIA |
| 0 | 0.75316 | 0.983213 | 0.180244 | 0 | purple | PCCB |
| 0 | 0.750836 | 0.966427 | 0.17183 | 0 | purple | FEN1 |
| 0 | 0.746693 | 0.973621 | 0.191528 | 0 | purple | SNCA |
| 0 | 0.741162 | 0.93765 | 0.188299 | 0 | purple | UNG |
| 0 | 0.733742 | 0.959233 | 0.067082 | 0 | purple | EPOR |
| 0 | 0.731682 | 0.822542 | 0.076376 | 0 | purple | CCNB1 |
| 0 | 0.731317 | 0.978417 | 0.193582 | 0 | purple | HNRNPAB |
| 0 | 0.730673 | 0.817746 | 0.01523 | 0 | purple | SELENBP1 |
| 0 | 0.728781 | 0.983213 | 0.189897 | 0 | purple | MCM2 |
| 0 | 0.707899 | 0.942446 | 0.133903 | 0 | purple | SLC43A3 |
| 0 | 0.706899 | 0.973621 | 0.139969 | 0 | purple | HMGN5 |
| 0 | 0.698122 | 0.961631 | 0.143393 | 0 | purple | GMNN |
| 0 | 0.683745 | 0.968825 | 0.139153 | 0 | purple | SLC38A5 |
| 0 | 0.681361 | 0.923261 | 0.021132 | 0 | purple | ADD2 |
| 0 | 0.675105 | 0.966427 | 0.186016 | 0 | purple | CENPW |
| 0 | 0.663213 | 0.872902 | 0.02772 | 0 | purple | DNAJA4 |
| 0 | 0.653082 | 0.973621 | 0.189082 | 0 | purple | FKBP4 |
| 0 | 0.649902 | 0.93765 | 0.135403 | 0 | purple | FADS2 |
| 0 | 0.63684 | 0.942446 | 0.113195 | 0 | purple | LXN |
| 0 | 0.629586 | 0.954436 | 0.107031 | 0 | purple | NMNAT3 |
| 0 | 0.6246 | 0.865707 | 0.022828 | 0 | purple | SPTA1 |
| 0 | 0.617171 | 0.604317 | 0.002348 | 0 | purple | GYPA |
| 0 | 0.617042 | 0.956835 | 0.189212 | 0 | purple | FSCN1 |
| 0 | 0.614876 | 0.901679 | 0.091051 | 0 | purple | TUBG1 |
| 0 | 0.611767 | 0.971223 | 0.196908 | 0 | purple | DTYMK |
| 0 | 0.611347 | 0.966427 | 0.201311 | 0 | purple | NOP16 |
| 0 | 0.610383 | 0.918465 | 0.074909 | 0 | purple | FHL2 |
| 0 | 0.608992 | 0.81295 | 0.084399 | 0 | purple | BIRC5 |
| 0 | 0.608984 | 0.940048 | 0.103868 | 0 | purple | PYCR1 |
| 0 | 0.600832 | 0.892086 | 0.08916 | 0 | purple | UBE2T |
| 0 | 0.599641 | 0.942446 | 0.117206 | 0 | purple | ARV1 |
| 0 | 0.596577 | 0.961631 | 0.127739 | 0 | purple | TPM1 |
| 0 | 0.592208 | 0.918465 | 0.141371 | 0 | purple | GINS2 |
| 0 | 0.585509 | 0.877698 | 0.098356 | 0 | purple | TPX2 |
| 0 | 0.585202 | 0.892086 | 0.027296 | 0 | purple | KEL |
| 0 | 0.584271 | 0.928058 | 0.164134 | 0 | purple | MCM4 |
| 0 | 0.569256 | 0.796163 | 0.085181 | 0 | purple | TOP2A |
| 0 | 0.566215 | 0.582734 | 0.063788 | 0 | purple | HIST1H3B |

| | | | | | | |
|---|---|---|---|---|---|---|
| 0 | 0.565753 | 0.911271 | 0.128196 | 0 | purple | ZWINT |
| 0 | 0.565497 | 0.942446 | 0.124315 | 0 | purple | PLA2G12A |
| 0 | 0.563269 | 0.916067 | 0.125946 | 0 | purple | CENPM |
| 0 | 0.561178 | 0.952038 | 0.177407 | 0 | purple | SMC2 |
| 0 | 0.556145 | 0.925659 | 0.127185 | 0 | purple | PCLAF |
| 0 | 0.55596 | 0.964029 | 0.189995 | 0 | purple | AK6 |
| 0 | 0.55499 | 0.752998 | 0.059907 | 0 | purple | CCNA2 |
| 0 | 0.553801 | 0.954436 | 0.18432 | 0 | purple | MAD2L1 |
| 0 | 0.553376 | 0.870504 | 0.032514 | 0 | purple | RFESD |
| 0 | 0.54861 | 0.508393 | 0.005185 | 0 | purple | HBA2 |
| 0 | 0.53878 | 0.976019 | 0.198115 | 0 | purple | HBS1L |
| 0 | 0.537113 | 0.733813 | 0.005348 | 0 | purple | EPB42 |
| 0 | 0.536543 | 0.971223 | 0.193419 | 0 | purple | MIR4435-2HG |
| 0 | 0.536004 | 0.714628 | 0.088508 | 0 | purple | UBE2C |
| 0 | 0.531878 | 0.932854 | 0.157416 | 0 | purple | RFC4 |
| 0 | 0.524998 | 0.880096 | 0.037438 | 0 | purple | MFSD2B |
| 0 | 0.524393 | 0.906475 | 0.149622 | 0 | purple | ALAD |
| 0 | 0.524381 | 0.906475 | 0.143589 | 0 | purple | TRAP1 |
| 0 | 0.522858 | 0.724221 | 0.065223 | 0 | purple | CDK1 |
| 0 | 0.521393 | 0.940048 | 0.181972 | 0 | purple | ACTL6A |
| 0 | 0.518579 | 0.800959 | 0.09066 | 0 | purple | MKI67 |
| 0 | 0.514077 | 0.947242 | 0.185592 | 0 | purple | STK25 |
| 0 | 0.512446 | 0.841727 | 0.039721 | 0 | purple | APOBEC3B |
| 0 | 0.509584 | 0.834532 | 0.107064 | 0 | purple | TK1 |
| 0 | 0.50894 | 0.930456 | 0.160057 | 0 | purple | SLC1A5 |
| 0 | 0.502416 | 0.925659 | 0.144273 | 0 | purple | PRKAB1 |
| 0 | 0.501657 | 0.925659 | 0.151024 | 0 | purple | WDR34 |
| 0 | 0.501291 | 0.923261 | 0.147469 | 0 | purple | CMBL |
| 4.61E-307 | 0.697042 | 0.98801 | 0.238912 | 2.30E-306 | purple | PSMG1 |
| 7.27E-307 | 0.628862 | 0.983213 | 0.21703 | 3.60E-306 | purple | MTHFD1 |
| 1.43E-303 | 0.808027 | 0.985612 | 0.227759 | 7.03E-303 | purple | PCNA |
| 4.53E-302 | 1.433056 | 1 | 0.429624 | 2.21E-301 | purple | TMEM14C |
| 9.30E-301 | 0.673628 | 0.978417 | 0.205877 | 4.50E-300 | purple | PLIN2 |
| 2.87E-297 | 1.044702 | 0.947242 | 0.210931 | 1.38E-296 | purple | MYC |
| 1.13E-296 | 0.654742 | 0.995204 | 0.239858 | 5.36E-296 | purple | GAR1 |
| 8.24E-295 | 0.901025 | 0.995204 | 0.298852 | 3.90E-294 | purple | TPGS2 |
| 5.79E-294 | 2.198922 | 1 | 0.544743 | 2.72E-293 | purple | PRDX2 |
| 5.28E-293 | 0.518823 | 0.954436 | 0.199322 | 2.46E-292 | purple | PSMD12 |
| 7.38E-293 | 0.502054 | 0.930456 | 0.194691 | 3.41E-292 | purple | KEAP1 |
| 7.64E-290 | 0.809507 | 0.964029 | 0.216606 | 3.50E-289 | purple | SLC25A37 |
| 2.56E-282 | 0.849334 | 1 | 0.293993 | 1.17E-281 | purple | PAICS |

| | | | | | | |
|---|---|---|---|---|---|---|
| 3.37E-280 | 0.659157 | 0.995204 | 0.255283 | 1.52E-279 | purple | BLVRA |
| 4.44E-279 | 0.681316 | 0.928058 | 0.205616 | 1.99E-278 | purple | PTTG1 |
| 1.13E-276 | 0.540995 | 0.973621 | 0.222998 | 5.03E-276 | purple | MPDU1 |
| 1.19E-275 | 0.509467 | 0.956835 | 0.203626 | 5.27E-275 | purple | RRM1 |
| 1.11E-274 | 0.509608 | 0.935252 | 0.195637 | 4.86E-274 | purple | NFE2 |
| 3.30E-274 | 0.578279 | 0.976019 | 0.227041 | 1.44E-273 | purple | HMGB3 |
| 4.80E-274 | 0.56424 | 0.961631 | 0.219736 | 2.08E-273 | purple | MPST |
| 1.12E-273 | 0.582446 | 0.968825 | 0.232879 | 4.83E-273 | purple | ECI2 |
| 1.39E-272 | 0.796464 | 0.985612 | 0.255609 | 5.92E-272 | purple | MCM3 |
| 3.90E-271 | 0.518881 | 0.930456 | 0.198898 | 1.65E-270 | purple | MCM6 |
| 2.96E-267 | 0.973315 | 0.997602 | 0.357846 | 1.24E-266 | purple | CDK4 |
| 2.00E-265 | 0.841396 | 1 | 0.279579 | 8.36E-265 | purple | CAST |
| 2.49E-265 | 0.504739 | 0.956835 | 0.218139 | 1.03E-264 | purple | MRTO4 |
| 1.21E-264 | 1.412544 | 1 | 0.697691 | 4.97E-264 | purple | ATP5IF1 |
| 3.23E-263 | 0.536169 | 0.973621 | 0.234738 | 1.33E-262 | purple | MRPL13 |
| 7.68E-262 | 0.542329 | 0.980815 | 0.238651 | 3.13E-261 | purple | MRPL27 |
| 2.87E-256 | 0.589927 | 0.985612 | 0.252935 | 1.16E-255 | purple | CISD2 |
| 3.70E-255 | 0.552398 | 0.918465 | 0.201017 | 1.49E-254 | purple | CKS1B |
| 1.35E-254 | -2.90683 | 0.815348 | 0.996119 | 5.41E-254 | purple | TMSB4X |
| 3.71E-254 | 0.722019 | 0.98801 | 0.285384 | 1.47E-253 | purple | NARF |
| 4.90E-254 | 0.694151 | 0.985612 | 0.279383 | 1.93E-253 | purple | PSMB5 |
| 5.91E-254 | 0.95765 | 0.997602 | 0.313984 | 2.31E-253 | purple | MCM7 |
| 1.13E-251 | 0.881876 | 0.995204 | 0.342258 | 4.40E-251 | purple | ATG3 |
| 2.64E-251 | 1.218412 | 1 | 0.523741 | 1.02E-250 | purple | HSPD1 |
| 7.00E-251 | 0.533483 | 0.961631 | 0.233662 | 2.69E-250 | purple | IARS |
| 1.95E-250 | 0.53444 | 0.961631 | 0.228411 | 7.45E-250 | purple | RIF1 |
| 2.24E-248 | 0.630301 | 0.985612 | 0.266208 | 8.49E-248 | purple | UQCC2 |
| 4.23E-247 | 1.082266 | 1 | 0.479292 | 1.60E-246 | purple | CCT6A |
| 1.05E-242 | 0.711868 | 0.990408 | 0.292362 | 3.95E-242 | purple | DKC1 |
| 6.82E-242 | 0.748614 | 0.997602 | 0.323082 | 2.54E-241 | purple | RUVBL2 |
| 1.09E-240 | 0.761089 | 0.997602 | 0.302342 | 4.02E-240 | purple | DNAJC9 |
| 1.35E-239 | 0.624561 | 0.992806 | 0.284373 | 4.98E-239 | purple | MTCH2 |
| 1.45E-239 | 0.522352 | 0.968825 | 0.243054 | 5.33E-239 | purple | IFRD2 |
| 8.48E-239 | 0.506675 | 0.973621 | 0.244293 | 3.09E-238 | purple | FARSA |
| 1.91E-235 | 0.807614 | 0.992806 | 0.292558 | 6.91E-235 | purple | MCM5 |
| 1.94E-235 | 0.588344 | 0.980815 | 0.269371 | 7.00E-235 | purple | UROS |
| 2.08E-235 | 0.963031 | 1 | 0.493282 | 7.43E-235 | purple | ATP5MC1 |
| 9.25E-234 | 0.898313 | 0.997602 | 0.408133 | 3.29E-233 | purple | FBXO7 |
| 1.68E-233 | 0.7131 | 0.992806 | 0.282481 | 5.94E-233 | purple | CYTOR |
| 5.61E-232 | 1.041553 | 1 | 0.710442 | 1.97E-231 | purple | RAN |
| 9.11E-232 | 0.840354 | 0.995204 | 0.339029 | 3.19E-231 | purple | HMGA1 |

| | | | | | | |
|---|---|---|---|---|---|---|
| 1.45E-231 | 0.546211 | 0.983213 | 0.265914 | 5.04E-231 | purple | PDHA1 |
| 7.17E-231 | 0.551149 | 0.985612 | 0.263403 | 2.48E-230 | purple | TMEM141 |
| 1.16E-229 | 0.857578 | 0.992806 | 0.373435 | 3.97E-229 | purple | PRDX3 |
| 1.55E-229 | -2.09197 | 0.863309 | 0.970291 | 5.29E-229 | purple | SH3BGRL3 |
| 1.85E-229 | 0.588988 | 0.98801 | 0.282481 | 6.30E-229 | purple | MRPS2 |
| 3.82E-229 | 0.576383 | 0.978417 | 0.261903 | 1.29E-228 | purple | MGST2 |
| 2.46E-228 | 0.951013 | 1 | 0.513599 | 8.28E-228 | purple | TMEM14B |
| 3.15E-228 | 0.753421 | 0.992806 | 0.341834 | 1.05E-227 | purple | MRPL37 |
| 5.10E-228 | 0.527922 | 0.980815 | 0.263469 | 1.70E-227 | purple | CHID1 |
| 6.15E-227 | 0.826497 | 0.997602 | 0.371413 | 2.03E-226 | purple | METAP2 |
| 6.54E-226 | 0.817342 | 0.995204 | 0.376174 | 2.15E-225 | purple | SLIRP |
| 9.46E-226 | 0.62639 | 0.992806 | 0.284927 | 3.10E-225 | purple | SUPT16H |
| 3.50E-225 | 0.507319 | 0.966427 | 0.247358 | 1.14E-224 | purple | SLC39A8 |
| 1.54E-224 | -2.00582 | 0.956835 | 0.986629 | 5.00E-224 | purple | HLA-A |
| 5.64E-222 | 0.761358 | 0.980815 | 0.317245 | 1.82E-221 | purple | SLC25A39 |
| 6.56E-222 | 0.899884 | 1 | 0.423265 | 2.10E-221 | purple | MPC2 |
| 9.04E-222 | -2.02642 | 0.973621 | 0.997717 | 2.88E-221 | purple | HLA-B |
| 3.58E-221 | 0.622031 | 0.98801 | 0.295819 | 1.14E-220 | purple | THOC3 |
| 8.54E-218 | -2.14843 | 0.613909 | 0.951474 | 2.70E-217 | purple | HLA-E |
| 9.33E-218 | -1.6283 | 1 | 0.999674 | 2.93E-217 | purple | B2M |
| 2.78E-217 | 0.539327 | 0.978417 | 0.27498 | 8.70E-217 | purple | SAMM50 |
| 1.27E-216 | 0.939572 | 1 | 0.550548 | 3.94E-216 | purple | HSPE1 |
| 7.82E-216 | 0.588268 | 0.98801 | 0.295656 | 2.42E-215 | purple | PPA2 |
| 9.27E-216 | 0.507108 | 0.956835 | 0.255283 | 2.86E-215 | purple | PSMD1 |
| 2.27E-215 | -1.35652 | 0.973621 | 0.9699 | 6.95E-215 | purple | ARHGDIB |
| 1.23E-214 | -2.81287 | 0.220624 | 0.916612 | 3.76E-214 | purple | VIM |
| 3.35E-213 | -1.91109 | 0.990408 | 0.993282 | 1.02E-212 | purple | HLA-C |
| 1.48E-212 | 1.025321 | 1 | 0.659568 | 4.46E-212 | purple | RPL22L1 |
| 2.51E-212 | 0.842648 | 1 | 0.706855 | 7.54E-212 | purple | ATP5MC3 |
| 3.32E-212 | 0.578899 | 0.978417 | 0.278372 | 9.92E-212 | purple | HADH |
| 4.07E-212 | 0.718943 | 0.997602 | 0.327191 | 1.21E-211 | purple | GLRX |
| 3.75E-211 | 0.654013 | 0.992806 | 0.330257 | 1.11E-210 | purple | TIMM8B |
| 6.51E-211 | 0.579167 | 0.978417 | 0.285514 | 1.92E-210 | purple | PRDX4 |
| 1.04E-210 | 0.752976 | 1 | 0.342486 | 3.05E-210 | purple | CD82 |
| 2.60E-210 | 1.288682 | 1 | 0.619815 | 7.59E-210 | purple | TUBB |
| 7.48E-209 | 0.520281 | 0.978417 | 0.275861 | 2.17E-208 | purple | EBNA1BP2 |
| 1.57E-205 | 0.648891 | 1 | 0.315941 | 4.55E-205 | purple | RPA3 |
| 4.42E-203 | 0.850935 | 1 | 0.553809 | 1.27E-202 | purple | PA2G4 |
| 6.97E-203 | 0.799267 | 1 | 0.428287 | 2.00E-202 | purple | TIMM13 |
| 7.48E-203 | 0.516885 | 0.848921 | 0.205648 | 2.13E-202 | purple | EIF1AY |
| 5.36E-200 | 0.61577 | 0.992806 | 0.323963 | 1.52E-199 | purple | PAFAH1B3 |

| | | | | | | |
|---|---|---|---|---|---|---|
| 8.55E-200 | 0.830059 | 1 | 0.473519 | 2.42E-199 | purple | NOP10 |
| 6.90E-199 | 0.702184 | 0.997602 | 0.35951 | 1.94E-198 | purple | HPRT1 |
| 8.23E-199 | 0.5637 | 0.990408 | 0.304396 | 2.31E-198 | purple | LRRC59 |
| 1.35E-198 | 0.605424 | 1 | 0.32592 | 3.76E-198 | purple | MPP1 |
| 5.78E-198 | 0.731395 | 1 | 0.362771 | 1.60E-197 | purple | RANBP1 |
| 9.66E-198 | 1.028215 | 0.997602 | 0.397176 | 2.67E-197 | purple | TUBB4B |
| 1.14E-197 | 0.753015 | 1 | 0.395806 | 3.15E-197 | purple | CCT5 |
| 1.59E-196 | 0.691181 | 1 | 0.373793 | 4.35E-196 | purple | STOML2 |
| 2.70E-195 | 0.555972 | 0.985612 | 0.306288 | 7.36E-195 | purple | APEH |
| 3.20E-195 | 0.644962 | 0.997602 | 0.351194 | 8.68E-195 | purple | ETFA |
| 3.81E-193 | 0.62753 | 0.995204 | 0.333062 | 1.03E-192 | purple | CENPX |
| 8.50E-193 | 1.003721 | 1 | 0.860455 | 2.29E-192 | purple | HSP90AA1 |
| 2.32E-192 | 0.658014 | 0.995204 | 0.335149 | 6.23E-192 | purple | PRKDC |
| 2.84E-192 | 0.570288 | 0.992806 | 0.309549 | 7.59E-192 | purple | ATIC |
| 7.69E-192 | 0.740506 | 1 | 0.423363 | 2.04E-191 | purple | VDAC3 |
| 1.58E-191 | 0.603164 | 0.995204 | 0.331888 | 4.18E-191 | purple | MRPL3 |
| 1.86E-191 | -1.24647 | 0.973621 | 0.948539 | 4.89E-191 | purple | FXYD5 |
| 2.87E-191 | -1.46477 | 1 | 0.998728 | 7.53E-191 | purple | TMSB10 |
| 8.78E-191 | 0.869754 | 1 | 0.918993 | 2.29E-190 | purple | HMGB1 |
| 9.14E-191 | 0.735034 | 1 | 0.419319 | 2.38E-190 | purple | MRPS34 |
| 1.62E-188 | 0.650134 | 0.997602 | 0.339877 | 4.19E-188 | purple | DNPH1 |
| 4.48E-188 | 0.672091 | 1 | 0.391045 | 1.16E-187 | purple | TAF9 |
| 1.52E-187 | -1.45329 | 0.992806 | 0.972672 | 3.91E-187 | purple | ACTG1 |
| 7.66E-187 | 0.571438 | 0.990408 | 0.317473 | 1.96E-186 | purple | POLD2 |
| 2.08E-186 | 0.518468 | 0.985612 | 0.293178 | 5.30E-186 | purple | NOLC1 |
| 2.22E-186 | 0.800038 | 1 | 0.50512 | 5.63E-186 | purple | CCT2 |
| 7.15E-186 | 1.106811 | 1 | 0.657677 | 1.81E-185 | purple | TUBA1B |
| 3.87E-185 | 0.59649 | 0.995204 | 0.337921 | 9.74E-185 | purple | PCBP1 |
| 5.00E-185 | 0.795378 | 1 | 0.488456 | 1.25E-184 | purple | EIF3A |
| 2.35E-183 | 0.697916 | 0.995204 | 0.368804 | 5.87E-183 | purple | AHCY |
| 3.70E-183 | 0.719497 | 1 | 0.441658 | 9.19E-183 | purple | PHB |
| 4.53E-182 | 0.976501 | 1 | 0.609542 | 1.12E-181 | purple | H2AFZ |
| 1.58E-180 | 0.618253 | 0.98801 | 0.348226 | 3.90E-180 | purple | COPS3 |
| 6.71E-179 | 0.767693 | 0.992806 | 0.357879 | 1.65E-178 | purple | TUBA1C |
| 2.98E-176 | 0.570247 | 0.978417 | 0.333942 | 7.29E-176 | purple | ADSL |
| 5.22E-176 | 0.656735 | 0.98801 | 0.351683 | 1.27E-175 | purple | C1QBP |
| 5.74E-176 | -1.69955 | 0.786571 | 0.908981 | 1.39E-175 | purple | CD37 |
| 8.66E-176 | 0.734737 | 1 | 0.653144 | 2.09E-175 | purple | SERBP1 |
| 5.87E-175 | -1.86583 | 0.143885 | 0.829703 | 1.41E-174 | purple | CORO1A |
| 3.81E-174 | 0.513504 | 0.985612 | 0.304559 | 9.13E-174 | purple | SCCPDH |
| 1.29E-173 | 0.563595 | 0.995204 | 0.344541 | 3.09E-173 | purple | SMS |

| | | | | | | |
|---|---|---|---|---|---|---|
| 1.33E-173 | 0.711954 | 1 | 0.462888 | 3.17E-173 | purple | SNRPE |
| 1.46E-172 | 0.821955 | 1 | 0.676983 | 3.47E-172 | purple | GYPC |
| 4.09E-172 | 0.735527 | 1 | 0.535514 | 9.67E-172 | purple | SNRPG |
| 4.29E-172 | 0.567507 | 0.990408 | 0.339649 | 1.01E-171 | purple | EIF4EBP1 |
| 4.43E-170 | 0.842903 | 1 | 0.609575 | 1.04E-169 | purple | YBX1 |
| 6.50E-169 | 0.566413 | 0.980815 | 0.335247 | 1.52E-168 | purple | GADD45GIP1 |
| 9.48E-169 | 0.735418 | 0.997602 | 0.358825 | 2.21E-168 | purple | AC084033.3 |
| 2.44E-168 | 0.640855 | 0.995204 | 0.403535 | 5.65E-168 | purple | MRPL51 |
| 2.75E-168 | 0.679487 | 1 | 0.705061 | 6.36E-168 | purple | NDUFS5 |
| 7.56E-167 | -0.50195 | 1 | 0.999348 | 1.74E-166 | purple | FAU |
| 1.63E-165 | 0.704245 | 1 | 0.54021 | 3.73E-165 | purple | SNRPD1 |
| 2.36E-165 | 0.674486 | 1 | 0.688788 | 5.39E-165 | purple | NDUFB9 |
| 6.09E-165 | 0.658551 | 1 | 0.47802 | 1.39E-164 | purple | NDUFAB1 |
| 7.98E-165 | 0.689922 | 1 | 0.427994 | 1.81E-164 | purple | IDH2 |
| 1.12E-164 | 0.711622 | 0.997602 | 0.407579 | 2.54E-164 | purple | NASP |
| 3.02E-163 | 0.542896 | 0.995204 | 0.360031 | 6.79E-163 | purple | C20orf27 |
| 3.36E-163 | 0.690462 | 1 | 0.643295 | 7.55E-163 | purple | ATP5MF |
| 5.49E-163 | -1.7807 | 0.194245 | 0.812973 | 1.23E-162 | purple | LSP1 |
| 1.02E-162 | 0.665392 | 1 | 0.457442 | 2.27E-162 | purple | CACYBP |
| 1.41E-162 | 0.516504 | 0.992806 | 0.347541 | 3.14E-162 | purple | BCCIP |
| 5.74E-162 | -2.20921 | 0.105516 | 0.796211 | 1.27E-161 | purple | CD52 |
| 6.05E-161 | 0.648261 | 1 | 0.431385 | 1.33E-160 | purple | NDUFB6 |
| 1.44E-160 | 0.641171 | 1 | 0.446224 | 3.16E-160 | purple | NUDC |
| 5.82E-160 | 0.675981 | 1 | 0.631164 | 1.27E-159 | purple | COX5A |
| 3.71E-158 | 0.513393 | 0.985612 | 0.343269 | 8.08E-158 | purple | EIF5B |
| 1.53E-157 | 0.562216 | 0.997602 | 0.369032 | 3.31E-157 | purple | SSRP1 |
| 6.01E-157 | 0.843542 | 1 | 0.531731 | 1.30E-156 | purple | DUT |
| 1.26E-156 | 0.611417 | 1 | 0.434483 | 2.72E-156 | purple | MRPL52 |
| 8.18E-155 | 0.595525 | 0.997602 | 0.423004 | 1.76E-154 | purple | TXN2 |
| 1.41E-154 | 0.721156 | 0.997602 | 0.448702 | 3.03E-154 | purple | CAT |
| 2.86E-154 | 0.678888 | 0.997602 | 0.512099 | 6.09E-154 | purple | SSB |
| 6.06E-154 | 0.704497 | 1 | 0.586225 | 1.29E-153 | purple | BSG |
| 2.92E-151 | 0.730098 | 1 | 0.65122 | 6.18E-151 | purple | EIF5A |
| 6.79E-151 | -1.73371 | 0.153477 | 0.782285 | 1.44E-150 | purple | IFITM2 |
| 7.93E-150 | 0.630597 | 1 | 0.61988 | 1.67E-149 | purple | UQCRQ |
| 1.06E-149 | 0.504 | 0.992806 | 0.366521 | 2.22E-149 | purple | POLR2F |
| 2.13E-149 | -2.08117 | 0.366906 | 0.824713 | 4.45E-149 | purple | TSC22D3 |
| 5.09E-149 | 0.696578 | 1 | 0.80844 | 1.06E-148 | purple | NCL |
| 4.08E-148 | 0.691253 | 1 | 0.538906 | 8.46E-148 | purple | ILF2 |
| 4.84E-148 | 0.67537 | 1 | 0.553059 | 1.00E-147 | purple | HNRNPF |
| 5.87E-148 | -0.53455 | 1 | 0.999543 | 1.21E-147 | purple | RPL30 |

| | | | | | | |
|---|---|---|---|---|---|---|
| 2.70E-146 | 0.613311 | 1 | 0.69593 | 5.55E-146 | purple | SOD1 |
| 2.01E-145 | 0.701756 | 1 | 0.47065 | 4.13E-145 | purple | STMN1 |
| 2.13E-145 | 0.607534 | 1 | 0.489434 | 4.35E-145 | purple | NDUFV2 |
| 2.65E-145 | 0.508126 | 0.985612 | 0.364108 | 5.39E-145 | purple | DECR1 |
| 7.98E-144 | 0.650232 | 1 | 0.821387 | 1.62E-143 | purple | S100A6 |
| 9.18E-144 | 1.352143 | 0.992806 | 0.58991 | 1.85E-143 | purple | HIST1H4C |
| 9.18E-144 | -0.97122 | 1 | 0.992467 | 1.85E-143 | purple | H3F3B |
| 1.02E-143 | 0.628399 | 1 | 0.683114 | 2.05E-143 | purple | PTGES3 |
| 1.33E-143 | -0.51996 | 1 | 0.999185 | 2.66E-143 | purple | RPL39 |
| 1.51E-143 | -0.63395 | 1 | 0.998956 | 3.02E-143 | purple | TPT1 |
| 5.74E-143 | 0.537286 | 0.997602 | 0.374837 | 1.14E-142 | purple | HDAC2 |
| 6.02E-143 | 0.617333 | 0.997602 | 0.484999 | 1.19E-142 | purple | SRI |
| 8.20E-143 | 0.539635 | 0.990408 | 0.377968 | 1.62E-142 | purple | AK2 |
| 1.26E-142 | -2.71418 | 0.889688 | 0.919286 | 2.49E-142 | purple | CD74 |
| 3.77E-142 | 0.552145 | 0.992806 | 0.408622 | 7.40E-142 | purple | TOMM5 |
| 1.14E-141 | 0.560725 | 0.863309 | 0.260534 | 2.24E-141 | purple | HIST1H1D |
| 2.16E-141 | -0.8022 | 1 | 0.996837 | 4.21E-141 | purple | EIF1 |
| 1.09E-139 | 0.502996 | 0.995204 | 0.385207 | 2.12E-139 | purple | MRPS15 |
| 8.41E-139 | 0.61963 | 0.997602 | 0.486597 | 1.63E-138 | purple | CCT3 |
| 2.16E-138 | 0.569021 | 0.995204 | 0.415243 | 4.17E-138 | purple | TCP1 |
| 1.30E-137 | 0.660329 | 1 | 0.917591 | 2.51E-137 | purple | NPM1 |
| 1.74E-137 | 0.569622 | 0.997602 | 0.459464 | 3.34E-137 | purple | NDUFC1 |
| 2.89E-136 | 0.509009 | 0.985612 | 0.387034 | 5.53E-136 | purple | PSMC3 |
| 5.43E-136 | 0.671143 | 1 | 0.581953 | 1.04E-135 | purple | TXN |
| 2.17E-134 | 0.593334 | 0.997602 | 0.498793 | 4.12E-134 | purple | RBX1 |
| 4.18E-134 | 0.587734 | 0.997602 | 0.635697 | 7.94E-134 | purple | TUFM |
| 5.57E-134 | 0.593243 | 1 | 0.626207 | 1.05E-133 | purple | ERH |
| 1.44E-132 | -0.87957 | 0.990408 | 0.904513 | 2.71E-132 | purple | COMMD6 |
| 1.45E-132 | 0.625601 | 1 | 0.695734 | 2.73E-132 | purple | ATP5F1B |
| 4.42E-129 | 0.50879 | 0.992806 | 0.414525 | 8.29E-129 | purple | PSMD6 |
| 6.08E-129 | -1.00992 | 1 | 0.999185 | 1.14E-128 | purple | MALAT1 |
| 1.15E-128 | -1.05782 | 0.961631 | 0.867728 | 2.15E-128 | purple | ARPC2 |
| 6.50E-128 | 0.529581 | 1 | 0.434027 | 1.21E-127 | purple | RNH1 |
| 8.44E-127 | -1.42025 | 0.064748 | 0.691625 | 1.56E-126 | purple | PKM |
| 1.85E-125 | 0.523372 | 1 | 0.471204 | 3.41E-125 | purple | NDUFA6 |
| 2.69E-124 | -1.14009 | 0.983213 | 0.926004 | 4.95E-124 | purple | LAPTM5 |
| 3.82E-124 | 0.505716 | 0.98801 | 0.416091 | 7.01E-124 | purple | TXNL1 |
| 3.63E-123 | 0.581293 | 1 | 0.45467 | 6.64E-123 | purple | YWHAE |
| 5.54E-123 | -1.32099 | 0.42446 | 0.777818 | 1.01E-122 | purple | GPSM3 |
| 8.71E-123 | 0.537362 | 1 | 0.688951 | 1.59E-122 | purple | SELENOH |
| 1.42E-122 | 0.572727 | 1 | 0.652328 | 2.58E-122 | purple | ATP5F1A |

| | | | | | | |
|---|---|---|---|---|---|---|
| 2.51E-122 | 0.50211 | 0.992806 | 0.417167 | 4.53E-122 | purple | ESD |
| 4.52E-122 | 0.554822 | 1 | 0.653698 | 8.15E-122 | purple | PHB2 |
| 2.89E-121 | 0.566926 | 0.997602 | 0.53959 | 5.21E-121 | purple | P4HB |
| 3.46E-121 | 0.524674 | 1 | 0.497522 | 6.21E-121 | purple | UQCRC1 |
| 4.19E-121 | 0.555596 | 1 | 0.60227 | 7.50E-121 | purple | SNRPB |
| 2.37E-120 | -1.05291 | 0.83693 | 0.841378 | 4.23E-120 | purple | GMFG |
| 7.10E-120 | 0.502809 | 0.992806 | 0.435494 | 1.26E-119 | purple | PSMB2 |
| 1.14E-119 | 0.536789 | 1 | 0.536721 | 2.02E-119 | purple | SNRPC |
| 1.16E-119 | 0.577973 | 0.992806 | 0.457246 | 2.06E-119 | purple | DYNLL1 |
| 2.46E-117 | -0.92215 | 1 | 0.980335 | 4.34E-117 | purple | UBC |
| 4.88E-117 | 0.517446 | 1 | 0.781829 | 8.59E-117 | purple | UQCRH |
| 6.78E-117 | -1.55556 | 0.352518 | 0.753359 | 1.19E-116 | purple | PTPRC |
| 4.97E-116 | 0.643114 | 1 | 0.677863 | 8.69E-116 | purple | HMGB2 |
| 3.27E-115 | 0.54951 | 1 | 0.528568 | 5.69E-115 | purple | CCT8 |
| 4.47E-115 | 0.553662 | 0.995204 | 0.543863 | 7.77E-115 | purple | NHP2 |
| 6.71E-115 | -1.21257 | 0.757794 | 0.840073 | 1.16E-114 | purple | LIMD2 |
| 7.24E-115 | 0.538013 | 0.997602 | 0.479096 | 1.25E-114 | purple | PPA1 |
| 1.34E-114 | -1.19454 | 0.997602 | 0.904872 | 2.32E-114 | purple | SARAF |
| 3.30E-114 | -1.63765 | 0.081535 | 0.665373 | 5.68E-114 | purple | HCST |
| 9.02E-114 | 0.517065 | 1 | 0.561179 | 1.55E-113 | purple | POMP |
| 2.59E-113 | 0.522942 | 1 | 0.481607 | 4.42E-113 | purple | NOP58 |
| 2.73E-110 | 0.51509 | 0.997602 | 0.525502 | 4.65E-110 | purple | RAD23A |
| 8.17E-109 | 0.601911 | 0.997602 | 0.482031 | 1.39E-108 | purple | IMPDH2 |
| 4.13E-108 | -2.07956 | 0.294964 | 0.713899 | 7.00E-108 | purple | ZFP36 |
| 7.40E-107 | -0.56352 | 1 | 0.98914 | 1.25E-106 | purple | EEF1D |
| 1.59E-106 | 0.512691 | 1 | 0.528437 | 2.69E-106 | purple | BANF1 |
| 1.96E-106 | 0.606363 | 1 | 0.867206 | 3.30E-106 | purple | HSP90AB1 |
| 8.29E-102 | 0.517483 | 1 | 0.678744 | 1.39E-101 | purple | PEBP1 |
| 4.22E-101 | 0.508088 | 1 | 0.601487 | 7.06E-101 | purple | FBL |
| 1.17E-100 | 0.549266 | 1 | 0.736434 | 1.95E-100 | purple | TPI1 |
| 3.49E-100 | -1.64 | 0.06235 | 0.61189 | 5.80E-100 | purple | CXCR4 |
| 5.01E-100 | 0.702428 | 1 | 0.858205 | 8.31E-100 | purple | HSPA8 |
| 3.32E-99 | 0.504282 | 1 | 0.51249 | 5.49E-99 | purple | RBBP4 |
| 2.08E-97 | -1.15238 | 0.110312 | 0.623206 | 3.43E-97 | purple | PSMB9 |
| 7.81E-97 | -0.55176 | 1 | 0.981574 | 1.29E-96 | purple | PFDN5 |
| 7.89E-94 | -0.94154 | 0.959233 | 0.854487 | 1.29E-93 | purple | RAC2 |
| 3.73E-93 | -2.01756 | 0.551559 | 0.780068 | 6.10E-93 | purple | JUN |
| 6.31E-92 | -1.01873 | 0.98801 | 0.893067 | 1.03E-91 | purple | MYL12A |
| 6.84E-91 | -1.49912 | 0.230216 | 0.64972 | 1.11E-90 | purple | ITGB2 |
| 9.85E-91 | -1.38554 | 0.064748 | 0.580127 | 1.60E-90 | purple | SELL |
| 1.46E-85 | -1.54256 | 0.959233 | 0.825757 | 2.36E-85 | purple | SRGN |

| | | | | | | |
|---|---|---|---|---|---|---|
| 7.47E-85 | 0.607333 | 1 | 0.824583 | 1.21E-84 | purple | MIF |
| 1.42E-74 | -2.11218 | 0.7506 | 0.771524 | 2.29E-74 | purple | JUNB |
| 7.37E-74 | -1.19827 | 0.071942 | 0.526024 | 1.19E-73 | purple | LITAF |
| 4.15E-73 | -0.64414 | 0.997602 | 0.902655 | 6.65E-73 | purple | DAZAP2 |
| 9.38E-73 | -1.078 | 0.685851 | 0.729324 | 1.50E-72 | purple | LCP1 |
| 1.30E-72 | -2.03107 | 0.17506 | 0.578333 | 2.08E-72 | purple | IFITM1 |
| 2.17E-72 | -0.97459 | 0.047962 | 0.503685 | 3.46E-72 | purple | ANXA6 |
| 1.15E-71 | -1.02459 | 0.458034 | 0.666612 | 1.83E-71 | purple | ICAM3 |
| 1.82E-71 | -1.56673 | 0.148681 | 0.555016 | 2.88E-71 | purple | AIF1 |
| 3.60E-70 | -1.32036 | 0.035971 | 0.488716 | 5.68E-70 | purple | CD69 |
| 4.65E-69 | -0.59834 | 0.980815 | 0.400176 | 7.32E-69 | purple | CST3 |
| 9.46E-69 | -0.53594 | 1 | 0.913547 | 1.49E-68 | purple | CCNI |
| 2.20E-67 | -1.65061 | 0.46283 | 0.655329 | 3.45E-67 | purple | HLA-DPB1 |
| 2.93E-66 | -1.31834 | 0.270983 | 0.589877 | 4.58E-66 | purple | EVL |
| 5.49E-65 | -0.84142 | 0.146283 | 0.529383 | 8.56E-65 | purple | CELF2 |
| 1.95E-64 | -1.23931 | 0.052758 | 0.473748 | 3.03E-64 | purple | TRBC2 |
| 5.41E-61 | -0.54379 | 1 | 0.915177 | 8.40E-61 | purple | CIRBP |
| 6.86E-61 | -1.05449 | 0.045564 | 0.453072 | 1.06E-60 | purple | CD48 |
| 7.91E-60 | -0.5351 | 1 | 0.913155 | 1.22E-59 | purple | ARPC3 |
| 8.74E-60 | -1.2421 | 0.127098 | 0.5015 | 1.35E-59 | purple | FYB1 |
| 9.68E-60 | -0.93931 | 0.647482 | 0.685984 | 1.49E-59 | purple | 6-Sep |
| 1.29E-58 | -0.8464 | 0.261391 | 0.563136 | 1.98E-58 | purple | WAS |
| 2.06E-57 | -1.45231 | 0.549161 | 0.667004 | 3.14E-57 | purple | KLF6 |
| 2.11E-55 | -0.86485 | 0.033573 | 0.420428 | 3.21E-55 | purple | CYTIP |
| 5.40E-55 | -0.51802 | 0.992806 | 0.827452 | 8.21E-55 | purple | ATP6V1G1 |
| 1.00E-54 | -1.03922 | 0.352518 | 0.58466 | 1.52E-54 | purple | 9-Sep |
| 7.10E-54 | -0.57535 | 1 | 0.994521 | 1.07E-53 | purple | RPS27 |
| 5.00E-53 | -1.09708 | 0.242206 | 0.531829 | 7.55E-53 | purple | BTG1 |
| 1.59E-52 | -1.5742 | 0.582734 | 0.651807 | 2.40E-52 | purple | HLA-DPA1 |
| 3.86E-52 | -1.85995 | 0.53717 | 0.642447 | 5.81E-52 | purple | HLA-DRB1 |
| 9.31E-52 | -1.01239 | 0.261391 | 0.535514 | 1.40E-51 | purple | RARRES3 |
| 1.16E-51 | -0.73135 | 0.995204 | 0.779872 | 1.74E-51 | purple | ZFAS1 |
| 1.71E-51 | -1.12787 | 0.115108 | 0.459073 | 2.55E-51 | purple | AHNAK |
| 5.86E-51 | -0.94292 | 0.251799 | 0.528307 | 8.72E-51 | purple | CYBA |
| 1.10E-50 | -1.08075 | 0.052758 | 0.410612 | 1.63E-50 | purple | SLC2A3 |
| 1.23E-50 | -0.72309 | 0.100719 | 0.446419 | 1.83E-50 | purple | RASGRP2 |
| 1.31E-50 | -0.93927 | 0.086331 | 0.433896 | 1.94E-50 | purple | HLA-F |
| 2.02E-50 | -1.81586 | 0.047962 | 0.406144 | 2.97E-50 | purple | CD3E |
| 2.17E-50 | -0.86276 | 0.779376 | 0.710181 | 3.20E-50 | purple | HCLS1 |
| 4.58E-50 | -1.61541 | 0.043165 | 0.401285 | 6.72E-50 | purple | LTB |
| 4.64E-50 | -1.26992 | 0.7506 | 0.712725 | 6.79E-50 | purple | IER2 |

| | | | | | | |
|---|---|---|---|---|---|---|
| 1.03E-49 | -1.74532 | 0.453237 | 0.610064 | 1.51E-49 | purple | S100A10 |
| 2.48E-49 | -0.73109 | 0.968825 | 0.77997 | 3.61E-49 | purple | C12orf57 |
| 2.52E-49 | -0.57968 | 0.997602 | 0.876565 | 3.66E-49 | purple | YWHAZ |
| 4.30E-49 | -0.77229 | 0.031175 | 0.386316 | 6.24E-49 | purple | C1orf162 |
| 6.99E-49 | -1.70846 | 0.640288 | 0.719149 | 1.01E-48 | purple | FOS |
| 6.55E-48 | -1.02534 | 0.035971 | 0.385142 | 9.45E-48 | purple | LST1 |
| 7.24E-48 | -0.73 | 0.040767 | 0.38736 | 1.04E-47 | purple | PPP1R18 |
| 1.35E-47 | -1.00259 | 0.184652 | 0.487966 | 1.94E-47 | purple | IFITM3 |
| 1.65E-47 | -1.18097 | 0.155875 | 0.468986 | 2.36E-47 | purple | CTSS |
| 3.13E-47 | -1.65374 | 0.038369 | 0.384262 | 4.48E-47 | purple | LGALS1 |
| 4.81E-47 | -0.5005 | 1 | 0.947071 | 6.86E-47 | purple | DDX5 |
| 8.74E-46 | -1.10391 | 0.035971 | 0.37275 | 1.24E-45 | purple | HLA-DQB1 |
| 1.80E-45 | -2.06893 | 0.06235 | 0.39261 | 2.55E-45 | purple | IL32 |
| 5.48E-45 | -0.72763 | 0.021583 | 0.355531 | 7.77E-45 | purple | IL16 |
| 6.67E-45 | -0.80721 | 0.508393 | 0.607781 | 9.42E-45 | purple | PSMB8 |
| 9.82E-45 | -0.64365 | 0.026379 | 0.358401 | 1.39E-44 | purple | SORL1 |
| 1.66E-44 | -1.27782 | 0.115108 | 0.425841 | 2.34E-44 | purple | TNFAIP3 |
| 4.79E-44 | -0.7264 | 0.074341 | 0.394273 | 6.72E-44 | purple | CYFIP2 |
| 1.40E-43 | -0.58964 | 0.047962 | 0.372 | 1.97E-43 | purple | PARVG |
| 1.46E-43 | -0.74156 | 0.071942 | 0.391371 | 2.03E-43 | purple | TPM4 |
| 1.73E-43 | -0.84679 | 0.311751 | 0.533981 | 2.41E-43 | purple | CD53 |
| 9.83E-43 | -1.10892 | 0.143885 | 0.436212 | 1.37E-42 | purple | GADD45B |
| 1.15E-42 | -0.67631 | 0.98801 | 0.804592 | 1.60E-42 | purple | GABARAP |
| 4.14E-42 | -0.68212 | 0.045564 | 0.362216 | 5.74E-42 | purple | MYO1G |
| 4.60E-42 | -0.79324 | 0.033573 | 0.350313 | 6.35E-42 | purple | PLEK |
| 2.25E-41 | -0.67166 | 0.021583 | 0.335507 | 3.11E-41 | purple | LAT2 |
| 5.11E-41 | -0.64839 | 0.023981 | 0.336616 | 7.03E-41 | purple | BIN1 |
| 9.80E-41 | -0.85524 | 0.153477 | 0.431907 | 1.34E-40 | purple | TBC1D10C |
| 2.17E-40 | -0.68011 | 0.038369 | 0.345584 | 2.97E-40 | purple | GLIPR1 |
| 3.51E-39 | -0.64471 | 0.019185 | 0.3208 | 4.80E-39 | purple | CAPG |
| 4.74E-39 | -1.52743 | 0.477218 | 0.58329 | 6.46E-39 | purple | HLA-DRB5 |
| 1.01E-38 | -0.55532 | 1 | 0.793341 | 1.37E-38 | purple | CLIC1 |
| 9.66E-38 | -2.19381 | 0.568345 | 0.602596 | 1.31E-37 | purple | HLA-DRA |
| 1.03E-37 | -0.63209 | 0.016787 | 0.310168 | 1.39E-37 | purple | PDE4B |
| 1.45E-37 | -1.03868 | 0.151079 | 0.415569 | 1.96E-37 | purple | DDIT4 |
| 2.18E-37 | -1.94752 | 0.035971 | 0.329246 | 2.94E-37 | purple | TYROBP |
| 3.18E-37 | -0.77026 | 0.103118 | 0.381066 | 4.27E-37 | purple | P2RY8 |
| 4.08E-37 | -0.889 | 0.431655 | 0.556222 | 5.48E-37 | purple | STK17B |
| 5.82E-37 | -0.82321 | 0.645084 | 0.250294 | 7.80E-37 | purple | IGLL1 |
| 6.28E-37 | -0.82569 | 0.047962 | 0.334366 | 8.40E-37 | purple | ISG20 |
| 9.87E-37 | -0.82022 | 0.021583 | 0.308766 | 1.32E-36 | purple | ARL4C |

| | | | | | | |
|---|---|---|---|---|---|---|
| 3.53E-36 | -0.53349 | 0.023981 | 0.308016 | 4.70E-36 | purple | PECAM1 |
| 3.86E-36 | -0.96857 | 0.040767 | 0.323506 | 5.13E-36 | purple | LCK |
| 4.04E-36 | -1.11052 | 0.040767 | 0.323213 | 5.36E-36 | purple | CD3D |
| 4.20E-36 | -0.84988 | 0.28777 | 0.48614 | 5.55E-36 | purple | PYCARD |
| 6.77E-36 | -1.40011 | 0.052758 | 0.333942 | 8.94E-36 | purple | AREG |
| 9.68E-36 | -0.60576 | 0.038369 | 0.319039 | 1.27E-35 | purple | TRIM22 |
| 9.90E-36 | -0.84177 | 0.016787 | 0.297776 | 1.30E-35 | purple | FCMR |
| 1.56E-35 | -1.02836 | 0.71223 | 0.687027 | 2.05E-35 | purple | ANXA1 |
| 1.69E-35 | -0.61436 | 0.057554 | 0.335247 | 2.21E-35 | purple | SASH3 |
| 1.85E-35 | -0.61088 | 0.059952 | 0.337692 | 2.42E-35 | purple | ARHGAP30 |
| 2.04E-35 | -0.67241 | 0.059952 | 0.337138 | 2.66E-35 | purple | RCSD1 |
| 2.31E-35 | -0.62927 | 0.095923 | 0.365869 | 3.00E-35 | purple | VAMP5 |
| 5.71E-35 | -2.25261 | 0.028777 | 0.309157 | 7.41E-35 | purple | NKG7 |
| 1.15E-34 | -0.52496 | 0.057554 | 0.330029 | 1.49E-34 | purple | FLI1 |
| 1.63E-34 | -0.50567 | 0.045564 | 0.319691 | 2.10E-34 | purple | ARMH1 |
| 1.76E-34 | -0.50147 | 0.045564 | 0.31943 | 2.27E-34 | purple | FNBP1 |
| 4.64E-34 | -0.89188 | 0.035971 | 0.307168 | 5.98E-34 | purple | SPOCK2 |
| 8.45E-34 | -1.30889 | 0.083933 | 0.350411 | 1.09E-33 | purple | S100A11 |
| 1.12E-33 | -0.8731 | 0.338129 | 0.499772 | 1.43E-33 | purple | PLAC8 |
| 1.81E-33 | -0.73968 | 0.213429 | 0.433049 | 2.31E-33 | purple | IDS |
| 2.16E-33 | -0.67007 | 0.01199 | 0.279024 | 2.76E-33 | purple | CSF3R |
| 4.63E-33 | -0.78012 | 0.151079 | 0.394045 | 5.90E-33 | purple | TRAF3IP3 |
| 9.20E-33 | -0.75066 | 0.673861 | 0.650241 | 1.17E-32 | purple | LY6E |
| 2.27E-32 | -0.73929 | 0.004796 | 0.263762 | 2.88E-32 | purple | SMIM24 |
| 2.34E-32 | -0.76602 | 0.019185 | 0.279481 | 2.96E-32 | purple | ZAP70 |
| 4.34E-32 | -0.68923 | 0.194245 | 0.417786 | 5.49E-32 | purple | CLEC2B |
| 1.32E-31 | -0.7495 | 0.009592 | 0.264349 | 1.67E-31 | purple | HOPX |
| 4.16E-31 | -0.89663 | 0.206235 | 0.42333 | 5.24E-31 | purple | BTG2 |
| 7.53E-31 | -0.84731 | 0.254197 | 0.444626 | 9.45E-31 | purple | VSIR |
| 1.04E-30 | -0.76112 | 0.014388 | 0.263762 | 1.30E-30 | purple | ALOX5AP |
| 1.46E-30 | -0.90941 | 0.007194 | 0.254696 | 1.82E-30 | purple | SPINK2 |
| 1.58E-30 | -0.96127 | 0.779376 | 0.639969 | 1.97E-30 | purple | TUBA1A |
| 1.63E-30 | -0.83348 | 0.254197 | 0.442538 | 2.03E-30 | purple | 1-Sep |
| 2.51E-30 | -0.72972 | 0.026379 | 0.27472 | 3.12E-30 | purple | IL10RA |
| 3.70E-30 | -0.66507 | 0.17506 | 0.396556 | 4.60E-30 | purple | ARHGAP45 |
| 1.17E-29 | -1.37002 | 0.047962 | 0.293373 | 1.45E-29 | purple | IL7R |
| 3.23E-29 | -0.8803 | 0.069544 | 0.310657 | 3.99E-29 | purple | HLA-DQA1 |
| 4.92E-29 | -0.86201 | 0.016787 | 0.25574 | 6.07E-29 | purple | GZMM |
| 5.03E-29 | -0.65322 | 0.045564 | 0.285742 | 6.19E-29 | purple | CRIP1 |
| 2.14E-28 | -0.65286 | 0.916067 | 0.738456 | 2.63E-28 | purple | ITM2B |
| 3.74E-28 | -0.59705 | 0.028777 | 0.263338 | 4.59E-28 | purple | MYO1F |

| | | | | | | |
|---|---|---|---|---|---|---|
| 4.99E-28 | -0.54826 | 0.160671 | 0.373859 | 6.11E-28 | purple | KCNAB2 |
| 5.81E-28 | -0.76457 | 0.729017 | 0.628033 | 7.09E-28 | purple | RNASET2 |
| 9.14E-28 | -0.77666 | 0.023981 | 0.254892 | 1.11E-27 | purple | CD3G |
| 1.23E-27 | -0.67599 | 0.227818 | 0.414949 | 1.50E-27 | purple | ARHGAP9 |
| 1.26E-27 | -1.28916 | 0.028777 | 0.260664 | 1.53E-27 | purple | FCER1G |
| 8.64E-27 | -0.73406 | 0.167866 | 0.372065 | 1.05E-26 | purple | RNF213 |
| 1.07E-26 | -0.95056 | 0.153477 | 0.362379 | 1.30E-26 | purple | NR4A2 |
| 1.24E-26 | -0.74557 | 0.038369 | 0.262914 | 1.50E-26 | purple | KLF2 |
| 1.69E-26 | -0.93302 | 0.477218 | 0.527883 | 2.03E-26 | purple | CCND3 |
| 1.96E-26 | -0.81161 | 0.033573 | 0.25649 | 2.36E-26 | purple | NCF1 |
| 4.01E-26 | -0.8259 | 0.100719 | 0.316723 | 4.81E-26 | purple | SAMHD1 |
| 1.55E-25 | -0.55721 | 0.06235 | 0.280296 | 1.86E-25 | purple | ANKRD44 |
| 2.88E-25 | -0.67453 | 0.261391 | 0.422319 | 3.44E-25 | purple | AKAP13 |
| 6.79E-25 | -0.63013 | 0.057554 | 0.270839 | 8.09E-25 | purple | PRKCH |
| 1.18E-24 | -0.59366 | 0.045564 | 0.258381 | 1.41E-24 | purple | PTPRE |
| 3.17E-24 | -1.06574 | 0.968825 | 0.791254 | 3.76E-24 | purple | TXNIP |
| 4.66E-24 | -0.58817 | 0.182254 | 0.36626 | 5.53E-24 | purple | SYTL1 |
| 5.26E-24 | -0.78795 | 0.088729 | 0.294156 | 6.23E-24 | purple | PIK3IP1 |
| 6.01E-24 | -0.51596 | 0.076739 | 0.283199 | 7.10E-24 | purple | PRKCB |
| 1.33E-23 | -0.79921 | 0.561151 | 0.547548 | 1.56E-23 | purple | STK4 |
| 3.06E-23 | -0.77493 | 0.786571 | 0.638501 | 3.60E-23 | purple | CNN2 |
| 1.02E-22 | -0.60009 | 0.103118 | 0.298656 | 1.20E-22 | purple | AKNA |
| 1.62E-22 | -1.11394 | 0.796163 | 0.607292 | 1.90E-22 | purple | SERPINB1 |
| 2.53E-22 | -0.72457 | 0.848921 | 0.65047 | 2.96E-22 | purple | CD44 |
| 8.27E-22 | -0.5752 | 0.088729 | 0.281861 | 9.66E-22 | purple | CD34 |
| 2.56E-21 | -0.72081 | 0.42446 | 0.48689 | 2.98E-21 | purple | WIPF1 |
| 3.08E-21 | -0.61547 | 0.992806 | 0.883838 | 3.58E-21 | purple | CALM1 |
| 4.32E-21 | -1.32953 | 0.280576 | 0.411655 | 5.01E-21 | purple | CD7 |
| 1.17E-20 | -0.73446 | 0.568345 | 0.536297 | 1.35E-20 | purple | ACAP1 |
| 2.49E-20 | -0.76383 | 0.441247 | 0.485586 | 2.89E-20 | purple | BIN2 |
| 3.76E-19 | -0.50905 | 0.079137 | 0.253816 | 4.34E-19 | purple | PDE4D |
| 7.64E-19 | -0.56699 | 0.251799 | 0.379207 | 8.81E-19 | purple | RABAC1 |
| 1.47E-18 | -0.52874 | 0.976019 | 0.723226 | 1.70E-18 | purple | ENO1 |
| 3.34E-18 | -0.599 | 0.115108 | 0.279807 | 3.84E-18 | purple | MGAT4A |
| 5.17E-18 | -0.65514 | 0.422062 | 0.468465 | 5.93E-18 | purple | PTPN6 |
| 8.10E-18 | -0.56738 | 0.860911 | 0.652068 | 9.27E-18 | purple | MSN |
| 1.62E-17 | -0.52517 | 0.208633 | 0.3464 | 1.85E-17 | purple | HSH2D |
| 1.73E-17 | -0.7325 | 0.330935 | 0.417428 | 1.98E-17 | purple | RIPOR2 |
| 6.39E-17 | -0.5348 | 0.997602 | 0.807657 | 7.27E-17 | purple | CD99 |
| 9.89E-17 | -0.79802 | 0.242206 | 0.361173 | 1.12E-16 | purple | ITM2C |
| 1.34E-16 | -0.57987 | 0.196643 | 0.332246 | 1.51E-16 | purple | TMEM243 |

| | | | | | | |
|---|---|---|---|---|---|---|
| 1.41E-16 | -0.73576 | 0.517986 | 0.498891 | 1.60E-16 | purple | ARRB2 |
| 1.64E-16 | -0.75979 | 0.64988 | 0.550678 | 1.85E-16 | purple | ARHGEF1 |
| 2.52E-16 | -0.59225 | 0.179856 | 0.318452 | 2.84E-16 | purple | RNF125 |
| 2.65E-16 | -1.48588 | 0.151079 | 0.298722 | 2.98E-16 | purple | CST7 |
| 2.80E-16 | -0.62059 | 0.848921 | 0.627902 | 3.15E-16 | purple | CAP1 |
| 1.84E-15 | -0.51209 | 0.980815 | 0.66446 | 2.07E-15 | purple | VAMP8 |
| 1.91E-15 | -0.66396 | 0.11271 | 0.25812 | 2.13E-15 | purple | FLT3LG |
| 2.46E-15 | -2.45324 | 0.103118 | 0.252968 | 2.75E-15 | purple | LYZ |
| 2.56E-15 | -0.56951 | 0.11271 | 0.257305 | 2.86E-15 | purple | ABLIM1 |
| 1.27E-14 | -0.60303 | 0.628297 | 0.53721 | 1.41E-14 | purple | TMEM123 |
| 1.46E-14 | -0.73677 | 0.41247 | 0.440093 | 1.62E-14 | purple | PIK3R1 |
| 1.54E-14 | -0.60925 | 0.280576 | 0.37125 | 1.71E-14 | purple | PBXIP1 |
| 1.74E-14 | -0.57355 | 0.290168 | 0.376728 | 1.93E-14 | purple | DDAH2 |
| 3.37E-14 | -0.87046 | 0.676259 | 0.553026 | 3.73E-14 | purple | LEPROTL1 |
| 1.29E-13 | -0.74795 | 0.690647 | 0.555309 | 1.43E-13 | purple | YPEL5 |
| 2.89E-13 | -0.61729 | 0.839329 | 0.605922 | 3.18E-13 | purple | DBI |
| 4.26E-13 | -0.50669 | 0.376499 | 0.415112 | 4.69E-13 | purple | TWF2 |
| 4.43E-13 | -0.61531 | 0.191847 | 0.304037 | 4.86E-13 | purple | TAGAP |
| 5.96E-13 | -0.79273 | 0.419664 | 0.444919 | 6.53E-13 | purple | IRF1 |
| 6.59E-13 | -0.54302 | 0.906475 | 0.649198 | 7.21E-13 | purple | KMT2E |
| 7.35E-13 | -0.69193 | 0.815348 | 0.611825 | 8.02E-13 | purple | MYH9 |
| 1.00E-12 | -0.58453 | 0.839329 | 0.617043 | 1.09E-12 | purple | TAPBP |
| 1.10E-12 | -0.58802 | 0.357314 | 0.406829 | 1.20E-12 | purple | LCP2 |
| 1.50E-12 | -0.51389 | 0.925659 | 0.641762 | 1.63E-12 | purple | ARHGDIA |
| 3.36E-12 | -0.58744 | 0.781775 | 0.582051 | 3.64E-12 | purple | ARPC5 |
| 3.89E-11 | -0.64149 | 0.390887 | 0.413188 | 4.21E-11 | purple | RHOH |
| 4.22E-11 | -0.53354 | 0.244604 | 0.325365 | 4.56E-11 | purple | RHOF |
| 6.05E-11 | -0.57758 | 0.580336 | 0.497913 | 6.53E-11 | purple | ARHGAP4 |
| 1.03E-10 | -0.85717 | 0.601918 | 0.497522 | 1.11E-10 | purple | HLA-DMA |
| 5.55E-10 | -0.99242 | 0.707434 | 0.561897 | 5.97E-10 | purple | CTSW |
| 9.16E-10 | -0.60275 | 0.436451 | 0.424406 | 9.82E-10 | purple | VAMP2 |
| 9.75E-10 | -0.75429 | 0.223022 | 0.30182 | 1.04E-09 | purple | ANXA2 |
| 1.73E-09 | -0.69343 | 0.681055 | 0.522208 | 1.85E-09 | purple | SLC9A3R1 |
| 4.40E-09 | -0.60437 | 0.268585 | 0.327811 | 4.69E-09 | purple | ANXA5 |
| 6.96E-09 | -0.82363 | 0.357314 | 0.373272 | 7.42E-09 | purple | MAP3K8 |
| 9.79E-09 | -0.80634 | 0.681055 | 0.535971 | 1.04E-08 | purple | FLNA |
| 1.40E-08 | -0.59437 | 0.434053 | 0.410644 | 1.48E-08 | purple | TNFRSF14 |
| 1.72E-08 | -0.73576 | 0.434053 | 0.428874 | 1.83E-08 | purple | FOSB |
| 4.00E-08 | -0.52314 | 0.477218 | 0.425874 | 4.24E-08 | purple | MAX |
| 4.40E-08 | -0.59953 | 0.788969 | 0.547776 | 4.64E-08 | purple | BST2 |
| 4.59E-08 | -0.92245 | 0.438849 | 0.422939 | 4.84E-08 | purple | DUSP1 |

| | | | | | | |
|---|---|---|---|---|---|---|
| 1.04E-07 | -0.57784 | 0.285372 | 0.322463 | 1.09E-07 | purple | CYTH1 |
| 1.07E-07 | -0.53346 | 0.297362 | 0.333779 | 1.13E-07 | purple | SLC38A1 |
| 2.31E-07 | -0.55439 | 0.429257 | 0.39998 | 2.43E-07 | purple | POLD4 |
| 3.46E-07 | -0.60838 | 0.292566 | 0.323278 | 3.62E-07 | purple | ADGRE5 |
| 4.55E-07 | -0.51394 | 0.503597 | 0.435886 | 4.75E-07 | purple | CMTM6 |
| 4.58E-07 | -0.51799 | 0.354916 | 0.359673 | 4.78E-07 | purple | TMC8 |
| 1.69E-06 | -0.51936 | 0.863309 | 0.586942 | 1.76E-06 | purple | CCNL1 |
| 1.88E-06 | -0.65865 | 0.582734 | 0.46204 | 1.95E-06 | purple | HIF1A |
| 2.66E-06 | -0.52596 | 0.527578 | 0.292786 | 2.76E-06 | purple | NAMPT |
| 1.26E-05 | -0.52463 | 0.606715 | 0.477726 | 1.31E-05 | purple | PNRC1 |
| 2.17E-05 | -0.77693 | 0.64988 | 0.479781 | 2.25E-05 | purple | MCL1 |
| 5.00E-05 | -0.57917 | 0.470024 | 0.402785 | 5.17E-05 | purple | ADD3 |
| 9.76E-05 | -0.54385 | 0.709832 | 0.501533 | 0.000101 | purple | LRRFIP1 |
| 0.000125 | -0.54853 | 0.647482 | 0.479455 | 0.000129 | purple | PPP1R2 |
| 0.000152 | -0.72613 | 0.306954 | 0.306875 | 0.000156 | purple | ID2 |
| 0.00019 | -0.58729 | 0.247002 | 0.267023 | 0.000195 | purple | LAT |
| 0.000912 | -0.53581 | 0.386091 | 0.345976 | 0.000935 | purple | AC007952.4 |
| 0.001896 | -0.82812 | 0.872902 | 0.603411 | 0.00194 | purple | NEAT1 |
| 0.002113 | -0.56984 | 0.647482 | 0.466345 | 0.002158 | purple | REL |
| 0.007977 | -0.60622 | 0.553957 | 0.418634 | 0.008134 | purple | IQGAP1 |
| 0.009329 | -0.51244 | 0.772182 | 0.515849 | 0.009498 | purple | FOXP1 |
| 0.010527 | -0.53748 | 0.609113 | 0.365575 | 0.0107 | purple | GRN |
| 0.015377 | -1.071 | 0.935252 | 0.618836 | 0.015603 | purple | SAT1 |
| 0.015975 | -0.6747 | 0.7506 | 0.492467 | 0.016184 | purple | AES |
| 0 | 2.259244 | 0.993002 | 0.215196 | 0 | yellow | LYZ |
| 0 | 2.080966 | 0.96501 | 0.090348 | 0 | yellow | FCN1 |
| 0 | 2.044011 | 0.975507 | 0.116721 | 0 | yellow | S100A9 |
| 0 | 2.025906 | 0.993002 | 0.379772 | 0 | yellow | CST3 |
| 0 | 1.936294 | 0.976907 | 0.113146 | 0 | yellow | S100A8 |
| 0 | 1.897291 | 0.9986 | 0.292864 | 0 | yellow | TYROBP |
| 0 | 1.745022 | 0.825752 | 0.074093 | 0 | yellow | SERPINA1 |
| 0 | 1.716375 | 0.996501 | 0.815459 | 0 | yellow | S100A6 |
| 0 | 1.665832 | 0.955913 | 0.607042 | 0 | yellow | SAT1 |
| 0 | 1.66248 | 0.895731 | 0.386483 | 0 | yellow | TIMP1 |
| 0 | 1.654976 | 0.949615 | 0.317786 | 0 | yellow | S100A11 |
| 0 | 1.612957 | 0.841847 | 0.078646 | 0 | yellow | VCAN |
| 0 | 1.608166 | 0.953114 | 0.351983 | 0 | yellow | LGALS1 |
| 0 | 1.599767 | 0.937719 | 0.224774 | 0 | yellow | FCER1G |
| 0 | 1.582227 | 0.987404 | 0.589674 | 0 | yellow | S100A10 |
| 0 | 1.566111 | 1 | 0.993828 | 0 | yellow | FTL |
| 0 | 1.550698 | 0.827852 | 0.11092 | 0 | yellow | LGALS3 |

| | | | | | | |
|---|---|---|---|---|---|---|
| 0 | 1.524895 | 0.892232 | 0.444186 | 0 | yellow | CTSS |
| 0 | 1.52399 | 0.811057 | 0.133718 | 0 | yellow | CD68 |
| 0 | 1.508412 | 0.857943 | 0.11436 | 0 | yellow | CSTA |
| 0 | 1.506251 | 0.745276 | 0.063975 | 0 | yellow | CD14 |
| 0 | 1.489057 | 0.960112 | 0.590011 | 0 | yellow | NEAT1 |
| 0 | 1.47802 | 0.943317 | 0.681168 | 0 | yellow | PSAP |
| 0 | 1.461079 | 0.920224 | 0.354445 | 0 | yellow | LST1 |
| 0 | 1.455244 | 0.968509 | 0.529374 | 0 | yellow | AIF1 |
| 0 | 1.452401 | 0.792162 | 0.119216 | 0 | yellow | TYMP |
| 0 | 1.439969 | 0.849545 | 0.164643 | 0 | yellow | COTL1 |
| 0 | 1.422204 | 0.53394 | 0.063706 | 0 | yellow | G0S2 |
| 0 | 1.420074 | 0.995101 | 0.80288 | 0 | yellow | S100A4 |
| 0 | 1.412522 | 0.70049 | 0.057365 | 0 | yellow | TMEM176B |
| 0 | 1.398331 | 0.783065 | 0.142419 | 0 | yellow | CFD |
| 0 | 1.391188 | 0.892932 | 0.400513 | 0 | yellow | DUSP1 |
| 0 | 1.390727 | 0.770469 | 0.132234 | 0 | yellow | CFP |
| 0 | 1.383207 | 0.804759 | 0.178909 | 0 | yellow | PLAUR |
| 0 | 1.381328 | 0.961512 | 0.6724 | 0 | yellow | TSPO |
| 0 | 1.352444 | 0.754374 | 0.075206 | 0 | yellow | S100A12 |
| 0 | 1.348817 | 0.836949 | 0.346284 | 0 | yellow | GRN |
| 0 | 1.338102 | 0.69839 | 0.072305 | 0 | yellow | MS4A6A |
| 0 | 1.335237 | 0.452064 | 0.038749 | 0 | yellow | APOBEC3A |
| 0 | 1.319738 | 0.9993 | 0.996358 | 0 | yellow | FTH1 |
| 0 | 1.281248 | 0.955913 | 0.62586 | 0 | yellow | HLA-DRB1 |
| 0 | 1.253646 | 0.771169 | 0.273034 | 0 | yellow | NAMPT |
| 0 | 1.244562 | 0.915325 | 0.565797 | 0 | yellow | HLA-DRB5 |
| 0 | 1.24165 | 0.730581 | 0.216714 | 0 | yellow | ATP2B1-AS1 |
| 0 | 1.218637 | 0.646606 | 0.060873 | 0 | yellow | CLEC7A |
| 0 | 1.211638 | 0.622113 | 0.186193 | 0 | yellow | CPVL |
| 0 | 1.176892 | 0.806158 | 0.276406 | 0 | yellow | ANXA2 |
| 0 | 1.17656 | 0.850945 | 0.532578 | 0 | yellow | NPC2 |
| 0 | 1.157352 | 0.53464 | 0.042358 | 0 | yellow | MAFB |
| 0 | 1.155181 | 0.76627 | 0.308546 | 0 | yellow | FCGRT |
| 0 | 1.127173 | 0.424073 | 0.027148 | 0 | yellow | LGALS2 |
| 0 | 1.113794 | 0.490553 | 0.075003 | 0 | yellow | MNDA |
| 0 | 1.088624 | 0.955213 | 0.585121 | 0 | yellow | HLA-DRA |
| 0 | 1.074533 | 0.588523 | 0.082018 | 0 | yellow | KLF4 |
| 0 | 1.068482 | 0.853744 | 0.464151 | 0 | yellow | MCL1 |
| 0 | 1.065584 | 0.321903 | 0.038817 | 0 | yellow | RETN |
| 0 | 1.059464 | 0.547936 | 0.03912 | 0 | yellow | TMEM176A |
| 0 | 1.052317 | 0.982505 | 0.82008 | 0 | yellow | SRGN |

| | | | | | | |
|---|---|---|---|---|---|---|
| 0 | 1.04913 | 0.644507 | 0.11318 | 0 | yellow | CTSH |
| 0 | 1.045348 | 0.601819 | 0.094361 | 0 | yellow | SLC11A1 |
| 0 | 1.042936 | 0.738279 | 0.307197 | 0 | yellow | ANXA5 |
| 0 | 1.040795 | 0.512246 | 0.062424 | 0 | yellow | FGL2 |
| 0 | 1.032983 | 0.63471 | 0.172332 | 0 | yellow | STXBP2 |
| 0 | 1.025284 | 0.596221 | 0.085256 | 0 | yellow | LILRB2 |
| 0 | 1.023902 | 0.547936 | 0.063132 | 0 | yellow | IGSF6 |
| 0 | 1.023488 | 0.474458 | 0.036052 | 0 | yellow | CD300E |
| 0 | 1.020773 | 0.961512 | 0.720558 | 0 | yellow | EMP3 |
| 0 | 1.01723 | 0.650805 | 0.228619 | 0 | yellow | SPI1 |
| 0 | 1.012024 | 0.948216 | 0.80025 | 0 | yellow | GABARAP |
| 0 | 1.010652 | 0.581526 | 0.142048 | 0 | yellow | CTSD |
| 0 | 1.00771 | 0.538139 | 0.051228 | 0 | yellow | IFI30 |
| 0 | 0.993005 | 0.771868 | 0.449885 | 0 | yellow | ASAH1 |
| 0 | 0.992857 | 0.522043 | 0.054431 | 0 | yellow | TNFAIP2 |
| 0 | 0.990497 | 0.395381 | 0.033185 | 0 | yellow | HMOX1 |
| 0 | 0.987321 | 0.507348 | 0.049575 | 0 | yellow | C5AR1 |
| 0 | 0.97729 | 0.575927 | 0.101174 | 0 | yellow | CYBB |
| 0 | 0.971387 | 0.829251 | 0.466815 | 0 | yellow | PYCARD |
| 0 | 0.9665 | 0.716585 | 0.303959 | 0 | yellow | GNAI2 |
| 0 | 0.966024 | 0.527642 | 0.051599 | 0 | yellow | PILRA |
| 0 | 0.964494 | 0.713086 | 0.340753 | 0 | yellow | CTSB |
| 0 | 0.952621 | 0.50035 | 0.096756 | 0 | yellow | CDKN1A |
| 0 | 0.943658 | 0.53464 | 0.06519 | 0 | yellow | NCF2 |
| 0 | 0.928604 | 0.515045 | 0.054027 | 0 | yellow | LILRA5 |
| 0 | 0.920923 | 0.496151 | 0.049744 | 0 | yellow | FPR1 |
| 0 | 0.914542 | 0.679496 | 0.255969 | 0 | yellow | CSF3R |
| 0 | 0.913014 | 0.76697 | 0.421725 | 0 | yellow | APLP2 |
| 0 | 0.911571 | 0.595521 | 0.201032 | 0 | yellow | RGS2 |
| 0 | 0.908819 | 0.619314 | 0.205989 | 0 | yellow | FMNL1 |
| 0 | 0.908521 | 0.596221 | 0.218839 | 0 | yellow | NINJ1 |
| 0 | 0.907943 | 0.578027 | 0.132504 | 0 | yellow | TNFRSF1B |
| 0 | 0.907092 | 0.847446 | 0.574059 | 0 | yellow | ATP6V0B |
| 0 | 0.90668 | 0.785164 | 0.425536 | 0 | yellow | VSIR |
| 0 | 0.904 | 0.480056 | 0.066572 | 0 | yellow | CEBPD |
| 0 | 0.899209 | 0.475857 | 0.042324 | 0 | yellow | LRP1 |
| 0 | 0.897983 | 0.571029 | 0.117463 | 0 | yellow | LY86 |
| 0 | 0.897318 | 0.440868 | 0.076926 | 0 | yellow | SGK1 |
| 0 | 0.894609 | 0.496151 | 0.063099 | 0 | yellow | TREM1 |
| 0 | 0.893558 | 0.353394 | 0.043336 | 0 | yellow | THBS1 |
| 0 | 0.892755 | 0.489153 | 0.043201 | 0 | yellow | SLC7A7 |

| | | | | | | |
|---|---|---|---|---|---|---|
| 0 | 0.885273 | 0.981805 | 0.705382 | 0 | yellow | FOS |
| 0 | 0.875128 | 0.993702 | 0.967658 | 0 | yellow | SH3BGRL3 |
| 0 | 0.863041 | 0.396081 | 0.044584 | 0 | yellow | AC020656.1 |
| 0 | 0.861546 | 0.981106 | 0.919263 | 0 | yellow | OAZ1 |
| 0 | 0.848032 | 0.602519 | 0.137225 | 0 | yellow | FGR |
| 0 | 0.847538 | 0.473058 | 0.11291 | 0 | yellow | CD36 |
| 0 | 0.837358 | 0.627712 | 0.237589 | 0 | yellow | PTPRE |
| 0 | 0.831897 | 1 | 0.998685 | 0 | yellow | TMSB10 |
| 0 | 0.830492 | 0.490553 | 0.131357 | 0 | yellow | ZNF385A |
| 0 | 0.822931 | 0.505248 | 0.124477 | 0 | yellow | C19orf38 |
| 0 | 0.816522 | 0.990203 | 0.903278 | 0 | yellow | VIM |
| 0 | 0.815796 | 0.378586 | 0.057433 | 0 | yellow | IER3 |
| 0 | 0.800559 | 0.424073 | 0.048125 | 0 | yellow | CDA |
| 0 | 0.799636 | 0.442267 | 0.05342 | 0 | yellow | PLBD1 |
| 0 | 0.780763 | 0.422673 | 0.04799 | 0 | yellow | CXCL16 |
| 0 | 0.773139 | 0.358293 | 0.019392 | 0 | yellow | PID1 |
| 0 | 0.75874 | 0.529741 | 0.149602 | 0 | yellow | UPP1 |
| 0 | 0.754182 | 0.991603 | 0.947187 | 0 | yellow | ATP5F1E |
| 0 | 0.752798 | 0.396081 | 0.064987 | 0 | yellow | MPEG1 |
| 0 | 0.74546 | 0.473058 | 0.122825 | 0 | yellow | PLEC |
| 0 | 0.744751 | 0.377187 | 0.075745 | 0 | yellow | ANPEP |
| 0 | 0.7434 | 0.43177 | 0.057972 | 0 | yellow | TGFBI |
| 0 | 0.74283 | 0.368789 | 0.0316 | 0 | yellow | SMIM25 |
| 0 | 0.731862 | 0.418474 | 0.045562 | 0 | yellow | CD86 |
| 0 | 0.718012 | 0.405878 | 0.073857 | 0 | yellow | CTSZ |
| 0 | 0.706047 | 0.426172 | 0.089404 | 0 | yellow | LILRA2 |
| 0 | 0.701528 | 0.391183 | 0.07841 | 0 | yellow | SLC43A2 |
| 0 | 0.697225 | 0.398181 | 0.06266 | 0 | yellow | RAB31 |
| 0 | 0.670937 | 0.996501 | 0.915149 | 0 | yellow | CD74 |
| 0 | 0.669561 | 0.373688 | 0.04799 | 0 | yellow | ITGAX |
| 0 | 0.669521 | 0.422673 | 0.094631 | 0 | yellow | STX11 |
| 0 | 0.647633 | 0.355493 | 0.063031 | 0 | yellow | ODF3B |
| 0 | 0.647609 | 0.959412 | 0.912148 | 0 | yellow | ARPC3 |
| 0 | 0.640571 | 0.988104 | 0.952044 | 0 | yellow | MYL6 |
| 0 | 0.636714 | 0.9986 | 0.98135 | 0 | yellow | SERF2 |
| 0 | 0.636373 | 0.361791 | 0.066808 | 0 | yellow | MIR22HG |
| 0 | 0.634722 | 0.29881 | 0.039559 | 0 | yellow | RBP7 |
| 0 | 0.633247 | 0.335899 | 0.042493 | 0 | yellow | CLEC4A |
| 0 | 0.616571 | 0.318404 | 0.057096 | 0 | yellow | MS4A7 |
| 0 | 0.60464 | 0.315605 | 0.054971 | 0 | yellow | FCGR2A |
| 0 | 0.602563 | 0.314906 | 0.027148 | 0 | yellow | LILRB3 |

| | | | | | | |
|---|---|---|---|---|---|---|
| 0 | 0.595815 | 0.343597 | 0.058512 | 0 | yellow | HCK |
| 0 | 0.589898 | 0.297411 | 0.032173 | 0 | yellow | STAB1 |
| 0 | 0.578533 | 0.256823 | 0.036659 | 0 | yellow | MCEMP1 |
| 0 | 0.577062 | 0.302309 | 0.043572 | 0 | yellow | LRRC25 |
| 0 | 0.562814 | 0.321204 | 0.060333 | 0 | yellow | FBP1 |
| 0 | 0.55606 | 0.257523 | 0.021719 | 0 | yellow | CLEC4E |
| 0 | 0.542978 | 0.289713 | 0.038547 | 0 | yellow | SULF2 |
| 0 | -0.52227 | 0.9993 | 0.99973 | 0 | yellow | RPL10 |
| 0 | -0.52833 | 0.996501 | 0.999191 | 0 | yellow | RPS23 |
| 0 | -0.54124 | 0.997201 | 0.999393 | 0 | yellow | RPS27A |
| 0 | -0.54557 | 0.988104 | 0.998617 | 0 | yellow | RPL7A |
| 0 | -0.58149 | 0.995101 | 0.998988 | 0 | yellow | RPS15A |
| 0 | -0.58566 | 0.941218 | 0.990658 | 0 | yellow | RPL23A |
| 0 | -0.60518 | 0.981106 | 0.996324 | 0 | yellow | RPL14 |
| 0 | -0.62015 | 0.9993 | 0.999393 | 0 | yellow | RPS12 |
| 0 | -0.62084 | 0.979006 | 0.99565 | 0 | yellow | RPS25 |
| 0 | -0.67584 | 0.937719 | 0.992884 | 0 | yellow | MT-ND4L |
| 0 | -0.70484 | 0.864941 | 0.97157 | 0 | yellow | EEF2 |
| 0 | -0.7371 | 0.996501 | 0.998921 | 0 | yellow | RPS3A |
| 0 | -0.74374 | 0.984605 | 0.996965 | 0 | yellow | RPS18 |
| 0 | -0.76428 | 0.909727 | 0.982396 | 0 | yellow | RPL10A |
| 0 | -0.8196 | 0.979006 | 0.994199 | 0 | yellow | RPS6 |
| 0 | -0.82063 | 0.96641 | 0.994132 | 0 | yellow | RPL5 |
| 0 | -0.82766 | 0.969209 | 0.995818 | 0 | yellow | RPS27 |
| 0 | -0.83361 | 0.997901 | 0.999696 | 0 | yellow | EEF1A1 |
| 0 | -0.88707 | 0.990903 | 0.998347 | 0 | yellow | RPS3 |
| 0 | -0.88861 | 0.988104 | 0.998314 | 0 | yellow | RPS4X |
| 0 | -0.92689 | 0.89853 | 0.98506 | 0 | yellow | RPSA |
| 0 | -0.96211 | 0.909727 | 0.98624 | 0 | yellow | RPS5 |
| 0 | -0.9659 | 0.284815 | 0.806489 | 0 | yellow | C12orf57 |
| 0 | -0.98152 | 0.871938 | 0.971267 | 0 | yellow | EEF1B2 |
| 0 | -1.01825 | 0.927222 | 0.991704 | 0 | yellow | RPL3 |
| 0 | -1.26225 | 0.442267 | 0.825105 | 0 | yellow | LDHB |
| 0 | -1.31054 | 0.689293 | 0.929752 | 0 | yellow | NPM1 |
| 0 | -1.51629 | 0.021693 | 0.589977 | 0 | yellow | CTSW |
| 5.40E-304 | 0.525356 | 0.262421 | 0.041582 | 1.29E-303 | yellow | TRIB1 |
| 6.41E-300 | 0.779909 | 0.893632 | 0.627074 | 1.53E-299 | yellow | TKT |
| 4.31E-298 | 0.863017 | 0.614416 | 0.255598 | 1.02E-297 | yellow | TNFSF13B |
| 5.45E-296 | -0.75658 | 0.850245 | 0.963274 | 1.29E-295 | yellow | HNRNPA1 |
| 8.93E-293 | 0.587538 | 0.286214 | 0.051767 | 2.09E-292 | yellow | DMXL2 |
| 4.60E-292 | -1.08464 | 0.212736 | 0.703899 | 1.07E-291 | yellow | GYPC |

| | | | | | | |
|---|---|---|---|---|---|---|
| 5.44E-291 | 0.500574 | 0.26522 | 0.044955 | 1.26E-290 | yellow | CSF1R |
| 1.32E-288 | 0.734476 | 0.457663 | 0.129772 | 3.04E-288 | yellow | JAML |
| 4.72E-285 | 0.845747 | 0.662001 | 0.298901 | 1.08E-284 | yellow | AGTRAP |
| 4.14E-279 | -0.86321 | 0.547936 | 0.875152 | 9.43E-279 | yellow | HSPA8 |
| 2.61E-278 | 0.685255 | 0.961512 | 0.726561 | 5.93E-278 | yellow | GSTP1 |
| 5.30E-278 | -0.65961 | 0.983205 | 0.995919 | 1.20E-277 | yellow | RPLP0 |
| 6.86E-277 | 0.506315 | 0.278516 | 0.050486 | 1.54E-276 | yellow | FAM49A |
| 1.15E-276 | 1.011451 | 0.736179 | 0.395724 | 2.57E-276 | yellow | NFKBIA |
| 2.14E-276 | -1.8508 | 0.083975 | 0.596486 | 4.74E-276 | yellow | IFITM1 |
| 9.08E-276 | -0.74504 | 0.759272 | 0.922703 | 2.01E-275 | yellow | HINT1 |
| 9.56E-275 | 0.559674 | 0.26592 | 0.047821 | 2.10E-274 | yellow | HBEGF |
| 3.36E-274 | 0.762802 | 0.517145 | 0.182787 | 7.35E-274 | yellow | ZYX |
| 5.83E-273 | 0.757432 | 0.864941 | 0.687306 | 1.27E-272 | yellow | LAMTOR4 |
| 2.86E-271 | 0.635463 | 0.374388 | 0.095407 | 6.19E-271 | yellow | GRINA |
| 5.73E-270 | 0.827487 | 0.776067 | 0.485802 | 1.23E-269 | yellow | ARRB2 |
| 7.51E-269 | -0.87293 | 0.23373 | 0.707237 | 1.61E-268 | yellow | 6-Sep |
| 4.97E-266 | -1.0029 | 0.660602 | 0.87903 | 1.06E-265 | yellow | HSP90AB1 |
| 5.05E-266 | 0.511605 | 0.273618 | 0.051936 | 1.07E-265 | yellow | EMILIN2 |
| 5.69E-265 | 0.588454 | 0.351994 | 0.086132 | 1.20E-264 | yellow | NLRP3 |
| 2.03E-258 | 0.782989 | 0.818055 | 0.524417 | 4.27E-258 | yellow | FLNA |
| 2.24E-258 | 0.965577 | 0.643107 | 0.290672 | 4.69E-258 | yellow | ID2 |
| 1.35E-256 | 0.783396 | 0.786564 | 0.527688 | 2.81E-256 | yellow | AP2S1 |
| 4.55E-255 | 0.511271 | 0.288314 | 0.059591 | 9.42E-255 | yellow | RXRA |
| 2.92E-254 | 0.751526 | 0.50035 | 0.176211 | 6.01E-254 | yellow | MIDN |
| 7.35E-250 | 0.519822 | 0.259622 | 0.048901 | 1.51E-249 | yellow | BASP1 |
| 1.53E-249 | 0.672189 | 0.887334 | 0.632369 | 3.12E-249 | yellow | ITGB2 |
| 2.95E-248 | 0.549588 | 0.314206 | 0.073385 | 5.98E-248 | yellow | PLXNB2 |
| 3.73E-248 | -1.15938 | 0.129461 | 0.607581 | 7.54E-248 | yellow | EVL |
| 8.33E-248 | 0.730425 | 0.537439 | 0.204708 | 1.67E-247 | yellow | FOSL2 |
| 1.95E-243 | 0.709526 | 0.615815 | 0.262546 | 3.90E-243 | yellow | BLVRB |
| 6.01E-243 | -1.2574 | 0.012596 | 0.490051 | 1.20E-242 | yellow | TRBC2 |
| 1.98E-241 | 0.570269 | 0.322603 | 0.077971 | 3.92E-241 | yellow | RELT |
| 3.53E-238 | -0.64637 | 0.685794 | 0.927425 | 6.96E-238 | yellow | CIRBP |
| 1.08E-237 | -1.28831 | 0.03219 | 0.50435 | 2.13E-237 | yellow | CD69 |
| 4.10E-237 | 0.704186 | 0.395381 | 0.112269 | 8.00E-237 | yellow | RHOB |
| 8.36E-237 | 0.807428 | 0.689293 | 0.366721 | 1.63E-236 | yellow | C1orf162 |
| 1.14E-236 | 0.718839 | 0.592022 | 0.244199 | 2.20E-236 | yellow | MYO1F |
| 5.00E-234 | 0.813271 | 0.76627 | 0.486038 | 9.64E-234 | yellow | HLA-DMA |
| 3.66E-232 | 0.859762 | 0.616515 | 0.302307 | 7.01E-232 | yellow | CAPG |
| 1.79E-231 | -0.80629 | 0.30091 | 0.70147 | 3.42E-231 | yellow | PEBP1 |
| 5.15E-230 | -0.71879 | 0.051085 | 0.527553 | 9.78E-230 | yellow | TSTD1 |

| | | | | | | |
|---|---|---|---|---|---|---|
| 3.29E-228 | 0.875957 | 0.862141 | 0.642655 | 6.22E-228 | yellow | HLA-DPB1 |
| 5.03E-222 | 0.843528 | 0.6669 | 0.353804 | 9.47E-222 | yellow | JUND |
| 5.83E-219 | 0.53427 | 0.287614 | 0.067044 | 1.09E-218 | yellow | BRI3 |
| 5.70E-218 | 0.814625 | 0.73268 | 0.414306 | 1.06E-217 | yellow | FOSB |
| 8.09E-217 | 0.536498 | 0.282715 | 0.066707 | 1.50E-216 | yellow | ASGR1 |
| 1.27E-216 | 0.734376 | 0.481456 | 0.176919 | 2.36E-216 | yellow | BCL2A1 |
| 4.92E-215 | 0.675261 | 0.509447 | 0.198975 | 9.07E-215 | yellow | THEMIS2 |
| 5.36E-214 | 0.663841 | 0.404479 | 0.134696 | 9.83E-214 | yellow | AP1S2 |
| 2.09E-213 | 0.793649 | 0.582225 | 0.279172 | 3.83E-213 | yellow | ATP2B1 |
| 5.58E-211 | 0.60887 | 0.893632 | 0.787974 | 1.02E-210 | yellow | ELOB |
| 2.58E-210 | -0.90327 | 0.643107 | 0.918791 | 4.68E-210 | yellow | SARAF |
| 1.56E-208 | 0.639726 | 0.439468 | 0.15692 | 2.82E-208 | yellow | MYO9B |
| 7.48E-206 | -0.66126 | 0.015395 | 0.446513 | 1.34E-205 | yellow | OCIAD2 |
| 2.16E-205 | 0.635565 | 0.40098 | 0.130919 | 3.85E-205 | yellow | EVI2B |
| 1.40E-204 | -1.53231 | 0.072778 | 0.497268 | 2.50E-204 | yellow | STMN1 |
| 1.59E-204 | 0.714441 | 0.601819 | 0.284905 | 2.82E-204 | yellow | TCIRG1 |
| 3.81E-204 | -0.81257 | 0.006998 | 0.43245 | 6.74E-204 | yellow | RHOH |
| 3.17E-203 | 0.608551 | 0.318404 | 0.089134 | 5.58E-203 | yellow | PPIF |
| 2.06E-202 | 0.623675 | 0.855143 | 0.683698 | 3.60E-202 | yellow | GNG5 |
| 5.67E-201 | -0.7508 | 0.002799 | 0.423108 | 9.88E-201 | yellow | ITM2A |
| 2.50E-198 | -0.9481 | 0.170749 | 0.576251 | 4.35E-198 | yellow | NUCB2 |
| 6.42E-198 | 0.81578 | 0.697691 | 0.452381 | 1.11E-197 | yellow | HIF1A |
| 1.74E-192 | -0.70262 | 0.572428 | 0.869553 | 3.00E-192 | yellow | RAC2 |
| 1.87E-191 | -1.84936 | 0.016795 | 0.41987 | 3.21E-191 | yellow | CD3E |
| 1.38E-190 | 0.66534 | 0.758572 | 0.513321 | 2.36E-190 | yellow | CYBA |
| 5.76E-190 | 0.833013 | 0.838348 | 0.641845 | 9.80E-190 | yellow | HLA-DPA1 |
| 3.16E-189 | 0.599338 | 0.419174 | 0.146567 | 5.35E-189 | yellow | ADA2 |
| 2.05E-188 | -1.41935 | 0.025892 | 0.428403 | 3.46E-188 | yellow | CD7 |
| 5.30E-188 | 0.593365 | 0.862141 | 0.752529 | 8.91E-188 | yellow | HIGD2A |
| 1.22E-187 | -0.69149 | 0.123863 | 0.556623 | 2.05E-187 | yellow | ACAP1 |
| 6.57E-186 | 0.767948 | 0.6662 | 0.423007 | 1.10E-185 | yellow | RILPL2 |
| 5.45E-184 | 0.553139 | 0.90063 | 0.800047 | 9.05E-184 | yellow | LSP1 |
| 7.89E-183 | 0.564772 | 0.342197 | 0.107851 | 1.31E-182 | yellow | TLE3 |
| 1.20E-179 | -2.09762 | 0.019594 | 0.405942 | 1.99E-179 | yellow | IL32 |
| 3.24E-179 | -0.97117 | 0.217635 | 0.613517 | 5.33E-179 | yellow | HIST1H4C |
| 2.25E-178 | 0.541552 | 0.355493 | 0.11237 | 3.67E-178 | yellow | SH3BP2 |
| 5.98E-178 | -0.68821 | 0.317705 | 0.679482 | 9.74E-178 | yellow | SNHG8 |
| 9.84E-178 | 0.698204 | 0.736179 | 0.440881 | 1.60E-177 | yellow | AHNAK |
| 9.81E-177 | 0.607648 | 0.379286 | 0.137293 | 1.59E-176 | yellow | ALDH2 |
| 2.43E-176 | -0.85054 | 0.030791 | 0.418589 | 3.91E-176 | yellow | CDK6 |
| 2.97E-175 | -0.62651 | 0.582925 | 0.822002 | 4.77E-175 | yellow | NCL |

| | | | | | | |
|---|---|---|---|---|---|---|
| 3.78E-175 | 0.512469 | 0.304409 | 0.089539 | 6.03E-175 | yellow | SH2B3 |
| 5.50E-175 | 0.734305 | 0.586424 | 0.307669 | 8.75E-175 | yellow | CARD16 |
| 6.40E-174 | -0.51039 | 0.678796 | 0.903312 | 1.01E-173 | yellow | HNRNPDL |
| 4.80E-172 | -0.59709 | 0.344997 | 0.717119 | 7.58E-172 | yellow | SOD1 |
| 4.82E-172 | 0.725931 | 0.686494 | 0.459902 | 7.58E-172 | yellow | CD63 |
| 2.52E-171 | 0.67614 | 0.643807 | 0.354951 | 3.95E-171 | yellow | HLA-DQB1 |
| 2.89E-170 | 0.631435 | 0.459762 | 0.192871 | 4.52E-170 | yellow | UBXN11 |
| 3.52E-169 | 0.593419 | 0.453464 | 0.182281 | 5.48E-169 | yellow | HLA-DMB |
| 1.13E-168 | 0.584791 | 0.850245 | 0.721368 | 1.75E-168 | yellow | COX8A |
| 6.87E-167 | 0.655437 | 0.776767 | 0.647713 | 1.06E-166 | yellow | ALDOA |
| 2.60E-165 | 0.653984 | 0.442267 | 0.192938 | 4.01E-165 | yellow | RAC1 |
| 1.38E-163 | 0.699044 | 0.864241 | 0.655841 | 2.12E-163 | yellow | KLF6 |
| 3.99E-161 | -0.57338 | 0.361092 | 0.703089 | 6.10E-161 | yellow | PTGES3 |
| 1.62E-160 | -0.59711 | 0.074178 | 0.460306 | 2.46E-160 | yellow | ARHGAP15 |
| 5.05E-160 | 0.721546 | 0.43317 | 0.190105 | 7.67E-160 | yellow | WARS |
| 2.16E-159 | -0.58757 | 0.020994 | 0.386955 | 3.27E-159 | yellow | BEX3 |
| 3.39E-159 | -1.11699 | 0.324703 | 0.639384 | 5.11E-159 | yellow | TUBB |
| 9.94E-159 | 0.58504 | 0.387684 | 0.154155 | 1.49E-158 | yellow | TNFSF13 |
| 1.11E-157 | 0.989385 | 0.458362 | 0.218333 | 1.66E-157 | yellow | HLA-DQA2 |
| 1.22E-157 | 0.623163 | 0.535339 | 0.239916 | 1.83E-157 | yellow | NCF1 |
| 3.69E-157 | 0.608687 | 0.488453 | 0.218097 | 5.48E-157 | yellow | GLIPR2 |
| 5.08E-157 | 0.587114 | 0.386284 | 0.152233 | 7.52E-157 | yellow | LYN |
| 1.17E-156 | 0.671681 | 0.464661 | 0.215061 | 1.72E-156 | yellow | NUP214 |
| 2.44E-156 | 0.567184 | 0.410777 | 0.164002 | 3.59E-156 | yellow | SCPEP1 |
| 1.44E-155 | -0.62006 | 0.36529 | 0.722717 | 2.11E-155 | yellow | SRSF7 |
| 3.28E-155 | -1.51216 | 0.064381 | 0.412485 | 4.79E-155 | yellow | LTB |
| 1.60E-153 | 0.501329 | 0.378586 | 0.134763 | 2.33E-153 | yellow | CD4 |
| 3.08E-153 | 0.58732 | 0.883835 | 0.699818 | 4.47E-153 | yellow | ZFP36 |
| 3.48E-153 | 0.637984 | 0.603219 | 0.351747 | 5.02E-153 | yellow | RNF130 |
| 9.56E-153 | -0.85934 | 0.169349 | 0.50435 | 1.38E-152 | yellow | IMPDH2 |
| 1.64E-151 | 0.671462 | 0.56683 | 0.317955 | 2.36E-151 | yellow | ARPC1B |
| 1.84E-151 | 0.569521 | 0.610917 | 0.299508 | 2.63E-151 | yellow | SAMHD1 |
| 2.65E-151 | -0.84719 | 0.025892 | 0.375658 | 3.78E-151 | yellow | ITM2C |
| 1.62E-150 | 0.630337 | 0.393982 | 0.159315 | 2.30E-150 | yellow | GNB2 |
| 1.79E-150 | -0.65929 | 0.006998 | 0.349015 | 2.53E-150 | yellow | SLC38A1 |
| 6.95E-150 | -0.72045 | 0.092372 | 0.456765 | 9.81E-150 | yellow | 1-Sep |
| 3.17E-149 | 0.750738 | 0.330301 | 0.116417 | 4.46E-149 | yellow | DUSP6 |
| 3.94E-149 | -0.53119 | 0.064381 | 0.429651 | 5.52E-149 | yellow | ZNF22 |
| 1.23E-147 | -1.17459 | 0.013996 | 0.351443 | 1.72E-147 | yellow | PRSS57 |
| 4.85E-147 | 0.622829 | 0.810357 | 0.737488 | 6.76E-147 | yellow | ITM2B |
| 5.77E-147 | 0.583655 | 0.369489 | 0.146027 | 8.01E-147 | yellow | PLXDC2 |

| | | | | | | |
|---|---|---|---|---|---|---|
| 1.97E-146 | 0.601333 | 0.869839 | 0.705686 | 2.73E-146 | yellow | IER2 |
| 2.27E-146 | 0.657711 | 0.573828 | 0.330737 | 3.13E-146 | yellow | PTP4A2 |
| 1.51E-144 | 0.61748 | 0.662701 | 0.408775 | 2.08E-144 | yellow | IQGAP1 |
| 2.81E-144 | -0.99281 | 0.004199 | 0.334918 | 3.84E-144 | yellow | LCK |
| 6.74E-144 | -0.54179 | 0.135759 | 0.505565 | 9.21E-144 | yellow | NOP58 |
| 7.11E-143 | -1.13593 | 0.006998 | 0.33448 | 9.68E-143 | yellow | CD3D |
| 7.20E-143 | -0.66271 | 0.20084 | 0.567719 | 9.77E-143 | yellow | PRDX2 |
| 1.61E-142 | -0.53261 | 0.488453 | 0.787502 | 2.18E-142 | yellow | EIF4B |
| 1.64E-142 | 0.642929 | 0.540238 | 0.304229 | 2.21E-142 | yellow | ATP6V0D1 |
| 2.31E-142 | -0.54753 | 0.349895 | 0.71233 | 3.10E-142 | yellow | EIF4A2 |
| 4.72E-142 | 0.701247 | 0.356193 | 0.134426 | 6.32E-142 | yellow | NR4A1 |
| 1.75E-141 | -0.5439 | 0.462561 | 0.77988 | 2.33E-141 | yellow | PNISR |
| 9.57E-141 | -0.55351 | 0.242127 | 0.592068 | 1.27E-140 | yellow | SRP9 |
| 3.98E-140 | -0.57376 | 0.116865 | 0.469513 | 5.29E-140 | yellow | PARP1 |
| 4.18E-140 | -0.71187 | 0.09937 | 0.444017 | 5.54E-140 | yellow | TBC1D10C |
| 5.48E-140 | 0.612656 | 0.421274 | 0.191623 | 7.22E-140 | yellow | CAPNS1 |
| 7.27E-138 | -0.51564 | 0.137159 | 0.507015 | 9.55E-138 | yellow | TAF7 |
| 1.04E-137 | 0.617699 | 0.421274 | 0.195366 | 1.36E-137 | yellow | PRELID1 |
| 1.35E-137 | 0.561569 | 0.776767 | 0.66498 | 1.76E-137 | yellow | ATP6V0C |
| 1.08E-136 | -0.73352 | 0.209237 | 0.571327 | 1.40E-136 | yellow | LEPROTL1 |
| 4.84E-136 | 0.602901 | 0.715185 | 0.551835 | 6.28E-136 | yellow | CNPY3 |
| 9.23E-136 | -0.5185 | 0.426872 | 0.723088 | 1.20E-135 | yellow | ST13 |
| 4.57E-135 | -0.54935 | 0.585724 | 0.794921 | 5.90E-135 | yellow | EIF3E |
| 1.31E-134 | 0.522819 | 0.848845 | 0.607311 | 1.69E-134 | yellow | TALDO1 |
| 7.66E-134 | -0.78876 | 0.054584 | 0.38247 | 9.83E-134 | yellow | AC084033.3 |
| 1.00E-133 | -0.91566 | 0.005598 | 0.317887 | 1.28E-133 | yellow | SPOCK2 |
| 3.82E-133 | 0.589716 | 0.720084 | 0.58738 | 4.87E-133 | yellow | ATP6V1F |
| 5.66E-133 | 0.644986 | 0.687194 | 0.52128 | 7.19E-133 | yellow | CSTB |
| 1.44E-132 | -0.59985 | 0.471659 | 0.726022 | 1.83E-132 | yellow | RAN |
| 5.11E-132 | 0.609718 | 0.635409 | 0.432281 | 6.46E-132 | yellow | RNH1 |
| 1.25E-131 | -0.59348 | 0.069279 | 0.405436 | 1.58E-131 | yellow | CYFIP2 |
| 2.78E-131 | -0.51883 | 0.242127 | 0.588088 | 3.49E-131 | yellow | RBMX |
| 8.42E-131 | 0.655478 | 0.545836 | 0.320383 | 1.05E-130 | yellow | GRK2 |
| 1.57E-130 | -0.71106 | 0.241428 | 0.544044 | 1.96E-130 | yellow | HSPD1 |
| 1.08E-129 | 0.54145 | 0.327502 | 0.125489 | 1.35E-129 | yellow | MAP2K3 |
| 1.26E-129 | 0.55217 | 0.816655 | 0.676784 | 1.56E-129 | yellow | PKM |
| 1.49E-129 | -0.71535 | 0.448565 | 0.73135 | 1.84E-129 | yellow | CALR |
| 3.79E-129 | 0.623916 | 0.644507 | 0.453325 | 4.68E-129 | yellow | GRB2 |
| 1.49E-128 | -0.56839 | 0.205038 | 0.526541 | 1.83E-128 | yellow | CCT2 |
| 2.07E-127 | -0.51163 | 0.321204 | 0.677762 | 2.54E-127 | yellow | 7-Sep |
| 1.29E-126 | -0.85306 | 0.008397 | 0.30777 | 1.57E-126 | yellow | FCMR |

| | | | | | | |
|---|---|---|---|---|---|---|
| 1.55E-124 | -0.60164 | 0.016095 | 0.319877 | 1.89E-124 | yellow | EGFL7 |
| 7.71E-124 | -0.63553 | 0 | 0.292729 | 9.38E-124 | yellow | CD34 |
| 1.35E-123 | -0.62686 | 0.002799 | 0.296169 | 1.64E-123 | yellow | FAM30A |
| 5.04E-123 | 0.664127 | 0.563331 | 0.359166 | 6.10E-123 | yellow | C20orf27 |
| 1.36E-122 | -0.90822 | 0.326802 | 0.617901 | 1.64E-122 | yellow | CXCR4 |
| 9.83E-122 | 0.612335 | 0.689993 | 0.529846 | 1.18E-121 | yellow | PGLS |
| 1.11E-121 | 0.623206 | 0.435269 | 0.224808 | 1.33E-121 | yellow | AMPD2 |
| 3.24E-120 | 0.62967 | 0.394682 | 0.182079 | 3.87E-120 | yellow | ANKRD13D |
| 5.13E-120 | 0.566541 | 0.708188 | 0.537535 | 6.12E-120 | yellow | PLP2 |
| 8.89E-120 | -1.40257 | 0.013996 | 0.303386 | 1.06E-119 | yellow | IL7R |
| 9.53E-120 | 0.600526 | 0.689293 | 0.475246 | 1.13E-119 | yellow | GLUL |
| 2.26E-119 | -0.54306 | 0.06788 | 0.384595 | 2.67E-119 | yellow | PBXIP1 |
| 8.92E-119 | 0.599161 | 0.386984 | 0.173985 | 1.05E-118 | yellow | RAB11FIP1 |
| 4.39E-118 | 0.556846 | 0.326102 | 0.122285 | 5.16E-118 | yellow | RNASE2 |
| 4.90E-116 | -0.5356 | 0.297411 | 0.589404 | 5.75E-116 | yellow | APEX1 |
| 1.47E-115 | 0.529324 | 0.413576 | 0.194523 | 1.72E-115 | yellow | TOM1 |
| 5.26E-115 | -0.61184 | 0.218334 | 0.542088 | 6.13E-115 | yellow | CCND3 |
| 7.90E-115 | -0.63369 | 0.186844 | 0.524518 | 9.18E-115 | yellow | NOSIP |
| 1.80E-114 | -0.76648 | 0.009797 | 0.288817 | 2.08E-114 | yellow | ZAP70 |
| 1.46E-113 | 0.560861 | 0.720784 | 0.602792 | 1.68E-113 | yellow | RHOG |
| 2.24E-111 | -0.75015 | 0.002099 | 0.27273 | 2.58E-111 | yellow | SMIM24 |
| 3.56E-111 | -0.76229 | 0.002799 | 0.273371 | 4.09E-111 | yellow | HOPX |
| 8.64E-110 | -0.68514 | 0.008397 | 0.278227 | 9.91E-110 | yellow | LBH |
| 8.80E-110 | 0.50373 | 0.303009 | 0.123567 | 1.01E-109 | yellow | BST1 |
| 5.27E-109 | 0.516642 | 0.371589 | 0.166734 | 6.01E-109 | yellow | GPCPD1 |
| 3.41E-107 | 0.587517 | 0.659902 | 0.49683 | 3.88E-107 | yellow | LRRFIP1 |
| 1.56E-106 | 0.519206 | 0.373688 | 0.175671 | 1.77E-106 | yellow | PRAM1 |
| 2.67E-106 | -0.92557 | 0.002799 | 0.263355 | 3.02E-106 | yellow | SPINK2 |
| 6.35E-106 | -0.79727 | 0.003499 | 0.26376 | 7.16E-106 | yellow | CD3G |
| 7.32E-106 | -0.57253 | 0.049685 | 0.329455 | 8.24E-106 | yellow | RNF125 |
| 7.86E-106 | -0.6058 | 0.005598 | 0.267402 | 8.83E-106 | yellow | ABLIM1 |
| 1.01E-105 | -1.39359 | 0.006298 | 0.267604 | 1.13E-105 | yellow | IGLL1 |
| 1.02E-105 | -0.87905 | 0.004199 | 0.264502 | 1.14E-105 | yellow | GZMM |
| 1.92E-105 | 0.567966 | 0.426172 | 0.223897 | 2.13E-105 | yellow | ZFAND5 |
| 2.35E-104 | 0.536605 | 0.356893 | 0.1638 | 2.61E-104 | yellow | PHACTR1 |
| 4.37E-104 | -1.46679 | 0.040588 | 0.309085 | 4.84E-104 | yellow | CST7 |
| 1.17E-103 | -0.50401 | 0.186144 | 0.504047 | 1.30E-103 | yellow | PRMT2 |
| 1.73E-103 | -0.69318 | 0.256123 | 0.570349 | 1.91E-103 | yellow | DNAJB1 |
| 5.26E-103 | -0.60943 | 0.0014 | 0.254924 | 5.78E-103 | yellow | SKAP1 |
| 8.14E-103 | 0.536991 | 0.513646 | 0.304465 | 8.93E-103 | yellow | SAT2 |
| 1.10E-102 | -0.61665 | 0.018894 | 0.279981 | 1.20E-102 | yellow | PRKCH |

| | | | | | | |
|---|---|---|---|---|---|---|
| 2.01E-102 | -0.55158 | 0.057383 | 0.335964 | 2.20E-102 | yellow | MCM7 |
| 1.95E-101 | -0.60885 | 0.263821 | 0.551228 | 2.12E-101 | yellow | DUT |
| 1.38E-98 | -0.85066 | 0.004899 | 0.251113 | 1.50E-98 | yellow | CD247 |
| 1.92E-98 | 0.569209 | 0.463961 | 0.282241 | 2.08E-98 | yellow | LGALS9 |
| 3.64E-97 | -0.7282 | 0.047586 | 0.30315 | 3.93E-97 | yellow | PIK3IP1 |
| 4.28E-97 | -0.6422 | 0.025892 | 0.278362 | 4.61E-97 | yellow | LAT |
| 9.38E-97 | 0.58101 | 0.442967 | 0.2615 | 1.01E-96 | yellow | PLSCR1 |
| 1.00E-95 | 0.58783 | 0.414976 | 0.235364 | 1.07E-95 | yellow | TBXAS1 |
| 1.64E-95 | 0.502306 | 0.740378 | 0.648928 | 1.76E-95 | yellow | CD44 |
| 4.66E-94 | 0.511979 | 0.347096 | 0.168353 | 4.97E-94 | yellow | CPPED1 |
| 4.68E-94 | 0.546166 | 0.517845 | 0.338493 | 4.97E-94 | yellow | PCBP1 |
| 7.42E-92 | 0.547696 | 0.514346 | 0.327634 | 7.87E-92 | yellow | KLF10 |
| 1.40E-91 | 0.522588 | 0.481456 | 0.279981 | 1.49E-91 | yellow | CAST |
| 1.09E-90 | -0.54677 | 0.118964 | 0.389788 | 1.15E-90 | yellow | P2RY8 |
| 1.90E-88 | -0.70376 | 0.623513 | 0.827971 | 2.00E-88 | yellow | TSC22D3 |
| 2.76E-86 | 0.536402 | 0.595521 | 0.442432 | 2.90E-86 | yellow | PGD |
| 9.51E-80 | -0.54779 | 0.230931 | 0.466916 | 9.96E-80 | yellow | CAT |
| 4.93E-77 | 0.508352 | 0.648705 | 0.513321 | 5.15E-77 | yellow | DDX21 |
| 7.23E-74 | 0.504478 | 0.359692 | 0.201167 | 7.54E-74 | yellow | MXD1 |
| 3.82E-73 | 0.500648 | 0.447166 | 0.285546 | 3.97E-73 | yellow | MGAT1 |
| 1.52E-72 | -0.56422 | 0.118265 | 0.340753 | 1.57E-72 | yellow | ISG20 |
| 6.12E-72 | -0.60149 | 0.057383 | 0.265749 | 6.33E-72 | yellow | FLT3LG |
| 7.40E-69 | 0.522298 | 0.477957 | 0.316471 | 7.63E-69 | yellow | PER1 |
| 9.46E-69 | 0.516479 | 0.451365 | 0.308242 | 9.74E-69 | yellow | BID |
| 1.19E-68 | -0.51583 | 0.860042 | 0.922973 | 1.22E-68 | yellow | HMGB1 |
| 2.79E-63 | -0.50128 | 0.069979 | 0.278868 | 2.86E-63 | yellow | CMC1 |
| 6.18E-62 | 0.508194 | 0.453464 | 0.312289 | 6.32E-62 | yellow | SOD2 |
| 2.33E-61 | -0.55766 | 0.174948 | 0.403177 | 2.37E-61 | yellow | SOX4 |
| 3.51E-61 | -0.69875 | 0.223233 | 0.431236 | 3.57E-61 | yellow | TNFAIP3 |
| 9.74E-55 | 0.743117 | 0.571728 | 0.479664 | 9.88E-55 | yellow | IFITM3 |
| 1.71E-49 | -0.85212 | 0.559132 | 0.66724 | 1.73E-49 | yellow | TUBA1B |
| 5.01E-47 | 0.514916 | 0.468859 | 0.361594 | 5.06E-47 | yellow | JPT1 |
| 3.27E-46 | 0.695697 | 0.26802 | 0.152806 | 3.30E-46 | yellow | IL1B |
| 8.17E-34 | -1.87091 | 0.174948 | 0.311682 | 8.21E-34 | yellow | NKG7 |
| 1.87E-32 | -0.55993 | 0.190343 | 0.336908 | 1.87E-32 | yellow | AREG |
| 5.76E-12 | -0.99234 | 0.296011 | 0.210509 | 5.76E-12 | yellow | HBB |
| 0 | 2.575127 | 0.982857 | 0.103702 | 0 | cyan | SERPINA1 |
| 0 | 2.416669 | 0.954286 | 0.071798 | 0 | cyan | FCGR3A |
| 0 | 2.239602 | 0.925714 | 0.064259 | 0 | cyan | MS4A7 |
| 0 | 2.132165 | 0.965714 | 0.103896 | 0 | cyan | LILRB2 |
| 0 | 2.080588 | 0.948571 | 0.065651 | 0 | cyan | C5AR1 |

| | | | | | | |
|---|---|---|---|---|---|---|
| 0 | 1.957825 | 0.891429 | 0.042322 | 0 | cyan | SMIM25 |
| 0 | 1.742946 | 0.874286 | 0.068951 | 0 | cyan | PILRA |
| 0 | 1.719064 | 0.902857 | 0.070536 | 0 | cyan | LILRA5 |
| 0 | 1.616546 | 0.811429 | 0.050799 | 0 | cyan | CSF1R |
| 0 | 1.609073 | 0.817143 | 0.045493 | 0 | cyan | HMOX1 |
| 0 | 1.554489 | 0.817143 | 0.067398 | 0 | cyan | HCK |
| 0 | 1.540148 | 0.851429 | 0.050961 | 0 | cyan | LRRC25 |
| 0 | 1.53102 | 0.337143 | 0.000226 | 0 | cyan | LYPD2 |
| 0 | 1.494511 | 0.788571 | 0.061121 | 0 | cyan | CXCL16 |
| 0 | 1.428068 | 0.828571 | 0.069339 | 0 | cyan | IFI30 |
| 0 | 1.257167 | 0.754286 | 0.059794 | 0 | cyan | SLC7A7 |
| 0 | 1.209395 | 0.662857 | 0.039636 | 0 | cyan | CTSL |
| 0 | 1.178165 | 0.577143 | 0.00948 | 0 | cyan | CDKN1C |
| 0 | 1.167978 | 0.291429 | 0.002394 | 0 | cyan | VMO1 |
| 0 | 1.055527 | 0.622857 | 0.029768 | 0 | cyan | LILRA1 |
| 0 | 0.966424 | 0.611429 | 0.03255 | 0 | cyan | HK3 |
| 0 | 0.838954 | 0.48 | 0.018249 | 0 | cyan | GPBAR1 |
| 0 | 0.775717 | 0.434286 | 0.016405 | 0 | cyan | CDH23 |
| 0 | 0.741448 | 0.405714 | 0.003365 | 0 | cyan | LINC02432 |
| 0 | 0.672961 | 0.365714 | 0.012651 | 0 | cyan | C3AR1 |
| 0 | 0.59978 | 0.354286 | 0.005662 | 0 | cyan | ZNF703 |
| 0 | 0.557395 | 0.354286 | 0.008898 | 0 | cyan | TPPP3 |
| 1.23E-289 | 1.332743 | 0.828571 | 0.078884 | 2.29E-288 | cyan | FGL2 |
| 1.83E-287 | 1.269809 | 0.862857 | 0.083414 | 3.28E-286 | cyan | CLEC7A |
| 1.69E-285 | 1.162677 | 0.64 | 0.049667 | 2.93E-284 | cyan | SIGLEC10 |
| 5.86E-285 | 1.339712 | 0.765714 | 0.068951 | 9.83E-284 | cyan | LILRB1 |
| 2.48E-277 | 0.567668 | 0.4 | 0.018249 | 4.02E-276 | cyan | SLC24A4 |
| 1.73E-256 | 0.830271 | 0.497143 | 0.030933 | 2.73E-255 | cyan | SCIMP |
| 7.67E-243 | 1.374071 | 0.897143 | 0.113279 | 1.17E-241 | cyan | SLC11A1 |
| 1.87E-236 | 2.012569 | 0.977143 | 0.16026 | 2.78E-235 | cyan | CD68 |
| 4.98E-228 | 1.174796 | 0.76 | 0.082961 | 7.20E-227 | cyan | NCF2 |
| 5.74E-221 | 0.921381 | 0.64 | 0.059438 | 8.07E-220 | cyan | CD86 |
| 4.48E-215 | 0.982049 | 0.571429 | 0.050476 | 6.12E-214 | cyan | OAS1 |
| 2.06E-210 | 0.71672 | 0.502857 | 0.03776 | 2.75E-209 | cyan | LILRB3 |
| 4.27E-209 | 1.239885 | 0.748571 | 0.083188 | 5.56E-208 | cyan | TMEM176B |
| 3.73E-207 | 0.847129 | 0.6 | 0.053129 | 4.73E-206 | cyan | CD300E |
| 1.04E-203 | 1.389213 | 0.594286 | 0.054714 | 1.29E-202 | cyan | APOBEC3A |
| 5.77E-202 | 0.860752 | 0.605714 | 0.057885 | 6.97E-201 | cyan | FAM49A |
| 1.08E-201 | 1.447125 | 0.931429 | 0.126027 | 1.27E-200 | cyan | FCN1 |
| 3.74E-200 | 1.503375 | 0.92 | 0.145797 | 4.32E-199 | cyan | TYMP |
| 4.90E-200 | 1.505122 | 0.931429 | 0.154242 | 5.53E-199 | cyan | FGR |

| | | | | | | |
|---|---|---|---|---|---|---|
| 1.41E-197 | 1.230974 | 0.685714 | 0.08445 | 1.56E-196 | cyan | TCF7L2 |
| 5.96E-196 | 1.318515 | 0.84 | 0.118941 | 6.46E-195 | cyan | CYBB |
| 6.90E-192 | 0.992136 | 0.68 | 0.076814 | 7.32E-191 | cyan | MPEG1 |
| 9.17E-192 | 0.61914 | 0.377143 | 0.02417 | 9.53E-191 | cyan | MS4A4A |
| 2.14E-190 | 0.564799 | 0.394286 | 0.025691 | 2.18E-189 | cyan | CHST15 |
| 4.08E-188 | 0.568458 | 0.422857 | 0.029962 | 4.08E-187 | cyan | SIRPB1 |
| 5.44E-187 | 0.908883 | 0.64 | 0.067042 | 5.34E-186 | cyan | FPR1 |
| 3.21E-186 | 1.625535 | 0.931429 | 0.167573 | 3.09E-185 | cyan | CFD |
| 1.58E-185 | 1.122658 | 0.742857 | 0.101275 | 1.49E-184 | cyan | LILRA2 |
| 2.84E-185 | 0.901652 | 0.525714 | 0.048825 | 2.64E-184 | cyan | SLC2A6 |
| 3.97E-182 | 0.990636 | 0.594286 | 0.063968 | 3.62E-181 | cyan | FCGR2A |
| 7.44E-182 | 2.406136 | 0.994286 | 0.253381 | 6.67E-181 | cyan | FCER1G |
| 5.16E-172 | 1.471875 | 0.874286 | 0.157542 | 4.54E-171 | cyan | CFP |
| 3.02E-168 | 1.025107 | 0.668571 | 0.08212 | 2.61E-167 | cyan | IGSF6 |
| 8.88E-168 | 1.307532 | 0.857143 | 0.139843 | 7.57E-167 | cyan | LGALS3 |
| 6.44E-165 | 1.752678 | 0.902857 | 0.189575 | 5.40E-164 | cyan | STXBP2 |
| 7.53E-161 | 1.173354 | 0.794286 | 0.13829 | 6.21E-160 | cyan | C19orf38 |
| 2.54E-160 | 0.500854 | 0.325714 | 0.02087 | 2.06E-159 | cyan | ADAP2 |
| 3.19E-159 | 1.472882 | 0.931429 | 0.191969 | 2.56E-158 | cyan | COTL1 |
| 1.01E-157 | 1.401347 | 0.897143 | 0.186922 | 7.99E-157 | cyan | BCL2A1 |
| 9.21E-157 | 1.217481 | 0.84 | 0.149097 | 7.15E-156 | cyan | TNFRSF1B |
| 2.11E-155 | 1.486899 | 0.788571 | 0.140976 | 1.62E-154 | cyan | NR4A1 |
| 1.73E-153 | 1.034059 | 0.674286 | 0.090824 | 1.31E-152 | cyan | MNDA |
| 3.41E-152 | 0.887328 | 0.531429 | 0.061671 | 2.54E-151 | cyan | SLC31A2 |
| 4.49E-152 | 0.647849 | 0.428571 | 0.038213 | 3.29E-151 | cyan | CD300C |
| 3.89E-149 | 0.620033 | 0.451429 | 0.040931 | 2.81E-148 | cyan | CX3CR1 |
| 3.98E-149 | 0.777899 | 0.56 | 0.062609 | 2.83E-148 | cyan | CDA |
| 7.77E-147 | 1.657159 | 0.851429 | 0.197599 | 5.46E-146 | cyan | WARS |
| 5.74E-144 | 2.21545 | 0.988571 | 0.343202 | 3.98E-143 | cyan | S100A11 |
| 1.01E-143 | 0.541914 | 0.382857 | 0.031644 | 6.94E-143 | cyan | LILRA6 |
| 7.00E-142 | 2.641973 | 0.982857 | 0.377047 | 4.73E-141 | cyan | LST1 |
| 3.05E-141 | 0.845651 | 0.565714 | 0.06976 | 2.03E-140 | cyan | ALOX5 |
| 7.52E-141 | 0.670664 | 0.537143 | 0.059827 | 4.95E-140 | cyan | TMEM176A |
| 5.54E-137 | 0.547098 | 0.36 | 0.029897 | 3.60E-136 | cyan | SNX18 |
| 5.15E-136 | 0.951926 | 0.571429 | 0.079143 | 3.30E-135 | cyan | CLEC12A |
| 6.06E-136 | 1.008672 | 0.68 | 0.109267 | 3.84E-135 | cyan | MTSS1 |
| 7.85E-133 | 0.988179 | 0.554286 | 0.077234 | 4.92E-132 | cyan | FAM110A |
| 1.51E-132 | 1.797834 | 0.994286 | 0.321523 | 9.34E-132 | cyan | TYROBP |
| 6.95E-131 | 0.871044 | 0.565714 | 0.077008 | 4.25E-130 | cyan | RHOC |
| 1.71E-130 | 0.912843 | 0.662857 | 0.106581 | 1.03E-129 | cyan | STX11 |
| 1.06E-121 | 0.701789 | 0.514286 | 0.062447 | 6.31E-121 | cyan | MAFB |

| | | | | | | |
|---|---|---|---|---|---|---|
| 1.35E-120 | 0.732701 | 0.485714 | 0.06303 | 7.98E-120 | cyan | ADGRE2 |
| 3.04E-120 | 0.549367 | 0.314286 | 0.026014 | 1.78E-119 | cyan | MTMR11 |
| 4.00E-120 | 0.808814 | 0.548571 | 0.075422 | 2.31E-119 | cyan | ABI3 |
| 6.92E-119 | 2.007942 | 0.988571 | 0.46182 | 3.95E-118 | cyan | CTSS |
| 6.93E-118 | 1.630229 | 0.902857 | 0.300815 | 3.91E-117 | cyan | PECAM1 |
| 1.21E-117 | 0.737609 | 0.485714 | 0.060571 | 6.76E-117 | cyan | ITGAX |
| 2.81E-117 | 0.744982 | 0.52 | 0.075131 | 1.55E-116 | cyan | CD300LF |
| 1.72E-114 | 1.930099 | 0.988571 | 0.547078 | 9.44E-114 | cyan | AIF1 |
| 5.48E-113 | 2.020141 | 0.971429 | 0.406717 | 2.97E-112 | cyan | TIMP1 |
| 3.66E-110 | 2.218449 | 1 | 0.620947 | 1.96E-109 | cyan | SAT1 |
| 6.25E-110 | 2.250676 | 0.988571 | 0.691549 | 3.32E-109 | cyan | PSAP |
| 8.43E-109 | 2.235159 | 0.96 | 0.481201 | 4.43E-108 | cyan | IFITM3 |
| 1.00E-108 | 1.068186 | 0.857143 | 0.204006 | 5.21E-108 | cyan | PLAUR |
| 1.31E-106 | 1.02887 | 0.708571 | 0.159904 | 6.75E-106 | cyan | LYN |
| 4.61E-106 | 1.461871 | 0.971429 | 0.46182 | 2.35E-105 | cyan | ASAH1 |
| 1.06E-103 | 0.851165 | 0.485714 | 0.069986 | 5.34E-103 | cyan | FBP1 |
| 4.26E-103 | 1.401065 | 0.537143 | 0.082767 | 2.13E-102 | cyan | G0S2 |
| 6.94E-101 | 0.787938 | 0.485714 | 0.074387 | 3.44E-100 | cyan | RNF144B |
| 8.56E-101 | 0.808475 | 0.502857 | 0.07772 | 4.20E-100 | cyan | TNFSF10 |
| 2.29E-100 | 1.761748 | 0.994286 | 0.994111 | 1.11E-99 | cyan | FTL |
| 2.88E-100 | 0.968984 | 0.714286 | 0.173526 | 1.39E-99 | cyan | CPPED1 |
| 5.98E-99 | 0.746324 | 0.508571 | 0.082055 | 2.85E-98 | cyan | PLXNB2 |
| 1.47E-98 | 1.365979 | 0.988571 | 0.544716 | 6.95E-98 | cyan | NPC2 |
| 1.70E-98 | 1.266331 | 0.897143 | 0.323788 | 7.95E-98 | cyan | ANXA5 |
| 2.72E-98 | 1.012355 | 0.691429 | 0.150294 | 1.26E-97 | cyan | CD83 |
| 3.55E-98 | 1.504481 | 1 | 0.996473 | 1.64E-97 | cyan | FTH1 |
| 3.92E-97 | 0.744045 | 0.56 | 0.099075 | 1.79E-96 | cyan | GNS |
| 1.51E-95 | 0.695834 | 0.377143 | 0.048437 | 6.85E-95 | cyan | CKB |
| 2.87E-95 | -1.21356 | 0.982857 | 0.998091 | 1.29E-94 | cyan | RPS3 |
| 4.91E-95 | 1.296033 | 0.874286 | 0.292662 | 2.18E-94 | cyan | NAMPT |
| 1.70E-93 | 0.971795 | 0.702857 | 0.163852 | 7.47E-93 | cyan | POU2F2 |
| 7.55E-93 | 1.427448 | 0.977143 | 0.404743 | 3.30E-92 | cyan | CST3 |
| 4.45E-92 | 1.08865 | 0.748571 | 0.198473 | 1.93E-91 | cyan | CASP1 |
| 6.00E-92 | 0.654895 | 0.474286 | 0.074225 | 2.58E-91 | cyan | SIDT2 |
| 4.54E-91 | 1.15 | 0.857143 | 0.311752 | 1.93E-90 | cyan | BID |
| 5.52E-91 | 0.948238 | 0.771429 | 0.196984 | 2.34E-90 | cyan | LYST |
| 6.61E-91 | 1.121345 | 0.994286 | 0.921698 | 2.77E-90 | cyan | OAZ1 |
| 1.21E-90 | -0.95575 | 0.994286 | 0.999029 | 5.05E-90 | cyan | RPL13 |
| 1.34E-90 | 1.128592 | 0.794286 | 0.244936 | 5.54E-90 | cyan | SPI1 |
| 5.06E-90 | 0.593589 | 0.451429 | 0.067948 | 2.07E-89 | cyan | RXRA |
| 1.64E-88 | 0.943362 | 0.714286 | 0.180806 | 6.68E-88 | cyan | UNC119 |

| | | | | | | |
|---|---|---|---|---|---|---|
| 2.59E-88 | 0.766911 | 0.605714 | 0.120818 | 1.05E-87 | cyan | SH3BP2 |
| 4.47E-88 | -1.32584 | 0.954286 | 0.988934 | 1.79E-87 | cyan | RPL3 |
| 5.17E-87 | 1.386974 | 0.92 | 0.439365 | 2.05E-86 | cyan | VSIR |
| 6.38E-87 | 1.430675 | 0.994286 | 0.604834 | 2.51E-86 | cyan | NEAT1 |
| 9.41E-87 | 1.424655 | 0.994286 | 0.810684 | 3.68E-86 | cyan | S100A4 |
| 9.48E-86 | -1.22222 | 0.977143 | 0.996506 | 3.68E-85 | cyan | RPS18 |
| 1.70E-85 | 1.445525 | 0.988571 | 0.772633 | 6.54E-85 | cyan | IFITM2 |
| 2.82E-85 | 1.002823 | 0.742857 | 0.219569 | 1.08E-84 | cyan | MYD88 |
| 3.20E-85 | 0.668425 | 0.514286 | 0.090403 | 1.22E-84 | cyan | SLC43A2 |
| 1.15E-84 | 0.950362 | 1 | 0.999644 | 4.33E-84 | cyan | ACTB |
| 1.37E-84 | -0.75477 | 0.994286 | 0.999321 | 5.14E-84 | cyan | RPS27A |
| 1.52E-84 | 1.096935 | 0.857143 | 0.296318 | 5.64E-84 | cyan | TCIRG1 |
| 2.12E-84 | 0.666584 | 0.445714 | 0.073384 | 7.81E-84 | cyan | SMCO4 |
| 2.83E-84 | -1.21039 | 0.977143 | 0.993593 | 1.04E-83 | cyan | RPS6 |
| 4.23E-83 | -0.82897 | 0.994286 | 0.999321 | 1.54E-82 | cyan | RPL32 |
| 1.17E-82 | -1.19797 | 0.925714 | 0.981395 | 4.24E-82 | cyan | RPSA |
| 1.54E-82 | -0.65146 | 0.994286 | 0.999482 | 5.53E-82 | cyan | RPL19 |
| 4.04E-81 | 1.371698 | 0.902857 | 0.431081 | 1.44E-80 | cyan | CTSC |
| 1.39E-80 | 1.2874 | 0.902857 | 0.465379 | 4.92E-80 | cyan | PTPN6 |
| 4.13E-80 | 0.574061 | 0.462857 | 0.075908 | 1.45E-79 | cyan | RAB31 |
| 6.65E-80 | 1.002629 | 0.971429 | 0.37627 | 2.32E-79 | cyan | LGALS1 |
| 8.89E-80 | -0.81114 | 0.994286 | 0.999418 | 3.08E-79 | cyan | RPS12 |
| 9.31E-80 | -1.20889 | 0.977143 | 0.995438 | 3.21E-79 | cyan | RPLP0 |
| 1.20E-79 | 0.971945 | 0.748571 | 0.240762 | 4.11E-79 | cyan | TBXAS1 |
| 1.32E-79 | 0.881159 | 0.748571 | 0.210218 | 4.49E-79 | cyan | THEMIS2 |
| 2.24E-79 | 0.889596 | 0.68 | 0.181971 | 7.56E-79 | cyan | PRAM1 |
| 3.94E-79 | 0.928676 | 0.982857 | 0.913933 | 1.32E-78 | cyan | ARPC3 |
| 1.04E-78 | -0.74445 | 1 | 0.997573 | 3.45E-78 | cyan | RPL37 |
| 1.41E-78 | -0.69872 | 0.994286 | 0.998835 | 4.69E-78 | cyan | RPS15A |
| 2.50E-78 | -0.61585 | 0.994286 | 0.999094 | 8.24E-78 | cyan | RPS15 |
| 3.72E-78 | 0.902123 | 0.994286 | 0.953472 | 1.22E-77 | cyan | MYL6 |
| 4.67E-78 | 1.155728 | 0.845714 | 0.297677 | 1.52E-77 | cyan | ANXA2 |
| 5.72E-78 | 1.038213 | 0.942857 | 0.480877 | 1.85E-77 | cyan | PYCARD |
| 1.08E-77 | -0.84845 | 0.971429 | 0.988481 | 3.48E-77 | cyan | RPL23A |
| 2.90E-77 | 0.904286 | 0.68 | 0.177118 | 9.24E-77 | cyan | SMAP2 |
| 3.07E-77 | 1.011898 | 0.76 | 0.233223 | 9.75E-77 | cyan | NINJ1 |
| 6.44E-77 | 1.290808 | 0.994286 | 0.822818 | 2.03E-76 | cyan | S100A6 |
| 1.04E-76 | -0.76659 | 0.988571 | 0.999126 | 3.26E-76 | cyan | RPS23 |
| 1.35E-76 | 0.537414 | 0.411429 | 0.062544 | 4.20E-76 | cyan | NAPSA |
| 1.97E-75 | -0.64334 | 0.994286 | 0.998738 | 6.10E-75 | cyan | RPS7 |
| 5.81E-75 | -0.96229 | 0.994286 | 0.997864 | 1.79E-74 | cyan | RPS4X |

| | | | | | | |
|---|---|---|---|---|---|---|
| 1.43E-74 | -0.79504 | 0.982857 | 0.994952 | 4.37E-74 | cyan | RPS25 |
| 1.76E-74 | 0.537328 | 0.445714 | 0.07306 | 5.35E-74 | cyan | TGFBI |
| 2.57E-74 | 1.173607 | 0.977143 | 0.730247 | 7.78E-74 | cyan | EMP3 |
| 2.63E-74 | 1.049886 | 0.834286 | 0.317576 | 7.92E-74 | cyan | CARD16 |
| 3.75E-74 | -0.66066 | 1 | 0.999353 | 1.12E-73 | cyan | RPS14 |
| 4.57E-74 | -0.83949 | 1 | 0.998803 | 1.36E-73 | cyan | RPS3A |
| 6.17E-74 | -1.1006 | 0.948571 | 0.982916 | 1.82E-73 | cyan | RPS5 |
| 2.76E-73 | 1.204719 | 0.828571 | 0.36391 | 8.10E-73 | cyan | JPT1 |
| 6.65E-73 | 1.039744 | 0.948571 | 0.60629 | 1.94E-72 | cyan | RHOG |
| 3.59E-71 | 0.521601 | 0.325714 | 0.044296 | 1.04E-70 | cyan | MARCKS |
| 9.82E-71 | 0.894608 | 0.531429 | 0.112955 | 2.84E-70 | cyan | CDKN1A |
| 1.01E-70 | -0.75661 | 0.988571 | 0.995276 | 2.90E-70 | cyan | RPS21 |
| 1.10E-70 | -0.69987 | 1 | 0.998123 | 3.14E-70 | cyan | RPL7A |
| 1.34E-70 | -0.88165 | 0.965714 | 0.980392 | 3.81E-70 | cyan | RPL37A |
| 2.53E-70 | 1.143235 | 0.92 | 0.526694 | 7.14E-70 | cyan | CSTB |
| 2.97E-70 | 1.04757 | 0.96 | 0.668479 | 8.33E-70 | cyan | ATP6V0C |
| 4.39E-70 | -0.63345 | 1 | 0.998609 | 1.23E-69 | cyan | RPL35A |
| 5.47E-70 | -0.79465 | 0.982857 | 0.993949 | 1.52E-69 | cyan | RPL9 |
| 9.41E-70 | 0.512775 | 0.657143 | 0.145668 | 2.60E-69 | cyan | CSTA |
| 9.81E-70 | 0.792605 | 0.657143 | 0.18443 | 2.70E-69 | cyan | CD300A |
| 1.20E-69 | 0.819077 | 1 | 0.993658 | 3.29E-69 | cyan | TMSB4X |
| 2.60E-69 | -0.69284 | 0.988571 | 0.998706 | 7.08E-69 | cyan | RPL18A |
| 3.33E-69 | -0.72041 | 1 | 0.999838 | 9.02E-69 | cyan | RPLP1 |
| 3.63E-69 | 0.550882 | 0.382857 | 0.060733 | 9.77E-69 | cyan | DMXL2 |
| 4.57E-69 | -0.93985 | 0.954286 | 0.979195 | 1.23E-68 | cyan | RPL10A |
| 8.01E-69 | -0.70275 | 0.994286 | 0.999515 | 2.14E-68 | cyan | RPS8 |
| 8.70E-69 | -0.64972 | 0.988571 | 0.99835 | 2.31E-68 | cyan | RPL29 |
| 1.27E-68 | 0.815198 | 0.68 | 0.203553 | 3.35E-68 | cyan | CPVL |
| 3.47E-68 | -0.59336 | 0.994286 | 0.999029 | 9.11E-68 | cyan | RPL18 |
| 4.73E-68 | 0.647519 | 0.605714 | 0.134958 | 1.24E-67 | cyan | CTSH |
| 4.68E-67 | -0.57778 | 1 | 0.99835 | 1.22E-66 | cyan | RPL6 |
| 6.79E-67 | 0.581992 | 0.474286 | 0.092409 | 1.76E-66 | cyan | HOTAIRM1 |
| 6.98E-67 | 0.761162 | 0.971429 | 0.906038 | 1.80E-66 | cyan | RHOA |
| 1.51E-66 | 1.031885 | 0.902857 | 0.496861 | 3.86E-66 | cyan | ARRB2 |
| 3.89E-66 | 1.09301 | 0.754286 | 0.27108 | 9.91E-66 | cyan | NFKBIZ |
| 5.01E-66 | 1.00257 | 0.931429 | 0.627677 | 1.27E-65 | cyan | RNASET2 |
| 5.14E-66 | 0.992606 | 0.851429 | 0.378891 | 1.30E-65 | cyan | C1orf162 |
| 7.49E-66 | 0.909843 | 0.8 | 0.298906 | 1.88E-65 | cyan | RNF149 |
| 1.11E-65 | -0.85655 | 0.988571 | 0.992882 | 2.78E-65 | cyan | RPL5 |
| 3.34E-65 | 0.980264 | 0.925714 | 0.538342 | 8.30E-65 | cyan | SYNGR2 |
| 3.86E-65 | -0.67659 | 0.994286 | 0.995632 | 9.56E-65 | cyan | RPL14 |

| | | | | | | |
|---|---|---|---|---|---|---|
| 4.56E-65 | 0.672079 | 0.508571 | 0.114347 | 1.12E-64 | cyan | C9orf72 |
| 9.60E-65 | -0.58681 | 1 | 0.999094 | 2.35E-64 | cyan | RPS28 |
| 3.85E-64 | 1.052493 | 0.965714 | 0.649097 | 9.39E-64 | cyan | HLA-DPA1 |
| 9.13E-64 | -0.72332 | 0.982857 | 0.995438 | 2.22E-63 | cyan | RPL15 |
| 1.88E-62 | 0.822522 | 0.697143 | 0.23057 | 4.56E-62 | cyan | ZFAND5 |
| 2.59E-62 | 1.016978 | 0.977143 | 0.605869 | 6.23E-62 | cyan | S100A10 |
| 3.07E-62 | 0.509453 | 0.371429 | 0.060506 | 7.36E-62 | cyan | LRP1 |
| 3.47E-62 | 0.781248 | 0.84 | 0.3267 | 8.28E-62 | cyan | FCGRT |
| 4.18E-62 | 1.064649 | 0.788571 | 0.320229 | 9.93E-62 | cyan | ADGRE5 |
| 9.68E-62 | -1.38372 | 0.754286 | 0.919627 | 2.29E-61 | cyan | NPM1 |
| 1.56E-61 | 0.891638 | 0.92 | 0.420339 | 3.67E-61 | cyan | DUSP1 |
| 3.17E-61 | 0.915451 | 0.811429 | 0.325633 | 7.42E-61 | cyan | SELPLG |
| 3.32E-61 | 0.828122 | 0.988571 | 0.683977 | 7.75E-61 | cyan | TSPO |
| 8.91E-61 | 0.983808 | 0.862857 | 0.439235 | 2.07E-60 | cyan | RNH1 |
| 2.83E-60 | -1.44529 | 0.182857 | 0.813758 | 6.55E-60 | cyan | CD99 |
| 6.31E-60 | 0.743268 | 0.56 | 0.140005 | 1.45E-59 | cyan | GBP2 |
| 7.70E-60 | 0.998307 | 0.965714 | 0.579693 | 1.76E-59 | cyan | HLA-DRB5 |
| 1.87E-59 | -1.56941 | 0.354286 | 0.810069 | 4.26E-59 | cyan | LDHB |
| 2.14E-59 | 0.996107 | 0.931429 | 0.652074 | 4.85E-59 | cyan | ALDOA |
| 3.79E-59 | 0.719036 | 0.548571 | 0.145603 | 8.57E-59 | cyan | ZNF385A |
| 3.93E-59 | -0.5555 | 1 | 0.999223 | 8.86E-59 | cyan | RPL34 |
| 1.53E-58 | 0.576782 | 0.411429 | 0.083123 | 3.44E-58 | cyan | SLC16A3 |
| 8.80E-58 | -0.65103 | 0.988571 | 0.993852 | 1.96E-57 | cyan | RPL36 |
| 1.87E-57 | -0.98932 | 0.942857 | 0.966835 | 4.15E-57 | cyan | EEF1B2 |
| 4.32E-57 | 0.7776 | 0.982857 | 0.806057 | 9.57E-57 | cyan | GABARAP |
| 7.77E-57 | 0.853482 | 0.811429 | 0.355303 | 1.71E-56 | cyan | CTSB |
| 1.08E-56 | 0.525775 | 0.422857 | 0.088429 | 2.36E-56 | cyan | NADK |
| 2.44E-56 | 0.879156 | 0.954286 | 0.637546 | 5.33E-56 | cyan | TKT |
| 5.36E-56 | 0.70279 | 0.497143 | 0.121756 | 1.17E-55 | cyan | CDC42EP3 |
| 5.97E-56 | 0.77823 | 0.8 | 0.328092 | 1.29E-55 | cyan | GRK2 |
| 6.28E-56 | 0.511412 | 0.394286 | 0.074128 | 1.36E-55 | cyan | TNFAIP2 |
| 8.15E-56 | -0.53888 | 1 | 0.999547 | 1.75E-55 | cyan | RPL30 |
| 9.67E-56 | -0.72054 | 0.937143 | 0.977318 | 2.07E-55 | cyan | RPL13A |
| 1.50E-55 | 0.578696 | 0.388571 | 0.07471 | 3.19E-55 | cyan | ODF3B |
| 1.72E-55 | 0.749139 | 0.634286 | 0.213551 | 3.66E-55 | cyan | AP2A1 |
| 4.06E-55 | 1.188788 | 0.737143 | 0.353232 | 8.59E-55 | cyan | CUX1 |
| 1.17E-54 | -0.74312 | 0.88 | 0.967191 | 2.47E-54 | cyan | RPL31 |
| 1.68E-54 | 1.498243 | 0.868571 | 0.472691 | 3.52E-54 | cyan | MT2A |
| 1.86E-54 | 0.763412 | 0.588571 | 0.177862 | 3.88E-54 | cyan | NAAA |
| 1.96E-54 | 0.642928 | 0.508571 | 0.123115 | 4.07E-54 | cyan | RHOB |
| 3.97E-54 | -0.63351 | 0.988571 | 0.999029 | 8.23E-54 | cyan | TPT1 |

| | | | | | | |
|---|---|---|---|---|---|---|
| 1.03E-53 | 1.003188 | 0.817143 | 0.434641 | 2.13E-53 | cyan | CMTM6 |
| 1.86E-53 | 0.541961 | 0.411429 | 0.087394 | 3.82E-53 | cyan | RELT |
| 1.90E-53 | -0.65065 | 0.977143 | 0.983725 | 3.89E-53 | cyan | RPL35 |
| 2.07E-53 | 0.706115 | 0.994286 | 0.968712 | 4.22E-53 | cyan | SH3BGRL3 |
| 2.16E-53 | 0.797602 | 1 | 0.826571 | 4.39E-53 | cyan | SRGN |
| 3.24E-53 | -0.66503 | 0.628571 | 0.150165 | 6.56E-53 | cyan | S100A8 |
| 5.44E-53 | 0.518856 | 0.44 | 0.100692 | 1.10E-52 | cyan | PIK3AP1 |
| 5.66E-53 | 0.592723 | 0.502857 | 0.126577 | 1.14E-52 | cyan | ARAP1 |
| 7.68E-53 | 0.753645 | 0.577143 | 0.182715 | 1.54E-52 | cyan | C15orf39 |
| 1.40E-52 | 0.835671 | 0.737143 | 0.302854 | 2.80E-52 | cyan | RGS19 |
| 2.05E-52 | 0.59739 | 1 | 0.979616 | 4.07E-52 | cyan | PFN1 |
| 2.29E-52 | 0.665585 | 0.474286 | 0.117356 | 4.53E-52 | cyan | DPEP2 |
| 3.24E-52 | -0.51518 | 0.8 | 0.247848 | 6.37E-52 | cyan | LYZ |
| 5.90E-52 | -0.67166 | 0.914286 | 0.969715 | 1.16E-51 | cyan | RPL27 |
| 7.45E-52 | 0.879559 | 0.72 | 0.304213 | 1.46E-51 | cyan | VASP |
| 7.49E-52 | 0.81665 | 0.971429 | 0.639164 | 1.46E-51 | cyan | HLA-DRB1 |
| 7.96E-52 | -0.55765 | 0.982857 | 0.993367 | 1.55E-51 | cyan | RPL22 |
| 1.13E-51 | -0.85122 | 0.988571 | 0.994629 | 2.19E-51 | cyan | RPS27 |
| 1.85E-51 | -0.91899 | 0.914286 | 0.958325 | 3.56E-51 | cyan | HNRNPA1 |
| 2.40E-51 | 0.772391 | 0.954286 | 0.868504 | 4.60E-51 | cyan | ARPC2 |
| 6.56E-51 | 0.852928 | 0.651429 | 0.241086 | 1.25E-50 | cyan | INSIG1 |
| 3.01E-50 | -0.56526 | 0.971429 | 0.99314 | 5.73E-50 | cyan | RPS16 |
| 8.30E-50 | 0.769511 | 0.925714 | 0.584708 | 1.58E-49 | cyan | ATP6V0B |
| 1.01E-49 | -0.936 | 0.811429 | 0.915777 | 1.90E-49 | cyan | HINT1 |
| 1.11E-49 | 0.804538 | 0.908571 | 0.646897 | 2.10E-49 | cyan | FKBP1A |
| 3.15E-49 | 0.628567 | 0.617143 | 0.179221 | 5.91E-49 | cyan | DOK2 |
| 6.45E-49 | 0.606979 | 0.731429 | 0.257523 | 1.21E-48 | cyan | MYO1F |
| 1.02E-48 | 0.749872 | 0.925714 | 0.690254 | 1.90E-48 | cyan | GNG5 |
| 1.98E-48 | 0.695415 | 0.702857 | 0.269656 | 3.67E-48 | cyan | TNFSF13B |
| 3.41E-48 | -0.51147 | 0.988571 | 0.994046 | 6.31E-48 | cyan | RPL24 |
| 3.59E-48 | 0.761183 | 0.72 | 0.300621 | 6.62E-48 | cyan | RAB24 |
| 8.38E-48 | 0.843468 | 0.88 | 0.583058 | 1.54E-47 | cyan | ARPC5 |
| 1.99E-47 | 0.583535 | 0.994286 | 0.99877 | 3.64E-47 | cyan | TMSB10 |
| 3.08E-47 | 0.654081 | 0.651429 | 0.216722 | 5.61E-47 | cyan | RGS2 |
| 3.20E-47 | 0.553675 | 0.445714 | 0.117291 | 5.81E-47 | cyan | NAGA |
| 8.94E-47 | 0.761408 | 0.794286 | 0.343526 | 1.62E-46 | cyan | PLEK |
| 4.67E-46 | 0.757181 | 0.76 | 0.339546 | 8.43E-46 | cyan | PTP4A2 |
| 6.26E-46 | 0.731135 | 0.948571 | 0.756487 | 1.13E-45 | cyan | HIGD2A |
| 1.32E-45 | 0.697948 | 0.874286 | 0.530835 | 2.37E-45 | cyan | SEC11A |
| 1.84E-45 | 0.697273 | 0.628571 | 0.237559 | 3.30E-45 | cyan | SEC14L1 |
| 2.30E-45 | 0.746822 | 0.948571 | 0.642367 | 4.09E-45 | cyan | ITGB2 |

| | | | | | | |
|---|---|---|---|---|---|---|
| 1.06E-44 | 0.69601 | 0.714286 | 0.31282 | 1.88E-44 | cyan | ATP6V0D1 |
| 1.54E-44 | 0.56148 | 0.514286 | 0.142755 | 2.72E-44 | cyan | JAML |
| 1.89E-44 | 0.776675 | 0.765714 | 0.355659 | 3.34E-44 | cyan | MYO1G |
| 3.43E-44 | 0.725608 | 0.514286 | 0.161878 | 6.02E-44 | cyan | CNIH4 |
| 3.87E-44 | 0.62798 | 0.748571 | 0.268686 | 6.77E-44 | cyan | IL10RA |
| 6.65E-44 | -0.65875 | 0.965714 | 0.966253 | 1.16E-43 | cyan | RPL4 |
| 8.68E-44 | 0.538119 | 0.417143 | 0.111402 | 1.51E-43 | cyan | ARRB1 |
| 8.69E-44 | 0.796706 | 0.828571 | 0.428622 | 1.51E-43 | cyan | ZEB2 |
| 9.75E-44 | 0.555977 | 0.422857 | 0.108361 | 1.68E-43 | cyan | PTPN18 |
| 2.78E-43 | 0.55571 | 0.497143 | 0.141332 | 4.79E-43 | cyan | EVI2B |
| 4.73E-43 | 0.606015 | 0.582857 | 0.195981 | 8.13E-43 | cyan | ZYX |
| 4.82E-43 | 0.756193 | 0.965714 | 0.706821 | 8.25E-43 | cyan | ZFP36 |
| 2.13E-42 | 0.771403 | 0.668571 | 0.298874 | 3.64E-42 | cyan | RAB10 |
| 3.03E-42 | 0.566212 | 0.462857 | 0.132401 | 5.14E-42 | cyan | RARA |
| 4.05E-42 | -0.54075 | 0.982857 | 0.992688 | 6.86E-42 | cyan | RACK1 |
| 5.73E-42 | 0.74346 | 0.691429 | 0.328642 | 9.68E-42 | cyan | MAPKAPK3 |
| 5.98E-42 | 0.849268 | 0.731429 | 0.360157 | 1.01E-41 | cyan | VAMP5 |
| 1.13E-41 | 0.684945 | 0.708571 | 0.307416 | 1.89E-41 | cyan | CSK |
| 2.10E-41 | 0.757755 | 0.914286 | 0.614638 | 3.52E-41 | cyan | PSMB9 |
| 2.35E-41 | 0.553619 | 0.588571 | 0.18841 | 3.91E-41 | cyan | ITGAL |
| 1.71E-40 | -0.5985 | 0.977143 | 0.97075 | 2.84E-40 | cyan | RPL7 |
| 2.34E-40 | 0.618133 | 0.514286 | 0.157089 | 3.87E-40 | cyan | ADA2 |
| 3.83E-40 | 0.746756 | 0.748571 | 0.39261 | 6.32E-40 | cyan | HSBP1 |
| 5.62E-40 | -0.52666 | 0.982857 | 0.986637 | 9.25E-40 | cyan | RPL21 |
| 1.23E-39 | -1.45876 | 0.017143 | 0.566945 | 2.02E-39 | cyan | CTSW |
| 1.28E-39 | 0.71715 | 0.977143 | 0.785867 | 2.10E-39 | cyan | CD52 |
| 4.47E-39 | 0.621825 | 0.72 | 0.320682 | 7.28E-39 | cyan | GNAI2 |
| 4.80E-39 | 0.587674 | 0.548571 | 0.195205 | 7.80E-39 | cyan | IMPDH1 |
| 2.01E-38 | 0.670504 | 0.92 | 0.709927 | 3.25E-38 | cyan | HCLS1 |
| 2.02E-38 | 0.585354 | 0.977143 | 0.980619 | 3.26E-38 | cyan | UBC |
| 2.09E-38 | 0.584113 | 0.485714 | 0.158869 | 3.37E-38 | cyan | NOTCH2 |
| 3.76E-38 | -0.98481 | 0.245714 | 0.68553 | 6.04E-38 | cyan | PEBP1 |
| 3.81E-38 | 0.603853 | 0.382857 | 0.10286 | 6.10E-38 | cyan | MBD2 |
| 6.69E-38 | 0.849767 | 0.874286 | 0.649291 | 1.07E-37 | cyan | LY6E |
| 1.07E-37 | 0.666115 | 0.925714 | 0.694169 | 1.71E-37 | cyan | LAMTOR4 |
| 1.23E-37 | 0.533127 | 0.405714 | 0.113926 | 1.94E-37 | cyan | SNX10 |
| 2.58E-37 | 0.589071 | 0.971429 | 0.857147 | 4.08E-37 | cyan | TPM3 |
| 3.19E-37 | -1.33074 | 0.285714 | 0.779784 | 5.03E-37 | cyan | JUN |
| 8.71E-37 | 0.66095 | 0.971429 | 0.700835 | 1.37E-36 | cyan | NAP1L1 |
| 1.33E-36 | 0.743965 | 0.68 | 0.316735 | 2.08E-36 | cyan | SOD2 |
| 1.64E-36 | 0.568168 | 0.708571 | 0.31337 | 2.57E-36 | cyan | AGTRAP |

| | | | | | | |
|---|---|---|---|---|---|---|
| 2.02E-36 | 0.619237 | 0.405714 | 0.124862 | 3.15E-36 | cyan | ICAM4 |
| 2.75E-36 | 0.528536 | 0.394286 | 0.113117 | 4.27E-36 | cyan | NUDT16 |
| 5.60E-36 | 0.58167 | 0.48 | 0.163107 | 8.67E-36 | cyan | TNFSF13 |
| 5.78E-36 | 0.511003 | 0.468571 | 0.14845 | 8.91E-36 | cyan | C11orf21 |
| 1.13E-35 | 0.654175 | 0.868571 | 0.536045 | 1.74E-35 | cyan | FLNA |
| 3.74E-35 | 0.737544 | 0.754286 | 0.418333 | 5.74E-35 | cyan | CD55 |
| 3.82E-35 | 0.587748 | 0.782857 | 0.373164 | 5.84E-35 | cyan | HERPUD1 |
| 5.12E-35 | 0.593528 | 0.742857 | 0.311396 | 7.81E-35 | cyan | SAMHD1 |
| 1.47E-34 | 0.509193 | 0.565714 | 0.206724 | 2.23E-34 | cyan | TSPAN14 |
| 1.91E-34 | 0.578093 | 0.502857 | 0.185045 | 2.90E-34 | cyan | CD151 |
| 2.52E-34 | -0.73027 | 0.828571 | 0.931243 | 3.82E-34 | cyan | RPS29 |
| 3.11E-34 | 0.699769 | 0.628571 | 0.29085 | 4.69E-34 | cyan | VMP1 |
| 5.11E-34 | 0.633563 | 0.594286 | 0.263347 | 7.67E-34 | cyan | BLVRA |
| 1.00E-33 | 0.699175 | 0.805714 | 0.502621 | 1.50E-33 | cyan | LRRFIP1 |
| 4.67E-33 | -0.85901 | 0.045714 | 0.539507 | 6.97E-33 | cyan | ACAP1 |
| 7.54E-33 | 0.618979 | 0.64 | 0.291011 | 1.12E-32 | cyan | MGAT1 |
| 9.07E-33 | 0.555863 | 0.565714 | 0.212548 | 1.35E-32 | cyan | MVP |
| 1.59E-32 | 0.58768 | 0.902857 | 0.738336 | 2.36E-32 | cyan | CAPZB |
| 1.90E-32 | 0.614438 | 0.96 | 0.803792 | 2.81E-32 | cyan | LSP1 |
| 2.26E-32 | 0.618511 | 0.457143 | 0.162816 | 3.33E-32 | cyan | PAK1 |
| 2.55E-32 | 0.526116 | 0.942857 | 0.749563 | 3.74E-32 | cyan | PRR13 |
| 3.09E-32 | 0.560913 | 0.965714 | 0.819679 | 4.52E-32 | cyan | CORO1A |
| 3.10E-32 | 0.71049 | 0.88 | 0.640491 | 4.53E-32 | cyan | TUBA1A |
| 3.60E-32 | 0.58531 | 0.577143 | 0.244904 | 5.24E-32 | cyan | RNF13 |
| 3.61E-32 | 0.653905 | 0.931429 | 0.651168 | 5.24E-32 | cyan | HLA-DPB1 |
| 5.44E-32 | 0.711706 | 0.725714 | 0.398531 | 7.88E-32 | cyan | POLD4 |
| 9.66E-32 | 0.519399 | 0.737143 | 0.366757 | 1.40E-31 | cyan | GRN |
| 1.05E-31 | 0.553599 | 0.577143 | 0.222999 | 1.51E-31 | cyan | FMNL1 |
| 1.08E-31 | 0.692187 | 0.908571 | 0.689704 | 1.55E-31 | cyan | PSME2 |
| 1.13E-31 | 0.632247 | 0.868571 | 0.605966 | 1.62E-31 | cyan | DRAP1 |
| 2.02E-31 | 0.503301 | 0.514286 | 0.198829 | 2.88E-31 | cyan | FAM45A |
| 2.14E-31 | 0.629226 | 0.902857 | 0.739921 | 3.05E-31 | cyan | ITM2B |
| 3.14E-31 | 0.566423 | 0.68 | 0.327412 | 4.46E-31 | cyan | ARPC1B |
| 4.87E-31 | -1.1173 | 0.125714 | 0.560053 | 6.90E-31 | cyan | NUCB2 |
| 4.94E-31 | 0.512748 | 0.965714 | 0.795153 | 6.98E-31 | cyan | CLIC1 |
| 5.68E-31 | 0.675692 | 0.862857 | 0.629554 | 8.01E-31 | cyan | CAP1 |
| 9.41E-31 | 0.507294 | 0.514286 | 0.189284 | 1.32E-30 | cyan | MIDN |
| 1.34E-30 | -0.96171 | 0.302857 | 0.68346 | 1.88E-30 | cyan | GYPC |
| 2.10E-30 | 0.592576 | 0.874286 | 0.653627 | 2.93E-30 | cyan | MSN |
| 2.13E-30 | 0.563439 | 0.445714 | 0.160131 | 2.97E-30 | cyan | PDLIM5 |
| 5.40E-30 | 0.662517 | 0.754286 | 0.413318 | 7.51E-30 | cyan | CYTIP |

| | | | | | | |
|---|---|---|---|---|---|---|
| 9.65E-30 | 0.562643 | 0.554286 | 0.22727 | 1.34E-29 | cyan | IFNGR1 |
| 1.29E-29 | 0.58281 | 0.685714 | 0.366757 | 1.78E-29 | cyan | C20orf27 |
| 1.44E-29 | -0.55492 | 0.754286 | 0.881803 | 1.99E-29 | cyan | SNRPD2 |
| 1.60E-29 | 0.536672 | 0.502857 | 0.201191 | 2.20E-29 | cyan | HK1 |
| 1.88E-29 | 0.553539 | 0.925714 | 0.727626 | 2.57E-29 | cyan | LCP1 |
| 2.17E-29 | 0.651309 | 0.834286 | 0.48311 | 2.98E-29 | cyan | GLUL |
| 2.23E-29 | -1.25089 | 0.034286 | 0.485181 | 3.05E-29 | cyan | CD69 |
| 2.43E-29 | -0.88415 | 0.885714 | 0.920274 | 3.30E-29 | cyan | HMGB1 |
| 2.79E-29 | -0.60911 | 0.834286 | 0.889698 | 3.79E-29 | cyan | RPL27A |
| 5.58E-29 | 0.527216 | 0.942857 | 0.738821 | 7.56E-29 | cyan | TPI1 |
| 1.08E-28 | -1.11261 | 0.182857 | 0.575422 | 1.46E-28 | cyan | SELL |
| 1.14E-28 | 0.609475 | 0.817143 | 0.535624 | 1.54E-28 | cyan | PGLS |
| 1.17E-28 | -1.25444 | 0.177143 | 0.597683 | 1.57E-28 | cyan | HIST1H4C |
| 1.19E-28 | 0.620262 | 0.765714 | 0.487543 | 1.60E-28 | cyan | MTPN |
| 1.28E-28 | -1.6909 | 0.051429 | 0.480166 | 1.70E-28 | cyan | STMN1 |
| 1.32E-28 | 0.619746 | 0.834286 | 0.557529 | 1.76E-28 | cyan | WAS |
| 1.53E-28 | -0.55031 | 0.868571 | 0.922216 | 2.03E-28 | cyan | RPL38 |
| 2.50E-28 | 0.683777 | 0.64 | 0.326733 | 3.32E-28 | cyan | BLOC1S1 |
| 2.75E-28 | 0.580698 | 0.594286 | 0.288876 | 3.64E-28 | cyan | LGALS9 |
| 3.02E-28 | -0.73523 | 0.674286 | 0.8278 | 3.98E-28 | cyan | MIF |
| 5.29E-28 | 0.554362 | 0.822857 | 0.522908 | 6.96E-28 | cyan | CYBA |
| 8.43E-28 | 0.601154 | 0.474286 | 0.197308 | 1.11E-27 | cyan | P2RX1 |
| 1.12E-27 | 0.622877 | 0.742857 | 0.459975 | 1.47E-27 | cyan | LTA4H |
| 1.74E-27 | 0.5306 | 0.737143 | 0.40429 | 2.27E-27 | cyan | LCP2 |
| 1.97E-27 | -1.08752 | 0.034286 | 0.470556 | 2.57E-27 | cyan | TRBC2 |
| 2.11E-27 | -0.83324 | 0.12 | 0.553291 | 2.75E-27 | cyan | PRDX2 |
| 3.00E-27 | 0.56373 | 0.822857 | 0.44548 | 3.90E-27 | cyan | CD48 |
| 3.25E-27 | 0.606553 | 0.525714 | 0.232835 | 4.21E-27 | cyan | AMPD2 |
| 4.60E-27 | 0.610693 | 0.897143 | 0.748269 | 5.94E-27 | cyan | PGK1 |
| 4.84E-27 | -0.66544 | 0.005714 | 0.429075 | 6.23E-27 | cyan | OCIAD2 |
| 7.03E-27 | 0.562301 | 0.382857 | 0.134569 | 9.02E-27 | cyan | IRAK3 |
| 7.52E-27 | -0.71403 | 0.497143 | 0.78412 | 9.63E-27 | cyan | C12orf57 |
| 8.83E-27 | 0.56404 | 0.542857 | 0.23785 | 1.13E-26 | cyan | RABGAP1L |
| 2.08E-26 | 0.553551 | 0.645714 | 0.324274 | 2.65E-26 | cyan | REEP5 |
| 3.22E-26 | 0.618742 | 0.902857 | 0.738659 | 4.10E-26 | cyan | CALM2 |
| 3.24E-26 | -0.51169 | 0.937143 | 0.966835 | 4.11E-26 | cyan | EEF2 |
| 4.64E-26 | 0.511231 | 0.937143 | 0.74691 | 5.87E-26 | cyan | PTPRC |
| 6.54E-26 | 0.560486 | 0.668571 | 0.365948 | 8.26E-26 | cyan | PARVG |
| 1.85E-25 | 0.53114 | 0.76 | 0.418527 | 2.33E-25 | cyan | IQGAP1 |
| 1.93E-25 | 0.508872 | 0.674286 | 0.362324 | 2.42E-25 | cyan | OSTF1 |
| 2.01E-25 | -0.93186 | 0.937143 | 0.965444 | 2.52E-25 | cyan | RPS26 |

| | | | | | | |
|---|---|---|---|---|---|---|
| 2.99E-25 | -0.79168 | 0.011429 | 0.415162 | 3.74E-25 | cyan | RHOH |
| 3.71E-25 | 0.524332 | 0.48 | 0.200608 | 4.62E-25 | cyan | CAPNS1 |
| 6.05E-25 | -0.85556 | 0.04 | 0.442277 | 7.52E-25 | cyan | 1-Sep |
| 6.47E-25 | -0.83513 | 0.794286 | 0.86941 | 8.03E-25 | cyan | HSP90AB1 |
| 7.30E-25 | 0.618498 | 0.514286 | 0.225458 | 9.04E-25 | cyan | ISG15 |
| 8.73E-25 | 0.553624 | 0.605714 | 0.305701 | 1.08E-24 | cyan | ERICH1 |
| 9.41E-25 | -0.80472 | 0.822857 | 0.892966 | 1.16E-24 | cyan | RPS2 |
| 2.02E-24 | 0.640058 | 0.617143 | 0.349511 | 2.48E-24 | cyan | ATG3 |
| 4.13E-24 | 0.523497 | 0.662857 | 0.339643 | 5.06E-24 | cyan | GLIPR1 |
| 4.64E-24 | 0.513255 | 0.411429 | 0.161522 | 5.67E-24 | cyan | ATP6V1B2 |
| 8.13E-24 | 0.538853 | 0.577143 | 0.268847 | 9.93E-24 | cyan | UTRN |
| 9.72E-24 | 0.535617 | 0.834286 | 0.576458 | 1.18E-23 | cyan | ATP6V0E1 |
| 1.09E-23 | -1.35574 | 0.022857 | 0.412088 | 1.32E-23 | cyan | CD7 |
| 2.47E-23 | -0.69354 | 0.017143 | 0.405973 | 2.99E-23 | cyan | ITM2A |
| 3.42E-23 | 0.603902 | 0.588571 | 0.291432 | 4.13E-23 | cyan | ATP2B1 |
| 9.28E-23 | -0.64161 | 0 | 0.372225 | 1.12E-22 | cyan | BEX3 |
| 1.12E-22 | 0.502682 | 0.788571 | 0.53818 | 1.35E-22 | cyan | AP2S1 |
| 1.54E-22 | -0.92787 | 0 | 0.369475 | 1.85E-22 | cyan | AC084033.3 |
| 2.00E-22 | 0.565319 | 0.617143 | 0.334595 | 2.40E-22 | cyan | GLRX |
| 5.61E-22 | -1.74984 | 0.045714 | 0.403352 | 6.70E-22 | cyan | CD3E |
| 8.26E-22 | -0.50374 | 0.794286 | 0.888598 | 9.85E-22 | cyan | PCBP2 |
| 1.45E-21 | 0.515449 | 0.697143 | 0.436582 | 1.72E-21 | cyan | VPS29 |
| 2.96E-21 | 0.536719 | 0.8 | 0.553161 | 3.52E-21 | cyan | STK17B |
| 5.82E-21 | -0.6008 | 0.565714 | 0.766421 | 6.89E-21 | cyan | PNISR |
| 7.29E-21 | -0.86833 | 0.011429 | 0.361548 | 8.62E-21 | cyan | ITM2C |
| 8.57E-21 | 0.505939 | 0.542857 | 0.276063 | 1.01E-20 | cyan | PCGF5 |
| 1.28E-20 | -0.59471 | 0.731429 | 0.811461 | 1.51E-20 | cyan | NCL |
| 2.09E-20 | -0.61967 | 0.36 | 0.647738 | 2.45E-20 | cyan | HMGN1 |
| 3.78E-20 | -0.69327 | 0.308571 | 0.577493 | 4.43E-20 | cyan | APEX1 |
| 6.84E-20 | 0.527691 | 0.788571 | 0.582411 | 7.99E-20 | cyan | ARL6IP5 |
| 1.08E-19 | -0.71682 | 0.148571 | 0.499547 | 1.26E-19 | cyan | ANXA6 |
| 1.49E-19 | -0.72368 | 0.142857 | 0.497929 | 1.74E-19 | cyan | AES |
| 3.41E-19 | -1.16849 | 0.011429 | 0.337766 | 3.96E-19 | cyan | PRSS57 |
| 3.64E-19 | -1.93615 | 0.068571 | 0.389989 | 4.22E-19 | cyan | IL32 |
| 8.37E-19 | -0.63175 | 0.468571 | 0.663949 | 9.67E-19 | cyan | SNHG8 |
| 1.30E-18 | -0.52232 | 0 | 0.317802 | 1.50E-18 | cyan | ARMH1 |
| 1.32E-18 | -0.527 | 0.531429 | 0.746845 | 1.52E-18 | cyan | RPL17 |
| 1.64E-18 | -0.55979 | 0.685714 | 0.785867 | 1.89E-18 | cyan | EIF3E |
| 2.47E-18 | -0.58555 | 0.451429 | 0.686792 | 2.83E-18 | cyan | 6-Sep |
| 2.83E-18 | -0.6262 | 0.017143 | 0.335081 | 3.24E-18 | cyan | SLC38A1 |
| 4.36E-18 | 0.530064 | 0.691429 | 0.45276 | 4.98E-18 | cyan | SDCBP |

| | | | | | | |
|---|---|---|---|---|---|---|
| 5.17E-18 | -1.10582 | 0.011429 | 0.321167 | 5.88E-18 | cyan | CD3D |
| 8.12E-18 | -0.50708 | 0.08 | 0.414741 | 9.22E-18 | cyan | ZNF22 |
| 9.34E-18 | -0.57436 | 0.177143 | 0.490326 | 1.06E-17 | cyan | NOP58 |
| 1.13E-17 | 0.503843 | 0.714286 | 0.47926 | 1.27E-17 | cyan | NOP10 |
| 1.35E-17 | -0.92199 | 0.011429 | 0.321459 | 1.52E-17 | cyan | LCK |
| 1.58E-17 | 0.537535 | 0.497143 | 0.238886 | 1.78E-17 | cyan | ATP2B1-AS1 |
| 1.66E-17 | -0.55208 | 0.331429 | 0.608393 | 1.87E-17 | cyan | FBL |
| 5.48E-17 | -0.82259 | 0.234286 | 0.49039 | 6.15E-17 | cyan | IMPDH2 |
| 7.61E-17 | 0.575832 | 0.697143 | 0.43655 | 8.51E-17 | cyan | ZFP36L1 |
| 7.83E-17 | 0.541034 | 0.697143 | 0.494305 | 8.74E-17 | cyan | WSB1 |
| 1.75E-16 | -0.80747 | 0.011429 | 0.305183 | 1.95E-16 | cyan | SPOCK2 |
| 2.90E-16 | -0.55032 | 0.051429 | 0.353006 | 3.22E-16 | cyan | CD82 |
| 2.90E-16 | -0.61147 | 0 | 0.284281 | 3.22E-16 | cyan | FAM30A |
| 2.92E-16 | -0.51309 | 0.205714 | 0.511454 | 3.23E-16 | cyan | ABHD14B |
| 4.95E-16 | -0.61626 | 0 | 0.280852 | 5.46E-16 | cyan | CD34 |
| 6.07E-16 | -0.56444 | 0.291429 | 0.574322 | 6.69E-16 | cyan | NUCKS1 |
| 1.06E-15 | -0.81826 | 0.017143 | 0.295574 | 1.16E-15 | cyan | FCMR |
| 2.10E-15 | -0.52551 | 0.062857 | 0.356889 | 2.30E-15 | cyan | ATF7IP2 |
| 3.56E-15 | -1.49398 | 0.022857 | 0.298292 | 3.89E-15 | cyan | CST7 |
| 5.39E-15 | -1.37733 | 0.022857 | 0.291594 | 5.89E-15 | cyan | IL7R |
| 9.06E-15 | -0.73818 | 0 | 0.261761 | 9.88E-15 | cyan | SMIM24 |
| 9.82E-15 | -0.64934 | 0.125714 | 0.402317 | 1.07E-14 | cyan | CDK6 |
| 1.01E-14 | -0.51941 | 0.182857 | 0.454831 | 1.10E-14 | cyan | PARP1 |
| 1.16E-14 | -0.56035 | 0.057143 | 0.33398 | 1.26E-14 | cyan | BIN1 |
| 1.72E-14 | -0.64152 | 0.097143 | 0.372711 | 1.86E-14 | cyan | FABP5 |
| 2.84E-14 | -0.57609 | 0.04 | 0.307416 | 3.07E-14 | cyan | EGFL7 |
| 6.28E-14 | -0.65919 | 0.508571 | 0.683168 | 6.76E-14 | cyan | HMGB2 |
| 6.43E-14 | -0.65145 | 0.594286 | 0.719051 | 6.91E-14 | cyan | CALR |
| 7.82E-14 | -0.73448 | 0.011429 | 0.262344 | 8.38E-14 | cyan | HOPX |
| 1.02E-13 | -0.89878 | 0.005714 | 0.252766 | 1.09E-13 | cyan | SPINK2 |
| 1.49E-13 | -0.7132 | 0.011429 | 0.261826 | 1.59E-13 | cyan | ALOX5AP |
| 1.76E-13 | -0.52975 | 0.28 | 0.513072 | 1.88E-13 | cyan | CCT2 |
| 2.08E-13 | -0.61875 | 0.022857 | 0.269365 | 2.21E-13 | cyan | PRKCH |
| 2.23E-13 | -1.33335 | 0.011429 | 0.256973 | 2.37E-13 | cyan | IGLL1 |
| 3.46E-13 | -0.55854 | 0.011429 | 0.256746 | 3.66E-13 | cyan | ABLIM1 |
| 4.84E-13 | -0.61378 | 0.12 | 0.378794 | 5.12E-13 | cyan | P2RY8 |
| 7.03E-13 | -1.20169 | 0.08 | 0.331586 | 7.42E-13 | cyan | AREG |
| 7.19E-13 | -0.83175 | 0.017143 | 0.253867 | 7.57E-13 | cyan | GZMM |
| 1.04E-12 | -0.67255 | 0.034286 | 0.277357 | 1.09E-12 | cyan | ZAP70 |
| 1.13E-12 | -0.73218 | 0.017143 | 0.253122 | 1.19E-12 | cyan | CD3G |
| 1.20E-12 | -0.61408 | 0.028571 | 0.267165 | 1.25E-12 | cyan | LBH |

| | | | | | | |
|---|---|---|---|---|---|---|
| 1.30E-12 | -0.56461 | 0.44 | 0.588397 | 1.36E-12 | cyan | TXN |
| 2.48E-12 | -0.63617 | 0.514286 | 0.674465 | 2.58E-12 | cyan | HSPA5 |
| 4.29E-12 | -0.54007 | 0.074286 | 0.317964 | 4.46E-12 | cyan | RNF125 |
| 9.30E-12 | -0.63562 | 0.062857 | 0.296124 | 9.65E-12 | cyan | CLEC11A |
| 1.48E-11 | -0.54833 | 0.051429 | 0.278878 | 1.53E-11 | cyan | MGAT4A |
| 1.67E-11 | -0.53912 | 0.091429 | 0.324468 | 1.72E-11 | cyan | MCM7 |
| 2.26E-11 | -0.607 | 0.36 | 0.531094 | 2.33E-11 | cyan | HSPD1 |
| 8.04E-11 | -0.54181 | 0.051429 | 0.267974 | 8.28E-11 | cyan | LAT |
| 2.16E-10 | -0.67215 | 0.331429 | 0.557173 | 2.22E-10 | cyan | DNAJB1 |
| 3.10E-10 | -0.73143 | 0.422857 | 0.605546 | 3.18E-10 | cyan | CXCR4 |
| 3.70E-10 | -0.85887 | 0.217143 | 0.422831 | 3.78E-10 | cyan | TNFAIP3 |
| 1.17E-09 | -0.52166 | 0.08 | 0.292726 | 1.19E-09 | cyan | HIST1H1C |
| 4.53E-09 | -0.74356 | 0.194286 | 0.393807 | 4.62E-09 | cyan | SOX4 |
| 6.04E-09 | -0.66539 | 0.217143 | 0.413124 | 6.14E-09 | cyan | DDIT4 |
| 7.42E-09 | -0.57968 | 0.4 | 0.538795 | 7.54E-09 | cyan | DUT |
| 2.40E-08 | -0.58875 | 0.268571 | 0.477189 | 2.43E-08 | cyan | HIST1H1E |
| 6.36E-08 | -0.97457 | 0.451429 | 0.57361 | 6.44E-08 | cyan | IFITM1 |
| 9.38E-08 | -0.5736 | 0.222857 | 0.406847 | 9.47E-08 | cyan | SLC2A3 |
| 3.66E-06 | -0.55504 | 0.16 | 0.292144 | 3.69E-06 | cyan | PIK3IP1 |
| 5.33E-06 | -0.58673 | 0.285714 | 0.446904 | 5.36E-06 | cyan | AL138963.3 |
| 0.00546 | -0.92755 | 0.405714 | 0.396428 | 0.005482 | cyan | LTB |
| 0.013491 | -1.73299 | 0.245714 | 0.305734 | 0.013517 | cyan | NKG7 |

**Sup. Tab. 5**: DE genes bone marrow
